# Embryonic Protein NODAL Regulates the Breast Tumour Microenvironment by Reprogramming Cancer-Derived Secretomes

**DOI:** 10.1101/2020.07.09.195842

**Authors:** Dylan Dieters-Castator, Paola Marino Dantonio, Matt Piaseczny, Guihua Zhang, Jiahui Liu, Miljan Kuljanin, Stephen Sherman, Michael Jewer, Katherine Quesnel, Eun Young Kang, Martin Köbel, Gabrielle M. Siegers, Andrew Leask, David Hess, Gilles Lajoie, Lynne-Marie Postovit

**Affiliations:** Department of Anatomy and Cell Biology, University of Western Ontario, London ON, Canada; Department of Biomedical and Molecular Sciences, Queen’s University, Kingston ON, Canada; Department of Oncology, University of Alberta, Edmonton AB, Canada; Robarts Research Institute, London ON, Canada; Department of Biochemistry, University of Western Ontario, London ON, Canada; Department of Physiology and Pharmacology, University of Western Ontario, London ON, Canada; Department of Pathology and Laboratory Medicine, University of Calgary, Foothills Medical Centre, Calgary, AB, Canada

**Author notes:** To whom correspondence should be addressed: Lynne-Marie Postovit, 563 Botterell Hall, Queen’s University, Kingston ON, Canada, K7L 2V7.

## Abstract

The tumour microenvironment (TME) is an important mediator of breast cancer progression. Cancer-associated fibroblasts (CAFs) constitute a major component of the TME and may originate from tissue-associated fibroblasts or infiltrating mesenchymal stromal cells (MSCs). The mechanisms by which cancer cells activate fibroblasts and recruit MSCs to the TME are largely unknown, but likely include deposition of a pro-tumourigenic secretome. The secreted embryonic protein NODAL is clinically associated with breast cancer stage and promotes tumour growth, metastasis, and vascularization. Herein, we show that NODAL expression correlates with the presence of activated fibroblasts in human triple negative breast cancers and that it directly induces CAF phenotypes. We further show that NODAL reprograms cancer cell secretomes by simultaneously altering levels of chemokines (e.g. CXCL1), cytokines (e.g. IL-6) and growth factors (e.g. PDGFRA), leading to alterations in MSC chemotaxis. We therefore demonstrate a hitherto unappreciated mechanism underlying the dynamic regulation of the TME.

## Introduction

Solid malignancies contain many non-transformed stromal cell types within the tumour microenvironment (TME). In breast cancer, a significant proportion of auxiliary cells, including cancer-associated fibroblasts (CAFs), pericytes, myoepithelial and endothelial cells, tumour-associated macrophages (TAMs) and other immune cell types, cooperate to promote pro-tumorigenic processes such as metastasis and drug resistance^1–3^. Dramatic gene expression changes in stromal cells are associated with the transformation of normal breast tissue to ductal carcinoma *in situ* (DCIS)^4,5^, and progression from DCIS to invasive ductal carcinoma (IDC) is marked by upregulated expression of extracellular matrix (ECM)-degrading proteases^5^, revealing an important role of TME components in breast cancer initiation and progression.

Among TME cell types, CAFs constitute the major stromal component of many breast cancers and have recently emerged as potential therapeutic targets^6–8^. Fibroblasts are the main producers of ECM and play fundamental roles in tissue repair, during which they acquire an activated myofibroblast phenotype characterized by α-smooth muscle actin (α-SMA) expression^9,10^. Fibroblasts are highly responsive to their microenvironment, interacting with and influencing a wide range of cells. For example, fibroblasts promote angiogenesis through the secretion of vascular endothelial growth factor A (VEGFA)^11^, coordinate immune response through cytokine and chemokine release^12,13^, and influence epithelial stem cells^14,15^. Ligands from the transforming growth factor beta (TGF-β) family are well-known fibroblast activators^16,17^, as are cytokines^18^ and ECM remodelling^19,20^. Extracellular factors, such as cytokines and ECM proteins, mediate the pro-tumorigenic behaviours of CAFs. For instance, CAF-derived CXCL12/stromal derived factor (SDF-1) can mobilize endothelial progenitor cells (EPCs) to increase vascularization of MCF-7 xenografts^21^. Moreover, subsets of CAFs can increase tumorigenesis and breast cancer stem cell (BCSC) enrichment by secreting interleukins (IL) IL-6, IL-8 and IL-1β^6,7,22^. Several recent studies have demonstrated that CAFs are heterogeneous and can be derived through activation of tissue-associated fibroblasts^23^, as well as the recruitment of mesenchymal stromal cells (MSCs)^24,25^. In fact, up to 20% of CAFs were derived from MSCs in a CXCL6/CXCR6-dependent manner in a mouse model of gastric cancer^26^. An orthotopic breast cancer model revealed that MSCs can be recruited to primary tumour sites and that TGF-β1 is involved in this process^27^. MSCs also acquire CAF-like phenotypes when cultured in tumour conditioned media or mixed with cancer cells in mouse xenografts^28–30^. The mechanisms underlying MSC recruitment are not fully understood, but it is becoming increasingly clear that this population may contribute to the CAF compartment in the TME.

Limited genetic alterations have been described in breast cancer-associated stromal cells^31–34^, suggesting that changes in gene expression observed in these cells are mainly due to epigenetic reprogramming^35,36^. For instance, breast cancer cells induce fibroblasts to secrete the ECM protein degrader ADAMTS1 through epigenetic changes^37^, demonstrating that the epigenetic reprogramming in stromal cells can be induced by cancer cells. The cancer secretome plays a vital role in the pro-tumorigenic effects of the TME, recruiting stromal cells and reprogramming them to support tumour progression^38–40^.

Several studies have uncovered tumour promoting roles for the secreted TGF-β superfamily member and embryonic morphogen NODAL^41,42^. NODAL expression, while primarily restricted to embryonic development and human embryonic stem cells (hESCs), has been observed in melanoma, glioblastoma, breast, pancreatic and hepatocellular cancers, amongst others^42,43^. In breast cancer, NODAL clinically correlates with stage and vascularization^44,45^. Moreover, NODAL expression emerges in breast cancers as they transition from DCIS into IDC^46^, wherein interactions between cancer and stromal cells are critical. NODAL inhibition reduces breast cancer-induced neovascularization and mitigates BCSC frequencies, tumour growth, and invasion^47–49^. NODAL may also play an essential role in remodeling the TME. For example, NODAL seems to induce a breast cancer secretome that promotes angiogenesis through regulation of the angiogenic factors platelet-derived growth factor (PDGF) and vascular endothelial growth factor (VEGF)^45^. Furthermore, NODAL expression is inversely correlated with susceptibility to gamma delta (γδ) T cell cytotoxicity, at least in part through decreased surface expression of the immune activating danger signal MHC class I polypeptide-related sequence A/B (MICA/B)^50^. CAFs from a gastric cancer mouse model have recently been shown to promote cancer cell proliferation and resistance to doxorubicin via NODAL secretion^51^ and NODAL appears to induce CAF-like phenotypes in mouse and human fibroblast cell lines^52^. The extent to which NODAL may affect CAF phenotypes in the breast TME has not, however, been explored.

In this study, we investigated the impact of NODAL on CAFs and MSCs within the triple negative breast cancer (TNBC) TME. We demonstrate that NODAL strongly correlates with CAFs in breast cancer patients and that this morphogen can directly signal to fibroblasts (but not to MSCs) to induce a CAF-associated phenotype. In addition, mass spectrometry-based proteomics of conditioned media derived from triple negative MDA-MB-231 and triple negative inflammatory SUM149 breast cancer cells demonstrated that NODAL is a potent regulator of the breast cancer secretome. Our analyses revealed cancer cell-type-specific alterations in several novel NODAL-regulated factors, including CXCL1, CXCL8, IL-6 and colony-stimulating factor 1 (CSF1), suggesting that NODAL may impact the ability of breast cancer cells to recruit a variety of stromal cell types. Accordingly, we found that MSC chemotaxis towards breast cancer cells is affected by NODAL-regulated factors such as IL-6. Collectively, these data reveal a previously unknown role for NODAL in the regulation of breast cancer TME.

## Results

### Cancer cells expressing NODAL associate with α-SMA-positive stromal cells in triple negative breast tumors

Because fibroblast activation (demarcated by α-SMA expression) and NODAL expression occur early during breast cancer progression^46^, we investigated whether these events could be correlated. We evaluated the expression and localization of NODAL and α-SMA in 41 primary tumour tissue samples from a cohort of 20 TNBC cases (**Table I**). Representative images are shown in **Fig. 1**. NODAL expression was observed in 92.7% of samples (38/41) (**Table I**), while α-SMA was detected in all slides. Stromal-associated α-SMA (**Fig. 1b, d**) was observed in all 38 NODAL-positive samples and the intensity of α-SMA staining was found to be increased in 94.7% (36/38) of regions with NODAL-positive cells as compared to NODAL-negative regions (**Table I**). Notably, α-SMA was also detected in areas that were negative for NODAL; however, in these instances, α-SMA delineated myoepithelial cells (**Fig. 1e, f**). Overall, these results reveal a strong association between NODAL and α-SMA expression in the stroma of TNBC patients, suggesting NODAL could have an impact on CAF phenotypes in breast cancer.

**Table I.** NODAL and α-SMA evaluation by immunohistochemistry.

|  | *n* (%) |
| --- | --- |
| **NODAL score – Median (range)** | 2 (0-3) |
| 0 | 3 (7.3) |
| 1 | 10 (24.4) |
| 2 | 19 (46.3) |
| 3 | 9 (22.0) |
| **NODAL percentage (%) – Median (range)** | 50 (5-100) |
| NA | 3 (7.3) |
| 5-20 | 12 (29.3) |
| 30-40 | 7 (17.1) |
| 60-80 | 14 (34.1) |
| 90-100 | 5 (12.2) |
| **NODAL distribution** |  |
| NA | 3 (7.3) |
| Diffuse | 17 (41.5) |
| Scattered | 20 (48.8) |
| Focal | 1 (2.4) |
| **α-SMA score – Median (range)** | 2 (1-3) |
| 1 | 3 (7.3) |
| 2 | 29 (70.7) |
| 3 | 9 (22.0) |
| **Intensity Association** |  |
| NA | 3 (NA) |
| Yes | 36 (94.7) |
| No | 2 (5.3) |

**Figure 1.**
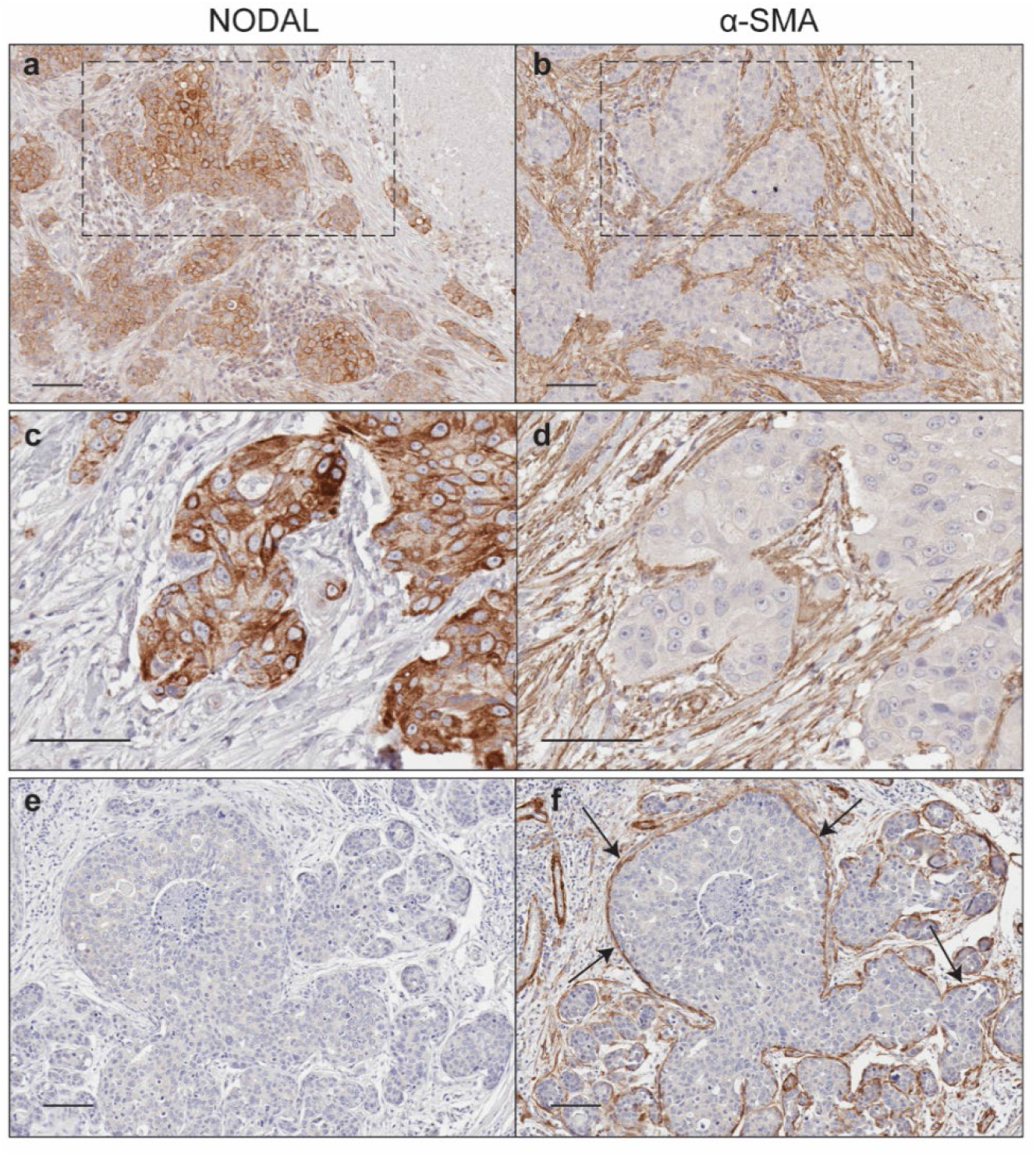
Breast cancer cells expressing NODAL reside adjacent to α-SMA-positive stromal cells. Representative images are shown wherein NODAL (a, c, e) and α-SMA (b, d, f) are stained in serial sections of tissue from triple negative breast cancer patients. **(a-d)** NODAL-positive breast cancer cells are surrounded by diffuse α-SMA+ stromal cells (for example in square). **(e, f)** In NODAL-negative sections, α-SMA is localized only to basement membranes (arrows). Bar equals 100 μm.

### NODAL induces fibroblast activation and chemotaxis

Since we found a consistent spatial association between NODAL and stromal α-SMA in human TNBC tissues, we decided to examine whether NODAL affects, directly or indirectly, breast cancer-induced fibroblast phenotypes (**Fig. 2**). MDA-MB-231 TNBC cells express high basal levels of NODAL, therefore we investigated if serum-free conditioned media (CM) of MDA-MB-231 cells stably expressing scrambled control (shControl) or NODAL knockdown (shNODAL) shRNA (**Fig. 2a**) can differentially impact primary fibroblasts. We also explored the effects of recombinant human NODAL (rhNODAL) on these cells. We detected a small, but statistically insignificant reduction in primary human foreskin fibroblast (HFF) chemotaxis towards CM from shNODAL versus shControl MDA-MB-231 cells (**Fig. 2b**), while rhNODAL (10 and 100 ng/mL) increased HFF chemotaxis (**Fig. 2c**), proliferation (**Fig. 2d**) and invasion (**Fig. 2e**). Further addressing fibroblast activation by NODAL, real-time RT-PCR revealed that rhNODAL (10 and 100ng/mL) induced expression of α-SMA, desmin, and connective tissue growth factor (CTGF) (**Fig. 2f**), which are CAF markers. In addition, we performed gene expression profiling on human dermal fibroblasts (HDFs) treated with 10ng/mL rhNODAL for 6h. Transcripts upregulated by at least 1.7 fold were analyzed in DAVID; gene clusters associated with the GO terms “wound healing”, “cell motion”, “extracellular matrix” and “growth factor” were significantly enriched (**Fig. 2g; Sup. Table 1**)^53^, suggesting that fibroblasts are indeed activated by NODAL.

**Figure 2.**
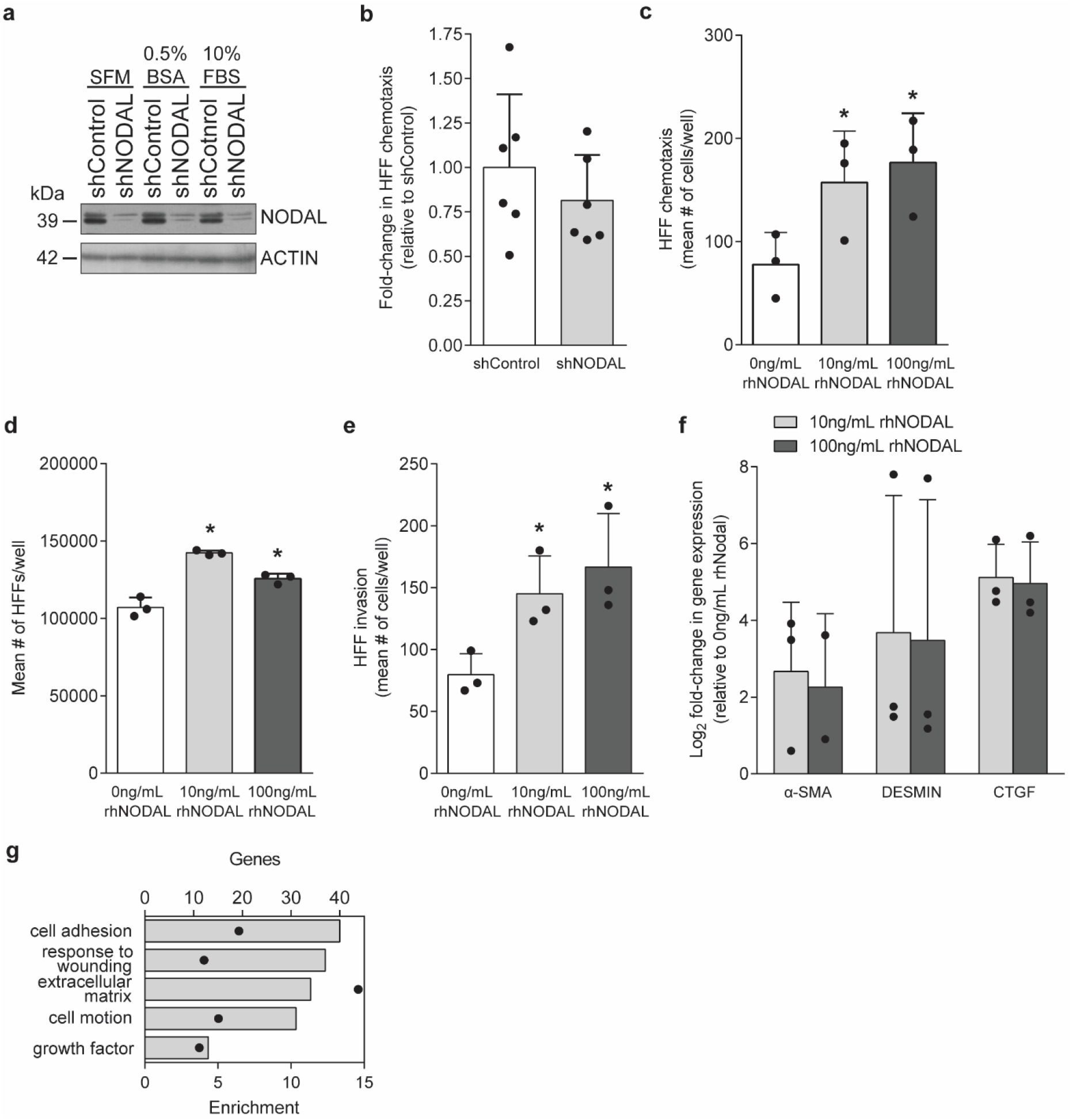
NODAL directly promotes phenotypes associated with activated fibroblasts in HFFs. **(a)** NODAL expression in MDA-MB-231 cells stably expressing scrambled (shControl) or NODAL knockdown (shNODAL) shRNA cultured in SFM, SFM+0.5%BSA (0.5%BSA), or complete media (10% FBS). **(b)** HFF chemotaxis towards CM from MDA-MB-231 cells (n=6). **(c-e)** Exposure to rhNODAL (10 and 100ng/mL) significantly increased HFF chemotaxis, proliferation, and invasion (n=3); Dunnett’s multiple comparison test, *p<0.05. **(f)** HFFs upregulated transcripts (*a-SMA, DESMIN* and *CTGF*) associated with activated fibroblasts following treatment with rhNODAL (10 and 100ng/mL; n=3, n=2 for α-SMA from 100ng/mL treatment). **(g)** Total genes (bars) upregulated by NODAL (10ng/mL) more than 1.7-fold in human dermal fibroblasts (HDFs) and their corresponding enrichment (black dots) following GO analysis in DAVID.

### The NODAL-regulated MDA-MB-231 secretome impacts MSC chemotaxis

The fibroblast population in breast cancer is highly heterogeneous and likely derived from different cell types^23,54^. Given the involvement of MSCs in tumour growth and neovascularization, and their probable contribution to the CAF population, we examined how NODAL affects the capacity of breast cancer cells to promote chemotaxis by comparing the ability of CM from shControl and shNODAL knockdown MDA-MB-231 cells to influence MSC chemotaxis (**Fig. 3**). Several primary human bone marrow-derived (BM-)MSC lines were utilized herein, some of which have been previously shown to form tubes *in vitro* and stimulate islet regeneration and revascularization *in vivo*^55,56^. In three out of four MSC lines, chemotaxis was significantly decreased (~1.8 to 3.5-fold) towards shNODAL CM as compared to shControl CM. We did not observe appreciable differences in proliferation or viability of MSCs cultured in CM for 24h, suggesting that the effects observed were not due to alterations in cell numbers, but rather a result of altered chemotaxis (**Sup. Fig. 1a**).

**Figure 3.**
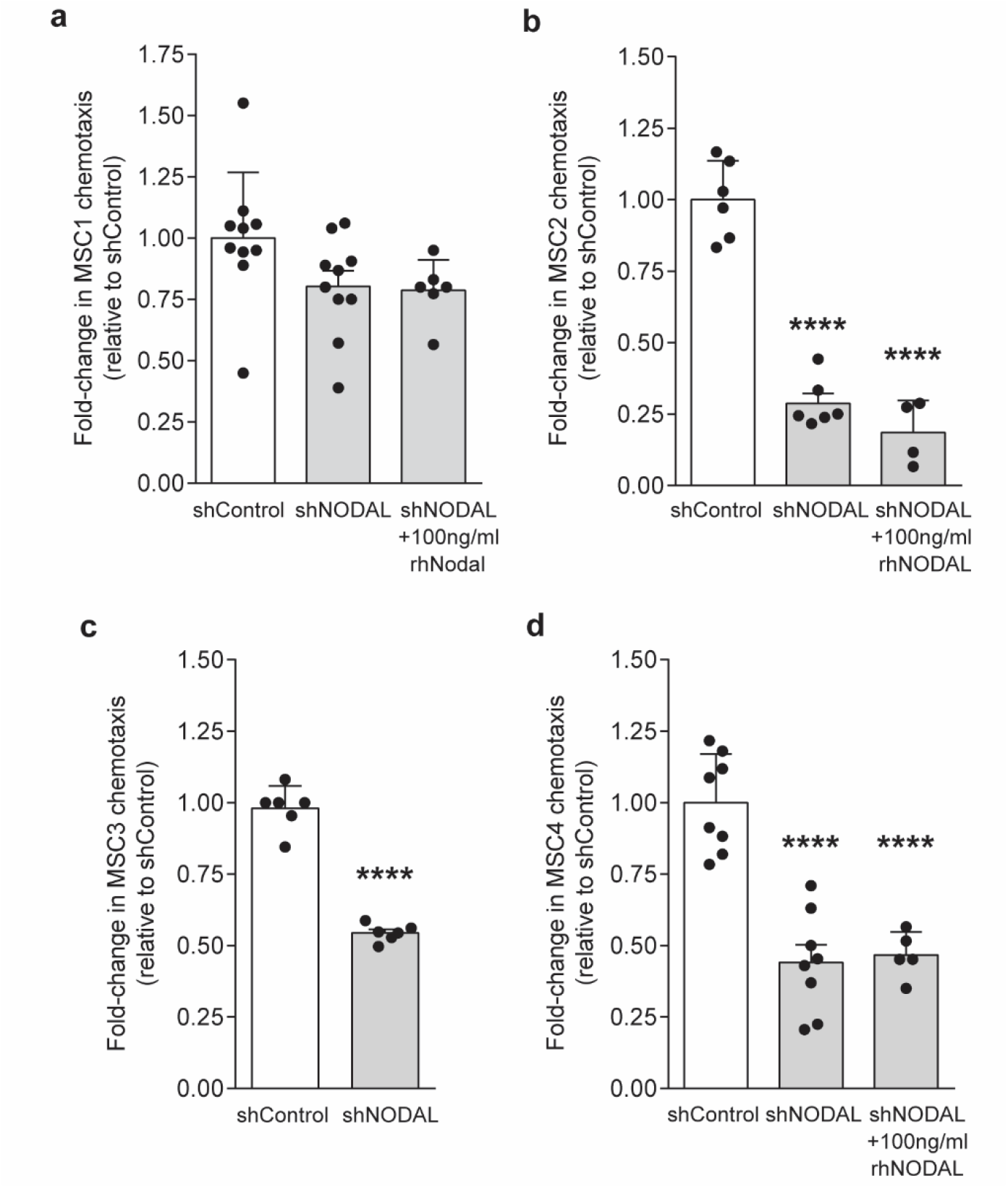
Conditioned media from NODAL-knockdown MDA-MB231 breast cancer cells indirectly modulates MSC chemotaxis. **(a-d)** Human bone-marrow derived MSC lines (MSC1-4) were plated onto fibronectin-coated transwells in the presence of CM (shControl or shNODAL +/-100ng/mL rhNODAL). MSC chemotaxis was significantly lower towards shNODAL CM compared to shControl CM after 24h and was not rescued by rhNODAL; ****p<0.0001, Dunnett’s multiple comparison test.

The reduction in MSC chemotaxis observed when NODAL was knocked down could not be rescued by the addition of 100ng/mL of rhNODAL (**Fig. 3a, b, d**) suggesting that MSCs are unable to sense this morphogen, perhaps due to an absence of receptor components. Hence, we performed real-time RT-PCR and western blotting for NODAL, its receptor (ALK4) and co-receptor (CRIPTO) on MSC lines (**Fig. 4a, b**). MSCs expressed moderate levels of NODAL and high levels of ALK4 at the transcript and protein level (**Fig. 4a, b**). *CRIPTO* mRNA expression approached the reliable limit of detection by quantitative real-time PCR (35 cycles) (**Fig. 4a**). Hence, while MSCs appear to make NODAL and to express NODAL receptors, they may not express enough CRIPTO to sense NODAL. Stimulation with 10 and 100 ng/mL rhNODAL did not affect canonical or non-canonical signalling through SMAD2 or ERK1/2 phosphorylation, respectively (**Fig. 4c**). In contrast to MSCs, which appeared unresponsive to NODAL, we found that rhNODAL (10 and 100ng/mL) caused an increase in both SMAD2 and ERK1/2 activation in fibroblasts (**Fig. 4d**), suggesting NODAL can directly promote fibroblast activation.

**Figure 4.**
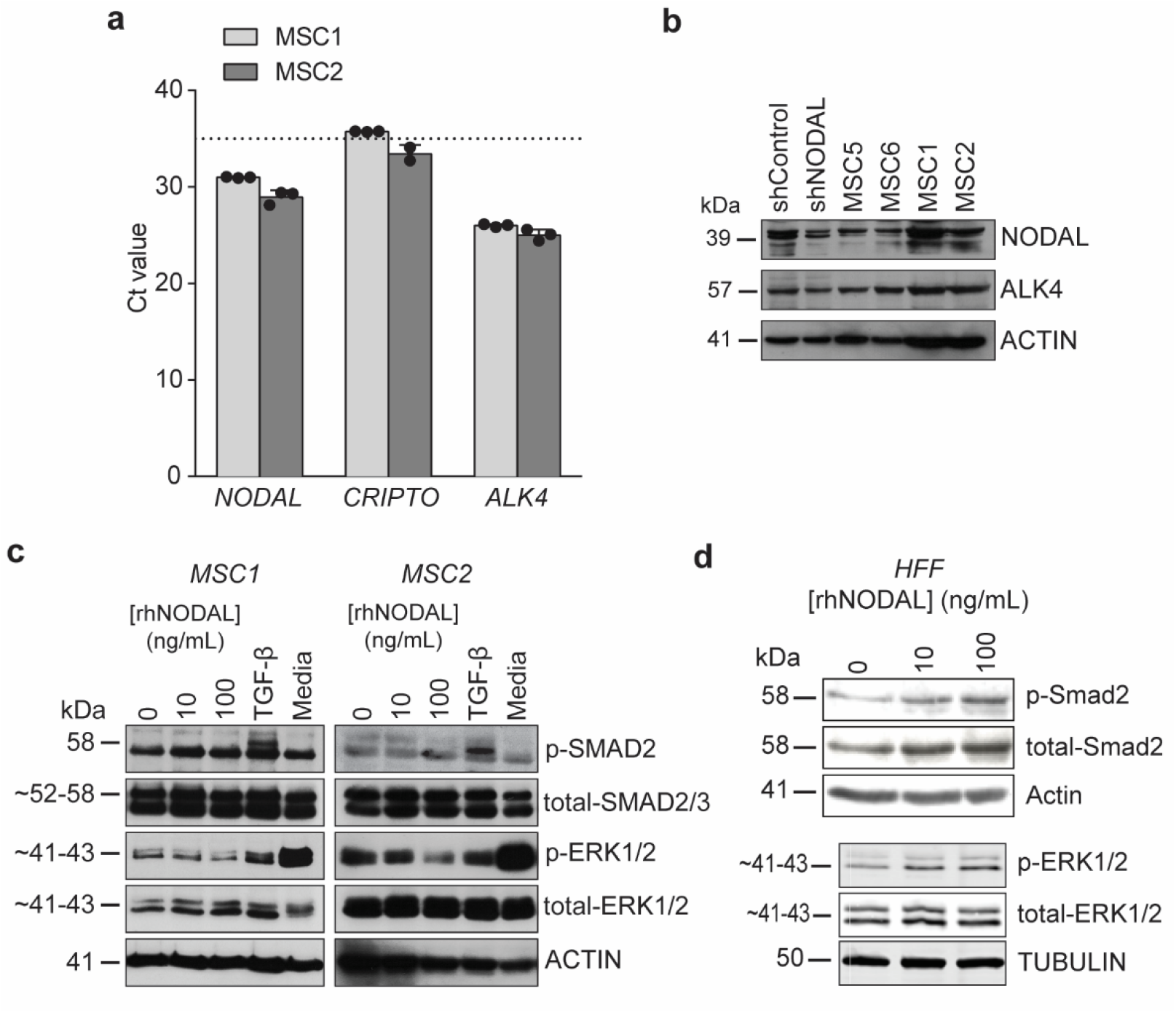
NODAL signalling in MSCs. **(a)** Real time PCR cycle threshold (Ct) values for *NODAL, ALK4* and *CRIPTO* in MSCs. Data are presented as mean Ct values ± SD from three biological replicates except for *CRIPTO* (n=2 for MSC2). High Ct values indicate low transcript expression with the horizontal dotted line corresponding to a Ct value of 35 or the reliable limit of detection. **(b)** Western blots showing expression of NODAL and ALK4 (receptor) in four MSC lines. shControl and shNODAL MDA-MB-231 cells were used as positive controls. **(c)** Serum-starved MSCs treated with varying concentrations of rhNODAL had no effect on downstream SMAD2 (p-SMAD2) or ERK1/2 (p-ERK1/2) activation. TGF-β treatment and cell culture media were used as positive controls for SMAD2 and ERK1/2 activation, respectively. **(d)** Stimulation of HFFs with rhNODAL activates SMAD2 and ERK1/2 phosphorylation in a dose-dependent fashion. Western blots are representative images taken from three biological replicates.

### Modulation of NODAL expression alters the breast cancer secretome

Mass spectrometry is a powerful tool for proteomic characterization of cancer cell lines and tissues^57,58^. To further elucidate the mechanisms through which NODAL may influence stromal cells, we employed high-resolution mass spectrometry to identify NODAL-regulated factors in serum-free CM from breast cancer cells. Stable isotopic labelling of amino acids in culture (SILAC) was combined with SDS-PAGE fractionation to determine relative changes in secreted proteins from shControl and shNODAL MDA-MB-231 cells. In total, this approach identified over 3200 proteins, which were reduced to ~1300 entries after filtering for proteins annotated with Gene Ontology Cellular Component (GOCC) terms containing “extracellular” and quantified in at least two out of three biological replicates (**Fig. 5a, Sup. Table 2**). Of those, 122 proteins were significantly different (p<0.05) between shControl and shNODAL CM (**Fig. 5a, Sup. Table 2**). From this list, 1D annotation enrichment in Perseus revealed a significant decrease in proteins involved in GO Biological Processes (GOBPs) associated with cell migration, inflammation and cytokine signalling following NODAL knockdown (**Fig. 5b, Sup. Table 3**)^59^. Alternatively, proteins matching to GOBP terms mRNA processes, protein localization and macromolecular complex disassembly were significantly increased. This observation was attributed to higher levels of ribosomal proteins (RPS and RPL members) shed by shNODAL MDA-MB-231 cells. We plotted Heavy/Light ratios (shNODAL/shControl) and their corresponding −log_10_ p-values for the ~1300 filtered extracellular proteins found in MDA-MB-231 CM (**Fig. 5c**). All proteins annotated with the aforementioned GOBP terms were highlighted in blue (depleted) or red (enriched); there was a clear trend towards a reduction in the secretion of inflammatory and chemotactic proteins following NODAL knockdown and an opposing increase in transcriptional and translational proteins. CXCL chemokines (CXCL1/3/8), IL-6 and CSF1 were significantly lower in shNODAL CM (p<0.05). Interleukin 11 (IL-11), on the other hand, was significantly higher (~1.85 fold, p<0.05). These factors have been associated with malignant phenotypes and may contribute to MSC chemotaxis given that they can promote chemotaxis of various immune cells and, in some cases, MSCs^60–62^. Similar to previous findings, PDGFA was significantly lower in shNODAL CM (2.31 fold)^45^. Proteomic findings were verified by ELISAs with CM from MDA-MB-231 cells for CXCL1, CXCL8, IL-6 and CSF1 (**Sup. Fig. 2a**). These results suggest that NODAL expression is associated with the secretion of inflammatory and chemotactic proteins by breast cancer cells.

**Figure 5.**
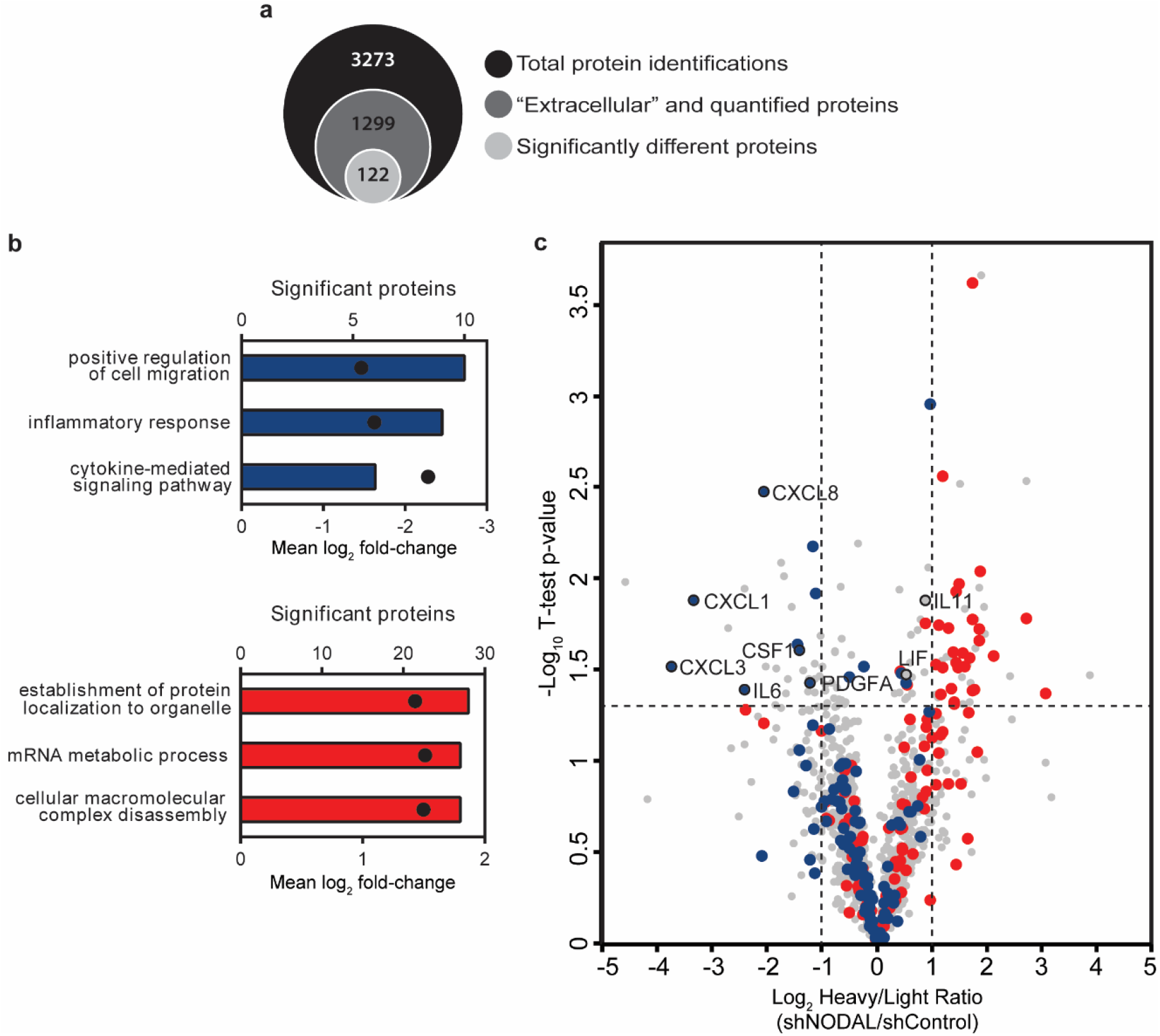
NODAL knockdown alters the MDA-MB-231 secretome. Extracellular proteins from serum-free shControl and shNODAL CM were analyzed by high-resolution mass spectrometry. **(a)** Venn diagram highlighting total protein identifications, number of “extracellular” and quantified proteins, and significantly different proteins between shControl and shNODAL CM (two-tailed, one sample t-test, p<0.05). **(b)** Number of significant proteins (bars) matching to a subset of significantly enriched (Benjamini Hochberg (BH) FDR threshold<0.02) GO biological processes (GOBPs). Mean log_2_ fold-changes in GOBPs are indicated by black dots. Blue and red bars highlight GOBPs decreased and increased in MDA-MB-231 CM following NODAL knockdown, respectively. **(c)** Volcano plot of quantified “extracellular” proteins. Negative and positive Log_2_ Heavy/Light ratios indicate proteins decreased and increased in MDA-MB-231 CM following NODAL knockdown, respectively (n=3). All proteins matching to corresponding GOBPs mentioned are highlighted in blue and red. Several cytokines and chemokines altered by NODAL are labelled in black. Vertical and horizontal dotted lines indicate Log_2_ fold-changes ≥2 and the −log_10_ p-value cut-off corresponding to p<0.05, respectively.

### IL-6 promotes MSC chemotaxis

Given that NODAL consistently altered CXCL1 and IL-6 levels in MDA-MB-231 CM, which were associated with differential MSC chemotaxis, we sought to determine whether receptors for these ligands were expressed by MSCs. While three MSC lines were highly positive for IL-6R based on flow cytometry (**Fig. 6a; Sup. Fig. 3a, b**), surface CXCR1 and CXCR2 expression could not be detected in any of the MSC lines by real-time PCR or flow cytometry (**Sup. Fig. 3c**; data not shown). Accordingly, treatment with 10 and 25ng/mL recombinant human IL-6 (rhIL-6) induced STAT3 phosphorylation in MSC2 cells, which could be blocked by the addition of an IL-6 neutralizing monoclonal antibody (mAb, **Fig. 6b**). Moreover, low doses of rhIL-6 (1 and 10ng/mL) significantly increased MSC2 chemotaxis by ~1.6 fold (p<0.05) although higher concentrations had no effect (**Fig. 6c**). Neutralizing IL-6 in shControl CM resulted in a small, but significant, decrease in MSC2 chemotaxis (Fig. 6d), while supplementing shNODAL CM with rhIL-6 (1ng/mL) increased MSC2 chemotaxis (**Fig. 6e**). These findings suggest that IL-6 may be involved in promoting MSC recruitment to breast cancers.

**Figure 6.**
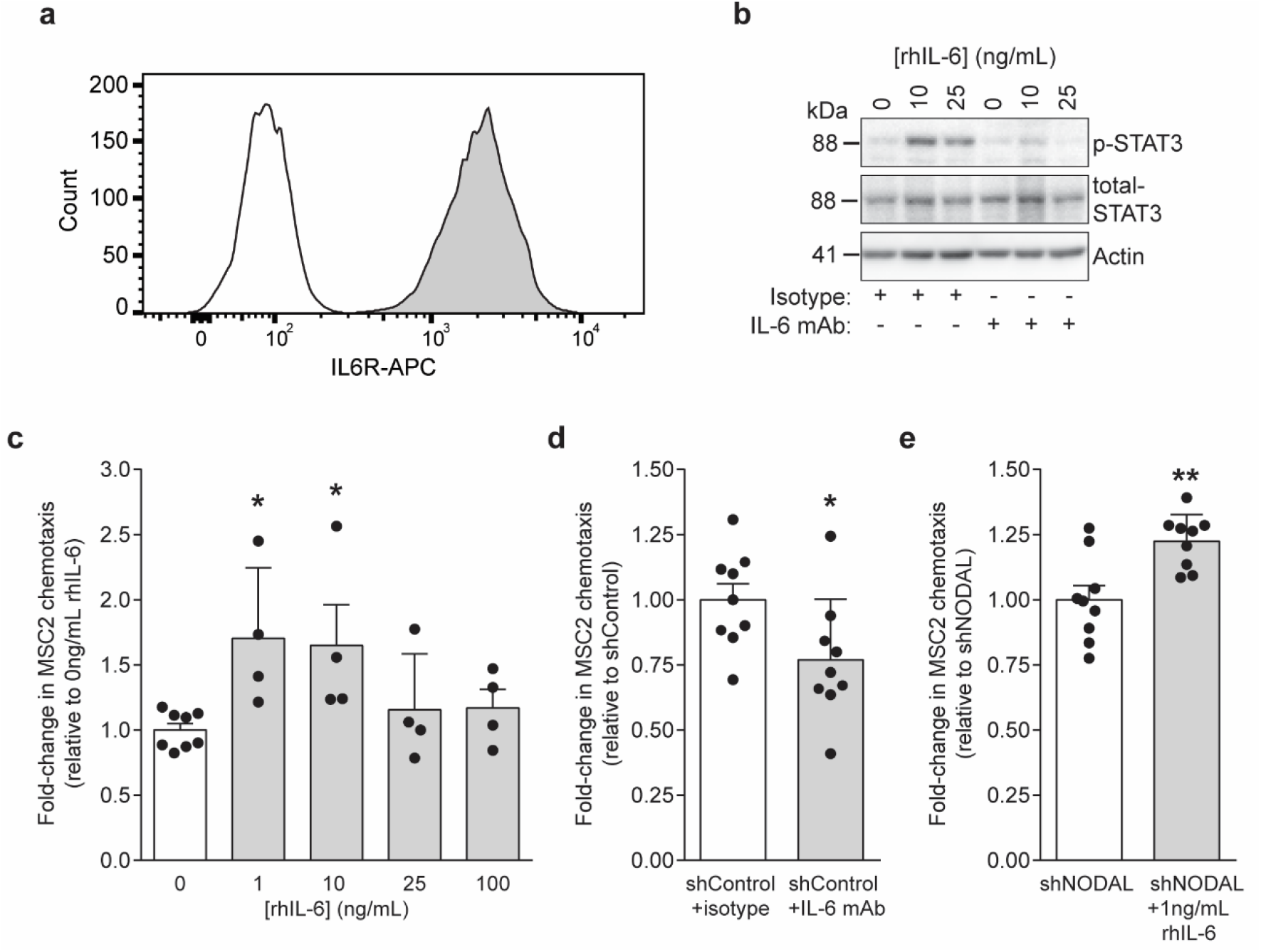
IL-6 contributes to MDA-MB-231 mediated MSC chemotaxis. **(a)** Flow cytometry showing nearly homogenous expression of the IL-6 receptor (IL-6R) by MSCs. **(b)** Stimulation with rhIL-6 induced phosphorylation of STAT3 in MSC2, which could be blocked by pre-incubation with an IL-6 neutralizing mAb. **(c)** MSC chemotaxis towards rhIL-6 after 24h (n=4-8). Low concentrations (1-10ng/mL) of rhIL-6 significantly induced MSC chemotaxis (Dunnett’s multiple comparison test, *p<0.05). **(d)** IL-6 neutralizing mAb significantly attenuated MSC chemotaxis. **(e)** Exogenous rhIL-6 significantly increased MSC chemotaxis towards shNODAL CM. Flow histogram and western blots are representative images from three biological replicates. Data are presented as mean fold-changes relative to controls ± SD. Black dots indicate replicate values and asterisks indicate significance differences (one-way ANOVA, Dunnett’s multiple comparison test for IL-6 dose response and two-tailed, two sample t-test for MDA-MB-231 treatments) in MSC chemotaxis compared to controls (* p<0.05, ** p<0.01).

### NODAL-induced reprogramming of the breast cancer secretome is context-dependent

NODAL/ACTIVIN regulates cell fate specification and phenotype by activating signal transduction pathways that directly affect transcription and mediate epigenetic modifications^63^. The ability of NODAL to broadly affect gene expression is context-dependent. Given that inflammatory breast cancer is marked by a discrete TME composition and function^64^, tumour cells and TME components in this breast cancer subtype may well cooperate under distinct cellular and signalling contexts. Therefore, we investigated the impact of NODAL on the secretome of SUM149 triple negative inflammatory breast cancer cells, which express low levels of NODAL, to determine if it differs from that of MDA-MB-231 cells.

In contrast to the knockdown model previously employed in MDA-MB-231 cells, we generated NODAL-overexpressing and green fluorescent protein (GFP) control SUM149 cells from which Strong Cation Exchange (SCX)-fractionated CM digests were obtained for label-free quantitative proteomics (**Fig. 7a**). Approximately 1500 proteins were annotated as “extracellular” and quantified in at least two out of three biological replicates, and 344 proteins were significantly different between NODAL and GFP expressing SUM149 cells (**Fig. 7b, Sup. Table 4**). GOBPs that were significantly enriched or depleted included terms associated with inflammation, cell migration/locomotion, translation, and transcription (**Fig. 7c, Sup. Table 5**). Unexpectedly, GOBPs depleted in shNODAL MDA-MB-231 samples were also depleted in NODAL overexpressing SUM149 CM. For instance, proteins matching the cytokine-mediated signalling pathway had a mean Log_2_ fold-change of −2.28 and −2.44 following NODAL knockdown and overexpression, respectively (**Fig. 5b and 7c**). Conversely, proteins matching “mRNA metabolic process” were increased significantly by NODAL knockdown in MDA-MB-231 and NODAL overexpression in SUM149 with mean Log_2_ fold-changes of 1.51 and 1.88, respectively. We also plotted Log_2_ protein fold-changes for SUM149 secretomes (NODAL-GFP) versus −log_10_ p-values and highlighted all proteins annotated with the aforementioned GOBPs (**Fig. 7d**). Several inflammatory and migratory factors decreased following NODAL overexpression while translational and transcriptional proteins were elevated. As in NODAL knockdown in MDA-MB-231 cells, CXCL1, CXCL3, IL-6 and CSF1 levels also decreased following NODAL overexpression in SUM149 cells. Highly similar proteomic results were also observed when comparing CM from NODAL overexpressing SUM149 cells to cells expressing an empty vector (EV) (**Sup. Fig. 4 and Sup. Tables 6 and 7**). In total, 56 proteins were significantly altered by NODAL in both MDA-MB-231 and SUM149 datasets; however, only a handful were associated with NODAL expression in a positive (CLU and CLSTN3) and negative (leukemia inhibitory factor [LIF] and neuropillin-2 [NRP2]) manner in both cell lines. Moreover, NODAL promoted a pro-angiogenic phenotype in both MDA-MB-231 and SUM149 cells. For example, although PDGFA was not detected in SUM149 CM, the angiogenic factors angiopoietin-1 (ANGPT1) and angiogenin (ANG) were significantly elevated in CM from NODAL-overexpressing SUM149 cells (Fig. 7d, Sup. Tables 4 and 6)^65,66^.

**Figure 7.**
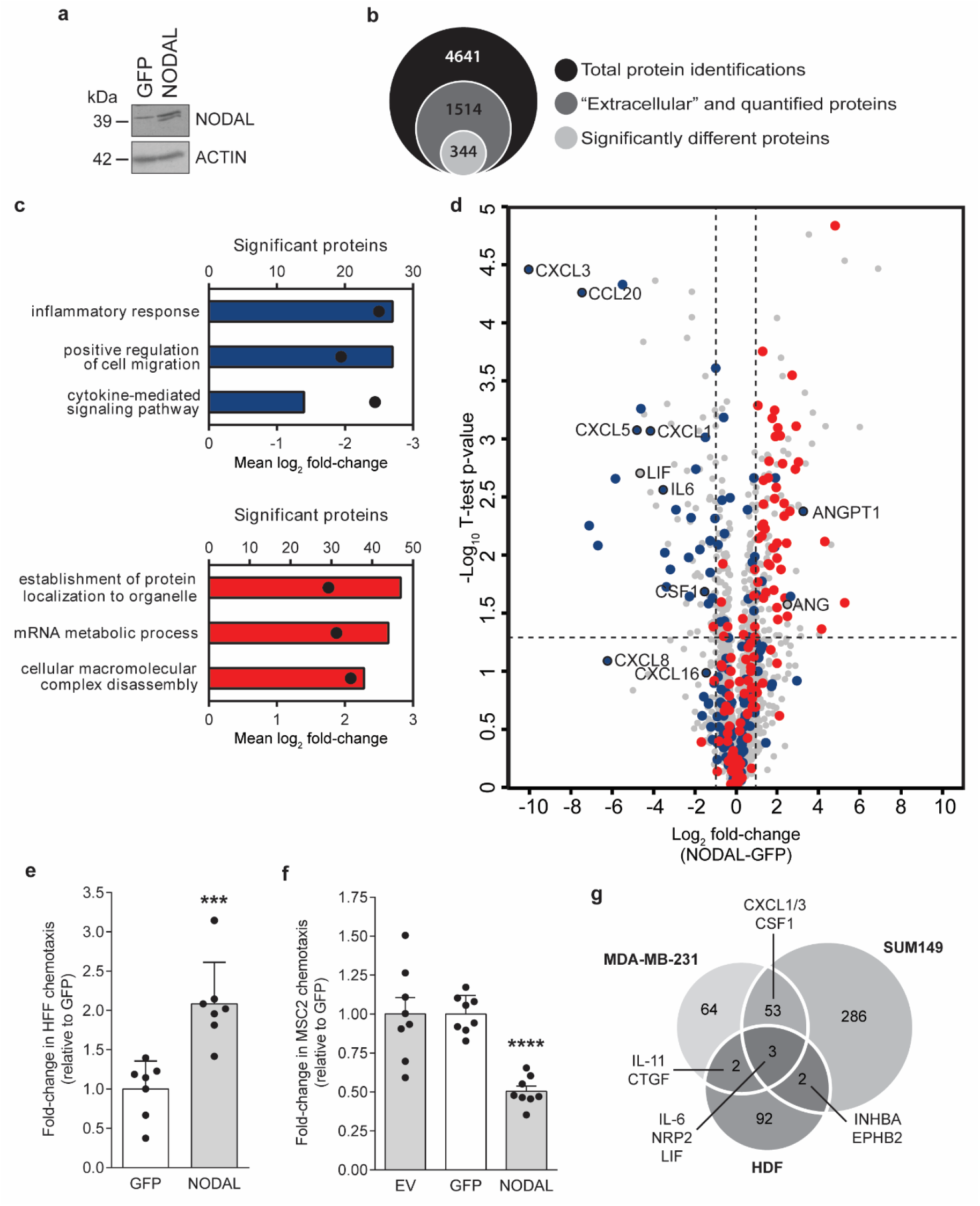
NODAL overexpression alters the SUM149 secretome and affects HFF and MSC chemotaxis. **(a)** NODAL expression in GFP- or NODAL-overexpressing SUM149 breast cancer cells. Extracellular proteins from serum-free CM (GFP or NODAL) were analyzed by mass spectrometry. **(b)** Venn diagram highlighting total protein identifications, “extracellular” and quantified proteins, and significantly different proteins between GFP and NODAL CM (two-tailed, two sample t-test, p<0.05). **(c)** Number of significant proteins (bars) matching to subset of significantly enriched GOBPs (BH FDR threshold<0.02). Mean Log_2_ fold-changes in GOBPs are indicated by black dots. Blue and red bars highlight GOBPs decreased and increased in SUM149 CM following NODAL overexpression, respectively. **(d)** Volcano plot of quantified “extracellular” proteins. Negative and positive Log_2_ fold-changes indicate proteins decreased and increased in SUM149 CM following NODAL overexpression, respectively (n=3). All proteins matching to corresponding GOBPs mentioned are highlighted in blue and red. Several cytokines, chemokines and growth factors altered by NODAL are labelled in black. Vertical and horizontal dotted lines indicate Log_2_ foldchanges ≥2 and the −log_10_ p-value cut-off corresponding to p<0.05, respectively. **(e)** NODAL overexpression increased HFF chemotaxis towards CM from SUM149 cells compared to GFP control (n=6); Two-sample t-test, ***p<0.001. **(f)** CM from NODAL-overexpressing SUM149 cells decreased MSC chemotaxis compared to empty vector (EV) and GFP controls. Data are presented as mean fold-changes relative to controls from a minimum of three biological replicates ± SD. Black dots indicate replicate values and asterisks indicate significance differences (one-way ANOVA, Dunnett’s multiple comparison test) in MSC chemotaxis compared controls (*** p<0.001, **** p<0.0001). **(g)** Overlap in proteins differentially expressed (increased or decreased) in MDA-MB-231 (shControl versus shNODAL), SUM149 (NODAL versus GFP) and HDF (rhNODAL-treated versus untreated) datasets.

In accordance with NODAL’s ability to directly activate fibroblasts, CM from NODAL overexpressing SUM149 cells significantly increased HFF chemotaxis compared to the GFP expressing control (**Fig. 7e**). Furthermore, in line with results suggesting that NODAL affects MSC chemotaxis indirectly by reprogramming the breast cancer secretome, CM derived from NODAL overexpressing SUM149 cells induced less chemotaxis in MSC2 cells compared to CM from the GFP expressing control cells (**Fig. 7f**). This reduced chemotaxis is likely attributable to lower levels of cytokines such as IL-6. Again, we confirmed by flow cytometry that differences in chemotaxis were not due to altered proliferation or viability (**Sup. Fig. 1b**; data not shown).

In a Venn diagram to assess relationships among differentially expressed proteins from MDA-MB-231, HDF, and SUM149 proteomics and microarray datasets, we found several factors consistently altered by NODAL, albeit some inversely correlated with NODAL levels (**Fig. 7g**). IL-6, LIF, and NRP2 were shared amongst all three datasets; however, CXCL1/3 appeared to be exclusively modulated in breast cancer cells. Hence, while NODAL indirectly affects MSC chemotaxis by altering the breast cancer secretome, NODAL can directly induce fibroblast activation. Moreover, certain key factors, such as IL-6 and LIF, are commonly affected by NODAL in all cell types investigated here.

### Differential signalling pathways may dictate cell type-dependent effects of NODAL

Our data demonstrate that the effects of NODAL are highly context-dependent. These differential responses may be due, in part, to which signal transduction pathways NODAL induces. Indeed, ELISAs showed that CXCL1 and IL-6 levels were substantially higher in GFP expressing SUM149 cells compared to MDA-MB-231 cell lines grown to similar confluence (**Sup. Fig. 2b, c**), suggesting different regulatory mechanisms between these cell lines. Canonically, NODAL triggers phosphorylation of SMAD2/3 via binding to its receptors ActRIIB/ALK(4/7) and co-receptor CRIPTO^67^. Phospho-SMAD2/3-SMAD4 heterodimers subsequently translocate into the nucleus to regulate the epigenetic status and transcription of target genes. NODAL can also signal non-canonically to activate ERK1/2, which is required for the induction of epithelial-mesenchymal transition (EMT) and invasion^49^.

Given the disparate effects of NODAL on cytokine secretion in MDA-MB-231 versus SUM149 cells, we hypothesized that NODAL may activate different signalling mediators like TGF-β in a cell-type dependent manner^68,69^. Accordingly, the activation of two documented mediators of NODAL signalling (SMAD2/3 and ERK1/2) were measured by western blotting in breast cancer cells wherein NODAL levels had been modified (**Fig. 8**). NODAL knockdown in MDA-MB-231 resulted in an expected and previously described reduction in both SMAD2/3 and ERK1/2 phosphorylation^49^. While overexpression of NODAL in SUM149 increased SMAD2 phosphorylation, a small reduction in ERK1/2 phosphorylation was observed. Moreover, constitutive SMAD2 activation was higher, while ERK1/2 was lower in SUM149 compared to MDA-MB-231 cells. Finally, we examined STAT3 activation, which occurs downstream of IL-6. In accordance with reduced IL-6 levels, NODAL knockdown reduced STAT3 phosphorylation in MDA-MB-231 cells. However, NODAL overexpression did not appear to affect STAT3 activation in SUM149 cells, perhaps due to the highly inflammatory nature of this cell line. These results suggest that some of the observed cell-type-specific effects of NODAL may relate to differential levels of SMAD2/3 and ERK1/2 activation induced by this ligand; MDA-MB-231 cells have higher ERK1/2 and lower SMAD2/3 basal activation compared to SUM149 cells. Hence, NODAL may preferentially signal through SMAD2/3 in SUM149 and ERK1/2 in MDA-MB-231 cells.

**Figure 8.**
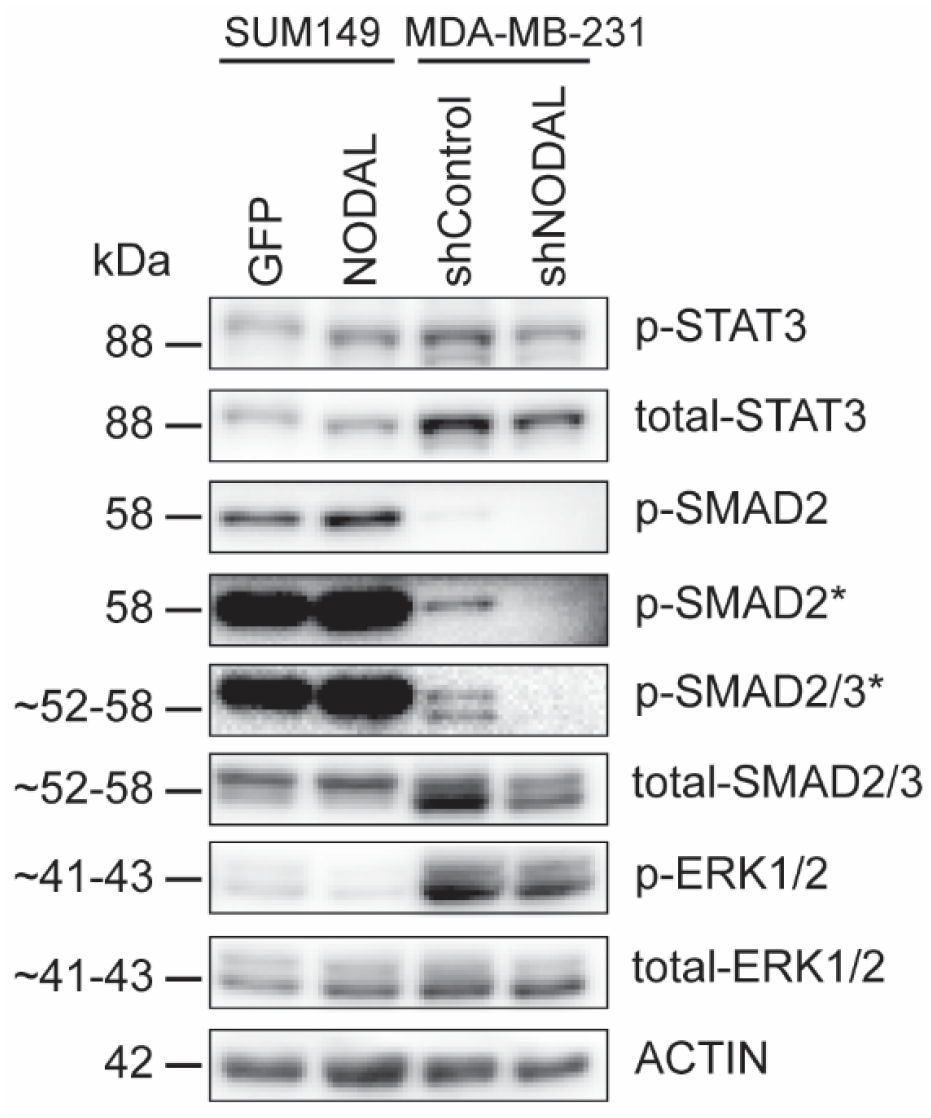
Effects of NODAL manipulation on signalling pathways in MDA-MB-231 and SUM149 cells. Western blotting revealed similarities and differences in activation of downstream pathways. NODAL expression (NODAL and shControl cell lines) was associated with increased phosphorylation of STAT3, SMAD2 and SMAD3. Basal levels of p-SMAD2 and p-ERK1/2 were substantially higher in SUM149 and MDA-MB-231 cell lines, respectively. p-ERK1/2 decreased slightly following NODAL overexpression in SUM149 cells. Western blots are representative images taken from three biological replicates and asterisks denote high contrast image settings.

## Discussion

The TME facilitates pro-tumourigenic processes, among which fibroblast activation – a common trait of many cancers, including breast carcinomas^70,71^ – plays an important role. Many factors activate fibroblasts, such as TGF-β, CXCL12/SDF-1, PDGFA/B, and IL-6^72–74^. NODAL has been shown to directly induce migration and invasion of breast, pancreatic, and hepatocellular cancer cell lines *in vitro*^43,49,75^. Moreover, ectopic overexpression of NODAL in breast cancer cells indirectly promotes endothelial tube formation by increasing the expression of pro-angiogenic proteins such as PDGFA^45^. Recent efforts have shown that NODAL alters breast cancer cell susceptibility to γδ T cell killing by acting on cancer cells to decrease recognizable antigens on the cell surface. Furthermore, long-term NODAL stimulation reduced Vδ2 T cell antigen receptor expression, suggesting activation of an as-of-yet unidentified signaling pathway in primary γδ T cells^50^. We build upon these studies by showing that NODAL may affect TME function and composition directly or indirectly by broadly regulating the breast cancer secretome. Specifically, we show that NODAL activates fibroblasts directly, but that it affects MSC chemotaxis indirectly, by reprogramming breast cancer cell secretomes.

We have shown herein that cells expressing the CAF marker α-SMA are spatially correlated with NODAL-positive cancer cells in human TNBC tissues, and that NODAL expression levels positively correlate to those of stromal α-SMA in these tissues. We demonstrate for the first time that NODAL signals directly on fibroblasts to induce an activated phenotype, characterized by increased proliferation rates, invasive capacity, and the expression of transcripts of known CAF markers. The origin of CAFs has been extensively debated over the years^76–79^, with evidence pointing to diverse sources such as resident tissue fibroblasts, bone marrow-derived MSCs, hematopoietic stem cells, epithelial cells that undergo epithelial-mesenchymal transition, and endothelial cells. A recent study has shown that the MDA-MB-231, but not MCF-7, secretome activates MSCs, converting them into tumour-associated MSCs^80^. We similarly show that although MSCs are unable to sense and respond to NODAL signals, they still undergo chemotaxis toward the NODAL-regulated breast cancer cell secretome.

Our robust proteomics approach allowed us to uncover dozens of secreted proteins that are affected by NODAL expression in breast cancer cells and may impact MSC recruitment to the breast TME. For these studies, we knocked down NODAL in claudin-low MDA-MB-231 cells that express high basal levels of NODAL, and overexpressed NODAL in SUM149, which represent inflammatory breast cancer cells and express low levels of NODAL. Consistent with the effects of NODAL *in vitro* and *in vivo*, the levels of several pro-angiogenic factors (PDGFA, ANGPT1, and ANG) in breast cancer CM were positively correlated with its expression^45^. However, we also made the seemingly paradoxical discovery that the expression of NODAL in MDA-MB-231 and SUM149 breast cancer cells oppositely regulates cytokines involved in chemotaxis. This difference may be coincident with the models chosen: MDA-MB-231 express relatively low levels of pro-inflammatory cytokines compared to SUM149, and thus the epigenetic regulation of the genes encoding these proteins may vary dramatically between the two cells lines.

Our discordant results are not uncommon for studies involving members of the TGF-β family, which function in a context-dependent manner. TGF-β1, for example, induces IL-6 production in PC3 and DU145 prostate cancer cells *via* SMAD2/TGFBRII and p38 MAPK^81^. Moreover, in MDA-MB-231 and MDA-MB-468 breast cancer cells, TGF-β1 stimulates IL-8 (CXCL8) and IL-11 secretion *via* SMAD3/TGFBRI and p38 MAPK^82^. However, in the Polyomavirus middle T antigen transformed mouse mammary carcinoma model, loss of TGF-β signalling results in the upregulation of CXCL1, CXCL5, and CCL20^83^. Remarkably, these factors decreased substantially in SUM149 CM following NODAL overexpression (**Sup. Tables 4 and 6**), thus suggesting negative regulatory roles for both NODAL and TGF-β. We did not observe significant differences in the levels of TGF-β1/2 between breast cancer lines (Sup. Tables 2, 4 and 6), hence the effects of NODAL were not likely mediated *via* alterations in TGF-β1/2. Taken together, both NODAL and TGF-β may differentially regulate chemokine and cytokine expression in cancer, depending on the context. This difference should be considered as treatment modalities designed to target these pathways evolve^84^.

Genes regulated by NODAL appear to be dictated, at least in part, by the accessibility of genomic regions, and NODAL induces histone modifications to affect gene expression^85^. Hence the differential effects of NODAL in MDA-MB-231 versus SUM149 cells may be due to differences in chromatin accessibility in the areas surrounding chemotactic and inflammatory cytokines. The differences observed may also be due to the ability of NODAL to activate ERK signaling in MDA-MB-231 cells but not in SUM149 cells. Several studies have demonstrated the role of ERK signaling in the upregulation of inflammatory cytokines such as IL-6^86,87^. Hence the effects of NODAL knockdown in MDA-MB-231 cells may be due to reduced ERK signalling.

While IL-6R was detected on MSC, CXCR1 and CXCR2 were not. Heterogeneity in MSC receptor expression has been reported in multiple studies and may be a product of culture conditions and donor heterogeneity^88,89^. For reference, Ponte *et al*. observed CXCR4 and CXCR5 but not CXCR1 or CXCR2 on human BM-MSC^90^. Chamberlain *et al*. also reported high expression for CXCR4 and CXCR5 but low to intermediate expression of CXCR1 and CXCR2, respectively^91^. Conversely, Ringe *et al*. extensively profiled chemokine receptors on human BM-MSC and detected CXCR1 and CXCR2 but noted loss of expression following ten passages^92^. While these pathways may play a role in MSC recruitment to tumours in breast cancer patients, we were unable to test this possibility.

In our hands, MDA-MB-231 cells produced less IL-6 and CXCL1 than those studied by Hartman *et al*., who investigated the role of cytokines in TNBC cell growth^60^. Notwithstanding, neutralizing IL-6 in MDA-MB-231 CM was sufficient to attenuate MSC chemotaxis^29,93,94^. We did not neutralize IL-6 in SUM149 CM; however, CM from either SUM149 or SUM159 breast cancer cells was previously shown to promote migration of aldehyde dehydrogenase-high MSC or macrophage-educated MSC in an IL-6 dependent manner^62,94^.

Although CXCR1/2 was not detected on MSC, differences in CXCL1 and CXCL8 levels following NODAL knockdown/overexpression remain important for cancer progression and trafficking of additional cell types and justify additional interrogation. For instance, CXCL1-mediated recruitment of CD11b+Gr1+ myeloid cells enhanced breast cancer cell survival, chemoresistance, and metastasis^61^. Moreover, obesity-associated CXCL1 expression in prostate tumours was linked to adipose-derived stromal cell migration *in vitro* and tumour engraftment *in vivo*^95^. Given the importance of NODAL-regulated cytokines in the TME, future studies interrogating the extent to which NODAL may modulate TME composition are warranted.

In summary, we demonstrate that NODAL directly activates stromal fibroblasts and that it reshapes the breast cancer secretome, affecting the deposition of factors such as IL-6, which may regulate the recruitment of MSCs, as well as other TME cell types (**Fig. 9**). Expanding our previous discovery that NODAL induces secretion of PDGF and VEGF by breast cancer cells, our present findings illuminate a hitherto unappreciated role for NODAL in the orchestration of the tumour microenvironment.

**Figure 9.**
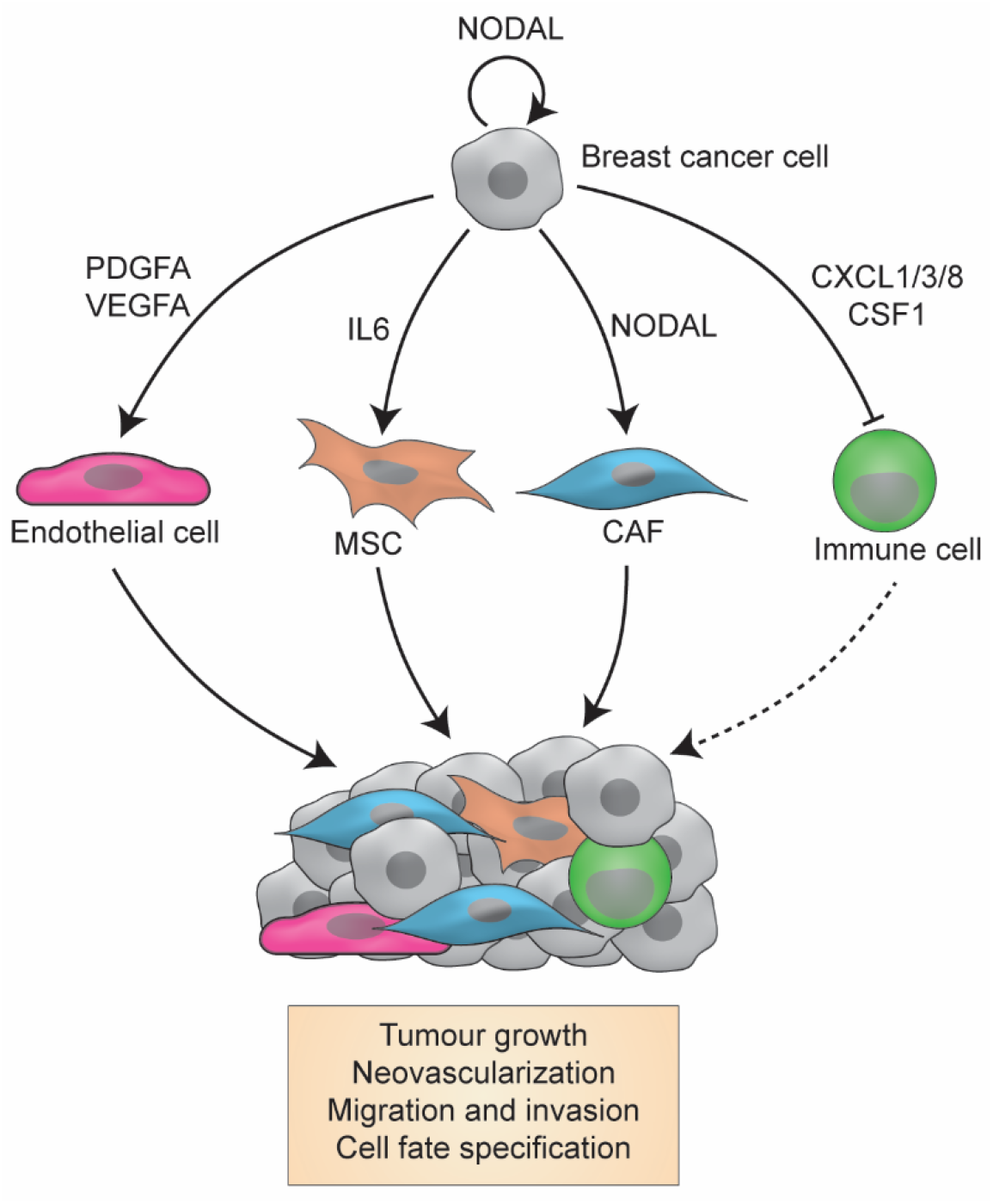
Proposed model for NODAL signalling in the breast cancer microenvironment. NODAL signals directly to breast cancer cells and CAFs, and indirectly regulates secretion of inflammatory, chemotactic and angiogenic factors by breast cancer cells, which act on endothelial and mesenchymal stromal cells and possibly immune cell types. Collectively, NODAL promotes tumorigenic phenotypes including tumour growth, neovascularization, cell migration and cell fate specification.

## Materials and Methods

### Patients and Tissues

We assessed 41 samples from 20 surgically resected TNBC tumors from cancer patients diagnosed at the Cross Cancer Institute, Edmonton, AB in 2017. This study was carried out in accordance with the recommendations of the Research Ethics Guidelines, Health Research Ethics Board of Alberta – Cancer Committee with written informed consent from all subjects. All subjects gave written informed consent in accordance with the Declaration of Helsinki.

### Immunohistochemistry

NODAL and α-SMA staining in TNBC tissues was performed as previously described for NODAL^45^. Briefly, tissue sections were deparaffinized in xylene and hydrated in graded ethanol. Antigen retrieval was performed with citrate buffer pH 6.0, followed by peroxidase and serum-free protein blocking. After incubation with primary antibodies (**Sup. Table 8**), slides were rinsed in TBS-T and treated with Envison+ HRP anti-mouse IgG (Dako, Glostrup, Denmark). Color was produced with 3,3’-Diaminobenzidine (DAB) substrate and counterstained with Mayer’s haematoxylin. Samples were dehydrated in graded alcohol and cover slipped with permanent mounting medium.

### Evaluation of NODAL and α-SMA staining

Light microscopy and semi-quantitative scoring were performed by two pathologists. The entirety of each slide was assessed. Scores for NODAL were 0, absent; 1, weak or very focal staining; 2, strong but focal or moderate intensity; and 3, strong and extensive staining. The score reflects the intensity of staining observed in the majority of cells. When scored 1-3, NODAL distribution was further identified as focal, diffuse or scattered, and an estimated proportion of tumour cells staining with NODAL was calculated (NODAL percentage in Table I). α-SMA was scored in the same manner on serial sections from the same cases. Intensity association was measured based on the extent to which α-SMA staining was increased in areas with NODAL-positive cells. Representative images were taken from a Nikon DS U3 camera on Nikon eclipse 80i microscrope (Nikon, Tokyo, Japan) at 400 x (500 px bar = 40 μm).

### Cell culture

MDA-MB-231 cells stably expressing scrambled control (shControl) or NODAL targeting (shNODAL) short hairpin RNAs as previously described and validated^45,48,49^ were maintained in DMEM/F12 (Gibco, Waltham, MA) supplemented with 10% FBS (Gibco) and 500ng/mL puromycin. To generate SUM149 cells stably expressing an empty vector (EV), green fluorescent protein (GFP) or NODAL, cells were transduced with lentiviral particles (GeneCopoeia, Rockville, MD) overnight then selected and maintained in HAM’s F10 (Gibco) supplemented with 10% FBS, 5μg/mL insulin (Santa Cruz Biotechnology, Dallas, TX), 1μg/mL hydrocortisone (Sigma-Aldrich, St. Louis, MO) and 100 ng/mL puromycin. Human BM-MSC lines were maintained in AmnioMAX™ with C100 supplement (Life Technologies, Carlsbad, CA); these lines were previously confirmed to express characteristic stromal markers (>95% CD90+, CD105+, and CD73+) and exhibit multipotent differentiation^56,96^. HFFs (Cascade Biologics, Portland, OR) were maintained in DMEM/F12 supplemented with 10% FBS and HDFs (ATCC, Manassas, VA) in DMEM supplemented with 10% FBS. For SILAC labelling, shControl and shNODAL MDA-MB-231 cells were cultured in DMEM F12 supplemented with dialyzed FBS (Life Technologies) containing light (Advanced ChemTech, Louisville, KY) or heavy (Cambridge Isotope Laboratories, Tewksbury, MA and Silantes GmbH, Germany) isotopes of arginine (0.398mM) and lysine (0.274mM) for at least nine days to achieve >90% label incorporation. SILAC media was additionally supplemented with 400 mg/L of proline (Sigma-Aldrich) to limit arginine to proline conversion^97^. CM was prepared by plating equal cell numbers onto flasks in culture media (Corning, Corning, NY). After 24h (MDA-MB-231 cells) or 48h (SUM149 cells), media was removed, and cells were thoroughly rinsed three times in PBS (with Ca2+ and Mg2+) to remove serum components. Cells were incubated in serum-free media (SFM) with 0.5% BSA for an additional 24h to generate CM (BSA was omitted for LC-MS samples). Conditions used to stimulate cells with rhNODAL and rhIL-6 are specified in the main text.

### Sample preparation for liquid chromatography-mass spectrometry (LC-MS)

CM (without BSA) were concentrated using 3 kDa molecular weight cut-off (MWCO) Amicon ultracentrifugal units (Millipore, Burlington, MA) and lyophilized overnight. The following day, CM was reconstituted in lysis buffer (8M urea, 50mM ammonium bicarbonate, 10mM dithiothreitol and 2% SDS), sonicated (3 × 0.5s pulses) with a probe sonicator (Level 1; Fisher Scientific, Waltham, MA) and quantified using a Pierce™ 660 nm assay (Thermo Fisher Scientific, Waltham, MA) with ionic detergent compatibility reagent. For SlLAC samples, light shControl and heavy shNODAL CM were pooled based on equal cell numbers and ~100μg protein were fractionated using SDS-PAGE on 12% acrylamide tris-glycine gels. In-gel digestion with trypsin (1:25 enzyme: protein ratio) was performed on 16-17 slices (fractions) from each lane in biological triplicate as previously described^98^. For label-free samples, ~50μg protein from SUM149 CM were precipitated in chloroform/methanol, digested overnight with trypsin (1:50 ratio) on a water bath shaker and fractionated on SCX StageTips as previously described^98–100^. Peptides were dried in a SpeedVac, reconstituted in 0.1% formic acid (FA; Fisher Scientific) and a volume corresponding to 1/10th of the total material recovered or 1 μg as determined by bicinchoninic acid (BCA) assay (Pierce™, Waltham, MA) was injected for each in-gel and SCX fraction, respectively.

### LC-MS

In-gel and SCX fractions were analyzed using a Q Exactive or Orbitrap Elite mass spectrometer (Thermo Fisher Scientific), respectively. Samples were injected using a nanoAcquity HPLC system (Waters, Milford, MA) and initially trapped on a Symmetry C18 Trap Column (5 μm, 180 μm x 20 mm) for 4 or 5 minutes in 99% Solvent A (Water/0.1% FA)/1% Solvent B (acetonitrile/0.1% FA) at a flow rate of 10 μl/min. Peptides were separated on an ACQUITY Peptide BEH C18 Column (130Å, 1.7μm, 75μm X 250mm) at a flow rate of 300 nL/min maintained at 35°C. The LC-MS gradient for in-gel digests consisted of 1-7% B over 1 minute and 7-37.5% B over 79 minutes. SCX fractions were separated using gradient consisting of 7.5% B over 1 minute, 25% B over 179 minutes, 32.5% B over 40 minutes and 60% B over 20 minutes. Column washing and re-equilibration was performed following each run and settings for data acquisition are outlined in **Sup. Table 9**.

### Data analysis and statistics

Raw MS files were searched in MaxQuant (1.5.2.8) with the Human Uniprot database (reviewed only; updated May 2014 with 40,550 entries)^101^. Missed cleavages were set to 3 and I=L. Cysteine carbamidomethylation was set as a fixed modification. Oxidation (M), n-terminal acetylation (protein), and deamidation (NQ) were used as variable modifications (max. number of modifications per peptide = 5) and min ratio count was set to 1. All other settings were left at default. The match-between-runs feature was utilized to maximize proteome coverage and quantitation between samples. Datasets were loaded into Perseus^59^ (version 1.5.5.3) and proteins identified by site, reverse and potential contaminants were removed^59^. Protein identifications with quantitative values in ≥2 biological replicates were retained for downstream analysis unless specified elsewhere. Missing values were imputed using a width of 0.3 and down shift of 1.8 for label free datasets. Statistical analysis was performed in Perseus or GraphPad Prism version 6.01 (San Diego, CA). All experiments were carried in at least three biological replicates unless specified otherwise. Where specified, replicate treatment values were normalised to the control group and relative fold-changes were reported. Two-tailed, one sample and two-sample t-tests (p<0.05) were performed to determine statistical differences unless more than two conditions were being compared and a one-way ANOVA using Dunnett’s multiple comparison test (p<0.05) was performed instead.

### Chemotaxis and invasion assays

MSCs were rinsed in warm PBS (with Ca2+ and Mg2+) and serum starved for ~2h in AmnioMAX™ prior to dissociation with trypsin for chemotaxis assays. In parallel, 8μM transwells (Falcon^®^, Corning, NY) were coated with 10μg/cm^2^ of bovine fibronectin (Sigma-Aldrich) in 100μL of PBS for 2h. After coating, excess solution was aspirated and 40K MSCs in 0.5mL of DMEM F12+0.5% BSA were plated in each transwell. HFFs were serum starved 24h prior to dissociation and plated at a density of 50K cells/transwell. For HFF chemotaxis and invasion assays, fibronectin and Matrigel™ (Corning) were omitted and included, respectively. To the bottom chamber, 1mL of DMEM/F12 + 0.5% BSA or CM was added +/- rhNODAL (R&D Systems, Minneapolis, MN), rhIL-6 (eBioscience, San Diego, CA), isotype or IL-6 neutralizing monoclonal antibodies (R&D Systems). After ~24h, transwells were rinsed in warm PBS and placed in cold methanol for 20 minutes to fix migrating cells. After fixing, transwells were rinsed in PBS and the inside membrane was thoroughly wiped with a cotton swab to remove non-migrated cells. Membranes were excised and mounted onto glass slides with ProLong™ Gold Antifade Mountant with DAPI (Invitrogen, Carlsbad, CA). Migrated cells were counted from at least 5-10 high power fields uniformly distributed across the entire membrane for each condition.

### Western blotting

Cells were thoroughly washed with PBS (with Ca2+ and Mg2+) and directly lysed on tissue culture plates in lysis buffer. Lysates recovered by pipetting were sonicated with a probe sonicator (20 × 0.5s pulses) to shear DNA and reduce viscosity. Equal protein amounts (15-25μg) were separated on hand cast 8-20% acrylamide Tris-glycine gels then transferred to Immobilon-P^®^ PVDF membranes (Millipore). Membranes were stained with amido black and rinsed in ddH2O for 5 minutes followed by blocking for 1h on rocker in 5% non-fat dry milk in TBST (Tris-buffered saline, 0.1% Tween 20) and overnight incubation in primary antibody at 4°C. Chemiluminescent detection was performed using film or a VersaDoc CCD camera with Clarity™ Western ECL Substrate and horseradish peroxidase-conjugated secondary antibodies (Bio-Rad, Hercules, CA) the next day. Antibody information is available in **Sup. Table 8**. Actin and Tubulin were used as loading controls. PVDF membranes were stippled in 0.2 M NaOH and re-probed when possible, otherwise western blots were run in duplicate.

### Real-time PCR

RNA was isolated from cells and treated with DNAse using a PerfectPure RNA cultured cell kit (5PRIME). RNA was quantified by NanoDrop™ (Thermo Fisher Scientific) and 2μg was reverse transcribed with a High Capacity cDNA Reverse Transcription kit (Applied Biosystems, Foster City, CA). Real-time PCR was performed with TaqMan™ Universal PCR Master Mix (Applied Biosystems) on a Bio-Rad CFX96/384 thermocycler. *HPRT1* or *RPLPO* were used as housekeeping genes to monitor variations between biological replicates. TaqMan™ primer probes were purchased from Applied Biosystems and are listed in **Sup. Table 10**.

### Flow cytometry

MSCs dissociated in 10mM EDTA/PBS solution for 5-10 minutes were resuspended in 5% FBS/PBS, counted, and pelleted at 450xg. Excess buffer was aspirated and MSCs were divided into 50-100K cell aliquots in 100μL of 5% FBS/PBS. Isotype controls and primary antibodies (**Sup. Table 8**) were added to cell suspensions and incubated for ~45 minutes in the dark on ice. Cell suspensions were washed in excess 5% FBS/PBS and pelleted to remove unbound antibody. Flow cytometry data was acquired on an LSR II (Becton Dickinson, Franklin Lakes, NJ) using FACSDiva (Becton Dickinson) at the London Regional Flow Cytometry Facility and analyzed with FlowJo (FlowJo LLC, Ashland, OR, Version 10.0.8r1). The gating strategy for live singlets was based on forward and side-scatter and is illustrated in **Sup. Fig. 5**. The CellTrace™ Violet Cell Proliferation assay (Thermo Fisher Scientific) was performed as instructed by the manufacturer. Briefly, MSCs were labelled in suspension with CellTrace™ Violet dye. After 20 min incubation at 37°C in the dark, MSCs were incubated for 5 min with culture medium to remove any free dye remaining in the solution. MSCs were pelleted, resuspended in fresh pre-warmed complete culture medium, and plated onto 6-well plates prior to incubation with CM.

### ELISAs

ELISA kits were purchased from eBioscience (IL-6) or R&D Systems (CXCL1, CXCL8 and CSF1) and performed according to the manufacturer’s specifications using CM derived from MDA-MB-231 and SUM149 cell lines.

### Gene expression profiling

HDFs were cultured until ~40-60% confluence, washed twice with PBS and incubated overnight in DMEM+0.5%FBS. The following day, cells were treated +/- rhNODAL (10 ng/mL) for 6h and RNA was harvested using TRIzol™ (Invitrogen). RNA was subjected to expression profiling at the London Regional Genomics Centre essentially as previously described^102,103^. RNA quality was assessed using an Agilent 2100 Bioanalyzer (Agilent Technologies, Santa Clara, CA) prior to preparing single stranded complimentary DNA (sscDNA) from 200ng of total RNA (Ambion WT Expression Kit for Affymetrix GeneChip Whole Transcript WT Expression Arrays; Applied Biosystems, Carlsbad, CA) according to the Affymetrix User Manual (Affymetrix, Santa Clara, CA). In total, 5.5μg of sscDNA was synthesized, converted into cRNA, end labeled and hybridized (16h at 45°C) to Human Gene 1.0 ST arrays. Liquid handling steps were performed by a GeneChip Fluidics Station 450 and GeneChips were scanned (GeneChip Scanner 3000 7G; Affymetrix) using Command Console v1.1 to generate Probe level (.CEL file) data. Gene level data was generated using the RMA algorithm^104^. Partek Genomics Suite v6.5 (St. Louis, MO) was used to determine gene level ANOVA p-values and fold-changes. Fold-changes were obtained by averaging data from two experiments (GeneSpring, Agilent). Fold-changes exceeding 1.7 in response to rhNODAL were required to identify a transcript as being altered (p<0.05). Altered genes were annotated using DAVID (version 6.7) and lists enriched >3.5 fold and comprised of >10 genes were reported.

## Acknowledgements

We thank Dr. Dean Betts (Western University), Dr. John Di Guglielmo (Western University) and Dr. Dwayne Jackson (Western University) for providing access to PCR and imaging equipment and Paula Pittock for technical support. The work was funded by operating grants from the CIHR and the Canadian Breast Cancer Foundation awarded to LMP.

