## Supplementary Material for "Embryonic Protein NODAL Regulates the Breast Tumour Microenvironment by Reprogramming Cancer-Derived Secretomes"

### Supplementary Figures and Tables

Supplementary Figure 1

a

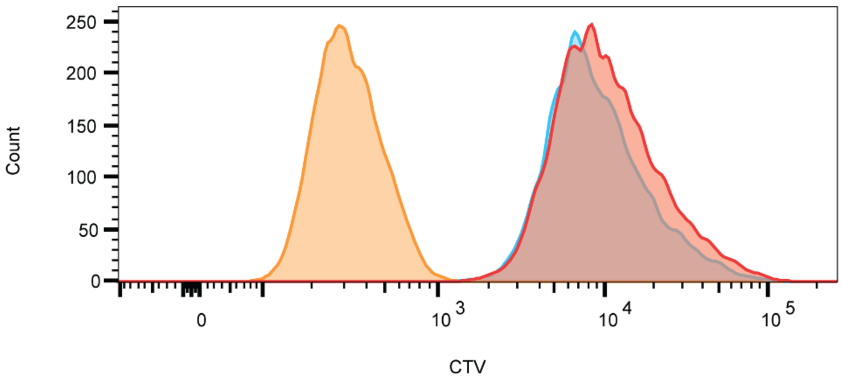


| | Sample Name | Median CTV intensity |
|---|---|---|
| | MSC in shControl CM | 9675 |
| | MSC in shNodal CM | 8326 |
| | Unstained MSC | 314 |

b

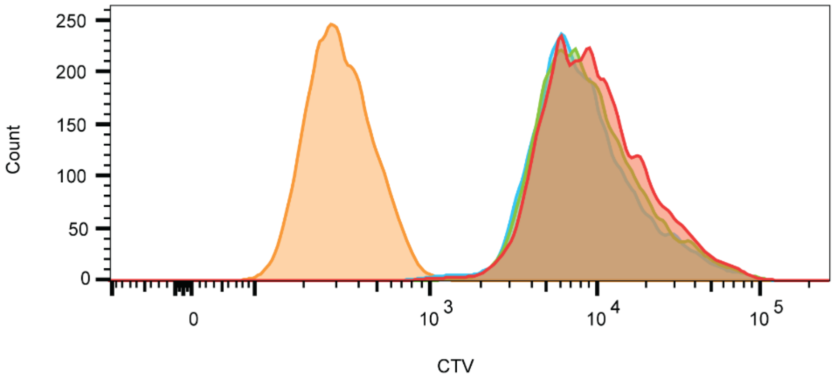


| | Sample Name | Median CTV intensity |
|---|---|---|
| | MSC in GFP CM | 9075 |
| | MSC in EV CM | 8245 |
| | MSC in NODAL CM | 7697 |
| | Unstained MSC | 314 |

**Supplementary Figure 1.** MDA-MB-231 and SUM149 CM have a negligible effect on MSC proliferation. MSCs were labelled with CellTrace Violet (CTV) in suspension and plated onto tissue culture plates. After ~48h, MSCs were rinsed in PBS and cultured for additional 24h in CM from MDA-MB-231 (shControl and shNODAL) or SUM149 (EV, GFP and NODAL) cells. MSCs were then trypsinized, pelleted and resuspended in 5% FBS/PBS. **(a, b)** Histograms and tables showing median CTV intensity of MSCs after

24h in CM from MDA-MB-231 and SUM149 conditions.  Histograms are representative images from three biological replicates.

Supplementary Figure 2

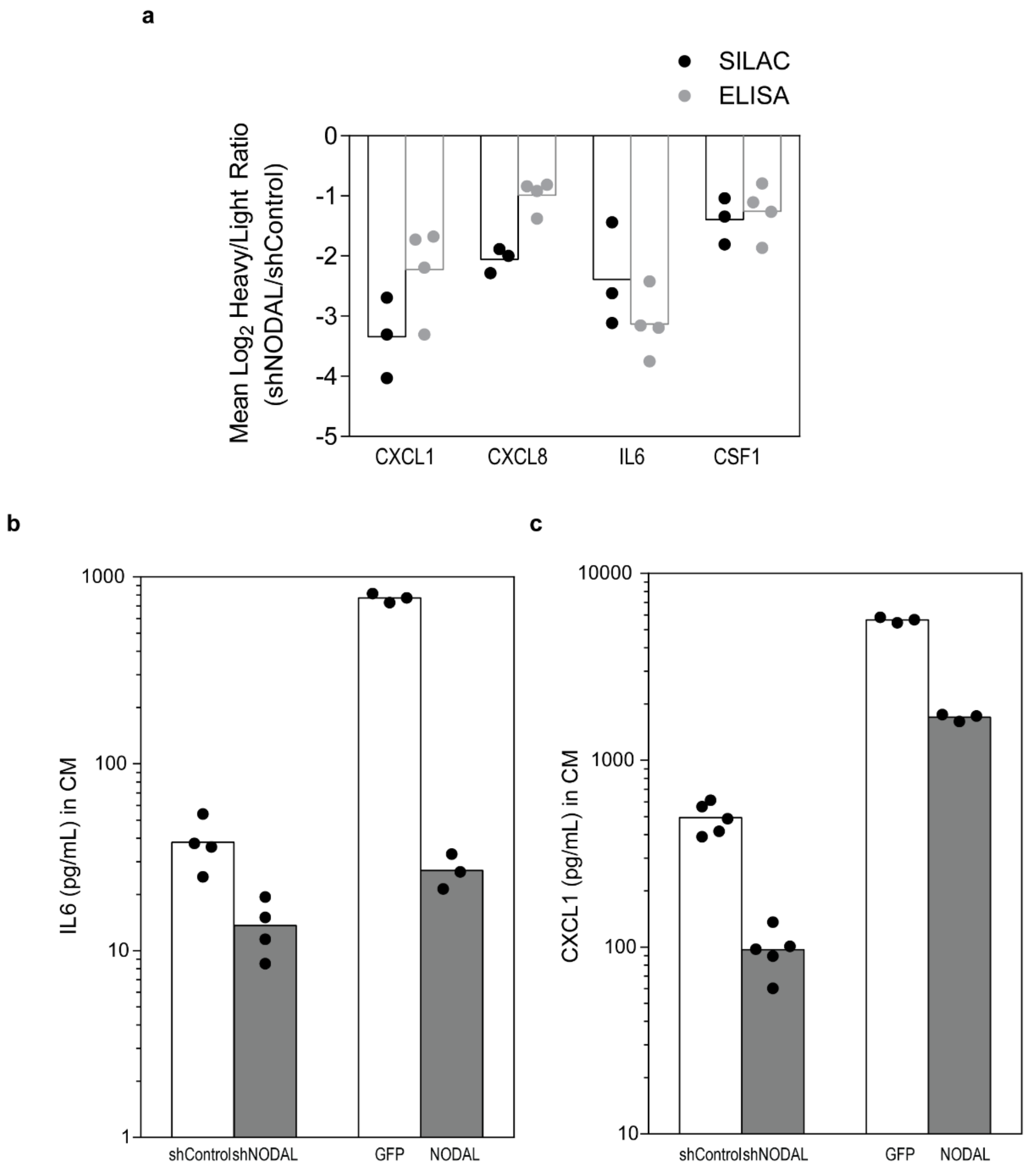


**Supplementary Figure 2.** Cytokine and chemokine levels in breast cancer CM. **(a)** SILAC quantification in MaxQuant and ELISAs report similar fold-changes in secreted factors between shControl and shNODAL MDA-MB-231 cells. **(b, c)** ELISAs reveal substantially higher levels of CXCL1 and IL-6 in CM derived from GFP expressing SUM149 cells compared to MDA-MB-231 cell lines. Black and grey dots indicate replicate values.

Supplementary Figure 3

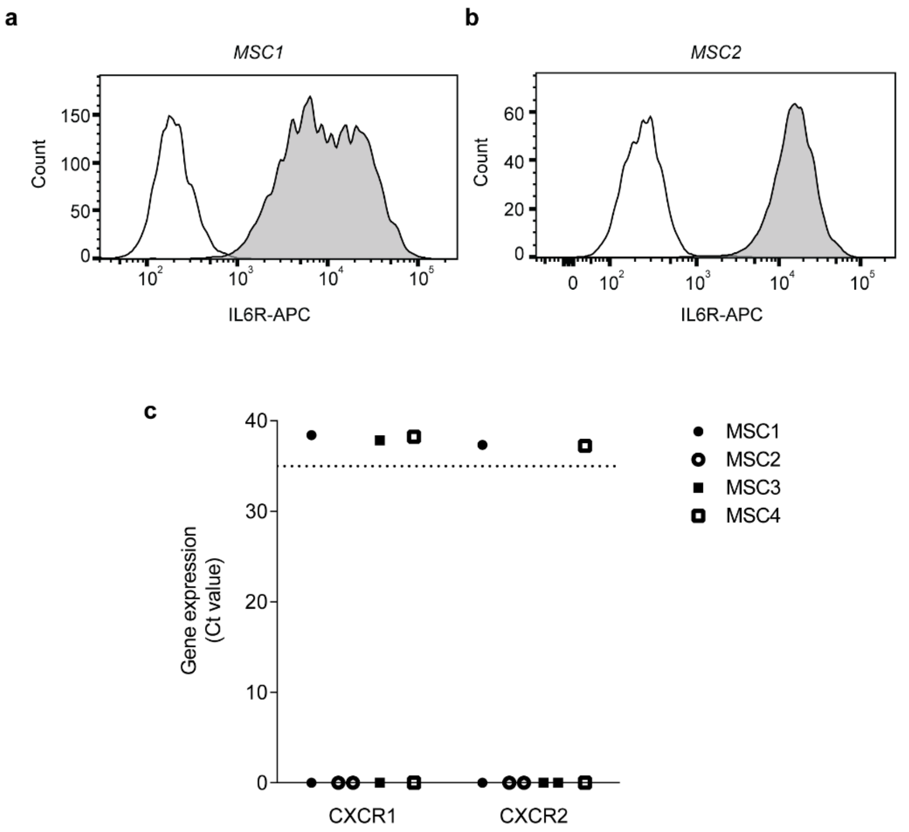


**Supplementary Figure 3.** MSCs express IL-6R but lack *CXCR1/2* gene expression. **(a, b)** Flow cytometry showing high surface expression of IL-6R (grey region) on two additional MSC lines compared to unstained MSCs (white region). **(c)** Real-time PCR results reveal no detectable transcript or reliable gene expression for *CXCR1* or *CXCR2* in four primary MSC lines. Individual data points correspond to real-time PCR readings from two biological replicates. A Ct value of 0 indicates no transcript was detected. The dotted line indicates a Ct value of 35 or the reliable limit of detection. Histograms are representative images from three biological replicates.

Supplementary Figure 4

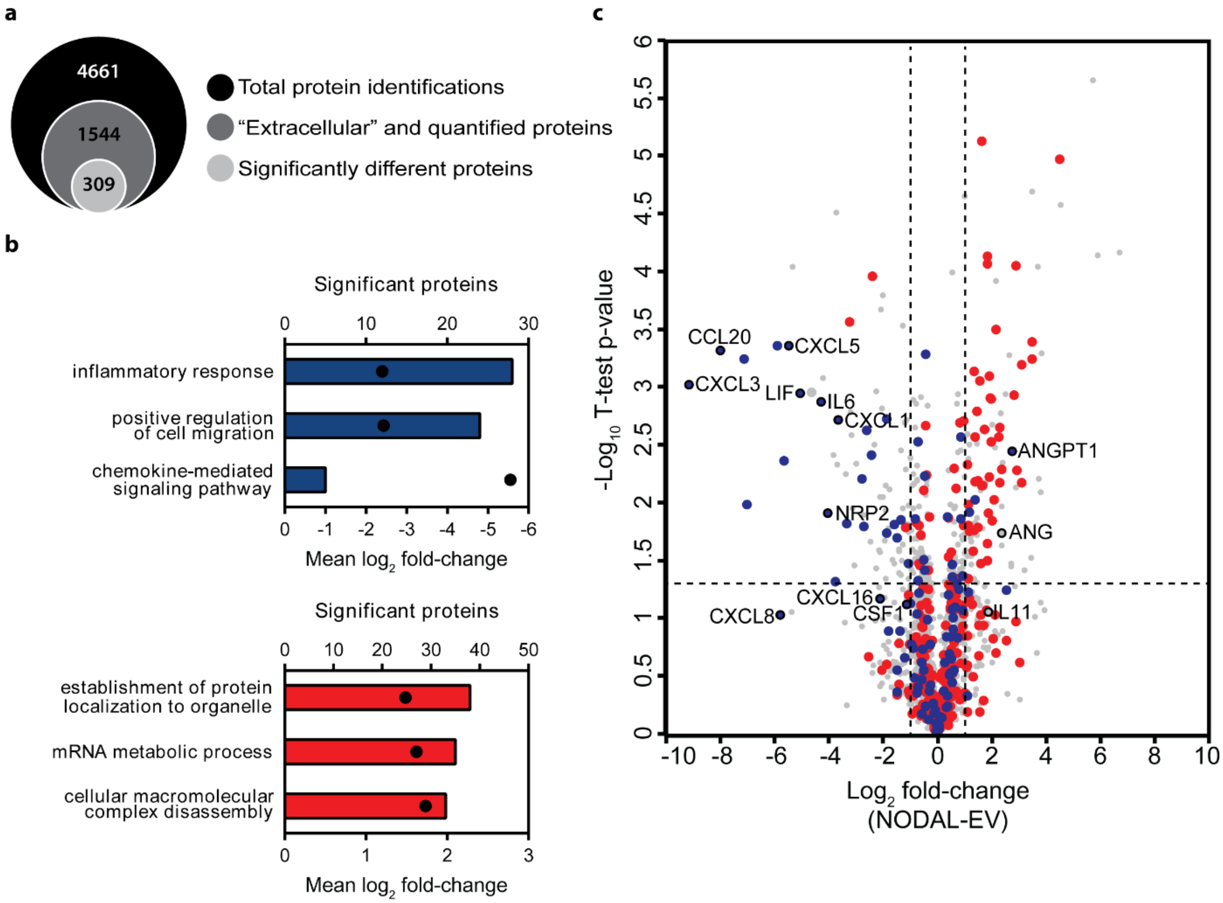


**Supplementary Figure 4.** NODAL overexpression alters the SUM149 secretome. Extracellular proteins (CM) from empty vector (EV) or Nodal overexpressing SUM149 cell lines were concentrated using 3kDa MWCO filter units, digested with trypsin, fractionated using SCX StageTips and analyzed on an Orbitrap Elite mass spectrometer. Raw files were searched with MaxQuant and protein lists were filtered and annotated in Perseus. **(a)** Venn diagram highlighting total protein identifications, "extracellular" and quantified proteins, and significantly different proteins between GFP and NODAL CM. A two-tailed, two sample t-test was used to identify differentially expressed proteins ($p<0.05$). **(c)** Number of significant proteins (bars) matching to significantly enriched GO biological processes (GOBPs). Mean $\log_2$ fold-changes in GOBPs are indicated by black dots. Blue and red bars highlight GOBPs decreased and increased in SUM149 CM following NODAL overexpression. **(d)** Volcano plot of quantified "extracellular" proteins. Negative and positive $\log_2$ fold-changes indicate proteins decreased and increased in SUM149 CM following NODAL overexpression, respectively (n=3). All proteins matching to corresponding GOBPs mentioned are highlighted in blue and red. Several cytokines, chemokines and growth factors altered by NODAL are labelled in black. Vertical and horizontal dotted lines indicate $log_2$ fold-changes ≥2 and the –$log_{10}$ p-value cut-off corresponding to $p<0.05$, respectively.

Supplementary Figure 5

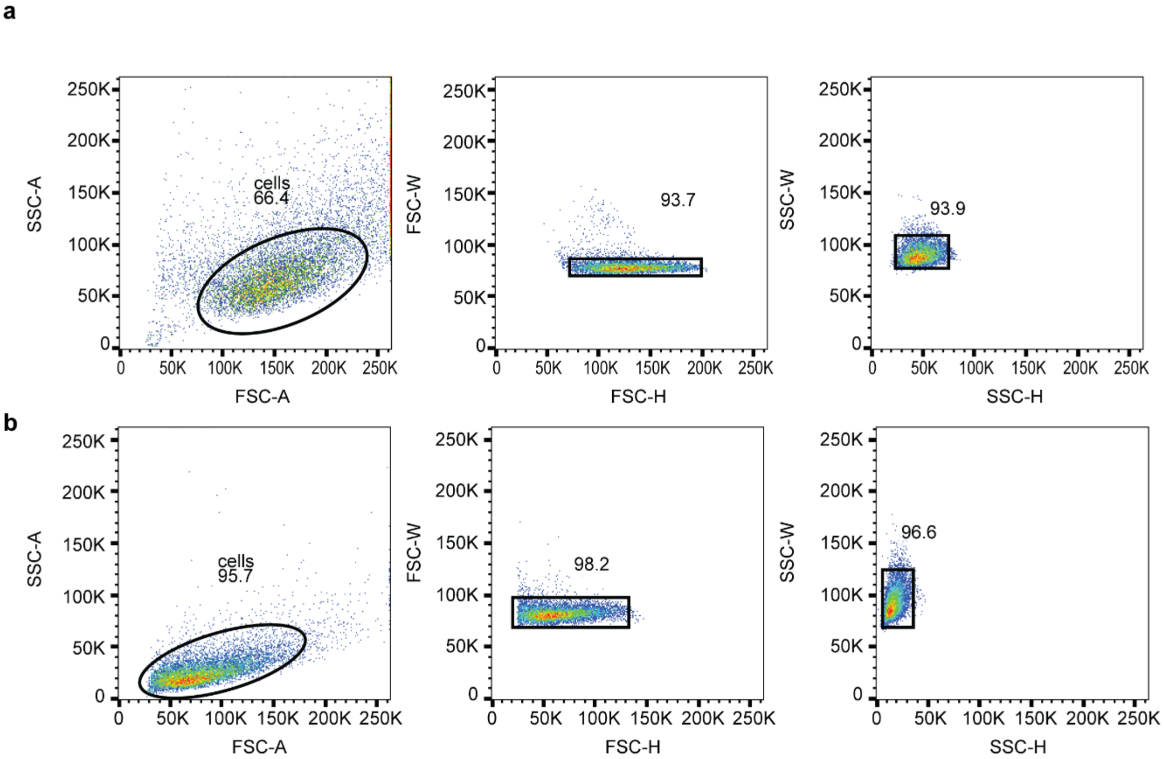


**Supplementary Figure 5.** Gating strategy for flow cytometry analysis. **(a, b)** Forward- and side-scatter gating method used to select live singlets for surface marker expression and CTV analyses.

**Sup. Table 1: Enriched gene clusters matching to transcripts elevated in HDFs following treatment with rhNODAL**

| extracellular matrix | enrichment 14.6 | | | | | | | |
|---|---|---|---|---|---|---|---|---|
| **yourlist:M20180608A7434721E10EE6586998A056CC** | **isomap:M20180608A7434721E10EE6586998A056CCD** | **Entry** | **Entry name** | **Status** | **Protein names** | **Gene names** | **Organism** | **Length** |
| NM_005099 | NM_005099 -> O75173-1 | O75173 | ATS4_HUMAN | reviewed | A disintegrin and metalloproteinase with thrombospondin motifs 4 (ADAM-TS 4) (ADAM | ADAMTS4 KIAA0688 UNQ769/P | Homo sapiens | 837 |
| NM_197941 | NM_197941 -> Q9UKP5-1 | Q9UKP5 | ATS6_HUMAN | reviewed | A disintegrin and metalloproteinase with thrombospondin motifs 6 (ADAM-TS 6) (ADAM | ADAMTS6 | Homo sapiens | 1117 |
| NM_004598 | | Q08629 | TICN1_HUMAN | reviewed | Testican-1 (Protein SPOCK) | SPOCK1 SPOCK TIC1 TICN1 | Homo sapiens | 439 |
| NM_000362 | | P35625 | TIMP3_HUMAN | reviewed | Metalloproteinase inhibitor 3 (Protein MIG-5) (Tissue inhibitor of metalloproteinases 3) ( | TIMP3 | Homo sapiens | 211 |
| NM_003391 | | P09544 | WNT2_HUMAN | reviewed | Protein Wnt-2 (Int-1-like protein 1) (Int-1-related protein) (IRP) | WNT2 INT1L1 IRP | Homo sapiens | 360 |
| NM_017680 | | Q9BXN1 | ASPN_HUMAN | reviewed | Asporin (Periodontal ligament-associated protein 1) (PLAP-1) | ASPN PLAP1 SLRR1C UNQ215/P | Homo sapiens | 380 |
| NM_003613 | | O75339 | CILP1_HUMAN | reviewed | Cartilage intermediate layer protein 1 (CILP-1) (Cartilage intermediate-layer protein) [Cle | CILP UNQ602/PRO1188 | Homo sapiens | 1184 |
| NM_000095 | NM_000095 -> P49747-1 | P49747 | COMP_HUMAN | reviewed | Cartilage oligomeric matrix protein (COMP) (Thrombospondin-5) (TSP5) | COMP | Homo sapiens | 757 |
| NM_138455 | NM_138455 -> Q96CG8-1 | Q96CG8 | CTHR1_HUMAN | reviewed | Collagen triple helix repeat-containing protein 1 (Protein NMTC1) | CTHRC1 UNQ762/PRO1550 | Homo sapiens | 243 |
| NM_001845 | | P02462 | CO4A1_HUMAN | reviewed | Collagen alpha-1(IV) chain [Cleaved into: Arresten] | COL4A1 | Homo sapiens | 1669 |
| NM_001846 | | P08572 | CO4A2_HUMAN | reviewed | Collagen alpha-2(IV) chain [Cleaved into: Canstatin] | COL4A2 | Homo sapiens | 1712 |
| NM_000093 | | P20908 | CO5A1_HUMAN | reviewed | Collagen alpha-1(V) chain | COL5A1 | Homo sapiens | 1838 |
| NM_000393 | | P05997 | CO5A2_HUMAN | reviewed | Collagen alpha-2(V) chain | COL5A2 | Homo sapiens | 1499 |
| NM_004369 | NM_004369 -> P12111-1 | P12111 | CO6A3_HUMAN | reviewed | Collagen alpha-3(VI) chain | COL6A3 | Homo sapiens | 3177 |
| NM_000094 | NM_000094 -> Q02388-1 | Q02388 | CO7A1_HUMAN | reviewed | Collagen alpha-1(VII) chain (Long-chain collagen) (LC collagen) | COL7A1 | Homo sapiens | 2944 |
| NM_001850 | | P27658 | CO8A1_HUMAN | reviewed | Collagen alpha-1(VIII) chain (Endothelial collagen) [Cleaved into: Vastatin] | COL8A1 C3orf7 | Homo sapiens | 744 |
| NM_000493 | | Q03692 | COAA1_HUMAN | reviewed | Collagen alpha-1(X) chain | COL10A1 | Homo sapiens | 680 |
| NM_001854 | NM_001854 -> P12107-1 | P12107 | COBA1_HUMAN | reviewed | Collagen alpha-1(XI) chain | COL11A1 COLL6 | Homo sapiens | 1806 |
| NM_001901 | | P29279 | CTGF_HUMAN | reviewed | Connective tissue growth factor (CCN family member 2) (Hypertrophic chondrocyte-spe | CTGF CCN2 HCS24 IGFBP8 | Homo sapiens | 349 |
| NM_001937 | | Q07507 | DERM_HUMAN | reviewed | Dermatopontin (Tyrosine-rich acidic matrix protein) (TRAMP) | DPT | Homo sapiens | 201 |
| NM_000501 | NM_000501 -> P15502-2 | P15502 | ELN_HUMAN | reviewed | Elastin (Tropoelastin) | ELN | Homo sapiens | 786 |
| NM_212482 | | P02751 | FINC_HUMAN | reviewed | Fibronectin (FN) (Cold-insoluble globulin) (CIG) [Cleaved into: Anastellin; Ugl-Y1; Ugl-Y2; | FN1 FN | Homo sapiens | 2386 |
| NM_013231 | | O43155 | FLRT2_HUMAN | reviewed | Leucine-rich repeat transmembrane protein FLRT2 (Fibronectin-like domain-containing l | FLRT2 KIAA0405 UNQ232/PRO2 | Homo sapiens | 660 |
| NM_031935 | NM_031935 -> Q96RW7-1 | Q96RW7 | HMCN1_HUMAN | reviewed | Hemicentin-1 (Fibulin-6) (FIBL-6) | HMCN1 FIBL6 | Homo sapiens | 5635 |
| NM_001884 | | P10915 | HPLN1_HUMAN | reviewed | Hyaluronan and proteoglycan link protein 1 (Cartilage-linking protein 1) (Cartilage-link pr | HAPLN1 CRTL1 | Homo sapiens | 354 |
| NM_000428 | | Q14767 | LTBP2_HUMAN | reviewed | Latent-transforming growth factor beta-binding protein 2 (LTBP-2) | LTBP2 C14orf141 LTBP3 | Homo sapiens | 1821 |
| NM_002425 | | P09238 | MMP10_HUMAN | reviewed | Stromelysin-2 (SL-2) (EC 3.4.24.22) (Matrix metalloproteinase-10) (MMP-10) (Transin-2) | MMP10 STMY2 | Homo sapiens | 476 |
| NM_002403 | NM_002403 -> P55001-1 | P55001 | MFAP2_HUMAN | reviewed | Microfibrillar-associated protein 2 (MFAP-2) (Microfibril-associated glycoprotein 1) (MA | MFAP2 MAGP1 | Homo sapiens | 183 |
| NM_001198695 | NM_001198695 -> P55083-2 | P55083 | MFAP4_HUMAN | reviewed | Microfibril-associated glycoprotein 4 | MFAP4 | Homo sapiens | 255 |
| NM_006475 | NM_006475 -> Q15063-1 | Q15063 | POSTN_HUMAN | reviewed | Periostin (PN) (Osteoblast-specific factor 2) (OSF-2) | POSTN OSF2 | Homo sapiens | 836 |
| NM_002160 | NM_002160 -> P24821-1 | P24821 | TENA_HUMAN | reviewed | Tenascin (TN) (Cytotactin) (GMEM) (GP 150-225) (Glioma-associated-extracellular matri | TNC HXB | Homo sapiens | 2201 |
| NM_000660 | | P01137 | TGFB1_HUMAN | reviewed | Transforming growth factor beta-1 (TGF-beta-1) [Cleaved into: Latency-associated pepti | TGFB1 TGFB | Homo sapiens | 390 |
| NM_000358 | | Q15582 | BGH3_HUMAN | reviewed | Transforming growth factor-beta-induced protein ig-h3 (Beta ig-h3) (Kerato-epithelin) (R | TGFBI BIGH3 | Homo sapiens | 683 |
| NM_004385 | NM_004385 -> P13611-1 | P13611 | CSPG2_HUMAN | reviewed | Versican core protein (Chondroitin sulfate proteoglycan core protein 2) (Chondroitin sulf | VCAN CSPG2 | Homo sapiens | 3396 |

| growth factor | enrichment 3.73 | | | | | | | |
|---|---|---|---|---|---|---|---|---|
| **yourlist:M20180608A7434721E10EE6586998A056CC** | **isomap:M20180608A7434721E10EE6586998A056CCD** | **Entry** | **Entry name** | **Status** | **Protein names** | **Gene names** | **Organism** | **Length** |
| NM_000800 | NM_000800 -> P05230-1 | P05230 | FGF1_HUMAN | reviewed | Fibroblast growth factor 1 (FGF-1) (Acidic fibroblast growth factor) (aFGF) (Endothelial ce | FGF1 FGFA | Homo sapiens | 155 |
| NM_001001557 | | Q6KF10 | GDF6_HUMAN | reviewed | Growth/differentiation factor 6 (GDF-6) (Bone morphogenetic protein 13) (BMP-13) (Gr | GDF6 BMP13 GDF16 | Homo sapiens | 455 |
| NM_001945 | | Q99075 | HBEGF_HUMAN | reviewed | Proheparin-binding EGF-like growth factor [Cleaved into: Heparin-binding EGF-like growt | HBEGF DTR DTS HEGFL | Homo sapiens | 208 |
| NM_002192 | | P08476 | INHBA_HUMAN | reviewed | Inhibin beta A chain (Activin beta-A chain) (Erythroid differentiation protein) (EDF) | INHBA | Homo sapiens | 426 |
| NM_031479 | | P58166 | INHBE_HUMAN | reviewed | Inhibin beta E chain (Activin beta-E chain) | INHBE | Homo sapiens | 350 |
| NM_001111283 | | Q13429 | Q13429_HUMAN | unreviewed | Insulin-like growth factor I (Fragment) | IGF1 IGF-I | Homo sapiens | 139 |
| NM_000641 | NM_000641 -> P20809-1 | P20809 | IL11_HUMAN | reviewed | Interleukin-11 (IL-11) (Adipogenesis inhibitory factor) (AGIF) (Oprelvekin) | IL11 | Homo sapiens | 199 |
| NM_000600 | | P05231 | IL6_HUMAN | reviewed | Interleukin-6 (IL-6) (B-cell stimulatory factor 2) (BSF-2) (CTL differentiation factor) (CDF) ( | IL6 IFNB2 | Homo sapiens | 212 |
| NM_002309 | NM_002309 -> P15018-1 | P15018 | LIF_HUMAN | reviewed | Leukemia inhibitory factor (LIF) (Differentiation-stimulating factor) (D factor) (Melanoma | LIF HILDA | Homo sapiens | 202 |
| NM_016205 | NM_016205 -> Q9NRA1-1 | Q9NRA1 | PDGFC_HUMAN | reviewed | Platelet-derived growth factor C (PDGF-C) (Fallotein) (Spinal cord-derived growth factor) | PDGFC SCDGF UNQ174/PRO200 | Homo sapiens | 345 |
| NM_000660 | | P01137 | TGFB1_HUMAN | reviewed | Transforming growth factor beta-1 (TGF-beta-1) [Cleaved into: Latency-associated pepti | TGFB1 TGFB | Homo sapiens | 390 |
| NM_001135599 | NM_001135599 -> P61812-2 | P61812 | TGFB2_HUMAN | reviewed | Transforming growth factor beta-2 (TGF-beta-2) (BSC-1 cell growth inhibitor) (Cetermin) | TGFB2 | Homo sapiens | 414 |
| NM_001025366 | NM_001025366 -> P15692-14 | P15692 | VEGFA_HUMAN | reviewed | Vascular endothelial growth factor A (VEGF-A) (Vascular permeability factor) (VPF) | VEGFA VEGF | Homo sapiens | 232 |

| cell motion | enrichment 5.03 | | | | | | | |
|---|---|---|---|---|---|---|---|---|
| **yourlist:M2018060883C3DD8CE55183C76102DC5D3A** | **isomap:M2018060883C3DD8CE55183C76102DC5D3A** | **Entry** | **Entry name** | **Status** | **Protein names** | **Gene names** | **Organism** | **Length** |
| NM_017449 | NM_017449 -> P29323-2 | P29323 | EPHB2_HUMAN | reviewed | Ephrin type-B receptor 2 (EC 2.7.10.1) (Developmentally-regulated Eph-related tyrosine | EPHB2 DRT EPHT3 EPTH3 ERK H | Homo sapiens | 1055 |
| NM_153487 | NM_153487 -> Q8NFP4-1 | Q8NFP4 | MDGA1_HUMAN | reviewed | MAM domain-containing glycosylphosphatidylinositol anchor protein 1 (GPI and MAM p | MDGA1 MAMDC3 | Homo sapiens | 955 |
| NM_001497 | NM_001497 -> P15291-1 | P15291 | B4GT1_HUMAN | reviewed | Beta-1,4-galactosyltransferase 1 (Beta-1,4-GalTase 1) (Beta4Gal-T1) (b4Gal-T1) (EC 2.4.1 | B4GALT1 GGTB2 | Homo sapiens | 398 |
| NM_004307 | | B4DJ88 | B4DJ88_HUMAN | unreviewed | cDNA FLJ52628, highly similar to Amyloid beta A4 protein-binding family B member 2 (Fe65-like protein) | | Homo sapiens | 471 |
| NM_001203 | NM_001203 -> O00238-1 | O00238 | BMR1B_HUMAN | reviewed | Bone morphogenetic protein receptor type-1B (BMP type-1B receptor) (BMPR-1B) (EC 2 | BMPR1B | Homo sapiens | 502 |
| NM_001792 | NM_001792 -> P19022-1 | P19022 | CADH2_HUMAN | reviewed | Cadherin-2 (CDw325) (Neural cadherin) (N-cadherin) (CD antigen CD325) | CDH2 CDHN NCAD | Homo sapiens | 906 |
| NM_138455 | NM_138455 -> Q96CG8-1 | Q96CG8 | CTHR1_HUMAN | reviewed | Collagen triple helix repeat-containing protein 1 (Protein NMTC1) | CTHRC1 UNQ762/PRO1550 | Homo sapiens | 243 |
| NM_000093 | | P20908 | CO5A1_HUMAN | reviewed | Collagen alpha-1(V) chain | COL5A1 | Homo sapiens | 1838 |
| NM_001901 | | P29279 | CTGF_HUMAN | reviewed | Connective tissue growth factor (CCN family member 2) (Hypertrophic chondrocyte-spe | CTGF CCN2 HCS24 IGFBP8 | Homo sapiens | 349 |
| NM_000399 | NM_000399 -> P11161-1 | P11161 | EGR2_HUMAN | reviewed | E3 SUMO-protein ligase EGR2 (EC 2.3.2.-) (AT591) (E3 SUMO-protein transferase ERG2) ( | EGR2 KROX20 | Homo sapiens | 476 |
| NM_001079675 | NM_001079675 -> P43268-1 | P43268 | ETV4_HUMAN | reviewed | ETS translocation variant 4 (Adenovirus E1A enhancer-binding protein) (E1A-F) (Polyoma | ETV4 E1AF PEA3 | Homo sapiens | 484 |
| NM_212482 | | P02751 | FINC_HUMAN | reviewed | Fibronectin (FN) (Cold-insoluble globulin) (CIG) [Cleaved into: Anastellin; Ugl-Y1; Ugl-Y2; | FN1 FN | Homo sapiens | 2386 |
| NM_001945 | | Q99075 | HBEGF_HUMAN | reviewed | Proheparin-binding EGF-like growth factor [Cleaved into: Heparin-binding EGF-like growt | HBEGF DTR DTS HEGFL | Homo sapiens | 208 |
| NM_181353 | NM_181353 -> P41134-2 | P41134 | ID1_HUMAN | reviewed | DNA-binding protein inhibitor ID-1 (Class B basic helix-loop-helix protein 24) (bHLHb24) ( | ID1 BHLHB24 ID | Homo sapiens | 155 |
| NM_001111283 | | Q13429 | Q13429_HUMAN | unreviewed | Insulin-like growth factor I (Fragment) | IGF1 IGF-I | Homo sapiens | 139 |
| NM_001004439 | NM_001004439 -> Q9UKX5-1 | Q9UKX5 | ITA11_HUMAN | reviewed | Integrin alpha-11 | ITGA11 MSTP018 | Homo sapiens | 1188 |
| NM_000600 | | P05231 | IL6_HUMAN | reviewed | Interleukin-6 (IL-6) (B-cell stimulatory factor 2) (BSF-2) (CTL differentiation factor) (CDF) ( | IL6 IFNB2 | Homo sapiens | 212 |
| NM_201266 | NM_201266 -> O60462-1 | O60462 | NRP2_HUMAN | reviewed | Neuropilin-2 (Vascular endothelial cell growth factor 165 receptor 2) | NRP2 VEGF165R2 | Homo sapiens | 931 |
| NM_003561 | | O15496 | PA2GX_HUMAN | reviewed | Group 10 secretory phospholipase A2 (EC 3.1.1.4) (Group X secretory phospholipase A2) | PLA2G10 | Homo sapiens | 165 |
| NM_002659 | NM_002659 -> Q03405-1 | Q03405 | UPAR_HUMAN | reviewed | Urokinase plasminogen activator surface receptor (U-PAR) (uPAR) (Monocyte activation | PLAUR MO3 UPAR | Homo sapiens | 335 |
| NM_006505 | | P15151 | PVR_HUMAN | reviewed | Poliovirus receptor (Nectin-like protein 5) (NECL-5) (CD antigen CD155) | PVR PVS | Homo sapiens | 417 |
| NM_004598 | | Q08629 | TICN1_HUMAN | reviewed | Testican-1 (Protein SPOCK) | SPOCK1 SPOCK TIC1 TICN1 | Homo sapiens | 439 |
| NM_003901 | | O95470 | SGPL1_HUMAN | reviewed | Sphingosine-1-phosphate lyase 1 (S1PL) (SP-lyase 1) (SPL 1) (hSPL) (EC 4.1.2.27) (Sphingo | SGPL1 KIAA1252 | Homo sapiens | 568 |
| NM_022648 | NM_022648 -> Q9HBL0-1 | Q9HBL0 | TENS1_HUMAN | reviewed | Tensin-1 | TNS1 TNS | Homo sapiens | 1735 |
| NM_000660 | | P01137 | TGFB1_HUMAN | reviewed | Transforming growth factor beta-1 (TGF-beta-1) [Cleaved into: Latency-associated pepti | TGFB1 TGFB | Homo sapiens | 390 |
| NM_001135599 | NM_001135599 -> P61812-2 | P61812 | TGFB2_HUMAN | reviewed | Transforming growth factor beta-2 (TGF-beta-2) (BSC-1 cell growth inhibitor) (Cetermin) | TGFB2 | Homo sapiens | 414 |
| NM_000366 | NM_000366 -> P09493-10 | P09493 | TPM1_HUMAN | reviewed | Tropomyosin alpha-1 chain (Alpha-tropomyosin) (Tropomyosin-1) | TPM1 C15orf13 TMSA | Homo sapiens | 284 |
| NM_016639 | NM_016639 -> Q9NP84-1 | Q9NP84 | TNR12_HUMAN | reviewed | Tumor necrosis factor receptor superfamily member 12A (Fibroblast growth factor-indu | TNFRSF12A FN14 | Homo sapiens | 129 |
| NM_170744 | NM_170744 -> Q8IZJ1-1 | Q8IZJ1 | UNC5B_HUMAN | reviewed | Netrin receptor UNC5B (Protein unc-5 homolog 2) (Protein unc-5 homolog B) (p53-regul | UNC5B P53RDL1 UNC5H2 UNQ1 | Homo sapiens | 945 |
| NM_004385 | NM_004385 -> P13611-1 | P13611 | CSPG2_HUMAN | reviewed | Versican core protein (Chondroitin sulfate proteoglycan core protein 2) (Chondroitin sulf | VCAN CSPG2 | Homo sapiens | 3396 |

| | | | | | | | | |
|---|---|---|---|---|---|---|---|---|
| NM_003391 | | P09544 | WNT2_HUMAN | reviewed | Protein Wnt-2 (Int-1-like protein 1) (Int-1-related protein) (IRP) | WNT2 INT1L1 IRP | Homo sapiens | 360 |

**response to wounding** **enrichment 4.05**

| yourlist:M201806089011D4373CD8E192AC56F03911 | isomap:M201806089011D4373CD8E192AC56F039114 | Entry | Entry name | Status | Protein names | Gene names | Organism | Length |
|---|---|---|---|---|---|---|---|---|
| NM_003485 | | Q15743 | OGR1_HUMAN | reviewed | Ovarian cancer G-protein coupled receptor 1 (OGR-1) (G-protein coupled receptor 68) (C | GPR68 OGR1 | Homo sapiens | 365 |
| NM_016931 | | Q9NPH5 | NOX4_HUMAN | reviewed | NADPH oxidase 4 (EC 1.6.3.-) (Kidney oxidase-1) (KOX-1) (Kidney superoxide-producing N | NOX4 RENOX | Homo sapiens | 578 |
| NM_000362 | | P35625 | TIMP3_HUMAN | reviewed | Metalloproteinase inhibitor 3 (Protein MIG-5) (Tissue inhibitor of metalloproteinases 3) ( | TIMP3 | Homo sapiens | 211 |
| NM_001497 | NM_001497 -> P15291-1 | P15291 | B4GT1_HUMAN | reviewed | Beta-1,4-galactosyltransferase 1 (Beta-1,4-GalTase 1) (Beta4Gal-T1) (b4Gal-T1) (EC 2.4.1 | B4GALT1 GGTB2 | Homo sapiens | 398 |
| NM_001203 | NM_001203 -> O00238-1 | O00238 | BMR1B_HUMAN | reviewed | Bone morphogenetic protein receptor type-1B (BMP type-1B receptor) (BMPR-1B) (EC 2 | BMPR1B | Homo sapiens | 502 |
| NM_000710 | | P46663 | BKRB1_HUMAN | reviewed | B1 bradykinin receptor (B1R) (BK-1 receptor) | BDKRB1 BRADYB1 | Homo sapiens | 353 |
| NM_001992 | | P25116 | PAR1_HUMAN | reviewed | Proteinase-activated receptor 1 (PAR-1) (Coagulation factor II receptor) (Thrombin recep | F2R CF2R PAR1 TR | Homo sapiens | 425 |
| NM_005242 | | P55085 | PAR2_HUMAN | reviewed | Proteinase-activated receptor 2 (PAR-2) (Coagulation factor II receptor-like 1) (G-protein | F2RL1 GPR11 PAR2 | Homo sapiens | 397 |
| NM_004101 | NM_004101 -> O00254-1 | O00254 | PAR3_HUMAN | reviewed | Proteinase-activated receptor 3 (PAR-3) (Coagulation factor II receptor-like 2) (Thrombir | F2RL2 PAR3 | Homo sapiens | 374 |
| NM_000093 | | P20908 | CO5A1_HUMAN | reviewed | Collagen alpha-1(V) chain | COL5A1 | Homo sapiens | 1838 |
| NM_001901 | | P29279 | CTGF_HUMAN | reviewed | Connective tissue growth factor (CCN family member 2) (Hypertrophic chondrocyte-spec | CTGF CCN2 HCS24 IGFBP8 | Homo sapiens | 349 |
| NM_004415 | NM_004415 -> P15924-1 | P15924 | DESP_HUMAN | reviewed | Desmoplakin (DP) (250/210 kDa paraneoplastic pemphigus antigen) | DSP | Homo sapiens | 2871 |
| NM_212482 | | P02751 | FINC_HUMAN | reviewed | Fibronectin (FN) (Cold-insoluble globulin) (CIG) [Cleaved into: Anastellin; Ugl-Y1; Ugl-Y2; I | FN1 FN | Homo sapiens | 2386 |
| NM_015973 | | P22466 | GALA_HUMAN | reviewed | Galanin peptides [Cleaved into: Galanin; Galanin message-associated peptide (GMAP)] | GAL GAL1 GALN GLNN | Homo sapiens | 123 |
| NM_002133 | | P09601 | HMOX1_HUMAN | reviewed | Heme oxygenase 1 (HO-1) (EC 1.14.14.18) | HMOX1 HO HO1 | Homo sapiens | 288 |
| NM_031935 | NM_031935 -> Q96RW7-1 | Q96RW7 | HMCN1_HUMAN | reviewed | Hemicentin-1 (Fibulin-6) (FIBL-6) | HMCN1 FIBL6 | Homo sapiens | 5635 |
| NM_001945 | | Q99075 | HBEGF_HUMAN | reviewed | Proheparin-binding EGF-like growth factor [Cleaved into: Heparin-binding EGF-like growt | HBEGF DTR DTS HEGFL | Homo sapiens | 208 |
| NM_002167 | | Q02535 | ID3_HUMAN | reviewed | DNA-binding protein inhibitor ID-3 (Class B basic helix-loop-helix protein 25) (bHLHb25) ( | ID3 1R21 BHLHB25 HEIR1 | Homo sapiens | 119 |
| NM_001111283 | | Q13429 | Q13429_HUMAN | unreviewed | Insulin-like growth factor I (Fragment) | IGF1 IGF-I | Homo sapiens | 139 |
| NM_000888 | NM_000888 -> P18564-1 | P18564 | ITB6_HUMAN | reviewed | Integrin beta-6 | ITGB6 | Homo sapiens | 788 |
| NM_173842 | NM_173842 -> P18510-1 | P18510 | IL1RA_HUMAN | reviewed | Interleukin-1 receptor antagonist protein (IL-1RN) (IL-1ra) (IRAP) (ICIL-1RA) (IL1 inhibitor) | IL1RN IL1F3 IL1RA | Homo sapiens | 177 |
| NM_000641 | NM_000641 -> P20809-1 | P20809 | IL11_HUMAN | reviewed | Interleukin-11 (IL-11) (Adipogenesis inhibitory factor) (AGIF) (Oprelvekin) | IL11 | Homo sapiens | 199 |
| NM_000600 | | P05231 | IL6_HUMAN | reviewed | Interleukin-6 (IL-6) (B-cell stimulatory factor 2) (BSF-2) (CTL differentiation factor) (CDF) ( | IL6 IFNB2 | Homo sapiens | 212 |
| NM_031419 | NM_031419 -> Q9BYH8-1 | Q9BYH8 | IKBZ_HUMAN | reviewed | NF-kappa-B inhibitor zeta (I-kappa-B-zeta) (IkB-zeta) (IkappaBzeta) (IL-1 inducible nuclea | NFKBIZ IKBZ INAP MAIL | Homo sapiens | 718 |
| NM_002659 | NM_002659 -> Q03405-1 | Q03405 | UPAR_HUMAN | reviewed | Urokinase plasminogen activator surface receptor (U-PAR) (uPAR) (Monocyte activation | PLAUR MO3 UPAR | Homo sapiens | 335 |
| NM_001040058 | NM_001040058 -> P10451-1 | P10451 | OSTP_HUMAN | reviewed | Osteopontin (Bone sialoprotein 1) (Nephropontin) (Secreted phosphoprotein 1) (SPP-1) ( | SPP1 BNSP OPN PSEC0156 | Homo sapiens | 314 |
| NM_000602 | NM_000602 -> P05121-1 | P05121 | PAI1_HUMAN | reviewed | Plasminogen activator inhibitor 1 (PAI) (PAI-1) (Endothelial plasminogen activator inhibit | SERPINE1 PAI1 PLANH1 | Homo sapiens | 402 |
| NM_002977 | | Q15858 | SCN9A_HUMAN | reviewed | Sodium channel protein type 9 subunit alpha (Neuroendocrine sodium channel) (hNE-Na | SCN9A NENA | Homo sapiens | 1988 |
| NM_001006946 | | P18827 | SDC1_HUMAN | reviewed | Syndecan-1 (SYND1) (CD antigen CD138) | SDC1 SDC | Homo sapiens | 310 |
| NM_002160 | NM_002160 -> P24821-1 | P24821 | TENA_HUMAN | reviewed | Tenascin (TN) (Cytotactin) (GMEM) (GP 150-225) (Glioma-associated-extracellular matri | TNC HXB | Homo sapiens | 2201 |
| NM_006528 | NM_006528 -> P48307-1 | P48307 | TFPI2_HUMAN | reviewed | Tissue factor pathway inhibitor 2 (TFPI-2) (Placental protein 5) (PP5) | TFPI2 | Homo sapiens | 235 |
| NM_000660 | | P01137 | TGFB1_HUMAN | reviewed | Transforming growth factor beta-1 (TGF-beta-1) [Cleaved into: Latency-associated peptic | TGFB1 TGFB | Homo sapiens | 390 |
| NM_001135599 | NM_001135599 -> P61812-2 | P61812 | TGFB2_HUMAN | reviewed | Transforming growth factor beta-2 (TGF-beta-2) (BSC-1 cell growth inhibitor) (Cetermin) | TGFB2 | Homo sapiens | 414 |
| NM_000366 | NM_000366 -> P09493-10 | P09493 | TPM1_HUMAN | reviewed | Tropomyosin alpha-1 chain (Alpha-tropomyosin) (Tropomyosin-1) | TPM1 C15orf13 TMSA | Homo sapiens | 284 |
| NM_003326 | NM_003326 -> P23510-1 | P23510 | TNFL4_HUMAN | reviewed | Tumor necrosis factor ligand superfamily member 4 (Glycoprotein Gp34) (OX40 ligand) ( | TNFSF4 TXGP1 | Homo sapiens | 183 |
| NM_007115 | | P98066 | TSG6_HUMAN | reviewed | Tumor necrosis factor-inducible gene 6 protein (Hyaluronate-binding protein) (TNF-stimu | TNFAIP6 TSG6 | Homo sapiens | 277 |
| NM_004385 | NM_004385 -> P13611-1 | P13611 | CSPG2_HUMAN | reviewed | Versican core protein (Chondroitin sulfate proteoglycan core protein 2) (Chondroitin sulf | VCAN CSPG2 | Homo sapiens | 3396 |

**cell adhesion** **enrichment 6.43**

| yourlist:M20180608A7434721E10EE6586998A056CCI | isomap:M20180608A7434721E10EE6586998A056CCD | Entry | Entry name | Status | Protein names | Gene names | Organism | Length |
|---|---|---|---|---|---|---|---|---|
| NM_003474 | NM_003474 -> O43184-1 | O43184 | ADA12_HUMAN | reviewed | Disintegrin and metalloproteinase domain-containing protein 12 (ADAM 12) (EC 3.4.24.- | ADAM12 MLTN UNQ346/PRO54 | Homo sapiens | 909 |
| NM_001129 | NM_001129 -> Q8IUX7-1 | Q8IUX7 | AEBP1_HUMAN | reviewed | Adipocyte enhancer-binding protein 1 (AE-binding protein 1) (Aortic carboxypeptidase-lil | AEBP1 ACLP | Homo sapiens | 1158 |
| NM_021155 | NM_021155 -> Q9NNX6-1 | Q9NNX6 | CD209_HUMAN | reviewed | CD209 antigen (C-type lectin domain family 4 member L) (Dendritic cell-specific ICAM-3- | CD209 CLEC4L | Homo sapiens | 404 |
| NM_002821 | NM_002821 -> Q13308-1 | Q13308 | PTK7_HUMAN | reviewed | Inactive tyrosine-protein kinase 7 (Colon carcinoma kinase 4) (CCK-4) (Protein-tyrosine k | PTK7 CCK4 | Homo sapiens | 1070 |
| NM_000459 | | Q02763 | TIE2_HUMAN | reviewed | Angiopoietin-1 receptor (EC 2.7.10.1) (Endothelial tyrosine kinase) (Tunica interna endot | TEK TIE2 VMCM VMCM1 | Homo sapiens | 1124 |
| NM_001497 | NM_001497 -> P15291-1 | P15291 | B4GT1_HUMAN | reviewed | Beta-1,4-galactosyltransferase 1 (Beta-1,4-GalTase 1) (Beta4Gal-T1) (b4Gal-T1) (EC 2.4.1 | B4GALT1 GGTB2 | Homo sapiens | 398 |
| NM_001143668 | | Q86SJ2 | AMGO2_HUMAN | reviewed | Amphoterin-induced protein 2 (AMIGO-2) (Alivin-1) (Differentially expressed in gastric ac | AMIGO2 ALI1 | Homo sapiens | 522 |
| NM_001203 | NM_001203 -> O00238-1 | O00238 | BMR1B_HUMAN | reviewed | Bone morphogenetic protein receptor type-1B (BMP type-1B receptor) (BMPR-1B) (EC 2 | BMPR1B | Homo sapiens | 502 |
| NM_001792 | NM_001792 -> P19022-1 | P19022 | CADH2_HUMAN | reviewed | Cadherin-2 (CDw325) (Neural cadherin) (N-cadherin) (CD antigen CD325) | CDH2 CDHN NCAD | Homo sapiens | 906 |
| NM_001795 | NM_001795 -> P33151-1 | P33151 | CADH5_HUMAN | reviewed | Cadherin-5 (7B4 antigen) (Vascular endothelial cadherin) (VE-cadherin) (CD antigen CD14 | CDH5 | Homo sapiens | 784 |
| NM_000095 | NM_000095 -> P49747-1 | P49747 | COMP_HUMAN | reviewed | Cartilage oligomeric matrix protein (COMP) (Thrombospondin-5) (TSP5) | COMP | Homo sapiens | 757 |
| NM_000093 | | P20908 | CO5A1_HUMAN | reviewed | Collagen alpha-1(V) chain | COL5A1 | Homo sapiens | 1838 |
| NM_004369 | NM_004369 -> P12111-1 | P12111 | CO6A3_HUMAN | reviewed | Collagen alpha-3(VI) chain | COL6A3 | Homo sapiens | 3177 |
| NM_000094 | NM_000094 -> Q02388-1 | Q02388 | CO7A1_HUMAN | reviewed | Collagen alpha-1(VII) chain (Long-chain collagen) (LC collagen) | COL7A1 | Homo sapiens | 2944 |
| NM_001850 | | P27658 | CO8A1_HUMAN | reviewed | Collagen alpha-1(VIII) chain (Endothelial collagen) [Cleaved into: Vastatin] | COL8A1 C3orf7 | Homo sapiens | 744 |
| NM_001854 | NM_001854 -> P12107-1 | P12107 | COBA1_HUMAN | reviewed | Collagen alpha-1(XI) chain | COL11A1 COLL6 | Homo sapiens | 1806 |
| NM_001901 | | P29279 | CTGF_HUMAN | reviewed | Connective tissue growth factor (CCN family member 2) (Hypertrophic chondrocyte-spec | CTGF CCN2 HCS24 IGFBP8 | Homo sapiens | 349 |
| NM_001937 | | Q07507 | DERM_HUMAN | reviewed | Dermatopontin (Tyrosine-rich acidic matrix protein) (TRAMP) | DPT | Homo sapiens | 201 |
| NM_006832 | NM_006832 -> Q96AC1-1 | Q96AC1 | FERM2_HUMAN | reviewed | Fermitin family homolog 2 (Kindlin-2) (Mitogen-inducible gene 2 protein) (MIG-2) (Plecks | FERMT2 KIND2 MIG2 PLEKHC1 | Homo sapiens | 680 |
| NM_212482 | | P02751 | FINC_HUMAN | reviewed | Fibronectin (FN) (Cold-insoluble globulin) (CIG) [Cleaved into: Anastellin; Ugl-Y1; Ugl-Y2; I | FN1 FN | Homo sapiens | 2386 |
| NM_013231 | | O43155 | FLRT2_HUMAN | reviewed | Leucine-rich repeat transmembrane protein FLRT2 (Fibronectin-like domain-containing l | FLRT2 KIAA0405 UNQ232/PRO2 | Homo sapiens | 660 |
| NM_005524 | | Q14469 | HES1_HUMAN | reviewed | Transcription factor HES-1 (Class B basic helix-loop-helix protein 39) (bHLHb39) (Hairy an | HES1 BHLHB39 HL HRY | Homo sapiens | 280 |
| NM_001884 | | P10915 | HPLN1_HUMAN | reviewed | Hyaluronan and proteoglycan link protein 1 (Cartilage-linking protein 1) (Cartilage-link pr | HAPLN1 CRTL1 | Homo sapiens | 354 |
| NM_001004439 | NM_001004439 -> Q9UKX5-1 | Q9UKX5 | ITA11_HUMAN | reviewed | Integrin alpha-11 | ITGA11 MSTP018 | Homo sapiens | 1188 |
| NM_000888 | NM_000888 -> P18564-1 | P18564 | ITB6_HUMAN | reviewed | Integrin beta-6 | ITGB6 | Homo sapiens | 788 |
| NM_016269 | NM_016269 -> Q9UJU2-1 | Q9UJU2 | LEF1_HUMAN | reviewed | Lymphoid enhancer-binding factor 1 (LEF-1) (T cell-specific transcription factor 1-alpha) ( | LEF1 | Homo sapiens | 399 |
| NM_001142393 | NM_001142393 -> Q14511-3 | Q14511 | CASL_HUMAN | reviewed | Enhancer of filamentation 1 (hEF1) (CRK-associated substrate-related protein) (CAS-L) (C | NEDD9 CASL CASS2 | Homo sapiens | 834 |
| NM_201266 | NM_201266 -> O60462-1 | O60462 | NRP2_HUMAN | reviewed | Neuropilin-2 (Vascular endothelial cell growth factor 165 receptor 2) | NRP2 VEGF165R2 | Homo sapiens | 931 |
| NM_016522 | NM_016522 -> Q9P121-1 | Q9P121 | NTRI_HUMAN | reviewed | Neurotrimin (hNT) (IgLON family member 2) | NTM IGLON2 NT UNQ297/PRO3 | Homo sapiens | 344 |
| NM_006475 | NM_006475 -> Q15063-1 | Q15063 | POSTN_HUMAN | reviewed | Periostin (PN) (Osteoblast-specific factor 2) (OSF-2) | POSTN OSF2 | Homo sapiens | 836 |
| NM_006505 | | P15151 | PVR_HUMAN | reviewed | Poliovirus receptor (Nectin-like protein 5) (NECL-5) (CD antigen CD155) | PVR PVS | Homo sapiens | 417 |
| NM_001105243 | NM_001105243 -> Q8TAB3-2 | Q8TAB3 | PCD19_HUMAN | reviewed | Protocadherin-19 | PCDH19 KIAA1313 | Homo sapiens | 1148 |
| NM_203487 | NM_203487 -> Q9HC56-1 | Q9HC56 | PCDH9_HUMAN | reviewed | Protocadherin-9 | PCDH9 | Homo sapiens | 1237 |
| NM_001040058 | NM_001040058 -> P10451-1 | P10451 | OSTP_HUMAN | reviewed | Osteopontin (Bone sialoprotein 1) (Nephropontin) (Secreted phosphoprotein 1) (SPP-1) ( | SPP1 BNSP OPN PSEC0156 | Homo sapiens | 314 |
| NM_004598 | | Q08629 | TICN1_HUMAN | reviewed | Testican-1 (Protein SPOCK) | SPOCK1 SPOCK TIC1 TICN1 | Homo sapiens | 439 |
| NM_002160 | NM_002160 -> P24821-1 | P24821 | TENA_HUMAN | reviewed | Tenascin (TN) (Cytotactin) (GMEM) (GP 150-225) (Glioma-associated-extracellular matri | TNC HXB | Homo sapiens | 2201 |
| NM_000358 | | Q15582 | BGH3_HUMAN | reviewed | Transforming growth factor-beta-induced protein ig-h3 (Beta ig-h3) (Kerato-epithelin) (R | TGFBI BIGH3 | Homo sapiens | 683 |
| NM_016639 | NM_016639 -> Q9NP84-1 | Q9NP84 | TNR12_HUMAN | reviewed | Tumor necrosis factor receptor superfamily member 12A (Fibroblast growth factor-induc | TNFRSF12A FN14 | Homo sapiens | 129 |
| NM_007115 | | P98066 | TSG6_HUMAN | reviewed | Tumor necrosis factor-inducible gene 6 protein (Hyaluronate-binding protein) (TNF-stimu | TNFAIP6 TSG6 | Homo sapiens | 277 |
| NM_004385 | NM_004385 -> P13611-1 | P13611 | CSPG2_HUMAN | reviewed | Versican core protein (Chondroitin sulfate proteoglycan core protein 2) (Chondroitin sulf | VCAN CSPG2 | Homo sapiens | 3396 |

**Sup. Table 2: Proteins differentially expressed between CM from shControl and shNODAL expressing MDA-MB-231 cells**

| Protein names | Gene names | T-test Significant | -Log T-test p-value | Log2 Heavy/Light ratio (shNodal/shControl) | Protein IDs | Majority protein IDs |
|---|---|---|---|---|---|---|
| 1,4-alpha-glucan-branching enzyme | GBE1 | | 0.364801 | 0.304593 | Q04446;E9PGM4 | Q04446;E9PGM4 |
| 10 kDa heat shock protein, mitochondrial | HSPE1 | | 0.956191 | 0.749295 | P61604;B8ZZL8;B8ZZ54 | P61604;B8ZZL8 |
| 14 kDa phosphohistidine phosphatase | PHPT1 | | 0.606235 | 0.268675 | Q9NRX4;Q9NRX4-2 | Q9NRX4;Q9NRX4-2 |
| 14-3-3 protein beta/alpha;14-3-3 protein beta/alpha, N-terminally processed | YWHAB | + | 1.94777 | 1.8601 | P31946;Q4VY19;Q4VY20 | P31946 |
| 14-3-3 protein beta/alpha;14-3-3 protein beta/alpha, N-terminally processed | YWHAB | + | 3.66263 | 1.89822 | P31946-2 | P31946-2 |
| 14-3-3 protein epsilon | YWHAE | | 0.608943 | 0.663179 | P62258;P62258-2;B4DJF2;I3L3T1;K7EIT4;K7EM20;I3L0W5 | P62258;P62258-2 |
| 14-3-3 protein eta | YWHAH | + | 1.42652 | 1.71052 | Q04917;A2IDB2;F8WEB6;A2IDB1 | Q04917;A2IDB2 |
| 14-3-3 protein gamma;14-3-3 protein gamma, N-terminally processed | YWHAG | | 0.261303 | 0.257777 | P61981 | P61981 |
| 14-3-3 protein sigma | SFN | + | 1.32225 | 1.68875 | P31947;P31947-2 | P31947;P31947-2 |
| 14-3-3 protein theta | YWHAQ | + | 1.46185 | 1.5442 | P27348;E9PG15 | P27348;E9PG15 |
| 14-3-3 protein zeta/delta | YWHAZ | | 1.04376 | 1.1354 | P63104;E7EX29;E7ESK7;B0AZS6;E9PD24;E7EVZ2;E5RIR4;P63104-2;B7Z | P63104;E7EX29;E7ESK7;B0AZS6 |
| 15 kDa selenoprotein | SEP15 | | 0.403181 | 0.483029 | O60613;A8MZD0;O60613-2 | O60613 |
| 2,3-cyclic-nucleotide 3-phosphodiesterase | CNP | | 0.380875 | -1.17101 | P09543-2;P09543;B4DFN6;K7ERC4;K7EN66;C9K0L8;K7ERZ0 | P09543-2;P09543 |
| 2,4-dienoyl-CoA reductase, mitochondrial | DECR1 | | 0.519543 | -0.251793 | B7Z6B8;Q16698;E5RJD2;E5RJG7;E5RFV2;H0YAW3 | B7Z6B8;Q16698;E5RJD2;E5RJG7;E5RFV2 |
| 26S protease regulatory subunit 10B | PSMC6 | | 0.113575 | 0.157327 | P62333;H0YJC0;H0YJS8;H0YJT1;H0YJE9;H0YJX2;H0YJD2;H0YJY8 | P62333;H0YJC0 |
| 26S protease regulatory subunit 8 | PSMC5 | | 0.244303 | 0.306844 | P62195;P62195-2;J3QQM1;J3QSA9;J3KRP2;J3QLH6;J3QSE0;J3QRW1;J3 | P62195;P62195-2;J3QQM1;J3QSA9;J3KRP2 |
| 26S proteasome non-ATPase regulatory subunit 1 | PSMD1 | | 0.159692 | 0.184627 | Q99460;Q99460-2;C9J9M4;H7BZR6;H7C378;F8WCE3 | Q99460;Q99460-2 |
| 26S proteasome non-ATPase regulatory subunit 11 | PSMD11 | | 0.39748 | 0.570746 | O00231;O00231-2;J3QRY4;J3QS13;J3KSW3 | O00231;O00231-2 |
| 26S proteasome non-ATPase regulatory subunit 12 | PSMD12 | | 0.0673075 | 0.0887761 | O00232;O00232-2;J3KTJ5;J3KSK1 | O00232;O00232-2 |
| 26S proteasome non-ATPase regulatory subunit 13 | PSMD13 | | 0.0837997 | 0.10856 | Q9UNM6;J3KNQ3;Q9UNM6-2;E9PL38;H0YD73;E9PQG3;E9PPD2;F8W8 | Q9UNM6;J3KNQ3;Q9UNM6-2;E9PL38;H0YD73 |
| 26S proteasome non-ATPase regulatory subunit 14 | PSMD14 | | 0.0689181 | 0.0898237 | O00487;C9JW37 | O00487 |
| 26S proteasome non-ATPase regulatory subunit 2 | PSMD2 | | 0.0996199 | 0.106264 | Q13200;E7EW34;E9PCS3;C9JPC0;H7C1H2;H7C2Q3;F8WBS8 | Q13200;E7EW34;E9PCS3 |
| 26S proteasome non-ATPase regulatory subunit 3 | PSMD3 | | 0.0641959 | 0.0747826 | O43242;B4DT72;H0YGV8 | O43242;B4DT72 |
| 26S proteasome non-ATPase regulatory subunit 6 | PSMD6 | | 0.0472352 | -0.0575581 | Q15008;C9IZE4;E9PHI9;Q15008-2;C9J0E9;H7C531;C9J7B7 | Q15008;C9IZE4;E9PHI9;Q15008-2 |
| 26S proteasome non-ATPase regulatory subunit 7 | PSMD7 | | 0.0685086 | -0.109966 | P51665;B4DXI8;H3BNT7;H3BTM8;H3BQV2 | P51665;B4DXI8;H3BNT7;H3BTM8 |
| 26S proteasome non-ATPase regulatory subunit 8 | PSMD8 | | 0.216851 | 0.251874 | R4GMR5;P48556;K7EJR3;K7EJC1;K7ERW6;K7ENY6 | R4GMR5;P48556;K7EJR3;K7EJC1 |
| 2-deoxynucleoside 5-phosphate N-hydrolase 1 | DNPH1 | | 0.802933 | 0.343481 | H0Y8X4;O43598;O43598-2 | H0Y8X4;O43598;O43598-2 |
| 3(2),5-bisphosphate nucleotidase 1 | BPNT1 | | 0.164469 | -0.134121 | O95861;O95861-4;O95861-2;A6NF51;O95861-3;F8W1J0;F8VVW8;F8VZ | O95861;O95861-4;O95861-2;A6NF51;O95861-3 |
| 3-hydroxybutyrate dehydrogenase type 2 | BDH2 | | 0.554613 | 0.776747 | Q9BUT1;D6RBF6;D6RFG2;D6R9U8;D6RIR6;Q9BUT1-2;D6R9P2;Q9BUT1 | Q9BUT1;D6RBF6;D6RFG2;D6R9U8;D6RIR6;Q9BUT1-2 |
| 3-hydroxyisobutyryl-CoA hydrolase, mitochondrial | HIBCH | + | 1.31213 | -0.591996 | Q6NVY1;Q6NVY1-2;B8ZZZ0;B9A058;H7C400;F8W8A6;H7C126;H7BYI7; | Q6NVY1;Q6NVY1-2;B8ZZZ0 |
| 3-mercaptopyruvate sulfurtransferase;Sulfurtransferase | MPST | | 0.564476 | 0.731704 | P25325;J3KPV7;B1AH49 | P25325;J3KPV7;B1AH49 |
| 40S ribosomal protein S10 | RPS10 | | 1.04591 | 1.82321 | P46783;F6U211;Q9NQ39 | P46783;F6U211 |
| 40S ribosomal protein S11 | RPS11 | + | 3.62091 | 1.74477 | P62280;M0QZC5;M0R1H6;M0R1H5 | P62280;M0QZC5 |
| 40S ribosomal protein S13 | RPS13 | | 1.26552 | 1.67141 | P62277;J3KMX5;E9PS50 | P62277;J3KMX5 |
| 40S ribosomal protein S14 | RPS14 | + | 1.59003 | 1.57099 | P62263;H0YB22;E5RH77 | P62263;H0YB22;E5RH77 |
| 40S ribosomal protein S15a | RPS15A | + | 1.31203 | 1.40898 | I3L3P7;P62244;I3L246;H3BV27;H3BT37;I3L303;H3BVC7 | I3L3P7;P62244;I3L246 |
| 40S ribosomal protein S16 | RPS16 | + | 1.59871 | 1.38809 | P62249;M0R210;M0R3H0;M0R1M5;Q6IPX4;M0QX76 | P62249;M0R210;M0R3H0;M0R1M5 |
| 40S ribosomal protein S17-like;40S ribosomal protein S17 | RPS17L;RPS17 | + | 1.51068 | 1.48451 | P0CW22;P08708;H0YN88;H0YK46;H0YN73;H3BNC9 | P0CW22;P08708;H0YN88;H0YK46 |
| 40S ribosomal protein S18 | RPS18 | + | 1.9713 | 1.5061 | P62269;J3JS69 | P62269 |
| 40S ribosomal protein S19 | RPS19 | + | 1.56457 | 1.68867 | P39019;M0R2L9;M0QXK4;M0QYF7;M0R140 | P39019 |
| 40S ribosomal protein S2 | RPS2 | + | 1.36619 | 1.15635 | P15880;H0YEN5;E9PQD7;I3L404;E9PM36;E9PMM9;E9PPT0;H3BNG3;H | P15880;H0YEN5;E9PQD7 |
| 40S ribosomal protein S20 | RPS20 | | 1.15968 | 1.19661 | P60866;P60866-2;E5RJX2;E5RIP1;G3XAN0 | P60866;P60866-2;E5RJX2;E5RIP1 |
| 40S ribosomal protein S25 | RPS25 | + | 2.5609 | 1.19157 | P62851 | P62851 |
| 40S ribosomal protein S26;Putative 40S ribosomal protein S26-like 1 | RPS26;RPS26P11 | + | 1.32408 | 1.41558 | P62854;Q5JNZ5 | P62854;Q5JNZ5 |
| 40S ribosomal protein S28 | RPS28 | | 0.576208 | 1.65511 | P62857 | P62857 |
| 40S ribosomal protein S29 | RPS29 | + | 1.38692 | 1.73468 | P62273;P62273-2 | P62273;P62273-2 |
| 40S ribosomal protein S3 | RPS3 | + | 1.51041 | 1.19483 | P23396;P23396-2;E9PL09;E9PPU1;F2Z2S8;H0YCJ7;H0YEU2;E9PQ96;E9 | P23396;P23396-2;E9PL09;E9PPU1;F2Z2S8;H0YCJ7;H0YEU2 |
| 40S ribosomal protein S30 | FAU | | 0.430587 | 1.44159 | E9PR30;P62861 | E9PR30;P62861 |
| 40S ribosomal protein S3a | RPS3A | + | 1.66032 | 1.86236 | P61247;D6RAT0;H0Y9Y4;D6RG13;D6R9B6;F5H4F9;D6RB09;E9PFI5;H0Y | P61247;D6RAT0;H0Y9Y4;D6RG13;D6R9B6;F5H4F9;D6RB09;E9PFI5;H0Y |
| 40S ribosomal protein S4, X isoform | RPS4X | + | 1.77532 | 1.7353 | P62701;Q8TD47;C9JEH7;P22090;A6NH36 | P62701 |
| 40S ribosomal protein S5;40S ribosomal protein S5, N-terminally processed | RPS5 | + | 1.7214 | 1.86909 | M0R0F0;P46782;M0R0R2;M0QZN2 | M0R0F0;P46782;M0R0R2;M0QZN2 |
| 40S ribosomal protein S7 | RPS7 | + | 1.54085 | 1.44479 | P62081;B5MCP9 | P62081;B5MCP9 |
| 40S ribosomal protein S8 | RPS8 | + | 1.51928 | 1.60583 | P62241;Q5JR95 | P62241;Q5JR95 |
| 40S ribosomal protein S9 | RPS9 | + | 1.39723 | 1.3532 | P46781;B5MCT8;C9JM19;F2Z3C0;A8MXK4 | P46781;B5MCT8;C9JM19 |
| 40S ribosomal protein SA | RPSA;RPSAP58 | + | 1.72823 | 1.31047 | C9J9K3;P08865;A6NE09;C9JQR9 | C9J9K3;P08865;A6NE09 |
| 45 kDa calcium-binding protein | SDF4 | | 0.816455 | -0.574285 | Q9BRK5;Q9BRK5-6;H0Y3T6;G3V1E2;Q9BRK5-4;Q9BRK5-3;Q9BRK5-2;Q | Q9BRK5;Q9BRK5-6;H0Y3T6;G3V1E2;Q9BRK5-4;Q9BRK5-3;Q9BRK5-2 |
| 4F2 cell-surface antigen heavy chain | SLC3A2 | | 0.0434188 | 0.0443413 | F5GZS6;J3KPF3;P08195-2;P08195-3;P08195;P08195-4;H0YFS2;F5H0E2; | F5GZS6;J3KPF3;P08195-2;P08195-3;P08195;P08195-4 |
| 4-trimethylaminobutyraldehyde dehydrogenase | ALDH9A1 | | 0.584093 | 0.507633 | P49189;B4DXY7 | P49189;B4DXY7 |
| 5(3)-deoxyribonucleotidase, cytosolic type | NT5C | | 0.0911618 | -0.0888188 | Q8TCD5;J3KRC4;J3KSX6;Q8TCD5-2;J3KSY6 | Q8TCD5;J3KRC4;J3KSX6;Q8TCD5-2 |
| 5-AMP-activated protein kinase subunit gamma-1 | PRKAG1 | | 0.441617 | -0.48497 | F8W9D1;F8VYY9;P54619-2;P54619;P54619-3;H0YIC9;H0YHF8;F8VPF5; | F8W9D1;F8VYY9;P54619-2;P54619;P54619-3;H0YIC9 |
| 5-nucleotidase | NT5E | | 0.30952 | -0.215192 | P21589-2;P21589;Q96B60;H0Y7R7;H0Y3X5 | P21589-2;P21589 |
| 60 kDa heat shock protein, mitochondrial | HSPD1 | | 0.0994209 | -0.0888914 | P10809;E7EXB4;E7ESH4;B7Z712;C9JL25;C9JCQ4;C9JL19;C9J0S9 | P10809 |
| 60S acidic ribosomal protein P0;60S acidic ribosomal protein P0-like | RPLP0;RPLP0P6 | | 0.633675 | 0.464242 | P05388;F8VWS0;Q3B7A4;F8VU65;F8VW21;F8VPE8;G3V210;F8VQY6;F8 | P05388;F8VWS0;Q3B7A4;F8VU65;F8VW21;F8VPE8;G3V210;F8VQY6;F |
| 60S acidic ribosomal protein P1 | RPLP1 | | 0.191356 | 0.19413 | P05386 | P05386 |
| 60S acidic ribosomal protein P2 | RPLP2 | | 0.626908 | 0.425869 | P05387;H0YDD8;P05386-2 | P05387 |
| 60S ribosomal protein L10a | RPL10A | + | 1.38862 | 1.78277 | P62906 | P62906 |
| 60S ribosomal protein L11 | RPL11 | | 1.18469 | 0.902691 | P62913 | P62913 |
| 60S ribosomal protein L12 | RPL12 | | 0.487651 | 0.659076 | P30050;P30050-2 | P30050;P30050-2 |
| 60S ribosomal protein L13a;Putative 60S ribosomal protein L13a protein RPL13AP3 | RPL13A;RPL13AP3 | | 0.871987 | 1.30043 | P40429;M0QYS1;Q6NVV1;M0QZU1 | P40429;M0QYS1;Q6NVV1 |
| 60S ribosomal protein L14 | RPL14 | | 0.876401 | 1.53843 | P50914;E7EPB3 | P50914;E7EPB3 |
| 60S ribosomal protein L22 | RPL22 | | 1.22621 | 0.608956 | P35268;K7ERI7;K7EMH1;K7EP65;K7EKS7;K7ELC4;K7EJT5 | P35268;K7ERI7;K7EMH1;K7EP65;K7EKS7;K7ELC4 |
| 60S ribosomal protein L23 | RPL23 | | 1.0776 | 0.85876 | P62829;C9JD32;J3KT29;B9ZVP7;J3KTJ3;J3QQT9 | P62829;C9JD32;J3KT29;B9ZVP7 |
| 60S ribosomal protein L23a | RPL23A | | 0.908812 | 0.627847 | P62750;H7BY10;K7EJV9;K7ERT8;A8MUS3;K7EMA7 | P62750;H7BY10;K7EJV9;K7ERT8;A8MUS3;K7EMA7 |
| 60S ribosomal protein L24 | RPL24 | | 1.22614 | 0.920367 | C9JXB8;C9JNW5;P83731 | C9JXB8;C9JNW5;P83731 |
| 60S ribosomal protein L26;60S ribosomal protein L26-like 1 | RPL26;KRBA2;RPL26L1 | | 0.735745 | 0.867563 | J3QRI7;J3QQQ9;J3QQV1;P61254;J3QRC4;J3KTJ8;E5RIT6;Q9UNX3;J3KS | J3QRI7;J3QQQ9;J3QQV1;P61254;J3QRC4;J3KTJ8;E5RIT6;Q9UNX3;J3KS |

| | | | | | | |
|---|---|---|---|---|---|---|
| 60S ribosomal protein L27 | RPL27 | + | 1.92947 | 1.44988 | P61353;K7ELC7;K7EQQ9;K7ERY7 | P61353;K7ELC7;K7EQQ9 |
| 60S ribosomal protein L28 | RPL28 | + | 1.5272 | 1.07005 | H0YMF4;H0YKD8;P46779-4;P46779-5;P46779;P46779-2;P46779-3;H0YL | H0YMF4;H0YKD8;P46779-4;P46779-5;P46779;P46779-2;P46779-3;H0Y |
| 60S ribosomal protein L3 | RPL3 | | 1.07245 | 0.495155 | P39023;G5E9G0;H7C3M2;B5MCW2;H7C422;F8WCR1 | P39023;G5E9G0;H7C3M2;B5MCW2;H7C422 |
| 60S ribosomal protein L30 | RPL30 | + | 1.75386 | 0.875907 | P62888;E5RI99;E5RJH3 | P62888;E5RI99 |
| 60S ribosomal protein L31 | RPL31 | + | 1.41577 | 0.560387 | P62899;H7C2W9;C9JU56;B7Z4E3;B7Z4C8;P62899-3;P62899-2;B8ZZK4 | P62899;H7C2W9;C9JU56;B7Z4E3;B7Z4C8;P62899-3;P62899-2;B8ZZK4 |
| 60S ribosomal protein L34 | RPL34 | | 0.832517 | 0.901368 | P49207 | P49207 |
| 60S ribosomal protein L35a | RPL35A | | 0.80106 | 0.839441 | P18077;C9K025;F8WBS5;F8WB72 | P18077;C9K025;F8WBS5;F8WB72 |
| 60S ribosomal protein L37a | RPL37A | | 1.1264 | 1.00004 | C9J4Z3;P61513;M0R0A1;Q6P4E4;E9PEL3;M0R2L6;G5E9R3;A6NKH3 | C9J4Z3;P61513;M0R0A1;Q6P4E4;E9PEL3;M0R2L6 |
| 60S ribosomal protein L4 | RPL4 | | 1.14569 | 1.16089 | P36578;H3BTP7;H3BU31;H3BM89 | P36578;H3BTP7;H3BU31;H3BM89 |
| 60S ribosomal protein L5 | RPL5 | + | 1.57268 | 2.12801 | P46777;Q5T7N0;R4GNJ2 | P46777 |
| 60S ribosomal protein L7 | RPL7 | | 1.26042 | 1.06854 | P18124;A8MUD9;C9JIJ5;C9JZ88 | P18124;A8MUD9 |
| 60S ribosomal protein L7a | RPL7A | + | 1.74179 | 1.12688 | P62424;Q5T8U2;Q5T8U3 | P62424;Q5T8U2;Q5T8U3 |
| 6-phosphogluconate dehydrogenase, decarboxylating | PGD | | 0.0538609 | 0.0492582 | B4DQJ8;P52209;F5H7U0;K7EM49;K7EMN2;K7EPF6;K7ELN9 | B4DQJ8;P52209;F5H7U0 |
| 6-phosphogluconolactonase | PGLS | | 0.429631 | -0.245328 | O95336;M0R261;M0R0U3;M0R1L2 | O95336;M0R261;M0R0U3 |
| 7,8-dihydro-8-oxoguanine triphosphatase | NUDT1 | | 0.0643923 | 0.0487939 | P36639-4;P36639-3;P36639-2;P36639;C9J361 | P36639-4;P36639-3;P36639-2;P36639 |
| 78 kDa glucose-regulated protein | HSPA5 | | 0.0556889 | 0.0541814 | P11021;Q5H8X8;O95399;O95399-2 | P11021 |
| A disintegrin and metalloproteinase with thrombospondin motifs 1 | ADAMTS1 | | 0.348238 | 0.48636 | Q9UHI8;H7C206;E5RI60;E5RJR7;Q8TE58 | Q9UHI8 |
| Acetyl-CoA acetyltransferase, cytosolic | ACAT2 | | 0.305833 | 0.181906 | Q9BWD1;B7Z233 | Q9BWD1;B7Z233 |
| Acetyl-CoA acetyltransferase, mitochondrial | ACAT1 | | 0.829919 | 0.574404 | P24752;G3XAB4;H0YEL7;E9PRQ6;E9PKF3 | P24752;G3XAB4 |
| Acid ceramidase;Acid ceramidase subunit alpha;Acid ceramidase subunit beta | ASAH1 | | 0.379284 | -0.140558 | E7EMM4;Q13510;Q13510-2;E7ERV9;Q13510-3 | E7EMM4;Q13510;Q13510-2;E7ERV9;Q13510-3 |
| Acidic leucine-rich nuclear phosphoprotein 32 family member B | ANP32B | | 0.280234 | -0.226962 | Q92688-2;Q92688;Q5T6W8;O95626 | Q92688-2;Q92688;Q5T6W8 |
| Actin, alpha skeletal muscle;Actin, alpha cardiac muscle 1;Actin, gamma-enteric smooth muscle;Actin, aor | ACTA1;ACTC1;ACTG2;ACTA2 | | 0.0669984 | 0.0820046 | P68133;P68032;Q5T8M8;A6NL76;P63267;P62736;Q5T8M7;P63267-2;F | P68133;P68032;Q5T8M8;A6NL76;P63267;P62736;Q5T8M7;P63267-2 |
| Actin, cytoplasmic 1;Actin, cytoplasmic 1, N-terminally processed | ACTB | | 0.783925 | 0.76532 | P60709;G5E9R0;E7EVS6;C9JZR7;C9JTX5 | P60709 |
| Actin, cytoplasmic 2;Actin, cytoplasmic 2, N-terminally processed | ACTG1 | | 0.263201 | 0.18897 | P63261;I3L3I0;I3L1U9;I3L4N8;I3L3R2;J3KT65;K7EM38;C9JUM1 | P63261;I3L3I0;I3L1U9 |
| Actin-related protein 2 | ACTR2 | | 0.0712601 | -0.0582011 | P61160;P61160-2;F5H6T1 | P61160;P61160-2;F5H6T1 |
| Actin-related protein 2/3 complex subunit 1A | ARPC1A | | 0.0961212 | 0.100146 | Q92747;E9PF58;B4DLQ7;F8WFD3 | Q92747;E9PF58;B4DLQ7 |
| Actin-related protein 2/3 complex subunit 1B | ARPC1B | | 0.0983774 | 0.0819602 | O15143;C9J4Z7;F8VXW2;C9JEY1;C9JQM8;C9JFG9;C9J6C8;C9JBJ7;C9K05 | O15143;C9J4Z7;F8VXW2;C9JEY1;C9JQM8;C9JFG9;C9J6C8 |
| Actin-related protein 2/3 complex subunit 2 | ARPC2 | | 0.0100804 | -0.00938658 | O15144;H7C3F9;C9JTV5;G5E9J0;G5E9S7 | O15144 |
| Actin-related protein 2/3 complex subunit 3 | ARPC3 | | 0.0121313 | -0.0130479 | O15145;C9JZD1;F8VR50 | O15145;C9JZD1 |
| Actin-related protein 2/3 complex subunit 4 | ARPC4;ARPC4-TTLL3 | | 0.225488 | -0.1809 | P59998;F8WCF6;P59998-3;F8WDD7;P59998-2;H7C0A3;P59998-4;R4GN | P59998;F8WCF6;P59998-3;F8WDD7;P59998-2;H7C0A3 |
| Actin-related protein 2/3 complex subunit 5 | ARPC5 | | 0.0554555 | -0.0464472 | O15511;B1ALC0;O15511-2 | O15511;B1ALC0;O15511-2 |
| Actin-related protein 2/3 complex subunit 5-like protein | ARPC5L | | 0.127094 | 0.112492 | Q9BPX5 | Q9BPX5 |
| Actin-related protein 3 | ACTR3 | | 0.10336 | -0.0800102 | P61158;B4DXW1;F5H3P5;Q9P1U1-2;Q9P1U1;Q9P1U1-3;F8WEW2;F8W | P61158;B4DXW1;F5H3P5 |
| Activated RNA polymerase II transcriptional coactivator p15 | SUB1 | | 0.88243 | -0.815452 | P53999;D6RC37;D6R970 | P53999 |
| Activator of 90 kDa heat shock protein ATPase homolog 1 | AHSA1 | | 0.526856 | -0.515488 | O95433;B4DUR9;G3V438;H0YJG7;H0YJU2;G3V3W9;H0YJ63;H0YJG6 | O95433;B4DUR9;G3V438;H0YJG7 |
| Acylamino-acid-releasing enzyme | APEH | | 0.0101517 | -0.00839581 | C9JIF9;P13798;H7C1U0;H7C393;C9JLK2;H0YFE5;F8WEH5 | C9JIF9;P13798;H7C1U0;H7C393 |
| Acyl-CoA-binding protein | DBI | | 0.112075 | -0.0583989 | P07108;P07108-3;P07108-2;P07108-4;P07108-5;P07108-6 | P07108;P07108-3;P07108-2;P07108-4;P07108-5 |
| Acyl-coenzyme A thioesterase 1;Acyl-coenzyme A thioesterase 2, mitochondrial | ACOT1;ACOT2 | | 0.159017 | -0.0988184 | Q86TX2;G3V4F2;P49753;B3KSA0;P49753-2 | Q86TX2;G3V4F2;P49753 |
| Acyl-coenzyme A thioesterase 13;Acyl-coenzyme A thioesterase 13, N-terminally processed | ACOT13 | | 0.116058 | -0.183115 | Q9NPJ3-2;Q9NPJ3 | Q9NPJ3-2;Q9NPJ3 |
| Acylphosphatase-1;Acylphosphatase | ACYP1;FBN3 | | 0.880324 | -0.785843 | P07311;G3V2U7;G3V3F8;G3V597;P07311-2 | P07311;G3V2U7 |
| Acyl-protein thioesterase 1 | LYPLA1 | | 0.206159 | -0.238667 | E5RGR0;O75608-2;O75608;E5RJ48;E5RI35 | E5RGR0;O75608-2;O75608 |
| Acyl-protein thioesterase 2 | LYPLA2 | | 0.232568 | -0.282983 | O95372;Q5QPN5;E7EPS4;Q5QPN9;Q5QPQ1;Q5QPQ0;Q5QPQ3;Q5QPQ | O95372;Q5QPN5;E7EPS4;Q5QPN9;Q5QPQ1;Q5QPQ0;Q5QPQ3;Q5QPC |
| ADAMTS-like protein 1 | ADAMTSL1 | | 0.455661 | -0.983615 | Q8N6G6-4;Q8N6G6;Q8N6G6-1;Q8N6G6-2;F8WEP3;A6NIB9 | Q8N6G6-4;Q8N6G6;Q8N6G6-1;Q8N6G6-2 |
| Adapter molecule crk | CRK | | 0.128352 | 0.097433 | P46108;I3L297 | P46108 |
| Adenine phosphoribosyltransferase | APRT | | 0.661564 | 0.666007 | P07741;P07741-2;H3BQZ9;H3BQB1;H3BQF1;H3BSW3 | P07741;P07741-2;H3BQZ9;H3BQB1;H3BQF1 |
| Adenosylhomocysteinase | AHCY | | 0.148416 | 0.109313 | P23526;P23526-2 | P23526;P23526-2 |
| Adenylate kinase 2, mitochondrial;Adenylate kinase 2, mitochondrial;Adenylate kinase 2, mitochondrial, I | AK2 | | 0.0643901 | -0.0537646 | F8W1A4;P54819-2;P54819;P54819-3;P54819-5;F8VZG5;F8VY04;G3V21 | F8W1A4;P54819-2;P54819;P54819-3;P54819-5;F8VZG5;F8VY04;G3V21 |
| Adenylate kinase 4, mitochondrial | AK4 | | 0.719328 | 0.364635 | P27144;D3DQ64 | P27144 |
| Adenylate kinase isoenzyme 1 | AK1 | | 0.433251 | -0.349813 | Q5T9B7;P00568;H0Y4J6;H0YID2;Q9Y6K8-2;Q9Y6K8-3;Q9Y6K8 | Q5T9B7;P00568;H0Y4J6 |
| Adenylosuccinate synthetase isozyme 2 | ADSS | | 0.164489 | 0.138836 | P30520;Q8N142;G3V5D8;G3V232;Q8N142-2 | P30520 |
| Adenylyl cyclase-associated protein 1 | CAP1 | | 0.0636118 | 0.0688287 | Q01518;Q5T0R1;Q5T0R4;Q5T0R3;Q5T0R2;Q5T0R5;Q5T0R7;Q5T0R6 | Q01518 |
| Adenylyl cyclase-associated protein 1 | CAP1 | | 0.191199 | 0.202888 | Q01518-2;Q5T0R9;Q5T0S3;Q5T0R8 | Q01518-2 |
| ADM;Adrenomedullin;Proadrenomedullin N-20 terminal peptide | ADM | | 0.803343 | 3.17594 | P35318;E9PL83;E9PQE6;E9PML4;E9PQT2 | P35318;E9PL83;E9PQE6 |
| ADP/ATP translocase 2;ADP/ATP translocase 2, N-terminally processed | SLC25A5 | | 0.472711 | -1.0384 | P05141 | P05141 |
| ADP-dependent glucokinase | ADPGK | | 0.0836305 | -0.273449 | Q9BRR6-2;Q9BRR6;Q9BRR6-4;Q9BRR6-3;H3BTH4;H3BN66;H3BRS6;Q9I | Q9BRR6-2;Q9BRR6;Q9BRR6-4;Q9BRR6-3 |
| ADP-ribose pyrophosphatase, mitochondrial | NUDT9 | | 0.324334 | 0.446484 | Q9BW91-2;Q9BW91;D6RAW2;H7C386;D6R8Z6 | Q9BW91-2;Q9BW91;D6RAW2;H7C386 |
| ADP-ribosylation factor 1 | ARF1 | | 0.408538 | -0.297111 | P84077;H0YGG7 | P84077 |
| ADP-ribosylation factor 4 | ARF4 | | 0.203063 | 0.137846 | C9JPM4;P18085;C9JAK5;U3KQF2 | C9JPM4;P18085;C9JAK5 |
| ADP-ribosylation factor 5 | ARF5 | | 1.21749 | -0.339172 | P84085;C9J1Z8;F8WDB3 | P84085;C9J1Z8 |
| ADP-ribosylation factor 6 | ARF6 | | 1.26969 | -1.35487 | P62330;P36406-3;P36406-2;P36406 | P62330 |
| ADP-ribosylation factor-like protein 1 | ARL1 | | 0.169907 | -0.174976 | B4DWW1;P40616;B4DZG7;F8VYN9;F8VP99;F8W1Z8;F8VP63 | B4DWW1;P40616;B4DZG7;F8VYN9 |
| ADP-ribosylation factor-like protein 2 | ARL2;ARL2-SNX15 | | 0.0564252 | -0.0645083 | P36404;V9GYD0;P36404-2 | P36404;V9GYD0;P36404-2 |
| ADP-ribosylation factor-like protein 3 | ARL3 | | 0.411571 | 0.620387 | P36405 | P36405 |
| ADP-ribosylation factor-like protein 8B;ADP-ribosylation factor-like protein 8A | ARL8B;ARL8A | | 0.737752 | -0.470766 | Q9NVJ2;B4DI85;Q96BM9 | Q9NVJ2;B4DI85;Q96BM9 |
| ADP-sugar pyrophosphatase | NUDT5 | | 0.202678 | 0.158052 | A6NFX8;Q9UKK9;A6NJU6;A6NCQ0;C9JYY9;H0YEY4 | A6NFX8;Q9UKK9;A6NJU6;A6NCQ0 |
| Aflatoxin B1 aldehyde reductase member 2 | AKR7A2 | | 0.712091 | 0.953597 | O43488;H3BLU7;Q8NHP1;H7C5H7;O95154 | O43488;H3BLU7 |
| Agrin;Agrin N-terminal 110 kDa subunit;Agrin C-terminal 110 kDa subunit;Agrin C-terminal 90 kDa fragme | AGRN | | 0.0534019 | -0.0797621 | O00468-6;O00468-7;O00468-3;O00468-5;O00468-4;O00468;O00468-2; | O00468-6;O00468-7;O00468-3;O00468-5;O00468-4;O00468;O00468-2 |
| Alanine--tRNA ligase, cytoplasmic | AARS | | 0.456056 | 0.548812 | P49588;H3BPK7 | P49588 |
| Alcohol dehydrogenase [NADP(+)] | AKR1A1 | | 0.175798 | 0.0996279 | P14550;Q5T621;V9GYG2;V9GYP9;G3V1C1;Q96JD6-4;Q96JD6 | P14550 |
| Alcohol dehydrogenase class-3 | ADH5 | | 0.121365 | -0.158888 | P11766;H0YAG8;D6RFE4;D6RAY0;D6R9G2 | P11766;H0YAG8 |
| Aldehyde dehydrogenase family 16 member A1 | ALDH16A1 | | 0.361579 | 0.469321 | Q8IZ83;Q8IZ83-3;F8WBH3;F5H4B6;Q8IZ83-2;M0QY06 | Q8IZ83;Q8IZ83-3;F8WBH3;F5H4B6 |
| Aldo-keto reductase family 1 member C3;Aldo-keto reductase family 1 member C1 | AKR1C3;AKR1C1 | | 0.457487 | -0.996684 | S4R3Z2;P42330;B4DL37;Q04828;S4R3D5;A6NHU4;H0Y804;B4DK69;P17 | S4R3Z2;P42330;B4DL37;Q04828 |
| Aldose 1-epimerase | GALM | | 0.919242 | -0.734812 | Q96C23;F8WB82;B8ZZ75;H7C320;H7C1B5 | Q96C23;F8WB82;B8ZZ75;H7C320 |
| Aldose reductase | AKR1B1 | | 0.291466 | 0.207497 | P15121;E9PCX2;E9PEF9;C9JRZ8-2;C9JRZ8 | P15121;E9PCX2 |
| Alpha/beta hydrolase domain-containing protein 14A | ABHD14A | | 0.146566 | -0.155101 | Q9BUJ0;C9JMW6;C9J7W8 | Q9BUJ0;C9JMW6;C9J7W8 |
| Alpha/beta hydrolase domain-containing protein 14B | ABHD14B | | 1.25356 | -0.912674 | Q96IU4;F8W9U3;B4DQI4 | Q96IU4;F8W9U3;B4DQI4 |
| Alpha-1,3-mannosyl-glycoprotein 2-beta-N-acetylglucosaminyltransferase | MGAT1 | | 0.961742 | 0.312339 | P26572;D6RF69;D6RA48;D6RBS3;D6RAK2;D6R9U2;D6RD15;D6RB69;D6 | P26572 |
| Alpha-1,6-mannosylglycoprotein 6-beta-N-acetylglucosaminyltransferase A | MGAT5 | | 0.982246 | -0.537479 | Q09328;C9J0J0;Q96EE4 | Q09328 |
| Alpha-2-macroglobulin receptor-associated protein | LRPAP1 | | 0.251227 | 0.285357 | P30533;D6REW6 | P30533 |

| | | | | | | |
|---|---|---|---|---|---|---|
| Alpha-actinin-1 | ACTN1 | | 0.536391 | -0.297314 | P12814;P12814-3;P12814-2;P12814-4;H9KV75;G3V2W4;G3V2N5;H7C5 | P12814;P12814-3;P12814-2;P12814-4;H9KV75 |
| Alpha-actinin-2 | ACTN2 | | 0.366813 | -0.179172 | P35609;B2RCS5 | P35609;B2RCS5 |
| Alpha-actinin-4 | ACTN4 | | 0.451611 | -0.241258 | O43707;O43707-2;F5GXS2;O43707-3;H7C144;K7EP19 | O43707;O43707-2 |
| Alpha-aminoadipic semialdehyde dehydrogenase | ALDH7A1 | | 0.0309596 | 0.0285306 | P49419-2;P49419;F8VS02;P49419-4;P49419-3;H0YHM6;F8VVF2 | P49419-2;P49419;F8VS02;P49419-4;P49419-3 |
| Alpha-centractin | ACTR1A | | 0.0432575 | 0.0521885 | P61163;R4GMT0;F5H3I4;B4DXP9;P42025 | P61163;R4GMT0;F5H3I4;B4DXP9 |
| Alpha-enolase | ENO1 | | 0.91729 | -0.539056 | P06733;P06733-2;K7EM90;E5RI09;E5RG95;K7EPM1;K7EKN2;E5RGZ4;K | P06733;P06733-2 |
| Alpha-galactosidase A | GLA | | 0.531658 | 0.180864 | P06280;V9GYN5 | P06280 |
| Alpha-L-iduronidase | IDUA | | 0.2747 | -0.367279 | D6REB5;P35475;B3KWK6;D6RBD5;D6R9D5;H0Y9B3 | D6REB5;P35475;B3KWK6 |
| Alpha-mannosidase 2 | MAN2A1 | | 0.856163 | -0.755309 | Q16706 | Q16706 |
| Alpha-N-acetylgalactosaminidase | NAGA | | 0.163683 | -0.138744 | P17050 | P17050 |
| Alpha-N-acetylglucosaminidase;Alpha-N-acetylglucosaminidase 82 kDa form;Alpha-N-acetylglucosaminid | NAGLU | | 0.426842 | -0.219381 | P54802;K7ENX5;K7EQH9 | P54802 |
| Alpha-soluble NSF attachment protein | NAPA | | 0.520496 | 0.460918 | P54920;M0R0Y2;M0R2M1;M0R058;M0R0I4;M0R213;M0R027;Q4G0M | P54920;M0R0Y2;M0R2M1 |
| Alpha-taxilin | TXLNA | | 0.564212 | 0.868731 | P40222 | P40222 |
| Aminoacyl tRNA synthase complex-interacting multifunctional protein 1;Endothelial monocyte-activating | AIMP1 | | 0.26471 | -0.288728 | Q12904;Q12904-2;D6R937 | Q12904;Q12904-2 |
| Aminoacylase-1 | ACY1;ABHD14A | | 0.0232952 | -0.0109537 | Q03154;B4DNW0;Q03154-4;Q03154-2;Q03154-3;C9JYZ0;F8WC59 | Q03154;B4DNW0;Q03154-4;Q03154-2;Q03154-3 |
| Aminopeptidase B | RNPEP | | 0.125433 | -0.0815268 | Q9H4A4;A6NKB8;C9JMZ3;H7C2T3 | Q9H4A4;A6NKB8 |
| Amyloid beta A4 protein;N-APP;Soluble APP-alpha;Soluble APP-beta;C99;Beta-amyloid protein 42;Beta-a | APP | | 0.632758 | 0.213141 | P05067-7;P05067-11;P05067-8;P05067-9;P05067;E9PEV0;P05067-3;P0 | P05067-7;P05067-11;P05067-8;P05067-9;P05067;E9PEV0;P05067-3;P0 |
| Amyloid-like protein 2 | APLP2 | | 0.0885932 | 0.0690782 | Q06481;Q06481-3;Q06481-6;Q06481-2;Q06481-4;F5GZY0;Q06481-5;F5 | Q06481;Q06481-3;Q06481-6;Q06481-2;Q06481-4;F5GZY0;Q06481-5;F |
| Angiogenin | ANG | | 0.629943 | -0.999497 | P03950 | P03950 |
| Ankyrin repeat and FYVE domain-containing protein 1 | ANKFY1 | | 0.161177 | 0.590611 | Q9P2R3;Q9P2R3-2;I3L3P8 | Q9P2R3;Q9P2R3-2 |
| Annexin A1 | ANXA1 | | 0.632998 | -0.611256 | P04083;Q5T3N1;Q5T3N0 | P04083 |
| Annexin A11 | ANXA11 | | 0.410691 | -0.736742 | P50995-2;P50995;H0Y6E1 | P50995-2;P50995 |
| Annexin A2;Annexin;Putative annexin A2-like protein | ANXA2;ANXA2P2 | | 1.07482 | -1.46761 | P07355;P07355-2;H0YN42;H0YMD0;H0YMU9;H0YM50;A6NMY6;H0YN | P07355;P07355-2;H0YN42;H0YMD0;H0YMU9;H0YM50;A6NMY6;H0YN |
| Annexin A3;Annexin | ANXA3 | | 0.522478 | -0.495703 | P12429;D6RA82;D6RFG5;D6RCA8;D6RAZ8;D6RFJ9 | P12429;D6RA82 |
| Annexin A4;Annexin | ANXA4 | | 0.548339 | -0.591859 | P09525;Q6P452;B4DDF9 | P09525;Q6P452;B4DDF9 |
| Annexin A5;Annexin | ANXA5 | | 0.595138 | -0.639504 | P08758;D6RBL5;D6RBE9;E9PHT9;D6RCN3 | P08758;D6RBL5;D6RBE9;E9PHT9 |
| Annexin A6;Annexin | ANXA6 | | 0.728659 | -0.332524 | A6NN80;P08133-2;P08133;E5RK69;E7EMC6;E5RFF0;H0YC77;E5RIU8;E5 | A6NN80;P08133-2;P08133;E5RK69;E7EMC6 |
| Annexin;Annexin A7 | ANXA7 | | 0.930895 | -0.493581 | B4DT77;P20073-2;P20073 | B4DT77;P20073-2;P20073 |
| Anthrax toxin receptor 1 | ANTXR1 | | 0.95885 | -0.561356 | Q9H6X2-5;Q9H6X2;Q9H6X2-3;Q9H6X2-4;Q9H6X2-6;Q9H6X2-2 | Q9H6X2-5;Q9H6X2;Q9H6X2-3;Q9H6X2-4;Q9H6X2-6;Q9H6X2-2 |
| Anthrax toxin receptor 2 | ANTXR2 | | 0.0457321 | -0.133373 | P58335-2;P58335;J3KPY9;P58335-4;P58335-3 | P58335-2;P58335;J3KPY9;P58335-4;P58335-3 |
| AP-1 complex subunit mu-1 | AP1M1 | | 0.256274 | -0.226198 | Q9BXS5;Q9BXS5-2;B4DDG7;K7EJL1;E7ENJ6;K7ENA7;K7EQX3;Q9Y6Q5;C | Q9BXS5;Q9BXS5-2;B4DDG7;K7EJL1;E7ENJ6 |
| AP-1 complex subunit sigma-1A | AP1S1 | | 0.247337 | 0.374094 | H7C1E4;P61966;P61966-2 | H7C1E4;P61966;P61966-2 |
| AP-2 complex subunit mu | AP2M1 | | 0.206657 | -0.199166 | E9PFW3;Q96CW1-2;Q96CW1;C9JJ47;C9JTK4;C9JGT8;C9JJD3;C9JPV8;H7 | E9PFW3;Q96CW1-2;Q96CW1;C9JJ47 |
| Apolipoprotein B-100;Apolipoprotein B-48 | APOB | | 0.427458 | 0.266688 | P04114;A8MUN2 | P04114 |
| Apoptosis regulator BAX | BAX | | 0.161388 | 0.136354 | Q07812-8;Q07812;Q07812-2;Q07812-5;Q07812-7;Q07812-6;Q07812-4 | Q07812-8;Q07812;Q07812-2;Q07812-5;Q07812-7 |
| Arginine--tRNA ligase, cytoplasmic | RARS | | 0.00179852 | -0.00298687 | P54136;P54136-2;F5H3T8;E5RJM9;E5RH09 | P54136;P54136-2;F5H3T8 |
| Arrestin domain-containing protein 1 | ARRDC1 | | 0.681792 | -0.664608 | Q5T370;Q8N5I2;Q5T371;F2Z3I4 | Q5T370;Q8N5I2;Q5T371;F2Z3I4 |
| Asparagine--tRNA ligase, cytoplasmic | NARS | | 0.81863 | 0.456047 | O43776;B4DN60;K7EIU7;K7EQ35;B4DG16;K7EMQ6;K7EPK2;K7EJ19 | O43776;B4DN60 |
| Aspartate aminotransferase, cytoplasmic | GOT1 | | 0.0116759 | -0.00815151 | P17174;B7Z7E9 | P17174;B7Z7E9 |
| Aspartate aminotransferase, mitochondrial | GOT2 | | 0.0655061 | 0.052191 | P00505;P00505-2 | P00505;P00505-2 |
| Aspartate--tRNA ligase, cytoplasmic | DARS | | 0.0212954 | 0.0329773 | P14868;Q68CR9;C9J7S3;C9JLC1;H7BZ35;H7C278;C9JQM9 | P14868;Q68CR9 |
| Aspartyl aminopeptidase | DNPEP | | 0.116551 | 0.131537 | E7ETB3;Q9ULA0;E7EMB6;C9JBE1;F8WAN0;B9ZVU2;E7EPX3;C9J1E2;E5 | E7ETB3;Q9ULA0;E7EMB6;C9JBE1 |
| Ataxin-10 | ATXN10 | | 0.111894 | 0.12357 | Q9UBB4;Q9UBB4-2;B1AHE4;H0Y7N2;B1AHE3 | Q9UBB4;Q9UBB4-2 |
| ATP synthase subunit alpha, mitochondrial | ATP5A1 | | 0.742555 | -0.508523 | P25705-2;P25705;P25705-3;K7EK77;K7ENJ4;K7ERX7;K7EJP1;K7EQH4;K | P25705-2;P25705;P25705-3;K7EK77 |
| ATP synthase subunit beta, mitochondrial;ATP synthase subunit beta | ATP5B | | 0.352314 | 0.13677 | P06576;F8VPV9;H0YH81;F8W079;F8W0P7;H0YI37;F8VQY0 | P06576;F8VPV9;H0YH81;F8W079 |
| ATP synthase subunit d, mitochondrial | ATP5H | | 0.526099 | 1.17509 | O75947-2;O75947;F5H608 | O75947-2;O75947;F5H608 |
| ATP synthase subunit gamma;ATP synthase subunit gamma, mitochondrial | ATP5C1 | | 0.347654 | -0.698622 | B4DL14;P36542-2;P36542 | B4DL14;P36542-2;P36542 |
| ATP synthase subunit O, mitochondrial | ATP5O | | 0.338716 | -0.787623 | P48047;H7C0C1;H7C086;H7C068 | P48047;H7C0C1 |
| ATPase ASNA1 | ASNA1 | | 0.692984 | 1.2633 | O43681;K7ERW9 | O43681;K7ERW9 |
| ATP-citrate synthase | ACLY | | 0.498275 | -0.305555 | P53396;P53396-2;B4E3P0;K7ESG8;K7EIE7 | P53396;P53396-2;B4E3P0 |
| ATP-dependent 6-phosphofructokinase, liver type | PFKL | | 0.231648 | -0.26136 | P17858;P17858-2;F8WEU2 | P17858;P17858-2 |
| ATP-dependent 6-phosphofructokinase, muscle type | PFKM | | 0.443382 | 0.636132 | P08237;P08237-3;P08237-2;F8VX13;F8VNX2;F8VZQ1;F8VP00;H0YHB8; | P08237;P08237-3;P08237-2 |
| ATP-dependent 6-phosphofructokinase, platelet type | PFKP | | 0.322118 | 0.413072 | Q01813;Q01813-2;B1APP6;B1APP8;Q5VSR5;H0Y757;V9GYV7;H0Y3Y3; | Q01813;Q01813-2 |
| ATP-dependent RNA helicase DDX19A;ATP-dependent RNA helicase DDX19B | DDX19A;DDX19B | | 0.562012 | 1.27674 | I3L0H8;Q9NUU7;H3BQK0;Q9UMR2-2;Q9UMR2-4;Q9UMR2;B4DRZ7;I3L | I3L0H8;Q9NUU7;H3BQK0;Q9UMR2-2;Q9UMR2-4;Q9UMR2;B4DRZ7;I3 |
| ATP-dependent RNA helicase DDX3X;ATP-dependent RNA helicase DDX3Y | DDX3X;DDX3Y | | 1.12646 | 0.205706 | O00571-2;O00571;B4DXX7;O15523 | O00571-2;O00571;B4DXX7;O15523 |
| ATP-dependent RNA helicase DHX36 | DHX36 | | 0.698881 | 0.501958 | F5GZS0;Q9H2U1-3;Q9H2U1-2;Q9H2U1;E7EWK3;H7C5F5 | F5GZS0;Q9H2U1-3;Q9H2U1-2;Q9H2U1;E7EWK3 |
| Attractin | ATRN | | 0.201505 | -0.126268 | O75882-3;O75882-2;O75882 | O75882-3;O75882-2;O75882 |
| Band 4.1-like protein 2 | EPB41L2 | | 0.944012 | 1.08233 | O43491;E9PHY5;O43491-4;E9PK52;O43491-3;E9PII3;O43491-2;Q6ZSX4 | O43491;E9PHY5;O43491-4;E9PK52;O43491-3;E9PII3;O43491-2 |
| Barrier-to-autointegration factor;Barrier-to-autointegration factor, N-terminally processed | BANF1 | | 1.04878 | 1.23614 | O75531 | O75531 |
| Basal cell adhesion molecule | BCAM | | 0.705498 | -0.758333 | P50895;K7ENU8 | P50895;K7ENU8 |
| Basement membrane-specific heparan sulfate proteoglycan core protein;Endorepellin;LG3 peptide | HSPG2 | | 0.277429 | 0.426926 | P98160;Q5SZJ2;Q5SZJ1;H7C4A6;Q5SZI9;H0Y5A9 | P98160 |
| Basigin | BSG | | 0.235345 | -0.261545 | P35613-2;P35613;P35613-4;I3L192 | P35613-2;P35613;P35613-4;I3L192 |
| Beta-1,3-galactosyl-O-glycosyl-glycoprotein beta-1,6-N-acetylglucosaminyltransferase | GCNT1 | | 0.154903 | 0.115265 | Q02742 | Q02742 |
| Beta-1,3-N-acetylglucosaminyltransferase lunatic fringe | LFNG | | 0.571437 | 0.217276 | Q8NES3;Q8NES3-3;Q8NES3-2;Q8NES3-4 | Q8NES3;Q8NES3-3;Q8NES3-2;Q8NES3-4 |
| Beta-1,4-galactosyltransferase 1;Lactose synthase A protein;N-acetyllactosamine synthase;Beta-N-acety | B4GALT1 | + | 1.51555 | -0.243741 | P15291-2;P15291;B4DMM8;Q86XA6 | P15291-2;P15291;B4DMM8 |
| Beta-1,4-galactosyltransferase 5 | B4GALT5 | | 0.334776 | -0.089197 | O43286;Q6NT09 | O43286 |
| Beta-2-microglobulin;Beta-2-microglobulin form pI 5.3 | B2M | | 0.781676 | -0.72638 | P61769;F5H6I0;H0YLF3 | P61769;F5H6I0;H0YLF3 |
| Beta-actin-like protein 2 | ACTBL2 | | 0.428862 | 0.427294 | Q562R1 | Q562R1 |
| Beta-galactosidase | GLB1 | | 0.123278 | -0.0552301 | P16278-3;P16278;E7EQ29;P16278-2;F8WF40;C9JF15;C9JWX1;C9J539;C | P16278-3;P16278;E7EQ29;P16278-2 |
| Beta-glucuronidase | GUSB | | 0.644113 | -0.227864 | P08236-2;P08236;P08236-3;F8WBK6;F2Z3L6 | P08236-2;P08236;P08236-3 |
| Beta-hexosaminidase subunit alpha;Beta-hexosaminidase | HEXA | | 0.769178 | -0.682682 | P06865;H3BS10;H3BP20;E9PGL4;H3BTD4;H3BU85;B4DKE7;H3BVH8;H3 | P06865;H3BS10;H3BP20;E9PGL4;H3BTD4;H3BU85 |
| Beta-hexosaminidase subunit beta;Beta-hexosaminidase subunit beta chain B;Beta-hexosaminidase subu | HEXB | | 0.128059 | -0.0693092 | P07686;Q5URX0;H0YA83;H0Y9B6;H0Y9M3;D6REQ8 | P07686;Q5URX0;H0YA83 |
| BH3-interacting domain death agonist;BH3-interacting domain death agonist p15;BH3-interacting domai | BID | | 0.778284 | -0.411856 | P55957;P55957-2;P55957-4;P55957-3 | P55957;P55957-2;P55957-4 |
| Bifunctional ATP-dependent dihydroxyacetone kinase/FAD-AMP lyase (cyclizing);ATP-dependent dihydro | DAK | | 0.0559321 | 0.0463586 | Q3LXA3;I3L252;H0YCY6 | Q3LXA3;I3L252;H0YCY6 |
| Bifunctional purine biosynthesis protein PURH;Phosphoribosylaminoimidazolecarboxamide formyltransf | ATIC | | 0.313605 | 0.248666 | P31939;P31939-2;F5GWY2;H7C1S2;H7C084;C9JLK0;F8WEF0;H7BZU3 | P31939;P31939-2;F5GWY2 |
| Biliverdin reductase A | BLVRA | | 0.250719 | -0.213452 | P53004;C9J1E1 | P53004 |
| Biogenesis of lysosome-related organelles complex 1 subunit 1 | BLOC1S1 | | 0.448524 | -0.453772 | G8JLQ3;P78537-2;P78537;F8W099;F8W606;F8W036;F8VP73 | G8JLQ3;P78537-2;P78537;F8W099;F8W606;F8W036;F8VP73 |
| Biotinidase | BTD | | 0.0818553 | -0.0787714 | F8W1Q3;P43251;B4DLJ9;A6NHF2;C9JSN9 | F8W1Q3;P43251;B4DLJ9;A6NHF2;C9JSN9 |
| Bis(5-adenosyl)-triphosphatase | FHIT | | 0.353416 | -0.661791 | P49789;E9PBZ0 | P49789;E9PBZ0 |

| | | | | | | |
|---|---|---|---|---|---|---|
| Bleomycin hydrolase | BLMH | | 1.17662 | -0.700321 | Q13867;K7ESE8;E7EMN3;J3KSD8;K7ES02;J3KS79;K7ENH5 | Q13867;K7ESE8;E7EMN3;J3KSD8;K7ES02 |
| BolA-like protein 2 | BOLA2 | | 0.989059 | 1.15416 | Q9H3K6;Q9H3K6-2;H3BV85;H3BTW0;H3BVE0 | Q9H3K6;Q9H3K6-2 |
| Bone morphogenetic protein 4 | BMP4 | | 0.120693 | 0.380463 | P12644;H0YLW3;H0YMP9;H0YM53 | P12644;H0YLW3 |
| Brain acid soluble protein 1 | BASP1 | | 0.041269 | 0.115261 | P80723;P80723-2 | P80723;P80723-2 |
| BRO1 domain-containing protein BROX | BROX | | 0.331269 | -0.300112 | Q5VW32;F5GXQ0;B7Z9G5;Q5VW33 | Q5VW32;F5GXQ0;B7Z9G5;Q5VW33 |
| BTB/POZ domain-containing protein KCTD12 | KCTD12 | | 0.885288 | -0.612889 | Q96CX2;Q6ZWB6 | Q96CX2 |
| C1GALT1-specific chaperone 1 | C1GALT1C1 | | 0.0874414 | 0.0951217 | Q96EU7 | Q96EU7 |
| C-1-tetrahydrofolate synthase, cytoplasmic;Methylenetetrahydrofolate dehydrogenase;Methenyltetrahy | MTHFD1 | | 0.60798 | 0.911561 | P11586;F5H2F4;V9GYY3;Q6UB35 | P11586;F5H2F4 |
| CAD protein;Glutamine-dependent carbamoyl-phosphate synthase;Aspartate carbamoyltransferase;Dihy | CAD | | 0.0656598 | -0.0477818 | F8VPD4;P27708;H7C2E4;H7BZB3;H7C3Z5 | F8VPD4;P27708 |
| Calcineurin B homologous protein 1 | CHP1 | | 0.276585 | 0.438551 | H0YKE7;F5GX29;H0YNG9;Q99653;H0YLY7 | H0YKE7;F5GX29;H0YNG9;Q99653;H0YLY7 |
| Calcium-binding protein 39 | CAB39 | | 0.29091 | 0.204218 | Q9Y376 | Q9Y376 |
| Calcyclin-binding protein | CACYBP | | 0.259336 | 0.294578 | Q9HB71;Q9HB71-3;B2ZWH1;Q9HB71-2 | Q9HB71;Q9HB71-3 |
| Calmodulin | CALM2;CALM1;CALM3 | | 0.238954 | -0.163872 | H0Y7A7;P62158;E7ETZ0;E7EMB3;G3V361;Q96HY3;F8WBR5;M0QZ52;G | H0Y7A7;P62158;E7ETZ0;E7EMB3;G3V361;Q96HY3 |
| Calnexin | CANX | | 0.672999 | -0.308247 | P27824;B4DGP8;B4E2T8;D6RAQ8;D6RAU8;D6RB85;D6RDP7;H0Y9Q7;D | P27824;B4DGP8;B4E2T8 |
| Calpain small subunit 1 | CAPNS1 | | 0.959248 | -0.741669 | P04632;K7ELJ7;U3KQE2;K7EIV0;K7EKD8;K7EM73;U3KPR7;K7EMQ1;Q9 | P04632;K7ELJ7;U3KQE2;K7EIV0;K7EKD8;K7EM73 |
| Calpain-1 catalytic subunit | CAPN1 | | 1.1952 | -0.810758 | P07384;E9PL37;E9PRM1;E9PLX0;E9PJJ3;E9PLC9;E9PMC6;E9PJA6;E9PS | P07384 |
| Calpain-2 catalytic subunit | CAPN2 | | 1.10397 | -0.543025 | P17655;P17655-2;C9JWY7 | P17655;P17655-2 |
| Calponin-2 | CNN2 | | 0.775199 | 0.886118 | B4DDF4;B4DUT8;Q99439;Q99439-2;H3BVI6;K7ES69;H3BQH0;U3KPS3 | B4DDF4;B4DUT8;Q99439;Q99439-2;H3BVI6;K7ES69 |
| Calreticulin | CALR | | 0.222186 | 0.137155 | P27797;K7EJB9;K7EL50 | P27797;K7EJB9 |
| Calretinin | CALB2 | | 0.318113 | 0.533096 | P22676;H3BN14;A6NER6;H3BMN5;B7Z9J4;P05937 | P22676;H3BN14;A6NER6 |
| Calsyntenin-1;Soluble Alc-alpha;CTF1-alpha | CLSTN1 | | 0.0414775 | -0.0175473 | B4E3Q1;O94985;Q5SR54 | B4E3Q1;O94985;Q5SR54 |
| Calsyntenin-1;Soluble Alc-alpha;CTF1-alpha | CLSTN1 | | 0.00243393 | -0.000497803 | O94985-2 | O94985-2 |
| Calsyntenin-3 | CLSTN3 | + | 1.31927 | -0.909779 | Q9BQT9;Q9BQT9-2;F5H5D7;F5H7C7;F5H5C6;F5H746;F5H172 | Q9BQT9;Q9BQT9-2 |
| Calumenin | CALU | + | 1.77489 | 0.798347 | O43852;O43852-3;O43852-2;O43852-4;O43852-5;H0Y875;O43852-15;( | O43852;O43852-3;O43852-2;O43852-4;O43852-5;H0Y875;O43852-15; |
| cAMP-dependent protein kinase catalytic subunit alpha | PRKACA;KIN27 | | 0.123116 | -0.117066 | P17612;P17612-2;Q15136;K7ERP6;K7ENJ5 | P17612;P17612-2;Q15136 |
| cAMP-dependent protein kinase catalytic subunit beta | PRKACB | | 0.580913 | 1.12591 | P22694-4;P22694-3;P22694;P22694-5;P22694-7;P22694-6;P22694-9;P2 | P22694-4;P22694-3;P22694;P22694-5;P22694-7;P22694-6;P22694-9;P |
| cAMP-dependent protein kinase type II-alpha regulatory subunit | PRKAR2A | | 1.12675 | 0.829289 | Q9BUB1;P13861;H7C1L0;H7C330;P31323;H7C4A9;C9J830 | Q9BUB1;P13861 |
| Carbohydrate sulfotransferase 14 | CHST14 | | 0.269812 | 0.226078 | Q8NCH0;H0YN65 | Q8NCH0;H0YN65 |
| Carbonic anhydrase 2 | CA2 | | 0.613435 | 0.542181 | P00918;E5RID5;E5RK37 | P00918 |
| Carbonyl reductase [NADPH] 1 | CBR1 | | 0.0246332 | 0.029714 | P16152;E9PQ63;A8MTM1;P16152-2 | P16152;E9PQ63;A8MTM1;P16152-2 |
| Carbonyl reductase [NADPH] 3 | CBR3 | | 0.829162 | 0.752569 | O75828 | O75828 |
| Carboxymethylenebutenolidase homolog | CMBL | + | 1.84519 | -1.5519 | Q96DG6 | Q96DG6 |
| Carboxypeptidase A4 | CPA4 | | 0.196511 | 0.119654 | Q9UI42;Q9UI42-2;B7Z5J4;C9J3P4;C9J6N7;F8WF19 | Q9UI42;Q9UI42-2;B7Z5J4 |
| Carboxypeptidase D | CPD | | 1.07316 | -0.806602 | O75976;O75976-2;J3QQJ4 | O75976;O75976-2 |
| Carboxypeptidase E | CPE | | 0.793495 | 0.496237 | P16870-2;P16870;D6R930;D6RF88;C9JE88;H0YAM0 | P16870-2;P16870 |
| Cartilage-associated protein | CRTAP | | 0.770983 | 1.07272 | O75718;C9JP16 | O75718;C9JP16 |
| Casein kinase II subunit beta | CSNK2B;CSNK2B-LY6G5B-1181 | | 0.134918 | 0.12204 | P67870;Q5SRQ6;Q5SRQ3 | P67870;Q5SRQ6;Q5SRQ3 |
| Catalase | CAT | | 0.525775 | -0.325987 | P04040 | P04040 |
| Cathepsin B;Cathepsin B light chain;Cathepsin B heavy chain | CTSB | + | 1.37463 | -1.13832 | P07858;E9PKQ7;E9PNL5;E9PHZ5;E9PLY3;E9PSG5;E9PQM1;E9PR54;E9P | P07858 |
| Cathepsin D;Cathepsin D light chain;Cathepsin D heavy chain | CTSD | | 0.338864 | 0.161583 | P07339;C9JH19;H7C469;F8WD96;H7C1V0;F8W787 | P07339;C9JH19;H7C469;F8WD96 |
| Cathepsin L1;Cathepsin L1 heavy chain;Cathepsin L1 light chain | CTSL | + | 1.30199 | -0.71968 | P07711;Q5T8F0;Q5NE16 | P07711;Q5T8F0 |
| Cathepsin O | CTSO | | 0.340157 | -0.213737 | P43234 | P43234 |
| Cathepsin Z | CTSZ | | 0.356626 | 0.181793 | Q9UBR2 | Q9UBR2 |
| Cation-independent mannose-6-phosphate receptor | IGF2R | | 0.496291 | 0.259943 | P11717;S4R328 | P11717 |
| CCR4-NOT transcription complex subunit 1 | CNOT1 | | 0.0593714 | 0.0473257 | A5YKK6-3;A5YKK6-2;A5YKK6;A5YKK6-4;B5MDN3;H3BMZ2;H3BNB1;H3E | A5YKK6-3;A5YKK6-2;A5YKK6;A5YKK6-4;B5MDN3 |
| CD151 antigen | CD151 | | 0.770376 | -0.816981 | P48509;E9PMR4;E9PRJ3;E9PP93;E9PK37;E9PJE8;E9PSA1;E9PJC8;E9PLZ | P48509;E9PMR4;E9PRJ3 |
| CD166 antigen | ALCAM | | 1.17326 | 0.612592 | Q13740-2;Q13740;F5GXJ9;H7C543;Q13740-4;Q13740-3 | Q13740-2;Q13740;F5GXJ9 |
| CD2-associated protein | CD2AP | | 0.716417 | 1.03155 | Q9Y5K6 | Q9Y5K6 |
| CD44 antigen | CD44 | | 0.0292079 | -0.0224705 | E7EPC6;P16070-18;P16070-12;P16070-14;P16070-13;P16070-11;P1607 | E7EPC6;P16070-18;P16070-12;P16070-14;P16070-13;P16070-11;P1607 |
| CD59 glycoprotein | CD59 | | 0.764047 | -0.440869 | E9PR17;E9PNW4;P13987;H0YET2;Q3C1V1 | E9PR17;E9PNW4;P13987;H0YET2 |
| CD63 antigen | CD63 | | 0.747388 | -0.382264 | F8VNT9;F8VV56;F8W022;F8VWK8;C9JV86;P08962-3;P08962-2;P08962 | F8VNT9;F8VV56;F8W022;F8VWK8;C9JV86;P08962-3;P08962-2;P08962 |
| CD81 antigen | CD81 | | 0.977882 | -1.13993 | E9PJK1;E9PRJ8;H0YDL9;H0YDJ9;E9PIF1;A6NMH8;P60033;E9PM31;H0Y | E9PJK1;E9PRJ8;H0YDL9;H0YDJ9;E9PIF1;A6NMH8;P60033 |
| CD9 antigen | CD9 | | 1.05242 | -1.43472 | A6NNI4;G8JLH6;P21926;F5GXT1 | A6NNI4;G8JLH6;P21926 |
| CD97 antigen;CD97 antigen subunit alpha;CD97 antigen subunit beta | CD97 | | 0.0226922 | 0.0177568 | P48960-2;P48960-3;P48960;B4DQ88;M0QY55;B4DWB8;M0R0K5;Q9UH | P48960-2;P48960-3;P48960 |
| Cdc42-interacting protein 4 | TRIP10 | | 0.306351 | 0.242588 | Q15642-2;Q15642;W4VSQ9;Q15642-4;Q15642-3;M0R2H7;M0R0F9;Q1 | Q15642-2;Q15642;W4VSQ9;Q15642-4;Q15642-3;M0R2H7;M0R0F9;Q1 |
| Cell division control protein 42 homolog | CDC42 | | 0.570282 | -0.409331 | P60953;E7ETU3;P60953-1;Q5JYX0;F8WET9;B1AH79;G3V4H1;G3V476;C | P60953;E7ETU3;P60953-1;Q5JYX0 |
| Cell surface glycoprotein MUC18 | MCAM | | 0.161997 | 0.149896 | P43121;P43121-2 | P43121;P43121-2 |
| Cellular retinoic acid-binding protein 2 | CRABP2 | | 0.1333 | -0.393824 | P29373;Q5SYZ4;B5MCB5;P29762 | P29373;Q5SYZ4 |
| Centrosomal protein of 55 kDa | CEP55 | | 0.741131 | -0.540031 | Q53EZ4;Q53EZ4-2;H0Y432 | Q53EZ4;Q53EZ4-2;H0Y432 |
| Ceroid-lipofuscinosis neuronal protein 5 | CLN5 | | 0.00572831 | 0.00321364 | O75503 | O75503 |
| Charged multivesicular body protein 1a | CHMP1A | | 0.126539 | 0.125679 | Q9HD42;F8VUA2;F8VVT7;F5H875;Q9HD42-2 | Q9HD42;F8VUA2;F8VVT7;F5H875 |
| Charged multivesicular body protein 1b | CHMP1B | | 0.364732 | -0.262077 | Q7LBR1 | Q7LBR1 |
| Charged multivesicular body protein 2a | CHMP2A | | 0.605233 | -0.541233 | O43633;M0R1T5;M0R1L7;M0QXX8;G3V4M1;G3V3F6;G3V599;Q96PC5; | O43633;M0R1T5;M0R1L7 |
| Charged multivesicular body protein 2b | CHMP2B | + | 1.52769 | 1.66322 | C9J0A7;Q9UQN3;Q9UQN3-2 | C9J0A7;Q9UQN3;Q9UQN3-2 |
| Charged multivesicular body protein 3 | CHMP3 | | 0.192651 | -0.174189 | Q9Y3E7;Q9Y3E7-4;Q9Y3E7-3;Q9Y3E7-2 | Q9Y3E7;Q9Y3E7-4;Q9Y3E7-3;Q9Y3E7-2 |
| Charged multivesicular body protein 4b | CHMP4B | + | 2.03633 | 1.89011 | Q9H444;Q96CF2 | Q9H444 |
| Charged multivesicular body protein 5 | CHMP5 | | 1.22923 | 1.16377 | Q9NZZ3;Q9NZZ3-2 | Q9NZZ3;Q9NZZ3-2 |
| Chitinase domain-containing protein 1 | CHID1 | | 0.42122 | 0.185055 | J3KNL3;Q9BWS9-3;Q9BWS9;Q9BWS9-2;E9PPA0;E9PI70;E9PIP0;E9PRL3 | J3KNL3;Q9BWS9-3;Q9BWS9;Q9BWS9-2;E9PPA0 |
| Chloride intracellular channel protein 1 | CLIC1 | | 0.151004 | 0.20815 | O00299 | O00299 |
| Chloride intracellular channel protein 4 | CLIC4 | | 0.0018862 | 0.00180126 | Q9Y696;A6PVS0;O15247 | Q9Y696 |
| Chondroitin sulfate proteoglycan 4 | CSPG4 | + | 1.9414 | -2.40629 | Q6UVK1 | Q6UVK1 |
| Chordin-like protein 1 | CHRDL1 | | 0.425135 | -0.71282 | Q9BU40-2;Q9BU40-4;Q9BU40-5;Q9BU40;Q9BU40-6;Q9BU40-3;D3YTA | Q9BU40-2;Q9BU40-4;Q9BU40-5;Q9BU40;Q9BU40-6;Q9BU40-3;D3YTA |
| Choriogonadotropin subunit beta variant 1;Choriogonadotropin subunit beta;Choriogonadotropin subuni | CGB2;CGB1;LHB;CGB | | 1.22732 | 2.45295 | M0R0E6;K7ELM3;J3QT02;J3KP00;J3KNZ1;H9KV56;A6NKQ9-2;P01233;P | M0R0E6;K7ELM3;J3QT02;J3KP00;J3KNZ1;H9KV56;A6NKQ9-2;P01233;F |
| Cilia- and flagella-associated protein 20 | CFAP20 | | 0.841717 | -0.751215 | Q9Y6A4;H3BPA3 | Q9Y6A4 |
| Citrate synthase;Citrate synthase, mitochondrial | CS | | 0.0276483 | -0.0231022 | B4DJV2;O75390;F8VPA1;F8W4S1;H0YH82;F8VPF9;F8W1S4;F8VRP1;F8V | B4DJV2;O75390 |
| C-Jun-amino-terminal kinase-interacting protein 4 | SPAG9 | | 0.125278 | -0.115377 | O60271-4;O60271;O60271-2;O60271-5;O60271-3;O60271-9;D6RHI8;H( | O60271-4;O60271;O60271-2;O60271-5;O60271-3;O60271-9 |
| Clathrin heavy chain 1 | CLTC | | 0.370819 | -0.212557 | Q00610-2;Q00610;J3KS13;K7EJJ5;J3KSQ2;J3KRF5;J3QL20;F5H5N6 | Q00610-2;Q00610 |
| Clusterin;Clusterin beta chain;Clusterin alpha chain;Clusterin | CLU | + | 2.08686 | -1.74706 | P10909-4;P10909;P10909-5;P10909-2;H0YLK8;H0YC35;E7ERK6;P10909 | P10909-4;P10909;P10909-5;P10909-2;H0YLK8;H0YC35;E7ERK6;P10909 |
| CMP-N-acetylneuraminate-beta-galactosamide-alpha-2,3-sialyltransferase 1 | ST3GAL1 | | 1.12542 | -0.885491 | Q11201;E5RHV6;E5RGL4 | Q11201;E5RHV6 |

| | | | | | | |
|---|---|---|---|---|---|---|
| CMP-N-acetylneuraminate-beta-galactosamide-alpha-2,3-sialyltransferase 2 | ST3GAL2 | | 0.866687 | -0.595456 | Q16842 | Q16842 |
| Coactosin-like protein | COTL1 | | 0.0908699 | -0.0482561 | Q14019;H3BT58 | Q14019;H3BT58 |
| Coatomer subunit alpha;Xenin;Proxenin | COPA | | 0.310833 | 0.375239 | P53621;P53621-2 | P53621;P53621-2 |
| Cofilin-1 | CFL1 | | 0.141198 | -0.125447 | P23528;E9PP50;E9PK25;G3V1A4;E9PQB7;E9PS23;E9PLJ3 | P23528;E9PP50;E9PK25;G3V1A4;E9PQB7;E9PS23;E9PLJ3 |
| Cofilin-2 | CFL2 | | 0.272686 | -0.180915 | Q9Y281;Q9Y281-3;F8WDN3;G3V2U0 | Q9Y281;Q9Y281-3 |
| Coiled-coil domain-containing protein 25 | CCDC25 | | 0.0994088 | 0.14369 | Q86WR0;Q0P663;G3V121;B7Z2L8;Q0VGD4 | Q86WR0;Q0P663 |
| Coiled-coil domain-containing protein 80 | CCDC80 | | 0.640132 | -0.604349 | Q76M96;Q76M96-2;H7C5K4;C9J8I6 | Q76M96;Q76M96-2 |
| Collagen alpha-1(IV) chain;Arresten | COL4A1 | | 1.06471 | -1.37002 | P02462-2;P02462;P29400;P29400-2 | P02462-2;P02462 |
| Collagen alpha-1(VI) chain | COL6A1 | + | 1.98133 | -4.57299 | P12109 | P12109 |
| Collagen alpha-1(XII) chain | COL12A1 | | 0.126661 | -0.120162 | D6RGG3;Q99715;Q99715-4;Q99715-2;H0Y4P7;H0Y5N9;H0Y991 | D6RGG3;Q99715;Q99715-4;Q99715-2 |
| Collagen alpha-1(XIII) chain | COL13A1 | | 0.126644 | 0.0690335 | E7EX21;E7ES50;E7ES56;E9PEG9;E7ES51;E7ES49;E7ES47;E7ES46;G5E98 | E7EX21;E7ES50;E7ES56;E9PEG9;E7ES51;E7ES49;E7ES47;E7ES46;G5E98 |
| Collagen alpha-1(XVIII) chain;Endostatin | COL18A1 | | 0.629171 | -1.14004 | P39060-2;P39060-1;P39060;H7BXV5;H7C457 | P39060-2;P39060-1;P39060;H7BXV5;H7C457 |
| Collagen alpha-2(IV) chain;Canstatin | COL4A2 | | 1.27946 | -1.25067 | P08572;A2A352 | P08572 |
| Collagen and calcium-binding EGF domain-containing protein 1 | CCBE1 | | 0.239669 | 0.259333 | Q6UXH8;K7EQ81;Q6UXH8-3 | Q6UXH8 |
| Complement C3;Complement C3 beta chain;C3-beta-c;Complement C3 alpha chain;C3a anaphylatoxin;Ac | C3 | | 0.478766 | -2.0884 | P01024;M0QXZ3;M0QYC8;M0R0Q9 | P01024 |
| Complement C4-A;Complement C4 beta chain;Complement C4-A alpha chain;C4a anaphylatoxin;C4b-A;C4 | C4A;C4B | | 0.0239277 | 0.0446728 | P0C0L4-2;P0C0L4;F5GXS0;P0C0L5 | P0C0L4-2;P0C0L4;F5GXS0;P0C0L5 |
| Complement component 1 Q subcomponent-binding protein, mitochondrial | C1QBP | | 0.278282 | 0.184591 | Q07021;I3L3Q7;I3L3B0 | Q07021;I3L3Q7;I3L3B0 |
| Complement decay-accelerating factor | CD55 | | 0.280485 | 0.309964 | H3BLV0;E9PSH2;B1AP13;P08174-4;P08174;P08174-3;P08174-5;P08174 | H3BLV0;E9PSH2;B1AP13;P08174-4;P08174;P08174-3;P08174-5;P0817 |
| Complement factor B;Complement factor B Ba fragment;Complement factor B Bb fragment | CFB | | 1.24514 | -2.17051 | B4E1Z4;P00751;E7ETN3;H7C5H1;P00751-2;C9JRT3 | B4E1Z4;P00751;E7ETN3;H7C5H1;P00751-2 |
| Complement factor H | CFH | + | 1.49948 | 1.50241 | P08603;F8WDX4;P08603-2;Q5TFM2;B1AKG0;Q03591;Q6NSD3;Q02985 | P08603 |
| Connective tissue growth factor | CTGF | + | 1.36401 | 1.07454 | P29279 | P29279 |
| COP9 signalosome complex subunit 4 | COPS4 | | 0.461665 | -0.305905 | Q9BT78;D6RFN0;D6RAX7;Q9BT78-2;D6RD63 | Q9BT78;D6RFN0;D6RAX7;Q9BT78-2;D6RD63 |
| COP9 signalosome complex subunit 6 | COPS6 | | 0.357312 | -0.250066 | E7EM64;Q7L5N1;H7C3T0 | E7EM64;Q7L5N1 |
| COP9 signalosome complex subunit 8 | COPS8 | | 0.321791 | -0.21727 | E9PGT6;Q99627-2;Q99627;B8ZZP3;H7C3S9 | E9PGT6;Q99627-2;Q99627;B8ZZP3;H7C3S9 |
| Copine-1 | CPNE1 | | 0.330874 | -0.209295 | B0QZ18;Q99829;F2Z2V0;A6PVH9;E7ENH5;Q5JX56;Q5JX58;Q5JX59;Q5J | B0QZ18;Q99829;F2Z2V0;A6PVH9;E7ENH5;Q5JX56;Q5JX58;Q5JX59;Q5J |
| Core histone macro-H2A.1;Histone H2A | H2AFY | | 1.20638 | -2.04989 | O75367-2;O75367-3;O75367;B4DJC3;D6RCF2;Q5SQT3;Q9P0M6 | O75367-2;O75367-3;O75367;B4DJC3 |
| Coronin-1B | CORO1B | | 0.459739 | 0.244593 | Q9BR76;F5H390;E7EW44;F5H0D2;A9Z1Z3-3 | Q9BR76 |
| Creatine kinase B-type | CKB | | 0.907808 | 1.99341 | P12277;H0YJG0;G3V4N7;G3V461;H0YJK0;G3V2I1 | P12277;H0YJG0 |
| Crk-like protein | CRKL | | 0.274185 | 0.237528 | P46109 | P46109 |
| CUB domain-containing protein 1 | CDCP1 | | 0.809972 | -0.714483 | Q9H5V8;Q9H5V8-2 | Q9H5V8;Q9H5V8-2 |
| Cullin-3 | CUL3 | | 0.324881 | 0.193384 | Q13618-2;Q13618;Q13618-3;H7C1L6;H7C399;Q9Y5S2 | Q13618-2;Q13618;Q13618-3 |
| Cullin-4B | CUL4B | | 0.052369 | 0.0616531 | K4DI93;Q13620-1;Q13620;Q13620-3;A6NE76 | K4DI93;Q13620-1;Q13620;Q13620-3;A6NE76 |
| Cullin-associated NEDD8-dissociated protein 1 | CAND1 | | 0.0902642 | 0.0905078 | Q86VP6;Q86VP6-2;H0YH27;Q86VP6-3;F5H6I6;O75155-2;O75155;F8WB | Q86VP6;Q86VP6-2 |
| C-X-C motif chemokine 3;GRO-gamma(5-73) | CXCL3 | + | 1.51743 | -3.73773 | P19876 | P19876 |
| C-X-C motif chemokine;Interleukin-8;MDNCF-a;Interleukin-8;IL-8(5-77);IL-8(6-77);IL-8(7-77);IL-8(8-77);IL-8 | CXCL8 | + | 2.47573 | -2.05726 | C9J4T6;P10145;P10145-2 | C9J4T6;P10145;P10145-2 |
| Cyclin-dependent kinase 1 | CDK1 | | 0.276866 | 0.404225 | P06493;P06493-2;E5RIU6 | P06493;P06493-2;E5RIU6 |
| Cystathionine gamma-lyase | CTH | | 0.0468488 | -0.0362186 | P32929;P32929-2;P32929-3 | P32929;P32929-2;P32929-3 |
| Cystatin-B | CSTB | | 0.340239 | -0.266664 | P04080 | P04080 |
| Cystatin-C | CST3 | | 0.502916 | -0.367875 | P01034 | P01034 |
| Cysteine and glycine-rich protein 1 | CSRP1 | | 0.0132956 | 0.00736134 | P21291;E9PS42;E9PND2;E9PP21 | P21291;E9PS42;E9PND2;E9PP21 |
| Cysteine-rich motor neuron 1 protein;Processed cysteine-rich motor neuron 1 protein | CRIM1 | | 0.0125318 | 0.0101895 | Q9NZV1;H7C2T6;H7C1Z0;H7C458 | Q9NZV1 |
| Cytochrome c oxidase subunit 2 | MT-CO2 | | 0.886242 | -2.29392 | P00403 | P00403 |
| Cytochrome c oxidase subunit 5A, mitochondrial | COX5A | + | 1.46488 | 2.42781 | P20674;H3BRM5;H3BV69;H3BNX8 | P20674;H3BRM5;H3BV69;H3BNX8 |
| Cytoplasmic aconitate hydratase | ACO1;IRP1 | | 0.310716 | -0.336854 | P21399;Q9HBB2;Q5VZA6;F5H2R8;F5H143;F5H3Z7;E9PEF1;Q6ZMR1;O0 | P21399;Q9HBB2 |
| Cytoplasmic dynein 1 heavy chain 1 | DYNC1H1 | | 0.0317545 | -0.0502497 | Q14204;H0YJ21 | Q14204 |
| Cytoplasmic FMR1-interacting protein 1 | CYFIP1 | | 0.316304 | -0.192165 | Q7L576;Q7L576-2;Q7L576-3;H0YL50;H0YLJ5;H0YL39;H0YN93 | Q7L576;Q7L576-2 |
| Cytosol aminopeptidase | LAP3 | | 0.142453 | 0.174996 | P28838-2;P28838;H0Y9Q1;H0Y983 | P28838-2;P28838 |
| Cytosolic acyl coenzyme A thioester hydrolase | ACOT7 | | 0.325325 | -0.376639 | O00154-4;O00154-6;O00154-7;O00154;O00154-5;B4DUX0;K7EKP8;O0 | O00154-4;O00154-6;O00154-7;O00154;O00154-5;B4DUX0;K7EKP8;O0 |
| Cytosolic Fe-S cluster assembly factor NUBP1 | NUBP1 | | 0.0736734 | 0.0971867 | P53384-2;P53384;I3L531;I3L3A0;I3L518 | P53384-2;P53384 |
| Cytosolic non-specific dipeptidase | CNDP2 | | 0.0538488 | -0.0484755 | Q96KP4;Q96KP4-2;J3QKT2;J3KSV5;J3QR27;J3QLU1;J3QKQ0;J3QRD0;J3 | Q96KP4;Q96KP4-2;J3QKT2 |
| D-3-phosphoglycerate dehydrogenase | PHGDH | | 0.111607 | -0.140624 | Q5SZU1;O43175 | Q5SZU1;O43175 |
| DCC-interacting protein 13-alpha | APPL1 | | 0.196658 | -0.390507 | Q9UKG1;C9JAB0;C9K0C4 | Q9UKG1;C9JAB0 |
| D-dopachrome decarboxylase;D-dopachrome decarboxylase-like protein | DDT;DDTL | | 0.10183 | -0.0757646 | P30046;B7Z522;B5MC82;A6NHG4;J3KQ18;H7C342 | P30046;B7Z522;B5MC82;A6NHG4;J3KQ18 |
| Dehydrogenase/reductase SDR family member 11 | DHRS11 | | 0.0214145 | 0.0348644 | Q6UWP2;K7END9;K7EK68;Q6UWP2-2;K7EJP4 | Q6UWP2;K7END9;K7EK68;Q6UWP2-2 |
| Deleted in autism protein 1 | C3orf58 | | 1.07203 | -0.667942 | Q8NDZ4;E7ET39;C9JY09;Q8NDZ4-2 | Q8NDZ4 |
| Delta(3,5)-Delta(2,4)-dienoyl-CoA isomerase, mitochondrial | ECH1 | | 0.0252035 | 0.0198645 | Q13011;M0R248;M0QZW4 | Q13011;M0R248 |
| Delta-aminolevulinic acid dehydratase | ALAD | | 0.887027 | -0.458361 | P13716;B7Z3I9;P13716-2;B7ZBK6 | P13716;B7Z3I9;P13716-2 |
| Deoxycytidylate deaminase | DCTD | | 0.0230801 | 0.0209997 | P32321;P32321-2;D6RBJ9;D6R9S0;D6RBN2;D6RAD7;D6RAR9;D6RC36;D | P32321;P32321-2;D6RBJ9 |
| Deoxyribonuclease-2-alpha | DNASE2 | | 0.169755 | -0.164451 | B7Z4K6;O00115;K7ENE5;REV__H3BPH3;REV__F8VR84;REV__F8VQQ3;I | B7Z4K6;O00115;K7ENE5 |
| Deoxyuridine 5-triphosphate nucleotidohydrolase, mitochondrial | DUT | | 0.204755 | 0.241957 | P33316-2;H0YNW5;H0YNJ9;H0YKC5;H0YMM5;H0YMP1 | P33316-2;H0YNW5;H0YNJ9;H0YKC5;H0YMM5 |
| Desmocollin-2 | DSC2 | | 0.417486 | -0.701581 | Q02487-2;Q02487;J3QRL9;Q14574-2 | Q02487-2;Q02487 |
| Desmoglein-2 | DSG2 | | 0.265217 | -0.167983 | Q14126;J3KSI6 | Q14126 |
| Desmoplakin | DSP | | 0.665672 | -0.945551 | P15924;P15924-3;P15924-2 | P15924;P15924-3;P15924-2 |
| Destrin | DSTN | | 0.227086 | 0.165362 | P60981;P60981-2;F6RFD5 | P60981;P60981-2;F6RFD5 |
| Diamine acetyltransferase 2 | SAT2 | | 0.0302213 | -0.06473 | I3L0W4;Q96F10;I3L2W7 | I3L0W4;Q96F10 |
| Dickkopf-related protein 1 | DKK1 | + | 1.48567 | 0.870732 | O94907;D6RGF1;D6RCC2;Q9UBU2 | O94907 |
| Dihydrolipoyllysine-residue succinyltransferase component of 2-oxoglutarate dehydrogenase complex, m | DLST | + | 1.56078 | 1.3921 | P36957;B7Z5W8;Q86SW4;H0YJF9;G3V5M3;G3V3F0 | P36957;B7Z5W8;Q86SW4 |
| Dihydropteridine reductase | QDPR | | 0.11015 | -0.128208 | P09417;P09417-2;B7Z415;D6RGG7;H0Y8F7;D6RHJ7 | P09417;P09417-2;B7Z415;D6RGG7 |
| Dihydropyrimidinase-related protein 2 | DPYSL2 | | 0.0707395 | 0.0633947 | Q16555-2;Q16555;E5RFU4;H0YBT4;E9PD68;Q14195;Q14194;Q14195-2 | Q16555-2;Q16555 |
| Di-N-acetylchitobiase | CTBS | | 0.0982058 | 0.115552 | Q01459 | Q01459 |
| Dipeptidyl peptidase 1;Dipeptidyl peptidase 1 exclusion domain chain;Dipeptidyl peptidase 1 heavy chain | CTSC | | 0.109711 | -0.0482666 | P53634;H0YCY8;P53634-2;P53634-3 | P53634;H0YCY8 |
| Dipeptidyl peptidase 2 | DPP7 | | 0.692608 | 0.140004 | Q9UHL4;R4GMR2;R4GNE8;R4GMU5;R4GMV4 | Q9UHL4 |
| Dipeptidyl peptidase 3 | DPP3 | | 0.011217 | -0.00794012 | Q9NY33;G3V1D3;G3V180;Q9NY33-4;Q9NY33-2;E9PQ14;E9PPK9;E9PKK | Q9NY33;G3V1D3;G3V180;Q9NY33-4 |
| Diphosphoinositol polyphosphate phosphohydrolase 1 | NUDT3 | | 0.224245 | -0.162051 | O95989 | O95989 |
| Disco-interacting protein 2 homolog B | DIP2B | | 0.0771905 | -0.0203612 | Q9P265;H0Y7Q3;Q9Y2E4 | Q9P265 |
| Disintegrin and metalloproteinase domain-containing protein 10 | ADAM10 | | 0.406462 | -0.540299 | O14672;B4DU28;H0YNC5;C9J9B4;H3BS53;H0YK87 | O14672;B4DU28 |
| Disintegrin and metalloproteinase domain-containing protein 15 | ADAM15 | + | 2.53277 | 2.72788 | Q13444-8;Q13444-2;Q13444-7;Q13444-6;Q13444-12;Q13444-5;Q1344 | Q13444-8;Q13444-2;Q13444-7;Q13444-6;Q13444-12;Q13444-5;Q1344 |
| Disintegrin and metalloproteinase domain-containing protein 9 | ADAM9 | | 0.586966 | 0.795388 | Q13443;A0AVL1;Q13443-2;F8WC54;C9J6H5;C9JPM3 | Q13443;A0AVL1;Q13443-2;F8WC54 |
| Disks large homolog 1 | DLG1 | | 0.255171 | -0.302698 | Q12959-8;Q12959-9;Q12959-5;Q12959-3;Q12959-6;Q12959-4;Q12959; | Q12959-8;Q12959-9;Q12959-5;Q12959-3;Q12959-6;Q12959-4;Q12959 |

| | | | | | | |
|---|---|---|---|---|---|---|
| DNA damage-binding protein 1 | DDB1 | | 0.442944 | -0.229036 | Q16531;F5GY55;B4DG00;F5H2L3;F5H581;F5H0Y5;F8WF81;F5GZ34;F5H | Q16531;F5GY55 |
| DnaJ homolog subfamily A member 1 | DNAJA1 | | 0.0750971 | 0.098775 | P31689;P31689-2;B7Z5C0 | P31689;P31689-2 |
| DnaJ homolog subfamily A member 2 | DNAJA2 | | 0.0485217 | -0.0428788 | O60884;I3L320;H3BMW5 | O60884 |
| DnaJ homolog subfamily B member 1 | DNAJB1 | | 0.213937 | 0.458026 | P25685;B4DX52;M0R080;M0QYT3;M0QXK0;M0R1D6;M0QZD0;M0R12 | P25685;B4DX52;M0R080 |
| DnaJ homolog subfamily B member 4 | DNAJB4 | | 0.161432 | 0.25033 | Q9UDY4 | Q9UDY4 |
| DnaJ homolog subfamily B member 6 | DNAJB6 | | 0.458444 | 0.630773 | O75190-2;C9J2C4;O75190-3;O75190;C9JDX6;F8WCZ4;C9J2P2;E9PH18; | O75190-2;C9J2C4;O75190-3;O75190;C9JDX6;F8WCZ4;C9J2P2;E9PH18; |
| DnaJ homolog subfamily C member 3 | DNAJC3 | | 0.213284 | -0.157178 | Q13217 | Q13217 |
| DnaJ homolog subfamily C member 7 | DNAJC7 | | 0.00282876 | -0.00699697 | Q99615-2;Q99615;K7EPP7;K7ESP1;K7EIH8;K7EJL5;K7EN19;K7EJV7;K7E | Q99615-2;Q99615;K7EPP7;K7ESP1;K7EIH8 |
| Dolichyl-diphosphooligosaccharide--protein glycosyltransferase subunit DAD1 | DAD1 | | 0.748909 | -0.631445 | F5GXX5;P61803;F5H895 | F5GXX5;P61803;F5H895 |
| Double-stranded RNA-binding protein Staufen homolog 1 | STAU1 | | 0.493296 | -0.594591 | Q5JW30;O95793-2;O95793-3;O95793 | Q5JW30;O95793-2;O95793-3;O95793 |
| Drebrin-like protein | DBNL | + | 1.31368 | 1.56244 | Q9UJU6;H0Y5J4;Q9UJU6-5 | Q9UJU6;H0Y5J4;Q9UJU6-5 |
| Dual specificity mitogen-activated protein kinase kinase 1 | MAP2K1 | | 0.13786 | 0.204436 | Q02750;Q02750-2;H3BRW9;G5E9C7 | Q02750;Q02750-2 |
| Dual specificity protein phosphatase 3 | DUSP3 | | 0.599212 | -0.447886 | P51452;Q8IYJ9;K7ES89;K7ELG5 | P51452;Q8IYJ9;K7ES89 |
| Dynactin subunit 2 | DCTN2 | | 0.78913 | 1.07907 | F5H2S7;Q13561;Q13561-3;Q13561-2;F5H223;F8W1I6;H0YI98;F8VW18; | F5H2S7;Q13561;Q13561-3;Q13561-2;F5H223;F8W1I6;H0YI98;F8VW18 |
| Dynamin-2 | DNM2 | | 0.370633 | 0.50205 | P50570-2;P50570-5;P50570;F5H4R9;P50570-3;P50570-4;K7EMQ3;Q05 | P50570-2;P50570-5;P50570;F5H4R9;P50570-3;P50570-4 |
| Dynein light chain 1, cytoplasmic | DYNLL1 | | 0.0142779 | 0.0142503 | P63167;F8VXL2;F8VRV5;F8VXI7 | P63167;F8VXL2;F8VRV5 |
| Dynein light chain roadblock-type 1;Dynein light chain roadblock-type 2 | DYNLRB1;DYNLRB2 | | 0.423737 | -0.2877 | Q9NP97;B4DFR2;B1AKR6;Q8TF09;H3BNG9;H3BPA0;Q7Z4M1;H3BQI1;C | Q9NP97;B4DFR2;B1AKR6;Q8TF09 |
| Dyslexia-associated protein KIAA0319-like protein | KIAA0319L | | 0.476648 | 0.0692776 | C9JVB2;B4DYG9;B1AN15;E7EN73;Q8IZA0-2;Q8IZA0;C9JQ56;H0Y4S8 | C9JVB2;B4DYG9;B1AN15;E7EN73;Q8IZA0-2;Q8IZA0;C9JQ56 |
| Dystonin | DST | | 0.0407517 | 0.0354708 | Q03001;E7ERU2;E9PEB9;F8W9J4;E7ERU0;E9PHM6;J3QTQ0;Q03001-8; | Q03001;E7ERU2;E9PEB9;F8W9J4;E7ERU0;E9PHM6;J3QTQ0;Q03001-8 |
| Dystonin | DST | | 0.373561 | -1.38193 | Q03001-3;Q6P0N6 | Q03001-3;Q6P0N6 |
| Dystroglycan;Alpha-dystroglycan;Beta-dystroglycan | DAG1 | | 0.158921 | -0.140325 | Q14118;C9JYS1;C9J196;C9JQL4 | Q14118 |
| E3 ubiquitin-protein ligase CHIP | STUB1 | | 0.954072 | 0.50191 | Q9UNE7;Q9UNE7-2;H3BS86;H3BTA3;H3BUD0 | Q9UNE7;Q9UNE7-2;H3BS86;H3BTA3 |
| E3 ubiquitin-protein ligase HUWE1 | HUWE1 | | 0.507498 | -0.650215 | Q7Z6Z7-2;Q7Z6Z7-3;Q7Z6Z7;H0Y5W0;Q5H962;H0Y659;F6WK14;H0Y7U | Q7Z6Z7-2;Q7Z6Z7-3;Q7Z6Z7;H0Y5W0 |
| E3 ubiquitin-protein ligase NEDD4 | NEDD4 | | 0.174252 | -0.133322 | P46934-4;H0Y8X6;H0Y8H4;H0Y9L1;K7EKL1 | P46934-4;H0Y8X6 |
| Early endosome antigen 1 | EEA1 | | 1.1836 | 0.646749 | Q15075 | Q15075 |
| Ectonucleotide pyrophosphatase/phosphodiesterase family member 1;Alkaline phosphodiesterase I;Nucl | ENPP1 | | 0.515934 | 0.382828 | P22413;E9PE72 | P22413 |
| EGF-containing fibulin-like extracellular matrix protein 1 | EFEMP1 | | 0.182719 | 0.0710706 | Q12805-2;Q12805-4;Q12805-3;Q12805;Q12805-5;Q580Q6 | Q12805-2;Q12805-4;Q12805-3;Q12805;Q12805-5 |
| EGF-like repeat and discoidin I-like domain-containing protein 3 | EDIL3 | | 0.0228747 | 0.0386312 | O43854-2;O43854 | O43854-2;O43854 |
| EH domain-containing protein 1 | EHD1 | | 0.0523988 | -0.047261 | Q9H4M9;C9JC03;C9J2Z4;Q9NZN3;C9JDQ8;C9JIJ3;C9IZH1;B4DFR5 | Q9H4M9;C9JC03;C9J2Z4 |
| EH domain-containing protein 4 | EHD4 | | 0.221658 | -0.644506 | Q9H223 | Q9H223 |
| Electron transfer flavoprotein subunit alpha, mitochondrial | ETFA | | 0.0329498 | -0.0311918 | P13804;P13804-2;H0YLU7;H0YK49;H0YL12;H0YNX6;H0YKF0;H0YL83;J3 | P13804;P13804-2;H0YLU7;H0YK49;H0YL12;H0YNX6;H0YKF0 |
| Electron transfer flavoprotein subunit beta | ETFB | | 0.027448 | -0.0334311 | P38117;P38117-2;M0QY67 | P38117;P38117-2 |
| Elongation factor 1-alpha 1;Putative elongation factor 1-alpha-like 3 | EEF1A1;EEF1A1P5 | | 0.0434574 | 0.044349 | P68104;Q5VTE0;Q5JR01;A6PW80 | P68104;Q5VTE0 |
| Elongation factor 1-gamma | EEF1G | | 0.0664572 | 0.021724 | P26641;B4DTG2 | P26641;B4DTG2 |
| Elongation factor 2 | EEF2 | | 0.480045 | 0.42751 | P13639 | P13639 |
| Elongation factor Tu, mitochondrial | TUFM | + | 1.36788 | 0.639702 | P49411;H3BNU3 | P49411 |
| Endophilin-A2 | SH3GL1 | | 0.816678 | 1.15991 | Q99961;Q99961-3;Q99961-2;M0QYE0;M0R0I3;M0R2K6;F5H5J3;Q9996 | Q99961;Q99961-3;Q99961-2;M0QYE0;M0R0I3 |
| Endophilin-B1 | SH3GLB1 | + | 1.39914 | 1.36738 | Q9Y371;Q9Y371-2;Q9Y371-3 | Q9Y371;Q9Y371-2 |
| Endoplasmic reticulum aminopeptidase 1 | ERAP1 | | 0.309377 | 0.195853 | Q9NZ08;Q9NZ08-2;H0Y9X5;D6RAL9 | Q9NZ08;Q9NZ08-2 |
| Endoplasmic reticulum resident protein 29 | ERP29 | | 0.0477918 | 0.0557033 | P30040;F8VY02;F8W1G0;P30040-2 | P30040;F8VY02 |
| Endoplasmic reticulum resident protein 44 | ERP44 | | 0.985308 | 0.774307 | Q9BS26 | Q9BS26 |
| Endoplasmin | HSP90B1 | | 0.257826 | 0.126563 | P14625;H0YIV0;Q58FF3;F8W026;F8VPC7 | P14625 |
| Endothelial cell-selective adhesion molecule | ESAM | | 0.863305 | -0.549783 | F8WDW9;Q96AP7;C9JIE7 | F8WDW9;Q96AP7 |
| Endothelial differentiation-related factor 1 | EDF1 | | 0.0537658 | 0.0683488 | O60869-2;O60869-3;O60869 | O60869-2;O60869-3;O60869 |
| Endothelial protein C receptor | PROCR | | 0.433908 | -0.154291 | Q9UNN8 | Q9UNN8 |
| Endothelin-1;Endothelin-1;Big endothelin-1 | EDN1 | | 0.666902 | -0.927339 | P05305 | P05305 |
| Endothelin-converting enzyme 1 | ECE1 | | 0.043308 | -0.0265103 | P42892-2;P42892-3;P42892-4;P42892;B4DKB2;E9PJG1 | P42892-2;P42892-3;P42892-4;P42892;B4DKB2 |
| Enolase-phosphatase E1 | ENOPH1 | | 1.06075 | -0.640592 | Q9UHY7;D6RA00;Q9UHY7-2 | Q9UHY7;D6RA00;Q9UHY7-2 |
| Enoyl-CoA delta isomerase 1, mitochondrial | ECI1;DCI | | 0.0555234 | 0.0500955 | P42126;Q96DC0;P42126-2;H3BS70 | P42126;Q96DC0;P42126-2;H3BS70 |
| Enoyl-CoA hydratase, mitochondrial | ECHS1 | | 0.984905 | 0.801191 | P30084 | P30084 |
| Envoplakin | EVPL | | 0.196555 | 0.244169 | K7EKI0;Q92817 | K7EKI0;Q92817 |
| Ephrin type-B receptor 2 | EPHB2 | | 0.226363 | 0.130293 | B1AKC9;P29323-2;P29323-3;P29323;Q6NVW1 | B1AKC9;P29323-2;P29323-3;P29323 |
| Ephrin type-B receptor 4 | EPHB4 | | 0.0301491 | 0.0423526 | Q96L35;P54760 | Q96L35;P54760 |
| Ephrin-A1;Ephrin-A1, secreted form | EFNA1 | | 1.09271 | -2.41015 | P20827;P20827-2 | P20827;P20827-2 |
| Ephrin-A5 | EFNA5 | | 0.156379 | -0.196528 | P52803;D6RDV5 | P52803;D6RDV5 |
| Ephrin-B1 | EFNB1 | | 1.27509 | -2.40255 | P98172 | P98172 |
| Epidermal growth factor receptor | EGFR | | 0.584224 | -0.482362 | P00533;E9PFD7;Q504U8;P00533-4;P00533-3;P00533-2;C9JYS6;B4DTR1 | P00533;E9PFD7;Q504U8;P00533-4;P00533-3 |
| Epidermal growth factor receptor kinase substrate 8-like protein 2 | EPS8L2 | | 0.19715 | 0.232221 | Q9H6S3;Q9H6S3-3;Q9H6S3-2;H0YD62;H0YF37;E9PNT0;E9PLH1 | Q9H6S3;Q9H6S3-3 |
| Epididymal secretory protein E1 | NPC2 | | 0.291491 | -0.254867 | P61916;J3KMY5;G3V3E8;E7EMS2;G3V3D1;H0YIZ1;G3V2V8;B4DQV7 | P61916;J3KMY5;G3V3E8;E7EMS2;G3V3D1;H0YIZ1;G3V2V8;B4DQV7 |
| Epididymis-specific alpha-mannosidase | MAN2B2 | | 0.367096 | -0.23769 | Q9Y2E5;E9PCD7;H0YA68;Q9Y2E5-2 | Q9Y2E5;E9PCD7;H0YA68;Q9Y2E5-2 |
| Epsin-3;Epsin-2 | EPN2;EPN3 | | 0.416737 | -0.248921 | I3L2B2;J3QLN2;D6RBR6;D6RFG3;D6REL6;B4DIQ9;I6L9I8;I3L2H1;F6PQP | I3L2B2;J3QLN2;D6RBR6;D6RFG3;D6REL6;B4DIQ9;I6L9I8;I3L2H1;F6PQP |
| Erlin-1;Erlin-2 | ERLIN1;ERLIN2 | | 0.509713 | 0.25514 | O75477;B0QZ43;E5RHW4;O94905;E5RJ09;O94905-2;O94905-3 | O75477;B0QZ43;E5RHW4;O94905 |
| Erythrocyte band 7 integral membrane protein | STOM | | 0.430788 | 0.542298 | B4E2V5;P27105;P27105-2;Q8TAV4-2;Q8TAV4 | B4E2V5;P27105 |
| Ester hydrolase C11orf54 | C11orf54 | | 0.336573 | -0.246451 | Q9H0W9-2;Q9H0W9;Q9H0W9-3;E9PQS1;E9PJU8;Q9H0W9-4;E9PPB5;E | Q9H0W9-2;Q9H0W9;Q9H0W9-3;E9PQS1;E9PJU8;Q9H0W9-4;E9PPB5 |
| Ethylmalonyl-CoA decarboxylase | ECHDC1 | | 0.406311 | 0.469812 | Q9NTX5-3;Q9NTX5-6;Q9NTX5-2;Q9NTX5;H0Y5L2;H0Y525;H0YCS9;J3KP | Q9NTX5-3;Q9NTX5-6;Q9NTX5-2;Q9NTX5;H0Y5L2 |
| Eukaryotic initiation factor 4A-I | EIF4A1 | | 0.195011 | 0.227926 | P60842;J3KT12;P60842-2;J3QS69;J3KTB5;J3KSZ0;J3QL43;J3KTN0;J3QR | P60842;J3KT12;P60842-2;J3QS69;J3KTB5;J3KSZ0;J3QL43 |
| Eukaryotic translation elongation factor 1 epsilon-1 | EEF1E1;EEF1E1-BLOC1S5 | | 0.0507017 | -0.088796 | O43324;D6RBD7;C9J1V9;O43324-2;H0YAL7;D6RCQ0 | O43324;D6RBD7;C9J1V9;O43324-2;H0YAL7 |
| Eukaryotic translation initiation factor 2 subunit 1 | EIF2S1 | | 0.398602 | 0.459469 | P05198;G3V4T5 | P05198;G3V4T5 |
| Eukaryotic translation initiation factor 2 subunit 3;Putative eukaryotic translation initiation factor 2 subun | EIF2S3;EIF2S3L | | 0.122105 | -0.130047 | P41091;Q2VIR3;F8W810;Q2VIR3-2;H7BZU1 | P41091;Q2VIR3;F8W810;Q2VIR3-2 |
| Eukaryotic translation initiation factor 2A;Eukaryotic translation initiation factor 2A, N-terminally processe | EIF2A | | 0.242673 | -0.713259 | B4DQ14;Q9BY44;F8WAE5;B4DF96;C9IZE1;Q9BY44-2;H7C5Q3;H7C5R5; | B4DQ14;Q9BY44;F8WAE5;B4DF96;C9IZE1 |
| Eukaryotic translation initiation factor 3 subunit B | EIF3B | | 0.34309 | 0.428809 | P55884;P55884-2;C9JZG1;C9JQN7 | P55884;P55884-2 |
| Eukaryotic translation initiation factor 3 subunit E | EIF3E | | 0.280365 | 0.44313 | P60228;E5RGA2;H0YBR5;E5RHS5;H0YAW4;E5RIT4;E5RII3;E5RIP5;E5RJ | P60228;E5RGA2 |
| Eukaryotic translation initiation factor 3 subunit H | EIF3S3;EIF3H | | 0.108473 | 0.0688304 | B3KS98;O15372;E5RJT0;E5RGU4;E5RFW7;E5RFH0;E5RHC7;E5RH59 | B3KS98;O15372;E5RJT0 |
| Eukaryotic translation initiation factor 3 subunit I | EIF3I | | 0.947406 | 1.26623 | Q13347;Q5TFK1 | Q13347 |
| Eukaryotic translation initiation factor 3 subunit K | EIF3K | | 0.391815 | 0.577598 | Q9UBQ5;B7ZAM9;K7ERF1;K7ES31;K7EMQ9;K7EQM4;K7EK53;B4DQ48 | Q9UBQ5;B7ZAM9;K7ERF1;K7ES31;K7EMQ9;K7EQM4 |
| Eukaryotic translation initiation factor 4E | EIF4E | | 0.213191 | 0.247948 | P06730;P06730-2;D6RBW1;P06730-3;H0Y8J7;D6RFJ3;D6RHE2;H0Y8X3; | P06730;P06730-2;D6RBW1;P06730-3;H0Y8J7 |
| Eukaryotic translation initiation factor 5A-1;Eukaryotic translation initiation factor 5A-1-like | EIF5A;EIF5AL1 | | 0.0963112 | 0.106826 | P63241;I3L397;I3L504;P63241-2;Q6IS14;F8WCJ1;C9J7B5;C9J4W5;Q9GZ | P63241;I3L397;I3L504;P63241-2;Q6IS14 |
| Eukaryotic translation initiation factor 6 | EIF6 | | 0.331953 | 0.388604 | P56537;P56537-2;B7ZBH1;F8WD20;F8WDS6 | P56537 |
| Exosome complex component RRP45 | EXOSC9 | | 0.239492 | 0.350094 | D6RIY6;Q06265;Q06265-2;D6R905;D6RA17;D6RAP4;Q06265-3;Q06265 | D6RIY6;Q06265;Q06265-2;D6R905;D6RA17;D6RAP4;Q06265-3;Q0626 |

| | | | | | | |
|---|---|---|---|---|---|---|
| Exostosin-2 | EXT2 | | 0.377481 | -0.504381 | Q93063;Q93063-3;Q93063-2;E9PNL9 | Q93063;Q93063-3;Q93063-2 |
| Exostosin-like 2;Processed exostosin-like 2 | EXTL2 | | 1.04831 | 0.524368 | F5GZK1;Q9UBQ6;C9JEG3;C9IYF5 | F5GZK1;Q9UBQ6;C9JEG3 |
| Exportin-2 | CSE1L | | 0.151156 | -0.188249 | P55060-3;P55060;P55060-4;B4DUC5;P55060-2 | P55060-3;P55060;P55060-4;B4DUC5 |
| Extracellular matrix protein 1 | ECM1 | + | 2.95399 | 0.972894 | Q16610;Q16610-4;Q16610-2;Q16610-3 | Q16610;Q16610-4;Q16610-2 |
| Extracellular serine/threonine protein kinase FAM20C | FAM20C | | 0.311874 | -0.548284 | Q8IXL6;K7EQL5;Q8IXL6-2;Q96MK3 | Q8IXL6 |
| Extracellular superoxide dismutase [Cu-Zn] | SOD3 | | 0.791419 | -4.17782 | P08294;M0R1V4 | P08294 |
| Ezrin | EZR | | 0.265401 | 0.191181 | P15311;E7EQR4 | P15311;E7EQR4 |
| F-actin-capping protein subunit alpha-1 | CAPZA1 | | 0.325125 | 0.317952 | P52907 | P52907 |
| F-actin-capping protein subunit alpha-2 | CAPZA2 | | 0.41476 | 0.426597 | P47755;F8W9N7;C9JUG7;P47755-2;F8WED3 | P47755;F8W9N7 |
| F-actin-capping protein subunit beta | CAPZB | | 0.0266596 | 0.0240938 | P47756-2;B1AK87;B1AK88;P47756;B1AK85;F6USW4;F6Q0E3 | P47756-2;B1AK87;B1AK88;P47756;B1AK85;F6USW4 |
| Fascin | FSCN1 | | 0.366007 | 0.294696 | Q16658;J3KNT0;C9JFC0;C9JPH9 | Q16658;J3KNT0 |
| Fatty acid synthase;[Acyl-carrier-protein] S-acetyltransferase;[Acyl-carrier-protein] S-malonyltransferase;3 | FASN | | 0.474708 | -0.373161 | P49327;J3KTF0 | P49327 |
| Ferritin heavy chain;Ferritin heavy chain, N-terminally processed;Ferritin | FTH1 | | 0.0862485 | 0.047799 | P02794;G3V192;G3V1D1;E9PRK8;E9PKY7;E9PPQ4;E9PQR3;E9PKM5;E9 | P02794;G3V192;G3V1D1 |
| Ferritin light chain | FTL | | 0.595431 | 0.463356 | P02792 | P02792 |
| Fibrillin-1 | FBN1 | | 0.437064 | 0.370443 | P35555;H0YND0;F6U495;E9PHW4 | P35555 |
| Fibroblast growth factor receptor;Fibroblast growth factor receptor 1 | FGFR1 | | 0.115574 | 0.0766302 | J3KNT4;P11362-20;P11362-7;P11362-2;P11362-19;P11362;P11362-21;I | J3KNT4;P11362-20;P11362-7;P11362-2;P11362-19;P11362;P11362-21; |
| Fibronectin;Anastellin;Ugl-Y1;Ugl-Y2;Ugl-Y3 | FN1 | | 0.358138 | -0.168685 | P02751-17;P02751-3;P02751;P02751-5;P02751-7;P02751-15;F8W7G7;I | P02751-17;P02751-3;P02751;P02751-5;P02751-7;P02751-15;F8W7G7; |
| Filamin-A | FLNA | | 0.757237 | 0.692208 | P21333-2;P21333;Q5HY54;E9PHF0;F8WE98;H0Y5C6;H0Y5F3;H7C2E7 | P21333-2;P21333;Q5HY54 |
| Filamin-B | FLNB | | 0.845605 | -0.562141 | O75369-2;O75369-9;O75369;O75369-6;O75369-3;E7EN95;O75369-7;O7 | O75369-2;O75369-9;O75369;O75369-6;O75369-3;E7EN95;O75369-7;O |
| Filamin-B | FLNB | | 0.983003 | -0.628443 | O75369-8;H7C5L4;Q8WXT3 | O75369-8 |
| Flavin reductase (NADPH) | BLVRB | | 0.434562 | -0.373756 | P30043;M0R192;M0QZL1 | P30043;M0R192 |
| Flotillin-1 | FLOT1 | | 0.432438 | -0.510374 | B4DVY7;O75955;A2AB09;A2ABJ5;A2AB12;A2AB10 | B4DVY7;O75955;A2AB09 |
| Follistatin | FST | | 0.00983582 | 0.00922087 | P19883-2;P19883;H0YA75;H0YAF9 | P19883-2;P19883;H0YA75 |
| Follistatin-related protein 1 | FSTL1 | | 0.868824 | -1.45021 | Q12841;B4DTT5;C9J5G4;H7C4W4 | Q12841;B4DTT5 |
| Follistatin-related protein 3 | FSTL3 | + | 1.71995 | 0.736654 | O95633-2;O95633;K7ES54 | O95633-2;O95633;K7ES54 |
| Four-jointed box protein 1 | FJX1 | | 0.526154 | -0.226029 | Q86VR8 | Q86VR8 |
| Fragile X mental retardation syndrome-related protein 2 | FXR2 | | 0.037594 | 0.0501756 | P51116;I3L1Z2;F8W871;A8MQB8;G3V0J0;Q06787-8;Q06787-6;Q06787 | P51116 |
| FRAS1-related extracellular matrix protein 2 | FREM2 | + | 1.4292 | -0.819044 | Q5SZK8-2;Q5SZK8 | Q5SZK8-2;Q5SZK8 |
| Fructose-1,6-bisphosphatase 1 | FBP1 | | 0.88039 | 1.08011 | P09467;Q5VZC3 | P09467 |
| Fructose-bisphosphate aldolase A | ALDOA | | 0.533704 | -0.281918 | P04075;J3KPS3;P04075-2;H3BQN4;H3BPS8;H3BUH7;H3BR04;H3BMQ8; | P04075;J3KPS3;P04075-2;H3BQN4;H3BPS8;H3BUH7 |
| Fructose-bisphosphate aldolase C | ALDOC | | 0.852939 | -0.410769 | P09972;A8MVZ9;J3KSV6;J3QKP5;C9J8F3;K7EKH5;J3QKK1 | P09972;A8MVZ9;J3KSV6;J3QKP5 |
| Fumarate hydratase, mitochondrial | FH | | 0.24688 | 0.202497 | P07954-2;P07954 | P07954-2;P07954 |
| Fumarylacetoacetase | FAH | | 0.946453 | -0.564142 | P16930;Q53XA7;H0YLC7;H3BNP8 | P16930;Q53XA7 |
| Galactocerebrosidase | GALC | | 0.7238 | 0.406901 | P54803;P54803-4;P54803-5;E7EPA4;P54803-3;G3V5E8;G3V255;G3V4M | P54803;P54803-4;P54803-5;E7EPA4;P54803-3;G3V5E8 |
| Galactokinase | GALK1 | | 0.112471 | -0.160921 | P51570;B4E1G6;K7ERJ9;K7EII7;K7ERN9 | P51570;B4E1G6 |
| Galactosylgalactosylxylosylprotein 3-beta-glucuronosyltransferase 3 | B3GAT3 | | 0.156915 | -0.187894 | O94766;G3V150;B7ZAB3;E9PNA1;Q9NPZ5 | O94766;G3V150;B7ZAB3 |
| Galectin-1 | LGALS1 | | 0.430943 | 0.395115 | P09382;F8WEI7 | P09382 |
| Galectin-3;Galectin | LGALS3 | | 0.474601 | -0.440664 | P17931;G3V3R6 | P17931;G3V3R6 |
| Galectin-3-binding protein | LGALS3BP | | 0.548089 | -0.625649 | Q08380;K7EP36;K7ESM3;K7EKQ5;K7ERZ6;K7EJY8;K7EJD3;K7ES75;K7EN | Q08380 |
| Galectin-8;Galectin | LGALS8 | | 0.151655 | -0.245476 | O00214-2;E9PN19;F6V2D4;O00214;H7BXD8;Q5T3P9;B1ANM0;E9PJ77;I | O00214-2;E9PN19;F6V2D4;O00214;H7BXD8 |
| Gamma-enolase;Enolase | ENO2 | | 0.996102 | -0.682521 | P09104;B7Z2X9;F5H0C8;U3KQP4;F5H1C3 | P09104;B7Z2X9;F5H0C8 |
| Gamma-glutamyl hydrolase | GGH | | 0.563435 | -0.290085 | Q92820 | Q92820 |
| Gamma-glutamylcyclotransferase | GGCT | | 0.184708 | 0.113213 | O75223;M0QZK8;B8ZZK2;H7BZK5;O75223-3;O75223-2;O75223-4 | O75223;M0QZK8;B8ZZK2 |
| Gamma-interferon-inducible lysosomal thiol reductase | IFI30 | | 0.649215 | 0.276933 | P13284;M0QZG3 | P13284 |
| Ganglioside GM2 activator;Ganglioside GM2 activator isoform short | GM2A | | 0.708339 | -0.885008 | P17900;H0YBY3;E5RJD0 | P17900;H0YBY3 |
| GDP-L-fucose synthase | TSTA3 | | 0.120838 | 0.149092 | Q13630;E9PKL9;E9PP14;E9PLH9;E9PP60;H0YE90;H0YCP7 | Q13630;E9PKL9;E9PP14;E9PLH9 |
| GDP-mannose 4,6 dehydratase | GMDS | | 0.0656703 | -0.0470225 | O60547-2;O60547 | O60547-2;O60547 |
| Gelsolin | GSN | | 0.390618 | -0.34737 | P06396-2;P06396-4;P06396-3;P06396;Q5T0H8;Q5T0H7;Q5T0H9;Q5T0I | P06396-2;P06396-4;P06396-3;P06396;Q5T0H8;Q5T0H7 |
| Glia-derived nexin | SERPINE2 | | 0.876102 | -0.662116 | P07093-2;P07093-3;C9JN98;C9K031 | P07093-2;P07093-3;C9JN98 |
| Glucosamine-6-phosphate isomerase 1 | GNPDA1 | | 0.104615 | -0.0750018 | P46926;B7Z3X4;D6RAY7;D6R9P4;E7EVU7;D6RFF8;D6RB13;D6R917;D6I | P46926;B7Z3X4;D6RAY7;D6R9P4;E7EVU7;D6RFF8;D6RB13 |
| Glucose-6-phosphate 1-dehydrogenase | G6PD | | 0.795322 | 0.917097 | P11413;P11413-2;P11413-3;E7EUI8;E7EM57;E9PD92 | P11413;P11413-2;P11413-3;E7EUI8 |
| Glucose-6-phosphate isomerase | GPI | | 0.635316 | -0.337631 | P06744;P06744-2;K7EQ48;K7EP41;K7EPY4;K7ENA0;K7ELR7;K7ERC6;K7 | P06744;P06744-2;K7EQ48 |
| Glucosylceramidase | GBA | | 1.0031 | 0.474361 | P04062-4;P04062-2;P04062;F5H241;P04062-5;P04062-3 | P04062-4;P04062-2;P04062;F5H241;P04062-5;P04062-3 |
| Glutamine--fructose-6-phosphate aminotransferase [isomerizing] 1 | GFPT1 | | 0.301709 | 0.343532 | Q06210-2;Q06210;E5RJP4 | Q06210-2;Q06210 |
| Glutaredoxin-1 | GLRX | | 0.951043 | -0.48694 | P35754 | P35754 |
| Glutaredoxin-3 | GLRX3 | | 0.754325 | 0.783214 | O76003 | O76003 |
| Glutathione peroxidase 1 | GPX1 | + | 1.69801 | 1.97648 | P07203;P07203-2 | P07203 |
| Glutathione peroxidase 3 | GPX3 | | 0.23555 | 0.500821 | P22352;H0YBE4;E5RG32;H0YC19 | P22352 |
| Glutathione reductase, mitochondrial | GSR | | 0.0639526 | -0.0471678 | P00390-2;P00390;P00390-3;P00390-4;P00390-5;H0YBD4;E5RI06;H0YC6 | P00390-2;P00390;P00390-3;P00390-4;P00390-5 |
| Glutathione S-transferase kappa 1 | GSTK1 | | 0.131448 | 0.112521 | E9PFN5;Q9Y2Q3;Q9Y2Q3-4;Q9Y2Q3-3;Q9Y2Q3-2;C9JNT3 | E9PFN5;Q9Y2Q3;Q9Y2Q3-4;Q9Y2Q3-3;Q9Y2Q3-2 |
| Glutathione S-transferase Mu 3 | GSTM3 | | 0.0749687 | 0.104137 | P21266 | P21266 |
| Glutathione S-transferase omega-1 | GSTO1 | | 0.0177894 | -0.0135337 | P78417;Q5TA02;P78417-3;P78417-2;Q5TA01 | P78417;Q5TA02;P78417-3;P78417-2;Q5TA01 |
| Glutathione S-transferase P | GSTP1 | | 0.155468 | -0.111342 | P09211;A8MX94 | P09211;A8MX94 |
| Glutathione S-transferase theta-2B;Glutathione S-transferase theta-2 | Em:AP000351.3;GSTT2B;GSTT2 | | 0.0163641 | 0.017848 | Q6ICJ4;P0CG30;B5MCL2;P0CG29;G5E9I8;G5E9P7;F8WCF3;F6UD80;C9J | Q6ICJ4;P0CG30;B5MCL2;P0CG29 |
| Glutathione synthetase | GSS | | 0.128209 | -0.0943567 | P48637;B7Z514;P48637-2 | P48637;B7Z514;P48637-2 |
| Glyceraldehyde-3-phosphate dehydrogenase | GAPDH | | 0.472495 | -0.275911 | P04406;P04406-2;E7EUT5 | P04406;P04406-2;E7EUT5 |
| Glycine--tRNA ligase | GARS | | 0.454618 | 0.467105 | P41250;H7C443 | P41250 |
| Glycogen phosphorylase, brain form | PYGB | | 0.116173 | -0.0955167 | P11216;H0Y4Z6 | P11216 |
| Glycogen phosphorylase, liver form;Phosphorylase | PYGL | | 0.584871 | -0.350006 | P06737-2;P06737;E9PK47 | P06737-2;P06737;E9PK47 |
| Glycogenin-1 | GYG1 | + | 1.46077 | 0.581775 | P46976-2;P46976;C9JQ42;G5E9W8;P46976-3;C9J8R8;C9J7C7 | P46976-2;P46976;C9JQ42;G5E9W8;P46976-3 |
| Glycolipid transfer protein | GLTP | | 0.244663 | -0.23007 | Q9NZD2;F5H0U5;H0YFS9;F5GZ49 | Q9NZD2;F5H0U5 |
| Glyoxalase domain-containing protein 4 | GLOD4 | | 0.200143 | 0.286976 | Q9HC38-2;Q9HC38;B7Z403;I3L3Q4;Q9HC38-3;I3L1F4;I3L1I0;I3NI27;I3N | Q9HC38-2;Q9HC38;B7Z403;I3L3Q4;Q9HC38-3 |
| Glyoxylate reductase/hydroxypyruvate reductase | GRHPR | | 0.191902 | -0.21006 | Q9UBQ7;U3KQ56;Q5T946;Q9H636 | Q9UBQ7;U3KQ56;Q5T946 |
| Glypican-1;Secreted glypican-1 | GPC1 | | 0.000106025 | -0.000194329 | P35052;H7C410;H7C024;H7BZE9;H7BZL4;C9J4Y6 | P35052;H7C410;H7C024 |
| Glypican-6;Secreted glypican-6 | GPC6 | | 0.88923 | -0.80908 | Q9Y625 | Q9Y625 |
| Golgi apparatus protein 1 | GLG1 | | 0.330538 | 0.208439 | Q92896;Q92896-2;Q92896-3;H3BM42;H3BQU9;H3BQT1;H3BRD8 | Q92896;Q92896-2;Q92896-3;H3BM42 |
| Golgi membrane protein 1 | GOLM1 | | 0.50981 | 0.131753 | Q8NBJ4-2;Q8NBJ4;F8WAT9;C9JYM4;C9J941 | Q8NBJ4-2;Q8NBJ4 |
| Golgin subfamily A member 7 | GOLGA7 | | 0.851953 | -1.05926 | Q7Z5G4;Q7Z5G4-3 | Q7Z5G4;Q7Z5G4-3 |
| G-protein coupled receptor 56;GPR56 N-terminal fragment;GPR56 C-terminal fragment | GPR56 | | 0.939837 | -1.14917 | Q9Y653-2;Q9Y653-3;Q9Y653;H3BRH0;F5H144;Q9Y653-5;H3BSJ6;H3BN | Q9Y653-2;Q9Y653-3;Q9Y653;H3BRH0;F5H144;Q9Y653-5;H3BSJ6 |

| | | | | | | |
|---|---|---|---|---|---|---|
| Granulins;Acrogranin;Paragranulin;Granulin-1;Granulin-2;Granulin-3;Granulin-4;Granulin-5;Granulin-6;Gr | GRN | | 0.556878 | -0.557323 | P28799;P28799-2;K7EKL3;K7EQ05;P28799-3;K7EMR1;K7EQI0;K7EPL0;K | P28799;P28799-2;K7EKL3;K7EQ05;P28799-3 |
| GRIP1-associated protein 1 | GRIPAP1 | | 0.355339 | 0.459929 | B1B0M1;Q4V328-4;Q4V328;Q4V328-2;Q4V328-3 | B1B0M1;Q4V328-4;Q4V328;Q4V328-2 |
| Group XV phospholipase A2 | PLA2G15 | | 0.250447 | -0.110385 | Q8NCC3;H3BMU8;B4DJW4;H3BPT3;B4DUD1;H3BM47;B4DPU0 | Q8NCC3;H3BMU8;B4DJW4;H3BPT3 |
| Growth arrest-specific protein 6 | GAS6 | | 0.663512 | 0.402539 | J3KP07;Q14393-2;Q14393;Q14393-3;Q14393-5;Q14393-4 | J3KP07;Q14393-2;Q14393;Q14393-3;Q14393-5;Q14393-4 |
| Growth factor receptor-bound protein 2 | GRB2 | | 0.929597 | -0.635898 | P62993;P62993-2;J3KT38;J3QRL5;J3QLF6 | P62993;P62993-2;J3KT38 |
| Growth-regulated alpha protein;GRO-alpha(4-73);GRO-alpha(5-73);GRO-alpha(6-73) | CXCL1 | + | 1.8829 | -3.34477 | P09341 | P09341 |
| GTPase NRas | NRAS | | 0.772 | -0.929556 | P01111 | P01111 |
| GTP-binding nuclear protein Ran | RAN | | 0.00966528 | -0.0116761 | B5MDF5;P62826;J3KQE5;F5H018;B4DV51;H0YFC6 | B5MDF5;P62826;J3KQE5;F5H018;B4DV51 |
| GTP-binding protein Rheb | RHEB | | 0.649136 | 0.356852 | C9J931;Q15382 | C9J931;Q15382 |
| GTP-binding protein SAR1a | SAR1A | | 0.0253123 | -0.019383 | Q9NR31;B4DQ19;H0Y5E8;Q5SQT8;Q9Y6B6;X1WI22;D6RAA2;Q9H029;D | Q9NR31;B4DQ19;H0Y5E8 |
| Guanidinoacetate N-methyltransferase | GAMT | | 0.0502929 | 0.0636095 | Q14353;Q14353-2;K7EM34 | Q14353;Q14353-2 |
| Guanine nucleotide-binding protein G(i) subunit alpha-1 | GNAI1 | | 0.668073 | -1.11149 | P63096;P63096-2;C9JPP4;C9JL77 | P63096;P63096-2;C9JPP4 |
| Guanine nucleotide-binding protein G(i) subunit alpha-2 | GNAI2 | | 0.494267 | -0.626165 | P04899;P04899-4;P04899-6;P04899-3;P04899-5;P04899-2;F5GZL8;P09 | P04899;P04899-4;P04899-6;P04899-3;P04899-5;P04899-2;F5GZL8 |
| Guanine nucleotide-binding protein G(I)/G(S)/G(O) subunit gamma-12 | GNG12 | | 0.488974 | -0.682445 | Q9UBI6 | Q9UBI6 |
| Guanine nucleotide-binding protein G(I)/G(S)/G(T) subunit beta-1 | GNB1 | | 0.617016 | -0.532121 | P62873;F6UT28;B1AKQ8;F6X3N5;F5H8J8;F5H100;F5H0S8;E9PCP0;P165 | P62873;F6UT28 |
| Guanine nucleotide-binding protein G(I)/G(S)/G(T) subunit beta-2 | GNB2 | | 0.218084 | -0.192141 | P62879;E7EP32;C9JIS1;C9JXA5;C9JZN1;B3KPU1 | P62879;E7EP32;C9JIS1;C9JXA5;C9JZN1;B3KPU1 |
| Guanine nucleotide-binding protein G(k) subunit alpha | GNAI3 | | 0.636318 | -0.849585 | P08754 | P08754 |
| Guanine nucleotide-binding protein G(s) subunit alpha isoforms short;Guanine nucleotide-binding proteir | GNAS | | 0.821279 | -0.94269 | P63092;P63092-4;P63092-3;P63092-2;Q5JWF2;Q5JWF2-2;S4R3E3;Q5JV | P63092;P63092-4;P63092-3;P63092-2;Q5JWF2;Q5JWF2-2 |
| Guanine nucleotide-binding protein subunit alpha-11 | GNA11 | | 0.432721 | -0.420503 | P29992;K7EL62 | P29992;K7EL62 |
| Guanine nucleotide-binding protein subunit alpha-13 | GNA13 | | 0.726429 | -1.282 | Q14344;F5H1G8;Q03113;E7EUG4;E9PC54;B7Z3F7;B3KXS2;P19086 | Q14344;F5H1G8 |
| Guanine nucleotide-binding protein subunit beta-2-like 1;Guanine nucleotide-binding protein subunit bet | GNB2L1 | | 0.754581 | 0.739182 | P63244;J3KPE3;D6RAC2;D6RHH4;H0Y8W2;D6REE5;H0YAF8;D6R9Z1;H0 | P63244;J3KPE3;D6RAC2;D6RHH4;H0Y8W2;D6REE5;H0YAF8;D6R9Z1;H( |
| Guanine nucleotide-binding protein subunit beta-4 | GNB4 | | 0.711987 | -0.497709 | Q9HAV0;C9JD14;H7C5J5 | Q9HAV0 |
| Haloacid dehalogenase-like hydrolase domain-containing protein 2 | HDHD2 | | 0.062558 | -0.0486401 | K7ER15;Q9H0R4;K7EMY7;Q9H0R4-2;K7EQD2;K7EJQ8;K7ENF3 | K7ER15;Q9H0R4;K7EMY7;Q9H0R4-2;K7EQD2 |
| HD domain-containing protein 2 | HDDC2 | | 0.0194414 | -0.0217245 | Q7Z4H3;Q7Z4H3-2;V9GYS5;V9GZ04;Q7Z4H3-3 | Q7Z4H3;Q7Z4H3-2;V9GYS5;V9GZ04 |
| Heat shock 70 kDa protein 13 | HSPA13 | | 0.0248689 | 0.019048 | P48723;F5GWS8 | P48723;F5GWS8 |
| Heat shock 70 kDa protein 1A/1B | HSPA1A | | 0.177594 | -0.151347 | P08107;P08107-2;E7EP94;V9GZ37;P48741 | P08107;P08107-2;E7EP94;V9GZ37 |
| Heat shock 70 kDa protein 4 | HSPA4 | | 0.413051 | -0.246867 | P34932 | P34932 |
| Heat shock cognate 71 kDa protein | HSPA8 | | 0.0926004 | 0.126599 | P11142;E9PKE3;E9PNE6;P11142-2;A8K7Q2;E9PN89;E9PS65;E9PLF4;E9I | P11142;E9PKE3;E9PNE6;P11142-2;A8K7Q2 |
| Heat shock protein 105 kDa | HSPH1 | | 0.351102 | -0.262239 | Q92598-2;Q92598;B4DYH1;Q92598-3;B4DY72;R4GN69;Q5TBM3 | Q92598-2;Q92598;B4DYH1;Q92598-3;B4DY72 |
| Heat shock protein 75 kDa, mitochondrial | TRAP1 | | 0.530785 | 0.854732 | F5H897;Q12931;I3L0K7;I3L239;I3L2D5;I3L253 | F5H897;Q12931;I3L0K7 |
| Heat shock protein HSP 90-alpha | HSP90AA1 | | 0.685104 | -0.506419 | P07900;P07900-2;Q86U12;G3V2J8;Q58FG0;Q58FG1 | P07900;P07900-2;Q86U12 |
| Heat shock protein HSP 90-beta | HSP90AB1 | | 0.280728 | -0.196497 | P08238;Q58FF7;Q58FF8;Q5T9W8 | P08238 |
| Heme-binding protein 1 | HEBP1 | | 0.540448 | -0.409445 | Q9NRV9;F5GWX2;H0YG71 | Q9NRV9 |
| Heme-binding protein 2 | HEBP2 | | 0.623038 | -0.326277 | Q9Y5Z4;Q9Y5Z4-2;Q5THN1;C9IZA0 | Q9Y5Z4;Q9Y5Z4-2 |
| Heparan-sulfate 6-O-sulfotransferase 2 | HS6ST2 | | 0.802856 | -0.4714 | J3KP99;Q96MM7-2;Q96MM7-3;Q96MM7;Q96MM7-4;B7Z5H6 | J3KP99;Q96MM7-2;Q96MM7-3;Q96MM7;Q96MM7-4;B7Z5H6 |
| Hepatocyte growth factor receptor | MET | | 0.313348 | 0.132935 | P08581;P08581-2;P08581-3;H7C174;C9JKM5;H7C130;B4DPY6 | P08581;P08581-2;P08581-3 |
| Hepatocyte growth factor-regulated tyrosine kinase substrate | HGS | | 0.139943 | 0.143926 | O14964-2;O14964;I3L1P5;I3L165;I3L1E3;I3L2H4 | O14964-2;O14964;I3L1P5 |
| Hepatoma-derived growth factor | HDGF | | 0.0891315 | 0.075794 | P51858;A8K8G0;P51858-2;P51858-3;O75475-3;O75475-2 | P51858;A8K8G0;P51858-2;P51858-3 |
| Heterogeneous nuclear ribonucleoprotein A1;Heterogeneous nuclear ribonucleoprotein A1, N-terminally | HNRNPA1;HNRNPA1L2 | | 0.25803 | 0.163939 | F8W6I7;P09651-2;P09651;P09651-3;F8VZ49;Q32P51;F8VTQ5;F8W646; | F8W6I7;P09651-2;P09651;P09651-3;F8VZ49;Q32P51;F8VTQ5 |
| Heterogeneous nuclear ribonucleoprotein D0 | HNRNPD | | 0.654318 | -0.583899 | Q14103-3;Q14103;H0Y8G5;D6RAF8;D6RD83 | Q14103-3;Q14103;H0Y8G5;D6RAF8 |
| Heterogeneous nuclear ribonucleoprotein D-like | HNRNPDL | | 0.675022 | 0.467187 | O14979-3;O14979-2;O14979 | O14979-3;O14979-2;O14979 |
| Heterogeneous nuclear ribonucleoprotein K | HNRNPK | | 0.236496 | 0.96726 | P61978;P61978-2;Q5T6W1;S4R457;S4R359 | P61978;P61978-2;Q5T6W1 |
| Heterogeneous nuclear ribonucleoprotein K | HNRNPK | | 0.354298 | 0.324403 | P61978-3;Q5T6W5;Q5T6W2 | P61978-3;Q5T6W5;Q5T6W2 |
| Heterogeneous nuclear ribonucleoprotein L | HNRNPL | | 0.21988 | -0.170012 | P14866;M0QXS5;P14866-2;M0R1W6;M0QYL7;M0R076;B4DVF8;H7C46 | P14866;M0QXS5;P14866-2 |
| Heterogeneous nuclear ribonucleoprotein M | HNRNPM | | 0.824993 | -0.580384 | P52272-2;P52272;M0R019;M0R2T0;M0QYQ7;M0R0N3;M0QY96;M0R2 | P52272-2;P52272;M0R019 |
| Heterogeneous nuclear ribonucleoproteins C1/C2 | HNRNPC | | 0.56315 | -0.268577 | G3V4W0;B4DY08;P07910-2;G3V4C1;P07910-4;P07910;G3V2Q1;G3V57 | G3V4W0;B4DY08;P07910-2;G3V4C1;P07910-4;P07910;G3V2Q1;G3V57 |
| High mobility group protein B1;Putative high mobility group protein B1-like 1 | HMGB1;HMGB1P1 | | 0.376183 | -0.385601 | P09429;Q5T7C4;Q5T7C6;Q5T7C0;B2RPK0;P23497 | P09429;Q5T7C4;Q5T7C6;Q5T7C0;B2RPK0 |
| High mobility group protein B2 | HMGB2 | | 0.114011 | -0.168933 | P26583;D6R9A6 | P26583;D6R9A6 |
| Hippocalcin-like protein 1;Neuron-specific calcium-binding protein hippocalcin | HPCAL1;HPCA | | 0.0713689 | 0.0936553 | P37235;E9PC71;P84074;P61601;H7BZC1;E5RGA9;C9JW46;E5RK89;E5RI | P37235;E9PC71;P84074 |
| Histamine N-methyltransferase | HNMT | | 0.0652713 | 0.118951 | P50135;P50135-3;P50135-2 | P50135 |
| Histidine triad nucleotide-binding protein 1 | HINT1 | | 0.651198 | 0.610482 | P49773;D6REP8;D6RE99;D6RD60;D6RC06 | P49773 |
| Histidine triad nucleotide-binding protein 3 | HINT3 | | 0.268159 | -0.53401 | Q9NQE9 | Q9NQE9 |
| Histone H2A type 1-C;Histone H2A type 3;Histone H2A type 1-B/E | HIST1H2AC;HIST3H2A;HIST1H2 | | 0.672254 | -0.869185 | Q93077;Q7L7L0;P04908 | Q93077;Q7L7L0;P04908 |
| Histone H2A type 1-J;Histone H2A type 1-H;Histone H2A.J;Histone H2A type 2-C;Histone H2A type 2-A;His | HIST1H2AJ;HIST1H2AH;H2AFJ; | + | 1.51931 | -0.994491 | Q99878;Q96KK5;Q9BTM1;Q16777;Q6FI13;P20671;P0C0S8;H0YFX9;Q9E | Q99878;Q96KK5;Q9BTM1;Q16777;Q6FI13;P20671;P0C0S8;H0YFX9;Q9 |
| Histone H2A.V;Histone H2A.Z;Histone H2A | H2AFV;H2AFZ | | 0.184468 | -0.633661 | Q71UI9;P0C0S5;C9J0D1;Q71UI9-3;Q71UI9-4;Q71UI9-2;C9J386 | Q71UI9;P0C0S5;C9J0D1;Q71UI9-3;Q71UI9-4;Q71UI9-2;C9J386 |
| Histone H2B type 2-E;Histone H2B type 1-B;Histone H2B type 1-O;Histone H2B type 1-J;Histone H2B type | HIST2H2BE;HIST1H2BB;HIST1H | | 0.841171 | -0.743387 | Q16778;P33778;P23527;P06899;Q8N257;Q6DRA6;Q6DN03 | Q16778;P33778;P23527;P06899;Q8N257 |
| Histone H2B;Histone H2B type 1-M;Histone H2B type 1-N;Histone H2B type 1-H;Histone H2B type 2-F;His | HIST1H2BN;HIST1H2BM;HIST1 | | 0.989416 | -0.871122 | U3KQK0;Q99879;Q99877;Q93079;Q5QNW6;P62807;P58876;Q5QNW6- | U3KQK0;Q99879;Q99877;Q93079;Q5QNW6;P62807;P58876;Q5QNW6 |
| Histone H3;Histone H3.3 | H3F3B;H3F3A | | 0.956309 | -0.622205 | K7EK07;P84243 | K7EK07;P84243 |
| Histone H4 | HIST1H4A | | 1.02021 | -0.676215 | P62805 | P62805 |
| HLA class I histocompatibility antigen, A-2 alpha chain;HLA class I histocompatibility antigen, A-69 alpha cl | HLA-A | | 0.219956 | 0.300305 | P01892;P10316;P01891;P30447;P05534;Q5SRN7;Q5SRN5;P16188;P04 | P01892;P10316;P01891;P30447;P05534 |
| HLA class I histocompatibility antigen, Cw-12 alpha chain;HLA class I histocompatibility antigen, Cw-16 alp | HLA-C | | 0.0327006 | 0.138883 | P30508;Q29960-2;Q29960;P30505 | P30508;Q29960-2;Q29960;P30505 |
| Hsc70-interacting protein;Putative protein FAM10A5;Putative protein FAM10A4 | ST13;ST13P5;ST13P4 | | 0.483068 | 0.805687 | P50502;Q8NFI4;H7C3I1;F6VDH7;Q8IZP2;F8WAQ7 | P50502;Q8NFI4;H7C3I1;F6VDH7;Q8IZP2 |
| Hsp90 co-chaperone Cdc37;Hsp90 co-chaperone Cdc37, N-terminally processed | CDC37 | | 1.10913 | 1.12294 | Q16543;K7EQA9;K7EKQ2;K7EL68;K7EIU0 | Q16543;K7EQA9;K7EKQ2 |
| Hyaluronan and proteoglycan link protein 3 | HAPLN3 | | 0.972455 | -0.547687 | H3BTH8;Q96S86;H3BRY4;H3BT26;P10915;Q86UW8 | H3BTH8;Q96S86 |
| Hydroxyacylglutathione hydrolase, mitochondrial | HAGH | | 0.780937 | -0.531843 | H3BPK3;Q16775-2;Q16775;H3BPQ4;H3BQW8;H3BV79;E7EN93 | H3BPK3;Q16775-2;Q16775;H3BPQ4 |
| Hypoxanthine-guanine phosphoribosyltransferase | HPRT1 | | 0.988769 | -0.611834 | P00492 | P00492 |
| Hypoxia up-regulated protein 1 | HYOU1 | | 1.04158 | 1.09962 | E9PL22;Q9Y4L1;E9PJ21;B7Z909;J3KTF1;J3QL06;J3QQH7;J3QLE9;J3KT27 | E9PL22;Q9Y4L1;E9PJ21;B7Z909 |
| Immunoglobulin superfamily member 8 | IGSF8 | | 0.479072 | -0.241877 | Q969P0-3;Q969P0;C9J8Z4;Q969P0-2 | Q969P0-3;Q969P0;C9J8Z4 |
| Importin subunit alpha-3 | KPNA4 | | 0.159986 | -0.143193 | O00629;H7C4F6 | O00629 |
| Importin subunit beta-1 | KPNB1 | | 0.740087 | -0.631369 | Q14974;Q14974-2;F5H4R7;J3KTM9;J3QR48;J3QKQ5;J3QRG4;J3KS06 | Q14974;Q14974-2;F5H4R7;J3KTM9 |
| Inhibin beta B chain | INHBB | | 0.395467 | -0.742448 | P09529 | P09529 |
| Inorganic pyrophosphatase | PPA1 | | 0.657133 | 0.575351 | Q15181;Q5SQT6;H0YAK2 | Q15181;Q5SQT6 |
| Inorganic pyrophosphatase 2, mitochondrial | PPA2 | | 0.425822 | 0.473942 | Q9H2U2;Q9H2U2-2;E2QRM6;H0Y9D8;Q9H2U2-3;D6R967;B4DFH3;Q9H | Q9H2U2;Q9H2U2-2;E2QRM6;H0Y9D8;Q9H2U2-3;D6R967;B4DFH3;Q9H |
| Inosine-5-monophosphate dehydrogenase 2 | IMPDH2 | | 0.590492 | 0.749563 | P12268;H0Y4R1;E7ETK5 | P12268;H0Y4R1 |
| Inositol monophosphatase 1 | IMPA1 | | 0.827731 | -0.679866 | P29218;P29218-3;H0YBL1;E5RG13;E5RI82;E5RIP7;P29218-2;E5RG94;E5 | P29218;P29218-3;H0YBL1;E5RG13;E5RI82;E5RIP7;P29218-2;E5RG94;E5 |
| Insulin-degrading enzyme | IDE | | 0.0882259 | 0.0719453 | P14735;P14735-2;Q5T5N3 | P14735;P14735-2 |
| Insulin-like growth factor-binding protein 1 | IGFBP1 | + | 1.53961 | 0.59832 | P08833;C9JXF9;C9J6H2 | P08833;C9JXF9;C9J6H2 |
| Insulin-like growth factor-binding protein 3 | IGFBP3 | | 0.318146 | -0.15021 | P17936;P17936-2;B3KWK7;H0Y5K2;H0Y485;C9JMX4 | P17936;P17936-2;B3KWK7;H0Y5K2;H0Y485;C9JMX4 |
| Insulin-like growth factor-binding protein 4 | IGFBP4 | | 0.778312 | -0.961047 | P22692;B4E351 | P22692;B4E351 |

| | | | | | | |
|---|---|---|---|---|---|---|
| Insulin-like growth factor-binding protein 6 | IGFBP6 | | 1.0868 | -0.982909 | P24592;F8VYK9;F8VVA5 | P24592;F8VYK9 |
| Insulin-like growth factor-binding protein 7 | IGFBP7 | | 1.222 | -0.796671 | Q16270;Q16270-2 | Q16270;Q16270-2 |
| Insulin-like growth factor-binding protein-like 1 | IGFBPL1 | | 0.485203 | -0.279524 | Q8WX77 | Q8WX77 |
| Integral membrane protein 2B;BRI2, membrane form;BRI2 intracellular domain;BRI2C, soluble form;Bri23 | ITM2B | | 0.0807336 | -0.233132 | Q9Y287;Q5W0A2;U3KQL7 | Q9Y287;Q5W0A2 |
| Integral membrane protein GPR180 | GPR180 | | 0.569463 | -0.0633093 | Q86V85 | Q86V85 |
| Integrin alpha-3;Integrin alpha-3 heavy chain;Integrin alpha-3 light chain | ITGA3 | | 0.976854 | -1.29408 | P26006;P26006-1;D6R9X8;H0YA49;B4DDT0;K7EMU3;H0YA32 | P26006;P26006-1 |
| Integrin alpha-6;Integrin alpha-6 heavy chain;Integrin alpha-6 light chain;Processed integrin alpha-6 | ITGA6 | | 0.813815 | -1.48592 | P23229-4;P23229-2;P23229-5;P23229-3;P23229-9;P23229-6;P23229;P2 | P23229-4;P23229-2;P23229-5;P23229-3;P23229-9;P23229-6;P23229;P |
| Integrin alpha-V;Integrin alpha-V heavy chain;Integrin alpha-V light chain | ITGAV | | 0.38576 | -1.13676 | P06756-3;P06756-2;P06756;H7BZG1 | P06756-3;P06756-2;P06756 |
| Integrin beta-1 | ITGB1 | | 0.748458 | -1.01002 | P05556;P05556-2;P05556-5;P05556-4;P05556-3;E7EQW5;C9JPK5;Q5T3 | P05556;P05556-2;P05556-5;P05556-4;P05556-3 |
| Intercellular adhesion molecule 1 | ICAM1 | + | 1.9153 | -1.11207 | P05362;K7EKL8;E7ESS4 | P05362;K7EKL8 |
| Interferon-induced transmembrane protein 1;Interferon-induced transmembrane protein 2;Interferon-in | IFITM2;IFITM3;IFITM1 | | 1.17267 | -0.859609 | H7BYV1;E9PS44;E9PQN9;P13164;Q01629;Q01628 | H7BYV1;E9PS44;E9PQN9;P13164;Q01629;Q01628 |
| Interleukin-1 receptor accessory protein | IL1RAP | | 0.0461989 | 0.0413501 | Q9NPH3-3;Q9NPH3-2;Q9NPH3;Q9NPH3-5;C9J1D9;C9J9W1;C9JT28;H7C | Q9NPH3-3;Q9NPH3-2;Q9NPH3;Q9NPH3-5 |
| Interleukin-11 | IL11 | + | 1.8815 | 0.892343 | P20809 | P20809 |
| Interleukin-18 | IL18 | | 0.719326 | 0.622093 | Q14116;Q14116-2 | Q14116;Q14116-2 |
| Interleukin-6 | IL6 | + | 1.38873 | -2.41229 | B5MCZ3;P05231;B5MC21;B5MC14;B4DNQ5;C9J5B0;B4DNV3 | B5MCZ3;P05231;B5MC21;B5MC14;B4DNQ5 |
| Interleukin-6 receptor subunit beta | IL6ST | | 0.943373 | -0.36892 | P40189;P40189-3;D6RH03;P40189-2;Q5FC05 | P40189;P40189-3 |
| Interstitial collagenase;22 kDa interstitial collagenase;27 kDa interstitial collagenase | MMP1 | | 0.827018 | -1.3219 | P03956 | P03956 |
| Intraflagellar transport protein 25 homolog | HSPB11 | | 0.186852 | -0.162374 | Q9Y547;A6NIR2 | Q9Y547;A6NIR2 |
| Isoamyl acetate-hydrolyzing esterase 1 homolog | IAH1 | + | 1.4639 | -1.04449 | Q2TAA2;H7C5G1;B4DMV3;C9JE02;C9JDY4;C9J5J2;F8WF34 | Q2TAA2;H7C5G1;B4DMV3;C9JE02 |
| Isochorismatase domain-containing protein 1 | ISOC1 | | 1.29481 | -0.931015 | Q96CN7;D6RGE2 | Q96CN7 |
| Isocitrate dehydrogenase [NADP] cytoplasmic | IDH1 | | 0.142723 | 0.130151 | O75874;C9J4N6;C9JLU6;C9JJE5 | O75874 |
| Isoleucine--tRNA ligase, cytoplasmic | IARS | | 0.0891844 | -0.155968 | P41252;J3KR24;Q5TCD1;Q5TCC5;Q5TCD2;Q5TCC4;Q5TCC6 | P41252;J3KR24 |
| IST1 homolog | IST1 | | 0.725162 | -0.479905 | H3BMU1;H3BUI0;P53990-2;P53990-3;P53990;P53990-4;P53990-5;J3KR | H3BMU1;H3BUI0;P53990-2;P53990-3;P53990;P53990-4;P53990-5;J3KI |
| Keratin, type I cytoskeletal 18 | KRT18 | | 0.419955 | 0.206741 | P05783;F8VZY9;CON__H-INV:HIT000015463 | P05783;F8VZY9 |
| Ketohexokinase | KHK | | 1.186 | -0.573413 | P50053;C9JDL1;P50053-2 | P50053;C9JDL1;P50053-2 |
| Kinesin-like protein KIF23 | KIF23 | + | 1.44827 | -1.12203 | H7BYN4;Q02241-2;Q02241;B4E1K0;H0YMJ4;H0YME6 | H7BYN4;Q02241-2;Q02241;B4E1K0 |
| Kunitz-type protease inhibitor 2 | SPINT2 | | 0.303547 | -0.39584 | O43291;K7EM91;O43291-2;K7ESI5;K7EKQ0;K7EKC8;K7ES27 | O43291;K7EM91;O43291-2;K7ESI5;K7EKQ0 |
| Lactadherin;Lactadherin short form;Medin | MFGE8 | | 0.298705 | -0.352851 | F5H7N9;Q08431;F5GZN3;Q08431-2;Q08431-3;H0YKS8 | F5H7N9;Q08431;F5GZN3;Q08431-2;Q08431-3;H0YKS8 |
| Lactoylglutathione lyase | GLO1 | | 1.08093 | -0.765679 | Q04760;Q04760-2 | Q04760;Q04760-2 |
| Lambda-crystallin homolog | CRYL1 | + | 1.49816 | -1.34551 | Q9Y2S2-2;Q9Y2S2 | Q9Y2S2-2;Q9Y2S2 |
| Laminin subunit alpha-5 | LAMA5 | | 0.620384 | -0.591567 | O15230;Q5JTP4;E2QRK0;F2Z2S4 | O15230 |
| Laminin subunit beta-2 | LAMB2 | | 0.424618 | -0.287775 | P55268;F5H520 | P55268 |
| Laminin subunit beta-3 | LAMB3 | | 0.218554 | -0.13242 | Q13751;Q5THA1;X1WI29 | Q13751 |
| Laminin subunit gamma-1 | LAMC1 | + | 1.40043 | -0.734101 | P11047;R4GNC7 | P11047 |
| Laminin subunit gamma-2 | LAMC2 | + | 1.68758 | -1.02278 | Q13753-2;Q13753 | Q13753-2;Q13753 |
| L-aminoadipate-semialdehyde dehydrogenase-phosphopantetheinyl transferase | AASDHPPT | | 0.124922 | 0.13504 | Q9NRN7;E9PLW6;E9PNF3;B4DDW7 | Q9NRN7;E9PLW6 |
| LanC-like protein 1 | LANCL1 | | 0.13734 | 0.14117 | O43813;E9PHS0;F8WDS9;H7C2E3 | O43813;E9PHS0 |
| Large neutral amino acids transporter small subunit 1 | SLC7A5 | | 0.212995 | -0.40472 | Q01650;H0YJ95;G3V4Z6;Q9UM01;Q92536 | Q01650 |
| Latent-transforming growth factor beta-binding protein 1 | LTBP1 | + | 1.93692 | 0.410383 | E7EV71;Q14766-2;Q14766-5;Q14766;Q14766-4;Q14766-3;C9JD84;E7EU | E7EV71;Q14766-2;Q14766-5;Q14766;Q14766-4;Q14766-3;C9JD84 |
| Latent-transforming growth factor beta-binding protein 3 | LTBP3 | | 0.247697 | -0.299268 | E9PKW1;Q9NS15;Q9NS15-2;F5GWC4;H0YC99;E9PJR2;H0YCX6;E9PR84; | E9PKW1;Q9NS15;Q9NS15-2 |
| Latent-transforming growth factor beta-binding protein 4 | LTBP4 | + | 1.72949 | -2.71189 | E7EUU1;E7ENG9;Q8N2S1-2;Q8N2S1-3;Q8N2S1;F5GXC9;Q8N2S1-4;M0C | E7EUU1;E7ENG9;Q8N2S1-2;Q8N2S1-3;Q8N2S1;F5GXC9 |
| Latexin | LXN | | 0.493033 | -0.419981 | Q9BS40 | Q9BS40 |
| Legumain | LGMN | | 0.83123 | -0.636727 | Q99538;Q86TV3;Q86TV2;G3V2T4;G3V4E4;G3V4P5;H0YJN9;G3V3Z4 | Q99538;Q86TV3;Q86TV2 |
| Leucine-rich repeat transmembrane protein FLRT2 | FLRT2 | | 0.456847 | 0.513131 | O43155 | O43155 |
| Leucine-rich repeat-containing protein 16A | LRRC16A | | 0.196128 | -0.208547 | Q5VZK9-2;Q5VZK9;Q5VZK9-3;Q5VZK9-4 | Q5VZK9-2;Q5VZK9;Q5VZK9-3 |
| Leucine-rich repeat-containing protein 57 | LRRC57 | | 0.662643 | -0.44663 | H3BSW0;Q8N9N7 | H3BSW0;Q8N9N7 |
| Leucyl-cystinyl aminopeptidase;Leucyl-cystinyl aminopeptidase, pregnancy serum form | LNPEP | + | 1.45976 | 0.639176 | Q9UIQ6-3;Q9UIQ6-2;Q9UIQ6 | Q9UIQ6-3;Q9UIQ6-2;Q9UIQ6 |
| Leukemia inhibitory factor | LIF | + | 1.47742 | 0.487814 | P15018 | P15018 |
| Leukocyte elastase inhibitor | SERPINB1 | | 0.460807 | -0.305713 | P30740;B4DNT0;C9J7N5;O75830 | P30740;B4DNT0 |
| Leukotriene A-4 hydrolase | LTA4H | | 0.41418 | -0.269897 | P09960;P09960-4;P09960-2;P09960-3;B4DEH5 | P09960;P09960-4;P09960-2;P09960-3 |
| LIM and SH3 domain protein 1 | LASP1 | | 0.630766 | 0.569044 | Q14847;Q14847-2;C9J9W2;Q14847-3;F6S2S5;K7ESD6;J3KSN1;O76041- | Q14847;Q14847-2;C9J9W2;Q14847-3 |
| Lipolysis-stimulated lipoprotein receptor | LSR | + | 1.3363 | 0.510316 | B4DKL4;S4R3V8;Q86X29-5;Q86X29-4;Q86X29;Q86X29-3;Q86X29-2 | B4DKL4;S4R3V8;Q86X29-5;Q86X29-4;Q86X29;Q86X29-3 |
| L-lactate dehydrogenase A chain | LDHA | | 0.944997 | -0.571753 | P00338;P00338-3;P00338-4;P00338-5;P00338-2;F5GXY2;F5GYU2;F5GX | P00338;P00338-3;P00338-4;P00338-5;P00338-2;F5GXY2 |
| L-lactate dehydrogenase B chain;L-lactate dehydrogenase | LDHB | | 0.911905 | -0.46765 | P07195;A8MW50;C9J7H8;F5H793 | P07195;A8MW50;C9J7H8 |
| Low molecular weight phosphotyrosine protein phosphatase | ACP1 | | 0.165845 | 0.180122 | P24666;P24666-3;G5E9R5;P24666-4;F2Z2Q9 | P24666;P24666-3;G5E9R5;P24666-4 |
| Low-density lipoprotein receptor | LDLR | | 0.411947 | 0.247091 | J3KMZ9;P01130-5;P01130;H0YMD1;P01130-3;P01130-4;P01130-6;P01 | J3KMZ9;P01130-5;P01130;H0YMD1;P01130-3;P01130-4;P01130-6;P01 |
| Low-density lipoprotein receptor-related protein 6 | LRP6 | | 0.157737 | -0.111143 | F5H7J9;O75581;H0YGW5;E9PHY1;F5H0Z3 | F5H7J9;O75581;H0YGW5 |
| L-xylulose reductase | DCXR | | 0.858136 | -0.51236 | J3KS22;Q7Z4W1;J3QS36;J3KRZ4;J3QS45;J3KSZ5;J3QL34 | J3KS22;Q7Z4W1;J3QS36;J3KRZ4;J3QS45;J3KSZ5;J3QL34 |
| Lymphocyte antigen 96 | LY96 | | 0.541012 | -0.610913 | Q9Y6Y9;E5RJJ7 | Q9Y6Y9;E5RJJ7 |
| Lymphocyte function-associated antigen 3 | CD58 | | 0.338883 | -0.714844 | B1AMW1;P19256-2;P19256-3;P19256;H0YDI1 | B1AMW1;P19256-2;P19256-3;P19256;H0YDI1 |
| Lysine--tRNA ligase | KARS | | 0.866194 | 0.662106 | Q15046;Q15046-2;H3BVA8;J3KRL2;H3BPV7;H3BRC9 | Q15046;Q15046-2 |
| Lysophospholipase-like protein 1 | LYPLAL1 | | 0.669666 | -0.357281 | Q5VWZ2;Q5VWZ2-2 | Q5VWZ2;Q5VWZ2-2 |
| Lysosomal acid lipase/cholesteryl ester hydrolase | LIPA | | 0.649138 | 0.429222 | P38571-2;P38571;Q5T073;Q5T770 | P38571-2;P38571;Q5T073 |
| Lysosomal alpha-glucosidase;76 kDa lysosomal alpha-glucosidase;70 kDa lysosomal alpha-glucosidase | GAA | | 0.60632 | -0.588849 | P10253;I3L0S5;I3L3L3;I3L2V9 | P10253 |
| Lysosomal alpha-mannosidase;Lysosomal alpha-mannosidase A peptide;Lysosomal alpha-mannosidase B | MAN2B1 | | 1.14212 | 0.467794 | O00754;O00754-2;M0QZ24;M0R174;M0QYZ1;M0R2P5;M0QZG6 | O00754;O00754-2 |
| Lysosomal protective protein;Lysosomal protective protein 32 kDa chain;Lysosomal protective protein 20 | CTSA | | 0.840767 | 0.326229 | P10619-2;P10619;Q5JZG9;Q5JZH0;U3KQU6;U3KQ41;U3KQF1 | P10619-2;P10619 |
| Lysosomal Pro-X carboxypeptidase | PRCP | | 0.724771 | -0.387325 | P42785;P42785-2;B7Z7Q6;E9PIG4;E9PQB5;E9PL85;E9PQN3;E9PKN6;E9 | P42785;P42785-2;B7Z7Q6;E9PIG4 |
| Lysosomal thioesterase PPT2 | PPT2 | | 0.434624 | -0.325587 | Q9UMR5;Q9UMR5-3;G8JLL2;Q9UMR5-2;A2ABN7;A2ABN6;B7Z424 | Q9UMR5;Q9UMR5-3;G8JLL2;Q9UMR5-2 |
| Lysosome-associated membrane glycoprotein 1 | LAMP1 | | 0.17658 | -0.0995962 | P11279;B4DWL3 | P11279;B4DWL3 |
| Lysosome-associated membrane glycoprotein 2 | LAMP2 | | 0.101575 | -0.0687386 | B4E2S7;P13473-2;P13473;P13473-3;B7Z2R9;H0YCG2 | B4E2S7;P13473-2;P13473;P13473-3;B7Z2R9 |
| Lysyl oxidase homolog 2 | LOXL2 | | 1.10971 | -1.28819 | Q9Y4K0;H0YAP6;E5RHH3;E5RFY0;E5RJL2;E5RI22 | Q9Y4K0 |
| Lysyl oxidase homolog 4 | LOXL4 | | 0.16266 | 0.148822 | Q96JB6 | Q96JB6 |
| Macrophage colony-stimulating factor 1;Processed macrophage colony-stimulating factor 1 | CSF1 | + | 1.60917 | -1.40213 | E9PKP4;E9PJA2;P09603-3;P09603-2;P09603;E9PQ08;H7BY18 | E9PKP4;E9PJA2;P09603-3;P09603-2;P09603;E9PQ08 |
| Macrophage migration inhibitory factor | MIF | | 0.257727 | -0.157098 | P14174 | P14174 |
| Macrophage-capping protein | CAPG | | 0.341196 | -0.198866 | P40121;P40121-2;E7ENU9;B8ZZL6;H7C0X8 | P40121;P40121-2;E7ENU9 |
| Magnesium-dependent phosphatase 1 | NEDD8-MDP1;MDP1 | | 0.654335 | -0.39006 | E9PL57;Q86V88;F8VSA6;E9PS38;Q86V88-3;Q86V88-2 | E9PL57;Q86V88 |
| Major prion protein | PRNP | | 0.142291 | -0.0894776 | A2A2V1;P04156-2;P04156 | A2A2V1;P04156-2;P04156 |
| Major vault protein | MVP | | 0.289179 | 0.417026 | Q14764;H3BP76;H3BUK7;H3BNF6;H3BRL2;H3BQK6 | Q14764 |
| Malate dehydrogenase, cytoplasmic;Malate dehydrogenase | MDH1 | | 0.284823 | -0.178375 | P40925;P40925-3;P40925-2;B9A041;B8ZZ51;C9JF79;C9JRL4;C9JLV6;C9I | P40925;P40925-3;P40925-2;B9A041;B8ZZ51;C9JF79 |

| | | | | | | |
|---|---|---|---|---|---|---|
| Malate dehydrogenase, mitochondrial;Malate dehydrogenase | MDH2 | | 0.429589 | 0.350277 | P40926;E9PDB2;G3XAL0 | P40926;E9PDB2;G3XAL0 |
| Mammalian ependymin-related protein 1 | UCC1;EPDR1 | | 0.402969 | 0.125136 | A4D1W8;Q9UM22;Q9UM22-3;D6RIH7;Q9UM22-2 | A4D1W8;Q9UM22;Q9UM22-3 |
| Mannose-1-phosphate guanyltransferase alpha | GMPPA | | 0.044877 | -0.0187369 | Q96IJ6;Q96IJ6-2;F8WD54;C9J255;C9JAH0 | Q96IJ6;Q96IJ6-2;F8WD54 |
| Mannose-1-phosphate guanyltransferase beta | GMPPB | | 0.205114 | 0.406932 | Q9Y5P6;Q9Y5P6-2 | Q9Y5P6;Q9Y5P6-2 |
| Mannose-6-phosphate isomerase | MPI | | 0.59286 | -0.75916 | P34949-2;P34949;F5GX71;H3BPB8;H3BUG1;H3BU66;H3BN01;H3BQX0; | P34949-2;P34949;F5GX71;H3BPB8 |
| Mannosyl-oligosaccharide glucosidase | MOGS | | 0.176995 | -0.0699604 | Q13724-2;Q13724;C9J8D4;C9JDQ1;B8ZZE2 | Q13724-2;Q13724 |
| MAP kinase-activated protein kinase 3 | MAPKAPK3 | | 0.044592 | 0.0732246 | Q16644;C9J8E1;C9JPW3;P49137-2;P49137;H7C0G6;Q96PN8;O43930;P! | Q16644;C9J8E1 |
| MARCKS-related protein | MARCKSL1 | | 1.19944 | 1.07543 | P49006 | P49006 |
| Matrilin-2 | MATN2 | | 1.11829 | -1.88587 | J3KN08;O00339-2;O00339;O00339-3;H0YBJ4;O00339-4;H0YBU5;H0YBN | J3KN08;O00339-2;O00339;O00339-3;H0YBJ4;O00339-4 |
| Matrilin-3 | MATN3 | | 0.612585 | -1.00158 | O15232-2;O15232 | O15232-2;O15232 |
| Matrix metalloproteinase-14 | MMP14 | | 0.272698 | 0.196743 | P50281;F8W1B7;Q9Y5R2;P51511 | P50281 |
| Melanotransferrin | MFI2 | | 0.0484078 | 0.052323 | P08582;P08582-2;C9JVN1 | P08582 |
| Membrane-associated progesterone receptor component 1 | PGRMC1 | | 1.29334 | -1.06969 | O00264;O00264-2 | O00264;O00264-2 |
| Mesencephalic astrocyte-derived neurotrophic factor | MANF | | 0.271921 | 0.431634 | P55145;H7C2D6 | P55145 |
| Mesothelin;Megakaryocyte-potentiating factor;Mesothelin, cleaved form | MSLN | + | 1.44927 | -1.22898 | Q13421-4;Q13421-3;Q13421-2;Q13421;H3BR90;H3BMA1;H3BUX1;H3B | Q13421-4;Q13421-3;Q13421-2;Q13421;H3BR90;H3BMA1;H3BUX1 |
| Metalloproteinase inhibitor 1 | TIMP1 | | 0.0385024 | -0.0162942 | P01033;Q5H9A7;Q5H9B5;Q5H9B4 | P01033;Q5H9A7 |
| Metalloproteinase inhibitor 2 | TIMP2 | | 1.00306 | -1.19367 | P16035;B4DFW2;K7EIX4;K7EL90 | P16035;B4DFW2;K7EIX4 |
| Metalloproteinase inhibitor 3 | TIMP3 | | 1.2931 | -1.7398 | P35625 | P35625 |
| Metastasis-suppressor KiSS-1;Metastin;Kisspeptin-14;Kisspeptin-13;Kisspeptin-10 | KISS1 | + | 1.69007 | 0.61298 | Q15726 | Q15726 |
| Meteorin | METRN | | 0.411186 | 0.283671 | Q9UJH8;I3L2T3;J3KMW6;H3BUM1 | Q9UJH8;I3L2T3;J3KMW6 |
| Meteorin-like protein | METRNL | | 1.07101 | -2.65403 | Q641Q3;B3KSJ5;I3L0R0 | Q641Q3;B3KSJ5 |
| Methionine adenosyltransferase 2 subunit beta | MAT2B | | 0.143729 | -0.103611 | Q9NZL9;Q9NZL9-2;Q9NZL9-4;Q9NZL9-3;H7C0X7;E5RJR3;Q9NZL9-5 | Q9NZL9;Q9NZL9-2;Q9NZL9-4;Q9NZL9-3;H7C0X7 |
| Methionine--tRNA ligase, cytoplasmic | MARS | | 0.0114349 | 0.024198 | P56192;A6NC17;B3KVK7;H0YHV5;H0YIP0;F8VS26;F8W0M7;F8W0S4;F8 | P56192;A6NC17;B3KVK7 |
| MHC class I polypeptide-related sequence A | MICA | | 0.801604 | 0.657901 | Q96QC4;Q5SS58;Q29983;H0Y615 | Q96QC4;Q5SS58;Q29983 |
| Microfibrillar-associated protein 2 | MFAP2 | | 0.150028 | -0.123652 | P55001-3;P55001-2;P55001 | P55001-3;P55001-2;P55001 |
| Microtubule-associated protein;Microtubule-associated protein 4 | MAP4 | | 1.11938 | 1.54659 | E7EVA0;P27816;P27816-6;P27816-2;B9ZVR1;E9PGM5;P27816-5;P2781 | E7EVA0;P27816;P27816-6;P27816-2;B9ZVR1;E9PGM5;P27816-5;P2781 |
| Midkine | MDK | | 0.548064 | -1.516 | E9PPJ5;E9PLM6;P21741-2;P21741 | E9PPJ5;E9PLM6;P21741-2;P21741 |
| MIT domain-containing protein 1 | MITD1 | | 0.693616 | -0.819548 | H7C3Q6;Q8WV92;B8ZZL5;F8WED5 | H7C3Q6;Q8WV92;B8ZZL5 |
| Mitochondrial import inner membrane translocase subunit Tim8 B | TIMM8B | | 0.157243 | 0.143265 | Q9Y5J9;G3XAN8;E9PIR3 | Q9Y5J9;G3XAN8 |
| Mitochondrial peptide methionine sulfoxide reductase | MSRA | | 0.119678 | -0.0957097 | Q9UJ68-2;Q9UJ68-3;Q9UJ68-4;Q9UJ68-5;Q9UJ68;H0YAN3;E5RIA9;E5R. | Q9UJ68-2;Q9UJ68-3;Q9UJ68-4;Q9UJ68-5;Q9UJ68;H0YAN3;E5RIA9 |
| Mitogen-activated protein kinase 1 | MAPK1 | | 0.316203 | -0.165772 | P28482;P28482-2;B3KR49;H0YDH9;E9PBK7;D6RFF2;K7ELV1;K7EN18;Q! | P28482;P28482-2 |
| Mitogen-activated protein kinase 14 | MAPK14 | | 0.087852 | -0.103285 | Q16539;B4E0K5;Q16539-3;Q16539-4;B5TY33;E7EX54;H7C4E2;C9JWQ4 | Q16539;B4E0K5;Q16539-3;Q16539-4 |
| Mitogen-activated protein kinase kinase kinase kinase 4 | MAP4K4 | | 0.631709 | -0.427749 | C9J840;G3XAA2;E7ESS2;G5E948;E7EX83;H7C0P6;E7ENQ1;H7C360;E7EI | C9J840;G3XAA2;E7ESS2;G5E948;E7EX83;H7C0P6;E7ENQ1;H7C360;E7E |
| MOB kinase activator 1A;MOB kinase activator 1B | MOB1A;MOB1B | | 0.111864 | -0.0895617 | Q9H8S9;Q7L9L4;Q7L9L4-2;Q9H8S9-2;H7BZ12;D6RCK3 | Q9H8S9;Q7L9L4;Q7L9L4-2;Q9H8S9-2;H7BZ12 |
| Moesin | MSN | | 0.800741 | -0.468455 | P26038;V9GZ54 | P26038 |
| Molybdopterin synthase sulfur carrier subunit | MOCS2 | | 0.398572 | 0.359278 | E9PKT9;O96033 | E9PKT9;O96033 |
| Multifunctional methyltransferase subunit TRM112-like protein | TRMT112 | | 0.620465 | 0.830374 | Q9UI30;F5GX77;Q9UI30-2;F5GYQ2;M0R242 | Q9UI30;F5GX77;Q9UI30-2;F5GYQ2;M0R242 |
| Multifunctional protein ADE2;Phosphoribosylaminoimidazole-succinocarboxamide synthase;Phosphoribc | PAICS | | 0.685817 | -0.6351 | P22234;E9PBS1;P22234-2;D6RF62 | P22234;E9PBS1;P22234-2;D6RF62 |
| Multiple inositol polyphosphate phosphatase 1 | MINPP1 | | 1.22248 | -0.571723 | Q9UNW1;Q9UNW1-4;Q9UNW1-3;Q9UNW1-2 | Q9UNW1;Q9UNW1-4 |
| Multivesicular body subunit 12A | MVB12A | | 0.238429 | 0.13402 | Q96EY5;E9PQA6;Q96EY5-3;E9PLL0;Q96EY5-2;M0QZK6;E9PLZ8;E9PPW: | Q96EY5;E9PQA6;Q96EY5-3;E9PLL0;Q96EY5-2 |
| Myeloid-associated differentiation marker | MYADM | | 0.842354 | -0.786737 | C9JJV6;C9JZL8;C9J5M0;C9JC07;Q96S97 | C9JJV6;C9JZL8;C9J5M0;C9JC07;Q96S97 |
| Myoferlin | MYOF | | 0.591658 | -0.467707 | F8W8J4;Q9NZM1-6;Q9NZM1;Q9NZM1-2;Q9NZM1-5 | F8W8J4;Q9NZM1-6;Q9NZM1;Q9NZM1-2;Q9NZM1-5 |
| Myosin light polypeptide 6 | MYL6;PDE6H | | 0.80486 | 0.704552 | P60660-2;F8W1R7;G3V1V0;P60660;G8JLA2;J3KND3;B7Z6Z4;G3V1Y7;F8 | P60660-2;F8W1R7;G3V1V0;P60660;G8JLA2;J3KND3;B7Z6Z4;G3V1Y7;F8 |
| Myosin regulatory light chain 12A;Myosin regulatory light chain 12B | MYL12A;MYL12B | | 0.608388 | 1.23941 | J3QRS3;P19105;O14950;J3KTJ1;P24844-2 | J3QRS3;P19105;O14950 |
| Myosin-9 | MYH9 | | 0.335626 | 0.263016 | P35579;P35579-2;F8W6L6;P35580;P35580-2;P35580-3;P35580-4;Q5BK | P35579;P35579-2 |
| Myotrophin | MTPN | | 0.663228 | 0.929555 | P58546;C9JL85 | P58546;C9JL85 |
| Myristoylated alanine-rich C-kinase substrate | MARCKS | | 0.459562 | 0.749095 | P29966 | P29966 |
| N(4)-(beta-N-acetylglucosaminyl)-L-asparaginase;Glycosylasparaginase alpha chain;Glycosylasparaginase | AGA | | 0.21367 | 0.240074 | P20933;H0Y9C7;H0Y8L9 | P20933;H0Y9C7 |
| N(G),N(G)-dimethylarginine dimethylaminohydrolase 1 | DDAH1 | | 0.03622 | 0.0247791 | O94760;B4DYP1;B4DGT0;O94760-2 | O94760;B4DYP1;B4DGT0;O94760-2 |
| N(G),N(G)-dimethylarginine dimethylaminohydrolase 2 | DDAH2 | | 0.0951978 | 0.0721047 | O95865;Q5SSV3;Q5SRR8;H0Y7N1 | O95865;Q5SSV3;Q5SRR8;H0Y7N1 |
| Na(+)/H(+) exchange regulatory cofactor NHE-RF1 | SLC9A3R1 | | 0.334012 | -0.247471 | O14745;J3QRP6;B3KY21;J3QRA3 | O14745;J3QRP6;B3KY21 |
| Na(+)/H(+) exchange regulatory cofactor NHE-RF2 | SLC9A3R2 | | 0.720047 | 0.921111 | Q15599-3;H3BUQ9;H3BN50;H3BQS0 | Q15599-3;H3BUQ9;H3BN50;H3BQS0 |
| N-acetyl-D-glucosamine kinase | NAGK | | 0.316039 | 0.380126 | Q9UJ70;H7C3G9;Q9UJ70-2;C9JEV6;H7C286;H7C1L7;H0YF44;H0YEB7;H( | Q9UJ70;H7C3G9;Q9UJ70-2;C9JEV6;H7C286 |
| N-acetylgalactosamine-6-sulfatase | GALNS | | 0.978957 | 0.540545 | P34059;F5H325;H3BP66 | P34059;F5H325 |
| N-acetylgalactosaminyltransferase 7 | GALNT7 | | 0.361305 | 0.236078 | Q86SF2;H0YAH3;H0YAK0;E9PBY3 | Q86SF2;H0YAH3 |
| N-acetylglucosamine-1-phosphotransferase subunit gamma | GNPTG | | 0.702738 | 0.0565188 | Q9UJJ9;E9PQQ5;H0YEA7 | Q9UJJ9 |
| N-acetylglucosamine-6-sulfatase | GNS | | 1.11476 | 0.402354 | P15586;H7C3P4;F6S8M0;B4DYH8;H0YFA9;F5H4C6 | P15586;H7C3P4;F6S8M0;B4DYH8;H0YFA9;F5H4C6 |
| N-acetyllactosaminide beta-1,3-N-acetylglucosaminyltransferase | B3GNT1 | | 0.971379 | -1.22414 | O43505 | O43505 |
| NAD(P)H-hydrate epimerase | APOA1BP | | 0.384878 | 0.21305 | Q8NCW5;Q8NCW5-2;Q5T3I4;Q5T3I3 | Q8NCW5;Q8NCW5-2 |
| NADH-cytochrome b5 reductase 3;NADH-cytochrome b5 reductase 3 membrane-bound form;NADH-cyto | CYB5R3 | | 0.384942 | -0.530994 | P00387-2;P00387;P00387-3;B1AHF3 | P00387-2;P00387;P00387-3;B1AHF3 |
| N-alpha-acetyltransferase 15, NatA auxiliary subunit | NAA15 | | 0.326296 | 0.472185 | Q9BXJ9;Q9BXJ9-4;Q6N069-5;Q6N069-4;Q6N069-3;Q6N069-2;Q6N069 | Q9BXJ9 |
| N-alpha-acetyltransferase 50 | NAA50 | | 0.435878 | 0.489597 | E7EQ69;Q9GZZ1;C9J5D1;Q9GZZ1-2;B0AZT5;C9J5J3;C9JZU6;F8WCK0 | E7EQ69;Q9GZZ1;C9J5D1;Q9GZZ1-2 |
| Nascent polypeptide-associated complex subunit alpha;Nascent polypeptide-associated complex subunit | NACA | | 0.358704 | 0.333312 | F8VZJ2;H0YHX9;Q13765;E9PAV3-2;E9PAV3;F8W0W4;F8W1N5;F8VNW4 | F8VZJ2;H0YHX9;Q13765;E9PAV3-2;E9PAV3;F8W0W4;F8W1N5;F8VNW |
| Nck-associated protein 1 | NCKAP1 | | 0.2606 | -0.190861 | Q9Y2A7;Q9Y2A7-2;F8W050;P55160 | Q9Y2A7;Q9Y2A7-2 |
| Nectin-2 | PVRL2 | | 0.848997 | -0.551121 | Q92692-2;Q92692;K7EKE8 | Q92692-2;Q92692;K7EKE8 |
| NEDD4 family-interacting protein 1 | NDFIP1 | | 0.416678 | -0.505458 | Q9BT67-2;Q9BT67 | Q9BT67-2;Q9BT67 |
| NEDD8-conjugating enzyme Ubc12 | UBE2M | | 0.0984678 | -0.0755428 | P61081;M0QX69;M0QYI6 | P61081 |
| Netrin-4 | NTN4 | | 0.251906 | -0.306802 | Q9HB63-3;Q9HB63-2;Q9HB63;F8W0I7 | Q9HB63-3;Q9HB63-2;Q9HB63 |
| Neudesin | NENF | | 0.0188774 | -0.0135175 | Q9UMX5 | Q9UMX5 |
| Neuroblast differentiation-associated protein AHNAK | AHNAK | | 0.985066 | 0.728373 | Q09666;E9PQE3;E9PJZ0;E9PKR9;E9PJC6;E9PLK4;Q09666-2 | Q09666 |
| Neurogenic locus notch homolog protein 2;Notch 2 extracellular truncation;Notch 2 intracellular domain | NOTCH2 | | 0.50247 | -0.314179 | Q04721;R4GN02;E9PLD5;Q7Z3S9-2;Q7Z3S9;Q7Z3S9-3 | Q04721 |
| Neuronal growth regulator 1 | NEGR1 | | 0.50846 | 0.241342 | Q7Z3B1;B4DI94;Q8N440 | Q7Z3B1;B4DI94;Q8N440 |
| Neuropilin-1 | NRP1 | | 0.260857 | -0.117848 | E9PEP6;O14786;E7EX60;Q5T7F0;Q5JWQ6;O14786-3;O14786-2;Q5JWQ | E9PEP6;O14786;E7EX60;Q5T7F0;Q5JWQ6;O14786-3;O14786-2;Q5JWC |
| Neuropilin-2 | NRP2 | + | 1.4797 | 0.440686 | F5H2X3;F5H1G4;O60462-4;O60462-5;O60462-2;O60462-3;O60462;O60 | F5H2X3;F5H1G4;O60462-4;O60462-5;O60462-2;O60462-3;O60462;O6( |
| Neuroserpin | SERPINI1 | | 0.11954 | 0.0749874 | Q99574;H7C5T9 | Q99574;H7C5T9 |
| Neutral alpha-glucosidase AB | GANAB | | 0.186576 | -0.102614 | Q14697;Q14697-2;F5H6X6;E9PKU7;E9PNH1;Q14697-3 | Q14697;Q14697-2;F5H6X6;E9PKU7 |
| Neutral amino acid transporter B(0) | SLC1A5 | | 0.351147 | -0.448191 | Q15758;M0QXM4;Q15758-3;Q15758-2;M0QX44;M0R144 | Q15758;M0QXM4;Q15758-3;Q15758-2 |
| NHL repeat-containing protein 3 | NHLRC3 | | 0.00594615 | 0.00537947 | C9J973;Q5JS37-2;Q5JS37 | C9J973;Q5JS37-2;Q5JS37 |
| Niban-like protein 1 | FAM129B | | 0.0708119 | -0.0780747 | Q96TA1-2;Q96TA1 | Q96TA1-2;Q96TA1 |

| | | | | | | |
|---|---|---|---|---|---|---|
| Nicastrin | NCSTN | | 0.147875 | -0.108481 | Q92542-2;Q92542;E7ENA9;F6Y097;H0Y6T7;Q5T205;H0Y3Z4;Q5T209;Q | Q92542-2;Q92542;E7ENA9;F6Y097;H0Y6T7;Q5T205;H0Y3Z4 |
| Nicotinamide phosphoribosyltransferase | NAMPT;NAMPTL | | 0.0807233 | 0.0870588 | P43490;Q5SYT8;C9JG65;F5H246;C9JF35 | P43490;Q5SYT8 |
| Nicotinate phosphoribosyltransferase | NAPRT | | 0.72419 | 0.557411 | Q6XQN6;Q6XQN6-3;C9JC60;G5E977;Q6XQN6-2;C9J8U2;H0YF31;H0YDA | Q6XQN6;Q6XQN6-3;C9JC60;G5E977;Q6XQN6-2;C9J8U2 |
| Niemann-Pick C1 protein | NPC1 | | 0.690319 | -0.656117 | O15118;K7EQ23;B4DET3;Q9UHC9-3;Q9UHC9-4;Q9UHC9-2;Q9UHC9 | O15118;K7EQ23;B4DET3 |
| Nitrilase homolog 1 | NIT1 | | 0.0865012 | -0.0888128 | Q86X76-2;Q86X76-4;Q86X76;Q86X76-3;Q86X76-5 | Q86X76-2;Q86X76-4;Q86X76;Q86X76-3;Q86X76-5 |
| NKG2D ligand 2 | ULBP2 | | 1.08411 | 0.466074 | Q9BZM5;C9JAK3;Q6H3X3-2;Q5VY80;Q6H3X3 | Q9BZM5 |
| Noggin | NOG | + | 1.55037 | 1.18833 | Q13253 | Q13253 |
| Non-specific lipid-transfer protein | SCP2 | | 1.16965 | 1.61084 | P22307-4;P22307-7;P22307-8;P22307;P22307-6;P22307-2;H0YF61;H0Y | P22307-4;P22307-7;P22307-8;P22307;P22307-6;P22307-2 |
| N-sulphoglucosamine sulphohydrolase | SGSH | | 0.136723 | 0.241869 | P51688;F5H6A3;I3NI22;I3L3T3;I3L2I6;I3L4B7;I3L2L4 | P51688;F5H6A3 |
| Nuclear mitotic apparatus protein 1 | NUMA1 | | 1.2755 | -1.21218 | Q14980-4;Q14980-3;Q14980-2;Q14980;F5H4J1;F5H6Y5;H0YFY6;Q1498 | Q14980-4;Q14980-3;Q14980-2;Q14980 |
| Nuclear receptor 2C2-associated protein | NR2C2AP | | 0.312887 | 0.339963 | Q86WQ0;B4DW92;U3KQ53;F5H0B6 | Q86WQ0;B4DW92;U3KQ53 |
| Nuclear transport factor 2 | NUTF2 | | 0.257551 | 0.163681 | P61970 | P61970 |
| Nuclease-sensitive element-binding protein 1 | YBX1 | | 0.0173774 | -0.0171681 | P67809;H0Y449;C9J5V9;Q9Y2T7 | P67809;H0Y449 |
| Nucleobindin-1 | NUCB1 | | 1.26699 | 0.999537 | Q02818;H7BZI1;C9JKZ2;C9JBD3;C9J3C1 | Q02818;H7BZI1 |
| Nucleobindin-2;Nesfatin-1 | NUCB2;DCAF5 | | 0.991823 | 3.07739 | P80303-2;P80303;E9PKG6;J3KQU0;E9PLR0;E9PLE9;H0YEG8;E9PM22;E9 | P80303-2;P80303;E9PKG6;J3KQU0;E9PLR0;E9PLE9 |
| Nucleolin | NCL | | 0.109735 | 0.0742949 | P19338;H7BY16;C9JYW2;C9JLB1;C9J1H7;C9JWL1 | P19338;H7BY16 |
| Nucleolysin TIAR | TIAL1 | | 0.482698 | 0.515379 | Q01085-2;Q01085;E7ETJ9;E7ETC0;A6NKZ9;Q49AS9;F8WE16 | Q01085-2;Q01085;E7ETJ9;E7ETC0 |
| Nucleoside diphosphate kinase 3;Nucleoside diphosphate kinase | NME3 | + | 1.57793 | -0.470989 | Q13232;H3BPR2;H3BPD9;H3BMQ7;H3BRA8 | Q13232;H3BPR2 |
| Nucleoside diphosphate kinase A | NME1 | + | 1.66775 | -0.635027 | P15531;P15531-2;E7ERL0;E5RHP0;C9K028 | P15531;P15531-2;E7ERL0;E5RHP0 |
| Nucleoside diphosphate kinase;Nucleoside diphosphate kinase B | NME1-NME2;NME2 | + | 1.60047 | -0.760929 | Q32Q12;P22392-2;J3KPD9;P22392;O60361 | Q32Q12;P22392-2;J3KPD9;P22392 |
| Nucleotide exchange factor SIL1 | SIL1 | | 1.00687 | 1.94424 | D6REA1;Q9H173;D6RIU8;D6RG16;D6RBP7;D6RAI3;A0RZB6 | D6REA1;Q9H173 |
| NudC domain-containing protein 2 | NUDCD2 | | 0.782553 | 0.992901 | Q8WVJ2;E5RFP0 | Q8WVJ2;E5RFP0 |
| Obg-like ATPase 1 | OLA1 | | 0.107024 | 0.124879 | J3KQ32;Q9NTK5;Q9NTK5-2;Q9NTK5-3;C9JTK6;C9JCJ9 | J3KQ32;Q9NTK5;Q9NTK5-2;Q9NTK5-3 |
| Olfactomedin-like protein 2A | OLFML2A | + | 1.3847 | -0.941487 | Q5JTM7;Q68BL7-2;Q68BL7;Q68BL7-3 | Q5JTM7;Q68BL7-2;Q68BL7;Q68BL7-3 |
| Omega-amidase NIT2 | NIT2 | | 0.326406 | 0.310805 | Q9NQR4;H7C579;F8WF70 | Q9NQR4;H7C579 |
| Osteoclast-stimulating factor 1 | OSTF1 | | 0.108964 | 0.128416 | Q92882 | Q92882 |
| Out at first protein homolog | OAF | | 0.470605 | 0.100415 | Q86UD1;E9PJ29 | Q86UD1 |
| Palmitoyl-protein thioesterase 1 | PPT1 | | 0.244678 | -0.20412 | P50897;E9PMG2;E9PIA8;P50897-2;Q5T0S4;E9PK48;E9PP28;E9PSE5 | P50897;E9PMG2;E9PIA8;P50897-2;Q5T0S4 |
| Parkinson disease 7 domain-containing protein 1 | PDDC1 | | 0.0914229 | -0.114834 | Q8NB37;B7Z1J9;Q8NB37-2;H0YF25;Q8NB37-3;E9PQD8;H0YE25;E9PLI5; | Q8NB37;B7Z1J9;Q8NB37-2;H0YF25;Q8NB37-3;E9PQD8;H0YE25;E9PLI5 |
| PDZ domain-containing protein GIPC1 | GIPC1 | | 0.149264 | -0.147566 | O14908;K7EM11;O14908-2;K7ELJ2;K7ESN1 | O14908;K7EM11;O14908-2 |
| Peflin | PEF1 | | 0.0584545 | -0.0400275 | Q9UBV8 | Q9UBV8 |
| Pentraxin-related protein PTX3 | PTX3 | + | 1.63877 | -1.4445 | P26022;R4GNA2;Q5W026;Q9BXX3 | P26022 |
| Peptidase M20 domain-containing protein 2 | PM20D2 | | 0.539861 | 0.428456 | Q8IYS1 | Q8IYS1 |
| Peptidyl-glycine alpha-amidating monooxygenase;Peptidylglycine alpha-hydroxylating monooxygenase;Pe | PAM | | 0.286984 | -0.235438 | P19021-4;P19021-3;P19021;P19021-5;P19021-6;F8WE90;P19021-2;A6N | P19021-4;P19021-3;P19021;P19021-5;P19021-6;F8WE90;P19021-2 |
| Peptidyl-prolyl cis-trans isomerase A;Peptidyl-prolyl cis-trans isomerase A, N-terminally processed;Peptid | PPIA | | 0.399487 | 0.319378 | P62937;F8WE65;C9J5S7;E5RIZ5;Q9Y536;F5H284 | P62937;F8WE65;C9J5S7 |
| Peptidyl-prolyl cis-trans isomerase B | PPIB | | 0.24719 | -0.192069 | P23284 | P23284 |
| Peptidyl-prolyl cis-trans isomerase C | PPIC | | 0.0578154 | -0.0670534 | P45877 | P45877 |
| Peptidyl-prolyl cis-trans isomerase FKBP1A;Peptidyl-prolyl cis-trans isomerase | FKBP1A | | 0.185876 | -0.142904 | P62942;Q5W0X3;Q1JUQ5;A8MSS1;Q5VVH2 | P62942;Q5W0X3 |
| Peptidyl-prolyl cis-trans isomerase FKBP2;Peptidyl-prolyl cis-trans isomerase | FKBP2 | | 0.348026 | 0.204838 | P26885;F5H0N4 | P26885;F5H0N4 |
| Peptidyl-prolyl cis-trans isomerase FKBP4;Peptidyl-prolyl cis-trans isomerase FKBP4, N-terminally process | FKBP4 | | 0.727192 | -0.531787 | Q02790;F5H1U3;H0YFG2;F5H120 | Q02790 |
| Peptidyl-prolyl cis-trans isomerase FKBP5 | FKBP5 | | 0.475579 | 0.472408 | Q13451;B7Z7Z8;Q13451-2 | Q13451;B7Z7Z8 |
| Peptidyl-prolyl cis-trans isomerase NIMA-interacting 4 | PIN4 | | 0.355885 | -0.34698 | Q9Y237;Q9Y237-2;H0Y4T6;J3KMW3;H0Y8P6;H0Y610;Q9Y237-3 | Q9Y237;Q9Y237-2;H0Y4T6;J3KMW3;H0Y8P6;H0Y610;Q9Y237-3 |
| Peptidyl-prolyl cis-trans isomerase-like 1 | PPIL1 | | 0.156771 | -0.130281 | Q9Y3C6 | Q9Y3C6 |
| Peroxidasin homolog | PXDN | + | 1.33129 | -0.634452 | Q92626;H7C1W1;Q92626-2;H7C300;H7C3W2 | Q92626;H7C1W1;Q92626-2 |
| Peroxiredoxin-1 | PRDX1 | + | 1.32401 | -0.568005 | Q06830 | Q06830 |
| Peroxiredoxin-2 | PRDX2 | | 0.0548703 | 0.0452942 | P32119;A6NIW5;P32119-2 | P32119;A6NIW5 |
| Peroxiredoxin-4 | PRDX4 | | 0.784533 | -0.383505 | Q13162;H7C3T4;A6NG45;A6NJJ0 | Q13162;H7C3T4 |
| Peroxiredoxin-5, mitochondrial | PRDX5 | | 1.26736 | 0.958068 | P30044-2;P30044;P30044-3;P30044-4 | P30044-2;P30044;P30044-3;P30044-4 |
| Peroxiredoxin-6 | PRDX6 | | 0.0591791 | 0.0469758 | P30041 | P30041 |
| Phosphate carrier protein, mitochondrial | SLC25A3 | | 0.728522 | -1.15734 | F8VVM2;Q00325-2;Q00325;F8VWQ0;F8VZL5;F8VWR4 | F8VVM2;Q00325-2;Q00325 |
| Phosphatidylethanolamine-binding protein 1;Hippocampal cholinergic neurostimulating peptide | PEBP1 | | 1.2053 | -0.476022 | P30086 | P30086 |
| Phosphatidylinositol transfer protein alpha isoform | PITPNA | | 0.00264295 | 0.00324797 | F5GWE5;Q00169;I3L459;I3L4H1;I3L4C0;I3NI10;I3L2X8;I3L471;I3L4U7;I3 | F5GWE5;Q00169;I3L459;I3L4H1;I3L4C0 |
| Phosphatidylinositol transfer protein beta isoform | PITPNB | | 0.0924521 | 0.0857678 | P48739;B7Z7Q0;P48739-2;B3KYB7;B3KYB6 | P48739;B7Z7Q0;P48739-2 |
| Phosphoenolpyruvate carboxykinase [GTP], mitochondrial | PCK2 | + | 2.19303 | -0.349025 | Q16822;B4DW73;H0YML5;H0YM31;Q16822-2;H0YMU6;H0YNG4;H0YM | Q16822;B4DW73;H0YML5;H0YM31 |
| Phosphoglucomutase-1 | PGM1 | | 0.244655 | -0.184276 | P36871;P36871-2;P36871-3 | P36871;P36871-2;P36871-3 |
| Phosphoglucomutase-2 | PGM2 | | 0.0185776 | -0.0127003 | Q96G03;F5H6V2;E7ENQ8;E9PD70 | Q96G03;F5H6V2 |
| Phosphoglycerate kinase 1 | PGK1 | | 0.737348 | -0.555247 | P00558;B7Z7A9;E7ERH5 | P00558;B7Z7A9;E7ERH5 |
| Phosphoglycerate mutase 1 | PGAM1 | | 0.197977 | -0.148758 | P18669;P15259;Q8N0Y7 | P18669 |
| Phospholipase A-2-activating protein | PLAA | | 0.173893 | -0.21754 | Q9Y263;E5RIM3;H0YC16;H0YAU9;H0YBW4 | Q9Y263;E5RIM3 |
| Phospholipase D3 | PLD3 | | 0.52343 | -0.278398 | Q8IV08;M0QZI4;M0R3G9;M0R2W7;E2QRG1;M0R1F7 | Q8IV08 |
| Phospholipid hydroperoxide glutathione peroxidase, mitochondrial;Glutathione peroxidase | GPX4 | | 0.853828 | 1.15775 | P36969-2;P36969;K7ERP4;K7ENB4;K7EKX7;K7EJ20;R4GNE4 | P36969-2;P36969;K7ERP4;K7ENB4;K7EKX7;K7EJ20 |
| Phosphomannomutase 2;Phosphomannomutase | PMM2 | | 0.162568 | 0.212165 | O15305;H3BV55;H3BRM0;F5H0W0;H3BPH4;B7Z419;B7Z6R0;H3BR08;H | O15305;H3BV55;H3BRM0;F5H0W0;H3BPH4 |
| Phosphomevalonate kinase | PMVK | | 0.757879 | 0.675325 | Q15126 | Q15126 |
| Phosphopantothenate--cysteine ligase | PPCS | | 0.79349 | -0.536114 | Q9HAB8;Q9HAB8-2;Q5VVM3 | Q9HAB8;Q9HAB8-2 |
| Phosphoribosylformylglycinamidine synthase | PFAS | | 0.0384188 | -0.0284976 | O15067;F5GWT9;H0YGH1;J3QSG0;J3QSH6;J3KTL4;J3QL39;J3KT98;J3KT | O15067;F5GWT9 |
| Phosphoserine aminotransferase | PSAT1 | | 0.279021 | -0.210641 | Q9Y617;Q9Y617-2 | Q9Y617;Q9Y617-2 |
| Phosphotriesterase-related protein | PTER | | 0.368398 | 0.358822 | Q96BW5-2;Q96BW5 | Q96BW5-2;Q96BW5 |
| Pituitary tumor-transforming gene 1 protein-interacting protein | PTTG1IP | | 0.273521 | -0.178469 | A8MXQ1;P53801;A8MZH8 | A8MXQ1;P53801;A8MZH8 |
| Plasma alpha-L-fucosidase | FUCA2 | | 0.925452 | 0.354012 | Q9BTY2;Q7Z6V1;Q7Z6V2 | Q9BTY2 |
| Plasminogen activator inhibitor 1 | SERPINE1 | | 0.201155 | -0.129432 | P05121;P05121-2 | P05121;P05121-2 |
| Plasminogen activator inhibitor 1 RNA-binding protein | SERBP1 | + | 1.37087 | 3.07636 | Q8NC51;Q8NC51-3;Q8NC51-2;Q8NC51-4 | Q8NC51;Q8NC51-3;Q8NC51-2;Q8NC51-4 |
| Plasminogen activator inhibitor 2 | SERPINB2 | | 0.0224844 | -0.063235 | P05120;E7EPJ9;E7ERB5;E9PDK7 | P05120 |
| Plastin-1 | PLS1 | | 0.17278 | 0.131942 | Q14651;C9JAM8 | Q14651;C9JAM8 |
| Platelet-activating factor acetylhydrolase IB subunit alpha | PAFAH1B1 | | 0.478626 | 0.462566 | P43034;B4DF38;I3L3N5;I3L495;P43034-2 | P43034 |
| Platelet-activating factor acetylhydrolase IB subunit beta | PAFAH1B2 | | 0.296329 | 0.361799 | P68402;P68402-3;P68402-2;P68402-4;J3KNE3 | P68402;P68402-3;P68402-2;P68402-4 |
| Platelet-activating factor acetylhydrolase IB subunit gamma | PAFAH1B3 | | 0.715306 | -0.496994 | Q15102;M0R389;M0R1K3;M0QZT2;M0QXS6;M0R323 | Q15102;M0R389;M0R1K3 |
| Platelet-derived growth factor C;Platelet-derived growth factor C, latent form;Platelet-derived growth fac | PDGFC | | 0.162699 | -0.154648 | Q9NRA1;Q9NRA1-2;B4E3A5;Q9NRA1-4;Q9NRA1-3;H0Y9N2 | Q9NRA1;Q9NRA1-2;B4E3A5;Q9NRA1-4 |
| Platelet-derived growth factor D;Platelet-derived growth factor D, latent form;Platelet-derived growth fac | PDGFD | | 0.39199 | 0.332716 | Q9GZP0-2;Q9GZP0;H0YD37 | Q9GZP0-2;Q9GZP0 |
| Platelet-derived growth factor subunit A | PDGFA | + | 1.42876 | -1.21462 | H7BYW6;P04085-2;P04085 | H7BYW6;P04085-2;P04085 |

| | | | | | | |
|---|---|---|---|---|---|---|
| Plectin | PLEC | | 0.225542 | -0.244888 | Q15149-4 | Q15149-4 |
| Plexin-B2 | PLXNB2 | | 0.203809 | -0.191722 | O15031;E2PU09;A6QRH1;H0Y7X5;H0Y6J7 | O15031 |
| Podocalyxin | PODXL | | 0.374499 | -0.238396 | F5GWY5;O00592-2;O00592;F5H7A2 | F5GWY5;O00592-2;O00592;F5H7A2 |
| Poliovirus receptor | PVR | | 0.220568 | -0.165546 | P15151-2;P15151-3;P15151-4;P15151 | P15151-2;P15151-3;P15151-4;P15151 |
| Poly(rC)-binding protein 1 | PCBP1 | | 0.18405 | -0.169682 | Q15365;F8VTZ0;H3BSP4;C9K0A2;F8WC71;C9IZV9;C9J7A9;C9JSA6;C9J5 | Q15365 |
| Poly(rC)-binding protein 2;Poly(rC)-binding protein 3 | PCBP2;PCBP3 | | 0.346794 | -0.31324 | Q15366-6;Q15366-3;B4DLC0;Q15366;Q15366-2;Q15366-7;Q15366-4;E | Q15366-6;Q15366-3;B4DLC0;Q15366;Q15366-2;Q15366-7;Q15366-4;E |
| Polyadenylate-binding protein 1 | PABPC1 | | 0.0961818 | 0.102032 | P11940;E7EQV3;E7ERJ7;P11940-2;H0YAR2;H0YAP2;H0YBN4;E5RGH3;E | P11940;E7EQV3;E7ERJ7;P11940-2;H0YAR2 |
| Polypeptide N-acetylgalactosaminyltransferase 2;Polypeptide N-acetylgalactosaminyltransferase 2 solubl | GALNT2 | | 0.0131268 | -0.0257087 | G3V1S6;Q10471;B7Z8V8 | G3V1S6;Q10471;B7Z8V8 |
| Polypyrimidine tract-binding protein 1 | PTBP1 | | 0.0274974 | -0.0326734 | P26599;P26599-2;P26599-3;K7ES59;A6NLN1;K7EKJ7;K7ELW5;K7EK45;C | P26599;P26599-2;P26599-3 |
| POTE ankyrin domain family member E;POTE ankyrin domain family member F | POTEE;POTEF | | 1.11175 | 0.633545 | Q6S8J3;A5A3E0;Q9BYX7;Q495V5;F8W967;D7UEQ8;A6NC16;F8W858;Q | Q6S8J3;A5A3E0 |
| Prefoldin subunit 2 | PFDN2 | | 0.88956 | 1.59112 | Q9UHV9 | Q9UHV9 |
| Probable serine carboxypeptidase CPVL | CPVL | | 1.00754 | 1.69794 | Q9H3G5;H7C0X5;H7C218;C9JLV0;C9J6L4;C9JI22 | Q9H3G5;H7C0X5 |
| Probable Xaa-Pro aminopeptidase 3 | XPNPEP3 | | 0.872455 | 0.44911 | Q9NQH7-2;Q9NQH7-4;Q9NQH7;Q9NQH7-3;Q9NQH7-5 | Q9NQH7-2;Q9NQH7-4;Q9NQH7;Q9NQH7-3;Q9NQH7-5 |
| Pro-cathepsin H;Cathepsin H mini chain;Cathepsin H;Cathepsin H heavy chain;Cathepsin H light chain | CTSH | | 0.856902 | -0.671398 | P09668;E9PKT6 | P09668 |
| Procollagen C-endopeptidase enhancer 2 | PCOLCE2 | | 0.448668 | 0.147838 | Q9UKZ9;C9JYX9;H7C5D5 | Q9UKZ9;C9JYX9 |
| Procollagen-lysine,2-oxoglutarate 5-dioxygenase 1 | PLOD1 | | 1.12305 | -0.474789 | Q02809;B4DR87;Q5JXB7;Q5JXB9;Q5JXB8;M0QZS0 | Q02809;B4DR87 |
| Procollagen-lysine,2-oxoglutarate 5-dioxygenase 2 | PLOD2 | | 0.350554 | -0.140943 | O00469-2;E7ETU9;O00469;B3KWS3;C9JXZ0;F8WEW3 | O00469-2;E7ETU9;O00469;B3KWS3 |
| Procollagen-lysine,2-oxoglutarate 5-dioxygenase 3 | PLOD3 | | 0.768211 | 0.607806 | O60568;H7C2V1;H7C2S8;C9JU11;H7C0B8;H7C2A8;C9JIX5 | O60568 |
| Profilin;Profilin-2 | PFN2 | | 0.605532 | -0.280906 | C9J0J7;G5E9Q6;P35080-2 | C9J0J7;G5E9Q6;P35080-2 |
| Profilin-1 | PFN1 | | 0.333963 | -0.210518 | P07737;K7EJ44;I3L3D5 | P07737;K7EJ44 |
| Programmed cell death 1 ligand 1 | CD274 | | 0.952445 | 0.730705 | Q9NZQ7-2;Q9NZQ7-3;Q9NZQ7 | Q9NZQ7-2;Q9NZQ7-3;Q9NZQ7 |
| Programmed cell death 6-interacting protein | PDCD6IP | | 0.546386 | -0.273949 | Q8WUM4;Q8WUM4-2;C9IZF9;F8WDK9;F8WEQ7;F8WBR8 | Q8WUM4;Q8WUM4-2 |
| Programmed cell death protein 10 | PDCD10 | | 1.00877 | 0.786048 | Q9BUL8;C9J6F3;C9J5C3;H7C5M9;C9JSA3;C9JND6;C9J363;C9J932;F8WD | Q9BUL8;C9J6F3;C9J5C3;H7C5M9;C9JSA3;C9JND6;C9J363 |
| Programmed cell death protein 5 | PDCD5 | | 0.0266286 | -0.0197342 | O14737;B4DE64;K7EQA1;K7ESJ4;Q3HM38 | O14737;B4DE64;K7EQA1;K7ESJ4 |
| Programmed cell death protein 6 | PDCD6 | | 0.971757 | -0.667236 | O75340;O75340-2;E7EPW6;H0Y9X3;O75340-3;D6RA21;Q86W51 | O75340;O75340-2 |
| Prohibitin | PHB | | 0.486015 | -1.18737 | P35232;C9JW96;C9JZ20;E7ESE2;E9PCW0;B4DY47;D6RBK0 | P35232;C9JW96;C9JZ20;E7ESE2;E9PCW0 |
| Prohibitin-2 | PHB2 | | 0.977597 | -1.36923 | J3KPX7;Q99623;F5GY37;Q99623-2;F5H3X6;F5GWA7;F5H0C5;F5H2D2;U | J3KPX7;Q99623;F5GY37;Q99623-2;F5H3X6;F5GWA7 |
| Proliferating cell nuclear antigen | PCNA | | 0.615514 | 0.784182 | P12004 | P12004 |
| Proliferation-associated protein 2G4 | PA2G4 | | 0.280317 | 0.311173 | Q9UQ80;F8VR77;F8W0A3;H0YIN7;F8VZ69 | Q9UQ80;F8VR77 |
| Proline synthase co-transcribed bacterial homolog protein | PROSC | | 0.540938 | 0.544423 | O94903;E5RFX7;E5RG77;H0YBG2;E5RFZ4 | O94903;E5RFX7;E5RG77 |
| Prolyl 3-hydroxylase 1 | LEPRE1 | | 0.338687 | 0.273755 | Q32P28;Q32P28-3;H7C0D1;H7C2W6;Q32P28-2;E2QRI1 | Q32P28;Q32P28-3 |
| Pro-neuregulin-1, membrane-bound isoform;Neuregulin-1 | NRG1 | | 0.76568 | 0.461941 | E7EX30;F8W9E3;H7BXY2;H0YBA3;Q02297-8;Q02297-4;Q02297-12;Q02 | E7EX30;F8W9E3;H7BXY2;H0YBA3;Q02297-8;Q02297-4;Q02297-12;Q0 |
| Proprotein convertase subtilisin/kexin type 9 | PCSK9 | + | 2.52041 | 1.51719 | Q8NBP7;F5GWF0;Q8NBP7-2;B4DEZ9 | Q8NBP7 |
| Prosaposin;Saposin-A;Saposin-B-Val;Saposin-B;Saposin-C;Saposin-D | PSAP | | 0.68523 | 0.307679 | B1AVU8;P07602 | B1AVU8;P07602 |
| Prosaposin;Saposin-A;Saposin-B-Val;Saposin-B;Saposin-C;Saposin-D | PSAP | | 0.0359717 | 0.0277158 | P07602-3 | P07602-3 |
| Prostaglandin E synthase 3 | PTGES3 | + | 1.55614 | 1.68933 | B4DP11;Q15185;B4DP21;B4DDC6;B4DHP2 | B4DP11;Q15185;B4DP21;B4DDC6 |
| Prostaglandin reductase 2 | PTGR2 | | 0.259294 | 0.277838 | Q8N8N7;G3V3Y1;G3V288;G3V2R9;Q8N8N7-2 | Q8N8N7;G3V3Y1;G3V288;G3V2R9;Q8N8N7-2 |
| Protease-associated domain-containing protein 1 | PRADC1 | | 1.02987 | 0.664443 | Q9BSG0 | Q9BSG0 |
| Proteasome activator complex subunit 1 | PSME1 | | 0.364058 | 0.336314 | Q06323;H0YNE3;Q06323-2;H0YKK6;H0YLU2 | Q06323;H0YNE3;Q06323-2 |
| Proteasome activator complex subunit 2 | PSME2 | | 0.0591941 | -0.0556672 | Q9UL46;H0YM70;H0YKU2;H0YLG1 | Q9UL46;H0YM70 |
| Proteasome subunit alpha type-1;Proteasome subunit alpha type | PSMA1 | | 0.190318 | -0.133163 | P25786;P25786-2;F5GX11;B4DEV8;F5H112 | P25786;P25786-2;F5GX11;B4DEV8 |
| Proteasome subunit alpha type-2 | PSMA2 | | 0.222186 | -0.139788 | P25787;C9JCK5;Q6MZI6;H3BT36;H7C402 | P25787;C9JCK5 |
| Proteasome subunit alpha type-3 | PSMA3 | | 0.206767 | -0.144723 | P25788-2;P25788;G3V4X5;H0YJ03;G3V3W4;G3V5N4 | P25788-2;P25788;G3V4X5 |
| Proteasome subunit alpha type-4;Proteasome subunit alpha type;Proteasome subunit beta type | PSMA4 | | 0.0602283 | -0.0556895 | P25789;H0YN18;H0YMZ1;H0YL69;P25789-2;H0YMA1;H0YKT8;H0YMI6; | P25789;H0YN18;H0YMZ1;H0YL69;P25789-2;H0YMA1;H0YKT8;H0YMI6 |
| Proteasome subunit alpha type-5 | PSMA5 | | 0.179564 | -0.116654 | P28066;P28066-2 | P28066;P28066-2 |
| Proteasome subunit alpha type-6;Proteasome subunit alpha type | PSMA6 | | 0.143043 | -0.105965 | P60900;G3V295;G3V5Z7;P60900-2;G3V3I1;P60900-3;G3V3U4;H0YJC4; | P60900;G3V295;G3V5Z7;P60900-2;G3V3I1;P60900-3 |
| Proteasome subunit alpha type-7 | PSMA7 | | 0.236863 | -0.1688 | O14818;H0Y586;O14818-4;Q8TAA3-2;Q8TAA3-5;Q8TAA3;F5GY34 | O14818;H0Y586 |
| Proteasome subunit beta type;Proteasome subunit beta type-9 | PSMB9 | + | 2.01366 | -1.68251 | A2ACR1;P28065-2;P28065;A2ACR0 | A2ACR1;P28065-2;P28065 |
| Proteasome subunit beta type-1 | PSMB1 | | 0.15575 | -0.107346 | P20618 | P20618 |
| Proteasome subunit beta type-2 | PSMB2 | | 0.277006 | -0.170299 | P49721 | P49721 |
| Proteasome subunit beta type-3 | PSMB3 | | 0.222542 | -0.142226 | P49720;J3KSM3;J3KRR2;J3QKR3 | P49720;J3KSM3;J3KRR2;J3QKR3 |
| Proteasome subunit beta type-4 | PSMB4 | | 0.165883 | -0.116929 | P28070 | P28070 |
| Proteasome subunit beta type-5 | PSMB5 | | 0.22557 | -0.140919 | P28074;P28074-3;H0YJM8;P28074-2 | P28074;P28074-3 |
| Proteasome subunit beta type-6 | PSMB6 | | 0.233305 | -0.145594 | P28072;I3L3X7 | P28072 |
| Proteasome subunit beta type-7 | PSMB7 | | 0.338102 | -0.194157 | Q99436;Q5TBG5;B4E0P1 | Q99436 |
| Proteasome subunit beta type-8 | PSMB8 | | 0.160204 | -0.126353 | P28062-2;P28062;Q5JNW7 | P28062-2;P28062;Q5JNW7 |
| Protein archease | ZBTB8OS | | 0.768728 | 0.559818 | Q8IWT0;A8K0B5;Q8IWT0-2;H7C3R6 | Q8IWT0;A8K0B5;Q8IWT0-2 |
| Protein CREG1 | CREG1 | | 1.11898 | -0.817456 | O75629 | O75629 |
| Protein CutA | CUTA | | 0.566685 | -0.219254 | O60888-3;O60888;O60888-2 | O60888-3;O60888;O60888-2 |
| Protein CYR61 | CYR61 | | 0.579128 | -0.504389 | O00622 | O00622 |
| Protein disulfide-isomerase | P4HB | | 0.436662 | -0.253131 | P07237;H7BZ94;F5H8J2;H0Y3Z3;I3L398;I3L312;I3NI03;I3L0S0;I3L3P5;I3 | P07237;H7BZ94;F5H8J2 |
| Protein disulfide-isomerase A3 | PDIA3 | | 0.579479 | -0.306088 | G5EA52;P30101 | G5EA52;P30101 |
| Protein disulfide-isomerase A6 | PDIA6 | | 0.131342 | 0.13582 | Q15084-3;Q15084;F8WA83;B5MCQ5;Q15084-2 | Q15084-3;Q15084;F8WA83;B5MCQ5;Q15084-2 |
| Protein DJ-1 | PARK7 | | 0.657458 | -0.332258 | Q99497;K7ELW0;K7EN27 | Q99497;K7ELW0;K7EN27 |
| Protein FAM3A | FAM3A | | 0.717948 | 0.355599 | P98173-3;P98173;Q5HY75;P98173-2;A6QRH7 | P98173-3;P98173;Q5HY75;P98173-2 |
| Protein FAM3C | FAM3C | | 0.129155 | 0.0590777 | Q92520;C9JP35;C9JMN4 | Q92520;C9JP35;C9JMN4 |
| Protein FAM49B | FAM49B | + | 1.53304 | -1.13679 | Q9NUQ9;Q9NW21;E5RI16;E5RJL8;E5RJE1;E5RIR8;E5RGI7;E5RK61;E5RF | Q9NUQ9;Q9NW21 |
| Protein Hikeshi | C11orf73 | | 0.262442 | -0.364868 | Q53FT3;E9PPG8;E9PJB2 | Q53FT3;E9PPG8 |
| Protein jagged-1 | JAG1 | | 0.266957 | -0.063825 | P78504;B4DYR1;Q9Y219-2;Q9Y219 | P78504;B4DYR1 |
| Protein kinase C and casein kinase substrate in neurons protein 2 | PACSIN2 | | 0.227949 | 0.280576 | Q9UNF0-2;Q9UNF0;B0QYG7;B0QYG8 | Q9UNF0-2;Q9UNF0 |
| Protein kinase C and casein kinase substrate in neurons protein 3 | PACSIN3 | | 0.0335818 | 0.0227473 | Q9UKS6;E9PIY1;E9PJ75;E9PNM9;E9PJ33;H0YDM6 | Q9UKS6;E9PIY1 |
| Protein lifeguard 3 | TMBIM1 | | 0.0568527 | 0.0498341 | C9JEN3;C9IYT2;F8WDY4;C9IZ27;C9JW19;C9JWV9;B4DUD2;Q969X1 | C9JEN3;C9IYT2;F8WDY4;C9IZ27;C9JW19;C9JWV9;B4DUD2;Q969X1 |
| Protein lin-7 homolog C | LIN7C | | 0.100585 | -0.0902159 | Q9NUP9;G3V1D4 | Q9NUP9;G3V1D4 |
| Protein MON2 homolog | MON2 | | 0.0856545 | 0.0637315 | F8W1Z6;Q7Z3U7-2;F8VWA6;Q7Z3U7-5;Q7Z3U7-4;Q7Z3U7;Q7Z3U7-3;F | F8W1Z6;Q7Z3U7-2;F8VWA6;Q7Z3U7-5;Q7Z3U7-4;Q7Z3U7;Q7Z3U7-3 |
| Protein NDNF | NDNF | | 0.856676 | 0.551001 | Q8TB73;D6R972;D6RF18 | Q8TB73 |
| Protein NDRG1 | NDRG1 | + | 1.45035 | -1.61468 | Q92597-3;Q92597-2;Q92597;E7ESM1;E9PDL6;E5RG99;E5RJY1;E5RIR1; | Q92597-3;Q92597-2;Q92597;E7ESM1;E9PDL6 |
| Protein Niban | FAM129A | | 0.217531 | 0.145204 | Q9BZQ8;H0Y7M9 | Q9BZQ8 |
| Protein NOV homolog | NOV | + | 1.50693 | -1.83301 | P48745 | P48745 |
| Protein phosphatase 1 regulatory subunit 7 | PPP1R7 | | 0.416969 | 0.63012 | H7C003;Q15435-2;Q15435;C9J177;B5MBZ8;C9JD73;B5MCY6;Q15435-4 | H7C003;Q15435-2;Q15435;C9J177;B5MBZ8;C9JD73;B5MCY6;Q15435- |

| | | | | | | |
|---|---|---|---|---|---|---|
| Protein S100-A10 | S100A10 | | 0.619927 | 0.873546 | P60903 | P60903 |
| Protein S100-A11;Protein S100-A11, N-terminally processed | S100A11 | | 0.0625391 | -0.0448924 | P31949 | P31949 |
| Protein S100-A13 | S100A13 | | 0.402836 | -0.358149 | Q99584 | Q99584 |
| Protein S100-A16 | S100A16 | | 0.403289 | -0.278648 | Q96FQ6 | Q96FQ6 |
| Protein S100-A6;Protein S100 | S100A6 | | 0.181794 | -0.133388 | P06703;R4GN98 | P06703;R4GN98 |
| Protein SEC13 homolog | SEC13 | | 0.303928 | 0.300915 | P55735;P55735-2;P55735-3;A8MXL6;E7ERC8 | P55735;P55735-2;P55735-3;A8MXL6 |
| Protein TFG | TFG | | 0.415765 | 0.474431 | Q92734-2;Q92734;C9JUE0;C9JJP5;C9JTY3 | Q92734-2;Q92734;C9JUE0;C9JJP5;C9JTY3 |
| Protein tweety homolog 3;Protein tweety homolog | TTYH3 | | 0.469597 | -0.561177 | Q9C0H2-3;Q9C0H2-2;Q9C0H2;Q9C0H2-4;H7C3T6 | Q9C0H2-3;Q9C0H2-2;Q9C0H2;Q9C0H2-4;H7C3T6 |
| Protein tyrosine phosphatase type IVA 2;Protein tyrosine phosphatase type IVA 1 | PTP4A2;PTP4A1 | + | 1.46189 | -0.503244 | Q12974;E9PMY3;E9PML8;E9PJC0;E9PRR9;Q12974-2;R4GN50;Q93096; | Q12974;E9PMY3;E9PML8;E9PJC0;E9PRR9;Q12974-2;R4GN50;Q93096; |
| Protein unc-119 homolog B | UNC119B | | 0.183116 | 0.392214 | A6NIH7;K7EJU3;K7EN86;C9JSK0;Q13432-2;Q13432 | A6NIH7 |
| Protein XRP2 | RP2 | | 1.04824 | -1.56949 | O75695 | O75695 |
| Protein-glutamine gamma-glutamyltransferase 2 | TGM2 | + | 1.37113 | -1.64697 | P21980;B4DIT7;F5H6P0;P21980-2;A2A299;P21980-3;A2A2A0;O43548-2 | P21980;B4DIT7;F5H6P0;P21980-2 |
| Protein-L-isoaspartate O-methyltransferase;Protein-L-isoaspartate(D-aspartate) O-methyltransferase | PCMT1 | | 0.112891 | -0.0774688 | H7BY58;P22061;P22061-2;B7Z972;C9J0F2;F8WDT3;F8WAV5;F8WAX2; | H7BY58;P22061;P22061-2;B7Z972 |
| Protein-lysine 6-oxidase | LOX | + | 1.34285 | -0.772052 | P28300 | P28300 |
| Prothymosin alpha;Prothymosin alpha, N-terminally processed;Thymosin alpha-1 | PTMA | | 0.016933 | 0.0170124 | B8ZZQ6;P06454-2;P06454;B8ZZA1;B8ZZW7;H7C2N1 | B8ZZQ6;P06454-2;P06454;B8ZZA1;B8ZZW7;H7C2N1 |
| Protocadherin Fat 1;Protocadherin Fat 1, nuclear form | FAT1 | | 0.666285 | -0.378865 | Q14517;D6RCE4 | Q14517 |
| Protocadherin Fat 4 | FAT4 | | 0.13654 | 0.0918136 | Q6V0I7-3;Q6V0I7;Q6V0I7-2 | Q6V0I7-3;Q6V0I7 |
| Protogenin | PRTG | | 0.362042 | -0.539412 | Q2VWP7;H0YLD1 | Q2VWP7 |
| Proto-oncogene tyrosine-protein kinase Src | SRC | | 0.82974 | -1.50863 | P12931;P12931-2 | P12931;P12931-2 |
| Pseudouridine-5-monophosphatase | HDHD1 | + | 1.37857 | -1.25513 | E7EVH9;Q08623-3;Q08623;Q08623-4;Q08623-2 | E7EVH9;Q08623-3;Q08623;Q08623-4;Q08623-2 |
| Pterin-4-alpha-carbinolamine dehydratase | PCBD1 | | 0.24098 | -0.171037 | P61457 | P61457 |
| Purine nucleoside phosphorylase | PNP | | 0.0339041 | -0.0230387 | P00491;G3V5M2;G3V2H3;G3V393 | P00491;G3V5M2 |
| Puromycin-sensitive aminopeptidase | NPEPPS | | 1.2322 | -0.748587 | E9PLK3;P55786;B7Z463;E5RJ24;E7EWZ2;I3L083;A6NEC2;F5GZY4;H0YA | E9PLK3;P55786;B7Z463 |
| Putative adenosylhomocysteinase 2;Adenosylhomocysteinase;Putative adenosylhomocysteinase 3 | AHCYL1;AHCYL2 | | 0.684027 | -0.917198 | O43865-2;O43865;D7UEQ7;H0Y8B3;F8W7N8;Q96HN2-4;Q96HN2-3;Q9 | O43865-2;O43865;D7UEQ7;H0Y8B3;F8W7N8;Q96HN2-4;Q96HN2-3;Q |
| Putative deoxyribose-phosphate aldolase | DERA | | 0.328623 | 0.406789 | Q9Y315;E9PPM8;G3V158;E9PML7;E9PMH9;E9PPK3;E9PJ44;E9PJB9 | Q9Y315;E9PPM8;G3V158;E9PML7 |
| Putative heat shock protein HSP 90-alpha A2 | HSP90AA2 | | 0.0800844 | 0.0994509 | Q14568 | Q14568 |
| Putative hydrolase RBBP9 | RBBP9 | | 0.10316 | -0.0535524 | O75884;O75884-2 | O75884;O75884-2 |
| Putative peptidyl-tRNA hydrolase PTRHD1 | PTRHD1 | | 0.390273 | -0.297475 | Q6GMV3 | Q6GMV3 |
| Putative phospholipase B-like 2;Putative phospholipase B-like 2 32 kDa form;Putative phospholipase B-lik | PLBD2 | | 0.307409 | -0.232056 | Q8NHP8;F5H5E2 | Q8NHP8;F5H5E2 |
| Pyridoxal kinase | PDXK | | 0.170291 | -0.214578 | O00764;F2Z2Y4;O00764-3;O00764-2;A8MV33 | O00764;F2Z2Y4;O00764-3;O00764-2 |
| Pyridoxal phosphate phosphatase | PDXP | | 0.0900895 | -0.0761975 | Q96GD0 | Q96GD0 |
| Pyridoxine-5-phosphate oxidase | PNPO | | 0.482417 | 0.731845 | Q9NVS9;B4E1D7;B4E0V0;B4E152;J3QQV6 | Q9NVS9;B4E1D7;B4E0V0;B4E152 |
| Pyruvate dehydrogenase E1 component subunit beta, mitochondrial | PDHB | | 0.349786 | -0.405753 | P11177-3;P11177-2;P11177;F8WF02;C9J634 | P11177-3;P11177-2;P11177;F8WF02;C9J634 |
| Pyruvate kinase PKM | PKM | | 0.0151949 | -0.013335 | P14618;P14618-3 | P14618;P14618-3 |
| Pyruvate kinase PKM;Pyruvate kinase | PKM;PKM2 | | 1.21928 | -0.410402 | P14618-2;H3BTN5;H3BQ34;Q504U3;H3BTJ2;H3BUW1;H3BT25;H3BU1 | P14618-2;H3BTN5;H3BQ34;Q504U3 |
| Quinone oxidoreductase | CRYZ | | 0.116218 | 0.0949714 | Q08257;Q08257-3;A6NP24;C9JH92;Q08257-2 | Q08257;Q08257-3;A6NP24;C9JH92 |
| Quinone oxidoreductase PIG3 | TP53I3 | | 0.300753 | 0.190243 | Q53FA7;Q53FA7-2;H7BZH6 | Q53FA7 |
| Rab GDP dissociation inhibitor beta | GDI2 | | 0.0172826 | -0.0119096 | P50395;E7EU23;P50395-2;Q5SX87;Q5SX86;V9GYF8;Q5SX90;Q5SX91;V | P50395;E7EU23;P50395-2;Q5SX87 |
| Radixin | RDX | | 0.440575 | 0.357679 | P35241;P35241-5;P35241-4;P35241-2;H0YE16;E9PNV3;P35241-3;E9PQ | P35241;P35241-5;P35241-4 |
| Raftlin | RFTN1 | | 0.283269 | -0.491948 | Q14699;G3XAJ6;C9JHG2;C9JWQ9;F8WAR1;C9JRN3 | Q14699;G3XAJ6;C9JHG2 |
| Ragulator complex protein LAMTOR1 | LAMTOR1 | | 0.374615 | -0.370269 | Q6IAA8;H0YFI1;F5H479;F5GX19;F5H3Y3 | Q6IAA8 |
| Ragulator complex protein LAMTOR3 | LAMTOR3 | | 0.293641 | -0.436414 | Q9UHA4;Q9UHA4-2 | Q9UHA4;Q9UHA4-2 |
| Rap1 GTPase-GDP dissociation stimulator 1 | RAP1GDS1 | | 0.0457513 | 0.0762743 | P52306-4;P52306;P52306-5;P52306-6;P52306-2;P52306-3;D6REZ0;D6R | P52306-4;P52306;P52306-5;P52306-6;P52306-2;P52306-3 |
| Ras GTPase-activating-like protein IQGAP1 | IQGAP1 | | 0.507652 | -0.375445 | P46940;H0YLE8;F5H7S7;E7EWC2;Q13576-2;J3KR91;Q13576;H0YKA5;E | P46940;H0YLE8 |
| Ras suppressor protein 1 | RSU1 | | 0.221169 | -0.166562 | Q15404;Q15404-2 | Q15404;Q15404-2 |
| Ras-related C3 botulinum toxin substrate 1;Ras-related C3 botulinum toxin substrate 3 | RAC1;RAC3 | | 0.985261 | -0.563952 | P63000;P63000-2;P60763;J3KSC4;J3QLK0 | P63000;P63000-2;P60763 |
| Ras-related C3 botulinum toxin substrate 2 | RAC2 | | 0.668285 | -0.395076 | P15153;B1AH77;B1AH80;B1AH78 | P15153;B1AH77;B1AH80;B1AH78 |
| Ras-related protein Rab-10 | RAB10 | | 0.645129 | -0.590226 | P61026 | P61026 |
| Ras-related protein Rab-11B;Ras-related protein Rab-11A | RAB11B;RAB11A | | 0.974772 | -0.456828 | Q15907;B4DMK0;J3KQP6;H3BMH2;H3BSC1;P62491-2;P62491;B4DQU5 | Q15907;B4DMK0;J3KQP6;H3BMH2;H3BSC1;P62491-2;P62491 |
| Ras-related protein Rab-13 | RAB13 | | 0.0568448 | -0.0532914 | P51153 | P51153 |
| Ras-related protein Rab-14 | RAB14 | | 0.518877 | -0.219973 | P61106 | P61106 |
| Ras-related protein Rab-18 | RAB18 | | 0.259442 | 0.207769 | Q9NP72;Q9NP72-2;Q5W0J0;H0Y6T8;Q9NP72-3 | Q9NP72;Q9NP72-2;Q5W0J0;H0Y6T8 |
| Ras-related protein Rab-1A | RAB1A | | 0.0995539 | -0.0834841 | P62820;B7Z8M7;E7END7;P62820-2;P62820-3;E7ETK2;P59190-2;P5919 | P62820;B7Z8M7;E7END7;P62820-2 |
| Ras-related protein Rab-1B;Putative Ras-related protein Rab-1C | RAB1B;RAB1C | | 0.0408059 | 0.035155 | Q9H0U4;E9PLD0;Q92928 | Q9H0U4;E9PLD0;Q92928 |
| Ras-related protein Rab-21 | RAB21 | | 0.509059 | 0.278364 | Q9UL25 | Q9UL25 |
| Ras-related protein Rab-22A | RAB22A | | 0.762961 | -0.447864 | Q9UL26;J3QR51;Q13636 | Q9UL26 |
| Ras-related protein Rab-23 | RAB23 | | 0.199525 | -0.421245 | Q9ULC3 | Q9ULC3 |
| Ras-related protein Rab-2A | RAB2A | | 0.199443 | -0.135658 | P61019;P61019-2;E9PKL7;H7C125;Q8WUD1;H0YD31;E9PE37;H0YDL5 | P61019;P61019-2;E9PKL7 |
| Ras-related protein Rab-32 | RAB32 | | 0.350614 | 0.309031 | Q13637;O14966;P57729 | Q13637 |
| Ras-related protein Rab-34 | RAB34 | | 0.508735 | -0.383394 | A8MYQ9;J3KQW8;Q9BZG1-2;Q9BZG1;C9JBG0;C9JY26;Q9BZG1-4;K7EIF | A8MYQ9;J3KQW8;Q9BZG1-2;Q9BZG1;C9JBG0;C9JY26;Q9BZG1-4;K7EIF |
| Ras-related protein Rab-35 | RAB35 | | 0.905224 | -1.0222 | Q15286;F5H157;Q15286-2;E9PI18;E9PJQ5;E9PRF7;E9PNB9;E9PS06;E9P | Q15286;F5H157;Q15286-2 |
| Ras-related protein Rab-5A | RAB5A | | 0.665256 | -0.304947 | B4DJA5;P20339;F8VVZ0;F8WCY6;F8WD79;F8VPW9;F8VWZ7 | B4DJA5;P20339 |
| Ras-related protein Rab-5B | RAB5B | | 0.779857 | -0.419318 | P61020;P61020-2;F8VUA5 | P61020;P61020-2;F8VUA5 |
| Ras-related protein Rab-5C | RAB5C | | 0.811945 | -0.473695 | P51148;P51148-2;K7ERI8;K7ERQ8;F8VVK3;K7ENY4;F8VWU4;K7EIP6;F8 | P51148;P51148-2;K7ERI8;K7ERQ8 |
| Ras-related protein Rab-6A;Ras-related protein Rab-6B | RAB6A;RAB6B | | 0.55241 | -0.369368 | P20340-2;P20340;H7BYW1;H0YGL6;P20340-4;Q9NRW1;J3KR73;P2034 | P20340-2;P20340;H7BYW1;H0YGL6;P20340-4;Q9NRW1 |
| Ras-related protein Rab-7a | RAB7A | | 0.583425 | -0.248116 | P51149;C9J592;C9J8S3;C9IZZ0;C9J4V0;C9J4S4;C9J7D1 | P51149;C9J592;C9J8S3;C9IZZ0;C9J4V0 |
| Ras-related protein Rab-8A | RAB8A | | 0.946518 | -0.585235 | P61006;B4DEK7;H0YMN7;M0R257;O95716;P20336;Q96E17;H0YLJ8;H0 | P61006;B4DEK7 |
| Ras-related protein Rab-8B | RAB8B | | 1.16636 | -1.01473 | H0YNE9;Q92930;F5GY21 | H0YNE9;Q92930;F5GY21 |
| Ras-related protein Rab-9A;Ras-related protein Rab-9B | RAB9A;RAB9B | | 1.02088 | -0.50412 | P51151;Q9NP90 | P51151;Q9NP90 |
| Ras-related protein Ral-A | RALA | | 0.645741 | -0.558727 | P11233;H7C3P7 | P11233;H7C3P7 |
| Ras-related protein Ral-B | RALB | | 0.526741 | -0.822757 | P11234;B4E040;Q6ZS74;C9J6B1;C9JQB3;C9JPE8;C9JYR1;F8WEQ6 | P11234;B4E040;Q6ZS74;C9J6B1;C9JQB3 |
| Ras-related protein Rap-1b;Ras-related protein Rap-1b-like protein;Ras-related protein Rap-1A | RAP1B;RAP1A | | 0.0815297 | -0.0898523 | P61224-3;P61224;E7ESV4;F5H7Y6;A6NIZ1;F5GX62;P61224-2;P61224-4; | P61224-3;P61224;E7ESV4;F5H7Y6;A6NIZ1;F5GX62;P61224-2;P61224-4 |
| Ras-related protein Rap-2b | RAP2B | | 0.666787 | -0.766543 | P61225 | P61225 |
| Ras-related protein R-Ras | RRAS | | 0.938843 | -0.987678 | P10301 | P10301 |
| Ras-related protein R-Ras2 | RRAS2 | | 0.774197 | -0.674565 | P62070;B7Z5Z2;P62070-3;P62070-2;E9PK85;E9PQ87;E9PQK5;E9PQC5 | P62070;B7Z5Z2;P62070-3;P62070-2;E9PK85 |
| Receptor expression-enhancing protein 5 | REEP5 | | 0.536538 | 0.655402 | Q00765;E2QRG8 | Q00765 |
| Receptor-type tyrosine-protein phosphatase eta | PTPRJ | | 0.790538 | -0.813604 | Q12913-2;Q12913;E9PPH3 | Q12913-2;Q12913 |
| Receptor-type tyrosine-protein phosphatase F | PTPRF | | 1.06064 | -1.23314 | P10586-2;P10586;H0Y6Z7;H0Y4H1;H0Y7Z9;F2Z3B8;H0Y380;A2A437;P2 | P10586-2;P10586;H0Y6Z7;H0Y4H1 |
| Receptor-type tyrosine-protein phosphatase gamma | PTPRG | | 0.557625 | -0.479435 | P23470-2;P23470 | P23470-2;P23470 |

| | | | | | | |
|---|---|---|---|---|---|---|
| Receptor-type tyrosine-protein phosphatase S | PTPRS | | 0.715152 | -0.802038 | G8JL96;E9PDC5;Q13332-6;Q13332-3;Q13332-4;Q13332-2;Q13332;Q13 | G8JL96;E9PDC5;Q13332-6;Q13332-3;Q13332-4;Q13332-2;Q13332;Q13 |
| Renin receptor | ATP6AP2 | + | 1.8307 | 1.60318 | B7Z9I3;O75787;H0Y750;B7Z1I9;H7C3E1;B7Z413;H7C240 | B7Z9I3;O75787;H0Y750;B7Z1I9;H7C3E1;B7Z413;H7C240 |
| Reticulon-3;Reticulon | RTN3 | | 0.0471127 | 0.0720066 | O95197-3;O95197-4;O95197-7;O95197-2;O95197;B7Z4M1;O95197-6;O | O95197-3;O95197-4;O95197-7;O95197-2;O95197;B7Z4M1;O95197-6;O |
| Reticulon-4;Reticulon | RTN4 | | 0.726893 | 0.717499 | Q9NQC3-2;F8W914;Q9NQC3-5;Q9NQC3;Q9NQC3-4;H7C106;Q9NQC3-3 | Q9NQC3-2;F8W914;Q9NQC3-5;Q9NQC3;Q9NQC3-4;H7C106;Q9NQC3- |
| Retinoic acid-induced protein 3 | GPRC5A | | 0.25934 | -0.306894 | F5GWG3;Q8NFJ5 | F5GWG3;Q8NFJ5 |
| Retinoid-inducible serine carboxypeptidase | SCPEP1 | | 0.687207 | -0.381836 | Q9HB40;Q9HB40-2;I3L4Z9;I3L4Z3;I3L506 | Q9HB40;Q9HB40-2 |
| Rho GDP-dissociation inhibitor 1 | ARHGDIA | | 0.0941135 | 0.101765 | P52565;J3KTF8;J3QQX2;J3KRE2;J3KRY1;P52565-2;J3KS60 | P52565;J3KTF8;J3QQX2;J3KRE2;J3KRY1;P52565-2 |
| Rho GDP-dissociation inhibitor 2 | ARHGDIB | | 0.0793362 | -0.127218 | P52566;F5H3P3;F5H6Q0;F5H2R5;H0YGX7 | P52566;F5H3P3;F5H6Q0;F5H2R5;H0YGX7 |
| Rho GTPase-activating protein 1 | ARHGAP1 | | 0.25679 | 0.412143 | Q07960;H0YE29;E9PNR6 | Q07960;H0YE29 |
| Rho-related GTP-binding protein RhoC | RHOC | | 0.826115 | -0.449448 | Q5JR08;P08134;E9PQH6;Q5JR07;Q5JR05;Q5JR06;E9PN11;U3KQA9;U3K | Q5JR08;P08134;E9PQH6;Q5JR07;Q5JR05;Q5JR06;E9PN11;U3KQA9;U3 |
| Rho-related GTP-binding protein RhoE | RND3 | | 0.256313 | 0.255954 | P61587;E9PFH1;Q53RZ3 | P61587;E9PFH1 |
| Rho-related GTP-binding protein RhoG | RHOG | | 0.871259 | -0.692481 | P84095 | P84095 |
| Ribonuclease 4 | RNASE4 | | 1.28245 | -2.38774 | P34096 | P34096 |
| Ribonuclease inhibitor | RNH1 | | 0.216414 | -0.156751 | P13489;H0YCR7;E9PIM9;E9PMJ3;E9PLZ3;E9PIK5;E9PMN0;E9PMA9;E9P | P13489 |
| Ribonuclease T2 | RNASET2 | | 0.843073 | -0.64245 | D6RHI9;O00584;D6REQ6;H0YAE9;O00584-2;J3QQ64 | D6RHI9;O00584;D6REQ6;H0YAE9 |
| Ribonucleoside-diphosphate reductase large subunit | RRM1 | | 0.363195 | -0.452323 | P23921;E9PD78;E9PL69;F5H861;E9PP77;H0YCY7 | P23921;E9PD78;E9PL69;F5H861 |
| Ribonucleoside-diphosphate reductase subunit M2 | RRM2 | | 0.590375 | -0.680854 | P31350;P31350-2;C9JXC1;Q7LG56-3 | P31350;P31350-2;C9JXC1 |
| Ribonucleoside-diphosphate reductase subunit M2 B | RRM2B | | 0.664666 | -0.546306 | H0YAV1;Q7LG56;Q7LG56-6;Q7LG56-2 | H0YAV1;Q7LG56;Q7LG56-6;Q7LG56-2 |
| Ribose-phosphate pyrophosphokinase 2 | PRPS2 | | 0.414841 | 0.596707 | P11908;P11908-2;H7C540;D3YTJ7;A6NMS2 | P11908;P11908-2;H7C540 |
| Ribosyldihydronicotinamide dehydrogenase [quinone] | NQO2 | | 0.136806 | -0.116434 | P16083;Q5TD07;Q5TD05;A2A2U4 | P16083;Q5TD07;Q5TD05 |
| Ribulose-phosphate 3-epimerase | RPE | | 0.208214 | 0.179693 | Q96AT9;E7EW52;B4E016;Q96AT9-2;C9J6A7;C9J9T0;C9IZU8;C9JPQ7;Q9 | Q96AT9;E7EW52;B4E016;Q96AT9-2;C9J6A7;C9J9T0;C9IZU8;C9JPQ7;Q |
| RING finger protein 11 | RNF11 | | 0.133939 | 0.246939 | Q9Y3C5 | Q9Y3C5 |
| RNA-binding motif protein, X chromosome;RNA-binding motif protein, X chromosome, N-terminally proc | RBMX | | 0.14862 | -0.139155 | P38159;H3BT71;P38159-2;H0Y6E7;H3BR27;P38159-3;B3KRG5;H3BUY5; | P38159;H3BT71;P38159-2;H0Y6E7 |
| Rootletin | CROCC | | 0.648956 | -0.637622 | Q5TZA2;Q5TZA2-2;B1AKD8 | Q5TZA2;Q5TZA2-2 |
| Roundabout homolog 1 | ROBO1 | | 0.920398 | 0.26335 | Q9Y6N7-6;Q9Y6N7-5;Q9Y6N7-4;Q9Y6N7-3;Q9Y6N7;Q9Y6N7-2;H7C4Y7 | Q9Y6N7-6;Q9Y6N7-5;Q9Y6N7-4;Q9Y6N7-3;Q9Y6N7;Q9Y6N7-2 |
| rRNA 2-O-methyltransferase fibrillarin | FBL | | 0.752783 | 0.257799 | P22087;M0R0P1;M0R299;M0QXL5;M0R2Q4;M0R2U2;M0R1H0;M0R2B | P22087;M0R0P1;M0R299;M0QXL5;M0R2Q4;M0R2U2;M0R1H0;M0R2E |
| RuvB-like 1 | RUVBL1 | | 0.353189 | 0.326265 | Q9Y265;Q9Y265-2;E7ETR0;H7C4G5;J3QLR1;H7C4I3 | Q9Y265;Q9Y265-2;E7ETR0 |
| RuvB-like 2 | RUVBL2 | | 0.421552 | 0.365104 | Q9Y230;B3KQ59;M0R0Y3;M0QXI6;M0R0Z0;M0QXZ7 | Q9Y230;B3KQ59;M0R0Y3 |
| SEC14-like protein 2 | SEC14L2 | | 0.240368 | -0.376282 | B3KRD8;O76054;O76054-5;O76054-4;C9JYY7;C9JZI9;H7C417 | B3KRD8;O76054;O76054-5;O76054-4 |
| Secreted and transmembrane protein 1 | SECTM1 | | 0.697309 | 0.656726 | J3KSN0;J3QQU6;J3QKK2;J3KTR4;Q8WVN6 | J3KSN0;J3QQU6;J3QKK2;J3KTR4;Q8WVN6 |
| Secretory carrier-associated membrane protein 3 | SCAMP3 | | 0.759231 | 0.362537 | O14828-2;O14828 | O14828-2;O14828 |
| Semaphorin-3C | SEMA3C | | 0.480116 | 0.298321 | F5H1Z7;Q99985;B4DRL8 | F5H1Z7;Q99985 |
| Semaphorin-4C | SEMA4C | | 0.123165 | -0.124891 | Q9C0C4 | Q9C0C4 |
| Sepiapterin reductase | SPR | | 0.457099 | -0.495714 | P35270 | P35270 |
| Septin-11 | SEPT11 | | 0.340671 | 0.469252 | D6RGI3;D6RER5;Q9NVA2;Q9NVA2-2;D6RDU5;H0Y961;B1AMS2;F5H1J5 | D6RGI3;D6RER5;Q9NVA2;Q9NVA2-2;D6RDU5 |
| Septin-2 | SEPT2 | | 0.590541 | 0.799201 | Q15019;Q15019-2;C9J938;C9JB25;C9IZU3;C9JZI2;C9JTV8;C9JFT1;F8WB | Q15019;Q15019-2 |
| Septin-7 | SEPT7 | | 0.497206 | 0.678466 | Q16181;E7ES33;E7EPK1;Q16181-2;H0Y3Y4;H0YFF6;F5GZE5;Q5JXL7;Q6 | Q16181;E7ES33;E7EPK1;Q16181-2;H0Y3Y4;H0YFF6 |
| Sequestosome-1 | SQSTM1 | | 0.702217 | 0.578118 | E7EMC7;Q13501;E9PFW8;E3W990;Q13501-2;D6RBF1;C9J6J8;C9JRJ8 | E7EMC7;Q13501;E9PFW8;E3W990;Q13501-2 |
| Serglycin | SRGN | + | 1.84146 | 1.95594 | P10124 | P10124 |
| Serine dehydratase-like | SDSL | | 0.00867294 | 0.0152058 | Q96GA7;F8VYZ3;H0YID3;H0YIA2 | Q96GA7;F8VYZ3 |
| Serine hydroxymethyltransferase, cytosolic | SHMT1 | | 0.0605935 | 0.0638655 | P34896-2;P34896;P34896-4;P34896-3;J3KRK5;J3KRZ5 | P34896-2;P34896;P34896-4;P34896-3 |
| Serine hydroxymethyltransferase, mitochondrial;Serine hydroxymethyltransferase | SHMT2 | | 0.334055 | -0.220731 | P34897-3;P34897;P34897-2;B4DLV4;H0YIZ0;G3V5L0;G3V2Y4;G3V4W5; | P34897-3;P34897;P34897-2;B4DLV4;H0YIZ0 |
| Serine incorporator 1 | SERINC1 | | 0.142583 | 0.0870815 | Q9NRX5 | Q9NRX5 |
| Serine protease 23 | PRSS23 | | 0.105997 | -0.074368 | O95084;B4E2J3;E9PRR2 | O95084;B4E2J3 |
| Serine protease HTRA1 | HTRA1 | | 1.05961 | 0.275931 | Q92743;H0Y7G9 | Q92743 |
| Serine protease HTRA2, mitochondrial | HTRA2 | | 0.0572994 | 0.0519061 | O43464-3;O43464;O43464-2;O43464-4;P83110-2;P83110 | O43464-3;O43464;O43464-2;O43464-4 |
| Serine/arginine-rich splicing factor 1 | SRSF1 | | 0.318094 | -0.551548 | J3KTL2;Q07955;Q07955-3;Q07955-2;J3KSR8;J3QQV5;J3KSW7 | J3KTL2;Q07955;Q07955-3;Q07955-2 |
| Serine/arginine-rich splicing factor 2 | SRSF2 | | 0.0132309 | 0.0252246 | J3QL05;J3KP15;B4DN89;Q01130;Q9BRL6-2;Q9BRL6 | J3QL05;J3KP15;B4DN89;Q01130 |
| Serine/arginine-rich splicing factor 7 | SRSF7 | | 0.291789 | -0.329265 | C9JAB2;Q16629-3;Q16629-2;Q16629-4;Q16629;F8WEA1 | C9JAB2;Q16629-3;Q16629-2;Q16629-4;Q16629 |
| Serine/threonine-protein kinase 10 | STK10 | | 0.186471 | -0.194359 | O94804 | O94804 |
| Serine/threonine-protein kinase 24;Serine/threonine-protein kinase 24 36 kDa subunit;Serine/threonine- | STK24 | | 0.98507 | 0.953265 | B4DR80;Q9Y6E0-2;Q9Y6E0;H0Y630;Q5JV98;Q5JV99 | B4DR80;Q9Y6E0-2;Q9Y6E0;H0Y630;Q5JV98 |
| Serine/threonine-protein kinase Chk1 | CHEK1 | | 0.541488 | -0.537784 | J3KN87;E7EPP6;O14757-2;O14757-3;O14757;E9PPA5 | J3KN87;E7EPP6;O14757-2;O14757-3;O14757;E9PPA5 |
| Serine/threonine-protein kinase MST4 | RBM4B;MST4 | | 0.829918 | 0.687189 | Q8NBY1;B4E0Y9;Q9P289;Q9P289-2;Q9P289-3;C9JJV0 | Q8NBY1;B4E0Y9;Q9P289;Q9P289-2;Q9P289-3 |
| Serine/threonine-protein phosphatase 2A 65 kDa regulatory subunit A alpha isoform | PPP2R1A | | 0.262252 | 0.237324 | P30153;F5H3X9;B3KQV6;C9J9C1;E9PH38;E9PPI5;E9PHZ6;M0QXG4;M0I | P30153;F5H3X9;B3KQV6;C9J9C1 |
| Serine/threonine-protein phosphatase 2A 65 kDa regulatory subunit A beta isoform | PPP2R1B | | 0.157765 | 0.278482 | P30154;P30154-2;P30154-5;P30154-4;P30154-3;J3KR29;H0YDG7 | P30154;P30154-2;P30154-5;P30154-4;P30154-3;J3KR29;H0YDG7 |
| Serine/threonine-protein phosphatase 2A activator | PPP2R4 | | 0.0370436 | -0.043781 | Q15257;A6PVN5;B4DZF8;Q15257-2;Q15257-3;Q5T949;Q5T948;A6PVN | Q15257;A6PVN5;B4DZF8;Q15257-2;Q15257-3;Q5T949;Q5T948;A6PVN |
| Serine/threonine-protein phosphatase 2A catalytic subunit alpha isoform | PPP2CA | | 0.11718 | 0.124482 | P67775;P67775-2 | P67775;P67775-2 |
| Serine/threonine-protein phosphatase 2A catalytic subunit beta isoform | PPP2CB | | 0.099258 | 0.124557 | P62714;E5RHC1;H0YC23;E7ESG8;E5RFI3;E5RHP4;H0YBN9;B7Z2C5;E5RJ | P62714 |
| Serine/threonine-protein phosphatase 2B catalytic subunit alpha isoform;Serine/threonine-protein phosp | PPP3CA | | 0.0745752 | 0.0617438 | Q08209-2;Q08209;Q08209-3;E7ETC2;Q08209-4;Q08209-5;E9PPC8;E9PI | Q08209-2;Q08209;Q08209-3;E7ETC2;Q08209-4;Q08209-5 |
| Serine/threonine-protein phosphatase CPPED1 | CPPED1 | | 0.952632 | 0.549433 | Q9BRF8;Q9BRF8-2;B4DQ68 | Q9BRF8;Q9BRF8-2 |
| Serine/threonine-protein phosphatase PP1-alpha catalytic subunit;Serine/threonine-protein phosphatase | PPP1CA | | 0.187273 | 0.228111 | P62136;P62136-2;P62136-3;E9PMD7;F5H1L6;F5H037 | P62136;P62136-2;P62136-3;E9PMD7 |
| Serine/threonine-protein phosphatase PP1-beta catalytic subunit;Serine/threonine-protein phosphatase | PPP1CB | | 0.0194582 | 0.0276166 | P62140;E7ETD8;C9JP48;C9J9S3 | P62140;E7ETD8 |
| Serine--tRNA ligase, cytoplasmic | SARS | | 0.3199 | -0.397023 | Q5T5C7;P49591 | Q5T5C7;P49591 |
| Serpin B6 | SERPINB6 | | 0.881077 | -0.468926 | P35237 | P35237 |
| Serpin B7 | SERPINB7 | | 0.627559 | -0.550426 | O75635-2;O75635;C9JA68;C9JZR8;C9JM00 | O75635-2;O75635 |
| Serpin B8 | SERPINB8 | | 0.139594 | -0.180625 | P50452;P50452-2;H7BXK7;C9JVA8;C9JTJ8;B4DTW2 | P50452;P50452-2;H7BXK7;C9JVA8 |
| Serpin B9 | SERPINB9 | | 0.874433 | -1.00674 | P50453 | P50453 |
| Serpin H1 | SERPINH1 | | 0.539593 | 0.421428 | P50454;E9PPV6;E9PKH2;E9PR70;E9PK86;E9PMI5;E9PNX1;E9PJH8;E9PI | P50454;E9PPV6;E9PKH2;E9PR70 |
| S-formylglutathione hydrolase | ESD | | 0.150027 | 0.153062 | P10768;H7BZT7;U3KQT1 | P10768;H7BZT7 |
| SH3 domain-binding glutamic acid-rich-like protein | SH3BGRL | | 0.191548 | -0.144711 | O75368 | O75368 |
| SH3 domain-binding glutamic acid-rich-like protein 2 | SH3BGRL2 | | 0.0305875 | 0.0406761 | Q9UJC5 | Q9UJC5 |
| SH3 domain-binding glutamic acid-rich-like protein 3 | SH3BGRL3 | | 0.214285 | 0.137309 | Q9H299;Q5T123 | Q9H299;Q5T123 |
| Sialate O-acetylesterase | SIAE | | 0.0179622 | 0.0193143 | Q9HAT2-2;Q9HAT2 | Q9HAT2-2;Q9HAT2 |
| Sialic acid synthase | NANS | | 0.0274016 | 0.0513862 | Q9NR45;Q5TBR0;Q5TBR1 | Q9NR45 |
| Sialidase-1 | NEU1 | + | 1.47104 | 0.470624 | Q99519;E9PIF4 | Q99519;E9PIF4 |
| Signal recognition particle 14 kDa protein | SRP14 | | 0.867808 | 1.07831 | P37108;H0YLA2;H0YLW0 | P37108;H0YLA2 |
| Signal recognition particle 9 kDa protein | SRP9 | | 0.948735 | 0.916014 | P49458;E9PE20;P49458-2 | P49458;E9PE20;P49458-2 |
| Single-stranded DNA-binding protein, mitochondrial | SSBP1 | | 0.462496 | -0.377621 | Q04837;E7EUY5;C9K0U8 | Q04837;E7EUY5;C9K0U8 |
| Slit homolog 1 protein | SLIT1 | | 1.04384 | -0.936143 | Q5T0V3;Q5T0V2;Q5T0V4;Q5T0V0;O75093-2;O75093 | Q5T0V3;Q5T0V2;Q5T0V4;Q5T0V0;O75093-2;O75093 |

| | | | | | | |
|---|---|---|---|---|---|---|
| Slit homolog 2 protein;Slit homolog 2 protein N-product;Slit homolog 2 protein C-product | SLIT2 | | 0.25776 | -1.55356 | O94813-3;O94813-2;O94813;E9PCX4 | O94813-3;O94813-2;O94813 |
| Small nuclear ribonucleoprotein E | SNRPE | | 0.453021 | 0.426131 | P62304;A6NHK2 | P62304;A6NHK2 |
| Small nuclear ribonucleoprotein Sm D2 | SNRPD2 | | 0.511742 | 0.468975 | P62316;K7ERG4;P62316-2;K7EJB5 | P62316;K7ERG4;P62316-2 |
| Small nuclear ribonucleoprotein Sm D3 | SNRPD3 | | 0.447539 | 0.347142 | B4DJP7;P62318;H3BT13 | B4DJP7;P62318;H3BT13 |
| Small nuclear ribonucleoprotein-associated proteins B and B;Small nuclear ribonucleoprotein-associated | SNRPB;SNRPN | | 0.760874 | 0.516508 | P14678-2;P14678;P14678-3;J3QLE5;P63162;B3KVR1;J3KRY3;S4R3P3;A | P14678-2;P14678;P14678-3;J3QLE5;P63162;B3KVR1;J3KRY3;S4R3P3 |
| Sodium channel and clathrin linker 1 | SCLT1 | | 0.728614 | -0.410962 | Q96NL6 | Q96NL6 |
| Sodium/potassium-transporting ATPase subunit alpha-1 | ATP1A1 | | 0.822221 | -0.923156 | P05023-3;P05023-4;P05023;P05023-2;M0R116;P13637;P13637-2;P136 | P05023-3;P05023-4;P05023;P05023-2 |
| Sodium/potassium-transporting ATPase subunit beta-1 | ATP1B1 | | 0.271722 | -0.4098 | B7Z9S8;A6NGH2;P05026-2;P05026;V9GYR2 | B7Z9S8;A6NGH2;P05026-2;P05026;V9GYR2 |
| Sodium/potassium-transporting ATPase subunit beta-3 | ATP1B3 | | 0.427037 | -0.674794 | P54709;C9JXZ1;C9JA36;B7Z1N7;H7C547;F8WBY4;C9J6S2 | P54709;C9JXZ1;C9JA36;B7Z1N7 |
| Soluble calcium-activated nucleotidase 1 | CANT1 | | 0.833504 | -1.41906 | Q8WVQ1-3;Q8WVQ1;K7EN15;Q8WVQ1-2 | Q8WVQ1-3;Q8WVQ1 |
| Solute carrier family 2, facilitated glucose transporter member 1 | SLC2A1 | | 0.0699605 | 0.0594101 | P11166;C9JIM8 | P11166;C9JIM8 |
| Sorbitol dehydrogenase | SORD | | 1.29455 | -0.758223 | Q00796;H0YLA4;H0YKB3;B7Z3A6 | Q00796;H0YLA4 |
| Sorcin | SRI | | 0.235801 | -0.172805 | C9J0K6;P30626-2;P30626;P30626-3;B4DHQ6 | C9J0K6;P30626-2;P30626;P30626-3;B4DHQ6 |
| Sorting nexin-12 | SNX12 | | 0.383613 | -0.299981 | Q9UMY4-2;Q9UMY4;Q9UMY4-3 | Q9UMY4-2;Q9UMY4;Q9UMY4-3 |
| Sorting nexin-2 | SNX2 | | 0.316718 | 0.51787 | O60749;O60749-2;D6RC15 | O60749;O60749-2 |
| Sorting nexin-3 | SNX3 | | 0.182921 | 0.277608 | O60493;O60493-2;O60493-4;O60493-3 | O60493;O60493-2;O60493-4;O60493-3 |
| Spectrin alpha chain, non-erythrocytic 1 | SPTAN1 | | 0.137471 | 0.123239 | Q13813;Q13813-3;A6NG51;Q13813-2 | Q13813;Q13813-3;A6NG51;Q13813-2 |
| Spectrin beta chain, non-erythrocytic 1 | SPTBN1 | | 0.163325 | 0.154855 | Q01082;Q01082-2;F8W6C1;E9PJZ2;REV__B4DHX2 | Q01082;Q01082-2 |
| Spectrin beta chain, non-erythrocytic 1 | SPTBN1 | | 0.857369 | 0.561373 | Q01082-3 | Q01082-3 |
| Spermine synthase | SMS | | 0.246709 | -0.164271 | P52788;P52788-2;B4DE40;H7C2R7 | P52788;P52788-2;B4DE40 |
| S-phase kinase-associated protein 1 | SKP1 | | 0.810299 | -0.871315 | E5RJR5;P63208;E7ERH2;F8W8N3;P63208-2;E5RGM3 | E5RJR5;P63208;E7ERH2;F8W8N3;P63208-2;E5RGM3 |
| Sphingomyelin phosphodiesterase | SMPD1 | + | 1.37313 | 0.415834 | P17405-4;P17405;E7EWE2;G3XAB5;P17405-3;P17405-2;H0YEP5;G3V1 | P17405-4;P17405;E7EWE2;G3XAB5;P17405-3;P17405-2;H0YEP5 |
| Src substrate cortactin | CTTN | + | 1.56866 | 1.26369 | Q14247;B4E358;E9PP90;E9PKG3;H0YCD9;H0YEV2 | Q14247 |
| STAM-binding protein | STAMBP | | 0.202929 | 0.135331 | O95630;C9JK83;F5H5B9;C9JEK5;C9JZ93 | O95630;C9JK83;F5H5B9 |
| Stanniocalcin-1 | STC1 | | 1.18113 | -1.8361 | P52823;B4DN22 | P52823;B4DN22 |
| Staphylococcal nuclease domain-containing protein 1 | SND1 | | 0.34093 | 0.41125 | Q7KZF4;H7C597 | Q7KZF4 |
| Stathmin | STMN1 | + | 1.78264 | 2.71714 | P16949;P16949-2;A2A2D0;E5RGX5;Q93045;Q93045-2;E5RIR6;E7EVN3; | P16949;P16949-2;A2A2D0 |
| STE20-like serine/threonine-protein kinase | SLK | | 0.0648562 | 0.0597684 | Q9H2G2-2;Q9H2G2 | Q9H2G2-2;Q9H2G2 |
| Steroid receptor RNA activator 1 | SRA1 | | 0.664649 | 0.563786 | Q9HD15;R4GMW4 | Q9HD15 |
| Stress-70 protein, mitochondrial | HSPA9 | | 0.437674 | 0.378874 | P38646;D6RJI2;D6RA73;H0YBG6;H0Y8S0 | P38646 |
| Stromal cell-derived factor 2 | SDF2 | | 0.0663666 | -0.122778 | Q99470;K7ELI9 | Q99470 |
| Stromelysin-1 | MMP3 | + | 1.51777 | -2.02161 | P08254;H7C139 | P08254 |
| Structural maintenance of chromosomes protein 2 | SMC2 | | 0.050427 | 0.0661511 | O95347;O95347-2;Q5T821 | O95347;O95347-2 |
| Structural maintenance of chromosomes protein 3 | SMC3 | | 0.3103 | 0.474691 | Q9UQE7 | Q9UQE7 |
| Succinyl-CoA ligase [ADP/GDP-forming] subunit alpha, mitochondrial | SUCLG1 | | 0.796769 | -0.496578 | P53597;H7C233 | P53597 |
| Sulfhydryl oxidase 1 | QSOX1 | + | 1.34684 | 0.546688 | O00391;O00391-2;A8MXT8 | O00391;O00391-2 |
| Sulfhydryl oxidase 2;Sulfhydryl oxidase | QSOX2 | | 0.0757771 | 0.0482942 | Q6ZRP7;H0Y430 | Q6ZRP7;H0Y430 |
| Superoxide dismutase [Cu-Zn] | SOD1 | | 0.424833 | -0.280469 | P00441;H7BYH4 | P00441;H7BYH4 |
| Superoxide dismutase [Mn], mitochondrial | SOD2 | | 0.253255 | 0.452965 | P04179-4;P04179;F5H3C5;F5H4R2;F5GYZ5;P04179-2;G8JLJ2;P04179-3; | P04179-4;P04179;F5H3C5;F5H4R2;F5GYZ5;P04179-2;G8JLJ2;P04179-3 |
| Sushi repeat-containing protein SRPX2 | SRPX2 | | 1.05916 | -1.41663 | O60687 | O60687 |
| Synaptic vesicle membrane protein VAT-1 homolog | VAT1 | | 0.935446 | -0.688806 | Q99536;B4DPX4;B0AZP7;K7ESA3;K7ERT7;K7ENX2;K7EM19;K7EJM4;K7 | Q99536;B4DPX4;B0AZP7;K7ESA3 |
| Synaptobrevin homolog YKT6 | YKT6 | | 0.117033 | -0.106836 | O15498;B4DR94;H7C3K7 | O15498;B4DR94 |
| Synaptogyrin-2 | SYNGR2 | + | 1.95416 | -0.655949 | O43760;Q3KQZ2;K7ELD9;K7ENG9;K7ELS8;K7EJ35 | O43760;Q3KQZ2;K7ELD9;K7ENG9;K7ELS8;K7EJ35 |
| Syndecan-1 | SDC1 | | 0.2637 | -0.184211 | E9PHH3;H7C1K4;P18827 | E9PHH3;H7C1K4;P18827 |
| Syndecan-4;Syndecan | SDC4 | | 0.0964208 | -0.100564 | P31431;B4E1S6;P31431-2 | P31431;B4E1S6;P31431-2 |
| Syntaxin-7 | STX7 | | 1.00419 | -0.381867 | O15400-2;O15400 | O15400-2;O15400 |
| Syntenin-1 | SDCBP | | 0.76187 | -0.367076 | O00560;O00560-3 | O00560;O00560-3 |
| Syntenin-1 | SDCBP | | 0.86426 | -0.429001 | O00560-2;G5EA09;B4DHN5;E9PBU7 | O00560-2;G5EA09;B4DHN5;E9PBU7 |
| Talin-1 | TLN1 | | 0.178953 | 0.169897 | Q9Y490;Q5TCU6 | Q9Y490;Q5TCU6 |
| Tax1-binding protein 3 | TAX1BP3 | | 0.788137 | 0.666201 | O14907 | O14907 |
| T-cell immunomodulatory protein | ITFG1 | | 0.973602 | 0.749833 | F5GXC5;Q8TB96;H3BUJ1 | F5GXC5;Q8TB96;H3BUJ1 |
| T-complex protein 1 subunit alpha | TCP1 | | 0.344267 | 0.42463 | P17987;E7EQR6;E7ERF2;F5H282;F5GZ03;F5H136;F5H676;F5GZI8;F5H7 | P17987;E7EQR6;E7ERF2;F5H282 |
| T-complex protein 1 subunit beta | CCT2 | | 0.289509 | 0.421413 | P78371;F5GWF6;P78371-2;F8VQ14 | P78371;F5GWF6;P78371-2;F8VQ14 |
| T-complex protein 1 subunit delta | CCT4 | | 0.213734 | 0.354978 | P50991;B7Z9L0;P50991-2;B7Z2F4 | P50991;B7Z9L0;P50991-2;B7Z2F4 |
| T-complex protein 1 subunit epsilon | CCT5 | | 0.37759 | 0.491058 | P48643;E7ENZ3;B7ZAR1;E9PCA1;P48643-2;D6RIZ7;H0Y914 | P48643;E7ENZ3;B7ZAR1;E9PCA1;P48643-2 |
| T-complex protein 1 subunit eta | CCT7 | | 0.17293 | 0.291998 | Q99832;Q99832-3;F5GZK5;Q99832-4;B7Z1C9;Q99832-2;F8WAM2;F8W | Q99832;Q99832-3;F5GZK5;Q99832-4;B7Z1C9;Q99832-2 |
| T-complex protein 1 subunit gamma | CCT3 | | 0.263954 | 0.329992 | P49368;P49368-2;B4DUR8;E9PRC8;Q5SZX6;E9PQ35;E9PM09;Q5SZW8; | P49368;P49368-2;B4DUR8 |
| T-complex protein 1 subunit theta | CCT8 | | 0.121159 | 0.114129 | P50990;P50990-2;P50990-3;H7C4C8;H7C2U0 | P50990;P50990-2;P50990-3 |
| T-complex protein 1 subunit zeta | CCT6A | | 0.382044 | 0.617651 | P40227;B4DPJ8;P40227-2;J3KRI6 | P40227;B4DPJ8;P40227-2 |
| Tenascin | TNC | + | 1.61825 | 1.79038 | P24821;E9PC84;P24821-3;F5H7V9;P24821-6 | P24821;E9PC84;P24821-3;F5H7V9;P24821-6 |
| Tetratricopeptide repeat protein 38 | TTC38 | | 0.180386 | 0.212257 | Q5R3I4;E7ES35;A9UJP8;H7C089;F8WBU8;H7C2L7 | Q5R3I4;E7ES35;A9UJP8 |
| Thiopurine S-methyltransferase | TPMT | | 0.616336 | 1.19247 | P51580 | P51580 |
| Thioredoxin | TXN | | 0.261471 | -0.180824 | P10599;P10599-2 | P10599;P10599-2 |
| Thioredoxin domain-containing protein 16 | TXNDC16 | | 0.866109 | -0.665729 | Q9P2K2 | Q9P2K2 |
| Thioredoxin domain-containing protein 17 | TXNDC17 | | 0.121865 | -0.0819244 | Q9BRA2;I3L2R6;I3L0K2;I3L3M7 | Q9BRA2;I3L2R6;I3L0K2;I3L3M7 |
| Thioredoxin domain-containing protein 5 | TXNDC5 | | 0.137588 | -0.0772298 | Q86UY0;Q8NBS9;Q8NBS9-2 | Q86UY0;Q8NBS9;Q8NBS9-2 |
| Thioredoxin reductase 1, cytoplasmic | TXNRD1;GML | + | 1.33951 | -0.696442 | B7Z2S5;Q16881-4;Q16881-3;Q16881;B2R5P6;E2QRB9;E9PNQ6;E7EW1 | B7Z2S5;Q16881-4;Q16881-3;Q16881;B2R5P6;E2QRB9;E9PNQ6;E7EW1 |
| Thioredoxin-dependent peroxide reductase, mitochondrial | PRDX3 | | 0.773547 | 0.458185 | P30048-2;P30048 | P30048-2;P30048 |
| Thioredoxin-like protein 1 | TXNL1 | | 0.0603398 | -0.067695 | O43396;K7ER96;G3V1K0;K7EML9;K7EPB7;K7EME7;K7EKG2 | O43396;K7ER96;G3V1K0 |
| Thiosulfate sulfurtransferase | TST | | 0.67501 | 0.48297 | Q16762;B1AH48 | Q16762 |
| THO complex subunit 4 | ALYREF | | 0.168962 | -0.506682 | Q86V81;E9PB61 | Q86V81;E9PB61 |
| Threonine--tRNA ligase, cytoplasmic | TARS | | 0.485693 | -0.336568 | P26639;P26639-2;G3XAN9;E7ERI3;D6R9F8;D6RCA5;D6RDJ6;D6RBR8;D | P26639;P26639-2;G3XAN9;E7ERI3 |
| Thrombospondin-1 | THBS1 | | 0.0746153 | -0.0548796 | P07996;A8MZG1;P35442 | P07996 |
| Tissue alpha-L-fucosidase | FUCA1 | | 0.378436 | -0.27929 | P04066 | P04066 |
| Tissue factor | F3 | | 0.430821 | -0.371262 | P13726;P13726-2 | P13726;P13726-2 |
| Tissue factor pathway inhibitor | TFPI | | 0.603044 | 0.215473 | P10646;C9JBB3;P10646-2;C9JKV3;C9JT76;C9JQ14;C9JP39;C9J103 | P10646;C9JBB3;P10646-2;C9JKV3 |
| Tissue-type plasminogen activator;Tissue-type plasminogen activator chain A;Tissue-type plasminogen ac | PLAT | | 0.736107 | -0.962846 | P00750;B4DN26;E7ESF4;B4DNJ1;P00750-4;P00750-3;K4DI88;P00750-2; | P00750;B4DN26;E7ESF4;B4DNJ1;P00750-4;P00750-3 |
| Titin | TTN | | 1.19164 | -0.595146 | Q8WZ42-5;Q8WZ42-11;Q8WZ42-4;Q8WZ42-7;Q8WZ42-2;Q8WZ42;Q8 | Q8WZ42-5;Q8WZ42-11;Q8WZ42-4;Q8WZ42-7;Q8WZ42-2;Q8WZ42;Q8 |
| Toll-interacting protein | TOLLIP | | 0.719359 | 0.582834 | Q9H0E2;B3KR28;E7EN89;F2Z2Y8;B3KXC6;E9PP67;E9PQ25;E9PNS3 | Q9H0E2;B3KR28;E7EN89;F2Z2Y8;B3KXC6;E9PP67;E9PQ25 |

| | | | | | | |
|---|---|---|---|---|---|---|
| Torsin-1A | TOR1A | | 0.0150967 | 0.0134233 | O14656;O14656-2 | O14656 |
| Torsin-1B | TOR1B | | 0.517025 | 0.236609 | O14657;H0Y7C8 | O14657 |
| Torsin-3A | TOR3A | | 0.959446 | 0.87882 | Q9H497;Q5W0C6;Q9H497-2;Q9H497-3 | Q9H497;Q5W0C6;Q9H497-2 |
| Transaldolase | TALDO1 | | 0.264207 | 0.195408 | P37837;F2Z393;E9PM01;E9PKI8 | P37837;F2Z393 |
| Transcription elongation factor B polypeptide 2 | TCEB2 | | 0.48991 | 0.627284 | Q15370;B8ZZU8;Q15370-2;I3L0M9 | Q15370;B8ZZU8;Q15370-2;I3L0M9 |
| Transferrin receptor protein 1;Transferrin receptor protein 1, serum form | TFRC | | 0.787155 | -0.362253 | P02786;G3V0E5;F5H6B1;H7C3V5;F8WBE5;Q9UP52-2;Q9UP52-3;Q9UP5 | P02786;G3V0E5;F5H6B1 |
| Transforming growth factor beta receptor type 3 | TGFBR3 | | 0.697004 | -2.5122 | Q03167-2;Q03167;E9PKY4 | Q03167-2;Q03167 |
| Transforming growth factor beta-1;Latency-associated peptide | TGFB1 | | 0.565392 | -0.664096 | P01137;M0R2S0 | P01137 |
| Transforming growth factor beta-2;Latency-associated peptide | TGFB2 | | 0.0946535 | -0.124942 | P61812;P61812-2 | P61812;P61812-2 |
| Transforming growth factor-beta-induced protein ig-h3 | TGFBI | | 1.15348 | -1.26086 | Q15582;G8JLA8;H0Y8L3;S4R3C6;H0Y9D7;H0Y8M8;H0YAB8;D6RBX4;H0 | Q15582;G8JLA8;H0Y8L3 |
| Transforming protein RhoA | RHOA | | 1.00222 | -0.482029 | P61586;C9JNR4;C9JX21;C9J1T2;C9JRM1 | P61586;C9JNR4;C9JX21 |
| Transgelin-2 | TAGLN2 | | 0.354854 | -0.24593 | P37802;C9J5W6;Q9UI15 | P37802 |
| Transitional endoplasmic reticulum ATPase | VCP | | 0.706709 | -0.497718 | P55072;C9IZA5;C9JUP7;K7EM02;K7EIJ8;Q8IYT4-2;Q8IYT4 | P55072 |
| Transketolase | TKT | | 0.508858 | -0.241638 | P29401;P29401-2;B4E022;E9PFF2;F8W888;F8WAX4;Q5TYJ8;P51854-1;I | P29401;P29401-2;B4E022 |
| Translationally-controlled tumor protein | TPT1 | | 0.0433517 | -0.0401488 | P13693;J3KPG2;Q5W0H4;P13693-2;E9PJF7;H0YCX0;Q9HAU6;Q56UQ5 | P13693;J3KPG2;Q5W0H4;P13693-2;E9PJF7 |
| Translocon-associated protein subunit delta | SSR4 | | 0.329578 | -0.21016 | P51571;A6NLM8 | P51571 |
| Transmembrane 9 superfamily member 2 | TM9SF2 | | 0.808491 | -1.17136 | Q99805 | Q99805 |
| Transmembrane emp24 domain-containing protein 10 | TMED10 | + | 1.59789 | -0.764146 | P49755;G3V2K7 | P49755;G3V2K7 |
| Transmembrane protein 109 | TMEM109 | | 0.921177 | 0.317019 | Q9BVC6 | Q9BVC6 |
| Transmembrane protein 132A | TMEM132A | + | 1.30481 | -1.84562 | Q24JP5;Q24JP5-2;Q24JP5-4;Q24JP5-3;H0YFQ2;H0YFB3;F5H765;H0YFQ | Q24JP5;Q24JP5-2;Q24JP5-4 |
| Transmembrane protein 2 | TMEM2 | | 0.867717 | 1.33855 | B4E1B9;Q9UHN6-2;Q9UHN6 | B4E1B9;Q9UHN6-2;Q9UHN6 |
| Transmembrane protein 59 | TMEM59 | | 0.171087 | -0.096702 | Q5T6Z8;Q9BXS4 | Q5T6Z8;Q9BXS4 |
| Transportin-1 | TNPO1 | | 0.250849 | -0.23384 | Q92973-2;Q92973;Q92973-3;S4R398;B4DSC0;E7EW37;K7ENW1;K7ESC | Q92973-2;Q92973;Q92973-3 |
| Trifunctional purine biosynthetic protein adenosine-3;Phosphoribosylamine--glycine ligase;Phosphoribos | GART | | 0.142323 | 0.135529 | P22102;P22102-2;F8WD69;B4DJ93;C9JBJ1;C9JTV6;C9JZG2;C9JKQ7;H7C | P22102;P22102-2 |
| Triosephosphate isomerase | TPI1 | | 1.01039 | -0.643715 | P60174-1;P60174;P60174-4;U3KPZ0;U3KQF3;U3KPS5 | P60174-1;P60174;P60174-4;U3KPZ0 |
| Tripeptidyl-peptidase 1 | TPP1 | | 0.069434 | 0.0528069 | O14773;O14773-2;E7EV34 | O14773 |
| Tropomyosin alpha-3 chain | TPM3 | | 0.589827 | 0.769076 | J3KN67;P06753;P06753-7;H0YL80;H0YKP3 | J3KN67;P06753;P06753-7 |
| Tropomyosin alpha-3 chain | TPM3 | | 0.348266 | 0.400577 | P06753-2;Q5VU59;P06753-3;P06753-5;P06753-6;P06753-4;Q5VU61;D6 | P06753-2;Q5VU59;P06753-3;P06753-5;P06753-6;P06753-4;Q5VU61 |
| Tropomyosin alpha-4 chain | TPM4 | | 0.183883 | 0.185934 | P67936 | P67936 |
| Trypsin-2;Putative trypsin-6 | PRSS2;PRSS3P2 | | 0.523889 | 0.595777 | P07478;Q8NHM4 | P07478;Q8NHM4 |
| Tryptophan--tRNA ligase, cytoplasmic;T1-TrpRS;T2-TrpRS | WARS | | 0.246112 | 0.336684 | P23381;P23381-2;G3V3H8;G3V3Y5;G3V277;H0YJP3;G3V423;G3V5U1;G | P23381;P23381-2 |
| Tsukushin | TSKU | | 0.217978 | -0.745193 | Q8WUA8;E9PLG7 | Q8WUA8 |
| Tubulin alpha-1B chain | TUBA1B | | 0.0728394 | 0.0369517 | P68363 | P68363 |
| Tubulin alpha-4A chain | TUBA4A | | 0.816248 | 1.7324 | A8MUB1;P68366;C9JEV8;C9JQ00;C9JJQ8;C9JDS9;C9JDL2 | A8MUB1;P68366 |
| Tubulin beta chain | TUBB | | 1.05323 | 0.673617 | P07437;Q5JP53;Q5ST81;E9PBJ4;F5H0I4;Q5SQY0;I3L2F9;A6NNZ2;M0R0 | P07437;Q5JP53;Q5ST81 |
| Tubulin beta-2A chain | TUBB2A | + | 1.34822 | 1.11541 | Q13885 | Q13885 |
| Tubulin beta-3 chain | TUBB3 | | 0.72007 | 0.759343 | Q13509;Q13509-2;G3V5W4;G3V2R8;G3V3R4;G3V2N6;G3V2A3;G3V3J6 | Q13509;Q13509-2 |
| Tubulin beta-4B chain | TUBB4B | | 0.876608 | 0.659419 | P68371;Q3ZCM7;M0R2D3;M0QY85;M0R0X0;M0QY37;M0QX14;M0QZL | P68371 |
| Tubulin beta-6 chain | TUBB6 | | 0.698438 | 0.451452 | Q9BUF5;K7ESM5;K7EL29;K7EN98;K7ESQ3;K7EQT3;K7EJ64;K7ERA8;K7E | Q9BUF5;K7ESM5 |
| Tubulin-specific chaperone A | TBCA | | 0.43126 | -0.270261 | E5RIW3;E5RJD8;O75347;E5RHG6;B4DT30;E5RIX8 | E5RIW3;E5RJD8;O75347;E5RHG6;B4DT30;E5RIX8 |
| Tubulointerstitial nephritis antigen-like | TINAGL1 | + | 1.41562 | -0.935264 | Q9GZM7-3;Q9GZM7;E7EUP2;Q9GZM7-2;B4DNS5 | Q9GZM7-3;Q9GZM7 |
| Tumor necrosis factor receptor superfamily member 1A;Tumor necrosis factor receptor superfamily mem | TNFRSF1A | | 0.45953 | -1.21387 | P19438;F5H6V7;F5H8A6;P19438-5;P19438-4;F5H061;P19438-2;F5H6Z2 | P19438;F5H6V7;F5H8A6;P19438-5;P19438-4;F5H061;P19438-2 |
| Tumor susceptibility gene 101 protein | TSG101 | | 1.03106 | -0.382482 | Q99816;F5H442;Q99816-2;J3QRU6;J3QKS4 | Q99816;F5H442;Q99816-2 |
| Tumor-associated calcium signal transducer 2 | TACSTD2 | | 0.261939 | 0.211914 | P09758 | P09758 |
| Twinfilin-2 | TWF2 | | 0.893689 | -0.624065 | Q6IBS0;D6RG15;H0Y858;H7C5I2;Q9NR96-4;F8W645 | Q6IBS0;D6RG15 |
| Twisted gastrulation protein homolog 1 | TWSG1 | | 0.407428 | -0.277754 | Q9GZX9;J3QS03;Q9GZX9-2 | Q9GZX9;J3QS03 |
| Tyrosine-protein kinase CSK | CSK | | 0.0114612 | 0.0174954 | P41240;H3BUM9;H3BU69 | P41240 |
| Tyrosine-protein kinase JAK1 | JAK1 | | 1.19821 | -1.17219 | P23458;K7EJR6;E9PM19;P29597 | P23458 |
| Tyrosine-protein kinase receptor UFO | AXL | | 0.46996 | -0.352125 | P30530-2;P30530;M0R0W6 | P30530-2;P30530;M0R0W6 |
| Tyrosine-protein kinase Yes | YES1 | | 0.841311 | -0.585284 | J3QRU1;P07947 | J3QRU1;P07947 |
| Tyrosine-protein phosphatase non-receptor type 23 | PTPN23 | + | 1.36905 | 0.3248 | B4DST5;Q9H3S7 | B4DST5;Q9H3S7 |
| Tyrosine--tRNA ligase, cytoplasmic;Tyrosine--tRNA ligase, cytoplasmic, N-terminally processed | YARS | | 0.241162 | 0.359395 | P54577 | P54577 |
| U6 snRNA-associated Sm-like protein LSm6 | LSM6 | | 0.400756 | 0.52606 | P62312 | P62312 |
| Ubiquitin carboxyl-terminal hydrolase 14;Ubiquitin carboxyl-terminal hydrolase | USP14 | | 0.727658 | -0.652053 | P54578-2;P54578;A6NJA2;B7Z4N8;J3QQT6;J3KS55;H0YA47 | P54578-2;P54578;A6NJA2;B7Z4N8 |
| Ubiquitin carboxyl-terminal hydrolase isozyme L3 | UCHL3 | | 0.469392 | 0.44782 | P15374;Q5TBK7 | P15374;Q5TBK7 |
| Ubiquitin thioesterase OTUB1 | OTUB1 | | 0.290357 | 0.207034 | Q96FW1;F5GYN4;F5H6Q1;J3KR44;F5GYJ8;F5H3F0;Q96FW1-2 | Q96FW1;F5GYN4;F5H6Q1;J3KR44;F5GYJ8 |
| Ubiquitin-40S ribosomal protein S27a;Ubiquitin;40S ribosomal protein S27a;Ubiquitin-60S ribosomal prot | RPS27A;UBB;UBC;UBA52;UBBI | | 0.361421 | -0.246308 | P62979;J3QS39;J3QTR3;F5H6Q2;F5GYU3;F5H2Z3;F5H265;B4DV12;F5H: | P62979;J3QS39;J3QTR3;F5H6Q2;F5GYU3;F5H2Z3;F5H265;B4DV12;F5H |
| Ubiquitin-conjugating enzyme E2 D2 | UBE2D2 | | 0.00510764 | 0.00411115 | P62837 | P62837 |
| Ubiquitin-conjugating enzyme E2 D3;Ubiquitin-conjugating enzyme E2 D2 | UBE2D3;UBE2D2 | | 0.324718 | 0.372309 | P61077;P61077-2;P61077-3;D6RA11;D6RFM0;D6RAH7;H9KV45;P62837 | P61077;P61077-2;P61077-3;D6RA11;D6RFM0;D6RAH7;H9KV45;P6283 |
| Ubiquitin-conjugating enzyme E2 G1;Ubiquitin-conjugating enzyme E2 G1, N-terminally processed | UBE2G1 | | 0.459183 | -0.362393 | P62253;I3L0Q0;I3L2Q4;I3L2H7;K7EPR7 | P62253;I3L0Q0;I3L2Q4 |
| Ubiquitin-conjugating enzyme E2 K | UBE2K | | 0.293758 | -0.198476 | P61086;B4DIZ2;D6RDM7;P61086-2;P61086-3;D6RFX1 | P61086;B4DIZ2;D6RDM7;P61086-2;P61086-3 |
| Ubiquitin-conjugating enzyme E2 L3 | UBE2L3 | | 0.208696 | -0.140426 | P68036;P68036-2;P61129 | P68036;P68036-2 |
| Ubiquitin-conjugating enzyme E2 N | UBE2N | | 0.225791 | -0.133726 | P61088;F8VQQ8;F8VSD4;F8VV71;F8VZ29 | P61088;F8VQQ8;F8VSD4;F8VV71 |
| Ubiquitin-conjugating enzyme E2 variant 1 | UBE2V1 | | 0.391613 | -0.269891 | Q13404;G3V2F7;Q13404-7;Q13404-2;Q13404-1;Q13404-8;Q13404-6 | Q13404;G3V2F7;Q13404-7;Q13404-2;Q13404-1;Q13404-8;Q13404-6 |
| Ubiquitin-conjugating enzyme E2 variant 2 | UBE2V2 | | 0.537123 | -0.287189 | Q15819;G3V113;H0YBP9;H0YBX6;E5RIF1;D6RG00 | Q15819;G3V113;H0YBP9 |
| Ubiquitin-fold modifier 1 | UFM1 | | 0.00862578 | 0.00828931 | H0Y614;P61960 | H0Y614;P61960 |
| Ubiquitin-fold modifier-conjugating enzyme 1 | UFC1 | | 0.103499 | 0.0961923 | Q9Y3C8 | Q9Y3C8 |
| Ubiquitin-like modifier-activating enzyme 1 | UBA1 | | 0.295634 | -0.317343 | P22314;Q5JRR6;Q5JRR9;Q5JRS0;Q5JRS2;Q5JRS1;Q5JRS3 | P22314 |
| Ubiquitin-like protein ISG15 | ISG15 | + | 2.1754 | -1.16935 | P05161 | P05161 |
| UDP-glucose 4-epimerase | GALE | | 0.622619 | 0.619561 | Q14376;Q5QPP4;Q5QPP3;Q5QPP1;Q5QPP9 | Q14376;Q5QPP4;Q5QPP3;Q5QPP1 |
| UDP-glucose 6-dehydrogenase | UGDH | | 0.0736166 | 0.123701 | O60701;O60701-2;O60701-3;E7ER83;E7ETF4;E7EV97;E7ER95;D6RHF4;I | O60701;O60701-2;O60701-3 |
| UDP-glucose:glycoprotein glucosyltransferase 1 | UGGT1 | | 0.164734 | -0.101742 | Q9NYU2-2;Q9NYU2;H7BZG0;F8WCI2;F8WCE6 | Q9NYU2-2;Q9NYU2 |
| UDP-glucuronic acid decarboxylase 1 | UXS1 | + | 2.06223 | 0.930237 | Q8NBZ7;Q8NBZ7-2;B3KV61;Q8NBZ7-3;C9JCB7;C9JFU6;C9JE50;B4E3U7; | Q8NBZ7;Q8NBZ7-2;B3KV61;Q8NBZ7-3 |
| UMP-CMP kinase | CMPK1 | | 0.231422 | 0.332437 | P30085;Q5T0D2;P30085-2 | P30085;Q5T0D2;P30085-2 |
| Uncharacterized protein C9orf142 | C9orf142 | | 0.389216 | -0.358148 | Q9BUH6;Q9BUH6-2 | Q9BUH6 |
| Unconventional myosin-Ic | MYO1C | | 0.836067 | -0.575696 | F5H6E2;O00159-2;O00159-3;O00159;I3L204;I3L168;I3L3F5;I3L4D4;I3L5 | F5H6E2;O00159-2;O00159-3;O00159 |
| UPF0160 protein MYG1, mitochondrial | C12orf10 | | 0.481579 | 0.64408 | Q9HB07;F8VQQ3;F8VR84;H3BPH3 | Q9HB07;F8VQQ3;F8VR84 |
| UPF0556 protein C19orf10 | C19orf10 | | 0.101531 | -0.0700821 | Q969H8;M0QXF7;M0QYN0 | Q969H8;M0QXF7;M0QYN0 |
| UPF0587 protein C1orf123 | C1orf123 | | 0.18248 | 0.149012 | Q9NWV4;B1ARP7 | Q9NWV4 |

| | | | | | | |
|---|---|---|---|---|---|---|
| Urokinase plasminogen activator surface receptor | PLAUR | | 0.694673 | -0.35741 | Q03405;M0R1I2;Q03405-3;M0QYR6;M0R0H5;M0R0Y4;M0R2E9;M0QY | Q03405;M0R1I2;Q03405-3;M0QYR6;M0R0H5;M0R0Y4;M0R2E9 |
| Urokinase-type plasminogen activator;Urokinase-type plasminogen activator long chain A;Urokinase-type | PLAU | | 0.911312 | -0.844747 | E7ET40;P00749-2;P00749;E7ESM2;S4R3G7 | E7ET40;P00749-2;P00749;E7ESM2 |
| UTP--glucose-1-phosphate uridylyltransferase | UGP2 | | 0.104975 | -0.0876808 | Q16851-2;C9JNZ1;C9JVG3;C9JWG0;C9JUW1;C9JTZ5;F2Z3H1;C9J3M0;F2 | Q16851-2;C9JNZ1 |
| Vacuolar protein sorting-associated protein 26A | VPS26A | | 0.0398308 | 0.0281954 | O75436;F5H4L7;G3V1N8;S4R3Q6;O75436-2;S4R2Y3 | O75436;F5H4L7;G3V1N8;S4R3Q6;O75436-2 |
| Vacuolar protein sorting-associated protein 28 homolog | VPS28 | + | 1.4932 | 0.425277 | Q9UK41;E9PQR7;Q9UK41-2;E9PM90;E9PI55;E9PR04;E9PLM9 | Q9UK41;E9PQR7;Q9UK41-2;E9PM90 |
| Vacuolar protein sorting-associated protein 29 | VPS29 | | 0.0286727 | 0.0327723 | Q9UBQ0;F8VXU5;Q9UBQ0-2 | Q9UBQ0;F8VXU5;Q9UBQ0-2 |
| Vacuolar protein sorting-associated protein 35 | VPS35 | | 0.14884 | 0.190012 | Q96QK1;I3L4S0;I3L4P4 | Q96QK1 |
| Vacuolar protein sorting-associated protein 37B | VPS37B | | 0.441796 | 0.218454 | Q9H9H4;F5H4M0;F5H1F6 | Q9H9H4;F5H4M0;F5H1F6 |
| Vacuolar protein sorting-associated protein 4A | VPS4A | | 0.201968 | -0.194529 | Q9UN37;I3L4J1;Q6PIW4-2;Q6PIW4 | Q9UN37 |
| Vacuolar protein sorting-associated protein VTA1 homolog | VTA1 | | 0.0846675 | 0.100453 | Q9NP79;Q5TGM0;E7ETQ7 | Q9NP79;Q5TGM0;E7ETQ7 |
| Vacuolar protein-sorting-associated protein 25 | VPS25 | | 0.159381 | -0.261107 | Q9BRG1;K7ENE3;K7EKV4;K7EP45 | Q9BRG1;K7ENE3 |
| Vacuolar-sorting protein SNF8 | SNF8 | | 0.0653756 | 0.0485667 | Q96H20-2;Q96H20;H0Y8S5;D6RBI1;D6RJ86;D6RFY6 | Q96H20-2;Q96H20;H0Y8S5;D6RBI1;D6RJ86;D6RFY6 |
| Vascular endothelial growth factor A | VEGFA | | 0.244718 | -0.0869885 | A2A2V4;H3BLW8;P15692-8;P15692-4;P15692-15;P15692-11;J3KPA4;P1 | A2A2V4;H3BLW8;P15692-8;P15692-4;P15692-15;P15692-11;J3KPA4;P |
| Vascular endothelial growth factor C | VEGFC | + | 1.42687 | 0.526829 | P49767 | P49767 |
| Vasodilator-stimulated phosphoprotein | VASP | | 0.337027 | 0.393793 | P50552;K7EM16;K7ENL7;K7ENR7;K7EQD0;K7EIG8 | P50552 |
| Vasorin | VASN | | 0.276717 | -0.319999 | Q6EMK4 | Q6EMK4 |
| Very low-density lipoprotein receptor | VLDLR | | 0.603569 | -0.790874 | P98155-2;P98155 | P98155-2;P98155 |
| Vesicle-associated membrane protein 7 | VAMP7 | | 0.0868588 | -0.130037 | P51809;P51809-2;P51809-3 | P51809;P51809-2;P51809-3 |
| Vesicle-fusing ATPase | NSF | | 0.501794 | 1.73282 | I3L0N3;B4DFA2;P46459;I3L2G1;K7EQD6 | I3L0N3;B4DFA2;P46459 |
| Vesicular integral-membrane protein VIP36 | LMAN2 | | 0.713272 | -1.2492 | D6RBV2;Q12907;D6RIU4;D6RDX1;D6RBH1 | D6RBV2;Q12907;D6RIU4;D6RDX1 |
| Vigilin | HDLBP | | 0.929882 | 1.44444 | Q00341;Q00341-2;H0Y394;H7C0A4;C9JIZ1;C9J5E5;C9JZI8;H7BZC3;C9JT | Q00341;Q00341-2;H0Y394 |
| Vimentin | VIM | + | 1.77788 | 1.79032 | P08670;B0YJC4;B0YJC5;Q5JVS8;H7C5W5;P17661;F8W835;Q16352;B4D | P08670;B0YJC4 |
| Vinculin | VCL | | 0.242963 | -0.171288 | P18206-2;P18206;Q5JQ13;B4DTM7;P18206-3 | P18206-2;P18206;Q5JQ13 |
| Vitamin K-dependent protein S | PROS1 | | 0.154626 | -0.186957 | P07225;G5E9F8;H7BXT0 | P07225;G5E9F8 |
| Voltage-dependent anion-selective channel protein 1 | VDAC1 | | 0.606821 | -1.31488 | P21796;C9JI87 | P21796;C9JI87 |
| Voltage-dependent anion-selective channel protein 2 | VDAC2 | | 0.381309 | -0.669625 | P45880-2;P45880;P45880-1;Q5JSD2;Q5JSD1;A2A3S1 | P45880-2;P45880;P45880-1;Q5JSD2;Q5JSD1 |
| V-type proton ATPase catalytic subunit A | ATP6V1A | | 0.0246647 | -0.0303817 | B7Z1R5;P38606;C9JA17;C9JVW8 | B7Z1R5;P38606 |
| V-type proton ATPase subunit B, brain isoform | ATP6V1B2 | | 0.314782 | 0.331168 | P21281;H0YC04;E5RGH6;C9JL73;P15313;H0YAT8;C9J5E3;C9JNS9;H0YC | P21281;H0YC04 |
| V-type proton ATPase subunit C 1 | ATP6V1C1 | | 0.637572 | -0.512082 | P21283;E7EV59;Q8NEY4-2;Q8NEY4 | P21283;E7EV59 |
| V-type proton ATPase subunit D | ATP6V1D | | 0.028022 | 0.0434691 | Q9Y5K8;G3V2S6;G3V2V6;H0YJH8;G3V559;H0YJS0;H0YJ55;G3V3T8;G3V | Q9Y5K8;G3V2S6;G3V2V6;H0YJH8;G3V559 |
| V-type proton ATPase subunit d 1 | ATP6V0D1 | | 0.086661 | 0.258442 | J3QL14;R4GN72;F5GYQ1;P61421 | J3QL14;R4GN72;F5GYQ1;P61421 |
| V-type proton ATPase subunit E 1 | ATP6V1E1 | | 0.571937 | 0.578182 | P36543-2;P36543;P36543-3;C9J8H1 | P36543-2;P36543;P36543-3;C9J8H1 |
| V-type proton ATPase subunit G 1 | ATP6V1G1 | | 0.661293 | 0.993431 | O75348;F2Z307;O95670-2;O95670 | O75348 |
| V-type proton ATPase subunit S1 | ATP6AP1 | + | 1.4687 | 3.87123 | Q15904;A6QRJ1;A6NLC6;H7C2Y8 | Q15904;A6QRJ1 |
| WD repeat-containing protein 1 | WDR1 | | 0.708056 | -0.352367 | O75083;O75083-3;D6RD66 | O75083;O75083-3;D6RD66 |
| Wiskott-Aldrich syndrome protein family member 2 | WASF2 | | 0.0416278 | 0.0347166 | Q9Y6W5;Q9Y6W5-2;Q9UPY6 | Q9Y6W5;Q9Y6W5-2 |
| Xaa-Pro aminopeptidase 1 | XPNPEP1 | | 0.548508 | 0.597601 | Q9NQW7-3;Q9NQW7;Q5T6H7;G8JLB2;Q9NQW7-2;Q5T6H2;Q5T6H3 | Q9NQW7-3;Q9NQW7;Q5T6H7;G8JLB2;Q9NQW7-2 |
| Xaa-Pro dipeptidase | PEPD | | 0.0253249 | 0.04159 | P12955;P12955-3;P12955-2;K7ES25;V9GY89;V9GYL0;V9GYE4 | P12955;P12955-3;P12955-2 |
| Zymogen granule protein 16 homolog B | ZG16B | | 0.64699 | -0.735983 | Q96DA0;I3L1H9;I3L3X0 | Q96DA0 |

**Sup. Table 3: MDA-MB-231 1D annotation enrichment (shNODAL versus shControl)**

| C: Type | C: Name | N: Size | N: Score | N: P value | N: Benj. Hoch. FDR | N: Mean | N: Median |
|---|---|---|---|---|---|---|---|
| **GOBP name** | **mRNA metabolic process** | **27** | **0.631189** | **5.96E-07** | **0.000106633** | **1.51085** | **1.44988** |
| GOCC name | cytosolic small ribosomal subunit | 19 | 0.604497 | 2.96E-05 | 0.000888652 | 1.50826 | 1.48451 |
| GOCC name | small ribosomal subunit | 19 | 0.604497 | 2.96E-05 | 0.000799787 | 1.50826 | 1.48451 |
| GOBP name | cellular macromolecular complex disassembly | 27 | 0.62963 | 6.36E-07 | 0.000104916 | 1.49754 | 1.44988 |
| GOBP name | cellular protein complex disassembly | 27 | 0.62963 | 6.36E-07 | 9.74E-05 | 1.49754 | 1.44988 |
| **GOBP name** | **macromolecular complex disassembly** | **27** | **0.62963** | **6.36E-07** | **9.09E-05** | **1.49754** | **1.44988** |
| GOBP name | protein complex disassembly | 27 | 0.62963 | 6.36E-07 | 8.52E-05 | 1.49754 | 1.44988 |
| GOBP name | establishment of protein localization in endoplasmic reticulum | 27 | 0.624951 | 7.69E-07 | 8.69E-05 | 1.46691 | 1.44988 |
| GOBP name | protein targeting to ER | 27 | 0.624951 | 7.69E-07 | 8.25E-05 | 1.46691 | 1.44988 |
| GOBP name | protein targeting to membrane | 27 | 0.624951 | 7.69E-07 | 7.86E-05 | 1.46691 | 1.44988 |
| GOCC name | ribonucleoprotein complex | 27 | 0.616374 | 1.09E-06 | 2.93E-04 | 1.45947 | 1.44988 |
| GOBP name | cotranslational protein targeting to membrane | 26 | 0.600962 | 2.75E-06 | 0.000245529 | 1.45063 | 1.44734 |
| GOBP name | mRNA catabolic process | 26 | 0.600962 | 2.75E-06 | 0.000235708 | 1.45063 | 1.44734 |
| GOBP name | nuclear-transcribed mRNA catabolic process | 26 | 0.600962 | 2.75E-06 | 0.000226642 | 1.45063 | 1.44734 |
| GOBP name | nuclear-transcribed mRNA catabolic process, nonsense-mediated decay | 26 | 0.600962 | 2.75E-06 | 0.000218248 | 1.45063 | 1.44734 |
| GOBP name | RNA catabolic process | 26 | 0.600962 | 2.75E-06 | 0.000210453 | 1.45063 | 1.44734 |
| GOBP name | SRP-dependent cotranslational protein targeting to membrane | 26 | 0.600962 | 2.75E-06 | 0.000203196 | 1.45063 | 1.44734 |
| GOBP name | translational initiation | 26 | 0.600962 | 2.75E-06 | 0.000196423 | 1.45063 | 1.44734 |
| GOBP name | translational termination | 26 | 0.600962 | 2.75E-06 | 0.000190087 | 1.45063 | 1.44734 |
| GOBP name | viral transcription | 26 | 0.600962 | 2.75E-06 | 0.000184147 | 1.45063 | 1.44734 |
| GOBP name | viral infectious cycle | 29 | 0.612162 | 6.87E-07 | 8.67E-05 | 1.43345 | 1.44988 |
| **GOBP name** | **establishment of protein localization to organelle** | **28** | **0.601824** | **1.42E-06** | **1.39E-04** | **1.42971** | **1.44734** |
| GOBP name | translation | 27 | 0.586745 | 3.47E-06 | 0.000218967 | 1.4206 | 1.44479 |
| GOBP name | translational elongation | 27 | 0.586745 | 3.47E-06 | 0.000212711 | 1.4206 | 1.44479 |
| GOCC name | ribosome | 22 | 0.538182 | 8.07E-05 | 0.00145274 | 1.40997 | 1.43019 |
| GOBP name | protein targeting | 32 | 0.629167 | 1.34E-07 | 3.20E-05 | 1.3962 | 1.4672 |
| GOBP name | cellular process involved in reproduction | 29 | 0.574342 | 3.19E-06 | 0.000207428 | 1.36856 | 1.44479 |
| GOBP name | viral reproductive process | 30 | 0.571739 | 2.72E-06 | 0.000253735 | 1.3486 | 1.44734 |
| GOBP name | intracellular protein transport | 35 | 0.61642 | 1.08E-07 | 3.32E-05 | 1.3384 | 1.44988 |
| GOBP name | RNA metabolic process | 39 | 0.632994 | 1.86E-08 | 3.99E-05 | 1.33496 | 1.44988 |
| GOBP name | establishment of protein localization | 39 | 0.599011 | 1.02E-07 | 4.39E-05 | 1.30318 | 1.44988 |
| GOBP name | protein transport | 39 | 0.599011 | 1.02E-07 | 3.66E-05 | 1.30318 | 1.44988 |
| GOBP name | viral reproduction | 34 | 0.590909 | 4.47E-07 | 8.72E-05 | 1.30171 | 1.4672 |
| GOBP name | intracellular transport | 39 | 0.629904 | 2.18E-08 | 2.34E-05 | 1.3014 | 1.48451 |
| GOBP name | gene expression | 36 | 0.609819 | 1.16E-07 | 3.12E-05 | 1.29989 | 1.4672 |
| GOBP name | RNA biosynthetic process | 38 | 0.604637 | 9.55E-08 | 5.13E-05 | 1.28913 | 1.44734 |
| GOBP name | cellular macromolecular complex subunit organization | 33 | 0.524685 | 8.96E-06 | 0.000480895 | 1.2408 | 1.41558 |
| GOBP name | nucleic acid metabolic process | 42 | 0.59881 | 5.93E-08 | 4.24E-05 | 1.23686 | 1.44734 |
| GOBP name | cellular macromolecule biosynthetic process | 34 | 0.490642 | 2.77E-05 | 0.00132244 | 1.20683 | 1.39853 |

| | | | | | | | |
|---|---|---|---|---|---|---|---|
| GOBP name | cellular macromolecule catabolic process | 31 | 0.476072 | 7.85E-05 | 0.00336888 | 1.19729 | 1.41558 |
| GOCC name | cell-substrate adherens junction | 31 | 0.51223 | 2.15E-05 | 0.00145019 | 1.13337 | 1.5061 |
| GOCC name | cell-substrate junction | 31 | 0.51223 | 2.15E-05 | 0.00116015 | 1.13337 | 1.5061 |
| GOCC name | focal adhesion | 31 | 0.51223 | 2.15E-05 | 0.000966793 | 1.13337 | 1.5061 |
| GOBP name | nucleobase-containing compound metabolic process | 47 | 0.56539 | 1.58E-07 | 3.40E-05 | 1.1227 | 1.41558 |
| GOBP name | reproductive process | 38 | 0.472431 | 3.07E-05 | 0.00143052 | 1.11876 | 1.39853 |
| GOCC name | adherens junction | 33 | 0.499489 | 2.36E-05 | 0.000911349 | 1.09842 | 1.5061 |
| GOCC name | cell junction | 37 | 0.497615 | 1.31E-05 | 0.00117996 | 1.07178 | 1.48451 |
| GOBP name | macromolecule biosynthetic process | 36 | 0.429587 | 0.000189205 | 0.00780833 | 1.06617 | 1.36599 |
| GOBP name | macromolecule catabolic process | 35 | 0.418719 | 0.000307993 | 0.0118027 | 1.06566 | 1.38809 |
| GOCC name | anchoring junction | 34 | 0.481283 | 3.93E-05 | 0.00088485 | 1.05894 | 1.4953 |
| GOBP name | cellular nitrogen compound metabolic process | 50 | 0.527778 | 7.59E-07 | 9.05E-05 | 1.03659 | 1.40054 |
| GOBP name | macromolecular complex subunit organization | 42 | 0.504167 | 5.02E-06 | 0.000291 | 1.03355 | 1.41228 |
| GOBP name | protein complex subunit organization | 42 | 0.504167 | 5.02E-06 | 0.000283343 | 1.03355 | 1.41228 |
| GOBP name | cellular component disassembly | 36 | 0.455426 | 7.57E-05 | 0.00338539 | 1.00592 | 1.43019 |
| GOBP name | cellular component disassembly at cellular level | 36 | 0.455426 | 7.57E-05 | 0.0033163 | 1.00592 | 1.43019 |
| GOBP name | establishment of localization in cell | 47 | 0.49844 | 3.81E-06 | 0.000226902 | 0.999501 | 1.41558 |
| GOBP name | cellular biosynthetic process | 49 | 0.473302 | 9.83E-06 | 0.000514592 | 0.979238 | 1.3439 |
| GOBP name | cellular catabolic process | 39 | 0.40068 | 0.000370546 | 0.0139507 | 0.974895 | 1.38809 |
| GOBP name | biosynthetic process | 50 | 0.466111 | 1.26E-05 | 0.00064168 | 0.969066 | 1.32719 |
| GOBP name | nitrogen compound metabolic process | 52 | 0.484066 | 5.09E-06 | 0.000280094 | 0.945762 | 1.39009 |
| GOBP name | catabolic process | 44 | 0.377622 | 0.000550433 | 0.0187497 | 0.917693 | 1.35564 |
| GOBP name | establishment of localization | 53 | 0.455291 | 1.71E-05 | 0.00085273 | 0.91126 | 1.40898 |
| GOBP name | transport | 53 | 0.455291 | 1.71E-05 | 0.00083335 | 0.91126 | 1.40898 |
| GOCC name | cytoplasm | 53 | 0.442713 | 2.90E-05 | 0.000979017 | 0.892769 | 1.36738 |
| GOCC name | cytosol | 56 | 0.434524 | 3.70E-05 | 0.000907597 | 0.837081 | 1.39853 |
| GOCC name | membrane | 74 | 0.518018 | 1.42E-06 | 1.92E-04 | 0.817715 | 1.28708 |
| GOCC name | macromolecular complex | 55 | 0.400814 | 0.000144766 | 0.00229923 | 0.814685 | 1.38809 |
| GOBP name | cellular macromolecule metabolic process | 66 | 0.419913 | 6.69E-05 | 0.00305673 | 0.774519 | 1.1932 |
| GOBP name | macromolecule metabolic process | 73 | 0.414593 | 0.000107731 | 0.00453316 | 0.669204 | 1.19157 |
| GOBP name | cellular metabolic process | 74 | 0.369932 | 0.000574795 | 0.0192736 | 0.668922 | 1.13132 |
| GOCC name | intracellular organelle | 74 | 0.354167 | 0.000978999 | 0.0125871 | 0.664588 | 1.13132 |
| GOCC name | intracellular organelle part | 82 | 0.44939 | 5.83E-05 | 0.00121146 | 0.660287 | 1.13132 |
| GOCC name | organelle part | 82 | 0.44939 | 5.83E-05 | 0.00112493 | 0.660287 | 1.13132 |
| GOCC name | cytoplasmic part | 91 | 0.449132 | 0.000194938 | 0.00292407 | 0.578454 | 1.07454 |
| GOCC name | membrane-bounded vesicle | 96 | 0.453526 | 0.000402246 | 0.00571612 | 0.522247 | 1.04415 |
| GOCC name | vesicle | 96 | 0.453526 | 0.000402246 | 0.00543032 | 0.522247 | 1.04415 |
| GOCC name | extracellular region | 34 | -0.457219 | 9.39E-05 | 0.0015846 | -0.614368 | -0.726891 |
| **GOBP name** | **positive regulation of cell migration** | **10** | **-0.678571** | **0.000390231** | **0.0144386** | **-1.46546** | **-1.30838** |
| GOBP name | positive regulation of cell motility | 10 | -0.678571 | 0.000390231 | 0.0141938 | -1.46546 | -1.30838 |
| GOBP name | positive regulation of locomotion | 10 | -0.678571 | 0.000390231 | 0.0139573 | -1.46546 | -1.30838 |

| | | | | | | | |
|---|---|---|---|---|---|---|---|
| **GOBP name** | **inflammatory response** | **9** | **-0.706981** | **0.000429785** | **0.01512** | **-1.6271** | **-1.4445** |
| GOBP name | positive regulation of leukocyte migration | 7 | -0.781366 | 0.000535642 | 0.0185401 | -1.91453 | -2.05726 |
| GOBP name | cellular response to cytokine stimulus | 6 | -0.876437 | 0.000304962 | 0.0123481 | -2.28263 | -2.23478 |
| **GOBP name** | **cytokine-mediated signaling pathway** | **6** | **-0.876437** | **0.000304962** | **0.0121194** | **-2.28263** | **-2.23478** |
| GOBP name | response to cytokine stimulus | 6 | -0.876437 | 0.000304962 | 0.0118991 | -2.28263 | -2.23478 |
| GOBP name | response to molecule of bacterial origin | 5 | -0.907692 | 0.000606565 | 0.0197225 | -2.50529 | -2.41229 |

**Sup. Table 4: Proteins differentially expressed between CM from NODAL and GFP expressing SUM149 cells**

| Protein names | Gene names | T-test Significant | -Log T-test p-value | Log2 T-test fold-change (NODAL-GFP) | Protein IDs | Majority protein IDs |
|---|---|---|---|---|---|---|
| C-X-C motif chemokine 3;GRO-gamma(5-73) | CXCL3 | + | 4.45583 | -10.034 | P19876 | P19876 |
| C-C motif chemokine 20;CCL20(1-67);CCL20(1-64);CCL20(2-70) | CCL20 | + | 4.25786 | -7.46399 | P78556-2;P78556 | P78556-2;P78556 |
| Protein S100-A8;Protein S100-A8, N-terminally processed | S100A8 | + | 2.25063 | -7.11032 | P05109 | P05109 |
| Protein S100-A7 | S100A7 | + | 2.08301 | -6.69118 | P31151;Q86SG5 | P31151 |
| Interleukin-8;MDNCF-a;Interleukin-8;IL-8(5-77);IL-8(6-77);IL-8(7-77);IL-8(8-77);IL-8(9-77 | CXCL8 | | 1.08917 | -6.21022 | P10145;P10145-2 | P10145;P10145-2 |
| Protein S100-A9 | S100A9 | + | 2.65623 | -5.81834 | P06702 | P06702 |
| Platelet basic protein;Connective tissue-activating peptide III;TC-2;Connective tissue-ac | PPBP | + | 4.33043 | -5.49326 | P02775 | P02775 |
| Cadherin-2 | CDH2 | | 0.833356 | -4.98238 | P19022-2 | P19022-2 |
| C-X-C motif chemokine 5;ENA-78(8-78);ENA-78(9-78) | CXCL5 | + | 3.07208 | -4.81573 | P42830 | P42830 |
| Leukemia inhibitory factor | LIF | + | 2.70781 | -4.65854 | P15018 | P15018 |
| Matrix metalloproteinase-9;67 kDa matrix metalloproteinase-9;82 kDa matrix metallop | MMP9 | + | 3.2615 | -4.62235 | P14780 | P14780 |
| Ras GTPase-activating-like protein IQGAP2 | IQGAP2 | + | 3.8339 | -4.47783 | Q13576-3;Q13576-2;Q13576 | Q13576-3;Q13576-2;Q13576 |
| Complement factor B;Complement factor B Ba fragment;Complement factor B Bb fragn | CFB | + | 3.20929 | -4.44468 | P00751;P00751-2 | P00751 |
| Annexin A6 | ANXA6 | | 0.973813 | -4.24747 | P08133-2;P08133 | P08133-2;P08133 |
| Growth-regulated alpha protein;GRO-alpha(4-73);GRO-alpha(5-73);GRO-alpha(6-73) | CXCL1 | + | 3.07163 | -4.13373 | P09341 | P09341 |
| Plasminogen activator inhibitor 2 | SERPINB2 | + | 2.53899 | -4.1254 | P05120 | P05120 |
| Aldehyde dehydrogenase family 1 member A3 | ALDH1A3 | + | 4.36277 | -3.9218 | P47895;P30837 | P47895 |
| Complement C1s subcomponent;Complement C1s subcomponent heavy chain;Comple | C1S | + | 2.42149 | -3.88904 | P09871 | P09871 |
| Interleukin-6 | IL6 | + | 2.56162 | -3.5127 | P05231 | P05231 |
| Plasminogen activator inhibitor 1 | SERPINE1 | + | 2.01844 | -3.46294 | P05121;P05121-2 | P05121;P05121-2 |
| Platelet factor 4;Platelet factor 4, short form | PF4 | + | 1.72871 | -3.37532 | P02776 | P02776 |
| Aldo-keto reductase family 1 member C3 | AKR1C3 | + | 3.30447 | -3.3372 | P42330;P42330-2;P17516 | P42330;P42330-2 |
| Serpin B4 | SERPINB4 | + | 2.09647 | -3.30859 | P48594 | P48594 |
| Aldo-keto reductase family 1 member C1 | AKR1C1 | | 1.26824 | -3.167 | Q04828;P52895-2 | Q04828 |
| Alpha-1-antichymotrypsin;Alpha-1-antichymotrypsin His-Pro-less | SERPINA3 | + | 1.87698 | -3.16523 | P01011;P01011-2 | P01011 |
| Elafin | PI3 | + | 3.12868 | -3.13577 | P19957 | P19957 |
| Superoxide dismutase [Mn], mitochondrial | SOD2 | + | 2.78079 | -2.97946 | P04179;P04179-4;P04179-2;P04179-3 | P04179;P04179-4;P04179-2;P04179-3 |
| Tumor necrosis factor-inducible gene 6 protein | TNFAIP6 | + | 2.39294 | -2.91023 | P98066 | P98066 |
| Urokinase-type plasminogen activator;Urokinase-type plasminogen activator long chai | PLAU | + | 2.30622 | -2.85086 | P00749-2;P00749 | P00749-2;P00749 |
| Tissue factor pathway inhibitor 2 | TFPI2 | | 1.18945 | -2.52764 | P48307-2;P48307 | P48307-2;P48307 |
| Tenascin | TNC | | 0.780948 | -2.45024 | P24821-4;P24821-2;P24821-6 | P24821-4;P24821-2;P24821-6 |
| Cathepsin S | CTSS | + | 1.58487 | -2.39814 | P25774;P25774-2 | P25774;P25774-2 |
| Transferrin receptor protein 1;Transferrin receptor protein 1, serum form | TFRC | + | 3.86704 | -2.37675 | P02786;Q9UP52-2;Q9UP52-3;Q9UP52 | P02786 |
| Transmembrane protein 132A | TMEM132A | + | 2.72324 | -2.35493 | Q24JP5;Q24JP5-2;Q24JP5-4;Q24JP5-3 | Q24JP5;Q24JP5-2;Q24JP5-4 |
| Ephrin type-B receptor 2 | EPHB2 | + | 2.44148 | -2.32883 | P29323-2;P29323-3;P29323;P54762-5;P54762 | P29323-2;P29323-3;P29323 |
| Neutrophil gelatinase-associated lipocalin | LCN2 | + | 1.61392 | -2.3228 | P80188 | P80188 |
| Serpin B3 | SERPINB3 | + | 1.97883 | -2.28858 | P29508;P29508-2 | P29508;P29508-2 |
| Neuropilin-2 | NRP2 | + | 1.64129 | -2.24013 | O60462-4;O60462-5;O60462-2;O60462-3;O60462;O6046 | O60462-4;O60462-5;O60462-2;O60462-3;O60462;C |
| Serpin B9 | SERPINB9 | + | 1.65968 | -2.22788 | P50453 | P50453 |
| Fructose-1,6-bisphosphatase 1 | FBP1 | + | 2.28452 | -2.22281 | P09467;O00757 | P09467 |
| Plastin-2 | LCP1 | + | 1.32689 | -2.19912 | P13796;P13796-2 | P13796 |
| Monocyte differentiation antigen CD14;Monocyte differentiation antigen CD14, urinary | CD14 | + | 2.32122 | -2.192 | P08571 | P08571 |
| Leukocyte elastase inhibitor | SERPINB1 | + | 4.04465 | -2.15331 | P30740;P30740-2 | P30740;P30740-2 |
| Nicotinamide phosphoribosyltransferase | NAMPT | + | 4.26721 | -2.15057 | P43490 | P43490 |
| Protein Niban | FAM129A | + | 1.95019 | -2.10339 | Q9BZQ8 | Q9BZQ8 |
| NEDD8 | NEDD8 | | 0.507178 | -2.037 | Q15843 | Q15843 |
| Guanine deaminase | GDA | + | 1.82171 | -2.00285 | Q9Y2T3;Q9Y2T3-3;Q9Y2T3-2 | Q9Y2T3;Q9Y2T3-3;Q9Y2T3-2 |
| Hyaluronan-binding protein 2;Hyaluronan-binding protein 2 50 kDa heavy chain;Hyalur | HABP2 | | 0.652081 | -1.9651 | Q14520-2;Q14520 | Q14520-2;Q14520 |
| CD82 antigen | CD82 | + | 2.40692 | -1.94721 | P27701-2;P27701 | P27701-2;P27701 |
| Interleukin-1 receptor antagonist protein | IL1RN | + | 2.73991 | -1.93222 | P18510-2;P18510-4;P18510;P18510-3 | P18510-2;P18510-4;P18510;P18510-3 |
| Plectin | PLEC | | 0.556944 | -1.8835 | Q15149-4;Q15149-7;Q15149-8;Q15149-9;Q15149-5;Q151 | Q15149-4;Q15149-7;Q15149-8;Q15149-9;Q15149-5 |
| Alpha-2-macroglobulin-like protein 1 | A2ML1 | + | 1.3412 | -1.86549 | A8K2U0;A8K2U0-2 | A8K2U0;A8K2U0-2 |
| Aldose reductase | AKR1B1 | + | 1.84506 | -1.85018 | P15121;C9JRZ8;C9JRZ8-2 | P15121 |
| Peflin | PEF1 | | 0.885485 | -1.84253 | Q9UBV8 | Q9UBV8 |
| Aldo-keto reductase family 1 member B10 | AKR1B10 | + | 3.06592 | -1.79265 | O60218 | O60218 |
| Protein jagged-1 | JAG1 | + | 2.73783 | -1.75701 | P78504-2;P78504;Q9Y219-2;Q9Y219 | P78504-2;P78504 |
| Fibronectin;Anastellin;Ugl-Y1;Ugl-Y2;Ugl-Y3 | FN1 | + | 2.04589 | -1.75002 | P02751;P02751-3;P02751-8;P02751-17;P02751-14;P0275 | P02751;P02751-3;P02751-8;P02751-17;P02751-14;I |
| Pappalysin-1 | PAPPA | + | 1.81811 | -1.73146 | Q13219 | Q13219 |
| Laminin subunit gamma-2 | LAMC2 | + | 1.57617 | -1.71596 | Q13753;Q13753-2 | Q13753;Q13753-2 |
| Vacuolar protein sorting-associated protein 28 homolog | VPS28 | | 0.391122 | -1.69068 | Q9UK41;Q9UK41-2 | Q9UK41;Q9UK41-2 |
| Interleukin-6 receptor subunit beta | IL6ST | | 0.616844 | -1.64292 | P40189;P40189-3;P40189-2 | P40189;P40189-3 |
| Protein-lysine 6-oxidase | LOX | + | 2.10284 | -1.63768 | P28300 | P28300 |
| PDZ domain-containing protein 11 | PDZD11 | | 0.518928 | -1.63535 | Q5EBL8;Q5EBL8-2 | Q5EBL8;Q5EBL8-2 |
| Laminin subunit beta-3 | LAMB3 | | 1.14548 | -1.61704 | Q13751 | Q13751 |
| Complement C1r subcomponent;Complement C1r subcomponent heavy chain;Complei | C1R | + | 3.18271 | -1.60561 | P00736 | P00736 |

| | | | | | | |
|---|---|---|---|---|---|---|
| Tumor necrosis factor receptor superfamily member 1A;Tumor necrosis factor receptor | TNFRSF1A | | 0.779294 | -1.55114 | P19438-5;P19438-4;P19438 | P19438-5;P19438-4;P19438 |
| Macrophage colony-stimulating factor 1;Processed macrophage colony-stimulating fact | CSF1 | + | 1.68735 | -1.50366 | P09603;P09603-3;P09603-2 | P09603;P09603-3;P09603-2 |
| Retinol-binding protein 4;Plasma retinol-binding protein(1-182);Plasma retinol-binding | RBP4 | | 0.538458 | -1.50251 | P02753 | P02753 |
| Inhibin beta A chain | INHBA | + | 3.54133 | -1.4892 | P08476 | P08476 |
| Coronin-1A | CORO1A | + | 3.01595 | -1.46872 | P31146 | P31146 |
| Acyl-protein thioesterase 2 | LYPLA2 | | 0.765337 | -1.45742 | O95372 | O95372 |
| Mitogen-activated protein kinase kinase kinase kinase 4 | MAP4K4 | | 0.541727 | -1.44867 | O95819-4;O95819-2;O95819;O95819-5;O95819-3;Q9UKE | O95819-4;O95819-2;O95819;O95819-5;O95819-3 |
| Quinone oxidoreductase PIG3 | TP53I3 | | 0.402445 | -1.44627 | Q53FA7;Q53FA7-2 | Q53FA7 |
| C-X-C motif chemokine 16 | CXCL16 | | 0.986525 | -1.44191 | Q9H2A7 | Q9H2A7 |
| Protein CREG1 | CREG1 | | 0.395443 | -1.42873 | O75629 | O75629 |
| A disintegrin and metalloproteinase with thrombospondin motifs 1 | ADAMTS1 | | 0.692282 | -1.35077 | Q9UHI8;Q8TE58 | Q9UHI8 |
| Interleukin-36 gamma | IL36G | + | 1.58363 | -1.34178 | Q9NZH8;Q9NZH8-2 | Q9NZH8;Q9NZH8-2 |
| Ferritin heavy chain;Ferritin heavy chain, N-terminally processed | FTH1 | | 0.588998 | -1.3389 | P02794 | P02794 |
| Interleukin-1 alpha | IL1A | | 0.725469 | -1.31996 | P01583 | P01583 |
| Latexin | LXN | | 0.718504 | -1.31834 | Q9BS40 | Q9BS40 |
| 5(3)-deoxyribonucleotidase, cytosolic type | NT5C | | 0.557491 | -1.28754 | Q8TCD5;Q8TCD5-2 | Q8TCD5;Q8TCD5-2 |
| Nephronectin | NPNT | + | 1.64682 | -1.28401 | Q6UXI9;Q6UXI9-3;Q6UXI9-6;Q6UXI9-5;Q6UXI9-4;Q6UXI9 | Q6UXI9;Q6UXI9-3;Q6UXI9-6;Q6UXI9-5;Q6UXI9-4;Q |
| Ester hydrolase C11orf54 | C11orf54 | + | 2.48134 | -1.27358 | Q9H0W9-2;Q9H0W9;Q9H0W9-3;Q9H0W9-4 | Q9H0W9-2;Q9H0W9;Q9H0W9-3;Q9H0W9-4 |
| Cytoplasmic aconitate hydratase | ACO1 | + | 2.50127 | -1.27083 | P21399 | P21399 |
| Argininosuccinate synthase | ASS1 | + | 1.85097 | -1.26313 | P00966 | P00966 |
| Urokinase plasminogen activator surface receptor | PLAUR | + | 2.84352 | -1.25309 | Q03405;Q03405-2;Q03405-3 | Q03405;Q03405-2;Q03405-3 |
| Syntaxin-binding protein 1 | STXBP1 | | 0.492179 | -1.24743 | P61764;P61764-2 | P61764;P61764-2 |
| Serine/threonine-protein phosphatase 2B catalytic subunit gamma isoform | PPP3CC | + | 2.09101 | -1.24593 | P48454-2;P48454;P48454-3 | P48454-2;P48454;P48454-3 |
| Intersectin-2 | ITSN2 | | 0.440196 | -1.2445 | Q9NZM3-4;Q9NZM3-3;Q9NZM3-2;Q9NZM3 | Q9NZM3-4;Q9NZM3-3;Q9NZM3-2;Q9NZM3 |
| Reticulon-3 | RTN3 | | 0.382061 | -1.24388 | O95197-3;O95197-4;O95197-7;O95197-2;O95197 | O95197-3;O95197-4;O95197-7;O95197-2;O95197 |
| Alpha-1-antitrypsin;Short peptide from AAT | SERPINA1 | + | 2.12193 | -1.23317 | P01009;P01009-2;P01009-3 | P01009;P01009-2;P01009-3 |
| Cystatin-M | CST6 | + | 1.41089 | -1.23042 | Q15828 | Q15828 |
| Nectin-2 | PVRL2 | | 0.395 | -1.22643 | Q92692-2;Q92692 | Q92692-2;Q92692 |
| Plexin-A1 | PLXNA1 | | 1.27198 | -1.22418 | Q9UIW2 | Q9UIW2 |
| Desmoglein-1 | DSG1 | | 0.410325 | -1.21731 | Q02413;Q02413-2 | Q02413 |
| Glutaredoxin-1 | GLRX | | 1.0734 | -1.21396 | P35754 | P35754 |
| Hepatocyte growth factor activator;Hepatocyte growth factor activator short chain;Hep | HGFAC | | 0.402347 | -1.20611 | Q04756 | Q04756 |
| Lysyl oxidase homolog 1 | LOXL1 | | 1.04491 | -1.19845 | Q08397 | Q08397 |
| Protein kinase C delta type;Protein kinase C delta type regulatory subunit;Protein kinas | PRKCD | | 0.519189 | -1.19758 | Q05655;Q05655-2 | Q05655;Q05655-2 |
| Spondin-1 | SPON1 | | 0.564635 | -1.19097 | Q9HCB6 | Q9HCB6 |
| Follistatin | FST | + | 1.57479 | -1.16516 | P19883-2;P19883 | P19883-2;P19883 |
| Periplakin | PPL | | 0.653281 | -1.1606 | O60437 | O60437 |
| Desmoglein-3 | DSG3 | | 0.770653 | -1.15979 | P32926 | P32926 |
| Pentraxin-related protein PTX3 | PTX3 | + | 1.62866 | -1.15546 | P26022 | P26022 |
| Protein BRICK1 | BRK1 | | 0.512777 | -1.14742 | Q8WUW1;Q8WUW1-2 | Q8WUW1;Q8WUW1-2 |
| Laminin subunit alpha-3 | LAMA3 | + | 1.38552 | -1.14044 | Q16787;Q16787-3 | Q16787;Q16787-3 |
| Calpain-7 | CAPN7 | | 0.407695 | -1.13027 | Q9Y6W3 | Q9Y6W3 |
| Acyl-protein thioesterase 1 | LYPLA1 | + | 1.35474 | -1.12943 | O75608-2;O75608 | O75608-2;O75608 |
| Antileukoproteinase | SLPI | + | 2.65784 | -1.1288 | P03973 | P03973 |
| 26S proteasome non-ATPase regulatory subunit 7 | PSMD7 | + | 1.57328 | -1.11132 | P51665 | P51665 |
| Golgin subfamily A member 4 | GOLGA4 | + | 1.38255 | -1.09832 | Q13439-3;Q13439-4;Q13439;Q13439-5 | Q13439-3;Q13439-4;Q13439;Q13439-5 |
| Insulin-like growth factor-binding protein 2 | IGFBP2 | + | 1.42192 | -1.08246 | P18065 | P18065 |
| BH3-interacting domain death agonist;BH3-interacting domain death agonist p15;BH3-i | BID | | 0.920856 | -1.07851 | P55957;P55957-2;P55957-4;P55957-3 | P55957;P55957-2 |
| Podocalyxin | PODXL | | 0.900324 | -1.06913 | O00592-2;O00592 | O00592-2;O00592 |
| WAP four-disulfide core domain protein 2 | WFDC2 | + | 1.67377 | -1.06049 | Q14508;Q14508-2;Q14508-3;Q14508-4;Q14508-5 | Q14508;Q14508-2;Q14508-3 |
| Hepatocyte growth factor-like protein;Hepatocyte growth factor-like protein alpha chai | MST1 | | 0.196156 | -1.05012 | P26927 | P26927 |
| Protocadherin Fat 1;Protocadherin Fat 1, nuclear form | FAT1 | + | 2.4662 | -1.02455 | Q14517 | Q14517 |
| Complement C3;Complement C3 beta chain;C3-beta-c;Complement C3 alpha chain;C3a | C3 | + | 2.3124 | -1.0245 | P01024 | P01024 |
| Plasma kallikrein;Plasma kallikrein heavy chain;Plasma kallikrein light chain | KLKB1 | | 0.305807 | -1.02079 | P03952 | P03952 |
| Serpin B6 | SERPINB6 | + | 3.24461 | -1.01737 | P35237 | P35237 |
| Aldehyde dehydrogenase, mitochondrial | ALDH2 | + | 1.79177 | -1.00677 | P05091;P05091-2 | P05091;P05091-2 |
| Actin-related protein 2/3 complex subunit 1B | ARPC1B | + | 3.10963 | -0.999318 | O15143 | O15143 |
| Coiled-coil domain-containing protein 132 | CCDC132 | | 1.16248 | -0.997663 | Q96JG6;Q96JG6-3;Q96JG6-2 | Q96JG6;Q96JG6-3;Q96JG6-2 |
| Suppressor of tumorigenicity 14 protein | ST14 | + | 2.79699 | -0.992232 | Q9Y5Y6 | Q9Y5Y6 |
| Pterin-4-alpha-carbinolamine dehydratase | PCBD1 | + | 1.32785 | -0.973577 | P61457 | P61457 |
| Cytidine deaminase | CDA | + | 1.67992 | -0.972997 | P32320 | P32320 |
| Filamin-B | FLNB | + | 3.6068 | -0.972551 | O75369-8;O75369;O75369-2;O75369-9;O75369-3;O7536 | O75369-8;O75369;O75369-2;O75369-9;O75369-3;C |
| UDP-glucose 6-dehydrogenase | UGDH | | 1.27015 | -0.968405 | O60701;O60701-2 | O60701;O60701-2 |
| Tubulointerstitial nephritis antigen-like | TINAGL1 | + | 1.47173 | -0.966138 | Q9GZM7;Q9GZM7-3;Q9GZM7-2 | Q9GZM7;Q9GZM7-3 |
| Endothelial protein C receptor | PROCR | + | 1.56511 | -0.960657 | Q9UNN8 | Q9UNN8 |
| Cystatin-B | CSTB | + | 3.20164 | -0.955949 | P04080 | P04080 |
| Serpin B8 | SERPINB8 | + | 1.98466 | -0.932871 | P50452;P50452-2;P50452-3 | P50452;P50452-2 |
| Collagenase 3 | MMP13 | | 0.921378 | -0.930181 | P45452 | P45452 |

| | | | | | | |
|---|---|---|---|---|---|---|
| Complement C1r subcomponent-like protein | C1RL | | 0.773602 | -0.930091 | Q9NZP8 | Q9NZP8 |
| SEC14-like protein 2 | SEC14L2 | + | 1.3576 | -0.929386 | O76054;O76054-5;O76054-4;Q9UDX4 | O76054;O76054-5;O76054-4 |
| Myotubularin-related protein 2 | MTMR2 | | 0.597748 | -0.923745 | Q13614-2;Q13614 | Q13614-2;Q13614 |
| Complement C1q tumor necrosis factor-related protein 3 | C1QTNF3 | | 0.726646 | -0.918636 | Q9BXJ4;Q9BXJ4-3;Q9BXJ4-2 | Q9BXJ4;Q9BXJ4-3;Q9BXJ4-2 |
| Programmed cell death protein 6 | PDCD6 | | 0.237442 | -0.916959 | O75340;O75340-2 | O75340;O75340-2 |
| Low-density lipoprotein receptor-related protein 6 | LRP6 | + | 1.79314 | -0.912401 | O75581 | O75581 |
| Nectin-4;Processed poliovirus receptor-related protein 4 | PVRL4 | | 0.250596 | -0.912107 | Q96NY8;Q96NY8-2 | Q96NY8 |
| Interferon-induced guanylate-binding protein 1 | GBP1 | | 0.607344 | -0.905563 | P32455;Q9H0R5-4;Q9H0R5 | P32455 |
| T-complex protein 1 subunit beta | CCT2 | | 1.27902 | -0.904424 | P78371;P78371-2 | P78371;P78371-2 |
| Collagen alpha-1(XI) chain | COL11A1 | | 0.701337 | -0.896329 | P12107-4;P12107-3;P12107;P12107-2 | P12107-4;P12107-3;P12107;P12107-2 |
| Serpin B7 | SERPINB7 | | 1.04336 | -0.895287 | O75635-2;O75635 | O75635-2;O75635 |
| Transcription elongation factor B polypeptide 2 | TCEB2 | | 0.919721 | -0.892812 | Q15370;Q15370-2 | Q15370;Q15370-2 |
| Glutamine--fructose-6-phosphate aminotransferase [isomerizing] 1 | GFPT1 | | 1.08493 | -0.891084 | Q06210-2;Q06210 | Q06210-2;Q06210 |
| Vascular endothelial growth factor B | VEGFB | | 0.381351 | -0.890736 | P49765-2;P49765 | P49765-2;P49765 |
| Heterogeneous nuclear ribonucleoprotein K | HNRNPK | | 0.137003 | -0.889065 | P61978-2;P61978 | P61978-2;P61978 |
| Angiopoietin-related protein 4 | ANGPTL4 | + | 1.31215 | -0.87749 | Q9BY76;Q9BY76-3;Q9BY76-2 | Q9BY76;Q9BY76-3;Q9BY76-2 |
| 5-nucleotidase | NT5E | | 0.413589 | -0.875699 | P21589-2;P21589 | P21589-2;P21589 |
| Bone morphogenetic protein 5 | BMP5 | | 0.401172 | -0.871629 | P22003-2;P22003 | P22003-2;P22003 |
| N-acetyl-D-glucosamine kinase | NAGK | + | 2.45222 | -0.866546 | Q9UJ70;Q9UJ70-2 | Q9UJ70;Q9UJ70-2 |
| Protein CYR61 | CYR61 | + | 2.09192 | -0.86489 | O00622 | O00622 |
| Migration and invasion enhancer 1 | MIEN1 | | 0.95321 | -0.862074 | Q9BRT3 | Q9BRT3 |
| T-complex protein 1 subunit delta | CCT4 | | 1.26442 | -0.839933 | P50991;P50991-2 | P50991;P50991-2 |
| Echinoderm microtubule-associated protein-like 2 | EML2 | | 0.910577 | -0.83886 | O95834;O95834-2;O95834-3;Q05BV3-4;Q05BV3-2;Q6ZM | O95834;O95834-2;O95834-3 |
| Vascular endothelial growth factor A | VEGFA | | 0.52164 | -0.838641 | P15692-8;P15692-4;P15692-3;P15692-2;P15692;P15692- | P15692-8;P15692-4;P15692-3;P15692-2;P15692;P1 |
| Coiled-coil domain-containing protein 80 | CCDC80 | | 0.690688 | -0.835498 | Q76M96;Q76M96-2;Q76M96-3 | Q76M96;Q76M96-2 |
| E3 ubiquitin-protein ligase NEDD4 | NEDD4 | | 0.39528 | -0.833963 | P46934-4;P46934-2;P46934;P46934-3 | P46934-4;P46934-2;P46934 |
| Proteasome subunit beta type-9 | PSMB9 | + | 2.18829 | -0.82756 | P28065-2;P28065 | P28065-2;P28065 |
| T-complex protein 1 subunit zeta | CCT6A | + | 1.8644 | -0.827452 | P40227;P40227-2;Q92526;Q92526-2;Q92526-3 | P40227;P40227-2 |
| Serine/threonine-protein phosphatase 2A activator | PPP2R4 | | 0.672894 | -0.824621 | Q15257-3;Q15257-2;Q15257;Q15257-4 | Q15257-3;Q15257-2;Q15257;Q15257-4 |
| Kin of IRRE-like protein 1 | KIRREL | + | 1.7692 | -0.824109 | Q96J84;Q96J84-2;Q96J84-3 | Q96J84;Q96J84-2;Q96J84-3 |
| Ephrin-A1;Ephrin-A1, secreted form | EFNA1 | | 0.459395 | -0.820918 | P20827;P20827-2 | P20827;P20827-2 |
| T-complex protein 1 subunit theta | CCT8 | + | 2.04816 | -0.817949 | P50990;P50990-2;P50990-3 | P50990;P50990-2;P50990-3 |
| Intraflagellar transport protein 25 homolog | HSPB11 | | 0.513106 | -0.814313 | Q9Y547 | Q9Y547 |
| Cystatin-A;Cystatin-A, N-terminally processed | CSTA | + | 1.30636 | -0.809388 | P01040 | P01040 |
| Dystonin | DST | | 0.604692 | -0.808778 | Q03001;Q03001-8;Q03001-9;Q03001-13;Q03001-10 | Q03001;Q03001-8 |
| Collagen alpha-1(II) chain;Collagen alpha-1(II) chain;Chondrocalcin | COL2A1 | | 0.967149 | -0.807041 | P02458-3;P02458-1;P02458 | P02458-3;P02458-1;P02458 |
| Ragulator complex protein LAMTOR2 | LAMTOR2 | | 1.01904 | -0.803217 | Q9Y2Q5;Q9Y2Q5-2;Q9Y2Q5-3 | Q9Y2Q5;Q9Y2Q5-2 |
| T-complex protein 1 subunit alpha | TCP1 | + | 1.59728 | -0.801014 | P17987 | P17987 |
| Macrophage-capping protein | CAPG | + | 2.39224 | -0.792356 | P40121;P40121-2 | P40121;P40121-2 |
| Nicotinate phosphoribosyltransferase | NAPRT | + | 2.32541 | -0.790056 | Q6XQN6;Q6XQN6-3;Q6XQN6-2 | Q6XQN6;Q6XQN6-3;Q6XQN6-2 |
| Protein archease | ZBTB8OS | + | 1.54055 | -0.778626 | Q8IWT0;Q8IWT0-2 | Q8IWT0;Q8IWT0-2 |
| Fumarylacetoacetase | FAH | | 0.67139 | -0.775288 | P16930;P16930-2 | P16930;P16930-2 |
| N(G),N(G)-dimethylarginine dimethylaminohydrolase 2 | DDAH2 | | 1.12563 | -0.769023 | O95865 | O95865 |
| Semaphorin-4D | SEMA4D | | 0.733251 | -0.767787 | Q92854-2;Q92854 | Q92854-2;Q92854 |
| Annexin A3 | ANXA3 | + | 1.42142 | -0.767557 | P12429 | P12429 |
| Titin | TTN | | 0.58389 | -0.760197 | Q8WZ42-5;Q8WZ42-2;Q8WZ42;Q8WZ42-8;Q8WZ42-13;( | Q8WZ42-5;Q8WZ42-2;Q8WZ42;Q8WZ42-8;Q8WZ4 |
| Leucine-rich repeat transmembrane protein FLRT3 | FLRT3 | + | 1.32131 | -0.755262 | Q9NZU0 | Q9NZU0 |
| Insulin-like growth factor II;Insulin-like growth factor II;Insulin-like growth factor II Ala-: | IGF2 | | 0.868534 | -0.7536 | P01344;P01344-2;P01344-3 | P01344;P01344-2;P01344-3 |
| Biliverdin reductase A | BLVRA | + | 1.37456 | -0.752351 | P53004 | P53004 |
| Chloride intracellular channel protein 4 | CLIC4 | | 0.828343 | -0.750121 | Q9Y696;O15247 | Q9Y696 |
| Matrix metalloproteinase-15 | MMP15 | | 1.21469 | -0.749917 | P51511;Q9Y5R2 | P51511 |
| Leucine-rich repeat-containing protein 16A | LRRC16A | | 0.412182 | -0.743399 | Q5VZK9-2;Q5VZK9;Q5VZK9-3;Q5VZK9-4 | Q5VZK9-2;Q5VZK9 |
| Reticulon-4 | RTN4 | | 0.84538 | -0.742446 | Q9NQC3-5;Q9NQC3;Q9NQC3-2;Q9NQC3-4;Q9NQC3-6 | Q9NQC3-5;Q9NQC3;Q9NQC3-2;Q9NQC3-4;Q9NQC |
| Ferritin light chain | FTL | | 0.358355 | -0.734725 | P02792 | P02792 |
| Pantetheinase | VNN1 | | 0.840509 | -0.726321 | O95497 | O95497 |
| Syntaxin-12 | STX12 | + | 1.76588 | -0.726131 | Q86Y82 | Q86Y82 |
| Choriogonadotropin subunit beta variant 1;Choriogonadotropin subunit beta;Choriogoi | CGB1;CGB;CGB2 | | 1.25167 | -0.721333 | A6NKQ9-2;P01233;P01233-2;A6NKQ9;Q6NT52 | A6NKQ9-2;P01233;P01233-2;A6NKQ9;Q6NT52 |
| Protein MON2 homolog | MON2 | + | 1.34488 | -0.719811 | Q7Z3U7-2;Q7Z3U7-6;Q7Z3U7-5;Q7Z3U7;Q7Z3U7-3 | Q7Z3U7-2;Q7Z3U7-6;Q7Z3U7-5;Q7Z3U7 |
| Aminoacyl tRNA synthase complex-interacting multifunctional protein 1;Endothelial mc | AIMP1 | | 1.03478 | -0.717545 | Q12904;Q12904-2 | Q12904;Q12904-2 |
| Arylsulfatase A;Arylsulfatase A component B;Arylsulfatase A component C | ARSA | | 0.674277 | -0.715984 | P15289-2;P15289 | P15289-2;P15289 |
| Signal recognition particle receptor subunit alpha | SRPR | + | 1.59391 | -0.714719 | P08240;P08240-2 | P08240;P08240-2 |
| Perilipin-2 | PLIN2 | | 0.67576 | -0.706142 | Q99541 | Q99541 |
| Alpha-actinin-1 | ACTN1 | + | 1.59475 | -0.703785 | P12814;P12814-3;P12814-2;P12814-4 | P12814;P12814-3;P12814-2;P12814-4 |
| Apoptosis regulator BAX | BAX | | 1.05793 | -0.699135 | Q07812-5;Q07812-8;Q07812;Q07812-2;Q07812-7;Q0781 | Q07812-5;Q07812-8;Q07812;Q07812-2;Q07812-7 |
| Sialic acid synthase | NANS | | 0.776992 | -0.697978 | Q9NR45 | Q9NR45 |
| AP-2 complex subunit mu | AP2M1 | | 0.828647 | -0.692143 | Q96CW1-2;Q96CW1 | Q96CW1-2;Q96CW1 |
| Osteoclast-stimulating factor 1 | OSTF1 | | 0.371091 | -0.692051 | Q92882 | Q92882 |
| Moesin | MSN | + | 3.26077 | -0.691112 | P26038;REV__Q9H446-2;REV__Q9H446 | P26038 |

| | | | | | | |
|---|---|---|---|---|---|---|
| Arachidonate 15-lipoxygenase B | ALOX15B | | 0.498898 | -0.690193 | O15296-3;O15296;O15296-2;O15296-4 | O15296-3;O15296;O15296-2;O15296-4 |
| Chondroitin sulfate proteoglycan 4 | CSPG4 | + | 2.53956 | -0.688801 | Q6UVK1 | Q6UVK1 |
| Eukaryotic translation elongation factor 1 epsilon-1 | EEF1E1 | | 1.06894 | -0.680758 | O43324;O43324-2 | O43324;O43324-2 |
| Tumor necrosis factor ligand superfamily member 13 | TNFSF13 | | 0.88086 | -0.680546 | O43508-2;O75888-5;O75888-4;O75888;O75888-3 | O43508-2;O75888-5;O75888-4;O75888;O75888-3 |
| Protein S100-A14 | S100A14 | | 0.716907 | -0.675877 | Q9HCY8 | Q9HCY8 |
| Probable ATP-dependent RNA helicase DDX23 | DDX23 | | 1.04341 | -0.675294 | Q9BUQ8 | Q9BUQ8 |
| Serine/threonine-protein phosphatase 2A 65 kDa regulatory subunit A beta isoform | PPP2R1B | | 0.638279 | -0.673712 | P30154;P30154-3;P30154-2;P30154-4;P30154-5 | P30154;P30154-3;P30154-2;P30154-4;P30154-5 |
| Gamma-enolase | ENO2 | + | 2.64099 | -0.673433 | P09104;P09104-2 | P09104;P09104-2 |
| Thioredoxin | TXN | | 0.451638 | -0.671707 | P10599;P10599-2 | P10599;P10599-2 |
| 26S proteasome non-ATPase regulatory subunit 12 | PSMD12 | | 1.06142 | -0.666801 | O00232;O00232-2 | O00232;O00232-2 |
| Ribosyldihydronicotinamide dehydrogenase [quinone] | NQO2 | + | 1.48564 | -0.662428 | P16083 | P16083 |
| 26S proteasome non-ATPase regulatory subunit 8 | PSMD8 | | 1.17185 | -0.661452 | P48556 | P48556 |
| Intercellular adhesion molecule 1 | ICAM1 | + | 2.47564 | -0.659642 | P05362 | P05362 |
| Dual specificity protein phosphatase 3 | DUSP3 | | 0.28813 | -0.659125 | P51452-2 | P51452-2 |
| Protein S100-P | S100P | | 0.526786 | -0.655245 | P25815 | P25815 |
| Prostasin;Prostasin light chain;Prostasin heavy chain | PRSS8 | | 0.483363 | -0.653887 | Q16651;Q16651-2 | Q16651;Q16651-2 |
| Ras-related protein Rab-1A | RAB1A | | 0.369611 | -0.650841 | P62820;P62820-2;P62820-3 | P62820;P62820-2;P62820-3 |
| Single-stranded DNA-binding protein, mitochondrial | SSBP1 | | 0.418287 | -0.649351 | Q04837 | Q04837 |
| Kunitz-type protease inhibitor 1 | SPINT1 | | 0.62164 | -0.647347 | O43278 | O43278 |
| Proteasome subunit beta type-8 | PSMB8 | | 0.427732 | -0.642673 | P28062-2;P28062 | P28062-2;P28062 |
| T-complex protein 1 subunit gamma | CCT3 | + | 1.52185 | -0.635544 | P49368;P49368-2 | P49368;P49368-2 |
| 15 kDa selenoprotein | SEP15 | | 0.218111 | -0.633457 | O60613;O60613-2 | O60613 |
| Proteasome activator complex subunit 2 | PSME2 | + | 1.55906 | -0.629313 | Q9UL46 | Q9UL46 |
| Syntenin-1 | SDCBP | + | 1.88309 | -0.62894 | O00560;O00560-3 | O00560;O00560-3 |
| Plastin-1 | PLS1 | | 0.759604 | -0.628364 | Q14651 | Q14651 |
| Cytosolic Fe-S cluster assembly factor NUBP1 | NUBP1 | | 0.334191 | -0.627654 | P53384-2;P53384 | P53384-2;P53384 |
| Dynein light chain 1, cytoplasmic | DYNLL1 | | 0.233533 | -0.62765 | P63167 | P63167 |
| Mitochondrial import receptor subunit TOM70 | TOMM70A | + | 1.9275 | -0.624317 | O94826 | O94826 |
| Msx2-interacting protein | SPEN | | 0.138011 | -0.624013 | Q96T58 | Q96T58 |
| Insulin-like growth factor-binding protein 7 | IGFBP7 | | 0.722375 | -0.623051 | Q16270-2;Q16270 | Q16270-2;Q16270 |
| EH domain-containing protein 1 | EHD1 | | 0.692925 | -0.622058 | Q9H4M9 | Q9H4M9 |
| Galactokinase | GALK1 | | 0.389049 | -0.615256 | P51570;P51570-2 | P51570;P51570-2 |
| Calpain-1 catalytic subunit | CAPN1 | + | 2.14187 | -0.613473 | P07384 | P07384 |
| Puromycin-sensitive aminopeptidase | NPEPPS | | 0.920677 | -0.612694 | P55786;P55786-2;A6NEC2;A6NEC2-2;A6NEC2-3 | P55786;P55786-2 |
| Deoxyribose-phosphate aldolase | DERA | + | 1.43321 | -0.611127 | Q9Y315 | Q9Y315 |
| Twinfilin-2 | TWF2 | + | 3.18805 | -0.610703 | Q6IBS0;Q9NR96-4 | Q6IBS0 |
| N(G),N(G)-dimethylarginine dimethylaminohydrolase 1 | DDAH1 | | 0.931276 | -0.60918 | O94760;O94760-2 | O94760;O94760-2 |
| Glutathione S-transferase A3 | GSTA3 | | 0.332843 | -0.607155 | Q16772 | Q16772 |
| Tyrosine-protein phosphatase non-receptor type 6 | PTPN6 | | 0.61751 | -0.60626 | P29350;P29350-3;P29350-4;P29350-2 | P29350;P29350-3;P29350-4;P29350-2 |
| Phospholipid scramblase 1 | PLSCR1 | | 0.343508 | -0.606218 | O15162-2;O15162 | O15162-2;O15162 |
| Galectin-8 | LGALS8 | | 0.854296 | -0.604326 | O00214;O00214-2 | O00214;O00214-2 |
| T-complex protein 1 subunit eta | CCT7 | + | 2.00821 | -0.602604 | Q99832;Q99832-3;Q99832-4;Q99832-2 | Q99832;Q99832-3;Q99832-4;Q99832-2 |
| SH3 domain-binding glutamic acid-rich-like protein 3 | SH3BGRL3 | | 1.0935 | -0.601634 | Q9H299 | Q9H299 |
| STAM-binding protein | STAMBP | | 0.845793 | -0.599161 | O95630;O95630-2 | O95630;O95630-2 |
| Programmed cell death protein 10 | PDCD10 | | 0.464444 | -0.59685 | Q9BUL8 | Q9BUL8 |
| Actin-related protein 2/3 complex subunit 3 | ARPC3 | + | 1.32663 | -0.593358 | O15145 | O15145 |
| Eukaryotic initiation factor 4A-I | EIF4A1 | + | 1.3036 | -0.592355 | P60842;P60842-2 | P60842;P60842-2 |
| V-type proton ATPase catalytic subunit A | ATP6V1A | | 0.699531 | -0.5919 | P38606-2;P38606 | P38606-2;P38606 |
| NKG2D ligand 2 | ULBP2 | | 0.906448 | -0.589724 | Q9BZM5;Q6H3X3-2;Q5VY80;Q6H3X3 | Q9BZM5 |
| Proteasome subunit alpha type-2 | PSMA2 | | 0.79727 | -0.588732 | P25787 | P25787 |
| WD repeat-containing protein 1 | WDR1 | + | 1.42025 | -0.587461 | O75083;O75083-3 | O75083;O75083-3 |
| Vinculin | VCL | + | 1.44744 | -0.587328 | P18206-2;P18206;P18206-3 | P18206-2;P18206 |
| Transaldolase | TALDO1 | | 0.555106 | -0.585161 | P37837 | P37837 |
| 26S protease regulatory subunit 8 | PSMC5 | | 0.844707 | -0.583152 | P62195;P62195-2;Q8NB90-3;Q8NB90-2;Q8NB90 | P62195;P62195-2 |
| 26S proteasome non-ATPase regulatory subunit 6 | PSMD6 | | 1.10889 | -0.58079 | Q15008;Q15008-4;Q15008-3;Q15008-2 | Q15008;Q15008-4;Q15008-3;Q15008-2 |
| Peptidyl-prolyl cis-trans isomerase FKBP1A | FKBP1A | | 0.346088 | -0.580031 | P62942 | P62942 |
| Argininosuccinate lyase | ASL | | 1.02578 | -0.579806 | P04424-2;P04424;P04424-3 | P04424-2;P04424;P04424-3 |
| Mannosyl-oligosaccharide 1,2-alpha-mannosidase IA | MAN1A1 | + | 1.95766 | -0.577739 | P33908;P33908-2 | P33908;P33908-2 |
| Eukaryotic translation initiation factor 2 subunit 3;Putative eukaryotic translation initiat | EIF2S3;EIF2S3L | | 1.07645 | -0.577408 | P41091;Q2VIR3;Q2VIR3-2 | P41091;Q2VIR3;Q2VIR3-2 |
| Programmed cell death 6-interacting protein | PDCD6IP | | 1.01255 | -0.575124 | Q8WUM4;Q8WUM4-2;Q8WUM4-3 | Q8WUM4;Q8WUM4-2 |
| Coronin-1B | CORO1B | + | 2.83543 | -0.570129 | Q9BR76 | Q9BR76 |
| Nascent polypeptide-associated complex subunit alpha;Nascent polypeptide-associatec | NACA | | 0.978043 | -0.565635 | Q13765;E9PAV3-2;E9PAV3;Q9BZK3 | Q13765;E9PAV3-2;E9PAV3 |
| 7,8-dihydro-8-oxoguanine triphosphatase | NUDT1 | | 0.561506 | -0.56513 | P36639-2;P36639-4;P36639-3;P36639 | P36639-2;P36639-4;P36639-3;P36639 |
| Cytoplasmic dynein 1 heavy chain 1 | DYNC1H1 | | 1.16979 | -0.564935 | Q14204 | Q14204 |
| Clathrin heavy chain 1 | CLTC | | 1.22345 | -0.563947 | Q00610-2;Q00610 | Q00610-2;Q00610 |
| Unconventional myosin-Ie | MYO1E | | 0.214732 | -0.562139 | Q12965 | Q12965 |
| Interleukin-1 receptor-like 1 | IL1RL1 | | 0.572164 | -0.561935 | Q01638-2;Q01638;Q01638-4;Q01638-3 | Q01638-2;Q01638;Q01638-4;Q01638-3 |
| 26S proteasome non-ATPase regulatory subunit 3 | PSMD3 | + | 1.53021 | -0.560316 | O43242;O43242-2 | O43242;O43242-2 |

| | | | | | | |
|---|---|---|---|---|---|---|
| Heparan-sulfate 6-O-sulfotransferase 2 | HS6ST2 | | 0.724842 | -0.559744 | Q96MM7-2;Q96MM7-3;Q96MM7;Q96MM7-4 | Q96MM7-2;Q96MM7-3;Q96MM7;Q96MM7-4 |
| Epithelial discoidin domain-containing receptor 1 | DDR1 | | 0.45953 | -0.558921 | Q08345-2;Q08345-6;Q08345;Q08345-5;Q08345-4 | Q08345-2;Q08345-6;Q08345;Q08345-5;Q08345-4 |
| Phosphatidylinositol transfer protein alpha isoform | PITPNA | | 0.353995 | -0.558137 | Q00169 | Q00169 |
| Ubiquitin-like protein ISG15 | ISG15 | + | 2.18376 | -0.55722 | P05161 | P05161 |
| Leukotriene A-4 hydrolase | LTA4H | + | 1.4289 | -0.55592 | P09960;P09960-4;P09960-2;P09960-3 | P09960;P09960-4;P09960-2;P09960-3 |
| Protein ITFG3 | ITFG3 | | 0.28626 | -0.553857 | Q9H0X4;Q9H0X4-2 | Q9H0X4;Q9H0X4-2 |
| Alpha-actinin-4 | ACTN4 | + | 1.99985 | -0.553582 | O43707;O43707-2;O43707-3 | O43707;O43707-2;O43707-3 |
| Acid sphingomyelinase-like phosphodiesterase 3b | SMPDL3B | | 0.690761 | -0.553114 | Q92485;Q92485-2 | Q92485;Q92485-2 |
| Actin-related protein 3 | ACTR3 | | 0.773276 | -0.552073 | P61158;Q9C0K3;Q9P1U1-3 | P61158 |
| Platelet-activating factor acetylhydrolase IB subunit alpha | PAFAH1B1 | | 1.26053 | -0.551915 | P43034;P43034-2 | P43034 |
| Breakpoint cluster region protein | BCR | | 0.569647 | -0.551232 | P11274-2;P11274 | P11274-2;P11274 |
| Tropomyosin alpha-4 chain | TPM4 | | 1.00169 | -0.551158 | P67936 | P67936 |
| Peptidyl-prolyl cis-trans isomerase-like 1 | PPIL1 | | 0.657066 | -0.551008 | Q9Y3C6 | Q9Y3C6 |
| F-actin-capping protein subunit alpha-1 | CAPZA1 | | 0.960484 | -0.548446 | P52907 | P52907 |
| Bleomycin hydrolase | BLMH | | 0.585147 | -0.547608 | Q13867 | Q13867 |
| Calpain small subunit 1 | CAPNS1 | | 0.767487 | -0.546758 | P04632;Q96L46 | P04632 |
| Mesencephalic astrocyte-derived neurotrophic factor | MANF | | 1.13682 | -0.544308 | P55145 | P55145 |
| 3(2),5-bisphosphate nucleotidase 1 | BPNT1 | | 1.08665 | -0.54212 | O95861;O95861-2;O95861-4 | O95861;O95861-2;O95861-4 |
| Transketolase | TKT | | 0.802137 | -0.540896 | P29401;P29401-2 | P29401;P29401-2 |
| Hydroxyacylglutathione hydrolase, mitochondrial | HAGH | | 0.296434 | -0.540231 | Q16775-2;Q16775;Q16775-3 | Q16775-2;Q16775 |
| 5-AMP-activated protein kinase subunit gamma-1 | PRKAG1 | | 1.06484 | -0.534566 | P54619-2;P54619;P54619-3 | P54619-2;P54619;P54619-3 |
| Diphosphoinositol polyphosphate phosphohydrolase 1 | NUDT3 | | 1.00093 | -0.534415 | O95989;Q96G61;Q8NFP7 | O95989 |
| Dehydrogenase/reductase SDR family member 4 | DHRS4 | | 0.394292 | -0.534353 | Q9BTZ2-3;Q9BTZ2-7;Q9BTZ2-2;Q9BTZ2-5;Q9BTZ2-8;Q9B | Q9BTZ2-3;Q9BTZ2-7;Q9BTZ2-2;Q9BTZ2-5;Q9BTZ2-8 |
| Nuclear receptor 2C2-associated protein | NR2C2AP | | 0.54954 | -0.532539 | Q86WQ0;Q86WQ0-2 | Q86WQ0;Q86WQ0-2 |
| E3 ubiquitin-protein ligase HUWE1 | HUWE1 | | 0.862569 | -0.531613 | Q7Z6Z7-3;Q7Z6Z7;Q7Z6Z7-2;A5YM72-2;A5YM72;A5YM7 | Q7Z6Z7-3;Q7Z6Z7;Q7Z6Z7-2 |
| Syntaxin-7 | STX7 | | 1.03995 | -0.530256 | O15400-2;O15400 | O15400-2;O15400 |
| Adenylyl cyclase-associated protein 1 | CAP1 | + | 1.73724 | -0.529403 | Q01518-2 | Q01518-2 |
| Coatomer subunit alpha;Xenin;Proxenin | COPA | | 0.974979 | -0.528547 | P53621;P53621-2 | P53621;P53621-2 |
| Actin-related protein 2/3 complex subunit 2 | ARPC2 | | 0.62285 | -0.526996 | O15144 | O15144 |
| 26S protease regulatory subunit 10B | PSMC6 | | 1.1946 | -0.526896 | P62333 | P62333 |
| Staphylococcal nuclease domain-containing protein 1 | SND1 | | 1.00981 | -0.526558 | Q7KZF4 | Q7KZF4 |
| Ubiquitin-conjugating enzyme E2 G1;Ubiquitin-conjugating enzyme E2 G1, N-terminally | UBE2G1 | + | 1.5919 | -0.525494 | P62253 | P62253 |
| Arginase-1 | ARG1 | | 0.639677 | -0.524035 | P05089;P05089-2;P05089-3 | P05089;P05089-2;P05089-3 |
| Coactosin-like protein | COTL1 | + | 1.66934 | -0.524022 | Q14019 | Q14019 |
| Prostaglandin reductase 1 | PTGR1 | | 1.23119 | -0.519964 | Q14914;Q14914-2 | Q14914;Q14914-2 |
| 26S proteasome non-ATPase regulatory subunit 2 | PSMD2 | | 1.27619 | -0.519462 | Q13200;Q13200-3;Q13200-2 | Q13200;Q13200-3;Q13200-2 |
| Ubiquitin carboxyl-terminal hydrolase 14 | USP14 | + | 1.34704 | -0.518799 | P54578;P54578-2;P54578-3 | P54578;P54578-2;P54578-3 |
| Niban-like protein 1 | FAM129B | | 0.922001 | -0.517822 | Q96TA1-2;Q96TA1 | Q96TA1-2;Q96TA1 |
| V-type proton ATPase subunit C 1 | ATP6V1C1 | + | 1.33181 | -0.516702 | P21283 | P21283 |
| Spectrin beta chain, non-erythrocytic 2 | SPTBN2 | | 0.763805 | -0.51538 | O15020;O15020-2 | O15020;O15020-2 |
| Apolipoprotein B-100;Apolipoprotein B-48 | APOB | | 0.604831 | -0.510304 | P04114 | P04114 |
| PDZ domain-containing protein GIPC1 | GIPC1 | | 0.639694 | -0.509687 | O14908;O14908-2;Q8TF64 | O14908;O14908-2 |
| Adenylate kinase isoenzyme 1 | AK1 | | 0.598426 | -0.508541 | P00568;Q9Y6K8-2;Q9Y6K8-3;Q9Y6K8 | P00568 |
| Ribonuclease UK114 | HRSP12 | | 0.820056 | -0.508447 | P52758 | P52758 |
| Plectin | PLEC | + | 1.45986 | -0.503785 | Q15149-2 | Q15149-2 |
| T-complex protein 1 subunit epsilon | CCT5 | + | 1.44384 | -0.501701 | P48643;P48643-2 | P48643;P48643-2 |
| Proteasome subunit alpha type-1 | PSMA1 | | 0.915393 | -0.501238 | P25786;P25786-2 | P25786;P25786-2 |
| Proteasome subunit alpha type-4 | PSMA4 | | 0.521146 | -0.499369 | P25789;P25789-2 | P25789;P25789-2 |
| Low-density lipoprotein receptor-related protein 8 | LRP8 | | 0.814005 | -0.499156 | Q14114-2;Q14114-3;Q14114;Q14114-4 | Q14114-2;Q14114-3;Q14114;Q14114-4 |
| Purine nucleoside phosphorylase | PNP | | 0.524351 | -0.498661 | P00491 | P00491 |
| Flavin reductase (NADPH) | BLVRB | | 0.732399 | -0.498236 | P30043 | P30043 |
| Ubiquitin-fold modifier-conjugating enzyme 1 | UFC1 | | 0.597537 | -0.497512 | Q9Y3C8 | Q9Y3C8 |
| Mitogen-activated protein kinase 1 | MAPK1 | | 0.493022 | -0.497304 | P28482-2;P28482 | P28482-2;P28482 |
| Xylosyltransferase 1 | XYLT1 | | 0.289796 | -0.496239 | Q86Y38 | Q86Y38 |
| F-actin-capping protein subunit beta | CAPZB | | 1.02234 | -0.495285 | P47756-2;P47756 | P47756-2;P47756 |
| Actin-related protein 2 | ACTR2 | + | 2.01556 | -0.492648 | P61160;P61160-2 | P61160;P61160-2 |
| Actin-related protein 2/3 complex subunit 1A | ARPC1A | | 1.02697 | -0.49238 | Q92747;Q92747-2 | Q92747;Q92747-2 |
| Hippocalcin-like protein 1 | HPCAL1 | | 1.23624 | -0.49198 | P37235;P84074 | P37235 |
| Proteasome subunit alpha type-7 | PSMA7 | | 1.18957 | -0.489724 | O14818;O14818-4;Q8TAA3-2;Q8TAA3-5;Q8TAA3 | O14818 |
| Vesicle-fusing ATPase | NSF | | 0.621708 | -0.489347 | P46459;P46459-2 | P46459;P46459-2 |
| Protein deglycase DJ-1 | PARK7 | | 1.10794 | -0.488703 | Q99497 | Q99497 |
| Insulin-like growth factor-binding protein 3 | IGFBP3 | | 0.958106 | -0.488008 | P17936;P17936-2 | P17936;P17936-2 |
| Glycogenin-1 | GYG1 | | 1.00875 | -0.487331 | P46976-2;P46976;P46976-3 | P46976-2;P46976;P46976-3 |
| 26S proteasome non-ATPase regulatory subunit 1 | PSMD1 | | 1.10943 | -0.486605 | Q99460;Q99460-2 | Q99460;Q99460-2 |
| Alcohol dehydrogenase [NADP(+)] | AKR1A1 | | 0.668459 | -0.486357 | P14550 | P14550 |
| Tyrosine-protein kinase Lyn | LYN | | 0.260087 | -0.485942 | P07948-2;P07948 | P07948-2;P07948 |
| Serpin B5 | SERPINB5 | | 0.600469 | -0.475894 | P36952;P36952-2 | P36952;P36952-2 |
| Putative helicase MOV-10 | MOV10 | | 0.697168 | -0.471884 | Q9HCE1;Q9HCE1-2 | Q9HCE1;Q9HCE1-2 |

| | | | | | | |
|---|---|---|---|---|---|---|
| Beta-1,4-galactosyltransferase 1;Lactose synthase A protein;N-acetyllactosamine synth | B4GALT1 | | 0.395671 | -0.46928 | P15291-2;P15291 | P15291-2;P15291 |
| Protein-L-isoaspartate(D-aspartate) O-methyltransferase | PCMT1 | | 0.537228 | -0.468789 | P22061;P22061-2 | P22061;P22061-2 |
| Major vault protein | MVP | | 0.379761 | -0.467739 | Q14764 | Q14764 |
| Actin-related protein 2/3 complex subunit 4 | ARPC4 | + | 1.57256 | -0.467178 | P59998;P59998-3;P59998-4;P59998-2 | P59998;P59998-3;P59998-4;P59998-2 |
| Proteasome subunit alpha type-5 | PSMA5 | | 0.439622 | -0.466092 | P28066;P28066-2 | P28066;P28066-2 |
| Cell surface glycoprotein MUC18 | MCAM | | 0.694537 | -0.464357 | P43121;P43121-2 | P43121 |
| Serine/threonine-protein kinase 10 | STK10 | | 0.231681 | -0.46423 | O94804 | O94804 |
| Ras suppressor protein 1 | RSU1 | | 0.418596 | -0.462973 | Q15404;Q15404-2 | Q15404;Q15404-2 |
| Aspartyl aminopeptidase | DNPEP | | 0.85958 | -0.462514 | Q9ULA0 | Q9ULA0 |
| Vasodilator-stimulated phosphoprotein | VASP | + | 1.53461 | -0.461053 | P50552 | P50552 |
| Xaa-Pro aminopeptidase 1 | XPNPEP1 | + | 3.32379 | -0.456177 | Q9NQW7-3;Q9NQW7;Q9NQW7-4;Q9NQW7-2 | Q9NQW7-3;Q9NQW7;Q9NQW7-4;Q9NQW7-2 |
| Semaphorin-3A | SEMA3A | | 0.456019 | -0.455661 | Q14563 | Q14563 |
| Protein FAM49B | FAM49B | | 0.569494 | -0.455098 | Q9NUQ9;Q9NUQ9-2 | Q9NUQ9 |
| 1,4-alpha-glucan-branching enzyme | GBE1 | | 0.826307 | -0.454873 | Q04446 | Q04446 |
| Uridine diphosphate glucose pyrophosphatase | NUDT14 | | 0.268648 | -0.454364 | O95848 | O95848 |
| Follistatin-related protein 3 | FSTL3 | | 0.395675 | -0.452367 | O95633-2;O95633 | O95633-2;O95633 |
| Cdc42-interacting protein 4 | TRIP10 | | 1.29509 | -0.44947 | Q15642-4;Q15642-2;Q15642-3;Q15642;Q15642-5 | Q15642-4;Q15642-2;Q15642-3;Q15642;Q15642-5 |
| 6-phosphogluconate dehydrogenase, decarboxylating | PGD | + | 1.89371 | -0.447404 | P52209-2;P52209 | P52209-2;P52209 |
| NEDD8-conjugating enzyme Ubc12 | UBE2M | + | 1.58624 | -0.444304 | P61081 | P61081 |
| Dipeptidyl peptidase 3 | DPP3 | | 1.05669 | -0.443406 | Q9NY33;Q9NY33-4;Q9NY33-2 | Q9NY33;Q9NY33-4 |
| Vasorin | VASN | | 0.623896 | -0.441916 | Q6EMK4 | Q6EMK4 |
| Glyoxalase domain-containing protein 4 | GLOD4 | + | 1.99126 | -0.439262 | Q9HC38-2;Q9HC38;Q9HC38-3 | Q9HC38-2;Q9HC38;Q9HC38-3 |
| Glutathione S-transferase theta-2B;Glutathione S-transferase theta-2 | GSTT2B;GSTT2 | | 1.1981 | -0.437384 | P0CG30;P0CG29 | P0CG30;P0CG29 |
| Protein kinase C and casein kinase substrate in neurons protein 2 | PACSIN2 | | 0.802812 | -0.436519 | Q9UNF0-2;Q9UNF0 | Q9UNF0-2;Q9UNF0 |
| L-lactate dehydrogenase A chain | LDHA | | 0.375086 | -0.431512 | P00338;P00338-3;P00338-4;P00338-5;P00338-2;Q6ZMR3 | P00338;P00338-3;P00338-4;P00338-5;P00338-2 |
| V-type proton ATPase subunit G 1 | ATP6V1G1 | | 0.176147 | -0.430048 | O75348;O95670-2;O95670 | O75348 |
| Proteasome subunit beta type-4 | PSMB4 | | 0.338354 | -0.429803 | P28070 | P28070 |
| 3-ketoacyl-CoA thiolase, mitochondrial | ACAA2 | | 0.438904 | -0.429774 | P42765 | P42765 |
| Elongation factor 1-alpha 1;Putative elongation factor 1-alpha-like 3 | EEF1A1;EEF1A1P5 | + | 1.59463 | -0.428654 | P68104;Q5VTE0;P68104-2 | P68104;Q5VTE0;P68104-2 |
| Actin-related protein 2/3 complex subunit 5 | ARPC5 | | 0.704768 | -0.427299 | O15511;O15511-2 | O15511;O15511-2 |
| Glucose-6-phosphate 1-dehydrogenase | G6PD | | 0.935202 | -0.427259 | P11413;P11413-2;P11413-3 | P11413;P11413-2;P11413-3 |
| Talin-1 | TLN1 | | 0.789601 | -0.427245 | Q9Y490 | Q9Y490 |
| Unconventional myosin-Ib | MYO1B | | 0.431753 | -0.425706 | O43795-2;O43795 | O43795-2;O43795 |
| Proteasome subunit beta type-7 | PSMB7 | | 0.690654 | -0.425467 | Q99436;Q99436-2 | Q99436;Q99436-2 |
| Thioredoxin reductase 1, cytoplasmic | TXNRD1 | | 1.12822 | -0.424983 | Q16881-4;Q16881-3;Q16881 | Q16881-4;Q16881-3;Q16881 |
| V-type proton ATPase subunit D | ATP6V1D | | 0.166027 | -0.424353 | Q9Y5K8 | Q9Y5K8 |
| Protein-glutamine gamma-glutamyltransferase 2 | TGM2 | | 0.811743 | -0.422038 | P21980;P21980-2;P21980-3 | P21980;P21980-2 |
| V-type proton ATPase subunit B, brain isoform | ATP6V1B2 | | 0.508887 | -0.419749 | P21281;P15313 | P21281 |
| Serine/threonine-protein kinase OSR1 | OXSR1 | | 0.365274 | -0.419466 | O95747 | O95747 |
| Eukaryotic translation initiation factor 2 subunit 1 | EIF2S1 | + | 1.31121 | -0.418898 | P05198 | P05198 |
| Cytosolic acyl coenzyme A thioester hydrolase | ACOT7 | | 0.776853 | -0.418505 | O00154-4;O00154-6;O00154-7;O00154;O00154-5;O0015 | O00154-4;O00154-6;O00154-7;O00154;O00154-5;C |
| Glutathione S-transferase omega-1 | GSTO1 | | 0.554145 | -0.418038 | P78417;P78417-3;P78417-2 | P78417;P78417-3;P78417-2 |
| Sorting nexin-9 | SNX9 | | 0.372774 | -0.417768 | Q9Y5X1 | Q9Y5X1 |
| UDP-glucose 4-epimerase | GALE | | 0.694017 | -0.414273 | Q14376;Q14376-2 | Q14376;Q14376-2 |
| Eukaryotic translation initiation factor 4E | EIF4E | | 0.395025 | -0.414222 | P06730;P06730-2;P06730-3;A6NMX2 | P06730;P06730-2;P06730-3 |
| Phosphoglycerate mutase 1 | PGAM1 | | 1.03311 | -0.413665 | P18669;P15259;Q8N0Y7 | P18669 |
| BRO1 domain-containing protein BROX | BROX | + | 1.87655 | -0.413186 | Q5VW32;Q5VW32-2 | Q5VW32;Q5VW32-2 |
| Connective tissue growth factor | CTGF | | 0.428414 | -0.413106 | P29279;P29279-2 | P29279;P29279-2 |
| Nitrilase homolog 1 | NIT1 | + | 1.6051 | -0.412734 | Q86X76-2;Q86X76-4;Q86X76;Q86X76-3;Q86X76-5 | Q86X76-2;Q86X76-4;Q86X76;Q86X76-3 |
| Spectrin beta chain, non-erythrocytic 1 | SPTBN1 | | 0.555562 | -0.412399 | Q01082;Q01082-2 | Q01082;Q01082-2 |
| Exosome complex component RRP45 | EXOSC9 | | 0.464868 | -0.412208 | Q06265;Q06265-2;Q06265-3;Q06265-4 | Q06265;Q06265-2;Q06265-3;Q06265-4 |
| Phosphoglucomutase-1 | PGM1 | | 1.13641 | -0.41071 | P36871;P36871-2;P36871-3 | P36871;P36871-2;P36871-3 |
| Proprotein convertase subtilisin/kexin type 5 | PCSK5 | | 0.20001 | -0.409564 | Q92824-2;Q92824 | Q92824-2;Q92824 |
| Cysteine and glycine-rich protein 1 | CSRP1 | + | 1.59373 | -0.406584 | P21291 | P21291 |
| Tumor susceptibility gene 101 protein | TSG101 | | 0.91535 | -0.405588 | Q99816;Q99816-2 | Q99816;Q99816-2 |
| Phosphatidylinositol transfer protein beta isoform | PITPNB | | 0.329863 | -0.402908 | P48739-3;P48739;P48739-2 | P48739-3;P48739;P48739-2 |
| Anterior gradient protein 2 homolog | AGR2 | | 0.302474 | -0.402391 | O95994;Q8TD06 | O95994 |
| Ezrin | EZR | | 0.682034 | -0.402023 | P15311 | P15311 |
| Heat shock 70 kDa protein 4 | HSPA4 | + | 1.38535 | -0.401814 | P34932;P34932-2 | P34932 |
| Glycolipid transfer protein | GLTP | | 0.640736 | -0.401189 | Q9NZD2 | Q9NZD2 |
| Exportin-2 | CSE1L | | 0.938372 | -0.401162 | P55060-3;P55060;P55060-4;P55060-2 | P55060-3;P55060;P55060-4 |
| Myosin-14 | MYH14 | | 0.425323 | -0.399674 | Q7Z406;Q7Z406-6;Q7Z406-2;Q7Z406-5;Q7Z406-4 | Q7Z406;Q7Z406-6;Q7Z406-2;Q7Z406-5;Q7Z406-4 |
| Calponin-2 | CNN2 | | 1.26644 | -0.396193 | Q99439;Q99439-2 | Q99439;Q99439-2 |
| GDP-mannose 4,6 dehydratase | GMDS | | 1.12005 | -0.39447 | O60547-2;O60547 | O60547-2;O60547 |
| Adenosine deaminase | ADA | + | 2.16917 | -0.393784 | P00813 | P00813 |
| Actin-related protein 2/3 complex subunit 5-like protein | ARPC5L | | 0.545304 | -0.393585 | Q9BPX5 | Q9BPX5 |
| CD166 antigen | ALCAM | | 1.09503 | -0.392412 | Q13740-2;Q13740;Q13740-4;Q13740-3 | Q13740-2;Q13740 |
| Cullin-associated NEDD8-dissociated protein 1 | CAND1 | + | 1.67516 | -0.392235 | Q86VP6;Q86VP6-2;Q86VP6-3;O75155-2;O75155 | Q86VP6;Q86VP6-2 |

| | | | | | | |
|---|---|---|---|---|---|---|
| Acyl-CoA-binding protein | DBI | | 0.293371 | -0.390301 | P07108;P07108-3;P07108-2;P07108-4;P07108-5;P07108- | P07108;P07108-3;P07108-2;P07108-4;P07108-5 |
| Proteoglycan 4;Proteoglycan 4 C-terminal part | PRG4 | | 0.365195 | -0.389193 | Q92954-5;Q92954-4;Q92954-3;Q92954-6;Q92954-2;Q92! | Q92954-5;Q92954-4;Q92954-3;Q92954-6;Q92954-2 |
| Mannose-1-phosphate guanyltransferase beta | GMPPB | | 0.447508 | -0.389181 | Q9Y5P6;Q9Y5P6-2 | Q9Y5P6;Q9Y5P6-2 |
| Glyoxylate reductase/hydroxypyruvate reductase | GRHPR | | 0.427305 | -0.38717 | Q9UBQ7 | Q9UBQ7 |
| Proteasome subunit beta type-1 | PSMB1 | | 0.256527 | -0.385682 | P20618 | P20618 |
| Bifunctional ATP-dependent dihydroxyacetone kinase/FAD-AMP lyase (cyclizing);ATP-d | DAK | | 0.480211 | -0.384111 | Q3LXA3;Q3LXA3-2 | Q3LXA3;Q3LXA3-2 |
| Matrix metalloproteinase-14 | MMP14 | | 1.29095 | -0.383586 | P50281 | P50281 |
| Fructose-bisphosphate aldolase C | ALDOC | | 0.662132 | -0.383394 | P09972 | P09972 |
| 26S proteasome non-ATPase regulatory subunit 14 | PSMD14 | | 0.399993 | -0.383108 | O00487 | O00487 |
| NAD(P)H-hydrate epimerase | APOA1BP | | 0.354806 | -0.383101 | Q8NCW5;Q8NCW5-2 | Q8NCW5;Q8NCW5-2 |
| F-box only protein 2 | FBXO2 | + | 1.70506 | -0.382987 | Q9UK22 | Q9UK22 |
| Profilin-1 | PFN1 | | 1.01365 | -0.382617 | P07737 | P07737 |
| Ubiquitin-conjugating enzyme E2 K | UBE2K | + | 1.84236 | -0.377942 | P61086;P61086-2;P61086-3 | P61086;P61086-2;P61086-3 |
| Mannose-6-phosphate isomerase | MPI | | 0.745899 | -0.375549 | P34949;P34949-2 | P34949;P34949-2 |
| Vacuolar protein sorting-associated protein 26A | VPS26A | | 1.1264 | -0.375186 | O75436;O75436-2 | O75436;O75436-2 |
| Tenascin | TNC | | 0.262683 | -0.374786 | P24821;P24821-3 | P24821;P24821-3 |
| Tubulin polymerization-promoting protein family member 3 | TPPP3 | | 0.215658 | -0.374145 | Q9BW30 | Q9BW30 |
| Calcium-binding protein 39 | CAB39 | | 0.488681 | -0.368824 | Q9Y376;Q9H9S4 | Q9Y376 |
| N-alpha-acetyltransferase 50 | NAA50 | | 0.486587 | -0.368096 | Q9GZZ1;Q9GZZ1-2 | Q9GZZ1;Q9GZZ1-2 |
| Putative hydrolase RBBP9 | RBBP9 | | 0.228103 | -0.366553 | O75884;O75884-2 | O75884;O75884-2 |
| Spectrin alpha chain, non-erythrocytic 1 | SPTAN1 | + | 1.56075 | -0.365627 | Q13813 | Q13813 |
| Vacuolar protein sorting-associated protein 29 | VPS29 | | 0.785793 | -0.363687 | Q9UBQ0;Q9UBQ0-2 | Q9UBQ0;Q9UBQ0-2 |
| Interferon regulatory factor 2-binding protein-like | IRF2BPL | | 0.24679 | -0.363591 | Q9H1B7 | Q9H1B7 |
| Mitogen-activated protein kinase 3 | MAPK3 | | 0.189035 | -0.362624 | P27361;P27361-2;P27361-3;P31152;Q16659 | P27361;P27361-2;P27361-3 |
| Hepatoma-derived growth factor | HDGF | | 0.869455 | -0.361905 | P51858;P51858-2;P51858-3 | P51858;P51858-2;P51858-3 |
| Protein S100-A13 | S100A13 | | 0.42977 | -0.361609 | Q99584 | Q99584 |
| Ubiquitin thioesterase OTUB1 | OTUB1 | + | 1.32141 | -0.359695 | Q96FW1;Q96FW1-2 | Q96FW1 |
| Receptor-type tyrosine-protein phosphatase F | PTPRF | + | 1.43879 | -0.358247 | P10586-2;P10586 | P10586-2;P10586 |
| Lysosome membrane protein 2 | SCARB2 | | 0.2264 | -0.357393 | Q14108;Q14108-2 | Q14108;Q14108-2 |
| SH3 domain-binding glutamic acid-rich-like protein | SH3BGRL | | 0.768572 | -0.355015 | O75368 | O75368 |
| Desmocollin-2 | DSC2 | | 0.534586 | -0.354986 | Q02487;Q02487-2 | Q02487;Q02487-2 |
| Neurogenic locus notch homolog protein 3;Notch 3 extracellular truncation;Notch 3 int | NOTCH3 | | 0.525102 | -0.354896 | Q9UM47 | Q9UM47 |
| Aminoacylase-1 | ACY1 | | 0.97274 | -0.354764 | Q03154;Q03154-3;Q03154-4;Q03154-2 | Q03154;Q03154-3;Q03154-4;Q03154-2 |
| Arrestin domain-containing protein 1 | ARRDC1 | | 0.682572 | -0.354616 | Q8N5I2 | Q8N5I2 |
| Heme-binding protein 2 | HEBP2 | | 0.245604 | -0.352982 | Q9Y5Z4;Q9Y5Z4-2 | Q9Y5Z4;Q9Y5Z4-2 |
| Phospholipid transfer protein | PLTP | | 0.194208 | -0.347058 | P55058;P55058-3;P55058-2 | P55058;P55058-3;P55058-2 |
| EF-hand domain-containing protein D2 | EFHD2 | | 0.751893 | -0.346107 | Q96C19;Q9BUP0-2;Q9BUP0 | Q96C19 |
| Glucose-6-phosphate isomerase | GPI | | 0.434694 | -0.34484 | P06744;P06744-2;Q8N196 | P06744;P06744-2 |
| Calpain-2 catalytic subunit | CAPN2 | | 1.18126 | -0.341704 | P17655;P17655-2 | P17655;P17655-2 |
| C-Jun-amino-terminal kinase-interacting protein 4 | SPAG9 | | 0.366306 | -0.340197 | O60271;O60271-4;O60271-2;O60271-9;O60271-3;O6027 | O60271;O60271-4;O60271-2;O60271-9;O60271-3;C |
| V-type proton ATPase subunit H | ATP6V1H | | 1.019 | -0.340005 | Q9UI12-2;Q9UI12 | Q9UI12-2;Q9UI12 |
| JmjC domain-containing protein 8 | JMJD8 | | 0.167195 | -0.339271 | Q96S16-2;Q96S16 | Q96S16-2;Q96S16 |
| Crk-like protein | CRKL | | 0.870323 | -0.338423 | P46109 | P46109 |
| Hepatocyte growth factor-regulated tyrosine kinase substrate | HGS | | 0.528964 | -0.337708 | O14964;O14964-2 | O14964;O14964-2 |
| Xaa-Pro dipeptidase | PEPD | + | 1.62428 | -0.336175 | P12955;P12955-2;P12955-3 | P12955;P12955-2;P12955-3 |
| Tyrosine-protein phosphatase non-receptor type substrate 1;Signal-regulatory protein I | SIRPA;SIRPB1 | | 0.294624 | -0.334324 | P78324-4;P78324;P78324-2;Q5TFQ8 | P78324-4;P78324;P78324-2;Q5TFQ8 |
| Ribonuclease inhibitor | RNH1 | | 0.790679 | -0.333783 | P13489 | P13489 |
| Mitochondrial peptide methionine sulfoxide reductase | MSRA | | 0.346465 | -0.333485 | Q9UJ68-2;Q9UJ68-3;Q9UJ68-5;Q9UJ68;Q9UJ68-4 | Q9UJ68-2;Q9UJ68-3;Q9UJ68-5;Q9UJ68;Q9UJ68-4 |
| Filamin-A | FLNA | + | 2.27392 | -0.333382 | P21333-2;P21333 | P21333-2;P21333 |
| Transitional endoplasmic reticulum ATPase | VCP | | 0.813654 | -0.331209 | P55072 | P55072 |
| COP9 signalosome complex subunit 4 | COPS4 | | 0.712135 | -0.330816 | Q9BT78;Q9BT78-2 | Q9BT78;Q9BT78-2 |
| Peptidyl-prolyl cis-trans isomerase FKBP2 | FKBP2 | | 0.711266 | -0.330671 | P26885 | P26885 |
| Serine/threonine-protein phosphatase PP1-beta catalytic subunit | PPP1CB | | 0.455301 | -0.330437 | P62140 | P62140 |
| Fibroblast growth factor-binding protein 1 | FGFBP1 | | 0.305908 | -0.329186 | Q14512 | Q14512 |
| Prolyl endopeptidase FAP;Antiplasmin-cleaving enzyme FAP, soluble form | FAP | | 0.622535 | -0.328833 | Q12884;Q12884-2 | Q12884 |
| Protein ERGIC-53 | LMAN1 | | 0.0741601 | -0.328311 | P49257 | P49257 |
| Enoyl-CoA delta isomerase 1, mitochondrial | ECI1 | | 0.261675 | -0.327847 | P42126;P42126-2 | P42126;P42126-2 |
| Aldehyde dehydrogenase family 16 member A1 | ALDH16A1 | | 0.414026 | -0.327311 | Q8IZ83-3;Q8IZ83;Q8IZ83-2 | Q8IZ83-3;Q8IZ83 |
| Synaptic vesicle membrane protein VAT-1 homolog | VAT1 | | 0.661327 | -0.3262 | Q99536;Q99536-3;Q99536-2 | Q99536;Q99536-3;Q99536-2 |
| Acylamino-acid-releasing enzyme | APEH | | 0.642578 | -0.325823 | P13798 | P13798 |
| Sulfhydryl oxidase 1 | QSOX1 | | 0.837868 | -0.325373 | O00391;O00391-2 | O00391;O00391-2 |
| AP-1 complex subunit sigma-1A | AP1S1 | | 0.425464 | -0.324925 | P61966;P61966-2 | P61966;P61966-2 |
| Isoleucine--tRNA ligase, cytoplasmic | IARS | | 0.456142 | -0.324422 | P41252 | P41252 |
| Cofilin-2 | CFL2 | | 0.666702 | -0.323152 | Q9Y281;Q9Y281-3 | Q9Y281;Q9Y281-3 |
| Serine/threonine-protein phosphatase 2A catalytic subunit alpha isoform | PPP2CA | | 0.890022 | -0.323039 | P67775;P67775-2 | P67775;P67775-2 |
| Adapter molecule crk | CRK | | 0.550382 | -0.320288 | P46108;P46108-2 | P46108;P46108-2 |
| Metalloproteinase inhibitor 4 | TIMP4 | | 1.04072 | -0.319143 | Q99727 | Q99727 |
| Aspartate aminotransferase, mitochondrial | GOT2 | | 0.286183 | -0.318821 | P00505;P00505-2 | P00505;P00505-2 |

| | | | | | | |
|---|---|---|---|---|---|---|
| Isoamyl acetate-hydrolyzing esterase 1 homolog | IAH1 | | 0.681791 | -0.316839 | Q2TAA2;Q2TAA2-2 | Q2TAA2;Q2TAA2-2 |
| Very-long-chain 3-oxoacyl-CoA reductase | HSD17B12 | | 0.102002 | -0.315612 | Q53GQ0 | Q53GQ0 |
| Glutathione S-transferase P | GSTP1 | | 1.21981 | -0.311709 | P09211 | P09211 |
| Syntaxin-binding protein 2 | STXBP2 | | 0.384689 | -0.310005 | Q15833-2;Q15833;Q15833-3 | Q15833-2;Q15833;Q15833-3 |
| V-type proton ATPase subunit d 1 | ATP6V0D1 | | 0.120723 | -0.308145 | P61421 | P61421 |
| Ubiquitin-like modifier-activating enzyme 1 | UBA1 | | 0.744333 | -0.306438 | P22314-2 | P22314-2 |
| 26S proteasome non-ATPase regulatory subunit 13 | PSMD13 | | 0.452462 | -0.305223 | Q9UNM6;Q9UNM6-2 | Q9UNM6;Q9UNM6-2 |
| Catenin beta-1 | CTNNB1 | | 0.76007 | -0.304719 | P35222;P35222-2 | P35222 |
| Glutathione synthetase | GSS | | 0.388001 | -0.302087 | P48637;P48637-2 | P48637;P48637-2 |
| Trifunctional purine biosynthetic protein adenosine-3;Phosphoribosylamine--glycine lig | GART | | 0.505196 | -0.299582 | P22102;P22102-2 | P22102;P22102-2 |
| 14 kDa phosphohistidine phosphatase | PHPT1 | | 0.408514 | -0.297334 | Q9NRX4;Q9NRX4-2 | Q9NRX4;Q9NRX4-2 |
| Proteasome subunit beta type-2 | PSMB2 | | 0.201621 | -0.295456 | P49721 | P49721 |
| Enolase-phosphatase E1 | ENOPH1 | | 0.406862 | -0.295233 | Q9UHY7;Q9UHY7-2 | Q9UHY7;Q9UHY7-2 |
| Phospholipase A-2-activating protein | PLAA | | 0.700914 | -0.295129 | Q9Y263 | Q9Y263 |
| Carbonyl reductase [NADPH] 1 | CBR1 | | 0.190032 | -0.294782 | P16152;P16152-2 | P16152;P16152-2 |
| Importin subunit beta-1 | KPNB1 | + | 2.49366 | -0.293734 | Q14974;Q14974-2 | Q14974;Q14974-2 |
| Nucleoside diphosphate kinase B | NME2 | | 0.349924 | -0.293087 | P22392-2;P22392 | P22392-2;P22392 |
| Extracellular serine/threonine protein kinase FAM20C | FAM20C | | 0.307794 | -0.292126 | Q8IXL6;Q8IXL6-2;Q96MK3 | Q8IXL6 |
| Proteasome subunit alpha type-6 | PSMA6 | | 0.208712 | -0.290852 | P60900;P60900-2;P60900-3 | P60900;P60900-2;P60900-3 |
| ATP-citrate synthase | ACLY | + | 1.83065 | -0.289495 | P53396;P53396-2;P53396-3 | P53396;P53396-2;P53396-3 |
| Phosphoglycerate kinase 1 | PGK1 | | 0.81664 | -0.287437 | P00558;P00558-2 | P00558;P00558-2 |
| Tyrosine--tRNA ligase, cytoplasmic;Tyrosine--tRNA ligase, cytoplasmic, N-terminally prc | YARS | | 0.709484 | -0.286285 | P54577 | P54577 |
| Lysine--tRNA ligase | KARS | | 0.58239 | -0.285632 | Q15046;Q15046-2 | Q15046;Q15046-2 |
| Nck-associated protein 1 | NCKAP1 | | 0.522263 | -0.285269 | Q9Y2A7;Q9Y2A7-2;P55160 | Q9Y2A7;Q9Y2A7-2 |
| Methionine--tRNA ligase, cytoplasmic | MARS | | 0.366113 | -0.284849 | P56192;P56192-2 | P56192;P56192-2 |
| Charged multivesicular body protein 1a | CHMP1A | | 0.0999742 | -0.284482 | Q9HD42 | Q9HD42 |
| MOB kinase activator 1A;MOB kinase activator 1B | MOB1A;MOB1B | | 0.199597 | -0.282708 | Q9H8S9;Q7L9L4;Q7L9L4-2;Q9H8S9-2 | Q9H8S9;Q7L9L4;Q7L9L4-2;Q9H8S9-2 |
| Rabankyrin-5 | ANKFY1 | | 0.556572 | -0.282214 | Q9P2R3;Q9P2R3-2;Q9P2R3-4 | Q9P2R3;Q9P2R3-2;Q9P2R3-4 |
| Alpha-enolase | ENO1 | | 0.289643 | -0.282038 | P06733 | P06733 |
| Fatty acid synthase;[Acyl-carrier-protein] S-acetyltransferase;[Acyl-carrier-protein] S-m | FASN | + | 2.22403 | -0.281244 | P49327 | P49327 |
| Polypeptide N-acetylgalactosaminyltransferase 3 | GALNT3 | | 0.586867 | -0.280232 | Q14435;Q14435-2 | Q14435 |
| STE20-like serine/threonine-protein kinase | SLK | | 0.115806 | -0.280031 | Q9H2G2;Q9H2G2-2 | Q9H2G2;Q9H2G2-2 |
| PRA1 family protein 3 | ARL6IP5 | | 0.131571 | -0.277478 | O75915 | O75915 |
| Syntaxin-binding protein 3 | STXBP3 | | 0.310385 | -0.274291 | O00186 | O00186 |
| LIM and SH3 domain protein 1 | LASP1 | | 0.847796 | -0.273146 | Q14847;Q14847-2;Q14847-3 | Q14847;Q14847-2;Q14847-3 |
| UPF0565 protein C2orf69 | C2orf69 | | 0.293682 | -0.272924 | Q8N8R5 | Q8N8R5 |
| 26S proteasome non-ATPase regulatory subunit 11 | PSMD11 | | 0.933595 | -0.272241 | O00231;O00231-2 | O00231;O00231-2 |
| Vacuolar protein sorting-associated protein 35 | VPS35 | | 0.489087 | -0.270819 | Q96QK1 | Q96QK1 |
| Structural maintenance of chromosomes protein 3 | SMC3 | | 0.615255 | -0.270624 | Q9UQE7 | Q9UQE7 |
| ATP-dependent 6-phosphofructokinase, platelet type | PFKP | | 0.427985 | -0.269496 | Q01813;Q01813-2 | Q01813;Q01813-2 |
| 3-hydroxyisobutyryl-CoA hydrolase, mitochondrial | HIBCH | | 0.301565 | -0.269377 | Q6NVY1;Q6NVY1-2 | Q6NVY1;Q6NVY1-2 |
| CLIP-associating protein 1 | CLASP1 | | 0.250856 | -0.269037 | Q7Z460-2;Q7Z460-4;Q7Z460-5;Q7Z460-3;Q7Z460 | Q7Z460-2;Q7Z460-4;Q7Z460-5;Q7Z460-3;Q7Z460 |
| Tumor necrosis factor receptor superfamily member 11B | TNFRSF11B | | 0.192004 | -0.268719 | O00300 | O00300 |
| Small nuclear ribonucleoprotein E | SNRPE | | 0.029539 | -0.268278 | P62304 | P62304 |
| Mitogen-activated protein kinase 14 | MAPK14 | | 1.00174 | -0.265547 | Q16539;Q16539-2;Q16539-3;Q16539-5;Q16539-4 | Q16539;Q16539-2;Q16539-3;Q16539-5;Q16539-4 |
| Proteasome subunit beta type-6 | PSMB6 | | 0.223925 | -0.264434 | P28072 | P28072 |
| Epidermal growth factor receptor kinase substrate 8-like protein 2 | EPS8L2 | | 0.182084 | -0.260222 | Q9H6S3;Q9H6S3-3;Q9H6S3-2 | Q9H6S3;Q9H6S3-3;Q9H6S3-2 |
| Metalloproteinase inhibitor 3 | TIMP3 | | 0.361962 | -0.259763 | P35625 | P35625 |
| Cadherin-1;E-Cad/CTF1;E-Cad/CTF2;E-Cad/CTF3 | CDH1 | | 0.409766 | -0.258619 | P12830;P12830-2 | P12830;P12830-2 |
| Adenylyl cyclase-associated protein 1 | CAP1 | | 0.0296987 | -0.258454 | Q01518 | Q01518 |
| DnaJ homolog subfamily B member 4 | DNAJB4 | | 0.373504 | -0.254965 | Q9UDY4 | Q9UDY4 |
| Multifunctional protein ADE2;Phosphoribosylaminoimidazole-succinocarboxamide synt | PAICS | | 0.238172 | -0.254773 | P22234;P22234-2 | P22234;P22234-2 |
| Hypoxanthine-guanine phosphoribosyltransferase | HPRT1 | | 0.345111 | -0.254695 | P00492 | P00492 |
| Protein S100-A11;Protein S100-A11, N-terminally processed | S100A11 | | 0.0676874 | -0.254214 | P31949 | P31949 |
| DCC-interacting protein 13-alpha | APPL1 | | 0.610904 | -0.252649 | Q9UKG1 | Q9UKG1 |
| Peptidyl-prolyl cis-trans isomerase FKBP4;Peptidyl-prolyl cis-trans isomerase FKBP4, N- | FKBP4 | | 1.16859 | -0.251085 | Q02790 | Q02790 |
| Mannan-binding lectin serine protease 1;Mannan-binding lectin serine protease 1 heav | MASP1 | | 0.19971 | -0.250343 | P48740-2;P48740-3;P48740;P48740-4 | P48740-2;P48740-3;P48740;P48740-4 |
| Syndecan-4 | SDC4 | | 0.249114 | -0.248973 | P31431;P31431-2 | P31431;P31431-2 |
| Proteasome subunit beta type-3 | PSMB3 | | 0.184563 | -0.246567 | P49720 | P49720 |
| Aminopeptidase B | RNPEP | | 0.562339 | -0.244968 | Q9H4A4 | Q9H4A4 |
| S-phase kinase-associated protein 1 | SKP1 | | 0.49589 | -0.244623 | P63208;P63208-2 | P63208;P63208-2 |
| Twisted gastrulation protein homolog 1 | TWSG1 | | 0.139671 | -0.243076 | Q9GZX9;Q9GZX9-2 | Q9GZX9 |
| Galectin-3 | LGALS3 | | 0.246037 | -0.242435 | P17931 | P17931 |
| Peptidase M20 domain-containing protein 2 | PM20D2 | | 0.583147 | -0.240839 | Q8IYS1 | Q8IYS1 |
| Paired amphipathic helix protein Sin3a | SIN3A | | 0.199346 | -0.240623 | Q96ST3;O75182-2 | Q96ST3 |
| Cofilin-1 | CFL1 | | 1.11848 | -0.240253 | P23528 | P23528 |
| Arginine--tRNA ligase, cytoplasmic | RARS | | 0.577493 | -0.240115 | P54136;P54136-2 | P54136;P54136-2 |
| Phosphoglucomutase-2 | PGM2 | | 0.477992 | -0.239581 | Q96G03;Q96G03-2 | Q96G03 |

| | | | | | |
|---|---|---|---|---|---|
| Insulin-like growth factor-binding protein 6 | IGFBP6 | 0.217632 | -0.239573 | P24592 | P24592 |
| Superoxide dismutase [Cu-Zn] | SOD1 | 0.23901 | -0.238618 | P00441 | P00441 |
| Cyclin-dependent kinase 6 | CDK6 | 0.21587 | -0.238553 | Q00534;P11802-2;P11802;Q96Q40-2;Q96Q40-4;Q96Q40 | Q00534 |
| Hsp90 co-chaperone Cdc37;Hsp90 co-chaperone Cdc37, N-terminally processed | CDC37 | 0.213523 | -0.237194 | Q16543 | Q16543 |
| Retinoic acid-induced protein 3 | GPRC5A | 0.407755 | -0.236999 | Q8NFJ5 | Q8NFJ5 |
| Nucleolysin TIAR | TIAL1 | 0.548387 | -0.235874 | Q01085 | Q01085 |
| Stanniocalcin-2 | STC2 | 0.125875 | -0.235281 | O76061 | O76061 |
| Adenylosuccinate synthetase isozyme 2 | ADSS | 0.883421 | -0.233067 | P30520 | P30520 |
| Protein Hikeshi | C11orf73 | 0.0938041 | -0.232478 | Q53FT3 | Q53FT3 |
| Heme oxygenase 1 | HMOX1 | 0.203796 | -0.2299 | P09601 | P09601 |
| Guanine nucleotide-binding protein subunit beta-2-like 1;Guanine nucleotide-binding p | GNB2L1 | 0.733283 | -0.229221 | P63244 | P63244 |
| Endoplasmic reticulum aminopeptidase 1 | ERAP1 | 0.216057 | -0.229092 | Q9NZ08;Q9NZ08-2 | Q9NZ08;Q9NZ08-2 |
| Transducin-like enhancer protein 3 | TLE3 | 0.630119 | -0.227466 | Q04726-2;Q04726-7;Q04726-3;Q04726-6;Q04726-5;Q04 | Q04726-2;Q04726-7;Q04726-3;Q04726-6;Q04726-5 |
| Sorting nexin-12 | SNX12 | 0.600327 | -0.227344 | Q9UMY4-2;Q9UMY4;Q9UMY4-3 | Q9UMY4-2;Q9UMY4;Q9UMY4-3 |
| Deoxycytidylate deaminase | DCTD | 0.313298 | -0.226928 | P32321;P32321-2 | P32321;P32321-2 |
| Transforming growth factor-beta-induced protein ig-h3 | TGFBI | 0.34165 | -0.224867 | Q15582 | Q15582 |
| Keratin, type I cytoskeletal 18 | KRT18 | 0.240432 | -0.224466 | P05783;CON__H-INV:HIT000015463 | P05783 |
| Beta-1,4-galactosyltransferase 5 | B4GALT5 | 0.128641 | -0.224364 | O43286;Q9UBX8-2;Q9UBX8 | O43286 |
| Malate dehydrogenase, cytoplasmic | MDH1 | 0.179037 | -0.223259 | P40925;P40925-3;P40925-2 | P40925;P40925-3;P40925-2 |
| Obg-like ATPase 1 | OLA1 | 0.502196 | -0.222987 | Q9NTK5;Q9NTK5-2 | Q9NTK5;Q9NTK5-2 |
| Interleukin-18 | IL18 | 0.436566 | -0.222687 | Q14116;Q14116-2 | Q14116;Q14116-2 |
| Annexin A8-like protein 2 | ANXA8L2 | 0.139855 | -0.221551 | Q5VT79;Q5VT79-2 | Q5VT79;Q5VT79-2 |
| Rho GTPase-activating protein 1 | ARHGAP1 | 0.89929 | -0.220899 | Q07960 | Q07960 |
| Rap1 GTPase-GDP dissociation stimulator 1 | RAP1GDS1 | 0.544109 | -0.220893 | P52306;P52306-5;P52306-4;P52306-2;P52306-6;P52306- | P52306;P52306-5;P52306-4;P52306-2;P52306-6;P5 |
| Profilin-2 | PFN2 | 0.553769 | -0.218723 | P35080-2 | P35080-2 |
| Tyrosine-protein phosphatase non-receptor type 23 | PTPN23 | 0.619575 | -0.217782 | Q9H3S7 | Q9H3S7 |
| Roundabout homolog 1 | ROBO1 | 1.26844 | -0.213984 | Q9Y6N7-6;Q9Y6N7-5;Q9Y6N7-4;Q9Y6N7-3;Q9Y6N7;Q9Y6 | Q9Y6N7-6;Q9Y6N7-5;Q9Y6N7-4;Q9Y6N7-3;Q9Y6N7 |
| L-lactate dehydrogenase B chain | LDHB | 0.293327 | -0.213484 | P07195 | P07195 |
| IST1 homolog | IST1 | 0.534593 | -0.213412 | P53990-2;P53990;P53990-4;P53990-3;P53990-5;P53990- | P53990-2;P53990;P53990-4;P53990-3;P53990-5 |
| Galectin-1 | LGALS1 | 0.57241 | -0.211021 | P09382 | P09382 |
| Rab GDP dissociation inhibitor beta | GDI2 | 1.09797 | -0.209865 | P50395;P50395-2 | P50395;P50395-2 |
| Fructose-bisphosphate aldolase A | ALDOA | 0.534883 | -0.207296 | P04075 | P04075 |
| Inosine-5-monophosphate dehydrogenase 2 | IMPDH2 | 0.198226 | -0.202982 | P12268 | P12268 |
| Heterogeneous nuclear ribonucleoprotein L | HNRNPL | 0.235826 | -0.202317 | P14866;P14866-2 | P14866;P14866-2 |
| Methionine adenosyltransferase 2 subunit beta | MAT2B | 0.221261 | -0.202044 | Q9NZL9;Q9NZL9-4;Q9NZL9-2;Q9NZL9-3;Q9NZL9-5 | Q9NZL9;Q9NZL9-4;Q9NZL9-2;Q9NZL9-3 |
| Complement component 1 Q subcomponent-binding protein, mitochondrial | C1QBP | 0.145243 | -0.201248 | Q07021 | Q07021 |
| Vacuolar protein-sorting-associated protein 36 | VPS36 | 0.0675532 | -0.200666 | Q86VN1;Q86VN1-2 | Q86VN1;Q86VN1-2 |
| N-alpha-acetyltransferase 15, NatA auxiliary subunit | NAA15 | 0.438254 | -0.198231 | Q9BXJ9;Q9BXJ9-4;Q6N069;Q6N069-5;Q6N069-4;Q6N069 | Q9BXJ9;Q9BXJ9-4 |
| Thioredoxin-like protein 1 | TXNL1 | 0.661288 | -0.19814 | O43396 | O43396 |
| Elongation factor 2 | EEF2 | 0.46964 | -0.197632 | P13639 | P13639 |
| Pyridoxal kinase | PDXK | 0.250564 | -0.197595 | O00764;O00764-3;O00764-2 | O00764;O00764-3;O00764-2 |
| Nucleoside diphosphate kinase 3 | NME3 | 0.076727 | -0.197157 | Q13232 | Q13232 |
| Septin-11 | SEPT11 | 0.432339 | -0.197036 | Q9NVA2;Q9NVA2-2 | Q9NVA2;Q9NVA2-2 |
| High mobility group protein B1 | HMGB1 | 0.48403 | -0.196104 | P09429;B2RPK0 | P09429 |
| Ubiquitin-conjugating enzyme E2 N | UBE2N | 0.476621 | -0.195862 | P61088 | P61088 |
| Dynamin-2 | DNM2 | 0.160601 | -0.195459 | P50570-2;P50570-5;P50570;P50570-3;P50570-4;Q9UQ16 | P50570-2;P50570-5;P50570;P50570-3;P50570-4 |
| S-formylglutathione hydrolase | ESD | 0.264748 | -0.195098 | P10768 | P10768 |
| Dihydropyrimidinase-related protein 2 | DPYSL2 | 0.232272 | -0.194909 | Q16555;Q16555-2;Q14195;Q14194;Q14194-2 | Q16555;Q16555-2 |
| Amiloride-sensitive amine oxidase [copper-containing] | AOC1 | 0.0759274 | -0.194607 | P19801;P19801-2 | P19801;P19801-2 |
| Glutathione reductase, mitochondrial | GSR | 0.327356 | -0.193757 | P00390-2;P00390;P00390-3;P00390-4;P00390-5 | P00390-2;P00390;P00390-3;P00390-4;P00390-5 |
| Ras GTPase-activating-like protein IQGAP1 | IQGAP1 | 0.340529 | -0.193365 | P46940 | P46940 |
| Thioredoxin domain-containing protein 16 | TXNDC16 | 0.0862747 | -0.193338 | Q9P2K2 | Q9P2K2 |
| Calmodulin | CALM1 | 0.149366 | -0.193222 | P62158 | P62158 |
| Kunitz-type protease inhibitor 2 | SPINT2 | 0.1533 | -0.190626 | O43291;O43291-2 | O43291;O43291-2 |
| Protein S100-A16 | S100A16 | 0.156299 | -0.190465 | Q96FQ6 | Q96FQ6 |
| Galectin-3-binding protein | LGALS3BP | 0.258562 | -0.187019 | Q08380 | Q08380 |
| Nectin-1 | PVRL1 | 0.184623 | -0.186567 | Q15223;Q15223-3;Q15223-2 | Q15223;Q15223-3;Q15223-2 |
| AP-1 complex subunit mu-1 | AP1M1 | 0.150338 | -0.185857 | Q9BXS5;Q9BXS5-2 | Q9BXS5;Q9BXS5-2 |
| Protein S100-A6 | S100A6 | 0.0472939 | -0.185453 | P06703 | P06703 |
| ADP-ribosylation factor-like protein 2 | ARL2 | 0.182818 | -0.185163 | P36404;P36404-2 | P36404;P36404-2 |
| Olfactomedin-like protein 3 | OLFML3 | 0.19283 | -0.184645 | Q9NRN5;Q9NRN5-2;Q9NRN5-3 | Q9NRN5;Q9NRN5-2 |
| Poly(rC)-binding protein 2 | PCBP2 | 0.288284 | -0.182471 | Q15366-3;Q15366-4;Q15366-6;Q15366-7;P57721-5;P577 | Q15366-3;Q15366-4;Q15366-6;Q15366-7 |
| Serine/threonine-protein kinase MRCK beta | CDC42BPB | 0.246272 | -0.181373 | Q9Y5S2 | Q9Y5S2 |
| Putative nucleoside diphosphate kinase | NME2P1 | 0.0286825 | -0.180725 | O60361 | O60361 |
| Electron transfer flavoprotein subunit alpha, mitochondrial | ETFA | 0.187728 | -0.180023 | P13804;P13804-2 | P13804;P13804-2 |
| Alpha/beta hydrolase domain-containing protein 14B | ABHD14B | 0.284748 | -0.179572 | Q96IU4;Q96IU4-2 | Q96IU4 |
| Heat shock protein HSP 90-beta | HSP90AB1 | 0.55662 | -0.179298 | P08238;Q58FF7 | P08238 |
| Ubiquitin-conjugating enzyme E2 D2;Ubiquitin-conjugating enzyme E2 D3 | UBE2D2;UBE2D3 | 0.0683688 | -0.176679 | P62837-2;P62837;P61077;P61077-2;P61077-3 | P62837-2;P62837;P61077;P61077-2;P61077-3 |

| | | | | | |
|---|---|---|---|---|---|
| Follistatin-related protein 1 | FSTL1 | 0.244059 | -0.175709 | Q12841;Q12841-2 | Q12841;Q12841-2 |
| Rho GDP-dissociation inhibitor 2 | ARHGDIB | 0.324734 | -0.171783 | P52566 | P52566 |
| Ganglioside GM2 activator;Ganglioside GM2 activator isoform short | GM2A | 0.119222 | -0.170817 | P17900 | P17900 |
| Glycogen phosphorylase, liver form | PYGL | 0.743906 | -0.170328 | P06737-2;P06737 | P06737-2;P06737 |
| Collagen alpha-2(I) chain | COL1A2 | 0.312386 | -0.1703 | P08123 | P08123 |
| Cartilage oligomeric matrix protein | COMP | 0.124058 | -0.170266 | P49747;P49747-2 | P49747;P49747-2 |
| Proteasome subunit beta type-5 | PSMB5 | 0.0872996 | -0.16979 | P28074;P28074-2 | P28074 |
| Eukaryotic translation initiation factor 2A;Eukaryotic translation initiation factor 2A, N-t | EIF2A | 0.237451 | -0.169299 | Q9BY44;Q9BY44-3;Q9BY44-4;Q9BY44-2 | Q9BY44;Q9BY44-3;Q9BY44-4;Q9BY44-2 |
| ADP-sugar pyrophosphatase | NUDT5 | 0.442387 | -0.169116 | Q9UKK9 | Q9UKK9 |
| Pyridoxal phosphate phosphatase | PDXP | 0.612297 | -0.168017 | Q96GD0 | Q96GD0 |
| Glia-derived nexin | SERPINE2 | 0.496813 | -0.167557 | P07093-2;P07093-3 | P07093-2;P07093-3 |
| Chloride intracellular channel protein 1 | CLIC1 | 0.176721 | -0.167422 | O00299 | O00299 |
| Sorting nexin-2 | SNX2 | 0.162111 | -0.16714 | O60749;O60749-2 | O60749;O60749-2 |
| Phosphopantothenate--cysteine ligase | PPCS | 0.396716 | -0.164241 | Q9HAB8;Q9HAB8-2 | Q9HAB8;Q9HAB8-2 |
| Putative adenosylhomocysteinase 2 | AHCYL1 | 0.137905 | -0.164176 | O43865;O43865-2;Q96HN2-4;Q96HN2-3;Q96HN2-2;Q96I | O43865;O43865-2 |
| LanC-like protein 1 | LANCL1 | 0.449343 | -0.163993 | O43813 | O43813 |
| Serpin H1 | SERPINH1 | 0.140865 | -0.163972 | P50454 | P50454 |
| Growth factor receptor-bound protein 2 | GRB2 | 0.305004 | -0.163184 | P62993;P62993-2 | P62993;P62993-2 |
| Protein SEC13 homolog | SEC13 | 0.22666 | -0.163158 | P55735;P55735-3;P55735-2 | P55735;P55735-3;P55735-2 |
| E3 ubiquitin-protein ligase LRSAM1 | LRSAM1 | 0.0570418 | -0.161114 | Q6UWE0-2;Q6UWE0 | Q6UWE0-2;Q6UWE0 |
| CAD protein;Glutamine-dependent carbamoyl-phosphate synthase;Aspartate carbamoy | CAD | 0.21305 | -0.160946 | P27708 | P27708 |
| Ubiquitin-conjugating enzyme E2 variant 2 | UBE2V2 | 0.182355 | -0.15995 | Q15819 | Q15819 |
| COP9 signalosome complex subunit 8 | COPS8 | 0.53134 | -0.157372 | Q99627-2;Q99627 | Q99627-2;Q99627 |
| Bifunctional purine biosynthesis protein PURH;Phosphoribosylaminoimidazolecarboxar | ATIC | 0.174186 | -0.156898 | P31939;P31939-2 | P31939;P31939-2 |
| Parkinson disease 7 domain-containing protein 1 | PDDC1 | 0.235272 | -0.156156 | Q8NB37;Q8NB37-3;Q8NB37-2 | Q8NB37;Q8NB37-3;Q8NB37-2 |
| InaD-like protein | INADL | 0.174816 | -0.154638 | Q8NI35-3;Q8NI35;Q8NI35-4;Q8NI35-2;Q8NI35-5 | Q8NI35-3;Q8NI35;Q8NI35-4;Q8NI35-2;Q8NI35-5 |
| Endothelial differentiation-related factor 1 | EDF1 | 0.135174 | -0.153671 | O60869-2;O60869-3;O60869 | O60869-2;O60869-3;O60869 |
| UPF0587 protein C1orf123 | C1orf123 | 0.193828 | -0.153135 | Q9NWV4 | Q9NWV4 |
| Omega-amidase NIT2 | NIT2 | 0.360389 | -0.151789 | Q9NQR4 | Q9NQR4 |
| CD276 antigen | CD276 | 0.211369 | -0.151373 | Q5ZPR3-4;Q5ZPR3;Q5ZPR3-3;Q5ZPR3-2 | Q5ZPR3-4;Q5ZPR3;Q5ZPR3-3;Q5ZPR3-2 |
| Importin subunit alpha-3 | KPNA4 | 0.117255 | -0.150389 | O00629 | O00629 |
| Alcohol dehydrogenase class-3 | ADH5 | 0.411495 | -0.148968 | P11766 | P11766 |
| ATP-dependent 6-phosphofructokinase, liver type | PFKL | 0.198166 | -0.148808 | P17858;P17858-2 | P17858;P17858-2 |
| Eukaryotic translation initiation factor 3 subunit K | EIF3K | 0.702593 | -0.147573 | Q9UBQ5;Q9UBQ5-2 | Q9UBQ5;Q9UBQ5-2 |
| Gamma-glutamylcyclotransferase | GGCT | 0.110504 | -0.145835 | O75223;O75223-3;O75223-2;O75223-4 | O75223;O75223-3 |
| Alpha-amylase 1;Pancreatic alpha-amylase;Alpha-amylase 2B | AMY1A;AMY2A;AMY2B | 0.180322 | -0.144612 | P04745;P04746;P19961;P19961-2;P04746-2 | P04745;P04746;P19961;P19961-2 |
| Nucleoside diphosphate kinase A | NME1 | 0.0706018 | -0.144183 | P15531;P15531-2 | P15531;P15531-2 |
| CCR4-NOT transcription complex subunit 1 | CNOT1 | 0.166649 | -0.142553 | A5YKK6;A5YKK6-2;A5YKK6-3;A5YKK6-4 | A5YKK6;A5YKK6-2;A5YKK6-3;A5YKK6-4 |
| Macrophage migration inhibitory factor | MIF | 0.0682716 | -0.141047 | P14174 | P14174 |
| Poliovirus receptor | PVR | 0.142911 | -0.138999 | P15151-3;P15151-2;P15151-4;P15151 | P15151-3;P15151-2;P15151-4;P15151 |
| Abl interactor 1 | ABI1 | 0.0475722 | -0.137393 | Q8IZP0-10;Q8IZP0-8;Q8IZP0-7;Q8IZP0-2;Q8IZP0-4;Q8IZP | Q8IZP0-10;Q8IZP0-8;Q8IZP0-7;Q8IZP0-2;Q8IZP0-4;( |
| Peroxiredoxin-4 | PRDX4 | 0.0673654 | -0.137126 | Q13162 | Q13162 |
| Polypyrimidine tract-binding protein 1 | PTBP1 | 0.283103 | -0.135583 | P26599;P26599-2;P26599-3 | P26599;P26599-2;P26599-3 |
| L-xylulose reductase | DCXR | 0.360543 | -0.135164 | Q7Z4W1 | Q7Z4W1 |
| Serine/threonine-protein phosphatase 2A 65 kDa regulatory subunit A alpha isoform | PPP2R1A | 0.317667 | -0.130686 | P30153 | P30153 |
| Cadherin-2 | CDH2 | 0.458044 | -0.129644 | P19022 | P19022 |
| Charged multivesicular body protein 2a | CHMP2A | 0.0437062 | -0.128326 | O43633 | O43633 |
| Ly6/PLAUR domain-containing protein 3 | LYPD3 | 0.192129 | -0.127672 | O95274 | O95274 |
| Delta-aminolevulinic acid dehydratase | ALAD | 0.153104 | -0.126851 | P13716;P13716-2 | P13716;P13716-2 |
| Platelet-activating factor acetylhydrolase IB subunit gamma | PAFAH1B3 | 0.207985 | -0.126385 | Q15102 | Q15102 |
| Myosin light polypeptide 6 | MYL6 | 0.09683 | -0.123692 | P60660-2 | P60660-2 |
| Protocadherin gamma-C3 | PCDHGC3 | 0.049848 | -0.123318 | Q9UN70-2;Q9UN70 | Q9UN70-2;Q9UN70 |
| Tyrosine-protein kinase CSK | CSK | 0.135646 | -0.12253 | P41240;P42679-3;P42679;P42679-2 | P41240 |
| Programmed cell death protein 5 | PDCD5 | 0.579289 | -0.121829 | O14737;O14737-2 | O14737;O14737-2 |
| Protein phosphatase 1 regulatory subunit 7 | PPP1R7 | 0.275593 | -0.121753 | Q15435;Q15435-2;Q15435-3;Q15435-5;Q15435-4 | Q15435;Q15435-2;Q15435-3;Q15435-5;Q15435-4 |
| Ras-related protein Rab-11B;Ras-related protein Rab-11A | RAB11B;RAB11A | 0.177548 | -0.120912 | Q15907;Q15907-2;P62491-2 | Q15907;Q15907-2;P62491-2 |
| Disks large homolog 1 | DLG1 | 0.489435 | -0.117709 | Q12959-4;Q12959-2 | Q12959-4;Q12959-2 |
| Delta(3,5)-Delta(2,4)-dienoyl-CoA isomerase, mitochondrial | ECH1 | 0.0637841 | -0.116508 | Q13011 | Q13011 |
| Malate dehydrogenase, mitochondrial | MDH2 | 0.0798694 | -0.115851 | P40926;P40926-2 | P40926;P40926-2 |
| Chondroadherin | CHAD | 0.0954477 | -0.115348 | O15335 | O15335 |
| Glutaredoxin-3 | GLRX3 | 0.186812 | -0.114956 | O76003 | O76003 |
| COP9 signalosome complex subunit 6 | COPS6 | 0.152017 | -0.114932 | Q7L5N1 | Q7L5N1 |
| 6-phosphogluconolactonase | PGLS | 0.284702 | -0.114755 | O95336 | O95336 |
| Nuclear transport factor 2 | NUTF2 | 0.22653 | -0.11444 | P61970 | P61970 |
| Glypican-6;Secreted glypican-6 | GPC6 | 0.0473248 | -0.110908 | Q9Y625;O75487-2;O75487 | Q9Y625 |
| Elongation factor 1-gamma | EEF1G | 0.311453 | -0.108678 | P26641;P26641-2 | P26641;P26641-2 |
| Guanine nucleotide-binding protein G(i) subunit alpha-1 | GNAI1 | 0.0611656 | -0.10766 | P63096;P63096-2 | P63096;P63096-2 |
| Peptidyl-prolyl cis-trans isomerase C | PPIC | 0.0783061 | -0.106673 | P45877 | P45877 |

| | | | | | |
|---|---|---|---|---|---|
| Aspartate--tRNA ligase, cytoplasmic | DARS | 0.558392 | -0.106453 | P14868;P14868-2 | P14868;P14868-2 |
| Integrin beta-4 | ITGB4 | 0.143789 | -0.105622 | P16144-4;P16144-2;P16144-3;P16144;P16144-5 | P16144-4;P16144-2;P16144-3;P16144;P16144-5 |
| Serine/threonine-protein phosphatase PP1-alpha catalytic subunit | PPP1CA | 0.411959 | -0.103896 | P62136;P62136-2;P62136-3 | P62136;P62136-2;P62136-3 |
| Insulin-degrading enzyme | IDE | 0.194782 | -0.103171 | P14735;P14735-2 | P14735;P14735-2 |
| Phosphomannomutase 2 | PMM2 | 0.232893 | -0.103046 | O15305;O15305-2 | O15305;O15305-2 |
| Sorbitol dehydrogenase | SORD | 0.22735 | -0.102677 | Q00796;Q00796-2 | Q00796 |
| Threonine--tRNA ligase, cytoplasmic | TARS | 0.332275 | -0.097496 | P26639;P26639-2;A2RTX5-2;A2RTX5 | P26639;P26639-2 |
| Glucosamine-6-phosphate isomerase 1 | GNPDA1 | 0.154105 | -0.0971025 | P46926;P46926-2 | P46926;P46926-2 |
| Carbonyl reductase [NADPH] 3 | CBR3 | 0.131267 | -0.0945034 | O75828 | O75828 |
| Heat shock protein 75 kDa, mitochondrial | TRAP1 | 0.198434 | -0.0939306 | Q12931-2;Q12931 | Q12931-2;Q12931 |
| Succinyl-CoA ligase [ADP-forming] subunit beta, mitochondrial | SUCLA2 | 0.095516 | -0.093565 | Q9P2R7-2;Q9P2R7 | Q9P2R7-2;Q9P2R7 |
| Synaptophysin-like protein 1 | SYPL1 | 0.0432974 | -0.0923754 | Q16563-2;Q16563 | Q16563-2;Q16563 |
| Translationally-controlled tumor protein | TPT1 | 0.0724641 | -0.0910136 | P13693;P13693-2;Q56UQ5 | P13693;P13693-2 |
| Inositol polyphosphate 5-phosphatase OCRL-1 | OCRL | 0.070959 | -0.0903676 | Q01968-2;Q01968 | Q01968-2;Q01968 |
| Radixin | RDX | 0.277741 | -0.0878048 | P35241;P35241-5;P35241-4;P35241-2 | P35241;P35241-5;P35241-4 |
| Neurogenic locus notch homolog protein 2;Notch 2 extracellular truncation;Notch 2 int | NOTCH2 | 0.0668601 | -0.0867303 | Q04721 | Q04721 |
| CMP-N-acetylneuraminate-beta-galactosamide-alpha-2,3-sialyltransferase 4 | ST3GAL4 | 0.0467228 | -0.0863113 | Q11206-3;Q11206-2;Q11206;Q11206-4;Q11206-7;Q1120 | Q11206-3;Q11206-2;Q11206;Q11206-4;Q11206-7;C |
| ATP-dependent RNA helicase DHX36 | DHX36 | 0.244909 | -0.0838712 | Q9H2U1-2;Q9H2U1;Q9H2U1-3 | Q9H2U1-2;Q9H2U1;Q9H2U1-3 |
| Heme-binding protein 1 | HEBP1 | 0.267218 | -0.0835787 | Q9NRV9 | Q9NRV9 |
| 60S acidic ribosomal protein P2 | RPLP2 | 0.147854 | -0.0803572 | P05387;P05386-2 | P05387 |
| Beta-centractin | ACTR1B | 0.0962445 | -0.0797952 | P42025 | P42025 |
| Alpha-1,6-mannosylglycoprotein 6-beta-N-acetylglucosaminyltransferase A | MGAT5 | 0.141701 | -0.079085 | Q09328 | Q09328 |
| Peptidyl-prolyl cis-trans isomerase B | PPIB | 0.0778556 | -0.0788021 | P23284 | P23284 |
| Enoyl-CoA hydratase, mitochondrial | ECHS1 | 0.125144 | -0.0784931 | P30084 | P30084 |
| Proteasome activator complex subunit 1 | PSME1 | 0.163587 | -0.0783043 | Q06323;Q06323-3;Q06323-2 | Q06323;Q06323-3;Q06323-2 |
| Protein disulfide-isomerase | P4HB | 0.0824068 | -0.0779425 | P07237 | P07237 |
| Integrin alpha-6;Integrin alpha-6 heavy chain;Integrin alpha-6 light chain;Processed inte | ITGA6 | 0.125655 | -0.0767899 | P23229-4;P23229-2;P23229-5;P23229-3;P23229-9;P2322! | P23229-4;P23229-2;P23229-5;P23229-3;P23229-9;F |
| Fascin | FSCN1 | 0.146321 | -0.0755596 | Q16658 | Q16658 |
| Plexin-B2 | PLXNB2 | 0.134706 | -0.07525 | O15031 | O15031 |
| Pyruvate kinase PKM | PKM | 0.156899 | -0.0745138 | P14618 | P14618 |
| Vacuolar protein sorting-associated protein VTA1 homolog | VTA1 | 0.13941 | -0.0730546 | Q9NP79;Q9NP79-2 | Q9NP79 |
| Alpha-taxilin | TXLNA | 0.046936 | -0.0712687 | P40222 | P40222 |
| Contactin-1 | CNTN1 | 0.281483 | -0.0692387 | Q12860-2;Q12860;Q12860-3 | Q12860-2;Q12860;Q12860-3 |
| Annexin A2;Putative annexin A2-like protein | ANXA2;ANXA2P2 | 0.128737 | -0.0687389 | P07355;P07355-2;A6NMY6 | P07355;P07355-2;A6NMY6 |
| Proteasome subunit alpha type-3 | PSMA3 | 0.0263909 | -0.0681025 | P25788-2;P25788 | P25788-2;P25788 |
| Protein disulfide-isomerase A3 | PDIA3 | 0.0413354 | -0.0644792 | P30101 | P30101 |
| Aspartate aminotransferase, cytoplasmic | GOT1 | 0.0563307 | -0.0638402 | P17174;P17174-2 | P17174;P17174-2 |
| Structural maintenance of chromosomes protein 2 | SMC2 | 0.0754785 | -0.0615368 | O95347;O95347-2 | O95347;O95347-2 |
| Ubiquitin-conjugating enzyme E2 variant 1 | UBE2V1 | 0.0863583 | -0.0605125 | Q13404;Q13404-8;Q13404-7;Q13404-2;Q13404-1;Q1340 | Q13404;Q13404-8;Q13404-7;Q13404-2;Q13404-1;C |
| Transportin-1 | TNPO1 | 0.0474416 | -0.0594953 | Q92973-2;Q92973;Q92973-3 | Q92973-2;Q92973;Q92973-3 |
| UMP-CMP kinase | CMPK1 | 0.0643089 | -0.0574462 | P30085;P30085-2 | P30085;P30085-2 |
| Tryptophan--tRNA ligase, cytoplasmic;T1-TrpRS;T2-TrpRS | WARS | 0.121495 | -0.0569839 | P23381;P23381-2 | P23381;P23381-2 |
| Calcyclin-binding protein | CACYBP | 0.323555 | -0.0567385 | Q9HB71;Q9HB71-3;Q9HB71-2 | Q9HB71;Q9HB71-3 |
| Cytoplasmic FMR1-interacting protein 1 | CYFIP1 | 0.0960489 | -0.0547918 | Q7L576;Q7L576-2;Q7L576-3 | Q7L576;Q7L576-2 |
| Myeloid-derived growth factor | MYDGF | 0.0877423 | -0.0544809 | Q969H8 | Q969H8 |
| Na(+)/H(+) exchange regulatory cofactor NHE-RF1 | SLC9A3R1 | 0.0291285 | -0.0544643 | O14745;O14745-2 | O14745;O14745-2 |
| Eukaryotic translation initiation factor 3 subunit B | EIF3B | 0.0950851 | -0.0521342 | P55884;P55884-2 | P55884;P55884-2 |
| Nesprin-2 | SYNE2 | 0.00515196 | -0.0510419 | Q8WXH0;Q8WXH0-2;Q8WXH0-7;Q8WXH0-4;Q8WXH0-3; | Q8WXH0;Q8WXH0-2 |
| Nidogen-2 | NID2 | 0.0479045 | -0.0507189 | Q14112-2 | Q14112-2 |
| DnaJ homolog subfamily C member 13 | DNAJC13 | 0.0561938 | -0.0498098 | O75165 | O75165 |
| Latent-transforming growth factor beta-binding protein 3 | LTBP3 | 0.0617076 | -0.048748 | Q9NS15 | Q9NS15 |
| Retinoic acid receptor responder protein 1 | RARRES1 | 0.0201936 | -0.0464331 | P49788;P49788-2 | P49788;P49788-2 |
| Cilia- and flagella-associated protein 20 | CFAP20 | 0.0347753 | -0.0456225 | Q9Y6A4 | Q9Y6A4 |
| Transgelin-2 | TAGLN2 | 0.103123 | -0.0454667 | P37802;P37802-2 | P37802;P37802-2 |
| Thioredoxin domain-containing protein 5 | TXNDC5 | 0.029062 | -0.0451946 | Q8NBS9;Q8NBS9-2 | Q8NBS9;Q8NBS9-2 |
| Ubiquitin carboxyl-terminal hydrolase isozyme L3 | UCHL3 | 0.078985 | -0.0449301 | P15374 | P15374 |
| Fumarate hydratase, mitochondrial | FH | 0.0435973 | -0.0443064 | P07954-2;P07954 | P07954-2;P07954 |
| Succinate dehydrogenase [ubiquinone] iron-sulfur subunit, mitochondrial | SDHB | 0.031972 | -0.04249 | P21912 | P21912 |
| Nuclear mitotic apparatus protein 1 | NUMA1 | 0.146723 | -0.042071 | Q14980;Q14980-2;Q14980-4;Q14980-3;Q14980-5 | Q14980;Q14980-2;Q14980-4;Q14980-3 |
| Alpha-centractin | ACTR1A | 0.0757336 | -0.0416686 | P61163 | P61163 |
| UTP--glucose-1-phosphate uridylyltransferase | UGP2 | 0.0935387 | -0.0404174 | Q16851 | Q16851 |
| Guanine nucleotide-binding protein-like 1 | GNL1 | 0.0655678 | -0.0390244 | P36915;P36915-2 | P36915;P36915-2 |
| Transforming growth factor beta-1;Latency-associated peptide | TGFB1 | 0.0440269 | -0.0376867 | P01137 | P01137 |
| Peroxiredoxin-1 | PRDX1 | 0.045309 | -0.0352542 | Q06830 | Q06830 |
| DNA damage-binding protein 1 | DDB1 | 0.0288865 | -0.0351957 | Q16531;Q16531-2 | Q16531 |
| Citrate synthase, mitochondrial | CS | 0.0222227 | -0.034373 | O75390 | O75390 |
| Acylphosphatase-1 | ACYP1 | 0.016961 | -0.0342948 | P07311;P07311-2 | P07311 |
| V-type proton ATPase subunit E 1 | ATP6V1E1 | 0.0186019 | -0.0311419 | P36543;P36543-2 | P36543;P36543-2 |

| | | | | | |
|---|---|---|---|---|---|
| Ubiquitin-conjugating enzyme E2 L3 | UBE2L3 | 0.0401545 | -0.0310103 | P68036;P68036-3;P68036-2 | P68036;P68036-3;P68036-2 |
| ATPase ASNA1 | ASNA1 | 0.0201397 | -0.0299416 | O43681 | O43681 |
| Junction plakoglobin | JUP | 0.0371158 | -0.0295296 | P14923 | P14923 |
| Fatty acid-binding protein, epidermal | FABP5 | 0.0590564 | -0.0291386 | Q01469;A8MUU1 | Q01469 |
| ATP-dependent RNA helicase DDX3X | DDX3X | 0.0131925 | -0.0255388 | O00571-2;O00571;Q9NQI0-3;Q9NQI0-2;Q9NQI0-4;Q9NQ | O00571-2;O00571 |
| D-dopachrome decarboxylase;D-dopachrome decarboxylase-like protein | DDT;DDTL | 0.0238381 | -0.0251452 | P30046;P30046-2;A6NHG4 | P30046;P30046-2;A6NHG4 |
| Activator of 90 kDa heat shock protein ATPase homolog 1 | AHSA1 | 0.0213792 | -0.0244382 | O95433;O95433-2 | O95433;O95433-2 |
| Heterogeneous nuclear ribonucleoprotein D0 | HNRNPD | 0.0502195 | -0.0226606 | Q14103-3;Q14103;Q14103-4;Q14103-2 | Q14103-3;Q14103;Q14103-4;Q14103-2 |
| Collagen alpha-1(I) chain | COL1A1 | 0.0179745 | -0.021939 | P02452;CON__Q862S4 | P02452 |
| Tetratricopeptide repeat protein 38 | TTC38 | 0.0312633 | -0.0213737 | Q5R3I4 | Q5R3I4 |
| DnaJ homolog subfamily B member 6 | DNAJB6 | 0.0223643 | -0.0210349 | O75190-3;O75190;O75190-2;Q8WWF6;O75190-4;Q7Z6W | O75190-3;O75190;O75190-2 |
| Apoptosis-associated speck-like protein containing a CARD | PYCARD | 0.0192404 | -0.0206159 | Q9ULZ3;Q9ULZ3-3;Q9ULZ3-2 | Q9ULZ3;Q9ULZ3-3;Q9ULZ3-2 |
| Peptidyl-prolyl cis-trans isomerase A;Peptidyl-prolyl cis-trans isomerase A, N-terminally | PPIA | 0.041066 | -0.0177555 | P62937;Q9Y536;F5H284 | P62937 |
| Pseudouridine-5-phosphatase | HDHD1 | 0.00448705 | -0.0157572 | Q08623-3;Q08623;Q08623-4;Q08623-2 | Q08623-3;Q08623;Q08623-4;Q08623-2 |
| Eukaryotic translation initiation factor 3 subunit H | EIF3H | 0.0141165 | -0.0151221 | O15372 | O15372 |
| Cytosolic non-specific dipeptidase | CNDP2 | 0.0240236 | -0.0133508 | Q96KP4;Q96KP4-2 | Q96KP4;Q96KP4-2 |
| Stress-70 protein, mitochondrial | HSPA9 | 0.0220444 | -0.0131429 | P38646 | P38646 |
| DnaJ homolog subfamily B member 1 | DNAJB1 | 0.00742283 | -0.0121517 | P25685;P25685-2 | P25685;P25685-2 |
| Coiled-coil domain-containing protein 25 | CCDC25 | 0.00931121 | -0.00976562 | Q86WR0;Q86WR0-2 | Q86WR0;Q86WR0-2 |
| Interleukin-1 receptor type 2;Interleukin-1 receptor type 2, membrane form;Interleukir | IL1R2 | 0.00527279 | -0.00961177 | P27930;P27930-2 | P27930;P27930-2 |
| Sepiapterin reductase | SPR | 0.00816665 | -0.00908597 | P35270 | P35270 |
| Tubulin alpha-4A chain | TUBA4A | 0.00791017 | -0.00732295 | P68366;P68366-2 | P68366;P68366-2 |
| Cullin-3 | CUL3 | 0.0134238 | -0.00716337 | Q13618-2;Q13618;Q13618-3 | Q13618-2;Q13618;Q13618-3 |
| Tubulin beta-3 chain | TUBB3 | 0.00454272 | -0.00669924 | Q13509;Q13509-2 | Q13509;Q13509-2 |
| Latent-transforming growth factor beta-binding protein 1 | LTBP1 | 0.00612057 | -0.00639598 | Q14766;Q14766-4;Q14766-2;Q14766-5;Q14766-3 | Q14766;Q14766-4;Q14766-2;Q14766-5;Q14766-3 |
| Alpha-1,3-mannosyl-glycoprotein 2-beta-N-acetylglucosaminyltransferase | MGAT1 | 0.00184292 | -0.00115458 | P26572 | P26572 |
| Metalloreductase STEAP4 | STEAP4 | 9.39E-05 | -0.000106812 | Q687X5;Q687X5-2 | Q687X5;Q687X5-2 |
| Heterogeneous nuclear ribonucleoprotein A1;Heterogeneous nuclear ribonucleoprotei | HNRNPA1 | 0.000479033 | 0.000240326 | P09651;P09651-2;P09651-3 | P09651;P09651-2;P09651-3 |
| CUB domain-containing protein 1 | CDCP1 | 0.00310576 | 0.00260607 | Q9H5V8;Q9H5V8-2 | Q9H5V8;Q9H5V8-2 |
| Electron transfer flavoprotein subunit beta | ETFB | 0.00173896 | 0.00274913 | P38117-2;P38117 | P38117-2;P38117 |
| Adenosylhomocysteinase | AHCY | 0.00413469 | 0.00386747 | P23526;P23526-2 | P23526;P23526-2 |
| V-type proton ATPase subunit F | ATP6V1F | 0.00186034 | 0.00542386 | Q16864;Q16864-2 | Q16864;Q16864-2 |
| Prenylcysteine oxidase 1 | PCYOX1 | 0.00223743 | 0.00802612 | Q9UHG3-2;Q9UHG3 | Q9UHG3-2;Q9UHG3 |
| Thrombospondin type-1 domain-containing protein 4 | THSD4 | 0.0174378 | 0.0115274 | Q6ZMP0;Q6ZMP0-4;Q6ZMP0-2 | Q6ZMP0;Q6ZMP0-4 |
| Phosphoserine aminotransferase | PSAT1 | 0.0248059 | 0.012921 | Q9Y617;Q9Y617-2 | Q9Y617;Q9Y617-2 |
| Transcription activator BRG1 | SMARCA4 | 0.017926 | 0.0132338 | P51532-5;P51532-2;P51532-3;P51532-4;P51532 | P51532-5;P51532-2;P51532-3;P51532-4;P51532 |
| Annexin A5 | ANXA5 | 0.0167973 | 0.014527 | P08758 | P08758 |
| Protein kinase C iota type | PRKCI | 0.0272265 | 0.0178324 | P41743 | P41743 |
| Desmoglein-2 | DSG2 | 0.051938 | 0.0201499 | Q14126 | Q14126 |
| Inositol monophosphatase 1 | IMPA1 | 0.0160025 | 0.0204875 | P29218;P29218-3;P29218-2 | P29218;P29218-3;P29218-2 |
| Brain-specific serine protease 4 | PRSS22 | 0.0278756 | 0.0210495 | Q9GZN4 | Q9GZN4 |
| Multivesicular body subunit 12A | MVB12A | 0.0590524 | 0.0226027 | Q96EY5;Q96EY5-3;Q96EY5-2 | Q96EY5;Q96EY5-3;Q96EY5-2 |
| Hsc70-interacting protein;Putative protein FAM10A4 | ST13;ST13P4 | 0.0746881 | 0.0259622 | P50502;Q8IZP2;Q8NFI4 | P50502;Q8IZP2 |
| Alpha-N-acetylgalactosaminidase | NAGA | 0.0240612 | 0.0264575 | P17050 | P17050 |
| GDP-L-fucose synthase | TSTA3 | 0.0199172 | 0.0274804 | Q13630 | Q13630 |
| Eukaryotic translation initiation factor 6 | EIF6 | 0.0232995 | 0.0291386 | P56537;P56537-2 | P56537;P56537-2 |
| Alpha-aminoadipic semialdehyde dehydrogenase | ALDH7A1 | 0.0385698 | 0.0305799 | P49419-2;P49419;P49419-4;P49419-3 | P49419-2;P49419;P49419-4 |
| Myosin-9 | MYH9 | 0.0243418 | 0.0315444 | P35579;P35579-2;Q9UKX2 | P35579;P35579-2 |
| Alpha-soluble NSF attachment protein | NAPA | 0.0782991 | 0.0326564 | P54920;Q9H115-3;Q9H115-2;Q9H115 | P54920 |
| Adenylate kinase 2, mitochondrial;Adenylate kinase 2, mitochondrial, N-terminally proc | AK2 | 0.031783 | 0.0351874 | P54819-2;P54819;P54819-5;P54819-3;P54819-6;P54819- | P54819-2;P54819;P54819-5;P54819-3;P54819-6;P5 |
| Heat shock protein HSP 90-alpha | HSP90AA1 | 0.0327638 | 0.0363083 | P07900;P07900-2;Q58FG0 | P07900;P07900-2 |
| Translocon-associated protein subunit delta | SSR4 | 0.0242938 | 0.0374699 | P51571 | P51571 |
| Ribulose-phosphate 3-epimerase | RPE | 0.0291645 | 0.0405725 | Q96AT9;Q96AT9-2;Q96AT9-5;Q96AT9-3;Q96AT9-4;Q2QD | Q96AT9;Q96AT9-2;Q96AT9-5;Q96AT9-3;Q96AT9-4 |
| Rab3 GTPase-activating protein catalytic subunit | RAB3GAP1 | 0.0518355 | 0.040638 | Q15042;Q15042-3;Q15042-4 | Q15042;Q15042-3;Q15042-4 |
| Complement decay-accelerating factor | CD55 | 0.0441445 | 0.041647 | P08174-4;P08174;P08174-3;P08174-5;P08174-2;P08174- | P08174-4;P08174;P08174-3;P08174-5;P08174-2;P0 |
| Ataxin-10 | ATXN10 | 0.0717848 | 0.0432529 | Q9UBB4;Q9UBB4-2 | Q9UBB4;Q9UBB4-2 |
| Protein disulfide-isomerase A6 | PDIA6 | 0.0323748 | 0.0440591 | Q15084-3;Q15084;Q15084-4;Q15084-5;Q15084-2 | Q15084-3;Q15084;Q15084-4;Q15084-5;Q15084-2 |
| Phosphatidylethanolamine-binding protein 1;Hippocampal cholinergic neurostimulatin | PEBP1 | 0.0397962 | 0.0447013 | P30086 | P30086 |
| MAGUK p55 subfamily member 5 | MPP5 | 0.0464293 | 0.0462564 | Q8N3R9-2;Q8N3R9 | Q8N3R9-2;Q8N3R9 |
| Triosephosphate isomerase | TPI1 | 0.0336888 | 0.0465476 | P60174-1;P60174;P60174-4 | P60174-1;P60174;P60174-4 |
| Desmoplakin | DSP | 0.079626 | 0.0470352 | P15924;P15924-3;P15924-2 | P15924;P15924-3;P15924-2 |
| Creatine kinase U-type, mitochondrial | CKMT1A | 0.0634609 | 0.0470606 | P12532;P17540 | P12532 |
| Proto-oncogene tyrosine-protein kinase Src | SRC | 0.0889244 | 0.0471516 | P12931;P12931-2;P06241-3;P06241-2;P06241;P08631-3; | P12931;P12931-2 |
| Complement factor H | CFH | 0.0427186 | 0.0471776 | P08603;P08603-2;Q02985-2;Q02985;Q03591 | P08603;P08603-2 |
| Peptidyl-prolyl cis-trans isomerase NIMA-interacting 4 | PIN4 | 0.0386425 | 0.0476182 | Q9Y237;Q9Y237-2;Q9Y237-3 | Q9Y237;Q9Y237-2 |
| Ubiquitin-fold modifier 1 | UFM1 | 0.122428 | 0.0477549 | P61960;P61960-2 | P61960;P61960-2 |
| Spermine synthase | SMS | 0.0786237 | 0.04813 | P52788;P52788-2 | P52788;P52788-2 |
| Receptor-type tyrosine-protein phosphatase gamma | PTPRG | 0.0651188 | 0.0520903 | P23470-2;P23470 | P23470-2;P23470 |

| | | | | | |
|---|---|---|---|---|---|
| Dipeptidyl peptidase 1;Dipeptidyl peptidase 1 exclusion domain chain;Dipeptidyl peptic | CTSC | 0.0686522 | 0.0540021 | P53634;P53634-2;P53634-3 | P53634 |
| Latent-transforming growth factor beta-binding protein 2 | LTBP2 | 0.0752707 | 0.0554498 | Q14767 | Q14767 |
| Gelsolin | GSN | 0.11217 | 0.055844 | P06396-2;P06396-4;P06396-3;P06396 | P06396-2;P06396-4;P06396-3;P06396 |
| ADP-ribose pyrophosphatase, mitochondrial | NUDT9 | 0.0575665 | 0.0565313 | Q9BW91-2;Q9BW91 | Q9BW91-2;Q9BW91 |
| CD151 antigen | CD151 | 0.0107794 | 0.0565999 | P48509 | P48509 |
| Isochorismatase domain-containing protein 1 | ISOC1 | 0.0374069 | 0.0571206 | Q96CN7 | Q96CN7 |
| Thiopurine S-methyltransferase | TPMT | 0.0257541 | 0.0608711 | P51580 | P51580 |
| Calreticulin | CALR | 0.024205 | 0.0635541 | P27797 | P27797 |
| Serine/threonine-protein kinase MARK2;MAP/microtubule affinity-regulating kinase 3 | MARK2;MARK3 | 0.108753 | 0.0648696 | Q7KZI7-10;Q7KZI7-13;Q7KZI7-7;Q7KZI7-2;Q7KZI7-15;Q7K | Q7KZI7-10;Q7KZI7-13;Q7KZI7-7;Q7KZI7-2;Q7KZI7-1 |
| High mobility group protein B2 | HMGB2 | 0.0586748 | 0.0655352 | P26583 | P26583 |
| Asparagine--tRNA ligase, cytoplasmic | NARS | 0.258895 | 0.0656446 | O43776;O43776-2 | O43776 |
| Sorcin | SRI | 0.0625537 | 0.0666663 | P30626-2;P30626;P30626-3 | P30626-2;P30626;P30626-3 |
| Ephrin type-A receptor 2 | EPHA2 | 0.185945 | 0.0676053 | P29317;P29317-2;Q15375-2;Q15375-4;Q15375;P54756-3 | P29317 |
| Thioredoxin-dependent peroxide reductase, mitochondrial | PRDX3 | 0.0472596 | 0.0696341 | P30048-2;P30048 | P30048-2;P30048 |
| Phosphatidylinositol 5-phosphate 4-kinase type-2 gamma | PIP4K2C | 0.0218032 | 0.0696608 | Q8TBX8-3;Q8TBX8;Q8TBX8-2 | Q8TBX8-3;Q8TBX8;Q8TBX8-2 |
| Caspase-14;Caspase-14 subunit p17, mature form;Caspase-14 subunit p10, mature forn | CASP14 | 0.0508589 | 0.0745805 | P31944 | P31944 |
| Protein Wnt-5a | WNT5A | 0.0643149 | 0.0755653 | P41221-2;P41221;O00755 | P41221-2;P41221 |
| Small nuclear ribonucleoprotein Sm D3 | SNRPD3 | 0.0387103 | 0.07831 | P62318-2;P62318 | P62318-2;P62318 |
| Core histone macro-H2A.1 | H2AFY | 0.0823976 | 0.0784683 | O75367-3;O75367;O75367-2;Q9P0M6 | O75367-3;O75367;O75367-2 |
| Eukaryotic translation initiation factor 3 subunit E | EIF3E | 0.196771 | 0.0788701 | P60228 | P60228 |
| 4-trimethylaminobutyraldehyde dehydrogenase | ALDH9A1 | 0.103114 | 0.079518 | P49189;P49189-2 | P49189;P49189-2 |
| Ribonucleoside-diphosphate reductase large subunit | RRM1 | 0.0966071 | 0.0798003 | P23921 | P23921 |
| Ras-related protein Rab-22A | RAB22A | 0.0216395 | 0.0823104 | Q9UL26 | Q9UL26 |
| F-actin-capping protein subunit alpha-2 | CAPZA2 | 0.0762328 | 0.0849177 | P47755;P47755-2 | P47755;P47755-2 |
| Beta-galactosidase | GLB1 | 0.434242 | 0.0868333 | P16278-3;P16278;P16278-2 | P16278-3;P16278;P16278-2 |
| COMM domain-containing protein 1 | COMMD1 | 0.0299733 | 0.0880807 | Q8N668;Q8N668-2 | Q8N668;Q8N668-2 |
| Cytosol aminopeptidase | LAP3 | 0.0454318 | 0.0886002 | P28838-2;P28838 | P28838-2;P28838 |
| N-acetyllactosaminide beta-1,3-N-acetylglucosaminyltransferase 2 | B3GNT2 | 0.165205 | 0.0889912 | Q9NY97-2;Q9NY97 | Q9NY97-2;Q9NY97 |
| Wiskott-Aldrich syndrome protein family member 2 | WASF2 | 0.0810366 | 0.0895138 | Q9Y6W5;Q9Y6W5-2 | Q9Y6W5;Q9Y6W5-2 |
| Vacuolar protein sorting-associated protein 4B | VPS4B | 0.0492131 | 0.0900459 | O75351 | O75351 |
| Protein NDRG1 | NDRG1 | 0.189523 | 0.0926641 | Q92597;Q92597-3;Q92597-2 | Q92597;Q92597-3;Q92597-2 |
| Peroxiredoxin-6 | PRDX6 | 0.843877 | 0.094628 | P30041 | P30041 |
| Protein tyrosine phosphatase type IVA 2;Protein tyrosine phosphatase type IVA 1 | PTP4A2;PTP4A1 | 0.177726 | 0.0948346 | Q12974;Q12974-3;Q12974-4;Q12974-2;Q93096 | Q12974;Q12974-3;Q12974-4;Q12974-2;Q93096 |
| Carbonic anhydrase 2 | CA2 | 0.121072 | 0.0969588 | P00918 | P00918 |
| Acetyl-CoA acetyltransferase, cytosolic | ACAT2 | 0.0598908 | 0.0978311 | Q9BWD1;Q9BWD1-2 | Q9BWD1;Q9BWD1-2 |
| Formin-binding protein 1-like | FNBP1L | 0.0613468 | 0.0983639 | Q5T0N5-3;Q5T0N5-4;Q5T0N5-2;Q5T0N5-5;Q5T0N5 | Q5T0N5-3;Q5T0N5-4;Q5T0N5-2;Q5T0N5-5;Q5T0N5 |
| Septin-7 | SEPT7 | 0.163461 | 0.100115 | Q16181;Q16181-2;Q6ZU15 | Q16181;Q16181-2 |
| Putative phospholipase B-like 2;Putative phospholipase B-like 2 32 kDa form;Putative p | PLBD2 | 0.15162 | 0.100552 | Q8NHP8;Q8NHP8-2 | Q8NHP8;Q8NHP8-2 |
| Laminin subunit beta-2 | LAMB2 | 0.174636 | 0.101008 | P55268;A4D0S4-2;A4D0S4-3;A4D0S4 | P55268 |
| ADP-ribosylation factor 1;ADP-ribosylation factor 3 | ARF1;ARF3 | 0.141926 | 0.101583 | P84077;P61204;P61204-2 | P84077;P61204;P61204-2 |
| Thrombospondin-4 | THBS4 | 0.117908 | 0.101636 | P35443 | P35443 |
| Proliferation-associated protein 2G4 | PA2G4 | 0.264758 | 0.102392 | Q9UQ80;Q9UQ80-2 | Q9UQ80;Q9UQ80-2 |
| Dipeptidyl peptidase 2 | DPP7 | 0.088269 | 0.102596 | Q9UHL4 | Q9UHL4 |
| U6 snRNA-associated Sm-like protein LSm6 | LSM6 | 0.0594113 | 0.105256 | P62312 | P62312 |
| Poly(rC)-binding protein 1 | PCBP1 | 0.238844 | 0.106972 | Q15365 | Q15365 |
| Proliferating cell nuclear antigen | PCNA | 0.213211 | 0.108044 | P12004 | P12004 |
| Cysteine-rich with EGF-like domain protein 2 | CRELD2 | 0.0468855 | 0.109776 | Q6UXH1-4;Q6UXH1;Q6UXH1-6;Q6UXH1-5;Q6UXH1-3;Q6 | Q6UXH1-4;Q6UXH1;Q6UXH1-6;Q6UXH1-5;Q6UXH1 |
| Isocitrate dehydrogenase [NADP] cytoplasmic | IDH1 | 0.095267 | 0.113555 | O75874 | O75874 |
| N-acetylgalactosamine-6-sulfatase | GALNS | 0.0871635 | 0.113683 | P34059 | P34059 |
| Serine/threonine-protein kinase 24;Serine/threonine-protein kinase 24 36 kDa subunit; | STK24 | 0.370776 | 0.114634 | Q9Y6E0-2;Q9Y6E0 | Q9Y6E0-2;Q9Y6E0 |
| Protein TFG | TFG | 0.280922 | 0.115533 | Q92734-4;Q92734-3;Q92734-2;Q92734 | Q92734-4;Q92734-3;Q92734-2;Q92734 |
| Beta-galactoside alpha-2,6-sialyltransferase 1 | ST6GAL1 | 0.0832671 | 0.115646 | P15907;P15907-2 | P15907;P15907-2 |
| E3 ubiquitin-protein ligase TRIM11 | TRIM11 | 0.0869933 | 0.115878 | Q96F44-3;Q96F44;Q96F44-2;Q6AZZ1-2;O00481-4;Q6AZZ | Q96F44-3;Q96F44;Q96F44-2 |
| cAMP-dependent protein kinase type II-alpha regulatory subunit | PRKAR2A | 0.134164 | 0.118212 | P13861-2;P13861 | P13861-2;P13861 |
| cAMP-dependent protein kinase catalytic subunit alpha | PRKACA | 0.144113 | 0.121074 | P17612-2;P17612;P22694-10;P22694-4;P22694-3;P22694 | P17612-2;P17612 |
| Heat shock protein 105 kDa | HSPH1 | 0.288919 | 0.121468 | Q92598-2 | Q92598-2 |
| Probable ATP-dependent RNA helicase DDX5 | DDX5 | 0.123894 | 0.1226 | P17844;P17844-2 | P17844;P17844-2 |
| Cation-independent mannose-6-phosphate receptor | IGF2R | 0.143446 | 0.123135 | P11717 | P11717 |
| Protein CutA | CUTA | 0.235846 | 0.12378 | O60888-3;O60888;O60888-2 | O60888-3;O60888;O60888-2 |
| NAD(P)H dehydrogenase [quinone] 1 | NQO1 | 0.115573 | 0.126308 | P15559-3;P15559-2;P15559 | P15559-3;P15559-2;P15559 |
| Tubulin-specific chaperone A | TBCA | 0.12965 | 0.127754 | O75347;O75347-2 | O75347;O75347-2 |
| Actin, alpha skeletal muscle;Actin, alpha cardiac muscle 1;Actin, gamma-enteric smootl | ACTA1;ACTC1;ACTG2;ACTA2 | 0.0553176 | 0.128433 | P68133;P68032;P63267;P62736;P63267-2 | P68133;P68032;P63267;P62736;P63267-2 |
| Inorganic pyrophosphatase 2, mitochondrial | PPA2 | 0.187517 | 0.131684 | Q9H2U2;Q9H2U2-2;Q9H2U2-6;Q9H2U2-3;Q9H2U2-4 | Q9H2U2;Q9H2U2-2;Q9H2U2-6;Q9H2U2-3 |
| Myoferlin | MYOF | 0.108631 | 0.133306 | Q9NZM1-6;Q9NZM1-3;Q9NZM1;Q9NZM1-5;Q9NZM1-2;( | Q9NZM1-6;Q9NZM1-3;Q9NZM1;Q9NZM1-5;Q9NZM |
| Endothelin-converting enzyme 1 | ECE1 | 0.233401 | 0.133386 | P42892-3;P42892-2;P42892-4;P42892 | P42892-3;P42892-2;P42892-4;P42892 |
| Annexin A7 | ANXA7 | 0.0992432 | 0.133838 | P20073-2;P20073 | P20073-2;P20073 |
| Alpha-mannosidase 2 | MAN2A1 | 0.233646 | 0.13416 | Q16706 | Q16706 |
| Sialidase-1 | NEU1 | 0.0948892 | 0.135345 | Q99519 | Q99519 |

| | | | | | |
|---|---|---|---|---|---|
| Lysosomal alpha-mannosidase;Lysosomal alpha-mannosidase A peptide;Lysosomal alpl | MAN2B1 | 0.137224 | 0.136643 | O00754-2;O00754 | O00754-2;O00754 |
| L-aminoadipate-semialdehyde dehydrogenase-phosphopantetheinyl transferase | AASDHPPT | 0.119876 | 0.137239 | Q9NRN7;Q9NRN7-2 | Q9NRN7 |
| Glycogen phosphorylase, brain form | PYGB | 0.331799 | 0.141006 | P11216 | P11216 |
| Quinone oxidoreductase | CRYZ | 0.137364 | 0.141148 | Q08257;Q08257-3;Q08257-2 | Q08257;Q08257-3;Q08257-2 |
| Suprabasin | SBSN | 0.106473 | 0.141841 | Q6UWP8;Q6UWP8-2 | Q6UWP8 |
| Ribose-phosphate pyrophosphokinase 2 | PRPS2 | 0.0983715 | 0.144843 | P11908;P11908-2 | P11908;P11908-2 |
| Acidic leucine-rich nuclear phosphoprotein 32 family member B | ANP32B | 0.0791274 | 0.145402 | Q92688-2;Q92688 | Q92688-2;Q92688 |
| Haloacid dehalogenase-like hydrolase domain-containing protein 2 | HDHD2 | 0.105627 | 0.145585 | Q9H0R4;Q9H0R4-2 | Q9H0R4;Q9H0R4-2 |
| Dihydropteridine reductase | QDPR | 0.654493 | 0.148022 | P09417;P09417-2 | P09417;P09417-2 |
| MARCKS-related protein | MARCKSL1 | 0.203681 | 0.14981 | P49006 | P49006 |
| Leucine zipper protein 1 | LUZP1 | 0.323789 | 0.150269 | Q86V48-2;Q86V48-3;Q86V48 | Q86V48-2;Q86V48-3;Q86V48 |
| Melanotransferrin | MFI2 | 0.2017 | 0.154842 | P08582;P08582-2 | P08582 |
| Ephrin type-B receptor 4 | EPHB4 | 0.24287 | 0.157076 | P54760;P54760-2;P54760-4;P54760-3 | P54760;P54760-2;P54760-4 |
| Cullin-4B | CUL4B | 0.27596 | 0.158169 | Q13620-1;Q13620;Q13620-3 | Q13620-1;Q13620;Q13620-3 |
| Cyclin-dependent kinase 1 | CDK1 | 0.780029 | 0.159341 | P06493;P06493-2 | P06493;P06493-2 |
| Polyadenylate-binding protein 1 | PABPC1 | 0.204943 | 0.159433 | P11940;P11940-2;Q9H361;Q96DU9-2;Q96DU9 | P11940;P11940-2 |
| Metalloproteinase inhibitor 1 | TIMP1 | 0.168568 | 0.159849 | P01033 | P01033 |
| Ectonucleoside triphosphate diphosphohydrolase 6 | ENTPD6 | 0.271732 | 0.163142 | O75354-2;O75354-3;O75354 | O75354-2;O75354-3;O75354 |
| Eukaryotic translation initiation factor 5A-1;Eukaryotic translation initiation factor 5A-1 | EIF5A;EIF5AL1 | 0.372415 | 0.164586 | P63241;P63241-2;Q6IS14 | P63241;P63241-2;Q6IS14 |
| Torsin-1B | TOR1B | 0.583645 | 0.165913 | O14657 | O14657 |
| Copine-1 | CPNE1 | 0.240287 | 0.166642 | Q99829 | Q99829 |
| Thioredoxin domain-containing protein 17 | TXNDC17 | 0.220635 | 0.167739 | Q9BRA2 | Q9BRA2 |
| Pyruvate kinase PKM | PKM | 0.0612076 | 0.169242 | P14618-2 | P14618-2 |
| Nucleolin | NCL | 0.203461 | 0.171552 | P19338 | P19338 |
| Cathepsin B;Cathepsin B light chain;Cathepsin B heavy chain | CTSB | 0.141034 | 0.172504 | P07858 | P07858 |
| Transmembrane protein 2 | TMEM2 | 0.0947586 | 0.17422 | Q9UHN6-2;Q9UHN6 | Q9UHN6-2;Q9UHN6 |
| Tropomyosin alpha-3 chain | TPM3 | 0.841816 | 0.174276 | P06753-2;P06753-5;P06753-7 | P06753-2;P06753-5 |
| Tuftelin-interacting protein 11 | TFIP11 | 0.060435 | 0.174634 | Q9UBB9;Q9UBB9-2 | Q9UBB9 |
| Destrin | DSTN | 0.187774 | 0.17628 | P60981;P60981-2 | P60981;P60981-2 |
| 78 kDa glucose-regulated protein | HSPA5 | 0.326038 | 0.178616 | P11021 | P11021 |
| Ig gamma-1 chain C region | IGHG1 | 0.209624 | 0.179293 | P01857 | P01857 |
| Leucine-rich alpha-2-glycoprotein | LRG1 | 0.127499 | 0.181745 | P02750 | P02750 |
| Integrin beta-1 | ITGB1 | 0.298653 | 0.182408 | P05556;P05556-2;P05556-5;P05556-4;P05556-3 | P05556;P05556-2;P05556-5;P05556-4;P05556-3 |
| Ethylmalonyl-CoA decarboxylase | ECHDC1 | 0.361434 | 0.185537 | Q9NTX5-6;Q9NTX5-2;Q9NTX5;Q9NTX5-3;Q9NTX5-5;Q9N | Q9NTX5-6;Q9NTX5-2;Q9NTX5;Q9NTX5-3 |
| Soluble calcium-activated nucleotidase 1 | CANT1 | 1.04549 | 0.186529 | Q8WVQ1-3;Q8WVQ1;Q8WVQ1-2 | Q8WVQ1-3;Q8WVQ1;Q8WVQ1-2 |
| Cathepsin Z | CTSZ | 0.235832 | 0.188894 | Q9UBR2 | Q9UBR2 |
| Vacuolar-sorting protein SNF8 | SNF8 | 0.910325 | 0.189185 | Q96H20-2;Q96H20 | Q96H20-2;Q96H20 |
| RuvB-like 2 | RUVBL2 | 0.483885 | 0.189917 | Q9Y230;Q9Y230-2 | Q9Y230;Q9Y230-2 |
| Plakophilin-1 | PKP1 | 0.159888 | 0.191995 | Q13835-2;Q13835 | Q13835-2;Q13835 |
| Heat shock protein 105 kDa | HSPH1 | 0.0331914 | 0.193596 | Q92598;Q92598-3;Q92598-4 | Q92598;Q92598-3;Q92598-4 |
| N-acetylglucosamine-6-sulfatase | GNS | 0.122961 | 0.197709 | P15586;P15586-2 | P15586;P15586-2 |
| Granulins;Acrogranin;Paragranulin;Granulin-1;Granulin-2;Granulin-3;Granulin-4;Granul | GRN | 0.243399 | 0.19839 | P28799;P28799-2;P28799-3 | P28799;P28799-2;P28799-3 |
| Voltage-dependent calcium channel subunit alpha-2/delta-1;Voltage-dependent calciu | CACNA2D1 | 0.0836576 | 0.200104 | P54289-5;P54289;P54289-4;P54289-2;P54289-3 | P54289-5;P54289;P54289-4;P54289-2;P54289-3 |
| Sorting nexin-3 | SNX3 | 0.269591 | 0.200207 | O60493;O60493-2;O60493-4;O60493-3 | O60493;O60493-2;O60493-4;O60493-3 |
| Interferon regulatory factor 6 | IRF6 | 0.285242 | 0.201723 | O14896-2;O14896 | O14896-2;O14896 |
| Glycine--tRNA ligase | GARS | 1.2428 | 0.202554 | P41250 | P41250 |
| RuvB-like 1 | RUVBL1 | 0.350406 | 0.203163 | Q9Y265;Q9Y265-2 | Q9Y265;Q9Y265-2 |
| Synaptobrevin homolog YKT6 | YKT6 | 0.206492 | 0.20365 | O15498;O15498-2 | O15498;O15498-2 |
| Tissue-type plasminogen activator;Tissue-type plasminogen activator chain A;Tissue-ty | PLAT | 0.242454 | 0.203901 | P00750;P00750-3;P00750-4;P00750-2 | P00750;P00750-3;P00750-4 |
| Protein PAXX | C9orf142 | 0.580078 | 0.207254 | Q9BUH6;Q9BUH6-2 | Q9BUH6 |
| DnaJ homolog subfamily A member 1 | DNAJA1 | 0.523077 | 0.208422 | P31689;P31689-2 | P31689;P31689-2 |
| Phosphoribosylformylglycinamidine synthase | PFAS | 0.318692 | 0.208724 | O15067 | O15067 |
| Liprin-alpha-1 | PPFIA1 | 0.373147 | 0.209432 | Q13136-2;Q13136;O75334-8;O75335;O75335-2;O75335- | Q13136-2;Q13136 |
| Actin, cytoplasmic 1;Actin, cytoplasmic 1, N-terminally processed | ACTB | 0.0245241 | 0.211264 | P60709 | P60709 |
| Acid ceramidase;Acid ceramidase subunit alpha;Acid ceramidase subunit beta | ASAH1 | 0.158109 | 0.212404 | Q13510;Q13510-2;Q13510-3 | Q13510;Q13510-2;Q13510-3 |
| THO complex subunit 4 | ALYREF | 0.203745 | 0.212582 | Q86V81 | Q86V81 |
| Phosphoenolpyruvate carboxykinase [GTP], mitochondrial | PCK2 | 0.331413 | 0.213646 | Q16822;Q16822-2;P35558;P35558-2 | Q16822 |
| 60 kDa heat shock protein, mitochondrial | HSPD1 | 0.314155 | 0.215056 | P10809;P10809-2 | P10809 |
| Ras-related protein Rab-6A | RAB6A | 0.551626 | 0.216735 | P20340-2;P20340-4;Q9H0N0 | P20340-2;P20340-4 |
| Peroxiredoxin-2 | PRDX2 | 0.292582 | 0.220803 | P32119;P32119-2 | P32119 |
| Golgi apparatus protein 1 | GLG1 | 0.334582 | 0.221144 | Q92896;Q92896-2;Q92896-3 | Q92896;Q92896-2;Q92896-3 |
| Prosaposin;Saposin-A;Saposin-B-Val;Saposin-B;Saposin-C;Saposin-D | PSAP | 0.286683 | 0.221645 | P07602;P07602-2;P07602-3 | P07602;P07602-2;P07602-3 |
| Proline synthase co-transcribed bacterial homolog protein | PROSC | 0.198565 | 0.224133 | O94903 | O94903 |
| Galactocerebrosidase | GALC | 0.137144 | 0.224326 | P54803;P54803-5;P54803-4;P54803-3 | P54803;P54803-5;P54803-4;P54803-3 |
| ADP-ribosylation factor 4 | ARF4 | 0.275172 | 0.227964 | P18085 | P18085 |
| Serine hydroxymethyltransferase, cytosolic | SHMT1 | 0.611901 | 0.23111 | P34896-2;P34896;P34896-3;P34896-4 | P34896-2;P34896;P34896-3;P34896-4 |
| Polypeptide N-acetylgalactosaminyltransferase 2;Polypeptide N-acetylgalactosaminyltr | GALNT2 | 0.553261 | 0.231823 | Q10471;Q10471-2 | Q10471 |
| Collagen alpha-1(XVIII) chain;Endostatin | COL18A1 | 0.129988 | 0.234053 | P39060-2;P39060-1;P39060 | P39060-2;P39060-1;P39060 |

| | | | | | | |
|---|---|---|---|---|---|---|
| Ephrin type-B receptor 3 | EPHB3 | | 0.420035 | 0.234845 | P54753 | P54753 |
| Low-density lipoprotein receptor | LDLR | | 0.230346 | 0.23654 | P01130-4;P01130-5;P01130;P01130-3;P01130-6;P01130- | P01130-4;P01130-5;P01130;P01130-3;P01130-6;P0 |
| UDP-glucuronic acid decarboxylase 1 | UXS1 | | 0.348982 | 0.239258 | Q8NBZ7;Q8NBZ7-2;Q8NBZ7-3 | Q8NBZ7;Q8NBZ7-2;Q8NBZ7-3 |
| Annexin A11 | ANXA11 | | 0.264599 | 0.242698 | P50995-2;P50995 | P50995-2;P50995 |
| Lysophospholipase-like protein 1 | LYPLAL1 | | 0.135913 | 0.24467 | Q5VWZ2-2;Q5VWZ2 | Q5VWZ2-2;Q5VWZ2 |
| Peripheral plasma membrane protein CASK | CASK | | 0.429647 | 0.246108 | O14936-3;O14936-4;O14936-6;O14936-2;O14936;O1493 | O14936-3;O14936-4;O14936-6;O14936-2;O14936;C |
| Beta-2-microglobulin;Beta-2-microglobulin form pI 5.3 | B2M | | 0.0597841 | 0.247098 | P61769 | P61769 |
| Dual specificity mitogen-activated protein kinase kinase 1 | MAP2K1 | | 0.395562 | 0.247432 | Q02750;Q02750-2 | Q02750;Q02750-2 |
| Membrane cofactor protein | CD46 | | 0.12847 | 0.250083 | P15529-7;P15529-4;P15529-6;P15529-3;P15529-5;P1552 | P15529-7;P15529-4;P15529-6;P15529-3;P15529-5;F |
| Ras-related protein Rab-1B | RAB1B | | 0.274427 | 0.250952 | Q9H0U4;Q92928;P59190-2;P59190 | Q9H0U4 |
| Myristoylated alanine-rich C-kinase substrate | MARCKS | | 0.297364 | 0.25212 | P29966 | P29966 |
| CD59 glycoprotein | CD59 | | 0.142798 | 0.253093 | P13987 | P13987 |
| Acetyl-CoA acetyltransferase, mitochondrial | ACAT1 | | 0.531134 | 0.253932 | P24752;P24752-2 | P24752 |
| Dystroglycan;Alpha-dystroglycan;Beta-dystroglycan | DAG1 | | 0.337609 | 0.254576 | Q14118 | Q14118 |
| Activated RNA polymerase II transcriptional coactivator p15 | SUB1 | | 0.367735 | 0.254799 | P53999 | P53999 |
| Receptor-type tyrosine-protein phosphatase S | PTPRS | | 0.505758 | 0.256873 | Q13332-6;Q13332-2;Q13332-3;Q13332-4;Q13332;Q1333 | Q13332-6;Q13332-2;Q13332-3;Q13332-4;Q13332;C |
| Collagen alpha-1(VI) chain | COL6A1 | | 1.09432 | 0.258362 | P12109 | P12109 |
| Procollagen-lysine,2-oxoglutarate 5-dioxygenase 2 | PLOD2 | | 0.840859 | 0.260856 | O00469-2;O00469;O00469-3 | O00469-2;O00469;O00469-3 |
| C-1-tetrahydrofolate synthase, cytoplasmic;Methylenetetrahydrofolate dehydrogenase | MTHFD1 | | 0.566749 | 0.268559 | P11586 | P11586 |
| Myotrophin | MTPN | | 0.106174 | 0.274252 | P58546 | P58546 |
| Transmembrane emp24 domain-containing protein 4 | TMED4 | | 0.203862 | 0.274844 | Q7Z7H5-3;Q7Z7H5-2;Q7Z7H5;Q9BVK6 | Q7Z7H5-3;Q7Z7H5-2;Q7Z7H5 |
| Annexin A1 | ANXA1 | | 0.304066 | 0.275207 | P04083 | P04083 |
| Growth arrest-specific protein 6 | GAS6 | | 0.365763 | 0.275347 | Q14393-2;Q14393-3;Q14393;Q14393-5;Q14393-4 | Q14393-2;Q14393-3;Q14393;Q14393-5;Q14393-4 |
| E3 ubiquitin-protein ligase Itchy homolog | ITCH | | 1.21281 | 0.275976 | Q96J02-2;Q96J02;Q96J02-3;Q9H0M0-6;Q9H0M0-3;Q9H0 | Q96J02-2;Q96J02;Q96J02-3 |
| Serine/threonine-protein kinase 26 | STK26 | | 0.230967 | 0.282371 | Q9P289;Q9P289-3;Q9P289-2;O00506-3;O00506-2 | Q9P289;Q9P289-3;Q9P289-2 |
| Protein NDRG3 | NDRG3 | | 0.206547 | 0.28303 | Q9UGV2-3;Q9UGV2-2;Q9UGV2 | Q9UGV2-3;Q9UGV2-2;Q9UGV2 |
| 60S acidic ribosomal protein P1 | RPLP1 | | 0.0831244 | 0.28352 | P05386 | P05386 |
| Pyridoxine-5-phosphate oxidase | PNPO | | 0.210618 | 0.284391 | Q9NVS9-3;Q9NVS9;Q9NVS9-2;Q9NVS9-4 | Q9NVS9-3;Q9NVS9;Q9NVS9-2;Q9NVS9-4 |
| Dynein light chain roadblock-type 1 | DYNLRB1 | | 0.408293 | 0.288337 | Q9NP97;Q8TF09;Q9NP97-2 | Q9NP97 |
| Disco-interacting protein 2 homolog B | DIP2B | | 0.528859 | 0.289 | Q9P265;Q9Y2E4 | Q9P265 |
| Bone morphogenetic protein 1 | BMP1 | | 0.448045 | 0.289352 | P13497;P13497-5;P13497-2;P13497-6;P13497-4;P13497- | P13497;P13497-5;P13497-2;P13497-6;P13497-4 |
| Collagen alpha-1(V) chain | COL5A1 | | 0.321471 | 0.292766 | P20908 | P20908 |
| Metalloproteinase inhibitor 2 | TIMP2 | | 0.328661 | 0.293979 | P16035 | P16035 |
| Dynactin subunit 2 | DCTN2 | | 1.05754 | 0.295323 | Q13561;Q13561-3;Q13561-2 | Q13561;Q13561-3;Q13561-2 |
| Tsukushin | TSKU | + | 1.37819 | 0.296133 | Q8WUA8 | Q8WUA8 |
| Mesothelin;Megakaryocyte-potentiating factor;Mesothelin, cleaved form | MSLN | | 0.796167 | 0.296608 | Q13421-4;Q13421-3;Q13421-2;Q13421 | Q13421-4;Q13421-3;Q13421-2;Q13421 |
| Semaphorin-3B | SEMA3B | | 0.472263 | 0.298947 | Q13214-2;Q13214 | Q13214-2;Q13214 |
| Rho-related GTP-binding protein RhoC | RHOC | | 0.631357 | 0.311713 | P08134 | P08134 |
| DnaJ homolog subfamily A member 2 | DNAJA2 | | 0.79865 | 0.313571 | O60884 | O60884 |
| Heterogeneous nuclear ribonucleoproteins C1/C2 | HNRNPC | + | 1.45276 | 0.315725 | P07910-2;P07910;P07910-4;B2RXH8 | P07910-2;P07910;P07910-4 |
| Peroxidasin homolog | PXDN | | 0.700918 | 0.315898 | Q92626;Q92626-2 | Q92626 |
| Calsyntenin-1;Soluble Alc-alpha;CTF1-alpha | CLSTN1 | | 0.585058 | 0.319099 | O94985-2;Q9H4D0 | O94985-2 |
| N-acetylgalactosaminyltransferase 7 | GALNT7 | | 0.39202 | 0.319513 | Q86SF2 | Q86SF2 |
| Tetraspanin-6 | TSPAN6 | | 0.0983039 | 0.319539 | O43657 | O43657 |
| Dihydrolipoyllysine-residue succinyltransferase component of 2-oxoglutarate dehydrog | DLST | | 1.00321 | 0.320367 | P36957;P36957-2 | P36957;P36957-2 |
| 14-3-3 protein gamma;14-3-3 protein gamma, N-terminally processed | YWHAG | | 0.329795 | 0.327328 | P61981 | P61981 |
| Probable serine carboxypeptidase CPVL | CPVL | | 0.212321 | 0.328783 | Q9H3G5 | Q9H3G5 |
| Cadherin-13 | CDH13 | | 0.396815 | 0.329259 | P55290;P55290-4;P55290-3;P55290-2;P55290-5 | P55290;P55290-4;P55290-3;P55290-2;P55290-5 |
| Vesicle-associated membrane protein 7 | VAMP7 | | 0.542406 | 0.330952 | P51809;P51809-3;P51809-2 | P51809;P51809-3;P51809-2 |
| Protein FAM3C | FAM3C | | 0.966302 | 0.331399 | Q92520 | Q92520 |
| UDP-glucose:glycoprotein glucosyltransferase 1 | UGGT1 | | 0.425392 | 0.33147 | Q9NYU2-2;Q9NYU2 | Q9NYU2-2;Q9NYU2 |
| Protein kinase C and casein kinase substrate in neurons protein 3 | PACSIN3 | | 0.993219 | 0.331851 | Q9UKS6 | Q9UKS6 |
| Tumor-associated calcium signal transducer 2 | TACSTD2 | | 0.436545 | 0.333757 | P09758 | P09758 |
| CD109 antigen | CD109 | | 1.25924 | 0.334249 | Q6YHK3;Q6YHK3-4;Q6YHK3-2;Q6YHK3-3 | Q6YHK3;Q6YHK3-4;Q6YHK3-2;Q6YHK3-3 |
| Glutathione peroxidase 1 | GPX1 | | 0.103698 | 0.336665 | P07203;P07203-2 | P07203;P07203-2 |
| Myotubularin | MTM1 | | 1.12703 | 0.338069 | Q13496;Q13496-2 | Q13496;Q13496-2 |
| Secretory carrier-associated membrane protein 3 | SCAMP3 | | 0.709096 | 0.340458 | O14828;O14828-2 | O14828;O14828-2 |
| CD44 antigen | CD44 | | 0.447278 | 0.343168 | P16070-8;P16070-4;P16070-3;P16070-7;P16070-5;P1607 | P16070-8;P16070-4;P16070-3;P16070-7;P16070-5;F |
| EH domain-containing protein 4 | EHD4 | | 0.501835 | 0.344904 | Q9H223 | Q9H223 |
| Ras-related protein Rab-5A | RAB5A | | 0.213542 | 0.345089 | P20339-2;P20339 | P20339-2;P20339 |
| Serine hydroxymethyltransferase, mitochondrial | SHMT2 | | 0.578163 | 0.345429 | P34897-3;P34897;P34897-2 | P34897-3;P34897;P34897-2 |
| Rho-related GTP-binding protein RhoG | RHOG | | 1.10746 | 0.34591 | P84095;Q9H4E5-2;P17081;Q9H4E5 | P84095 |
| Unconventional myosin-VI | MYO6 | | 0.569679 | 0.346335 | Q9UM54-5;Q9UM54-2;Q9UM54-6;Q9UM54-4;Q9UM54- | Q9UM54-5;Q9UM54-2;Q9UM54-6;Q9UM54-4;Q9U |
| Integrin beta-like protein 1 | ITGBL1 | | 0.364109 | 0.346441 | O95965;O95965-3;O95965-2 | O95965;O95965-3;O95965-2 |
| EH domain-containing protein 2 | EHD2 | | 0.317627 | 0.348938 | Q9NZN4;Q9NZN4-2 | Q9NZN4;Q9NZN4-2 |
| Iron/zinc purple acid phosphatase-like protein | PAPL | | 0.396034 | 0.349515 | Q6ZNF0 | Q6ZNF0 |
| Inorganic pyrophosphatase | PPA1 | | 0.364356 | 0.351744 | Q15181 | Q15181 |
| Gamma-glutamyl hydrolase | GGH | | 0.337404 | 0.35234 | Q92820 | Q92820 |

| | | | | | | |
|---|---|---|---|---|---|---|
| Cystatin-C | CST3 | | 0.916874 | 0.355019 | P01034 | P01034 |
| Protein PRRC2A | PRRC2A | | 0.711437 | 0.355405 | P48634;P48634-3;P48634-2;P48634-4 | P48634;P48634-3;P48634-2;P48634-4 |
| Mannose-1-phosphate guanyltransferase alpha | GMPPA | | 0.28405 | 0.357674 | Q96IJ6;Q96IJ6-2 | Q96IJ6;Q96IJ6-2 |
| NudC domain-containing protein 2 | NUDCD2 | | 0.55553 | 0.358499 | Q8WVJ2 | Q8WVJ2 |
| Ribonuclease T2 | RNASET2 | | 0.438479 | 0.358845 | O00584;O00584-2 | O00584 |
| CD9 antigen | CD9 | | 0.455568 | 0.358934 | P21926 | P21926 |
| Casein kinase II subunit beta | CSNK2B | | 0.316783 | 0.358984 | P67870 | P67870 |
| Endoglin | ENG | | 0.533487 | 0.359017 | P17813-2;P17813 | P17813-2;P17813 |
| Brain-specific angiogenesis inhibitor 1-associated protein 2-like protein 1 | BAIAP2L1 | + | 1.58756 | 0.360483 | Q9UHR4 | Q9UHR4 |
| Epididymis-specific alpha-mannosidase | MAN2B2 | | 0.246381 | 0.360483 | Q9Y2E5;Q9Y2E5-2 | Q9Y2E5;Q9Y2E5-2 |
| CD81 antigen | CD81 | | 0.542759 | 0.362174 | P60033 | P60033 |
| Peroxiredoxin-5, mitochondrial | PRDX5 | | 0.522352 | 0.362372 | P30044-2;P30044;P30044-3;P30044-4 | P30044-2;P30044;P30044-3;P30044-4 |
| Lysosomal protective protein;Lysosomal protective protein 32 kDa chain;Lysosomal pro | CTSA | | 0.361694 | 0.362809 | P10619-2;P10619 | P10619-2;P10619 |
| Junctional adhesion molecule A | F11R | | 0.760965 | 0.372002 | Q9Y624;Q9Y624-2 | Q9Y624;Q9Y624-2 |
| Galactosylgalactosylxylosylprotein 3-beta-glucuronosyltransferase 3 | B3GAT3 | | 0.8427 | 0.374172 | O94766-2;O94766 | O94766-2;O94766 |
| Heterogeneous nuclear ribonucleoprotein K | HNRNPK | + | 1.31517 | 0.376399 | P61978-3 | P61978-3 |
| Ras-related protein Rab-14 | RAB14 | | 0.40217 | 0.376515 | P61106 | P61106 |
| BTB/POZ domain-containing protein KCTD12 | KCTD12 | | 0.937723 | 0.377078 | Q96CX2;Q68DU8 | Q96CX2 |
| 2-deoxynucleoside 5-phosphate N-hydrolase 1 | DNPH1 | | 0.310036 | 0.377256 | O43598;O43598-2 | O43598;O43598-2 |
| Tubulin beta-8 chain | TUBB8 | | 0.186924 | 0.380165 | Q3ZCM7 | Q3ZCM7 |
| Glucosylceramidase | GBA | + | 1.45208 | 0.381786 | P04062-4;P04062-2;P04062;P04062-5 | P04062-4;P04062-2;P04062;P04062-5 |
| Lysosomal alpha-glucosidase;76 kDa lysosomal alpha-glucosidase;70 kDa lysosomal alp | GAA | | 0.577031 | 0.381814 | P10253 | P10253 |
| Protein FAM3A | FAM3A | | 0.20261 | 0.38195 | P98173-3;P98173;P98173-2 | P98173-3;P98173;P98173-2 |
| Serine--tRNA ligase, cytoplasmic | SARS | | 0.771046 | 0.387611 | P49591 | P49591 |
| Ubiquitin-40S ribosomal protein S27a;Ubiquitin;40S ribosomal protein S27a;Ubiquitin-6 | RPS27A;UBA52;UBB;UBC | | 0.808271 | 0.389105 | P62979;P62987;P0CG47;P0CG48 | P62979;P62987;P0CG47;P0CG48 |
| 72 kDa type IV collagenase;PEX | MMP2 | | 0.825819 | 0.390863 | P08253;P08253-3;P08253-2 | P08253;P08253-3;P08253-2 |
| Beta-glucuronidase | GUSB | | 0.361067 | 0.392321 | P08236-2;P08236;P08236-3 | P08236-2;P08236;P08236-3 |
| Hepatocyte growth factor receptor | MET | | 0.583431 | 0.393422 | P08581;P08581-2;P08581-3;P42685-2;P42685 | P08581;P08581-2;P08581-3 |
| Collagen alpha-1(VII) chain | COL7A1 | | 0.431224 | 0.396221 | Q02388-2;Q02388 | Q02388-2;Q02388 |
| NADH-cytochrome b5 reductase 3;NADH-cytochrome b5 reductase 3 membrane-bounc | CYB5R3 | | 0.173032 | 0.399291 | P00387-2;P00387;P00387-3 | P00387-2;P00387;P00387-3 |
| Septin-2 | SEPT2 | | 1.03959 | 0.401474 | Q15019;Q15019-3;Q15019-2 | Q15019;Q15019-3;Q15019-2 |
| Neuropilin-1 | NRP1 | | 0.799879 | 0.402635 | O14786;O14786-3;O14786-2 | O14786;O14786-3;O14786-2 |
| Cartilage intermediate layer protein 2;Cartilage intermediate layer protein 2 C1;Cartilag | CILP2 | + | 1.7758 | 0.404655 | Q8IUL8 | Q8IUL8 |
| Exostosin-2 | EXT2 | + | 1.84229 | 0.406155 | Q93063;Q93063-3;Q93063-2 | Q93063;Q93063-3;Q93063-2 |
| Early endosome antigen 1 | EEA1 | | 0.877358 | 0.406907 | Q15075 | Q15075 |
| Protein NOV homolog | NOV | | 0.36545 | 0.40712 | P48745 | P48745 |
| Prolyl 3-hydroxylase 1 | LEPRE1 | | 0.502012 | 0.409311 | Q32P28-4;Q32P28;Q32P28-3;Q32P28-2 | Q32P28-4;Q32P28;Q32P28-3 |
| ADP-ribosylation factor 5 | ARF5 | | 0.306945 | 0.410351 | P84085 | P84085 |
| Glutathione S-transferase kappa 1 | GSTK1 | | 0.125261 | 0.413534 | Q9Y2Q3;Q9Y2Q3-4;Q9Y2Q3-3;Q9Y2Q3-2 | Q9Y2Q3;Q9Y2Q3-4;Q9Y2Q3-3;Q9Y2Q3-2 |
| Carbohydrate sulfotransferase 14 | CHST14 | | 0.571248 | 0.416014 | Q8NCH0 | Q8NCH0 |
| ADP-ribosylation factor 6 | ARF6 | | 0.281714 | 0.41615 | P62330 | P62330 |
| Epithelial cell adhesion molecule | EPCAM | | 0.804741 | 0.418941 | P16422 | P16422 |
| Pyruvate dehydrogenase E1 component subunit beta, mitochondrial | PDHB | | 0.212827 | 0.419844 | P11177-2;P11177;P11177-3 | P11177-2;P11177;P11177-3 |
| Integrin alpha-V;Integrin alpha-V heavy chain;Integrin alpha-V light chain | ITGAV | | 0.725427 | 0.420615 | P06756-3;P06756-2;P06756 | P06756-3;P06756-2;P06756 |
| Beta-hexosaminidase subunit beta;Beta-hexosaminidase subunit beta chain B;Beta-hex | HEXB | | 0.352902 | 0.421417 | P07686 | P07686 |
| Slit homolog 2 protein;Slit homolog 2 protein N-product;Slit homolog 2 protein C-produ | SLIT2 | + | 1.51508 | 0.424559 | O94813-3;O94813-2;O94813 | O94813-3;O94813-2;O94813 |
| Neural Wiskott-Aldrich syndrome protein | WASL | | 1.01633 | 0.427678 | O00401 | O00401 |
| Fibulin-1 | FBLN1 | | 0.548169 | 0.428183 | P23142 | P23142 |
| Catalase | CAT | | 0.28549 | 0.429274 | P04040 | P04040 |
| Protein scribble homolog | SCRIB | | 0.78452 | 0.429354 | Q14160;Q14160-3 | Q14160;Q14160-3 |
| Alpha-galactosidase A | GLA | + | 1.71192 | 0.430349 | P06280 | P06280 |
| Succinyl-CoA ligase [ADP/GDP-forming] subunit alpha, mitochondrial | SUCLG1 | | 0.132397 | 0.431101 | P53597 | P53597 |
| Laminin subunit gamma-1 | LAMC1 | | 1.26558 | 0.433041 | P11047 | P11047 |
| Legumain | LGMN | | 0.747977 | 0.435304 | Q99538;Q99538-2;Q99538-3 | Q99538;Q99538-2;Q99538-3 |
| Anthrax toxin receptor 1 | ANTXR1 | | 0.443507 | 0.436981 | Q9H6X2-4;Q9H6X2-6;Q9H6X2-2;Q9H6X2-5;Q9H6X2;Q9H | Q9H6X2-4;Q9H6X2-6;Q9H6X2-2;Q9H6X2-5;Q9H6X2 |
| Adipocyte plasma membrane-associated protein | APMAP | | 0.131833 | 0.437038 | Q9HDC9;Q9HDC9-2 | Q9HDC9;Q9HDC9-2 |
| HLA class I histocompatibility antigen, Cw-12 alpha chain;HLA class I histocompatibility | HLA-C | | 0.615545 | 0.437312 | P30508;Q07000;P30510;P30505;P30504;Q9TNN7;P3050 | P30508;Q07000;P30510;P30505;P30504;Q9TNN7;P |
| Neuroserpin | SERPINI1 | | 0.780462 | 0.439078 | Q99574 | Q99574 |
| Dermcidin;Survival-promoting peptide;DCD-1 | DCD | | 0.294824 | 0.441775 | P81605;P81605-2 | P81605;P81605-2 |
| Eukaryotic translation initiation factor 3 subunit I | EIF3I | | 0.873494 | 0.443611 | Q13347 | Q13347 |
| Transmembrane 9 superfamily member 2 | TM9SF2 | | 0.784701 | 0.447373 | Q99805 | Q99805 |
| Brain acid soluble protein 1 | BASP1 | | 0.206239 | 0.448147 | P80723;P80723-2 | P80723;P80723-2 |
| Disintegrin and metalloproteinase domain-containing protein 10 | ADAM10 | | 0.603036 | 0.45492 | O14672;O14672-2 | O14672 |
| Latent-transforming growth factor beta-binding protein 4 | LTBP4 | | 0.362019 | 0.461634 | Q8N2S1-2;Q8N2S1-3;Q8N2S1;Q8N2S1-4 | Q8N2S1-2;Q8N2S1-3;Q8N2S1 |
| Claudin-4;Claudin-9;Claudin-6;Claudin-3 | CLDN4;CLDN9;CLDN6;CLDN3 | | 0.606699 | 0.463561 | O14493;O95484;P56747;O15551 | O14493;O95484;P56747;O15551 |
| Adenine phosphoribosyltransferase | APRT | | 0.318773 | 0.464629 | P07741;P07741-2 | P07741;P07741-2 |
| Cathepsin L1;Cathepsin L1 heavy chain;Cathepsin L1 light chain | CTSL | | 0.365163 | 0.466297 | P07711 | P07711 |
| Tubulin alpha-1B chain | TUBA1B | | 0.635745 | 0.468148 | P68363;P68363-2 | P68363;P68363-2 |

| | | | | | | |
|---|---|---|---|---|---|---|
| Amyloid beta A4 protein;N-APP;Soluble APP-alpha;Soluble APP-beta;C99;Beta-amyloid | APP | | 1.09988 | 0.471545 | P05067-7;P05067-11;P05067-8;P05067-9;P05067;P05067 | P05067-7;P05067-11;P05067-8;P05067-9;P05067;P( |
| Myosin-10 | MYH10 | | 0.419026 | 0.47719 | P35580;P35580-5;P35580-2;P35580-3;P35580-4 | P35580;P35580-5;P35580-2;P35580-3;P35580-4 |
| Endoplasmin | HSP90B1 | | 0.638977 | 0.479343 | P14625;Q58FF3 | P14625 |
| Coiled-coil and C2 domain-containing protein 1A | CC2D1A | | 0.0992478 | 0.481985 | Q6P1N0;Q6P1N0-2 | Q6P1N0;Q6P1N0-2 |
| Sushi repeat-containing protein SRPX2 | SRPX2 | | 1.10212 | 0.487501 | O60687 | O60687 |
| Ras-related protein Rab-2A | RAB2A | | 0.871714 | 0.490524 | P61019;P61019-2 | P61019;P61019-2 |
| 2,3-cyclic-nucleotide 3-phosphodiesterase | CNP | | 0.846187 | 0.495551 | P09543-2;P09543 | P09543-2;P09543 |
| Neurotrypsin | PRSS12 | | 0.302887 | 0.49979 | P56730 | P56730 |
| Ras-related C3 botulinum toxin substrate 1;Ras-related C3 botulinum toxin substrate 3 | RAC1;RAC3 | | 0.934027 | 0.500016 | P63000;P63000-2;P60763 | P63000;P63000-2;P60763 |
| Attractin | ATRN | | 0.731793 | 0.501323 | O75882-3;O75882-2;O75882 | O75882-3;O75882-2;O75882 |
| Calnexin | CANX | | 0.895756 | 0.504735 | P27824;P27824-2;P27824-3 | P27824;P27824-2;P27824-3 |
| Semaphorin-3C | SEMA3C | | 0.827151 | 0.50709 | Q99985;Q99985-2 | Q99985 |
| Palmitoyl-protein thioesterase 1 | PPT1 | | 0.539308 | 0.507821 | P50897;P50897-2 | P50897 |
| 14-3-3 protein epsilon | YWHAE | | 1.18017 | 0.516141 | P62258;P62258-2 | P62258;P62258-2 |
| Vacuolar protein sorting-associated protein 37B | VPS37B | + | 1.40601 | 0.519004 | Q9H9H4 | Q9H9H4 |
| Endophilin-A2 | SH3GL1 | | 0.705805 | 0.520447 | Q99961;Q99961-3;Q99961-2;Q99962 | Q99961;Q99961-3;Q99961-2 |
| Lipolysis-stimulated lipoprotein receptor | LSR | | 1.22638 | 0.522907 | Q86X29-6;Q86X29-5;Q86X29-4;Q86X29;Q86X29-2 | Q86X29-6;Q86X29-5;Q86X29-4;Q86X29;Q86X29-2 |
| Integrin alpha-3;Integrin alpha-3 heavy chain;Integrin alpha-3 light chain | ITGA3 | | 0.621662 | 0.524813 | P26006;P26006-1 | P26006;P26006-1 |
| Myosin light chain 6B | MYL6B | | 0.47992 | 0.528196 | P14649 | P14649 |
| Beta-hexosaminidase subunit alpha | HEXA | | 0.527432 | 0.53344 | P06865;P06865-2 | P06865 |
| Cytochrome b-c1 complex subunit 2, mitochondrial | UQCRC2 | | 0.296163 | 0.538822 | P22695 | P22695 |
| Prothymosin alpha;Prothymosin alpha, N-terminally processed;Thymosin alpha-1 | PTMA | | 0.226116 | 0.544098 | P06454-2;P06454 | P06454-2;P06454 |
| Signal recognition particle 14 kDa protein | SRP14 | | 0.63997 | 0.544463 | P37108 | P37108 |
| Ephrin-B1 | EFNB1 | | 0.542554 | 0.547613 | P98172 | P98172 |
| Ras-related protein Rap-1b;Ras-related protein Rap-1b-like protein | RAP1B | | 1.0519 | 0.548137 | P61224;P61224-3;A6NIZ1;P61224-2;P61224-4 | P61224;P61224-3;A6NIZ1;P61224-2;P61224-4 |
| Guanine nucleotide-binding protein G(I)/G(S)/G(T) subunit beta-1 | GNB1 | | 0.848564 | 0.548887 | P62873;P62873-2;P16520-2;P16520 | P62873;P62873-2 |
| Laminin subunit beta-1 | LAMB1 | + | 2.38991 | 0.54907 | P07942 | P07942 |
| Tripeptidyl-peptidase 1 | TPP1 | | 1.27221 | 0.549673 | O14773;O14773-2 | O14773;O14773-2 |
| FAD-linked sulfhydryl oxidase ALR | GFER | | 0.422079 | 0.550163 | P55789;P55789-2 | P55789;P55789-2 |
| Antithrombin-III | SERPINC1 | | 0.393899 | 0.553295 | P01008 | P01008 |
| Heat shock 70 kDa protein 13 | HSPA13 | + | 1.43923 | 0.557641 | P48723 | P48723 |
| 60S ribosomal protein L23 | RPL23 | | 0.633171 | 0.558228 | P62829 | P62829 |
| Chitinase domain-containing protein 1 | CHID1 | | 0.419555 | 0.558278 | Q9BWS9-3;Q9BWS9;Q9BWS9-2 | Q9BWS9-3;Q9BWS9;Q9BWS9-2 |
| Heat shock-related 70 kDa protein 2 | HSPA2 | | 0.221285 | 0.558393 | P54652 | P54652 |
| ESF1 homolog | ESF1 | + | 2.00317 | 0.565652 | Q9H501 | Q9H501 |
| Tissue alpha-L-fucosidase | FUCA1 | | 0.738319 | 0.567139 | P04066 | P04066 |
| Na(+)/H(+) exchange regulatory cofactor NHE-RF2 | SLC9A3R2 | | 0.320342 | 0.567506 | Q15599;Q15599-2;Q15599-3 | Q15599;Q15599-2;Q15599-3 |
| Tubulin beta-2A chain | TUBB2A | | 0.5085 | 0.572454 | Q13885 | Q13885 |
| Calumenin | CALU | | 0.240356 | 0.57436 | O43852-2;O43852-4;O43852-8;O43852-7 | O43852-2;O43852-4;O43852-8 |
| Charged multivesicular body protein 3 | CHMP3 | | 0.716689 | 0.57679 | Q9Y3E7;Q9Y3E7-3;Q9Y3E7-2;Q9Y3E7-4 | Q9Y3E7;Q9Y3E7-3;Q9Y3E7-2;Q9Y3E7-4 |
| Deoxyribonuclease-2-alpha | DNASE2 | | 0.370514 | 0.578481 | O00115-2;O00115 | O00115-2;O00115 |
| Beta-1,3-galactosyl-O-glycosyl-glycoprotein beta-1,6-N-acetylglucosaminyltransferase | GCNT1 | | 0.947034 | 0.57856 | Q02742 | Q02742 |
| Heat shock cognate 71 kDa protein | HSPA8 | | 0.917617 | 0.580604 | P11142;P11142-2 | P11142;P11142-2 |
| Sulfhydryl oxidase 2 | QSOX2 | | 0.482095 | 0.580877 | Q6ZRP7 | Q6ZRP7 |
| Cell division control protein 42 homolog | CDC42 | | 0.295206 | 0.583375 | P60953;P60953-1 | P60953;P60953-1 |
| Epidermal growth factor receptor | EGFR | | 1.29724 | 0.583742 | P00533;P00533-4;P00533-3;P00533-2;P04626-3;P04626- | P00533;P00533-4;P00533-3;P00533-2 |
| Elongation factor Tu, mitochondrial | TUFM | | 0.279925 | 0.584024 | P49411 | P49411 |
| Cytochrome b5 | CYB5A | | 0.199087 | 0.584469 | P00167-2;P00167;P00167-3 | P00167-2;P00167;P00167-3 |
| Extracellular sulfatase Sulf-2 | SULF2 | | 0.247641 | 0.585459 | Q8IWU5-2;Q8IWU5 | Q8IWU5-2;Q8IWU5 |
| CD2-associated protein | CD2AP | | 1.17654 | 0.585803 | Q9Y5K6 | Q9Y5K6 |
| Cathepsin D;Cathepsin D light chain;Cathepsin D heavy chain | CTSD | | 0.796926 | 0.588294 | P07339 | P07339 |
| Multiple inositol polyphosphate phosphatase 1 | MINPP1 | + | 2.52129 | 0.596851 | Q9UNW1;Q9UNW1-4;Q9UNW1-3;Q9UNW1-2 | Q9UNW1;Q9UNW1-4 |
| Activity-dependent neuroprotector homeobox protein | ADNP | | 0.476954 | 0.598028 | Q9H2P0 | Q9H2P0 |
| Coxsackievirus and adenovirus receptor | CXADR | | 0.289141 | 0.598256 | P78310-5;P78310-7;P78310-2;P78310-6;P78310;P78310- | P78310-5;P78310-7;P78310-2;P78310-6;P78310;P7 |
| Hyaluronan and proteoglycan link protein 3 | HAPLN3 | | 1.16507 | 0.598358 | Q96S86;Q86UW8 | Q96S86 |
| Pro-cathepsin H;Cathepsin H mini chain;Cathepsin H;Cathepsin H heavy chain;Cathepsi | CTSH | | 1.22607 | 0.600762 | P09668 | P09668 |
| Di-N-acetylchitobiase | CTBS | | 0.535549 | 0.604013 | Q01459 | Q01459 |
| Cochlin | COCH | | 1.27989 | 0.60657 | O43405;O43405-2 | O43405;O43405-2 |
| Out at first protein homolog | OAF | | 1.01899 | 0.607599 | Q86UD1 | Q86UD1 |
| Hypoxia up-regulated protein 1 | HYOU1 | | 0.680817 | 0.608551 | Q9Y4L1;Q9Y4L1-2;Q9BXT5 | Q9Y4L1;Q9Y4L1-2 |
| HLA class I histocompatibility antigen, A-68 alpha chain;HLA class I histocompatibility ai | HLA-A | + | 1.62099 | 0.608813 | P01891;P10316;P01892 | P01891;P10316;P01892 |
| CD97 antigen;CD97 antigen subunit alpha;CD97 antigen subunit beta | CD97 | | 0.74639 | 0.610181 | P48960;P48960-2;P48960-3;Q9UHX3-4;Q9UHX3-6;Q9UH | P48960;P48960-2;P48960-3 |
| Laminin subunit alpha-4 | LAMA4 | + | 1.90761 | 0.610181 | Q16363-2;Q16363;Q16363-3 | Q16363-2;Q16363 |
| Ceroid-lipofuscinosis neuronal protein 5 | CLN5 | + | 1.88384 | 0.61027 | O75503 | O75503 |
| 60S acidic ribosomal protein P0;60S acidic ribosomal protein P0-like | RPLP0;RPLP0P6 | | 1.20849 | 0.613361 | P05388;P05388-2;Q8NHW5 | P05388;P05388-2;Q8NHW5 |
| Fibroblast growth factor receptor 4 | FGFR4 | | 0.749116 | 0.617094 | P22455-2;P22455;P22455-3 | P22455-2;P22455;P22455-3 |
| Choline transporter-like protein 2 | SLC44A2 | | 0.849817 | 0.617787 | Q8IWA5;Q8IWA5-3;Q8IWA5-2 | Q8IWA5;Q8IWA5-3;Q8IWA5-2 |
| DnaJ homolog subfamily C member 7 | DNAJC7 | + | 1.354 | 0.618056 | Q99615;Q99615-2 | Q99615;Q99615-2 |

| | | | | | | |
|---|---|---|---|---|---|---|
| Kazal-type serine protease inhibitor domain-containing protein 1 | KAZALD1 | | 0.635947 | 0.620739 | Q96I82-2;Q96I82 | Q96I82-2;Q96I82 |
| Endonuclease domain-containing 1 protein | ENDOD1 | | 0.823544 | 0.62108 | O94919 | O94919 |
| Microfibrillar-associated protein 2 | MFAP2 | | 0.845617 | 0.621428 | P55001-3;P55001-2;P55001 | P55001-3;P55001-2;P55001 |
| Tubulin beta-4B chain | TUBB4B | | 1.05223 | 0.627434 | P68371 | P68371 |
| Vesicular integral-membrane protein VIP36 | LMAN2 | + | 1.4446 | 0.629328 | Q12907 | Q12907 |
| Sodium/potassium-transporting ATPase subunit beta-1 | ATP1B1 | + | 1.37298 | 0.630488 | P05026-2;P05026 | P05026-2;P05026 |
| Cystathionine gamma-lyase | CTH | | 1.28501 | 0.631925 | P32929;P32929-3;P32929-2 | P32929;P32929-3;P32929-2 |
| Sialate O-acetylesterase | SIAE | | 0.344567 | 0.637812 | Q9HAT2;Q9HAT2-2 | Q9HAT2;Q9HAT2-2 |
| Testican-2 | SPOCK2 | | 0.258003 | 0.638385 | Q92563;Q92563-2 | Q92563 |
| Guanine nucleotide-binding protein G(I)/G(S)/G(T) subunit beta-2 | GNB2 | + | 1.38621 | 0.642486 | P62879;P62879-2 | P62879 |
| Calcyphosin | CAPS | | 0.328807 | 0.642662 | Q13938;Q13938-3 | Q13938;Q13938-3 |
| Isocitrate dehydrogenase [NADP], mitochondrial | IDH2 | | 0.319209 | 0.644094 | P48735-2;P48735 | P48735-2;P48735 |
| Glyceraldehyde-3-phosphate dehydrogenase | GAPDH | | 0.849109 | 0.644327 | P04406;O14556 | P04406 |
| Epididymal secretory protein E1 | NPC2 | | 0.533911 | 0.644341 | P61916;P61916-2 | P61916;P61916-2 |
| BolA-like protein 2 | BOLA2 | | 0.814737 | 0.644624 | Q9H3K6;Q9H3K6-2 | Q9H3K6;Q9H3K6-2 |
| Basigin | BSG | + | 1.67315 | 0.648691 | P35613-2;P35613;P35613-4 | P35613-2;P35613;P35613-4 |
| Heterogeneous nuclear ribonucleoprotein D-like | HNRNPDL | | 1.08324 | 0.649928 | O14979-3;O14979-2;O14979 | O14979-3;O14979-2;O14979 |
| 14-3-3 protein sigma | SFN | | 1.05749 | 0.65054 | P31947;P31947-2 | P31947;P31947-2 |
| Immunoglobulin superfamily member 8 | IGSF8 | + | 2.33469 | 0.650947 | Q969P0;Q969P0-3;Q969P0-2 | Q969P0;Q969P0-3 |
| Serine/threonine-protein phosphatase CPPED1 | CPPED1 | | 0.375231 | 0.657131 | Q9BRF8;Q9BRF8-2 | Q9BRF8;Q9BRF8-2 |
| Gamma-interferon-inducible lysosomal thiol reductase | IFI30 | | 0.524965 | 0.662207 | P13284 | P13284 |
| Myosin regulatory light chain 12A;Myosin regulatory light chain 12B;Myosin regulatory | MYL12A;MYL12B;MYL9 | + | 1.78827 | 0.663832 | P19105;O14950;P24844;P24844-2 | P19105;O14950;P24844 |
| Semaphorin-4C | SEMA4C | | 0.585719 | 0.665507 | Q9C0C4 | Q9C0C4 |
| Tubulin beta-6 chain | TUBB6 | | 0.611603 | 0.669537 | Q9BUF5 | Q9BUF5 |
| GTP-binding nuclear protein Ran | RAN | | 0.973493 | 0.67064 | P62826 | P62826 |
| Endoplasmic reticulum resident protein 44 | ERP44 | | 0.592283 | 0.671442 | Q9BS26 | Q9BS26 |
| Histone H3.1 | HIST1H3A | | 0.450804 | 0.67148 | P68431 | P68431 |
| Lactoylglutathione lyase | GLO1 | | 0.365332 | 0.673131 | Q04760;Q04760-2 | Q04760;Q04760-2 |
| ATP synthase subunit beta, mitochondrial | ATP5B | | 0.524086 | 0.674501 | P06576 | P06576 |
| Collagen alpha-1(XVI) chain | COL16A1 | + | 1.36359 | 0.674849 | Q07092-2;Q07092 | Q07092-2;Q07092 |
| Ras-related protein Rab-5C | RAB5C | + | 2.07856 | 0.67551 | P51148;P51148-2 | P51148;P51148-2 |
| Rho GDP-dissociation inhibitor 1 | ARHGDIA | | 0.731179 | 0.681178 | P52565;P52565-2 | P52565;P52565-2 |
| Deoxyuridine 5-triphosphate nucleotidohydrolase, mitochondrial | DUT | + | 1.57401 | 0.682332 | P33316-2 | P33316-2 |
| Sequestosome-1 | SQSTM1 | + | 2.04993 | 0.685971 | Q13501;Q13501-2 | Q13501;Q13501-2 |
| Protein lin-7 homolog C | LIN7C | | 0.309153 | 0.691073 | Q9NUP9 | Q9NUP9 |
| Retinoid-inducible serine carboxypeptidase | SCPEP1 | + | 2.17842 | 0.691281 | Q9HB40;Q9HB40-2 | Q9HB40;Q9HB40-2 |
| Multifunctional methyltransferase subunit TRM112-like protein | TRMT112 | | 0.787961 | 0.691437 | Q9UI30;Q9UI30-2 | Q9UI30;Q9UI30-2 |
| Ribonucleases P/MRP protein subunit POP1 | POP1 | | 0.700776 | 0.691651 | Q99575 | Q99575 |
| Cathepsin F | CTSF | | 0.900908 | 0.69676 | Q9UBX1 | Q9UBX1 |
| Ras-related protein Rab-34 | RAB34 | | 0.578417 | 0.697611 | Q9BZG1-4;Q9BZG1;Q9BZG1-2 | Q9BZG1-4;Q9BZG1;Q9BZG1-2 |
| ADAMTS-like protein 4 | ADAMTSL4 | | 0.856599 | 0.698805 | Q6UY14;Q6UY14-3;Q6UY14-2 | Q6UY14;Q6UY14-3;Q6UY14-2 |
| Exostosin-like 2;Processed exostosin-like 2 | EXTL2 | | 0.847198 | 0.700127 | Q9UBQ6 | Q9UBQ6 |
| Aminopeptidase N | ANPEP | | 0.852788 | 0.70032 | P15144 | P15144 |
| Neutral alpha-glucosidase AB | GANAB | | 1.0771 | 0.704225 | Q14697;Q14697-2;Q14697-3 | Q14697;Q14697-2 |
| Small nuclear ribonucleoprotein Sm D2 | SNRPD2 | | 1.03221 | 0.706999 | P62316;P62316-2 | P62316;P62316-2 |
| Desmocollin-3 | DSC3 | | 0.814077 | 0.70871 | Q14574-2;Q14574 | Q14574-2;Q14574 |
| Lysosomal acid lipase/cholesteryl ester hydrolase | LIPA | | 0.803562 | 0.709424 | P38571-2;P38571 | P38571-2;P38571 |
| Amyloid-like protein 2 | APLP2 | + | 1.87184 | 0.70984 | Q06481;Q06481-3;Q06481-6;Q06481-2;Q06481-4;Q0648 | Q06481;Q06481-3;Q06481-6;Q06481-2;Q06481-4;C |
| Lysosomal Pro-X carboxypeptidase | PRCP | | 0.783989 | 0.711066 | P42785;P42785-2 | P42785;P42785-2 |
| Low molecular weight phosphotyrosine protein phosphatase | ACP1 | | 0.744408 | 0.712656 | P24666;P24666-3;P24666-4 | P24666;P24666-3;P24666-4 |
| Serine/arginine-rich splicing factor 7 | SRSF7 | | 1.24222 | 0.713938 | Q16629-3;Q16629-2;Q16629-4;Q16629 | Q16629-3;Q16629-2;Q16629-4;Q16629 |
| Alanine--tRNA ligase, cytoplasmic | AARS | | 1.01275 | 0.714821 | P49588;P49588-2 | P49588;P49588-2 |
| Cytoskeleton-associated protein 4 | CKAP4 | | 0.553218 | 0.718084 | Q07065 | Q07065 |
| Guanine nucleotide-binding protein G(q) subunit alpha | GNAQ | | 1.23422 | 0.719673 | P50148 | P50148 |
| Catechol O-methyltransferase | COMT | | 0.397386 | 0.720147 | P21964-2;P21964 | P21964-2;P21964 |
| RNA-binding motif protein, X chromosome;RNA-binding motif protein, X chromosome, | RBMX;RBMXL1 | | 0.999234 | 0.720343 | P38159;P38159-2;Q96E39;P38159-3;Q8N7X1 | P38159;P38159-2;Q96E39;P38159-3 |
| Platelet-activating factor acetylhydrolase IB subunit beta | PAFAH1B2 | + | 1.47046 | 0.721202 | P68402;P68402-3;P68402-2;P68402-4 | P68402;P68402-3;P68402-2;P68402-4 |
| 4F2 cell-surface antigen heavy chain | SLC3A2 | + | 2.09344 | 0.723815 | P08195-2 | P08195-2 |
| Nuclear receptor coactivator 5 | NCOA5 | + | 2.19274 | 0.726887 | Q9HCD5 | Q9HCD5 |
| Sushi, von Willebrand factor type A, EGF and pentraxin domain-containing protein 1 | SVEP1 | | 0.594049 | 0.729256 | Q4LDE5;Q4LDE5-4;Q4LDE5-2;Q4LDE5-3 | Q4LDE5;Q4LDE5-4 |
| Prohibitin-2 | PHB2 | | 0.716587 | 0.736169 | Q99623;Q99623-2 | Q99623;Q99623-2 |
| ADP-ribosylation factor-like protein 3 | ARL3 | | 0.242721 | 0.736561 | P36405 | P36405 |
| Nuclease-sensitive element-binding protein 1 | YBX1 | | 0.85365 | 0.744354 | P67809;Q9Y2T7 | P67809 |
| Unconventional myosin-Ic | MYO1C | | 1.02776 | 0.744487 | O00159;O00159-2;O00159-3 | O00159;O00159-2;O00159-3 |
| Lysosome-associated membrane glycoprotein 1 | LAMP1 | | 0.167059 | 0.745335 | P11279;P11279-2 | P11279;P11279-2 |
| Insulin-like growth factor-binding protein 4 | IGFBP4 | | 0.828114 | 0.745459 | P22692;P22692-2 | P22692;P22692-2 |
| Liprin-beta-1 | PPFIBP1 | | 0.720287 | 0.749741 | Q86W92-4;Q86W92-2;Q86W92;Q86W92-3;Q86W92-5;Q | Q86W92-4;Q86W92-2;Q86W92;Q86W92-3 |
| Band 4.1-like protein 2 | EPB41L2 | | 1.20663 | 0.752801 | O43491;O43491-3;O43491-2;Q9Y2J2-3;Q9Y2J2-2;Q9Y2J2- | O43491;O43491-3;O43491-2 |

| | | | | | | |
|---|---|---|---|---|---|---|
| Peptidyl-prolyl cis-trans isomerase FKBP5;Peptidyl-prolyl cis-trans isomerase FKBP5, N- | FKBP5 | | 1.11946 | 0.753358 | Q13451;Q13451-2 | Q13451 |
| Vigilin | HDLBP | | 1.14377 | 0.758053 | Q00341;Q00341-2 | Q00341;Q00341-2 |
| Ras-related protein Rab-21 | RAB21 | | 0.940349 | 0.760103 | Q9UL25 | Q9UL25 |
| Secreted and transmembrane protein 1 | SECTM1 | | 0.115682 | 0.760923 | Q8WVN6 | Q8WVN6 |
| Protein delta homolog 2 | DLK2 | | 0.404011 | 0.769952 | Q6UY11;Q6UY11-2 | Q6UY11;Q6UY11-2 |
| Golgi membrane protein 1 | GOLM1 | + | 2.43247 | 0.771112 | Q8NBJ4-2;Q8NBJ4 | Q8NBJ4-2;Q8NBJ4 |
| Calcineurin B homologous protein 1 | CHP1 | | 0.308089 | 0.771997 | Q99653 | Q99653 |
| Meteorin-like protein | METRNL | | 0.964663 | 0.772937 | Q641Q3;Q641Q3-2 | Q641Q3;Q641Q3-2 |
| Angiotensin-converting enzyme;Angiotensin-converting enzyme, soluble form | ACE | | 0.68964 | 0.773439 | P12821-2;P12821;P12821-4;P12821-3 | P12821-2;P12821;P12821-4;P12821-3 |
| 60S ribosomal protein L11 | RPL11 | | 0.766075 | 0.778795 | P62913;P62913-2 | P62913;P62913-2 |
| Neuronal growth regulator 1 | NEGR1 | | 0.301088 | 0.77973 | Q7Z3B1;Q7Z3B1-2 | Q7Z3B1;Q7Z3B1-2 |
| Extracellular matrix protein 1 | ECM1 | + | 1.94109 | 0.781787 | Q16610;Q16610-4;Q16610-2;Q16610-3 | Q16610;Q16610-4;Q16610-2 |
| Carboxypeptidase E | CPE | | 0.708253 | 0.783791 | P16870-2;P16870 | P16870-2;P16870 |
| Integrin beta-5 | ITGB5 | | 1.19589 | 0.784426 | P18084 | P18084 |
| Interleukin-11 | IL11 | | 0.634092 | 0.793004 | P20809;P20809-2 | P20809;P20809-2 |
| Neudesin | NENF | | 0.250038 | 0.79354 | Q9UMX5 | Q9UMX5 |
| Tax1-binding protein 3 | TAX1BP3 | | 0.648789 | 0.795256 | O14907 | O14907 |
| Ras-related protein Ral-B | RALB | | 0.296347 | 0.795618 | P11234;P11234-2;P11234-3 | P11234;P11234-2;P11234-3 |
| Ras-related protein Rab-3D | RAB3D | | 0.530996 | 0.795824 | O95716;P20337;Q96E17 | O95716 |
| Aflatoxin B1 aldehyde reductase member 2 | AKR7A2 | | 0.943399 | 0.799364 | O43488;Q8NHP1;O95154 | O43488 |
| 60S ribosomal protein L3 | RPL3 | | 0.68881 | 0.799809 | P39023;Q92901 | P39023 |
| Brain-specific angiogenesis inhibitor 1-associated protein 2 | BAIAP2 | + | 1.43808 | 0.802214 | Q9UQB8-3;Q9UQB8-5;Q9UQB8-4;Q9UQB8-6;Q9UQB8-2; | Q9UQB8-3;Q9UQB8-5;Q9UQB8-4;Q9UQB8-6;Q9UC |
| Cartilage-associated protein | CRTAP | + | 1.57389 | 0.805752 | O75718 | O75718 |
| Annexin A4 | ANXA4 | | 1.11221 | 0.809198 | P09525;P09525-2 | P09525;P09525-2 |
| Microtubule-associated protein 4 | MAP4 | + | 1.55066 | 0.809337 | P27816;P27816-6;P27816-2;P27816-4 | P27816;P27816-6;P27816-2 |
| Endoplasmic reticulum resident protein 29 | ERP29 | | 0.895903 | 0.810873 | P30040;P30040-2 | P30040 |
| DnaJ homolog subfamily C member 5 | DNAJC5 | + | 1.31494 | 0.81352 | Q9H3Z4-2;Q9H3Z4 | Q9H3Z4-2;Q9H3Z4 |
| Biotinidase | BTD | | 0.642883 | 0.816919 | P43251-4;P43251;P43251-3;P43251-2 | P43251-4;P43251;P43251-3;P43251-2 |
| Actin, cytoplasmic 2;Actin, cytoplasmic 2, N-terminally processed | ACTG1 | + | 1.7375 | 0.819396 | P63261 | P63261 |
| Monocarboxylate transporter 1 | SLC16A1 | | 1.20024 | 0.819632 | P53985;P53985-2 | P53985;P53985-2 |
| Scaffold attachment factor B2 | SAFB2 | | 0.378458 | 0.822214 | Q14151;Q14151-2 | Q14151 |
| Group XV phospholipase A2 | PLA2G15 | + | 2.62306 | 0.823538 | Q8NCC3;Q8NCC3-2 | Q8NCC3 |
| MAGUK p55 subfamily member 6 | MPP6 | | 0.772389 | 0.824258 | Q9NZW5 | Q9NZW5 |
| pre-mRNA 3 end processing protein WDR33 | WDR33 | | 0.702902 | 0.830235 | Q9C0J8 | Q9C0J8 |
| UPF0160 protein MYG1, mitochondrial | C12orf10 | | 0.70019 | 0.83797 | Q9HB07 | Q9HB07 |
| Carboxypeptidase A4 | CPA4 | | 0.933224 | 0.840123 | Q9UI42;Q9UI42-2 | Q9UI42;Q9UI42-2 |
| HD domain-containing protein 2 | HDDC2 | | 0.287997 | 0.840743 | Q7Z4H3-2;Q7Z4H3;Q7Z4H3-3 | Q7Z4H3-2;Q7Z4H3 |
| Protein FAM65A | FAM65A | | 0.656755 | 0.845614 | Q6ZS17-2;Q6ZS17;Q6ZS17-3;Q6ZS17-4 | Q6ZS17-2;Q6ZS17;Q6ZS17-3;Q6ZS17-4 |
| Basement membrane-specific heparan sulfate proteoglycan core protein;Endorepellin;l | HSPG2 | | 1.18775 | 0.848293 | P98160 | P98160 |
| PDZ and LIM domain protein 2 | PDLIM2 | | 0.288081 | 0.852352 | Q96JY6;Q96JY6-5;Q96JY6-3;Q96JY6-4;Q96JY6-2 | Q96JY6;Q96JY6-5;Q96JY6-3;Q96JY6-4 |
| Disks large homolog 3 | DLG3 | | 0.40917 | 0.853764 | Q92796-3;Q92796-2;Q92796 | Q92796-3;Q92796-2;Q92796 |
| Ras-related protein Rab-7a | RAB7A | + | 1.65048 | 0.858945 | P51149 | P51149 |
| Tubulin beta chain | TUBB | | 0.762693 | 0.860978 | P07437;A6NNZ2 | P07437 |
| Collagen alpha-2(IV) chain;Canstatin | COL4A2 | | 1.17943 | 0.86403 | P08572 | P08572 |
| Ubiquitin-associated domain-containing protein 1 | UBAC1 | | 0.725894 | 0.864772 | Q9BSL1 | Q9BSL1 |
| Scavenger receptor class B member 1 | SCARB1 | + | 2.66461 | 0.868472 | Q8WTV0-4;Q8WTV0-5;Q8WTV0-2;Q8WTV0 | Q8WTV0-4;Q8WTV0-5;Q8WTV0-2;Q8WTV0 |
| Vascular endothelial growth factor C | VEGFC | + | 1.52231 | 0.868708 | P49767 | P49767 |
| Guanine nucleotide-binding protein G(k) subunit alpha | GNAI3 | + | 1.69112 | 0.869493 | P08754 | P08754 |
| Guanine nucleotide-binding protein subunit alpha-11 | GNA11 | | 1.1704 | 0.870876 | P29992 | P29992 |
| Heterogeneous nuclear ribonucleoprotein M | HNRNPM | | 1.12642 | 0.873547 | P52272-2;P52272 | P52272-2;P52272 |
| Plasma membrane calcium-transporting ATPase 1 | ATP2B1 | | 1.07787 | 0.88448 | P20020-6;P20020-3;P20020-4;P20020;P20020-5;P20020- | P20020-6;P20020-3;P20020-4;P20020;P20020-5;P2( |
| TBC1 domain family member 15 | TBC1D15 | | 0.448965 | 0.887882 | Q8TC07-2;Q8TC07-3;Q8TC07 | Q8TC07-2;Q8TC07-3;Q8TC07 |
| Thrombospondin-1 | THBS1 | + | 1.98309 | 0.890755 | P07996;P07996-2;P35442 | P07996;P07996-2 |
| Histidine triad nucleotide-binding protein 1 | HINT1 | + | 1.32848 | 0.891631 | P49773 | P49773 |
| Catenin delta-1 | CTNND1 | + | 2.13144 | 0.893737 | O60716-21;O60716-19;O60716-18;O60716-17;O60716-1 | O60716-21;O60716-19;O60716-18;O60716-17;O607 |
| Copine-3 | CPNE3 | + | 2.88143 | 0.89846 | O75131;Q86YQ8-2;Q9HCH3-2;Q96FN4-2;Q8IYJ1;Q96A23 | O75131 |
| Thyroid hormone receptor-associated protein 3 | THRAP3 | + | 1.38071 | 0.900817 | Q9Y2W1 | Q9Y2W1 |
| Dickkopf-related protein 1 | DKK1 | | 0.479363 | 0.909244 | O94907;Q9UBU2 | O94907 |
| Ribonuclease 4 | RNASE4 | | 1.29489 | 0.910049 | P34096 | P34096 |
| Leucine-rich repeat-containing protein 57 | LRRC57 | + | 1.59147 | 0.910583 | Q8N9N7 | Q8N9N7 |
| Ras-related protein Rab-8A | RAB8A | | 0.817747 | 0.918429 | P61006;P61006-2 | P61006;P61006-2 |
| Histone H1.5 | HIST1H1B | + | 1.87547 | 0.918882 | P16401 | P16401 |
| Beta-1,4-glucuronyltransferase 1 | B4GAT1 | + | 1.60174 | 0.91902 | O43505 | O43505 |
| Deoxyribonuclease-1-like 1 | DNASE1L1 | | 1.2942 | 0.919041 | P49184 | P49184 |
| Semaphorin-3F | SEMA3F | + | 1.84994 | 0.919669 | Q13275 | Q13275 |
| Bone marrow stromal antigen 2 | BST2 | | 0.641252 | 0.922394 | Q10589-2;Q10589 | Q10589-2;Q10589 |
| Plasma alpha-L-fucosidase | FUCA2 | | 0.694579 | 0.93188 | Q9BTY2;Q9BTY2-2 | Q9BTY2 |
| 40S ribosomal protein S20 | RPS20 | | 0.862505 | 0.935731 | P60866;P60866-2 | P60866;P60866-2 |

| | | | | | | |
|---|---|---|---|---|---|---|
| Prostaglandin E synthase 3 | PTGES3 | | 0.755739 | 0.937551 | Q15185-4;Q15185;Q15185-3;Q15185-2 | Q15185-4;Q15185;Q15185-3 |
| Serine/arginine-rich splicing factor 1 | SRSF1 | | 0.688504 | 0.943972 | Q07955;Q07955-3;Q07955-2 | Q07955;Q07955-3;Q07955-2 |
| Procollagen-lysine,2-oxoglutarate 5-dioxygenase 1 | PLOD1 | + | 2.54415 | 0.944916 | Q02809;Q02809-2 | Q02809;Q02809-2 |
| Ras-related protein Rab-18 | RAB18 | + | 2.38595 | 0.949959 | Q9NP72;Q9NP72-2;Q9NP72-3 | Q9NP72;Q9NP72-2;Q9NP72-3 |
| ATP synthase subunit alpha, mitochondrial | ATP5A1 | | 0.813455 | 0.954983 | P25705;P25705-2;P25705-3 | P25705;P25705-2;P25705-3 |
| Ras-related protein Rab-35 | RAB35 | + | 1.40021 | 0.959343 | Q15286;Q15286-2 | Q15286;Q15286-2 |
| Interleukin-1 receptor accessory protein | IL1RAP | | 1.2477 | 0.965558 | Q9NPH3-2;Q9NPH3;Q9NPH3-5;Q9NPH3-3 | Q9NPH3-2;Q9NPH3;Q9NPH3-5;Q9NPH3-3 |
| ADP/ATP translocase 2;ADP/ATP translocase 2, N-terminally processed | SLC25A5 | | 0.552804 | 0.968266 | P05141;Q9H0C2 | P05141 |
| Non-specific lipid-transfer protein | SCP2 | | 1.29509 | 0.97025 | P22307-6;P22307-2;P22307-4;P22307-7;P22307-8;P2230 | P22307-6;P22307-2;P22307-4;P22307-7;P22307-8;P |
| Choline transporter-like protein 1 | SLC44A1 | | 1.27845 | 0.970283 | Q8WWI5-3;Q8WWI5-2;Q8WWI5 | Q8WWI5-3;Q8WWI5-2;Q8WWI5 |
| Proprotein convertase subtilisin/kexin type 6 | PCSK6 | + | 2.58267 | 0.975119 | P29122-2;P29122-8;P29122;P29122-7;P29122-5;P29122- | P29122-2;P29122-8;P29122;P29122-7;P29122-5;P2 |
| Collagen alpha-2(VI) chain | COL6A2 | | 0.898911 | 0.978001 | P12110;P12110-3;P12110-2 | P12110;P12110-3;P12110-2 |
| Guanine nucleotide-binding protein G(i) subunit alpha-2 | GNAI2 | + | 1.46674 | 0.986579 | P04899-4;P04899;P04899-6;P04899-3;P04899-5;P04899- | P04899-4;P04899;P04899-6;P04899-3;P04899-5;P0 |
| Lysyl oxidase homolog 2 | LOXL2 | + | 1.88099 | 0.987504 | Q9Y4K0 | Q9Y4K0 |
| D-3-phosphoglycerate dehydrogenase | PHGDH | + | 1.33551 | 0.990645 | O43175 | O43175 |
| Platelet-derived growth factor C;Platelet-derived growth factor C, latent form;Platelet-c | PDGFC | | 1.28017 | 0.991541 | Q9NRA1;Q9NRA1-2;Q9NRA1-4;Q9NRA1-3 | Q9NRA1;Q9NRA1-2;Q9NRA1-4 |
| Tissue factor | F3 | + | 1.65679 | 0.996784 | P13726;P13726-2 | P13726;P13726-2 |
| 10 kDa heat shock protein, mitochondrial | HSPE1 | | 0.929325 | 0.999439 | P61604 | P61604 |
| Laminin subunit alpha-5 | LAMA5 | + | 1.71437 | 1.00278 | O15230;O15230-2 | O15230 |
| ATP-dependent 6-phosphofructokinase, muscle type | PFKM | | 0.304256 | 1.01128 | P08237-2;P08237;P08237-3 | P08237-2;P08237;P08237-3 |
| Glypican-1;Secreted glypican-1 | GPC1 | + | 2.43342 | 1.0142 | P35052 | P35052 |
| HBS1-like protein | HBS1L | | 0.66306 | 1.01525 | Q9Y450;Q9Y450-4;Q9Y450-2 | Q9Y450;Q9Y450-4 |
| Serine/threonine-protein phosphatase 2A catalytic subunit beta isoform | PPP2CB | | 0.602364 | 1.01765 | P62714 | P62714 |
| Barrier-to-autointegration factor;Barrier-to-autointegration factor, N-terminally proces | BANF1 | | 0.566481 | 1.01865 | O75531 | O75531 |
| Drebrin-like protein | DBNL | + | 1.61361 | 1.02319 | Q9UJU6-2;Q9UJU6-3;Q9UJU6-6;Q9UJU6-4 | Q9UJU6-2;Q9UJU6-3;Q9UJU6-6;Q9UJU6-4 |
| Small nuclear ribonucleoprotein-associated proteins B and B;Small nuclear ribonucleop | SNRPB;SNRPN | | 0.818424 | 1.0323 | P14678-2;P63162;P14678;P63162-2;P14678-3 | P14678-2;P63162;P14678;P63162-2;P14678-3 |
| Acyl-coenzyme A thioesterase 13;Acyl-coenzyme A thioesterase 13, N-terminally proce | ACOT13 | | 0.332289 | 1.03523 | Q9NPJ3;Q9NPJ3-2 | Q9NPJ3;Q9NPJ3-2 |
| Agrin;Agrin N-terminal 110 kDa subunit;Agrin C-terminal 110 kDa subunit;Agrin C-term | AGRN | + | 1.52417 | 1.03614 | O00468-6;O00468-7;O00468-3;O00468-5;O00468-4;O004 | O00468-6;O00468-7;O00468-3;O00468-5;O00468-4 |
| Transforming protein RhoA | RHOA | | 0.493676 | 1.0388 | P61586 | P61586 |
| SPARC-related modular calcium-binding protein 1 | SMOC1 | + | 1.30838 | 1.0409 | Q9H4F8;Q9H4F8-2 | Q9H4F8;Q9H4F8-2 |
| Ras-related protein Rap-2b | RAP2B | + | 1.6698 | 1.04254 | P61225 | P61225 |
| Nicastrin | NCSTN | | 0.861725 | 1.04282 | Q92542-2;Q92542 | Q92542-2;Q92542 |
| Stanniocalcin-1 | STC1 | | 0.885145 | 1.04438 | P52823;P52823-2 | P52823;P52823-2 |
| Prostatic acid phosphatase;PAPf39 | ACPP | | 1.07193 | 1.04484 | P15309;P15309-2;P15309-3 | P15309;P15309-2;P15309-3 |
| 60S ribosomal protein L12 | RPL12 | + | 3.28491 | 1.04618 | P30050;P30050-2 | P30050;P30050-2 |
| 60S ribosomal protein L31 | RPL31 | + | 1.76981 | 1.04929 | P62899;P62899-3;P62899-2 | P62899;P62899-3;P62899-2 |
| GRIP1-associated protein 1 | GRIPAP1 | | 0.80075 | 1.05417 | Q4V328-4;Q4V328;Q4V328-3;Q4V328-2 | Q4V328-4;Q4V328;Q4V328-3;Q4V328-2 |
| Ras-related protein R-Ras2 | RRAS2 | | 1.00247 | 1.05475 | P62070;P62070-4;P62070-3;P62070-2 | P62070;P62070-4;P62070-3;P62070-2 |
| Collagen alpha-3(VI) chain | COL6A3 | | 0.336871 | 1.06322 | P12111-2;P12111;P12111-4;P12111-5;P12111-3 | P12111-2;P12111;P12111-4;P12111-5 |
| Procollagen-lysine,2-oxoglutarate 5-dioxygenase 3 | PLOD3 | + | 1.92803 | 1.06859 | O60568 | O60568 |
| Tyrosine-protein kinase Yes | YES1 | | 0.385946 | 1.07225 | P07947 | P07947 |
| SPARC | SPARC | | 1.11976 | 1.08848 | P09486 | P09486 |
| Signal recognition particle 9 kDa protein | SRP9 | + | 2.142 | 1.09123 | P49458;P49458-2 | P49458;P49458-2 |
| Kinesin-like protein KIF23 | KIF23 | | 1.26725 | 1.09677 | Q02241-2;Q02241-3 | Q02241-2;Q02241-3 |
| 3-mercaptopyruvate sulfurtransferase | MPST | | 0.323827 | 1.09845 | P25325;P25325-2 | P25325;P25325-2 |
| Beta-1,3-N-acetylglucosaminyltransferase lunatic fringe | LFNG | | 0.963925 | 1.09862 | Q8NES3;Q8NES3-3;Q8NES3-2;Q8NES3-4 | Q8NES3;Q8NES3-3;Q8NES3-2;Q8NES3-4 |
| Proprotein convertase subtilisin/kexin type 9 | PCSK9 | + | 1.6467 | 1.11211 | Q8NBP7;Q8NBP7-2 | Q8NBP7 |
| ATP-dependent RNA helicase DDX19A;ATP-dependent RNA helicase DDX19B | DDX19A;DDX19B | | 0.514039 | 1.12288 | Q9NUU7;Q9NUU7-2;Q9UMR2-2;Q9UMR2-4;Q9UMR2;Q9 | Q9NUU7;Q9NUU7-2;Q9UMR2-2;Q9UMR2-4;Q9UM |
| Cathepsin L2 | CTSV | | 1.18197 | 1.12702 | O60911 | O60911 |
| Alpha-internexin | INA | | 0.576654 | 1.12744 | Q16352 | Q16352 |
| Cysteine-rich motor neuron 1 protein;Processed cysteine-rich motor neuron 1 protein | CRIM1 | + | 2.26447 | 1.12774 | Q9NZV1 | Q9NZV1 |
| Voltage-dependent anion-selective channel protein 2 | VDAC2 | | 0.465648 | 1.15859 | P45880-2;P45880;P45880-1 | P45880-2;P45880;P45880-1 |
| Transforming growth factor beta-2;Latency-associated peptide | TGFB2 | | 1.17794 | 1.16277 | P61812 | P61812 |
| Peptidyl-glycine alpha-amidating monooxygenase;Peptidylglycine alpha-hydroxylating | PAM | + | 1.92388 | 1.16411 | P19021-4;P19021-3;P19021-6;P19021;P19021-5;P19021- | P19021-4;P19021-3;P19021-6;P19021;P19021-5;P1 |
| Large neutral amino acids transporter small subunit 1 | SLC7A5 | | 1.17031 | 1.17194 | Q01650;Q9UM01 | Q01650 |
| Synapse-associated protein 1 | SYAP1 | | 0.758768 | 1.17696 | Q96A49 | Q96A49 |
| Renin receptor | ATP6AP2 | | 0.925088 | 1.18426 | O75787-2;O75787 | O75787-2;O75787 |
| Acyl-coenzyme A thioesterase 11 | ACOT11 | | 0.355538 | 1.21388 | Q8WXI4-2;Q8WXI4 | Q8WXI4-2;Q8WXI4 |
| Cysteine-rich secretory protein LCCL domain-containing 2 | CRISPLD2 | | 0.686551 | 1.21491 | Q9H0B8-3;Q9H0B8-2;Q9H0B8;O43692 | Q9H0B8-3;Q9H0B8-2;Q9H0B8 |
| Fibrillin-1 | FBN1 | + | 1.5369 | 1.23091 | P35555 | P35555 |
| Toll-interacting protein | TOLLIP | + | 1.77411 | 1.23458 | Q9H0E2;Q9H0E2-2 | Q9H0E2 |
| 60S ribosomal protein L35a | RPL35A | + | 2.24644 | 1.23705 | P18077 | P18077 |
| Guanine nucleotide-binding protein G(s) subunit alpha isoforms short;Guanine nucleoti | GNAS | | 1.04058 | 1.24047 | P63092;P63092-4;Q5JWF2 | P63092;P63092-4;Q5JWF2 |
| 40S ribosomal protein S3 | RPS3 | + | 2.16155 | 1.24106 | P23396;P23396-2 | P23396;P23396-2 |
| Deleted in malignant brain tumors 1 protein | DMBT1 | + | 1.91866 | 1.25277 | Q9UGM3-4;Q9UGM3-8;Q9UGM3-2;Q9UGM3-5;Q9UGM3 | Q9UGM3-4;Q9UGM3-8;Q9UGM3-2;Q9UGM3-5;Q9U |
| Protein kinase C-binding protein NELL2 | NELL2 | + | 2.82912 | 1.27128 | Q99435-4;Q99435;Q99435-2;Q99435-3 | Q99435-4;Q99435;Q99435-2;Q99435-3 |
| Decorin | DCN | | 1.24031 | 1.27449 | P07585;P07585-4;P07585-2;P07585-3;P07585-5 | P07585;P07585-4 |
| 60S ribosomal protein L23a | RPL23A | | 0.897436 | 1.27474 | P62750 | P62750 |

| | | | | | | |
|---|---|---|---|---|---|---|
| Synaptosomal-associated protein 23 | SNAP23 | | 0.7218 | 1.27528 | O00161;O00161-2 | O00161;O00161-2 |
| Serine protease inhibitor Kazal-type 5;Hemofiltrate peptide HF6478;Hemofiltrate pepti | SPINK5 | + | 2.22661 | 1.27815 | Q9NQ38;Q9NQ38-3;Q9NQ38-2 | Q9NQ38;Q9NQ38-3;Q9NQ38-2 |
| Ras-related protein Ral-A | RALA | + | 1.65892 | 1.27846 | P11233 | P11233 |
| Tyrosine-protein kinase receptor UFO | AXL | | 1.20927 | 1.28418 | P30530-2;P30530 | P30530-2;P30530 |
| Procollagen C-endopeptidase enhancer 2 | PCOLCE2 | + | 1.43351 | 1.28687 | Q9UKZ9 | Q9UKZ9 |
| Adipogenesis regulatory factor | ADIRF | | 0.908983 | 1.2975 | Q15847 | Q15847 |
| Major prion protein | PRNP | | 0.980111 | 1.29871 | P04156-2;P04156 | P04156-2;P04156 |
| 40S ribosomal protein SA | RPSA | + | 3.75375 | 1.3002 | P08865 | P08865 |
| Fibulin-2 | FBLN2 | | 0.729279 | 1.31302 | P98095;P98095-2 | P98095;P98095-2 |
| 40S ribosomal protein S3a | RPS3A | + | 1.63295 | 1.32225 | P61247 | P61247 |
| 40S ribosomal protein S15a | RPS15A | + | 2.64391 | 1.32623 | P62244 | P62244 |
| Guanine nucleotide-binding protein G(I)/G(S)/G(O) subunit gamma-12 | GNG12 | | 1.19363 | 1.32647 | Q9UBI6 | Q9UBI6 |
| Histone H2A type 2-C;Histone H2A type 2-A | HIST2H2AC;HIST2H2AA3 | + | 1.82033 | 1.32989 | Q16777;Q6FI13 | Q16777;Q6FI13 |
| 40S ribosomal protein S4, X isoform;40S ribosomal protein S4, Y isoform 1;40S ribosom | RPS4X;RPS4Y1;RPS4Y2 | + | 2.43889 | 1.33 | P62701;P22090;Q8TD47 | P62701;P22090;Q8TD47 |
| Ephrin-A5 | EFNA5 | | 0.471408 | 1.33258 | P52803 | P52803 |
| Ras-related protein Rap-1A | RAP1A | + | 1.73401 | 1.33611 | P62834 | P62834 |
| Zinc-alpha-2-glycoprotein | AZGP1 | | 0.726334 | 1.3379 | P25311 | P25311 |
| 40S ribosomal protein S16 | RPS16 | + | 2.26924 | 1.34334 | P62249 | P62249 |
| G-protein coupled receptor 56;GPR56 N-terminal fragment;GPR56 C-terminal fragment | GPR56 | | 0.398057 | 1.3489 | Q9Y653-2;Q9Y653-3;Q9Y653;Q9Y653-4 | Q9Y653-2;Q9Y653-3;Q9Y653 |
| Matrilysin | MMP7 | | 0.590145 | 1.35008 | P09237 | P09237 |
| E3 ubiquitin-protein ligase CHIP | STUB1 | + | 2.68668 | 1.35446 | Q9UNE7;Q9UNE7-2 | Q9UNE7;Q9UNE7-2 |
| Collagen alpha-1(XII) chain | COL12A1 | + | 1.70504 | 1.35843 | Q99715;Q99715-4;Q99715-2 | Q99715;Q99715-4;Q99715-2 |
| Amphiregulin | AREG | | 0.929774 | 1.36872 | P15514 | P15514 |
| Prefoldin subunit 2 | PFDN2 | + | 1.78545 | 1.3744 | Q9UHV9 | Q9UHV9 |
| Versican core protein | VCAN | + | 1.34725 | 1.37536 | P13611-2;P13611;P13611-5;P13611-4;P13611-3 | P13611-2;P13611;P13611-5;P13611-4;P13611-3 |
| Mammalian ependymin-related protein 1 | EPDR1 | | 1.27087 | 1.37977 | Q9UM22;Q9UM22-3;Q9UM22-2 | Q9UM22;Q9UM22-3 |
| Protein S100-A10 | S100A10 | | 0.964217 | 1.38393 | P60903 | P60903 |
| Furin | FURIN | + | 1.82676 | 1.3852 | P09958 | P09958 |
| 60S ribosomal protein L22 | RPL22 | + | 2.22517 | 1.40683 | P35268 | P35268 |
| Vesicle-associated membrane protein 3 | VAMP3 | | 0.980342 | 1.40867 | Q15836;P63027;P23763-2;P23763-3;P23763 | Q15836 |
| Voltage-dependent anion-selective channel protein 1 | VDAC1 | | 0.534099 | 1.427 | P21796 | P21796 |
| Syndecan-1 | SDC1 | | 0.384585 | 1.43632 | P18827 | P18827 |
| Polyadenylate-binding protein 2 | PABPN1 | + | 1.67836 | 1.44588 | Q86U42-2;Q86U42;Q92843-2 | Q86U42-2;Q86U42;Q92843-2 |
| Lipoprotein lipase | LPL | + | 2.42215 | 1.46024 | P06858 | P06858 |
| Sodium/potassium-transporting ATPase subunit alpha-1 | ATP1A1 | + | 2.14461 | 1.46144 | P05023-4;P05023;P05023-3;P05023-2;P20648;P54707;P5 | P05023-4;P05023;P05023-3;P05023-2 |
| Endothelial lipase | LIPG | + | 1.97063 | 1.4739 | Q9Y5X9;Q9Y5X9-2 | Q9Y5X9;Q9Y5X9-2 |
| Secreted frizzled-related protein 1 | SFRP1 | | 0.894607 | 1.50574 | Q8N474 | Q8N474 |
| Calsyntenin-3 | CLSTN3 | + | 1.66024 | 1.5062 | Q9BQT9;Q9BQT9-2 | Q9BQT9;Q9BQT9-2 |
| 45 kDa calcium-binding protein | SDF4 | + | 2.86638 | 1.51188 | Q9BRK5;Q9BRK5-6;Q9BRK5-2;Q9BRK5-4;Q9BRK5-3;Q9BR | Q9BRK5;Q9BRK5-6;Q9BRK5-2 |
| 14-3-3 protein eta | YWHAH | + | 2.48526 | 1.52437 | Q04917 | Q04917 |
| POTE ankyrin domain family member E;POTE ankyrin domain family member F | POTEE;POTEF | | 0.148579 | 1.53328 | Q6S8J3;A5A3E0;Q6S8J3-3;Q6S8J3-2 | Q6S8J3;A5A3E0 |
| Sodium/potassium-transporting ATPase subunit beta-3 | ATP1B3 | + | 2.85688 | 1.54437 | P54709;P54709-2 | P54709;P54709-2 |
| 40S ribosomal protein S10;Putative 40S ribosomal protein S10-like | RPS10;RPS10P5 | + | 1.9227 | 1.55345 | P46783;Q9NQ39 | P46783;Q9NQ39 |
| V-type proton ATPase subunit S1 | ATP6AP1 | + | 2.0191 | 1.55588 | Q15904 | Q15904 |
| Ras-related protein Rab-10 | RAB10 | | 0.635688 | 1.56163 | P61026 | P61026 |
| BolA-like protein 3 | BOLA3 | | 1.26396 | 1.56354 | Q53S33 | Q53S33 |
| Histone H4 | HIST1H4A | + | 2.07886 | 1.56538 | P62805 | P62805 |
| 40S ribosomal protein S2 | RPS2 | + | 2.66518 | 1.5662 | P15880 | P15880 |
| Calumenin | CALU | | 1.06175 | 1.58469 | O43852;O43852-3;O43852-6;O43852-10;O43852-5;O438 | O43852;O43852-3;O43852-6;O43852-10;O43852-5; |
| 14-3-3 protein zeta/delta | YWHAZ | + | 2.80965 | 1.58742 | P63104;P63104-2 | P63104;P63104-2 |
| Kallikrein-6 | KLK6 | + | 1.79803 | 1.59086 | Q92876;Q92876-2 | Q92876;Q92876-2 |
| Complement factor I;Complement factor I heavy chain;Complement factor I light chain | CFI | | 1.09124 | 1.59267 | P05156 | P05156 |
| Double-stranded RNA-binding protein Staufen homolog 1 | STAU1 | + | 2.69968 | 1.59887 | O95793-2;O95793-3;O95793 | O95793-2;O95793-3;O95793 |
| Epidermal growth factor receptor kinase substrate 8 | EPS8 | + | 1.35707 | 1.60087 | Q12929;Q12929-2 | Q12929;Q12929-2 |
| Nucleobindin-1 | NUCB1 | + | 1.53249 | 1.60514 | Q02818 | Q02818 |
| Sodium-coupled neutral amino acid transporter 1 | SLC38A1 | | 0.706206 | 1.60724 | Q9H2H9 | Q9H2H9 |
| Neutral amino acid transporter B(0) | SLC1A5 | | 1.26662 | 1.60915 | Q15758;Q15758-3;Q15758-2 | Q15758;Q15758-3;Q15758-2 |
| Inhibin alpha chain | INHA | + | 2.16314 | 1.61193 | P05111 | P05111 |
| Solute carrier family 2, facilitated glucose transporter member 1 | SLC2A1 | + | 2.0629 | 1.61242 | P11166 | P11166 |
| 40S ribosomal protein S17-like;40S ribosomal protein S17 | RPS17L;RPS17 | + | 1.90915 | 1.61981 | P0CW22;P08708 | P0CW22;P08708 |
| Histone H2B type 1-L;Histone H2B type 1-M;Histone H2B type 1-N;Histone H2B type 1-I | HIST1H2BL;HIST1H2BM;HIST1 | | 1.18624 | 1.66001 | Q99880;Q99879;Q99877;Q93079;Q5QNW6;P62807;P58 | Q99880;Q99879;Q99877;Q93079;Q5QNW6;P62807 |
| Platelet-derived growth factor D;Platelet-derived growth factor D, latent form;Platelet- | PDGFD | | 0.832234 | 1.66035 | Q9GZP0;Q9GZP0-2 | Q9GZP0;Q9GZP0-2 |
| Lymphocyte function-associated antigen 3 | CD58 | | 0.799906 | 1.68591 | P19256-2;P19256-3;P19256 | P19256-2;P19256-3;P19256 |
| Lysozyme C | LYZ | | 0.870456 | 1.71903 | P61626 | P61626 |
| 40S ribosomal protein S18 | RPS18 | + | 3.18124 | 1.73926 | P62269 | P62269 |
| Apolipoprotein C-III | APOC3 | | 0.891264 | 1.76406 | P02656 | P02656 |
| Prominin-2 | PROM2 | | 1.16667 | 1.76466 | Q8N271;Q8N271-2;Q8N271-3 | Q8N271;Q8N271-2 |
| Medium-chain specific acyl-CoA dehydrogenase, mitochondrial | ACADM | | 0.557541 | 1.76504 | P11310;P11310-2 | P11310;P11310-2 |

| | | | | | | |
|---|---|---|---|---|---|---|
| 60S ribosomal protein L28 | RPL28 | + | 2.0612 | 1.7812 | P46779-4;P46779-5;P46779;P46779-2;P46779-3 | P46779-4;P46779-5;P46779;P46779-2;P46779-3 |
| Specifically androgen-regulated gene protein | SARG | | 0.640229 | 1.78881 | Q9BW04;Q9BW04-2 | Q9BW04;Q9BW04-2 |
| Alpha-N-acetylglucosaminidase;Alpha-N-acetylglucosaminidase 82 kDa form;Alpha-N-a | NAGLU | | 0.861466 | 1.80469 | P54802 | P54802 |
| N-acetylglucosamine-1-phosphotransferase subunit gamma | GNPTG | | 0.701873 | 1.80887 | Q9UJJ9 | Q9UJJ9 |
| M-phase phosphoprotein 8 | MPHOSPH8 | | 0.945386 | 1.81624 | Q99549;Q99549-2 | Q99549;Q99549-2 |
| N(4)-(beta-N-acetylglucosaminyl)-L-asparaginase;Glycosylasparaginase alpha chain;Gly | AGA | | 0.268153 | 1.81756 | P20933 | P20933 |
| 40S ribosomal protein S25 | RPS25 | + | 1.69954 | 1.84373 | P62851 | P62851 |
| Cytochrome c oxidase subunit 4 isoform 1, mitochondrial | COX4I1 | | 0.941443 | 1.84641 | P13073 | P13073 |
| GTP-binding protein SAR1a | SAR1A | | 1.1409 | 1.8533 | Q9NR31;Q9NR31-2 | Q9NR31;Q9NR31-2 |
| Disintegrin and metalloproteinase domain-containing protein 9 | ADAM9 | + | 2.06647 | 1.85868 | Q13443-2;Q13443 | Q13443-2;Q13443 |
| Collagen alpha-1(IV) chain;Arresten | COL4A1 | + | 1.5398 | 1.86003 | P02462-2;P02462;P29400;P29400-2 | P02462-2;P02462 |
| 60S ribosomal protein L30 | RPL30 | + | 2.48583 | 1.8627 | P62888 | P62888 |
| Lactadherin;Lactadherin short form;Medin | MFGE8 | + | 2.35334 | 1.87777 | Q08431;Q08431-3 | Q08431;Q08431-3 |
| 40S ribosomal protein S14 | RPS14 | + | 3.24777 | 1.8806 | P62263 | P62263 |
| Heat shock protein beta-1 | HSPB1 | + | 2.66395 | 1.90115 | P04792 | P04792 |
| 40S ribosomal protein S11 | RPS11 | + | 3.02094 | 1.91575 | P62280 | P62280 |
| Integral membrane protein 2B;BRI2, membrane form;BRI2 intracellular domain;BRI2C, | ITM2B | | 0.683189 | 1.93668 | Q9Y287;Q9Y287-2 | Q9Y287 |
| Histone H2B type 2-E;Histone H2B type 1-B;Histone H2B type 1-O;Histone H2B type 1-J; | HIST2H2BE;HIST1H2BB;HIST1I | + | 1.30878 | 1.95634 | Q16778;P33778;P23527;P06899;Q8N257 | Q16778;P33778;P23527;P06899;Q8N257 |
| 60S ribosomal protein L7a | RPL7A | + | 2.57988 | 1.9598 | P62424 | P62424 |
| Membrane-associated progesterone receptor component 1 | PGRMC1 | + | 1.99592 | 1.96099 | O00264;O00264-2 | O00264;O00264-2 |
| Serine protease HTRA1 | HTRA1 | + | 2.77852 | 1.96641 | Q92743 | Q92743 |
| Clusterin;Clusterin beta chain;Clusterin alpha chain | CLU | + | 4.0419 | 1.97297 | P10909-4;P10909;P10909-5;P10909-2;P10909-3 | P10909-4;P10909;P10909-5;P10909-2;P10909-3 |
| 60S ribosomal protein L14 | RPL14 | + | 2.10569 | 1.97923 | P50914 | P50914 |
| 40S ribosomal protein S7 | RPS7 | + | 1.97254 | 1.98832 | P62081 | P62081 |
| 40S ribosomal protein S5;40S ribosomal protein S5, N-terminally processed | RPS5 | | 1.06508 | 1.99532 | P46782 | P46782 |
| 40S ribosomal protein S13 | RPS13 | + | 1.5472 | 1.99759 | P62277 | P62277 |
| 60S ribosomal protein L7 | RPL7 | + | 3.09338 | 2.01086 | P18124 | P18124 |
| Flotillin-1 | FLOT1 | | 0.94866 | 2.02284 | O75955;O75955-2 | O75955;O75955-2 |
| 40S ribosomal protein S26;Putative 40S ribosomal protein S26-like 1 | RPS26;RPS26P11 | + | 1.44697 | 2.02425 | P62854;Q5JNZ5 | P62854;Q5JNZ5 |
| Sphingomyelin phosphodiesterase | SMPD1 | + | 1.31143 | 2.08474 | P17405-3;P17405-2;P17405-4;P17405 | P17405-3;P17405-2;P17405-4;P17405 |
| Calcium-regulated heat stable protein 1 | CARHSP1 | | 0.614858 | 2.11051 | Q9Y2V2 | Q9Y2V2 |
| 60S ribosomal protein L15 | RPL15 | + | 3.02959 | 2.12705 | P61313;P61313-2 | P61313 |
| Disintegrin and metalloproteinase domain-containing protein 15 | ADAM15 | + | 3.70181 | 2.16345 | Q13444-10;Q13444-9;Q13444-8;Q13444-2;Q13444-7;Q1 | Q13444-10;Q13444-9;Q13444-8;Q13444-2;Q13444- |
| Charged multivesicular body protein 2b | CHMP2B | | 1.24113 | 2.16346 | Q9UQN3-2;Q9UQN3 | Q9UQN3-2;Q9UQN3 |
| Ladinin-1 | LAD1 | + | 2.51549 | 2.16377 | O00515 | O00515 |
| GTPase NRas | NRAS | + | 1.87561 | 2.16594 | P01111 | P01111 |
| Prohibitin | PHB | | 0.757786 | 2.19979 | P35232;P35232-2 | P35232 |
| Protein LAP2 | ERBB2IP | + | 3.53812 | 2.23781 | Q96RT1-6;Q96RT1-4;Q96RT1-5;Q96RT1-3;Q96RT1-9;Q96 | Q96RT1-6;Q96RT1-4;Q96RT1-5;Q96RT1-3;Q96RT1- |
| 60S ribosomal protein L5 | RPL5 | + | 2.78972 | 2.26522 | P46777 | P46777 |
| 14-3-3 protein theta | YWHAQ | + | 2.89859 | 2.2897 | P27348 | P27348 |
| 60S ribosomal protein L27 | RPL27 | + | 2.44514 | 2.3138 | P61353 | P61353 |
| 40S ribosomal protein S8 | RPS8 | + | 2.33364 | 2.31473 | P62241 | P62241 |
| 60S ribosomal protein L13a;Putative 60S ribosomal protein L13a protein RPL13AP3 | RPL13A;RPL13AP3 | + | 1.63323 | 2.36535 | P40429;Q6NVV1 | P40429;Q6NVV1 |
| Tyrosyl-DNA phosphodiesterase 2 | TDP2 | | 0.928412 | 2.38062 | O95551;O95551-2;O95551-3 | O95551;O95551-2;O95551-3 |
| Tropomyosin alpha-3 chain | TPM3 | | 0.352874 | 2.38795 | P06753 | P06753 |
| Complement factor D | CFD | + | 2.37179 | 2.40422 | P00746 | P00746 |
| Carboxypeptidase M | CPM | | 1.25506 | 2.41133 | P14384 | P14384 |
| 40S ribosomal protein S9 | RPS9 | + | 2.10532 | 2.43929 | P46781 | P46781 |
| ATP synthase subunit d, mitochondrial | ATP5H | | 1.28309 | 2.46128 | O75947-2;O75947 | O75947-2;O75947 |
| 60S ribosomal protein L26-like 1;60S ribosomal protein L26 | RPL26L1;RPL26 | + | 1.47177 | 2.47356 | Q9UNX3;P61254 | Q9UNX3;P61254 |
| Voltage-dependent anion-selective channel protein 3 | VDAC3 | + | 1.40752 | 2.49384 | Q9Y277;Q9Y277-2 | Q9Y277;Q9Y277-2 |
| Angiogenin | ANG | + | 1.57868 | 2.49403 | P03950 | P03950 |
| EGF-containing fibulin-like extracellular matrix protein 1 | EFEMP1 | + | 3.00072 | 2.50884 | Q12805-2;Q12805-4;Q12805-3;Q12805 | Q12805-2;Q12805-4;Q12805-3;Q12805 |
| Src substrate cortactin | CTTN | + | 2.86822 | 2.53832 | Q14247-3;Q14247;Q14247-2 | Q14247-3;Q14247;Q14247-2 |
| Nidogen-1 | NID1 | | 0.978879 | 2.54128 | P14543;P14543-2 | P14543;P14543-2 |
| Tolloid-like protein 1 | TLL1 | | 1.14773 | 2.56701 | O43897;O43897-2 | O43897 |
| Ras-related protein Rab-13 | RAB13 | | 1.04771 | 2.5694 | P51153 | P51153 |
| Serine/arginine-rich splicing factor 2 | SRSF2 | + | 2.37415 | 2.59658 | Q01130-2;Q01130;Q9BRL6-2;Q9BRL6 | Q01130-2;Q01130 |
| Deleted in autism protein 1 | C3orf58 | | 1.21598 | 2.63525 | Q8NDZ4;Q8NDZ4-2 | Q8NDZ4;Q8NDZ4-2 |
| Bone morphogenetic protein 2 | BMP2 | + | 1.64703 | 2.64754 | P12643 | P12643 |
| Neuroblast differentiation-associated protein AHNAK | AHNAK | + | 2.41265 | 2.68459 | Q09666;Q09666-2 | Q09666 |
| 60S ribosomal protein L4 | RPL4 | + | 3.55083 | 2.71661 | P36578 | P36578 |
| 40S ribosomal protein S19 | RPS19 | + | 2.7397 | 2.86223 | P39019 | P39019 |
| Insulin-like growth factor-binding protein 5 | IGFBP5 | | 1.04468 | 2.88658 | P24593 | P24593 |
| 60S ribosomal protein L10a | RPL10A | + | 3.10717 | 2.90163 | P62906 | P62906 |
| EGF-containing fibulin-like extracellular matrix protein 1 | EFEMP1 | + | 1.86539 | 2.90952 | Q12805-5 | Q12805-5 |
| Secretogranin-2;Secretoneurin;Manserin | SCG2 | | 0.914753 | 2.945 | P13521 | P13521 |
| Secretogranin-1;PE-11;GAWK peptide;CCB peptide | CHGB | + | 2.43199 | 3.01518 | P05060 | P05060 |

| | | | | | | |
|---|---|---|---|---|---|---|
| Plasminogen activator inhibitor 1 RNA-binding protein | SERBP1 | + | 2.79977 | 3.0193 | Q8NC51-3;Q8NC51-4 | Q8NC51-3;Q8NC51-4 |
| Charged multivesicular body protein 5 | CHMP5 | + | 2.42011 | 3.0225 | Q9NZZ3;Q9NZZ3-2 | Q9NZZ3;Q9NZZ3-2 |
| Rac GTPase-activating protein 1 | RACGAP1 | + | 1.56563 | 3.07854 | Q9H0H5 | Q9H0H5 |
| Spondin-2 | SPON2 | + | 1.3712 | 3.11207 | Q9BUD6 | Q9BUD6 |
| Serine protease 23 | PRSS23 | | 1.29926 | 3.15292 | O95084;O95084-2 | O95084;O95084-2 |
| Angiopoietin-1 | ANGPT1 | + | 2.37364 | 3.26633 | Q15389-2;Q15389 | Q15389-2;Q15389 |
| Melanoma-derived growth regulatory protein | MIA | + | 3.39737 | 3.34167 | Q16674 | Q16674 |
| 14-3-3 protein beta/alpha;14-3-3 protein beta/alpha, N-terminally processed | YWHAB | + | 2.78593 | 3.3555 | P31946-2;P31946 | P31946-2;P31946 |
| Biglycan | BGN | + | 4.76192 | 3.53261 | P21810 | P21810 |
| Protein ELYS | AHCTF1 | + | 2.95721 | 3.66532 | Q8WYP5-3;Q8WYP5;Q8WYP5-2 | Q8WYP5-3;Q8WYP5;Q8WYP5-2 |
| C1q-related factor | C1QL1 | + | 3.22488 | 3.72069 | O75973;Q5VWW1-3;Q5VWW1-2;Q5VWW1;Q7Z5L3 | O75973 |
| Nucleobindin-2;Nesfatin-1 | NUCB2 | + | 1.3176 | 4.03824 | P80303-2;P80303 | P80303-2;P80303 |
| 60S ribosomal protein L24 | RPL24 | + | 1.36276 | 4.15014 | P83731 | P83731 |
| Charged multivesicular body protein 4b | CHMP4B | + | 2.11502 | 4.30521 | Q9H444 | Q9H444 |
| WNT1-inducible-signaling pathway protein 2 | WISP2 | + | 3.11243 | 4.34776 | O76076;O76076-2 | O76076 |
| Microfibrillar-associated protein 1 | MFAP1 | + | 2.08636 | 4.60037 | P55081 | P55081 |
| Stathmin | STMN1 | + | 4.83247 | 4.78368 | P16949;P16949-2 | P16949;P16949-2 |
| 40S ribosomal protein S28 | RPS28 | + | 1.5864 | 5.24325 | P62857 | P62857 |
| Neurosecretory protein VGF;Neuroendocrine regulatory peptide-1;Neuroendocrine reg | VGF | + | 4.53305 | 5.25135 | O15240 | O15240 |
| Matrix Gla protein | MGP | + | 3.10029 | 5.98888 | P08493;P08493-2 | P08493;P08493-2 |
| Vimentin | VIM | + | 4.46272 | 6.88996 | P08670;P41219;P41219-2 | P08670 |

**Sup. Table 5: SUM149 1D annotation enrichment (NODAL versus GFP)**

| C: Type | C: Name | N: Size | N: Score | N: P value | N: Benj. Hoch. FDR | N: Mean | N: Median |
|---|---|---|---|---|---|---|---|
| GOBP name | cellular component biogenesis | 12 | 0.719378 | 2.29E-05 | 0.00114383 | 2.22683 | 1.98378 |
| GOBP name | cellular component biogenesis at cellular level | 12 | 0.719378 | 2.29E-05 | 0.00112817 | 2.22683 | 1.98378 |
| GOBP name | ribonucleoprotein complex biogenesis | 12 | 0.719378 | 2.29E-05 | 0.00111292 | 2.22683 | 1.98378 |
| GOBP name | rRNA metabolic process | 13 | 0.711829 | 1.34E-05 | 0.000704955 | 2.19372 | 1.98832 |
| GOBP name | rRNA processing | 13 | 0.711829 | 1.34E-05 | 0.000694738 | 2.19372 | 1.98832 |
| GOCC name | cytosolic large ribosomal subunit | 17 | 0.712898 | 7.18E-07 | 4.27E-05 | 2.09685 | 2.01086 |
| GOCC name | large ribosomal subunit | 17 | 0.712898 | 7.18E-07 | 3.73E-05 | 2.09685 | 2.01086 |
| **GOBP name** | **cellular macromolecular complex disassembly** | **38** | **0.739766** | **1.02E-13** | **9.12E-11** | **2.08636** | **1.93777** |
| GOBP name | cellular protein complex disassembly | 38 | 0.739766 | 1.02E-13 | 7.30E-11 | 2.08636 | 1.93777 |
| GOBP name | macromolecular complex disassembly | 38 | 0.739766 | 1.02E-13 | 6.08E-11 | 2.08636 | 1.93777 |
| GOBP name | protein complex disassembly | 38 | 0.739766 | 1.02E-13 | 5.21E-11 | 2.08636 | 1.93777 |
| GOBP name | nuclear-transcribed mRNA catabolic process, nonsense-mediated decay | 37 | 0.727793 | 4.72E-13 | 1.41E-10 | 2.01346 | 1.91575 |
| GOBP name | translational termination | 37 | 0.727793 | 4.72E-13 | 1.30E-10 | 2.01346 | 1.91575 |
| GOBP name | viral transcription | 37 | 0.727793 | 4.72E-13 | 1.21E-10 | 2.01346 | 1.91575 |
| GOBP name | ncRNA metabolic process | 14 | 0.631169 | 6.32E-05 | 0.00280111 | 1.98141 | 1.98378 |
| GOBP name | ncRNA processing | 14 | 0.631169 | 6.32E-05 | 0.00276695 | 1.98141 | 1.98378 |
| GOBP name | establishment of protein localization in endoplasmic reticulum | 40 | 0.705757 | 3.95E-13 | 1.42E-10 | 1.97949 | 1.89818 |
| GOBP name | protein targeting to ER | 40 | 0.705757 | 3.95E-13 | 1.29E-10 | 1.97949 | 1.89818 |
| GOBP name | viral infectious cycle | 43 | 0.711968 | 4.25E-14 | 7.63E-11 | 1.974 | 1.8806 |
| GOBP name | translational elongation | 38 | 0.706054 | 1.25E-12 | 2.99E-10 | 1.94919 | 1.89818 |
| GOCC name | cytosolic small ribosomal subunit | 20 | 0.664198 | 6.17E-07 | 5.13E-05 | 1.94258 | 1.79149 |
| GOCC name | small ribosomal subunit | 20 | 0.664198 | 6.17E-07 | 4.28E-05 | 1.94258 | 1.79149 |
| GOBP name | cotranslational protein targeting to membrane | 39 | 0.693821 | 1.71E-12 | 3.84E-10 | 1.91986 | 1.8806 |
| GOBP name | SRP-dependent cotranslational protein targeting to membrane | 39 | 0.693821 | 1.71E-12 | 3.61E-10 | 1.91986 | 1.8806 |
| GOBP name | mRNA catabolic process | 39 | 0.692812 | 1.84E-12 | 3.67E-10 | 1.91812 | 1.8806 |
| GOBP name | nuclear-transcribed mRNA catabolic process | 39 | 0.692812 | 1.84E-12 | 3.48E-10 | 1.91812 | 1.8806 |
| GOBP name | RNA catabolic process | 39 | 0.692812 | 1.84E-12 | 3.30E-10 | 1.91812 | 1.8806 |
| GOBP name | viral reproductive process | 46 | 0.682375 | 9.33E-14 | 1.12E-10 | 1.88435 | 1.85321 |
| GOBP name | translational initiation | 39 | 0.681715 | 4.11E-12 | 5.91E-10 | 1.88428 | 1.8806 |
| **GOBP name** | **mRNA metabolic process** | **44** | **0.679848** | **3.25E-13** | **1.46E-10** | **1.87637** | **1.87165** |
| GOBP name | protein targeting to membrane | 42 | 0.663671 | 3.17E-12 | 4.95E-10 | 1.86069 | 1.87165 |
| GOCC name | ribosome | 16 | 0.668826 | 6.24E-06 | 0.000288336 | 1.8538 | 1.97406 |
| GOBP name | cellular macromolecular complex subunit organization | 53 | 0.669325 | 9.10E-15 | 3.27E-11 | 1.83749 | 1.91575 |
| GOCC name | ribonucleoprotein complex | 46 | 0.64984 | 1.29E-12 | 5.39E-10 | 1.7747 | 1.81246 |
| GOBP name | **establishment of protein localization to organelle** | 47 | 0.631492 | 3.47E-12 | 5.19E-10 | 1.75716 | 1.84373 |
| GOBP name | translation | 42 | 0.633712 | 2.83E-11 | 3.38E-09 | 1.74844 | 1.85321 |
| GOBP name | protein targeting | 52 | 0.60735 | 2.98E-12 | 5.09E-10 | 1.68797 | 1.81246 |
| GOBP name | RNA processing | 20 | 0.512037 | 0.000121235 | 0.00500268 | 1.63914 | 1.75021 |
| GOBP name | viral reproduction | 57 | 0.562076 | 2.03E-11 | 2.51E-09 | 1.63666 | 1.73926 |

| | | | | | | | |
|---|---|---|---|---|---|---|---|
| GOBP name | cellular macromolecular complex assembly | 20 | 0.495988 | 0.00019676 | 0.00751455 | 1.61116 | 1.86217 |
| GOBP name | protein complex subunit organization | 66 | 0.551995 | 3.12E-12 | 5.09E-10 | 1.54554 | 1.81246 |
| GOBP name | cellular process involved in reproduction | 49 | 0.544517 | 1.03E-09 | 1.03E-07 | 1.53247 | 1.73926 |
| GOBP name | intracellular protein transport | 59 | 0.562415 | 1.04E-11 | 1.44E-09 | 1.5315 | 1.61981 |
| GOBP name | macromolecular complex subunit organization | 71 | 0.560233 | 3.51E-13 | 1.40E-10 | 1.53075 | 1.73926 |
| GOBP name | gene expression | 57 | 0.547772 | 6.44E-11 | 7.00E-09 | 1.4702 | 1.61981 |
| GOBP name | RNA biosynthetic process | 57 | 0.563176 | 1.86E-11 | 2.38E-09 | 1.4663 | 1.61981 |
| GOCC name | nucleolus | 28 | 0.502486 | 1.05E-05 | 0.000434898 | 1.45443 | 1.89818 |
| GOBP name | cellular component disassembly | 60 | 0.498944 | 1.25E-09 | 1.21E-07 | 1.4054 | 1.59301 |
| GOBP name | cellular component disassembly at cellular level | 60 | 0.498944 | 1.25E-09 | 1.18E-07 | 1.4054 | 1.59301 |
| GOBP name | intracellular transport | 70 | 0.520125 | 1.85E-11 | 2.46E-09 | 1.39797 | 1.53891 |
| GOBP name | RNA metabolic process | 62 | 0.52814 | 7.41E-11 | 7.83E-09 | 1.3858 | 1.57681 |
| GOBP name | protein transport | 73 | 0.501289 | 4.87E-11 | 5.64E-09 | 1.34955 | 1.55345 |
| GOBP name | cellular macromolecule catabolic process | 54 | 0.476756 | 2.65E-08 | 2.16E-06 | 1.33155 | 1.48014 |
| GOBP name | nucleic acid metabolic process | 68 | 0.504369 | 1.17E-10 | 1.20E-08 | 1.29869 | 1.53891 |
| GOBP name | establishment of protein localization | 78 | 0.486794 | 6.21E-11 | 6.97E-09 | 1.29471 | 1.4656 |
| GOBP name | macromolecule catabolic process | 61 | 0.459654 | 1.78E-08 | 1.53E-06 | 1.28148 | 1.35446 |
| GOBP name | macromolecule biosynthetic process | 68 | 0.472293 | 1.60E-09 | 1.48E-07 | 1.24111 | 1.3911 |
| GOBP name | cellular macromolecule biosynthetic process | 62 | 0.456417 | 1.83E-08 | 1.52E-06 | 1.20741 | 1.42635 |
| GOBP name | nucleobase-containing compound metabolic process | 76 | 0.410153 | 4.80E-08 | 3.52E-06 | 1.06122 | 1.3489 |
| GOBP name | establishment of localization in cell | 96 | 0.358535 | 2.49E-07 | 1.68E-05 | 0.922635 | 1.28933 |
| GOCC name | nuclear part | 74 | 0.281181 | 0.000210348 | 0.00673114 | 0.856789 | 0.909849 |
| GOBP name | nitrogen compound metabolic process | 99 | 0.328963 | 1.78E-06 | 0.000108008 | 0.852238 | 1.04618 |
| GOBP name | cellular nitrogen compound metabolic process | 89 | 0.320908 | 6.54E-06 | 0.000366864 | 0.843289 | 1.23705 |
| GOBP name | transport | 122 | 0.352681 | 6.22E-08 | 4.38E-06 | 0.840379 | 1.23906 |
| GOBP name | establishment of localization | 127 | 0.356218 | 3.50E-08 | 2.79E-06 | 0.835741 | 1.11211 |
| GOBP name | biosynthetic process | 99 | 0.333581 | 1.27E-06 | 7.85E-05 | 0.818381 | 1.23705 |
| GOCC name | macromolecular complex | 137 | 0.322896 | 3.97E-07 | 4.12E-05 | 0.816903 | 1.00278 |
| GOCC name | cell-substrate junction | 63 | 0.276959 | 0.000589744 | 0.0163556 | 0.814809 | 1.32225 |
| GOBP name | cellular biosynthetic process | 97 | 0.32877 | 2.08E-06 | 0.000124478 | 0.814118 | 1.24106 |
| GOBP name | reproductive process | 79 | 0.283305 | 0.000132129 | 0.0053297 | 0.80117 | 1.32623 |
| GOCC name | adherens junction | 65 | 0.272015 | 0.000635748 | 0.0155571 | 0.796719 | 1.27846 |
| GOCC name | anchoring junction | 65 | 0.272015 | 0.000635748 | 0.0146928 | 0.796719 | 1.27846 |
| GOBP name | cellular component organization at cellular level | 146 | 0.345164 | 4.44E-08 | 3.39E-06 | 0.779866 | 1.00849 |
| GOBP name | cellular component organization or biogenesis at cellular level | 146 | 0.345164 | 4.44E-08 | 3.32E-06 | 0.779866 | 1.00849 |
| GOBP name | cellular component organization | 170 | 0.362542 | 6.08E-09 | 5.46E-07 | 0.747921 | 0.996711 |
| GOBP name | cellular component organization or biogenesis | 170 | 0.362542 | 6.08E-09 | 5.33E-07 | 0.747921 | 0.996711 |
| GOBP name | cellular catabolic process | 81 | 0.288363 | 8.68E-05 | 0.00366697 | 0.739984 | 1.23705 |
| GOCC name | cell junction | 79 | 0.261619 | 0.000415781 | 0.0123546 | 0.738402 | 1.04929 |
| GOCC name | intracellular organelle part | 199 | 0.361012 | 1.07E-08 | 2.24E-06 | 0.675695 | 0.869493 |
| GOBP name | cellular macromolecule metabolic process | 135 | 0.298742 | 2.88E-06 | 0.000169355 | 0.662306 | 0.986579 |

| | | | | | | | |
|---|---|---|---|---|---|---|---|
| GOBP name | cellular protein metabolic process | 105 | 0.268619 | 7.25E-05 | 0.00313369 | 0.661126 | 1.04618 |
| GOBP name | catabolic process | 90 | 0.254156 | 0.000339217 | 0.0117095 | 0.649519 | 1.04773 |
| GOCC name | organelle part | 204 | 0.339356 | 8.90E-08 | 1.23E-05 | 0.634718 | 0.863709 |
| GOCC name | membrane | 195 | 0.276269 | 1.13E-05 | 0.0004261 | 0.536919 | 0.868708 |
| GOBP name | macromolecule metabolic process | 158 | 0.259698 | 3.30E-05 | 0.00154002 | 0.511898 | 0.931968 |
| GOCC name | intracellular membrane-bounded organelle | 167 | 0.213843 | 0.000607141 | 0.0157857 | 0.49972 | 0.81352 |
| GOBP name | positive regulation of biological process | 141 | -0.222164 | 0.000456686 | 0.0153225 | -0.427352 | -0.444304 |
| GOBP name | regulation of response to stimulus | 97 | -0.248967 | 0.000326167 | 0.0113683 | -0.696654 | -0.444304 |
| GOBP name | negative regulation of molecular function | 41 | -0.355389 | 0.000220963 | 0.00801271 | -0.807783 | -0.82756 |
| GOBP name | response to stress | 92 | -0.308144 | 1.21E-05 | 0.000650051 | -0.820064 | -0.581744 |
| GOCC name | extracellular region | 81 | -0.311928 | 2.18E-05 | 0.000756207 | -0.903124 | -0.912401 |
| GOBP name | negative regulation of catalytic activity | 37 | -0.385685 | 0.00012652 | 0.00516143 | -0.914761 | -0.87749 |
| GOBP name | immune system process | 72 | -0.302083 | 8.08E-05 | 0.00345189 | -0.975128 | -0.620008 |
| GOBP name | regulation of response to stress | 41 | -0.358126 | 0.000197491 | 0.00746308 | -1.04604 | -0.62894 |
| GOBP name | positive regulation of response to stimulus | 58 | -0.312515 | 0.000174487 | 0.00673557 | -1.14132 | -0.644478 |
| GOBP name | regulation of immune system process | 52 | -0.337197 | 0.000106796 | 0.0044581 | -1.1872 | -0.675377 |
| GOBP name | regulation of locomotion | 45 | -0.352657 | 0.000136421 | 0.00544168 | -1.25472 | -0.691112 |
| GOBP name | defense response | 59 | -0.402795 | 1.11E-06 | 7.01E-05 | -1.29462 | -1.0245 |
| GOBP name | regulation of cell motility | 42 | -0.357616 | 0.000172827 | 0.00674401 | -1.32637 | -0.729335 |
| GOBP name | regulation of cell migration | 41 | -0.366981 | 0.000136606 | 0.00538916 | -1.3746 | -0.767557 |
| GOBP name | negative regulation of hydrolase activity | 25 | -0.512476 | 1.98E-05 | 0.00101304 | -1.38989 | -1.06049 |
| GOBP name | response to cytokine stimulus | 26 | -0.415578 | 0.0004255 | 0.0144108 | -1.43645 | -1.10286 |
| GOBP name | negative regulation of peptidase activity | 21 | -0.54268 | 3.08E-05 | 0.00147221 | -1.46681 | -1.23042 |
| GOBP name | immune response | 51 | -0.435455 | 6.91E-07 | 4.51E-05 | -1.53945 | -1.0245 |
| GOBP name | negative regulation of endopeptidase activity | 18 | -0.593729 | 2.22E-05 | 0.00112439 | -1.60904 | -1.2318 |
| **GOBP name** | **positive regulation of immune system process** | **36** | **-0.413781** | **4.84E-05** | **0.00220144** | **-1.61531** | **-1.05348** |
| GOBP name | regulation of peptidase activity | 27 | -0.529618 | 4.90E-06 | 0.000279197 | -1.63624 | -1.23042 |
| GOBP name | regulation of endopeptidase activity | 25 | -0.592476 | 8.06E-07 | 5.17E-05 | -1.84988 | -1.23317 |
| GOBP name | positive regulation of locomotion | 28 | -0.417722 | 0.000248266 | 0.00882449 | -1.85845 | -1.48619 |
| **GOBP name** | **positive regulation of cell migration** | **27** | **-0.431008** | **0.000200542** | **0.00749944** | **-1.943** | **-1.50366** |
| GOBP name | positive regulation of cell motility | 27 | -0.431008 | 0.000200542 | 0.00742213 | -1.943 | -1.50366 |
| GOBP name | regulation of response to external stimulus | 33 | -0.47072 | 8.71E-06 | 0.000474016 | -1.98605 | -1.25309 |
| GOBP name | response to wounding | 32 | -0.585537 | 4.88E-08 | 3.50E-06 | -2.19463 | -1.57067 |
| GOBP name | positive regulation of response to external stimulus | 22 | -0.463862 | 0.000271947 | 0.00957146 | -2.30151 | -1.9509 |
| **GOBP name** | **cytokine-mediated signaling pathway** | **14** | **-0.563636** | **0.000353602** | **0.0120898** | **-2.44117** | **-1.15717** |
| **GOBP name** | **inflammatory response** | **27** | **-0.638977** | **3.54E-08** | **2.76E-06** | **-2.50001** | **-1.75002** |
| GOBP name | acute inflammatory response | 8 | -0.758185 | 0.000246796 | 0.00885996 | -2.5783 | -1.84112 |
| GOBP name | regulation of leukocyte migration | 17 | -0.594531 | 3.57E-05 | 0.00164398 | -2.89795 | -3.37532 |
| GOBP name | response to molecule of bacterial origin | 22 | -0.675607 | 1.14E-07 | 7.88E-06 | -3.12271 | -3.17739 |
| GOBP name | response to lipopolysaccharide | 20 | -0.682716 | 2.98E-07 | 1.98E-05 | -3.31659 | -3.41913 |
| GOBP name | regulation of leukocyte chemotaxis | 12 | -0.680723 | 6.16E-05 | 0.00276371 | -3.4032 | -3.48782 |

| | | | | | | | |
|---|---|---|---|---|---|---|---|
| GOBP name | positive regulation of leukocyte chemotaxis | 11 | -0.738466 | 3.08E-05 | 0.00145321 | -3.75118 | -3.5127 |
| GOBP name | positive regulation of leukocyte migration | 13 | -0.736463 | 6.64E-06 | 0.000366711 | -3.79897 | -3.5127 |
| GOBP name | cell chemotaxis | 13 | -0.763886 | 2.97E-06 | 0.000172152 | -4.09714 | -4.13373 |
| GOBP name | chemokine-mediated signaling pathway | 5 | -0.965782 | 0.000209077 | 0.00765906 | -5.5704 | -4.81573 |

**Sup. Table 6: Proteins differentially expressed between CM from NODAL and EV expressing SUM149 cells**

| Protein names | Gene names | T-test Significant | -Log T-test p-value | Log2 T-test fold-change (NODAL-EV) | Protein IDs | Majority protein IDs |
|---|---|---|---|---|---|---|
| Vimentin | VIM | + | 4.16185 | 6.70008 | P08670;P41219;P41219-2 | P08670 |
| Matrix Gla protein | MGP | + | 4.143 | 5.88634 | P08493;P08493-2 | P08493;P08493-2 |
| Neurosecretory protein VGF;Neuroendocrine regulatory peptide-1;Neuroendocrine regulatory peptide-2;Antim | VGF | + | 5.65461 | 5.72969 | O15240 | O15240 |
| Melanoma-derived growth regulatory protein | MIA | + | 4.57681 | 4.54431 | Q16674 | Q16674 |
| Stathmin | STMN1 | + | 4.97419 | 4.50183 | P16949;P16949-2 | P16949;P16949-2 |
| Nucleobindin-2;Nesfatin-1 | NUCB2 | | 1.07118 | 3.9263 | P80303-2;P80303 | P80303-2;P80303 |
| Secretogranin-1;PE-11;GAWK peptide;CCB peptide | CHGB | + | 3.29225 | 3.81956 | P05060 | P05060 |
| EGF-containing fibulin-like extracellular matrix protein 1 | EFEMP1 | | 1.13368 | 3.79935 | Q12805-5 | Q12805-5 |
| Microfibrillar-associated protein 1 | MFAP1 | + | 2.08846 | 3.78647 | P55081 | P55081 |
| C1q-related factor | C1QL1 | + | 2.20469 | 3.72135 | O75973;Q5VWW1-3;Q5VWW1-2;Q5VWW1;Q7Z5L3 | O75973 |
| WNT1-inducible-signaling pathway protein 2 | WISP2 | + | 4.04063 | 3.71122 | O76076;O76076-2 | O76076 |
| Tropomyosin alpha-3 chain | TPM3 | | 0.701583 | 3.67077 | P06753 | P06753 |
| Tyrosyl-DNA phosphodiesterase 2 | TDP2 | | 1.00809 | 3.5857 | O95551;O95551-2;O95551-3 | O95551;O95551-2;O95551-3 |
| 40S ribosomal protein S28 | RPS28 | + | 3.2433 | 3.50069 | P62857 | P62857 |
| V-type proton ATPase subunit S1 | ATP6AP1 | + | 3.38992 | 3.48031 | Q15904 | Q15904 |
| Biglycan | BGN | + | 4.69021 | 3.47491 | P21810 | P21810 |
| CD151 antigen | CD151 | + | 1.52259 | 3.44048 | P48509 | P48509 |
| Serine protease 23 | PRSS23 | + | 1.41673 | 3.19419 | O95084;O95084-2 | O95084;O95084-2 |
| Charged multivesicular body protein 5 | CHMP5 | + | 2.16838 | 3.09315 | Q9NZZ3;Q9NZZ3-2 | Q9NZZ3;Q9NZZ3-2 |
| 14-3-3 protein beta/alpha;14-3-3 protein beta/alpha, N-terminally processed | YWHAB | + | 3.19389 | 3.09175 | P31946-2;P31946 | P31946-2;P31946 |
| POTE ankyrin domain family member E;POTE ankyrin domain family member F | POTEE;POTEF | | 0.294131 | 3.03939 | Q6S8J3;A5A3E0;Q6S8J3-3;Q6S8J3-2 | Q6S8J3;A5A3E0 |
| 60S ribosomal protein L24 | RPL24 | | 0.615141 | 3.03722 | P83731 | P83731 |
| Charged multivesicular body protein 4b | CHMP4B | + | 2.27693 | 2.92235 | Q9H444 | Q9H444 |
| Neuroblast differentiation-associated protein AHNAK | AHNAK | + | 2.12884 | 2.9046 | Q09666;Q09666-2 | Q09666 |
| Secretogranin-2;Secretoneurin;Manserin | SCG2 | | 0.972181 | 2.89027 | P13521 | P13521 |
| Plasminogen activator inhibitor 1 RNA-binding protein | SERBP1 | + | 4.05277 | 2.87416 | Q8NC51-3;Q8NC51-4 | Q8NC51-3;Q8NC51-4 |
| Myotrophin | MTPN | | 1.27585 | 2.86717 | P58546 | P58546 |
| 60S ribosomal protein L10a | RPL10A | + | 2.92877 | 2.82303 | P62906 | P62906 |
| Angiopoietin-1 | ANGPT1 | + | 2.44066 | 2.73521 | Q15389-2;Q15389 | Q15389-2;Q15389 |
| Carboxypeptidase M | CPM | + | 1.47204 | 2.72182 | P14384 | P14384 |
| Adipogenesis regulatory factor | ADIRF | + | 1.45079 | 2.69921 | Q15847 | Q15847 |
| Lipoprotein lipase | LPL | + | 3.27642 | 2.654 | P06858 | P06858 |
| Actin, cytoplasmic 1;Actin, cytoplasmic 1, N-terminally processed | ACTB | | 0.342614 | 2.57707 | P60709 | P60709 |
| 40S ribosomal protein S10;Putative 40S ribosomal protein S10-like | RPS10;RPS10P5 | | 0.806673 | 2.54175 | P46783;Q9NQ39 | P46783;Q9NQ39 |
| Bone morphogenetic protein 2 | BMP2 | | 1.24266 | 2.52853 | P12643 | P12643 |
| EGF-containing fibulin-like extracellular matrix protein 1 | EFEMP1 | + | 2.93921 | 2.50379 | Q12805-2;Q12805-4;Q12805-3;Q12805 | Q12805-2;Q12805-4;Q12805-3;Q12805 |
| Insulin-like growth factor-binding protein 5 | IGFBP5 | | 0.888685 | 2.40462 | P24593 | P24593 |
| Disintegrin and metalloproteinase domain-containing protein 15 | ADAM15 | + | 2.34254 | 2.39849 | Q13444-10;Q13444-9;Q13444-8;Q13444-2;Q13444-7;Q13444-6;Q13 | Q13444-10;Q13444-9;Q13444-8;Q13444-2;Q13444-7;Q13 |
| Tyrosine-protein kinase receptor UFO | AXL | + | 1.73559 | 2.37908 | P30530-2;P30530 | P30530-2;P30530 |
| Serine/arginine-rich splicing factor 2 | SRSF2 | + | 2.29099 | 2.3615 | Q01130-2;Q01130;Q9BRL6-2;Q9BRL6 | Q01130-2;Q01130 |
| M-phase phosphoprotein 8 | MPHOSPH8 | + | 1.62914 | 2.35042 | Q99549;Q99549-2 | Q99549;Q99549-2 |
| BolA-like protein 3 | BOLA3 | + | 1.96632 | 2.34847 | Q53S33 | Q53S33 |
| Angiogenin | ANG | + | 1.72565 | 2.32347 | P03950 | P03950 |
| Ladinin-1 | LAD1 | + | 2.77554 | 2.31549 | O00515 | O00515 |
| 40S ribosomal protein S9 | RPS9 | + | 2.16926 | 2.29296 | P46781 | P46781 |
| Protein LAP2 | ERBB2IP | + | 2.65256 | 2.2915 | Q96RT1-6;Q96RT1-4;Q96RT1-5;Q96RT1-3;Q96RT1-9;Q96RT1-2;Q96 | Q96RT1-6;Q96RT1-4;Q96RT1-5;Q96RT1-3;Q96RT1-9;Q96 |
| Src substrate cortactin | CTTN | + | 2.56561 | 2.2471 | Q14247-3;Q14247;Q14247-2 | Q14247-3;Q14247;Q14247-2 |
| Deleted in autism protein 1 | C3orf58 | | 0.850649 | 2.21231 | Q8NDZ4;Q8NDZ4-2 | Q8NDZ4;Q8NDZ4-2 |
| Serine/arginine-rich splicing factor 7 | SRSF7 | | 0.697322 | 2.17067 | Q16629-3;Q16629-2;Q16629-4;Q16629 | Q16629-3;Q16629-2;Q16629-4;Q16629 |
| 14-3-3 protein theta | YWHAQ | + | 3.49775 | 2.1489 | P27348 | P27348 |
| Serine protease HTRA1 | HTRA1 | + | 3.91946 | 2.14285 | Q92743 | Q92743 |
| Complement factor D | CFD | + | 2.47681 | 2.12866 | P00746 | P00746 |
| Apolipoprotein C-III | APOC3 | | 1.02887 | 2.11481 | P02656 | P02656 |
| Kallikrein-6 | KLK6 | | 0.887989 | 2.11104 | Q92876;Q92876-2 | Q92876;Q92876-2 |
| 40S ribosomal protein S8 | RPS8 | + | 2.02379 | 2.08555 | P62241 | P62241 |
| Calcium-regulated heat stable protein 1 | CARHSP1 | | 0.818074 | 2.03383 | Q9Y2V2 | Q9Y2V2 |
| 60S ribosomal protein L14 | RPL14 | + | 1.84315 | 2.0283 | P50914 | P50914 |
| Integral membrane protein 2B;BRI2, membrane form;BRI2 intracellular domain;BRI2C, soluble form;Bri23 pepti | ITM2B | | 0.611898 | 2.01522 | Q9Y287;Q9Y287-2 | Q9Y287 |
| Double-stranded RNA-binding protein Staufen homolog 1 | STAU1 | + | 2.6766 | 2.00126 | O95793-2;O95793-3;O95793 | O95793-2;O95793-3;O95793 |
| 40S ribosomal protein S19 | RPS19 | + | 2.89509 | 1.98496 | P39019 | P39019 |
| 60S ribosomal protein L5 | RPL5 | + | 2.52215 | 1.97386 | P46777 | P46777 |
| 40S ribosomal protein S11 | RPS11 | + | 2.90398 | 1.95423 | P62280 | P62280 |
| 60S ribosomal protein L7a | RPL7A | + | 2.21964 | 1.91715 | P62424 | P62424 |
| 60S ribosomal protein L4 | RPL4 | + | 3.09002 | 1.90763 | P36578 | P36578 |
| Kazal-type serine protease inhibitor domain-containing protein 1 | KAZALD1 | | 0.745783 | 1.88817 | Q96I82-2;Q96I82 | Q96I82-2;Q96I82 |
| Interleukin-11 | IL11 | | 1.04955 | 1.8622 | P20809;P20809-2 | P20809;P20809-2 |
| 60S ribosomal protein L27 | RPL27 | + | 1.91134 | 1.8583 | P61353 | P61353 |
| 60S ribosomal protein L13a;Putative 60S ribosomal protein L13a protein RPL13AP3 | RPL13A;RPL13AP3 | + | 1.64327 | 1.85587 | P40429;Q6NVV1 | P40429;Q6NVV1 |
| 60S ribosomal protein L30 | RPL30 | + | 4.12975 | 1.85349 | P62888 | P62888 |
| Large neutral amino acids transporter small subunit 1 | SLC7A5 | | 0.950331 | 1.85252 | Q01650;Q9UM01 | Q01650 |
| ADAMTS-like protein 4 | ADAMTSL4 | | 0.719628 | 1.85121 | Q6UY14;Q6UY14-3;Q6UY14-2 | Q6UY14;Q6UY14-3;Q6UY14-2 |
| Charged multivesicular body protein 2b | CHMP2B | + | 1.49622 | 1.84067 | Q9UQN3-2;Q9UQN3 | Q9UQN3-2;Q9UQN3 |
| Clusterin;Clusterin beta chain;Clusterin alpha chain | CLU | + | 4.06882 | 1.83264 | P10909-4;P10909;P10909-5;P10909-2;P10909-3 | P10909-4;P10909;P10909-5;P10909-2;P10909-3 |
| ATP synthase subunit d, mitochondrial | ATP5H | | 1.14125 | 1.83158 | O75947-2;O75947 | O75947-2;O75947 |
| 40S ribosomal protein S5;40S ribosomal protein S5, N-terminally processed | RPS5 | | 1.06562 | 1.81139 | P46782 | P46782 |
| Lactadherin;Lactadherin short form;Medin | MFGE8 | + | 2.17164 | 1.78483 | Q08431;Q08431-3 | Q08431;Q08431-3 |
| MARCKS-related protein | MARCKSL1 | | 0.807976 | 1.72743 | P49006 | P49006 |

| | | | | | | |
|---|---|---|---|---|---|---|
| HBS1-like protein | HBS1L | | 1.27252 | 1.72049 | Q9Y450;Q9Y450-4;Q9Y450-2 | Q9Y450;Q9Y450-4 |
| 40S ribosomal protein S14 | RPS14 | + | 2.63391 | 1.71861 | P62263 | P62263 |
| Vacuolar protein sorting-associated protein 28 homolog | VPS28 | | 0.286233 | 1.70799 | Q9UK41;Q9UK41-2 | Q9UK41;Q9UK41-2 |
| 60S ribosomal protein L26-like 1;60S ribosomal protein L26 | RPL26L1;RPL26 | | 0.937059 | 1.7075 | Q9UNX3;P61254 | Q9UNX3;P61254 |
| Neurotrypsin | PRSS12 | | 0.746757 | 1.68498 | P56730 | P56730 |
| 40S ribosomal protein S26;Putative 40S ribosomal protein S26-like 1 | RPS26;RPS26P11 | + | 2.14446 | 1.66537 | P62854;Q5JNZ5 | P62854;Q5JNZ5 |
| Epidermal growth factor receptor kinase substrate 8 | EPS8 | + | 1.77493 | 1.66463 | Q12929;Q12929-2 | Q12929;Q12929-2 |
| 40S ribosomal protein S17-like;40S ribosomal protein S17 | RPS17L;RPS17 | + | 2.14641 | 1.64508 | P0CW22;P08708 | P0CW22;P08708 |
| 14-3-3 protein zeta/delta | YWHAZ | + | 5.12993 | 1.62481 | P63104;P63104-2 | P63104;P63104-2 |
| Complement factor I;Complement factor I heavy chain;Complement factor I light chain | CFI | | 0.903687 | 1.62103 | P05156 | P05156 |
| Tubulin beta-8 chain | TUBB8 | | 0.588102 | 1.61595 | Q3ZCM7 | Q3ZCM7 |
| 40S ribosomal protein S13 | RPS13 | | 1.02667 | 1.61007 | P62277 | P62277 |
| 40S ribosomal protein S25 | RPS25 | + | 1.47034 | 1.59818 | P62851 | P62851 |
| Plasma kallikrein;Plasma kallikrein heavy chain;Plasma kallikrein light chain | KLKB1 | | 0.262155 | 1.598 | P03952 | P03952 |
| Nucleobindin-1 | NUCB1 | + | 2.10864 | 1.58898 | Q02818 | Q02818 |
| BolA-like protein 2 | BOLA2 | | 1.10854 | 1.56744 | Q9H3K6;Q9H3K6-2 | Q9H3K6;Q9H3K6-2 |
| Calumenin | CALU | | 1.1063 | 1.56449 | O43852;O43852-3;O43852-6;O43852-10;O43852-5;O43852-13;O438 | O43852;O43852-3;O43852-6;O43852-10;O43852-5;O438 |
| 60S ribosomal protein L15 | RPL15 | + | 3.0548 | 1.55735 | P61313;P61313-2 | P61313 |
| Heterogeneous nuclear ribonucleoprotein K | HNRNPK | | 0.188151 | 1.55049 | P61978-2;P61978 | P61978-2;P61978 |
| Proteoglycan 4;Proteoglycan 4 C-terminal part | PRG4 | | 0.864212 | 1.54258 | Q92954-5;Q92954-4;Q92954-3;Q92954-6;Q92954-2;Q92954 | Q92954-5;Q92954-4;Q92954-3;Q92954-6;Q92954-2;Q92 |
| Coxsackievirus and adenovirus receptor | CXADR | | 0.63576 | 1.52565 | P78310-5;P78310-7;P78310-2;P78310-6;P78310;P78310-4;P78310-3 | P78310-5;P78310-7;P78310-2;P78310-6;P78310;P78310- |
| GTPase NRas | NRAS | | 0.675672 | 1.50724 | P01111 | P01111 |
| Protein ELYS | AHCTF1 | | 0.940654 | 1.50655 | Q8WYP5-3;Q8WYP5;Q8WYP5-2 | Q8WYP5-3;Q8WYP5;Q8WYP5-2 |
| Major prion protein | PRNP | | 1.16788 | 1.50409 | P04156-2;P04156 | P04156-2;P04156 |
| Alpha-N-acetylglucosaminidase;Alpha-N-acetylglucosaminidase 82 kDa form;Alpha-N-acetylglucosaminidase 77 | NAGLU | | 0.586376 | 1.50107 | P54802 | P54802 |
| Renin receptor | ATP6AP2 | | 1.00826 | 1.49124 | O75787-2;O75787 | O75787-2;O75787 |
| 40S ribosomal protein S2 | RPS2 | + | 2.19125 | 1.48491 | P15880 | P15880 |
| 60S ribosomal protein L7 | RPL7 | + | 1.78874 | 1.48332 | P18124 | P18124 |
| Tolloid-like protein 1 | TLL1 | | 1.1342 | 1.47912 | O43897;O43897-2 | O43897 |
| 14-3-3 protein eta | YWHAH | + | 2.78958 | 1.47085 | Q04917 | Q04917 |
| 45 kDa calcium-binding protein | SDF4 | + | 2.50547 | 1.45898 | Q9BRK5;Q9BRK5-6;Q9BRK5-2;Q9BRK5-4;Q9BRK5-3;Q9BRK5-5 | Q9BRK5;Q9BRK5-6;Q9BRK5-2 |
| Rac GTPase-activating protein 1 | RACGAP1 | | 0.664132 | 1.45481 | Q9H0H5 | Q9H0H5 |
| Solute carrier family 2, facilitated glucose transporter member 1 | SLC2A1 | + | 1.72132 | 1.44536 | P11166 | P11166 |
| Tubulin beta-6 chain | TUBB6 | + | 1.30798 | 1.44404 | Q9BUF5 | Q9BUF5 |
| Calumenin | CALU | | 0.98197 | 1.44366 | O43852-2;O43852-4;O43852-8;O43852-7 | O43852-2;O43852-4;O43852-8 |
| Endothelial lipase | LIPG | + | 1.87023 | 1.43627 | Q9Y5X9;Q9Y5X9-2 | Q9Y5X9;Q9Y5X9-2 |
| Amiloride-sensitive amine oxidase [copper-containing] | AOC1 | | 0.385458 | 1.42988 | P19801;P19801-2 | P19801;P19801-2 |
| Barrier-to-autointegration factor;Barrier-to-autointegration factor, N-terminally processed | BANF1 | | 1.17427 | 1.42731 | O75531 | O75531 |
| Lymphocyte function-associated antigen 3 | CD58 | | 0.583972 | 1.4058 | P19256-2;P19256-3;P19256 | P19256-2;P19256-3;P19256 |
| Collagen alpha-1(XII) chain | COL12A1 | + | 1.76932 | 1.39675 | Q99715;Q99715-4;Q99715-2 | Q99715;Q99715-4;Q99715-2 |
| Heat shock protein beta-1 | HSPB1 | + | 2.02611 | 1.39545 | P04792 | P04792 |
| 40S ribosomal protein S4, X isoform;40S ribosomal protein S4, Y isoform 1;40S ribosomal protein S4, Y isoform 2 | RPS4X;RPS4Y1;RPS4Y2 | + | 2.1764 | 1.38121 | P62701;P22090;Q8TD47 | P62701;P22090;Q8TD47 |
| 40S ribosomal protein S18 | RPS18 | + | 2.56925 | 1.37536 | P62269 | P62269 |
| HD domain-containing protein 2 | HDDC2 | | 0.339883 | 1.3734 | Q7Z4H3-2;Q7Z4H3;Q7Z4H3-3 | Q7Z4H3-2;Q7Z4H3 |
| Spondin-2 | SPON2 | + | 1.8277 | 1.3725 | Q9BUD6 | Q9BUD6 |
| Mammalian ependymin-related protein 1 | EPDR1 | + | 1.34076 | 1.34762 | Q9UM22;Q9UM22-3;Q9UM22-2 | Q9UM22;Q9UM22-3 |
| 40S ribosomal protein SA | RPSA | + | 3.13268 | 1.33589 | P08865 | P08865 |
| Histone H2B type 1-L;Histone H2B type 1-M;Histone H2B type 1-N;Histone H2B type 1-H;Histone H2B type 2-F;H | HIST1H2BL;HIST1H2BM;HIST1H2BN;HIST1 | + | 1.75839 | 1.33313 | Q99880;Q99879;Q99877;Q93079;Q5QNW6;P62807;P58876;P57053 | Q99880;Q99879;Q99877;Q93079;Q5QNW6;P62807;P588 |
| 40S ribosomal protein S3 | RPS3 | + | 1.57951 | 1.32241 | P23396;P23396-2 | P23396;P23396-2 |
| E3 ubiquitin-protein ligase CHIP | STUB1 | + | 2.57024 | 1.32112 | Q9UNE7;Q9UNE7-2 | Q9UNE7;Q9UNE7-2 |
| ADP-ribosylation factor 5 | ARF5 | | 0.490027 | 1.31465 | P84085 | P84085 |
| Histone H4 | HIST1H4A | + | 1.725 | 1.30201 | P62805 | P62805 |
| Procollagen C-endopeptidase enhancer 2 | PCOLCE2 | + | 1.73337 | 1.285 | Q9UKZ9 | Q9UKZ9 |
| Versican core protein | VCAN | | 1.1957 | 1.27225 | P13611-2;P13611;P13611-5;P13611-4;P13611-3 | P13611-2;P13611;P13611-5;P13611-4;P13611-3 |
| Transforming growth factor beta-2;Latency-associated peptide | TGFB2 | | 1.11844 | 1.26698 | P61812 | P61812 |
| Fibrillin-1 | FBN1 | | 1.22613 | 1.26419 | P35555 | P35555 |
| Sequestosome-1 | SQSTM1 | | 0.843313 | 1.26102 | Q13501;Q13501-2 | Q13501;Q13501-2 |
| Furin | FURIN | + | 1.66322 | 1.25992 | P09958 | P09958 |
| Tax1-binding protein 3 | TAX1BP3 | + | 1.46556 | 1.25195 | O14907 | O14907 |
| Histone H2B type 2-E;Histone H2B type 1-B;Histone H2B type 1-O;Histone H2B type 1-J;Histone H2B type 3-B | HIST2H2BE;HIST1H2BB;HIST1H2BO;HIST1H2BJ;HIST3H2B | | 1.07662 | 1.24787 | Q16778;P33778;P23527;P06899;Q8N257 | Q16778;P33778;P23527;P06899;Q8N257 |
| Band 4.1-like protein 2 | EPB41L2 | + | 1.54171 | 1.23934 | O43491;O43491-3;O43491-2;Q9Y2J2-3;Q9Y2J2-2;Q9Y2J2-4;Q9Y2J2 | O43491;O43491-3;O43491-2 |
| Ras-related protein Ral-A | RALA | + | 1.69223 | 1.2208 | P11233 | P11233 |
| Dermcidin;Survival-promoting peptide;DCD-1 | DCD | | 0.50646 | 1.2139 | P81605;P81605-2 | P81605;P81605-2 |
| Procollagen-lysine,2-oxoglutarate 5-dioxygenase 1 | PLOD1 | + | 2.50052 | 1.17174 | Q02809;Q02809-2 | Q02809;Q02809-2 |
| Polyadenylate-binding protein 2 | PABPN1 | | 1.16459 | 1.16548 | Q86U42-2;Q86U42;Q92843-2 | Q86U42-2;Q86U42;Q92843-2 |
| 60S ribosomal protein L22 | RPL22 | + | 1.75328 | 1.16472 | P35268 | P35268 |
| Alpha-internexin | INA | | 0.533079 | 1.16287 | Q16352 | Q16352 |
| Tissue factor | F3 | + | 1.9145 | 1.16097 | P13726;P13726-2 | P13726;P13726-2 |
| Secreted frizzled-related protein 1 | SFRP1 | | 0.727838 | 1.152 | Q8N474 | Q8N474 |
| SPARC | SPARC | | 1.22799 | 1.14822 | P09486 | P09486 |
| Sodium/potassium-transporting ATPase subunit beta-3 | ATP1B3 | + | 2.37643 | 1.1459 | P54709;P54709-2 | P54709;P54709-2 |
| 40S ribosomal protein S16 | RPS16 | + | 1.98207 | 1.14249 | P62249 | P62249 |
| MAGUK p55 subfamily member 6 | MPP6 | | 0.507704 | 1.1419 | Q9NZW5 | Q9NZW5 |
| Scaffold attachment factor B2 | SAFB2 | | 0.853859 | 1.12811 | Q14151;Q14151-2 | Q14151 |
| Deleted in malignant brain tumors 1 protein | DMBT1 | + | 1.80255 | 1.12694 | Q9UGM3-4;Q9UGM3-8;Q9UGM3-2;Q9UGM3-5;Q9UGM3-3;Q9UGM | Q9UGM3-4;Q9UGM3-8;Q9UGM3-2;Q9UGM3-5;Q9UGM3 |
| 40S ribosomal protein S3a | RPS3A | + | 1.34409 | 1.12287 | P61247 | P61247 |
| Protein PRRC2A | PRRC2A | + | 1.54162 | 1.11913 | P48634;P48634-3;P48634-2;P48634-4 | P48634;P48634-3;P48634-2;P48634-4 |
| Small nuclear ribonucleoprotein E | SNRPE | | 0.184697 | 1.11792 | P62304 | P62304 |
| Tyrosine-protein kinase Yes | YES1 | | 0.361976 | 1.11619 | P07947 | P07947 |
| Signal recognition particle 9 kDa protein | SRP9 | + | 2.32554 | 1.1036 | P49458;P49458-2 | P49458;P49458-2 |

| | | | | | | |
|---|---|---|---|---|---|---|
| Sodium/potassium-transporting ATPase subunit alpha-1 | ATP1A1 | + | 2.59225 | 1.10266 | P05023-4;P05023;P05023-3;P05023-2;P20648;P54707;P54707-2 | P05023-4;P05023;P05023-3;P05023-2 |
| Neutral amino acid transporter B(0) | SLC1A5 | | 0.866907 | 1.10264 | Q15758;Q15758-3;Q15758-2 | Q15758;Q15758-3;Q15758-2 |
| Syndecan-1 | SDC1 | | 0.325421 | 1.09862 | P18827 | P18827 |
| 40S ribosomal protein S7 | RPS7 | | 0.588899 | 1.08763 | P62081 | P62081 |
| Protein S100-A13 | S100A13 | | 0.846283 | 1.08647 | Q99584 | Q99584 |
| Proprotein convertase subtilisin/kexin type 9 | PCSK9 | + | 1.90683 | 1.08486 | Q8NBP7;Q8NBP7-2 | Q8NBP7 |
| Ras-related protein Rab-3D | RAB3D | | 0.574422 | 1.07523 | O95716;P20337;Q96E17 | O95716 |
| Histone H2A type 2-C;Histone H2A type 2-A | HIST2H2AC;HIST2H2AA3 | | 1.28812 | 1.063 | Q16777;Q6FI13 | Q16777;Q6FI13 |
| GRIP1-associated protein 1 | GRIPAP1 | | 0.65283 | 1.04843 | Q4V328-4;Q4V328;Q4V328-3;Q4V328-2 | Q4V328-4;Q4V328;Q4V328-3;Q4V328-2 |
| Guanine nucleotide-binding protein G(s) subunit alpha isoforms short;Guanine nucleotide-binding protein G(s) | GNAS | | 1.20355 | 1.03573 | P63092;P63092-4;Q5JWF2 | P63092;P63092-4;Q5JWF2 |
| Ephrin-B1 | EFNB1 | | 0.33212 | 1.03315 | P98172 | P98172 |
| Decorin | DCN | | 1.04143 | 1.02779 | P07585;P07585-4;P07585-2;P07585-3;P07585-5 | P07585;P07585-4 |
| Prefoldin subunit 2 | PFDN2 | | 1.05922 | 1.02677 | Q9UHV9 | Q9UHV9 |
| Epididymal secretory protein E1 | NPC2 | | 0.92612 | 1.01867 | P61916;P61916-2 | P61916;P61916-2 |
| Procollagen-lysine,2-oxoglutarate 5-dioxygenase 3 | PLOD3 | + | 1.5875 | 1.01851 | O60568 | O60568 |
| Serine protease inhibitor Kazal-type 5;Hemofiltrate peptide HF6478;Hemofiltrate peptide HF7665 | SPINK5 | + | 1.37471 | 1.01649 | Q9NQ38;Q9NQ38-3;Q9NQ38-2 | Q9NQ38;Q9NQ38-3;Q9NQ38-2 |
| Synapse-associated protein 1 | SYAP1 | | 0.768897 | 1.00822 | Q96A49 | Q96A49 |
| Non-specific lipid-transfer protein | SCP2 | + | 2.22155 | 0.999908 | P22307-6;P22307-2;P22307-4;P22307-7;P22307-8;P22307 | P22307-6;P22307-2;P22307-4;P22307-7;P22307-8;P2230 |
| Heat shock-related 70 kDa protein 2 | HSPA2 | | 0.591428 | 0.995817 | P54652 | P54652 |
| ATP-dependent RNA helicase DDX19A;ATP-dependent RNA helicase DDX19B | DDX19A;DDX19B | | 0.350415 | 0.993593 | Q9NUU7;Q9NUU7-2;Q9UMR2-2;Q9UMR2-4;Q9UMR2;Q9UMR2-3 | Q9NUU7;Q9NUU7-2;Q9UMR2-2;Q9UMR2-4;Q9UMR2;Q |
| Guanine nucleotide-binding protein G(k) subunit alpha | GNAI3 | + | 1.72512 | 0.990112 | P08754 | P08754 |
| Proprotein convertase subtilisin/kexin type 6 | PCSK6 | + | 4.64708 | 0.988755 | P29122-2;P29122-8;P29122;P29122-7;P29122-5;P29122-4;P29122-3 | P29122-2;P29122-8;P29122;P29122-7;P29122-5;P29122- |
| Amphiregulin | AREG | | 0.645072 | 0.988731 | P15514 | P15514 |
| UPF0160 protein MYG1, mitochondrial | C12orf10 | | 0.975467 | 0.987309 | Q9HB07 | Q9HB07 |
| Nidogen-1 | NID1 | + | 1.56371 | 0.985182 | P14543;P14543-2 | P14543;P14543-2 |
| 60S ribosomal protein L12 | RPL12 | + | 2.70331 | 0.977729 | P30050;P30050-2 | P30050;P30050-2 |
| 40S ribosomal protein S15a | RPS15A | + | 1.80516 | 0.966807 | P62244 | P62244 |
| 60S acidic ribosomal protein P0;60S acidic ribosomal protein P0-like | RPLP0;RPLP0P6 | | 1.03389 | 0.966304 | P05388;P05388-2;Q8NHW5 | P05388;P05388-2;Q8NHW5 |
| Serine/arginine-rich splicing factor 1 | SRSF1 | | 0.843312 | 0.96483 | Q07955;Q07955-3;Q07955-2 | Q07955;Q07955-3;Q07955-2 |
| Cartilage-associated protein | CRTAP | | 1.06709 | 0.955451 | O75718 | O75718 |
| Catechol O-methyltransferase | COMT | | 0.48518 | 0.954522 | P21964-2;P21964 | P21964-2;P21964 |
| Matrilysin | MMP7 | | 0.468191 | 0.953667 | P09237 | P09237 |
| Drebrin-like protein | DBNL | | 1.19552 | 0.950605 | Q9UJU6-2;Q9UJU6-3;Q9UJU6-6;Q9UJU6-4 | Q9UJU6-2;Q9UJU6-3;Q9UJU6-6;Q9UJU6-4 |
| Ribonuclease UK114 | HRSP12 | | 0.278367 | 0.948102 | P52758 | P52758 |
| Agrin;Agrin N-terminal 110 kDa subunit;Agrin C-terminal 110 kDa subunit;Agrin C-terminal 90 kDa fragment;Agr | AGRN | + | 1.44067 | 0.942369 | O00468-6;O00468-7;O00468-3;O00468-5;O00468-4;O00468;O00468 | O00468-6;O00468-7;O00468-3;O00468-5;O00468-4;O004 |
| Toll-interacting protein | TOLLIP | | 1.06704 | 0.94127 | Q9H0E2;Q9H0E2-2 | Q9H0E2 |
| Ras-related protein Rab-5A | RAB5A | | 0.930556 | 0.935305 | P20339-2;P20339 | P20339-2;P20339 |
| Tissue alpha-L-fucosidase | FUCA1 | | 1.19256 | 0.929621 | P04066 | P04066 |
| Sphingomyelin phosphodiesterase | SMPD1 | | 0.79355 | 0.929311 | P17405-3;P17405-2;P17405-4;P17405 | P17405-3;P17405-2;P17405-4;P17405 |
| Specifically androgen-regulated gene protein | SARG | + | 1.675 | 0.927429 | Q9BW04;Q9BW04-2 | Q9BW04;Q9BW04-2 |
| Platelet-derived growth factor C;Platelet-derived growth factor C, latent form;Platelet-derived growth factor C, | PDGFC | + | 1.49379 | 0.924768 | Q9NRA1;Q9NRA1-2;Q9NRA1-4;Q9NRA1-3 | Q9NRA1;Q9NRA1-2;Q9NRA1-4 |
| N(4)-(beta-N-acetylglucosaminyl)-L-asparaginase;Glycosylasparaginase alpha chain;Glycosylasparaginase beta c | AGA | | 0.182163 | 0.920842 | P20933 | P20933 |
| Histidine triad nucleotide-binding protein 1 | HINT1 | | 1.21678 | 0.918433 | P49773 | P49773 |
| Disintegrin and metalloproteinase domain-containing protein 9 | ADAM9 | + | 1.36408 | 0.917042 | Q13443-2;Q13443 | Q13443-2;Q13443 |
| D-3-phosphoglycerate dehydrogenase | PHGDH | + | 1.3719 | 0.916485 | O43175 | O43175 |
| Protein S100-A10 | S100A10 | | 1.10906 | 0.91636 | P60903 | P60903 |
| Sodium-coupled neutral amino acid transporter 1 | SLC38A1 | | 0.690417 | 0.914975 | Q9H2H9 | Q9H2H9 |
| Cathepsin L2 | CTSV | | 0.891749 | 0.896697 | O60911 | O60911 |
| Cysteine-rich motor neuron 1 protein;Processed cysteine-rich motor neuron 1 protein | CRIM1 | + | 1.4853 | 0.896274 | Q9NZV1 | Q9NZV1 |
| Kinesin-like protein KIF23 | KIF23 | | 1.00445 | 0.893436 | Q02241-2;Q02241-3 | Q02241-2;Q02241-3 |
| Semaphorin-3F | SEMA3F | + | 1.70687 | 0.888206 | Q13275 | Q13275 |
| Ephrin type-B receptor 3 | EPHB3 | | 0.953061 | 0.887864 | P54753 | P54753 |
| Membrane-associated progesterone receptor component 1 | PGRMC1 | + | 1.32845 | 0.88661 | O00264;O00264-2 | O00264;O00264-2 |
| Medium-chain specific acyl-CoA dehydrogenase, mitochondrial | ACADM | | 0.2151 | 0.877925 | P11310;P11310-2 | P11310;P11310-2 |
| Peptidyl-glycine alpha-amidating monooxygenase;Peptidylglycine alpha-hydroxylating monooxygenase;Peptidy | PAM | + | 1.40202 | 0.873694 | P19021-4;P19021-3;P19021-6;P19021;P19021-5;P19021-2 | P19021-4;P19021-3;P19021-6;P19021;P19021-5;P19021- |
| 60S ribosomal protein L28 | RPL28 | | 0.606254 | 0.868966 | P46779-4;P46779-5;P46779;P46779-2;P46779-3 | P46779-4;P46779-5;P46779;P46779-2;P46779-3 |
| Out at first protein homolog | OAF | | 1.17659 | 0.863754 | Q86UD1 | Q86UD1 |
| Thrombospondin-1 | THBS1 | + | 1.85956 | 0.859889 | P07996;P07996-2;P35442 | P07996;P07996-2 |
| 10 kDa heat shock protein, mitochondrial | HSPE1 | | 0.654812 | 0.859843 | P61604 | P61604 |
| Scavenger receptor class B member 1 | SCARB1 | + | 2.56439 | 0.858676 | Q8WTV0-4;Q8WTV0-5;Q8WTV0-2;Q8WTV0 | Q8WTV0-4;Q8WTV0-5;Q8WTV0-2;Q8WTV0 |
| Prostatic acid phosphatase;PAPf39 | ACPP | | 0.697283 | 0.852455 | P15309;P15309-2;P15309-3 | P15309;P15309-2;P15309-3 |
| Alanine--tRNA ligase, cytoplasmic | AARS | | 1.16061 | 0.852341 | P49588;P49588-2 | P49588;P49588-2 |
| Galactocerebrosidase | GALC | | 0.388945 | 0.851137 | P54803;P54803-5;P54803-4;P54803-3 | P54803;P54803-5;P54803-4;P54803-3 |
| Actin, cytoplasmic 2;Actin, cytoplasmic 2, N-terminally processed | ACTG1 | + | 2.45128 | 0.847438 | P63261 | P63261 |
| Calsyntenin-3 | CLSTN3 | | 1.13741 | 0.843019 | Q9BQT9;Q9BQT9-2 | Q9BQT9;Q9BQT9-2 |
| 60S acidic ribosomal protein P1 | RPLP1 | | 0.297752 | 0.840542 | P05386 | P05386 |
| Inhibin alpha chain | INHA | | 1.23545 | 0.826862 | P05111 | P05111 |
| 60S ribosomal protein L35a | RPL35A | + | 2.69127 | 0.822754 | P18077 | P18077 |
| Glypican-1;Secreted glypican-1 | GPC1 | + | 1.42464 | 0.821934 | P35052 | P35052 |
| EH domain-containing protein 2 | EHD2 | | 0.379647 | 0.810353 | Q9NZN4;Q9NZN4-2 | Q9NZN4;Q9NZN4-2 |
| Aminopeptidase N | ANPEP | | 1.00037 | 0.810072 | P15144 | P15144 |
| ESF1 homolog | ESF1 | | 0.790558 | 0.808058 | Q9H501 | Q9H501 |
| Cathepsin F | CTSF | | 1.03192 | 0.792982 | Q9UBX1 | Q9UBX1 |
| Choline transporter-like protein 1 | SLC44A1 | | 0.977954 | 0.792798 | Q8WWI5-3;Q8WWI5-2;Q8WWI5 | Q8WWI5-3;Q8WWI5-2;Q8WWI5 |
| Nuclease-sensitive element-binding protein 1 | YBX1 | | 0.859448 | 0.788767 | P67809;Q9Y2T7 | P67809 |
| Ephrin-A5 | EFNA5 | | 0.396214 | 0.786824 | P52803 | P52803 |
| Tubulin beta chain | TUBB | | 0.67187 | 0.783001 | P07437;A6NNZ2 | P07437 |
| Guanine nucleotide-binding protein G(I)/G(S)/G(O) subunit gamma-12 | GNG12 | + | 1.35182 | 0.781808 | Q9UBI6 | Q9UBI6 |
| Ras-related protein R-Ras2 | RRAS2 | | 1.24926 | 0.781384 | P62070;P62070-4;P62070-3;P62070-2 | P62070;P62070-4;P62070-3;P62070-2 |
| Vigilin | HDLBP | | 1.06789 | 0.772038 | Q00341;Q00341-2 | Q00341;Q00341-2 |

| | | | | | | |
|---|---|---|---|---|---|---|
| Deoxyribonuclease-1-like 1 | DNASE1L1 | | 1.15793 | 0.766008 | P49184 | P49184 |
| CD97 antigen;CD97 antigen subunit alpha;CD97 antigen subunit beta | CD97 | | 0.833138 | 0.762168 | P48960;P48960-2;P48960-3;Q9UHX3-4;Q9UHX3-6;Q9UHX3-3;Q9UH | P48960;P48960-2;P48960-3 |
| Copine-3 | CPNE3 | + | 2.93537 | 0.752402 | O75131;Q86YQ8-2;Q9HCH3-2;Q96FN4-2;Q8IYJ1;Q96A23;O95741;Q | O75131 |
| Heterogeneous nuclear ribonucleoprotein D-like | HNRNPDL | + | 1.34391 | 0.747575 | O14979-3;O14979-2;O14979 | O14979-3;O14979-2;O14979 |
| Multiple inositol polyphosphate phosphatase 1 | MINPP1 | + | 2.43116 | 0.746555 | Q9UNW1;Q9UNW1-4;Q9UNW1-3;Q9UNW1-2 | Q9UNW1;Q9UNW1-4 |
| Cytochrome b5 | CYB5A | | 0.303153 | 0.741828 | P00167-2;P00167;P00167-3 | P00167-2;P00167;P00167-3 |
| Biotinidase | BTD | | 0.625783 | 0.741309 | P43251-4;P43251;P43251-3;P43251-2 | P43251-4;P43251;P43251-3;P43251-2 |
| Cochlin | COCH | + | 1.63951 | 0.736099 | O43405;O43405-2 | O43405;O43405-2 |
| Prominin-2 | PROM2 | + | 1.38317 | 0.734939 | Q8N271;Q8N271-2;Q8N271-3 | Q8N271;Q8N271-2 |
| Collagen alpha-3(VI) chain | COL6A3 | | 0.211015 | 0.734273 | P12111-2;P12111;P12111-4;P12111-5;P12111-3 | P12111-2;P12111;P12111-4;P12111-5 |
| Low-density lipoprotein receptor-related protein 8 | LRP8 | | 0.416112 | 0.732301 | Q14114-2;Q14114-3;Q14114;Q14114-4 | Q14114-2;Q14114-3;Q14114;Q14114-4 |
| Low molecular weight phosphotyrosine protein phosphatase | ACP1 | | 0.868379 | 0.732042 | P24666;P24666-3;P24666-4 | P24666;P24666-3;P24666-4 |
| Glyceraldehyde-3-phosphate dehydrogenase | GAPDH | | 0.93717 | 0.726862 | P04406;O14556 | P04406 |
| Small nuclear ribonucleoprotein Sm D2 | SNRPD2 | | 1.13182 | 0.726559 | P62316;P62316-2 | P62316;P62316-2 |
| Nuclear receptor coactivator 5 | NCOA5 | | 1.21795 | 0.722407 | Q9HCD5 | Q9HCD5 |
| N-acetylglucosamine-1-phosphotransferase subunit gamma | GNPTG | | 0.971432 | 0.722364 | Q9UJJ9 | Q9UJJ9 |
| Sushi, von Willebrand factor type A, EGF and pentraxin domain-containing protein 1 | SVEP1 | | 0.500461 | 0.718399 | Q4LDE5;Q4LDE5-4;Q4LDE5-2;Q4LDE5-3 | Q4LDE5;Q4LDE5-4 |
| Desmocollin-3 | DSC3 | | 0.948318 | 0.718005 | Q14574-2;Q14574 | Q14574-2;Q14574 |
| Laminin subunit alpha-5 | LAMA5 | | 1.08735 | 0.713342 | O15230;O15230-2 | O15230 |
| Brain-specific angiogenesis inhibitor 1-associated protein 2 | BAIAP2 | + | 1.4879 | 0.712244 | Q9UQB8-3;Q9UQB8-5;Q9UQB8-4;Q9UQB8-6;Q9UQB8-2;Q9UQB8 | Q9UQB8-3;Q9UQB8-5;Q9UQB8-4;Q9UQB8-6;Q9UQB8-2; |
| GTP-binding nuclear protein Ran | RAN | + | 1.33714 | 0.712137 | P62826 | P62826 |
| 4F2 cell-surface antigen heavy chain | SLC3A2 | + | 2.85913 | 0.710527 | P08195-2 | P08195-2 |
| Guanine nucleotide-binding protein G(i) subunit alpha-2 | GNAI2 | | 1.18485 | 0.704288 | P04899-4;P04899;P04899-6;P04899-3;P04899-5;P04899-2;P09471-2 | P04899-4;P04899;P04899-6;P04899-3;P04899-5;P04899- |
| Extracellular matrix protein 1 | ECM1 | + | 1.30208 | 0.703943 | Q16610;Q16610-4;Q16610-2;Q16610-3 | Q16610;Q16610-4;Q16610-2 |
| Charged multivesicular body protein 3 | CHMP3 | | 0.695766 | 0.703116 | Q9Y3E7;Q9Y3E7-3;Q9Y3E7-2;Q9Y3E7-4 | Q9Y3E7;Q9Y3E7-3;Q9Y3E7-2;Q9Y3E7-4 |
| Prostaglandin E synthase 3 | PTGES3 | | 0.38654 | 0.697091 | Q15185-4;Q15185;Q15185-3;Q15185-2 | Q15185-4;Q15185;Q15185-3 |
| GTP-binding protein SAR1a | SAR1A | | 1.20281 | 0.695621 | Q9NR31;Q9NR31-2 | Q9NR31;Q9NR31-2 |
| Angiotensin-converting enzyme;Angiotensin-converting enzyme, soluble form | ACE | | 0.813739 | 0.693733 | P12821-2;P12821;P12821-4;P12821-3 | P12821-2;P12821;P12821-4;P12821-3 |
| Secreted and transmembrane protein 1 | SECTM1 | | 0.119305 | 0.688817 | Q8WVN6 | Q8WVN6 |
| Unconventional myosin-Ic | MYO1C | | 1.07547 | 0.687992 | O00159;O00159-2;O00159-3 | O00159;O00159-2;O00159-3 |
| Small nuclear ribonucleoprotein-associated proteins B and B;Small nuclear ribonucleoprotein-associated protei | SNRPB;SNRPN | | 0.364712 | 0.687346 | P14678-2;P63162;P14678;P63162-2;P14678-3 | P14678-2;P63162;P14678;P63162-2;P14678-3 |
| Galactosylgalactosylxylosylprotein 3-beta-glucuronosyltransferase 3 | B3GAT3 | | 1.28939 | 0.685775 | O94766-2;O94766 | O94766-2;O94766 |
| Ras-related protein Rab-7a | RAB7A | + | 2.12023 | 0.683753 | P51149 | P51149 |
| Vesicle-associated membrane protein 3 | VAMP3 | | 1.09541 | 0.683476 | Q15836;P63027;P23763-2;P23763-3;P23763 | Q15836 |
| Microtubule-associated protein 4 | MAP4 | | 1.09513 | 0.679576 | P27816;P27816-6;P27816-2;P27816-4 | P27816;P27816-6;P27816-2 |
| Hypoxia up-regulated protein 1 | HYOU1 | | 0.814494 | 0.670403 | Q9Y4L1;Q9Y4L1-2;Q9BXT5 | Q9Y4L1;Q9Y4L1-2 |
| SPARC-related modular calcium-binding protein 1 | SMOC1 | | 0.743317 | 0.66667 | Q9H4F8;Q9H4F8-2 | Q9H4F8;Q9H4F8-2 |
| Serine/threonine-protein phosphatase 2A catalytic subunit beta isoform | PPP2CB | | 0.278666 | 0.665106 | P62714 | P62714 |
| Guanine nucleotide-binding protein subunit alpha-11 | GNA11 | + | 1.53264 | 0.664857 | P29992 | P29992 |
| Interleukin-1 receptor accessory protein | IL1RAP | | 1.08967 | 0.663954 | Q9NPH3-2;Q9NPH3;Q9NPH3-5;Q9NPH3-3 | Q9NPH3-2;Q9NPH3;Q9NPH3-5;Q9NPH3-3 |
| Carbohydrate sulfotransferase 14 | CHST14 | | 1.21778 | 0.655191 | Q8NCH0 | Q8NCH0 |
| Beta-1,4-glucuronyltransferase 1 | B4GAT1 | + | 1.38023 | 0.654705 | O43505 | O43505 |
| Heat shock cognate 71 kDa protein | HSPA8 | | 1.11292 | 0.651967 | P11142;P11142-2 | P11142;P11142-2 |
| Semaphorin-3B | SEMA3B | | 1.05491 | 0.651698 | Q13214-2;Q13214 | Q13214-2;Q13214 |
| Guanine nucleotide-binding protein G(q) subunit alpha | GNAQ | | 0.791043 | 0.649918 | P50148 | P50148 |
| Immunoglobulin superfamily member 8 | IGSF8 | + | 1.96962 | 0.647927 | Q969P0;Q969P0-3;Q969P0-2 | Q969P0;Q969P0-3 |
| Myosin regulatory light chain 12A;Myosin regulatory light chain 12B;Myosin regulatory light polypeptide 9 | MYL12A;MYL12B;MYL9 | | 1.29201 | 0.64459 | P19105;O14950;P24844;P24844-2 | P19105;O14950;P24844 |
| Tubulin beta-2A chain | TUBB2A | | 0.663801 | 0.640981 | Q13885 | Q13885 |
| Platelet-activating factor acetylhydrolase IB subunit beta | PAFAH1B2 | | 1.09595 | 0.638561 | P68402;P68402-3;P68402-2;P68402-4 | P68402;P68402-3;P68402-2;P68402-4 |
| Di-N-acetylchitobiase | CTBS | | 0.495399 | 0.63294 | Q01459 | Q01459 |
| Transforming protein RhoA | RHOA | | 0.295103 | 0.629893 | P61586 | P61586 |
| Ras-related protein Rap-1A | RAP1A | | 1.20145 | 0.626631 | P62834 | P62834 |
| Vacuolar-sorting protein SNF8 | SNF8 | | 0.801317 | 0.624791 | Q96H20-2;Q96H20 | Q96H20-2;Q96H20 |
| Golgi membrane protein 1 | GOLM1 | | 1.19859 | 0.623992 | Q8NBJ4-2;Q8NBJ4 | Q8NBJ4-2;Q8NBJ4 |
| Histone H1.5 | HIST1H1B | + | 1.32332 | 0.622911 | P16401 | P16401 |
| Guanine nucleotide-binding protein G(I)/G(S)/G(T) subunit beta-2 | GNB2 | + | 1.34776 | 0.622672 | P62879;P62879-2 | P62879 |
| Phospholipid scramblase 1 | PLSCR1 | | 0.548121 | 0.6219 | O15162-2;O15162 | O15162-2;O15162 |
| RNA-binding motif protein, X chromosome;RNA-binding motif protein, X chromosome, N-terminally processed; | RBMX;RBMXL1 | | 0.791365 | 0.621355 | P38159;P38159-2;Q96E39;P38159-3;Q8N7X1 | P38159;P38159-2;Q96E39;P38159-3 |
| Extracellular sulfatase Sulf-2 | SULF2 | | 0.226514 | 0.620626 | Q8IWU5-2;Q8IWU5 | Q8IWU5-2;Q8IWU5 |
| Tubulin alpha-1B chain | TUBA1B | | 0.795145 | 0.615896 | P68363;P68363-2 | P68363;P68363-2 |
| Retinoid-inducible serine carboxypeptidase | SCPEP1 | + | 1.95044 | 0.61308 | Q9HB40;Q9HB40-2 | Q9HB40;Q9HB40-2 |
| TBC1 domain family member 4 | TBC1D4 | | 0.245255 | 0.609925 | O60343-2;O60343-3;O60343;O60343-5 | O60343-2;O60343-3;O60343 |
| 2,3-cyclic-nucleotide 3-phosphodiesterase | CNP | + | 1.83609 | 0.605516 | P09543-2;P09543 | P09543-2;P09543 |
| Mesothelin;Megakaryocyte-potentiating factor;Mesothelin, cleaved form | MSLN | | 0.892653 | 0.603085 | Q13421-4;Q13421-3;Q13421-2;Q13421 | Q13421-4;Q13421-3;Q13421-2;Q13421 |
| Hsc70-interacting protein;Putative protein FAM10A4 | ST13;ST13P4 | | 0.542252 | 0.602049 | P50502;Q8IZP2;Q8NFI4 | P50502;Q8IZP2 |
| Ubiquitin-40S ribosomal protein S27a;Ubiquitin;40S ribosomal protein S27a;Ubiquitin-60S ribosomal protein L4 | RPS27A;UBA52;UBB;UBC | + | 2.29881 | 0.601476 | P62979;P62987;P0CG47;P0CG48 | P62979;P62987;P0CG47;P0CG48 |
| Tripeptidyl-peptidase 1 | TPP1 | | 1.0027 | 0.596851 | O14773;O14773-2 | O14773;O14773-2 |
| Catenin delta-1 | CTNND1 | + | 1.72493 | 0.596547 | O60716-21;O60716-19;O60716-18;O60716-17;O60716-13;O60716-1 | O60716-21;O60716-19;O60716-18;O60716-17;O60716-1 |
| Lysosomal acid lipase/cholesteryl ester hydrolase | LIPA | | 0.531412 | 0.595415 | P38571-2;P38571 | P38571-2;P38571 |
| Guanine nucleotide-binding protein G(I)/G(S)/G(T) subunit beta-1 | GNB1 | | 1.24105 | 0.595069 | P62873;P62873-2;P16520-2;P16520 | P62873;P62873-2 |
| Stanniocalcin-1 | STC1 | | 0.389288 | 0.585224 | P52823;P52823-2 | P52823;P52823-2 |
| Ras-related protein Rap-2b | RAP2B | | 1.10899 | 0.585089 | P61225 | P61225 |
| Interleukin-18 | IL18 | | 1.00097 | 0.583324 | Q14116;Q14116-2 | Q14116;Q14116-2 |
| BTB/POZ domain-containing protein KCTD12 | KCTD12 | | 1.2776 | 0.582349 | Q96CX2;Q68DU8 | Q96CX2 |
| Epithelial cell adhesion molecule | EPCAM | + | 1.42003 | 0.574908 | P16422 | P16422 |
| Ceroid-lipofuscinosis neuronal protein 5 | CLN5 | + | 1.48603 | 0.57415 | O75503 | O75503 |
| Peroxiredoxin-5, mitochondrial | PRDX5 | | 0.90193 | 0.571432 | P30044-2;P30044;P30044-3;P30044-4 | P30044-2;P30044;P30044-3;P30044-4 |
| Ras-related protein Rab-2A | RAB2A | | 1.1642 | 0.567778 | P61019;P61019-2 | P61019;P61019-2 |
| Cystathionine gamma-lyase | CTH | | 1.17581 | 0.56744 | P32929;P32929-3;P32929-2 | P32929;P32929-3;P32929-2 |
| 60S ribosomal protein L11 | RPL11 | | 1.10202 | 0.565161 | P62913;P62913-2 | P62913;P62913-2 |

| | | | | | | |
|---|---|---|---|---|---|---|
| Amyloid-like protein 2 | APLP2 | + | 1.53459 | 0.562834 | Q06481;Q06481-3;Q06481-6;Q06481-2;Q06481-4;Q06481-5 | Q06481;Q06481-3;Q06481-6;Q06481-2;Q06481-4;Q0648 |
| Integrin beta-5 | ITGB5 | | 1.13209 | 0.559928 | P18084 | P18084 |
| Alpha-galactosidase A | GLA | + | 1.35476 | 0.5567 | P06280 | P06280 |
| Cell division control protein 42 homolog | CDC42 | | 0.285101 | 0.554398 | P60953;P60953-1 | P60953;P60953-1 |
| Ribonucleases P/MRP protein subunit POP1 | POP1 | | 0.481975 | 0.552158 | Q99575 | Q99575 |
| Beta-centractin | ACTR1B | | 0.480835 | 0.551655 | P42025 | P42025 |
| Neuronal growth regulator 1 | NEGR1 | | 0.142114 | 0.551594 | Q7Z3B1;Q7Z3B1-2 | Q7Z3B1;Q7Z3B1-2 |
| Slit homolog 2 protein;Slit homolog 2 protein N-product;Slit homolog 2 protein C-product | SLIT2 | + | 1.53591 | 0.551544 | O94813-3;O94813-2;O94813 | O94813-3;O94813-2;O94813 |
| Dickkopf-related protein 1 | DKK1 | | 0.24274 | 0.549509 | O94907;Q9UBU2 | O94907 |
| Rho GDP-dissociation inhibitor 1 | ARHGDIA | | 0.671968 | 0.54824 | P52565;P52565-2 | P52565;P52565-2 |
| Na(+)/H(+) exchange regulatory cofactor NHE-RF1 | SLC9A3R1 | | 0.250111 | 0.547192 | O14745;O14745-2 | O14745;O14745-2 |
| Cadherin-13 | CDH13 | | 0.43954 | 0.544096 | P55290;P55290-4;P55290-3;P55290-2;P55290-5 | P55290;P55290-4;P55290-3;P55290-2;P55290-5 |
| Ras-related protein Rab-8A | RAB8A | | 0.520468 | 0.540238 | P61006;P61006-2 | P61006;P61006-2 |
| 60S ribosomal protein L23 | RPL23 | | 0.588597 | 0.53758 | P62829 | P62829 |
| Ras-related C3 botulinum toxin substrate 1;Ras-related C3 botulinum toxin substrate 3 | RAC1;RAC3 | | 1.07755 | 0.53441 | P63000;P63000-2;P60763 | P63000;P63000-2;P60763 |
| V-type proton ATPase subunit G 1 | ATP6V1G1 | | 0.156025 | 0.53363 | O75348;O95670-2;O95670 | O75348 |
| Tsukushin | TSKU | + | 3.99384 | 0.532166 | Q8WUA8 | Q8WUA8 |
| Glutathione peroxidase 1 | GPX1 | | 0.121763 | 0.532145 | P07203;P07203-2 | P07203;P07203-2 |
| Sushi repeat-containing protein SRPX2 | SRPX2 | + | 1.35474 | 0.530633 | O60687 | O60687 |
| E3 ubiquitin-protein ligase Itchy homolog | ITCH | + | 1.46485 | 0.529387 | Q96J02-2;Q96J02;Q96J02-3;Q9H0M0-6;Q9H0M0-3;Q9H0M0 | Q96J02-2;Q96J02;Q96J02-3 |
| 3-mercaptopyruvate sulfurtransferase | MPST | | 0.19024 | 0.528825 | P25325;P25325-2 | P25325;P25325-2 |
| Voltage-dependent calcium channel subunit alpha-2/delta-1;Voltage-dependent calcium channel subunit alpha | CACNA2D1 | | 0.156755 | 0.528313 | P54289-5;P54289;P54289-4;P54289-2;P54289-3 | P54289-5;P54289;P54289-4;P54289-2;P54289-3 |
| Collagen alpha-2(IV) chain;Canstatin | COL4A2 | | 0.606387 | 0.526088 | P08572 | P08572 |
| Microfibrillar-associated protein 2 | MFAP2 | | 0.42975 | 0.525621 | P55001-3;P55001-2;P55001 | P55001-3;P55001-2;P55001 |
| Heat shock 70 kDa protein 13 | HSPA13 | | 1.18713 | 0.524949 | P48723 | P48723 |
| EH domain-containing protein 4 | EHD4 | | 0.743232 | 0.524766 | Q9H223 | Q9H223 |
| Ras-related protein Rab-34 | RAB34 | | 1.13358 | 0.524054 | Q9BZG1-4;Q9BZG1;Q9BZG1-2 | Q9BZG1-4;Q9BZG1;Q9BZG1-2 |
| Ras-related protein Rab-10 | RAB10 | | 0.31885 | 0.521789 | P61026 | P61026 |
| Ras-related protein Rab-18 | RAB18 | + | 1.56673 | 0.5212 | Q9NP72;Q9NP72-2;Q9NP72-3 | Q9NP72;Q9NP72-2;Q9NP72-3 |
| ADP/ATP translocase 2;ADP/ATP translocase 2, N-terminally processed | SLC25A5 | | 0.133698 | 0.518997 | P05141;Q9H0C2 | P05141 |
| Cathepsin D;Cathepsin D light chain;Cathepsin D heavy chain | CTSD | | 0.624537 | 0.516768 | P07339 | P07339 |
| Transmembrane 9 superfamily member 2 | TM9SF2 | | 1.11132 | 0.516155 | Q99805 | Q99805 |
| Deoxyuridine 5-triphosphate nucleotidohydrolase, mitochondrial | DUT | | 1.24446 | 0.513254 | P33316-2 | P33316-2 |
| Sulfhydryl oxidase 2 | QSOX2 | | 0.323526 | 0.511272 | Q6ZRP7 | Q6ZRP7 |
| Cytochrome b-c1 complex subunit 2, mitochondrial | UQCRC2 | | 0.27367 | 0.505379 | P22695 | P22695 |
| Ethylmalonyl-CoA decarboxylase | ECHDC1 | | 0.598174 | 0.503607 | Q9NTX5-6;Q9NTX5-2;Q9NTX5;Q9NTX5-3;Q9NTX5-5;Q9NTX5-4 | Q9NTX5-6;Q9NTX5-2;Q9NTX5;Q9NTX5-3 |
| Vacuolar protein sorting-associated protein 37B | VPS37B | | 1.23481 | 0.501122 | Q9H9H4 | Q9H9H4 |
| Leucine-rich repeat-containing protein 57 | LRRC57 | | 1.05775 | 0.501006 | Q8N9N7 | Q8N9N7 |
| Attractin | ATRN | | 0.647934 | 0.498952 | O75882-3;O75882-2;O75882 | O75882-3;O75882-2;O75882 |
| Activated RNA polymerase II transcriptional coactivator p15 | SUB1 | + | 1.32851 | 0.498196 | P53999 | P53999 |
| Heterogeneous nuclear ribonucleoprotein M | HNRNPM | | 0.734071 | 0.495966 | P52272-2;P52272 | P52272-2;P52272 |
| Endophilin-A2 | SH3GL1 | | 0.47808 | 0.489436 | Q99961;Q99961-3;Q99961-2;Q99962 | Q99961;Q99961-3;Q99961-2 |
| Thyroid hormone receptor-associated protein 3 | THRAP3 | | 0.457097 | 0.485825 | Q9Y2W1 | Q9Y2W1 |
| Lysosomal Pro-X carboxypeptidase | PRCP | | 0.523563 | 0.484334 | P42785;P42785-2 | P42785;P42785-2 |
| Annexin A5 | ANXA5 | | 0.973664 | 0.484 | P08758 | P08758 |
| DnaJ homolog subfamily C member 7 | DNAJC7 | | 1.23918 | 0.482401 | Q99615;Q99615-2 | Q99615;Q99615-2 |
| Hyaluronan and proteoglycan link protein 3 | HAPLN3 | | 1.06565 | 0.481927 | Q96S86;Q86UW8 | Q96S86 |
| Laminin subunit alpha-4 | LAMA4 | + | 1.85322 | 0.477543 | Q16363-2;Q16363;Q16363-3 | Q16363-2;Q16363 |
| Ras-related protein Rab-5C | RAB5C | | 1.09568 | 0.475718 | P51148;P51148-2 | P51148;P51148-2 |
| Signal recognition particle 14 kDa protein | SRP14 | | 0.654149 | 0.473392 | P37108 | P37108 |
| Ras-related protein Rab-1B | RAB1B | | 0.533452 | 0.471816 | Q9H0U4;Q92928;P59190-2;P59190 | Q9H0U4 |
| Acidic leucine-rich nuclear phosphoprotein 32 family member B | ANP32B | | 0.36468 | 0.47173 | Q92688-2;Q92688 | Q92688-2;Q92688 |
| pre-mRNA 3 end processing protein WDR33 | WDR33 | | 0.237551 | 0.46952 | Q9C0J8 | Q9C0J8 |
| Ubiquitin-associated domain-containing protein 1 | UBAC1 | | 0.452227 | 0.467112 | Q9BSL1 | Q9BSL1 |
| 5-AMP-activated protein kinase subunit gamma-1 | PRKAG1 | | 0.201125 | 0.466297 | P54619-2;P54619;P54619-3 | P54619-2;P54619;P54619-3 |
| Early endosome antigen 1 | EEA1 | + | 1.98305 | 0.465885 | Q15075 | Q15075 |
| Vascular endothelial growth factor C | VEGFC | | 0.838565 | 0.46486 | P49767 | P49767 |
| Exostosin-like 2;Processed exostosin-like 2 | EXTL2 | | 0.515108 | 0.464602 | Q9UBQ6 | Q9UBQ6 |
| Junctional adhesion molecule A | F11R | | 1.08619 | 0.462921 | Q9Y624;Q9Y624-2 | Q9Y624;Q9Y624-2 |
| Vesicular integral-membrane protein VIP36 | LMAN2 | | 1.07361 | 0.460894 | Q12907 | Q12907 |
| Serine/threonine-protein kinase 26 | STK26 | | 0.178578 | 0.453911 | Q9P289;Q9P289-3;Q9P289-2;O00506-3;O00506-2 | Q9P289;Q9P289-3;Q9P289-2 |
| Protein TFG | TFG | + | 1.42267 | 0.451517 | Q92734-4;Q92734-3;Q92734-2;Q92734 | Q92734-4;Q92734-3;Q92734-2;Q92734 |
| UDP-glucuronic acid decarboxylase 1 | UXS1 | + | 1.53068 | 0.449612 | Q8NBZ7;Q8NBZ7-2;Q8NBZ7-3 | Q8NBZ7;Q8NBZ7-2;Q8NBZ7-3 |
| Lysosomal alpha-glucosidase;76 kDa lysosomal alpha-glucosidase;70 kDa lysosomal alpha-glucosidase | GAA | | 0.725413 | 0.447122 | P10253 | P10253 |
| L-aminoadipate-semialdehyde dehydrogenase-phosphopantetheinyl transferase | AASDHPPT | | 0.38266 | 0.446108 | Q9NRN7;Q9NRN7-2 | Q9NRN7 |
| 60 kDa heat shock protein, mitochondrial | HSPD1 | | 0.804176 | 0.443909 | P10809;P10809-2 | P10809 |
| Plasma membrane calcium-transporting ATPase 1 | ATP2B1 | | 0.702674 | 0.442226 | P20020-6;P20020-3;P20020-4;P20020;P20020-5;P20020-2;Q01814-( | P20020-6;P20020-3;P20020-4;P20020;P20020-5;P20020- |
| Receptor-type tyrosine-protein phosphatase S | PTPRS | | 0.711774 | 0.441515 | Q13332-6;Q13332-2;Q13332-3;Q13332-4;Q13332;Q13332-7;Q1333 | Q13332-6;Q13332-2;Q13332-3;Q13332-4;Q13332;Q1333 |
| Testican-2 | SPOCK2 | | 0.152198 | 0.43903 | Q92563;Q92563-2 | Q92563 |
| Cystatin-C | CST3 | | 1.0089 | 0.435443 | P01034 | P01034 |
| Ribose-phosphate pyrophosphokinase 2 | PRPS2 | | 0.22334 | 0.434012 | P11908;P11908-2 | P11908;P11908-2 |
| Annexin A1 | ANXA1 | | 0.512524 | 0.430728 | P04083 | P04083 |
| Pro-cathepsin H;Cathepsin H mini chain;Cathepsin H;Cathepsin H heavy chain;Cathepsin H light chain | CTSH | | 0.690497 | 0.430039 | P09668 | P09668 |
| Brain acid soluble protein 1 | BASP1 | | 0.250665 | 0.427827 | P80723;P80723-2 | P80723;P80723-2 |
| Polypeptide N-acetylgalactosaminyltransferase 2;Polypeptide N-acetylgalactosaminyltransferase 2 soluble form | GALNT2 | | 0.970905 | 0.426908 | Q10471;Q10471-2 | Q10471 |
| Zinc-alpha-2-glycoprotein | AZGP1 | | 0.131081 | 0.4262 | P25311 | P25311 |
| Lysyl oxidase homolog 2 | LOXL2 | | 0.588881 | 0.423677 | Q9Y4K0 | Q9Y4K0 |
| Eukaryotic translation initiation factor 3 subunit I | EIF3I | | 0.821396 | 0.421582 | Q13347 | Q13347 |
| 60S ribosomal protein L31 | RPL31 | | 0.585583 | 0.421282 | P62899;P62899-3;P62899-2 | P62899;P62899-3;P62899-2 |
| Histone H3.1 | HIST1H3A | | 0.328402 | 0.420103 | P68431 | P68431 |

| | | | | | | |
|---|---|---|---|---|---|---|
| Tropomyosin alpha-3 chain | TPM3 | + | 2.27941 | 0.419803 | P06753-2;P06753-5;P06753-7 | P06753-2;P06753-5 |
| Heterogeneous nuclear ribonucleoproteins C1/C2 | HNRNPC | + | 1.8707 | 0.418627 | P07910-2;P07910;P07910-4;B2RXH8 | P07910-2;P07910;P07910-4 |
| 14-3-3 protein sigma | SFN | | 0.450453 | 0.417952 | P31947;P31947-2 | P31947;P31947-2 |
| CD2-associated protein | CD2AP | | 1.27456 | 0.416094 | Q9Y5K6 | Q9Y5K6 |
| Basigin | BSG | | 1.29359 | 0.416065 | P35613-2;P35613;P35613-4 | P35613-2;P35613;P35613-4 |
| Basement membrane-specific heparan sulfate proteoglycan core protein;Endorepellin;LG3 peptide | HSPG2 | | 0.460854 | 0.415011 | P98160 | P98160 |
| Beta-1,3-galactosyl-O-glycosyl-glycoprotein beta-1,6-N-acetylglucosaminyltransferase | GCNT1 | | 0.955167 | 0.414658 | Q02742 | Q02742 |
| 14-3-3 protein epsilon | YWHAE | + | 1.53085 | 0.41313 | P62258;P62258-2 | P62258;P62258-2 |
| Protein FAM3C | FAM3C | | 0.711871 | 0.409992 | Q92520 | Q92520 |
| Glucosylceramidase | GBA | | 1.07769 | 0.408901 | P04062-4;P04062-2;P04062;P04062-5 | P04062-4;P04062-2;P04062;P04062-5 |
| CD81 antigen | CD81 | | 0.940219 | 0.405912 | P60033 | P60033 |
| Exostosin-2 | EXT2 | + | 1.33284 | 0.399758 | Q93063;Q93063-3;Q93063-2 | Q93063;Q93063-3;Q93063-2 |
| Fibroblast growth factor receptor 4 | FGFR4 | | 1.05676 | 0.397097 | P22455-2;P22455;P22455-3 | P22455-2;P22455;P22455-3 |
| Neuroserpin | SERPINI1 | | 0.928944 | 0.397024 | Q99574 | Q99574 |
| Ribonuclease 4 | RNASE4 | | 0.370642 | 0.39596 | P34096 | P34096 |
| Desmoglein-2 | DSG2 | | 1.24571 | 0.392988 | Q14126 | Q14126 |
| Integrin alpha-V;Integrin alpha-V heavy chain;Integrin alpha-V light chain | ITGAV | | 1.20005 | 0.390507 | P06756-3;P06756-2;P06756 | P06756-3;P06756-2;P06756 |
| Protein ERGIC-53 | LMAN1 | | 0.0730643 | 0.385923 | P49257 | P49257 |
| Myosin-10 | MYH10 | | 0.348564 | 0.38419 | P35580;P35580-5;P35580-2;P35580-3;P35580-4 | P35580;P35580-5;P35580-2;P35580-3;P35580-4 |
| Acyl-coenzyme A thioesterase 13;Acyl-coenzyme A thioesterase 13, N-terminally processed | ACOT13 | | 0.4484 | 0.381991 | Q9NPJ3;Q9NPJ3-2 | Q9NPJ3;Q9NPJ3-2 |
| Integrin alpha-3;Integrin alpha-3 heavy chain;Integrin alpha-3 light chain | ITGA3 | | 0.239018 | 0.381604 | P26006;P26006-1 | P26006;P26006-1 |
| Adenine phosphoribosyltransferase | APRT | | 0.211438 | 0.381079 | P07741;P07741-2 | P07741;P07741-2 |
| HLA class I histocompatibility antigen, A-68 alpha chain;HLA class I histocompatibility antigen, A-69 alpha chain; | HLA-A | | 0.66394 | 0.380684 | P01891;P10316;P01892 | P01891;P10316;P01892 |
| CD109 antigen | CD109 | | 0.843147 | 0.380519 | Q6YHK3;Q6YHK3-4;Q6YHK3-2;Q6YHK3-3 | Q6YHK3;Q6YHK3-4;Q6YHK3-2;Q6YHK3-3 |
| Wiskott-Aldrich syndrome protein family member 2 | WASF2 | | 0.397936 | 0.379501 | Q9Y6W5;Q9Y6W5-2 | Q9Y6W5;Q9Y6W5-2 |
| Beta-hexosaminidase subunit alpha | HEXA | | 0.353315 | 0.378465 | P06865;P06865-2 | P06865 |
| 60S ribosomal protein L23a | RPL23A | | 0.437751 | 0.378327 | P62750 | P62750 |
| Activity-dependent neuroprotector homeobox protein | ADNP | | 0.340527 | 0.378191 | Q9H2P0 | Q9H2P0 |
| Beta-1,3-N-acetylglucosaminyltransferase lunatic fringe | LFNG | | 0.368839 | 0.378121 | Q8NES3;Q8NES3-3;Q8NES3-2;Q8NES3-4 | Q8NES3;Q8NES3-3;Q8NES3-2;Q8NES3-4 |
| Laminin subunit beta-1 | LAMB1 | + | 1.87801 | 0.377888 | P07942 | P07942 |
| Perilipin-2 | PLIN2 | | 0.408283 | 0.377842 | Q99541 | Q99541 |
| Disintegrin and metalloproteinase domain-containing protein 10 | ADAM10 | | 0.331241 | 0.375726 | O14672;O14672-2 | O14672 |
| Monocarboxylate transporter 1 | SLC16A1 | | 0.599425 | 0.373586 | P53985;P53985-2 | P53985;P53985-2 |
| Carboxypeptidase A4 | CPA4 | | 0.590226 | 0.37288 | Q9UI42;Q9UI42-2 | Q9UI42;Q9UI42-2 |
| Flotillin-1 | FLOT1 | | 0.197063 | 0.369479 | O75955;O75955-2 | O75955;O75955-2 |
| Liprin-beta-1 | PPFIBP1 | | 0.276661 | 0.369446 | Q86W92-4;Q86W92-2;Q86W92;Q86W92-3;Q86W92-5;Q86ND30-3;C | Q86W92-4;Q86W92-2;Q86W92;Q86W92-3 |
| 60S ribosomal protein L3 | RPL3 | | 0.301047 | 0.36233 | P39023;Q92901 | P39023 |
| Spermine synthase | SMS | | 0.906314 | 0.359696 | P52788;P52788-2 | P52788;P52788-2 |
| Laminin subunit gamma-1 | LAMC1 | | 1.02057 | 0.358428 | P11047 | P11047 |
| Carboxypeptidase E | CPE | | 0.242078 | 0.356491 | P16870-2;P16870 | P16870-2;P16870 |
| Collagen alpha-1(V) chain | COL5A1 | | 0.461025 | 0.35587 | P20908 | P20908 |
| 40S ribosomal protein S20 | RPS20 | | 0.240639 | 0.355433 | P60866;P60866-2 | P60866;P60866-2 |
| Plasma alpha-L-fucosidase | FUCA2 | | 0.238598 | 0.350957 | Q9BTY2;Q9BTY2-2 | Q9BTY2 |
| Nucleolin | NCL | | 0.511675 | 0.350502 | P19338 | P19338 |
| Ras-related protein Rab-35 | RAB35 | | 0.348124 | 0.349195 | Q15286;Q15286-2 | Q15286;Q15286-2 |
| Liprin-alpha-1 | PPFIA1 | | 0.381419 | 0.347407 | Q13136-2;Q13136;O75334-8;O75335;O75335-2;O75335-1 | Q13136-2;Q13136 |
| Epidermal growth factor receptor | EGFR | | 0.836747 | 0.344268 | P00533;P00533-4;P00533-3;P00533-2;P04626-3;P04626-2;P04626-6 | P00533;P00533-4;P00533-3;P00533-2 |
| Protein S100-A11;Protein S100-A11, N-terminally processed | S100A11 | | 0.0740513 | 0.343428 | P31949 | P31949 |
| Ribonucleoside-diphosphate reductase large subunit | RRM1 | | 0.447972 | 0.343397 | P23921 | P23921 |
| Vacuolar protein sorting-associated protein 4B | VPS4B | | 0.185216 | 0.339942 | O75351 | O75351 |
| Growth arrest-specific protein 6 | GAS6 | | 0.465138 | 0.337636 | Q14393-2;Q14393-3;Q14393;Q14393-5;Q14393-4 | Q14393-2;Q14393-3;Q14393;Q14393-5;Q14393-4 |
| Small nuclear ribonucleoprotein Sm D3 | SNRPD3 | | 0.174408 | 0.33638 | P62318-2;P62318 | P62318-2;P62318 |
| Septin-2 | SEPT2 | | 0.464727 | 0.335874 | Q15019;Q15019-3;Q15019-2 | Q15019;Q15019-3;Q15019-2 |
| Dystroglycan;Alpha-dystroglycan;Beta-dystroglycan | DAG1 | | 0.384047 | 0.334189 | Q14118 | Q14118 |
| Casein kinase II subunit beta | CSNK2B | | 0.317767 | 0.333699 | P67870 | P67870 |
| THO complex subunit 4 | ALYREF | | 0.535216 | 0.332042 | Q86V81 | Q86V81 |
| 14-3-3 protein gamma;14-3-3 protein gamma, N-terminally processed | YWHAG | | 0.35186 | 0.331355 | P61981 | P61981 |
| FAD-linked sulfhydryl oxidase ALR | GFER | | 0.169907 | 0.327251 | P55789;P55789-2 | P55789;P55789-2 |
| Semaphorin-4C | SEMA4C | | 0.175375 | 0.326539 | Q9C0C4 | Q9C0C4 |
| Protein NDRG3 | NDRG3 | | 0.222731 | 0.322659 | Q9UGV2-3;Q9UGV2-2;Q9UGV2 | Q9UGV2-3;Q9UGV2-2;Q9UGV2 |
| Prolyl 3-hydroxylase 1 | LEPRE1 | | 0.287564 | 0.322451 | Q32P28-4;Q32P28;Q32P28-3;Q32P28-2 | Q32P28-4;Q32P28;Q32P28-3 |
| Collagen alpha-1(XVIII) chain;Endostatin | COL18A1 | | 0.230006 | 0.322362 | P39060-2;P39060-1;P39060 | P39060-2;P39060-1;P39060 |
| Eukaryotic translation initiation factor 6 | EIF6 | | 0.289848 | 0.320943 | P56537;P56537-2 | P56537;P56537-2 |
| Serine--tRNA ligase, cytoplasmic | SARS | | 0.661734 | 0.319481 | P49591 | P49591 |
| Choline transporter-like protein 2 | SLC44A2 | | 0.41178 | 0.317801 | Q8IWA5;Q8IWA5-3;Q8IWA5-2 | Q8IWA5;Q8IWA5-3;Q8IWA5-2 |
| Neural Wiskott-Aldrich syndrome protein | WASL | | 0.733123 | 0.314313 | O00401 | O00401 |
| Phosphoenolpyruvate carboxykinase [GTP], mitochondrial | PCK2 | | 0.836953 | 0.314276 | Q16822;Q16822-2;P35558;P35558-2 | Q16822 |
| Cullin-4B | CUL4B | | 0.383503 | 0.31372 | Q13620-1;Q13620;Q13620-3 | Q13620-1;Q13620;Q13620-3 |
| Prohibitin | PHB | | 0.189298 | 0.312202 | P35232;P35232-2 | P35232 |
| Nicastrin | NCSTN | | 0.653329 | 0.310422 | Q92542-2;Q92542 | Q92542-2;Q92542 |
| Vesicle-associated membrane protein 7 | VAMP7 | | 0.444202 | 0.309407 | P51809;P51809-3;P51809-2 | P51809;P51809-3;P51809-2 |
| Neudesin | NENF | | 0.0588943 | 0.306525 | Q9UMX5 | Q9UMX5 |
| Protein FAM3A | FAM3A | | 0.211759 | 0.301637 | P98173-3;P98173;P98173-2 | P98173-3;P98173;P98173-2 |
| Copine-1 | CPNE1 | | 0.465026 | 0.299199 | Q99829 | Q99829 |
| Core histone macro-H2A.1 | H2AFY | | 0.324809 | 0.297087 | O75367-3;O75367;O75367-2;Q9P0M6 | O75367-3;O75367;O75367-2 |
| Lysosomal protective protein;Lysosomal protective protein 32 kDa chain;Lysosomal protective protein 20 kDa c | CTSA | | 0.338368 | 0.296038 | P10619-2;P10619 | P10619-2;P10619 |
| Peptidyl-prolyl cis-trans isomerase FKBP5;Peptidyl-prolyl cis-trans isomerase FKBP5, N-terminally processed | FKBP5 | | 0.581484 | 0.294635 | Q13451;Q13451-2 | Q13451 |
| RuvB-like 2 | RUVBL2 | | 0.939958 | 0.293062 | Q9Y230;Q9Y230-2 | Q9Y230;Q9Y230-2 |
| Lactoylglutathione lyase | GLO1 | | 0.156835 | 0.29155 | Q04760;Q04760-2 | Q04760;Q04760-2 |
| Proline synthase co-transcribed bacterial homolog protein | PROSC | | 0.253546 | 0.290252 | O94903 | O94903 |

| | | | | | | |
|---|---|---|---|---|---|---|
| Protein delta homolog 2 | DLK2 | | 0.145228 | 0.289474 | Q6UY11;Q6UY11-2 | Q6UY11;Q6UY11-2 |
| Mitogen-activated protein kinase 3 | MAPK3 | | 0.0977661 | 0.288752 | P27361;P27361-2;P27361-3;P31152;Q16659 | P27361;P27361-2;P27361-3 |
| Myosin light chain 6B | MYL6B | | 0.16213 | 0.285799 | P14649 | P14649 |
| Protein S100-P | S100P | | 0.199597 | 0.284824 | P25815 | P25815 |
| Myotubularin | MTM1 | | 0.568385 | 0.283435 | Q13496;Q13496-2 | Q13496;Q13496-2 |
| Cathepsin L1;Cathepsin L1 heavy chain;Cathepsin L1 light chain | CTSL | | 0.196039 | 0.283063 | P07711 | P07711 |
| Heat shock protein 105 kDa | HSPH1 | | 0.997467 | 0.282816 | Q92598-2 | Q92598-2 |
| Rho-related GTP-binding protein RhoC | RHOC | | 0.233147 | 0.279313 | P08134 | P08134 |
| Prohibitin-2 | PHB2 | | 0.177231 | 0.27896 | Q99623;Q99623-2 | Q99623;Q99623-2 |
| Thioredoxin domain-containing protein 17 | TXNDC17 | | 0.486334 | 0.27755 | Q9BRA2 | Q9BRA2 |
| UDP-glucose:glycoprotein glucosyltransferase 1 | UGGT1 | | 0.585184 | 0.27608 | Q9NYU2-2;Q9NYU2 | Q9NYU2-2;Q9NYU2 |
| Choriogonadotropin subunit beta variant 1;Choriogonadotropin subunit beta;Choriogonadotropin subunit beta | CGB1;CGB;CGB2 | | 0.118167 | 0.274721 | A6NKQ9-2;P01233;P01233-2;A6NKQ9;Q6NT52 | A6NKQ9-2;P01233;P01233-2;A6NKQ9;Q6NT52 |
| Inorganic pyrophosphatase | PPA1 | | 0.313212 | 0.274356 | Q15181 | Q15181 |
| Multifunctional methyltransferase subunit TRM112-like protein | TRMT112 | | 0.871714 | 0.272526 | Q9UI30;Q9UI30-2 | Q9UI30;Q9UI30-2 |
| Bone marrow stromal antigen 2 | BST2 | | 0.473404 | 0.266944 | Q10589-2;Q10589 | Q10589-2;Q10589 |
| Tumor-associated calcium signal transducer 2 | TACSTD2 | | 0.33981 | 0.263434 | P09758 | P09758 |
| Brain-specific angiogenesis inhibitor 1-associated protein 2-like protein 1 | BAIAP2L1 | + | 1.31791 | 0.263363 | Q9UHR4 | Q9UHR4 |
| Synaptosomal-associated protein 23 | SNAP23 | | 0.257952 | 0.263055 | O00161;O00161-2 | O00161;O00161-2 |
| Tubulin beta-3 chain | TUBB3 | | 0.206562 | 0.262657 | Q13509;Q13509-2 | Q13509;Q13509-2 |
| Semaphorin-3C | SEMA3C | | 0.413371 | 0.262479 | Q99985;Q99985-2 | Q99985 |
| Neuropilin-1 | NRP1 | | 0.612879 | 0.259585 | O14786;O14786-3;O14786-2 | O14786;O14786-3;O14786-2 |
| Metalloproteinase inhibitor 1 | TIMP1 | | 0.33058 | 0.259057 | P01033 | P01033 |
| Collagen alpha-2(VI) chain | COL6A2 | | 0.224481 | 0.258616 | P12110;P12110-3;P12110-2 | P12110;P12110-3;P12110-2 |
| Cytochrome c oxidase subunit 4 isoform 1, mitochondrial | COX4I1 | | 0.314221 | 0.258608 | P13073 | P13073 |
| Platelet-derived growth factor D;Platelet-derived growth factor D, latent form;Platelet-derived growth factor D, | PDGFD | | 0.528936 | 0.253407 | Q9GZP0;Q9GZP0-2 | Q9GZP0;Q9GZP0-2 |
| Peptidyl-prolyl cis-trans isomerase A;Peptidyl-prolyl cis-trans isomerase A, N-terminally processed | PPIA | | 0.872237 | 0.252938 | P62937;Q9Y536;F5H284 | P62937 |
| Glycogen phosphorylase, brain form | PYGB | | 0.784469 | 0.251602 | P11216 | P11216 |
| Protein FAM65A | FAM65A | | 0.113823 | 0.247946 | Q6ZS17-2;Q6ZS17;Q6ZS17-3;Q6ZS17-4 | Q6ZS17-2;Q6ZS17;Q6ZS17-3;Q6ZS17-4 |
| Ras-related protein Rap-1b;Ras-related protein Rap-1b-like protein | RAP1B | | 0.482586 | 0.247775 | P61224;P61224-3;A6NIZ1;P61224-2;P61224-4 | P61224;P61224-3;A6NIZ1;P61224-2;P61224-4 |
| Meteorin-like protein | METRNL | | 0.21734 | 0.247005 | Q641Q3;Q641Q3-2 | Q641Q3;Q641Q3-2 |
| Amyloid beta A4 protein;N-APP;Soluble APP-alpha;Soluble APP-beta;C99;Beta-amyloid protein 42;Beta-amyloid | APP | | 0.347893 | 0.246878 | P05067-7;P05067-11;P05067-8;P05067-9;P05067;P05067-3;P05067- | P05067-7;P05067-11;P05067-8;P05067-9;P05067;P05067 |
| Isocitrate dehydrogenase [NADP], mitochondrial | IDH2 | | 0.0728409 | 0.24676 | P48735-2;P48735 | P48735-2;P48735 |
| COMM domain-containing protein 1 | COMMD1 | | 0.0921621 | 0.244539 | Q8N668;Q8N668-2 | Q8N668;Q8N668-2 |
| Disco-interacting protein 2 homolog B | DIP2B | | 0.522755 | 0.242558 | Q9P265;Q9Y2E4 | Q9P265 |
| Lipolysis-stimulated lipoprotein receptor | LSR | | 0.302084 | 0.236006 | Q86X29-6;Q86X29-5;Q86X29-4;Q86X29;Q86X29-2 | Q86X29-6;Q86X29-5;Q86X29-4;Q86X29;Q86X29-2 |
| ATP synthase subunit alpha, mitochondrial | ATP5A1 | | 0.183763 | 0.234687 | P25705;P25705-2;P25705-3 | P25705;P25705-2;P25705-3 |
| Peroxidasin homolog | PXDN | | 0.489969 | 0.23369 | Q92626;Q92626-2 | Q92626 |
| Collagen alpha-1(VII) chain | COL7A1 | | 0.223751 | 0.233516 | Q02388-2;Q02388 | Q02388-2;Q02388 |
| Heterogeneous nuclear ribonucleoprotein K | HNRNPK | | 0.679967 | 0.231523 | P61978-3 | P61978-3 |
| Ras-related protein Rab-21 | RAB21 | | 0.481851 | 0.230612 | Q9UL25 | Q9UL25 |
| Unconventional myosin-VI | MYO6 | | 0.279629 | 0.23051 | Q9UM54-5;Q9UM54-2;Q9UM54-6;Q9UM54-4;Q9UM54-1;Q9UM54 | Q9UM54-5;Q9UM54-2;Q9UM54-6;Q9UM54-4;Q9UM54- |
| 2-deoxynucleoside 5-phosphate N-hydrolase 1 | DNPH1 | | 0.267217 | 0.227362 | O43598;O43598-2 | O43598;O43598-2 |
| Pyridoxine-5-phosphate oxidase | PNPO | | 0.194783 | 0.22581 | Q9NVS9-3;Q9NVS9;Q9NVS9-2;Q9NVS9-4 | Q9NVS9-3;Q9NVS9;Q9NVS9-2;Q9NVS9-4 |
| ATP synthase subunit gamma, mitochondrial | ATP5C1 | | 0.103469 | 0.224099 | P36542-2;P36542 | P36542-2;P36542 |
| Deoxyribonuclease-2-alpha | DNASE2 | | 0.11096 | 0.221865 | O00115-2;O00115 | O00115-2;O00115 |
| Protein scribble homolog | SCRIB | | 0.336724 | 0.221328 | Q14160;Q14160-3 | Q14160;Q14160-3 |
| Sodium/potassium-transporting ATPase subunit beta-1 | ATP1B1 | | 0.430282 | 0.219508 | P05026-2;P05026 | P05026-2;P05026 |
| Ras-related protein Rab-14 | RAB14 | | 0.23957 | 0.218705 | P61106 | P61106 |
| Apoptosis-associated speck-like protein containing a CARD | PYCARD | | 0.367208 | 0.216398 | Q9ULZ3;Q9ULZ3-3;Q9ULZ3-2 | Q9ULZ3;Q9ULZ3-3;Q9ULZ3-2 |
| Collagen alpha-1(XVI) chain | COL16A1 | | 0.860001 | 0.215581 | Q07092-2;Q07092 | Q07092-2;Q07092 |
| Proliferation-associated protein 2G4 | PA2G4 | | 0.24752 | 0.212151 | Q9UQ80;Q9UQ80-2 | Q9UQ80;Q9UQ80-2 |
| CMP-N-acetylneuraminate-beta-galactosamide-alpha-2,3-sialyltransferase 4 | ST3GAL4 | | 0.175267 | 0.211861 | Q11206-3;Q11206-2;Q11206;Q11206-4;Q11206-7;Q11206-6;Q1120 | Q11206-3;Q11206-2;Q11206;Q11206-4;Q11206-7;Q1120 |
| Legumain | LGMN | | 0.391374 | 0.211585 | Q99538;Q99538-2;Q99538-3 | Q99538;Q99538-2;Q99538-3 |
| Endoplasmic reticulum resident protein 29 | ERP29 | | 0.178369 | 0.211337 | P30040;P30040-2 | P30040 |
| ATP synthase subunit beta, mitochondrial | ATP5B | | 0.122316 | 0.209651 | P06576 | P06576 |
| Gamma-interferon-inducible lysosomal thiol reductase | IFI30 | | 0.161589 | 0.209503 | P13284 | P13284 |
| Endoplasmic reticulum resident protein 44 | ERP44 | | 0.149878 | 0.208014 | Q9BS26 | Q9BS26 |
| Cation-independent mannose-6-phosphate receptor | IGF2R | | 0.312784 | 0.20708 | P11717 | P11717 |
| Alpha-N-acetylgalactosaminidase | NAGA | | 0.187549 | 0.20292 | P17050 | P17050 |
| Eukaryotic translation initiation factor 5A-1;Eukaryotic translation initiation factor 5A-1-like | EIF5A;EIF5AL1 | | 0.467835 | 0.202707 | P63241;P63241-2;Q6IS14 | P63241;P63241-2;Q6IS14 |
| Tryptophan--tRNA ligase, cytoplasmic;T1-TrpRS;T2-TrpRS | WARS | | 0.465387 | 0.201168 | P23381;P23381-2 | P23381;P23381-2 |
| Heat shock protein HSP 90-alpha | HSP90AA1 | | 0.18573 | 0.201092 | P07900;P07900-2;Q58FG0 | P07900;P07900-2 |
| D-dopachrome decarboxylase;D-dopachrome decarboxylase-like protein | DDT;DDTL | | 0.175654 | 0.199349 | P30046;P30046-2;A6NHG4 | P30046;P30046-2;A6NHG4 |
| Proliferating cell nuclear antigen | PCNA | | 0.452035 | 0.198625 | P12004 | P12004 |
| Endoplasmin | HSP90B1 | | 0.281013 | 0.197854 | P14625;Q58FF3 | P14625 |
| RuvB-like 1 | RUVBL1 | | 0.51095 | 0.196767 | Q9Y265;Q9Y265-2 | Q9Y265;Q9Y265-2 |
| Tubulin alpha-4A chain | TUBA4A | | 0.226116 | 0.195845 | P68366;P68366-2 | P68366;P68366-2 |
| Voltage-dependent anion-selective channel protein 3 | VDAC3 | | 0.233605 | 0.195322 | Q9Y277;Q9Y277-2 | Q9Y277;Q9Y277-2 |
| Latent-transforming growth factor beta-binding protein 4 | LTBP4 | | 0.122179 | 0.193731 | Q8N2S1-2;Q8N2S1-3;Q8N2S1;Q8N2S1-4 | Q8N2S1-2;Q8N2S1-3;Q8N2S1 |
| Acetyl-CoA acetyltransferase, cytosolic | ACAT2 | | 0.162105 | 0.188358 | Q9BWD1;Q9BWD1-2 | Q9BWD1;Q9BWD1-2 |
| Heme-binding protein 2 | HEBP2 | | 0.0885223 | 0.187332 | Q9Y5Z4;Q9Y5Z4-2 | Q9Y5Z4;Q9Y5Z4-2 |
| Nucleolysin TIAR | TIAL1 | | 0.203958 | 0.186389 | Q01085 | Q01085 |
| Actin, alpha skeletal muscle;Actin, alpha cardiac muscle 1;Actin, gamma-enteric smooth muscle;Actin, aortic sm | ACTA1;ACTC1;ACTG2;ACTA2 | | 0.073746 | 0.186045 | P68133;P68032;P63267;P62736;P63267-2 | P68133;P68032;P63267;P62736;P63267-2 |
| Chitinase domain-containing protein 1 | CHID1 | | 0.107336 | 0.185781 | Q9BWS9-3;Q9BWS9;Q9BWS9-2 | Q9BWS9-3;Q9BWS9;Q9BWS9-2 |
| Phosphoribosylformylglycinamidine synthase | PFAS | | 0.228485 | 0.185548 | O15067 | O15067 |
| Prosaposin;Saposin-A;Saposin-B-Val;Saposin-B;Saposin-C;Saposin-D | PSAP | | 0.215229 | 0.180792 | P07602;P07602-2;P07602-3 | P07602;P07602-2;P07602-3 |
| Heat shock protein 105 kDa | HSPH1 | | 0.0370429 | 0.180488 | Q92598;Q92598-3;Q92598-4 | Q92598;Q92598-3;Q92598-4 |
| Protein PAXX | C9orf142 | | 1.1968 | 0.179085 | Q9BUH6;Q9BUH6-2 | Q9BUH6 |
| Ribulose-phosphate 3-epimerase | RPE | | 0.174098 | 0.178612 | Q96AT9;Q96AT9-2;Q96AT9-5;Q96AT9-3;Q96AT9-4;Q2QD12 | Q96AT9;Q96AT9-2;Q96AT9-5;Q96AT9-3;Q96AT9-4 |
| ADP-ribosylation factor 1;ADP-ribosylation factor 3 | ARF1;ARF3 | | 0.272042 | 0.177205 | P84077;P61204;P61204-2 | P84077;P61204;P61204-2 |

| | | | | | |
|---|---|---|---|---|---|
| Phosphoserine aminotransferase | PSAT1 | 0.508973 | 0.174833 | Q9Y617;Q9Y617-2 | Q9Y617;Q9Y617-2 |
| Unconventional myosin-Ie | MYO1E | 0.225021 | 0.172787 | Q12965 | Q12965 |
| Protein kinase C and casein kinase substrate in neurons protein 3 | PACSIN3 | 0.438769 | 0.169776 | Q9UKS6 | Q9UKS6 |
| DnaJ homolog subfamily A member 2 | DNAJA2 | 0.587243 | 0.169208 | O60884 | O60884 |
| Tetratricopeptide repeat protein 38 | TTC38 | 0.187685 | 0.16857 | Q5R3I4 | Q5R3I4 |
| Arginase-1 | ARG1 | 0.0631314 | 0.167013 | P05089;P05089-2;P05089-3 | P05089;P05089-2;P05089-3 |
| Complement decay-accelerating factor | CD55 | 0.228033 | 0.165493 | P08174-4;P08174;P08174-3;P08174-5;P08174-2;P08174-6;P08174-7 | P08174-4;P08174;P08174-3;P08174-5;P08174-2;P08174- |
| Superoxide dismutase [Cu-Zn] | SOD1 | 0.149177 | 0.163946 | P00441 | P00441 |
| Fascin | FSCN1 | 0.284203 | 0.163617 | Q16658 | Q16658 |
| Dynein light chain roadblock-type 1 | DYNLRB1 | 0.147754 | 0.163363 | Q9NP97;Q8TF09;Q9NP97-2 | Q9NP97 |
| Sialate O-acetylesterase | SIAE | 0.0901997 | 0.160819 | Q9HAT2;Q9HAT2-2 | Q9HAT2;Q9HAT2-2 |
| Connective tissue growth factor | CTGF | 0.13941 | 0.160135 | P29279;P29279-2 | P29279;P29279-2 |
| Rho-related GTP-binding protein RhoG | RHOG | 0.47602 | 0.158797 | P84095;Q9H4E5-2;P17081;Q9H4E5 | P84095 |
| Thioredoxin-dependent peroxide reductase, mitochondrial | PRDX3 | 0.110184 | 0.157525 | P30048-2;P30048 | P30048-2;P30048 |
| Tubulin beta-4B chain | TUBB4B | 0.257878 | 0.156483 | P68371 | P68371 |
| N-acetylgalactosaminyltransferase 7 | GALNT7 | 0.194842 | 0.154994 | Q86SF2 | Q86SF2 |
| Polyadenylate-binding protein 1 | PABPC1 | 0.173299 | 0.154179 | P11940;P11940-2;Q9H361;Q96DU9-2;Q96DU9 | P11940;P11940-2 |
| Protein CutA | CUTA | 0.135191 | 0.153554 | O60888-3;O60888;O60888-2 | O60888-3;O60888;O60888-2 |
| Pyridoxal kinase | PDXK | 0.187077 | 0.151665 | O00764;O00764-3;O00764-2 | O00764;O00764-3;O00764-2 |
| Palmitoyl-protein thioesterase 1 | PPT1 | 0.170656 | 0.150434 | P50897;P50897-2 | P50897 |
| Procollagen-lysine,2-oxoglutarate 5-dioxygenase 2 | PLOD2 | 0.458031 | 0.149417 | O00469-2;O00469;O00469-3 | O00469-2;O00469;O00469-3 |
| Protein Wnt-5a | WNT5A | 0.137747 | 0.14911 | P41221-2;P41221;O00755 | P41221-2;P41221 |
| DnaJ homolog subfamily B member 1 | DNAJB1 | 0.104138 | 0.148718 | P25685;P25685-2 | P25685;P25685-2 |
| Dynactin subunit 2 | DCTN2 | 0.50438 | 0.147945 | Q13561;Q13561-3;Q13561-2 | Q13561;Q13561-3;Q13561-2 |
| N-acetylglucosamine-6-sulfatase | GNS | 0.0868606 | 0.147603 | P15586;P15586-2 | P15586;P15586-2 |
| Metalloreductase STEAP4 | STEAP4 | 0.106183 | 0.145528 | Q687X5;Q687X5-2 | Q687X5;Q687X5-2 |
| Eukaryotic translation initiation factor 3 subunit E | EIF3E | 0.524106 | 0.144288 | P60228 | P60228 |
| Glia-derived nexin | SERPINE2 | 0.663331 | 0.142665 | P07093-2;P07093-3 | P07093-2;P07093-3 |
| Ras-related protein Ral-B | RALB | 0.0463249 | 0.142605 | P11234;P11234-2;P11234-3 | P11234;P11234-2;P11234-3 |
| Sialidase-1 | NEU1 | 0.0957555 | 0.140352 | Q99519 | Q99519 |
| Mitochondrial peptide methionine sulfoxide reductase | MSRA | 0.106889 | 0.139701 | Q9UJ68-2;Q9UJ68-3;Q9UJ68-5;Q9UJ68;Q9UJ68-4 | Q9UJ68-2;Q9UJ68-3;Q9UJ68-5;Q9UJ68;Q9UJ68-4 |
| G-protein coupled receptor 56;GPR56 N-terminal fragment;GPR56 C-terminal fragment | GPR56 | 0.190749 | 0.138696 | Q9Y653-2;Q9Y653-3;Q9Y653;Q9Y653-4 | Q9Y653-2;Q9Y653-3;Q9Y653 |
| Serine hydroxymethyltransferase, mitochondrial | SHMT2 | 0.191622 | 0.136386 | P34897-3;P34897;P34897-2 | P34897-3;P34897;P34897-2 |
| Metalloproteinase inhibitor 4 | TIMP4 | 0.477096 | 0.135798 | Q99727 | Q99727 |
| Annexin A4 | ANXA4 | 0.278449 | 0.134169 | P09525;P09525-2 | P09525;P09525-2 |
| Protein Wiz | WIZ | 0.0813114 | 0.131461 | O95785-4;O95785-3;O95785;O95785-2 | O95785-4;O95785-3;O95785;O95785-2 |
| Ras-related protein Rab-6A | RAB6A | 0.325883 | 0.131032 | P20340-2;P20340-4;Q9H0N0 | P20340-2;P20340-4 |
| Glycine--tRNA ligase | GARS | 0.804957 | 0.124692 | P41250 | P41250 |
| Asparagine--tRNA ligase, cytoplasmic | NARS | 0.498315 | 0.124633 | O43776;O43776-2 | O43776 |
| Fatty acid-binding protein, epidermal | FABP5 | 0.186303 | 0.123712 | Q01469;A8MUU1 | Q01469 |
| Low-density lipoprotein receptor | LDLR | 0.101571 | 0.123542 | P01130-4;P01130-5;P01130;P01130-3;P01130-6;P01130-2 | P01130-4;P01130-5;P01130;P01130-3;P01130-6;P01130- |
| Enolase-phosphatase E1 | ENOPH1 | 0.287419 | 0.12328 | Q9UHY7;Q9UHY7-2 | Q9UHY7;Q9UHY7-2 |
| DnaJ homolog subfamily A member 1 | DNAJA1 | 0.266823 | 0.121534 | P31689;P31689-2 | P31689;P31689-2 |
| Collagen alpha-2(I) chain | COL1A2 | 0.312358 | 0.118563 | P08123 | P08123 |
| Haloacid dehalogenase-like hydrolase domain-containing protein 2 | HDHD2 | 0.11047 | 0.115992 | Q9H0R4;Q9H0R4-2 | Q9H0R4;Q9H0R4-2 |
| Poly(rC)-binding protein 1 | PCBP1 | 0.270255 | 0.113301 | Q15365 | Q15365 |
| Insulin-degrading enzyme | IDE | 0.188932 | 0.112371 | P14735;P14735-2 | P14735;P14735-2 |
| Eukaryotic translation initiation factor 3 subunit B | EIF3B | 0.190127 | 0.108802 | P55884;P55884-2 | P55884;P55884-2 |
| Serine/threonine-protein kinase 24;Serine/threonine-protein kinase 24 36 kDa subunit;Serine/threonine-protei | STK24 | 0.353047 | 0.108633 | Q9Y6E0-2;Q9Y6E0 | Q9Y6E0-2;Q9Y6E0 |
| Leucine zipper protein 1 | LUZP1 | 0.22118 | 0.107738 | Q86V48-2;Q86V48-3;Q86V48 | Q86V48-2;Q86V48-3;Q86V48 |
| Serine/threonine-protein phosphatase CPPED1 | CPPED1 | 0.318733 | 0.106202 | Q9BRF8;Q9BRF8-2 | Q9BRF8;Q9BRF8-2 |
| Inositol monophosphatase 1 | IMPA1 | 0.073067 | 0.10331 | P29218;P29218-3;P29218-2 | P29218;P29218-3;P29218-2 |
| 60S acidic ribosomal protein P2 | RPLP2 | 0.480321 | 0.103232 | P05387;P05386-2 | P05387 |
| Calsyntenin-1;Soluble Alc-alpha;CTF1-alpha | CLSTN1 | 0.129357 | 0.103157 | O94985-2;Q9H4D0 | O94985-2 |
| Isochorismatase domain-containing protein 1 | ISOC1 | 0.0604505 | 0.102001 | Q96CN7 | Q96CN7 |
| Collagen alpha-1(IV) chain;Arresten | COL4A1 | 0.076073 | 0.098079 | P02462-2;P02462;P29400;P29400-2 | P02462-2;P02462 |
| 78 kDa glucose-regulated protein | HSPA5 | 0.144738 | 0.097976 | P11021 | P11021 |
| Claudin-4;Claudin-9;Claudin-6;Claudin-3 | CLDN4;CLDN9;CLDN6;CLDN3 | 0.13326 | 0.0967687 | O14493;O95484;P56747;O15551 | O14493;O95484;P56747;O15551 |
| Tubulin-specific chaperone A | TBCA | 0.113912 | 0.0946782 | O75347;O75347-2 | O75347;O75347-2 |
| Insulin-like growth factor-binding protein 7 | IGFBP7 | 0.0982981 | 0.0930405 | Q16270-2;Q16270 | Q16270-2;Q16270 |
| Thioredoxin-like protein 1 | TXNL1 | 0.169888 | 0.0925668 | O43396 | O43396 |
| Gelsolin | GSN | 0.283626 | 0.0922794 | P06396-2;P06396-4;P06396-3;P06396 | P06396-2;P06396-4;P06396-3;P06396 |
| Catalase | CAT | 0.0474583 | 0.0882696 | P04040 | P04040 |
| Beta-glucuronidase | GUSB | 0.0609645 | 0.0873 | P08236-2;P08236;P08236-3 | P08236-2;P08236;P08236-3 |
| Peripheral plasma membrane protein CASK | CASK | 0.124869 | 0.0872676 | O14936-3;O14936-4;O14936-6;O14936-2;O14936;O14936-5 | O14936-3;O14936-4;O14936-6;O14936-2;O14936;O1493 |
| Acid ceramidase;Acid ceramidase subunit alpha;Acid ceramidase subunit beta | ASAH1 | 0.06126 | 0.0865707 | Q13510;Q13510-2;Q13510-3 | Q13510;Q13510-2;Q13510-3 |
| Putative phospholipase B-like 2;Putative phospholipase B-like 2 32 kDa form;Putative phospholipase B-like 2 45 | PLBD2 | 0.177969 | 0.0848986 | Q8NHP8;Q8NHP8-2 | Q8NHP8;Q8NHP8-2 |
| Dihydropteridine reductase | QDPR | 1.00685 | 0.084053 | P09417;P09417-2 | P09417;P09417-2 |
| Septin-7 | SEPT7 | 0.1077 | 0.0829042 | Q16181;Q16181-2;Q6ZU15 | Q16181;Q16181-2 |
| ADP-ribosylation factor 6 | ARF6 | 0.0501611 | 0.0820287 | P62330 | P62330 |
| CAD protein;Glutamine-dependent carbamoyl-phosphate synthase;Aspartate carbamoyltransferase;Dihydroor | CAD | 0.129186 | 0.0798384 | P27708 | P27708 |
| Peroxiredoxin-1 | PRDX1 | 0.117061 | 0.0790354 | Q06830 | Q06830 |
| Multifunctional protein ADE2;Phosphoribosylaminoimidazole-succinocarboxamide synthase;Phosphoribosylam | PAICS | 0.0574431 | 0.0786438 | P22234;P22234-2 | P22234;P22234-2 |
| Group XV phospholipase A2 | PLA2G15 | 0.124508 | 0.0772495 | Q8NCC3;Q8NCC3-2 | Q8NCC3 |
| 4-trimethylaminobutyraldehyde dehydrogenase | ALDH9A1 | 0.14898 | 0.0769526 | P49189;P49189-2 | P49189;P49189-2 |
| ATP-dependent RNA helicase DDX3X | DDX3X | 0.0461385 | 0.0745182 | O00571-2;O00571;Q9NQI0-3;Q9NQI0-2;Q9NQI0-4;Q9NQI0 | O00571-2;O00571 |
| Ribonuclease T2 | RNASET2 | 0.0862853 | 0.073925 | O00584;O00584-2 | O00584 |
| Nebulin | NEB | 0.0358504 | 0.0737604 | P20929-4;P20929;P20929-3;P20929-2 | P20929-4;P20929;P20929-3;P20929-2 |
| NudC domain-containing protein 2 | NUDCD2 | 0.0992715 | 0.0734488 | Q8WVJ2 | Q8WVJ2 |
| Inorganic pyrophosphatase 2, mitochondrial | PPA2 | 0.0923131 | 0.0717964 | Q9H2U2;Q9H2U2-2;Q9H2U2-6;Q9H2U2-3;Q9H2U2-4 | Q9H2U2;Q9H2U2-2;Q9H2U2-6;Q9H2U2-3 |

| | | | | | |
|---|---|---|---|---|---|
| Keratin, type I cytoskeletal 18 | KRT18 | 0.0827539 | 0.0705725 | P05783;CON__H-INV:HIT000015463 | P05783 |
| Peroxiredoxin-2 | PRDX2 | 0.0807586 | 0.0693817 | P32119;P32119-2 | P32119 |
| Annexin A7 | ANXA7 | 0.0649602 | 0.0680917 | P20073-2;P20073 | P20073-2;P20073 |
| Pyruvate kinase PKM | PKM | 0.155762 | 0.0673434 | P14618 | P14618 |
| Beta-hexosaminidase subunit beta;Beta-hexosaminidase subunit beta chain B;Beta-hexosaminidase subunit be | HEXB | 0.0455479 | 0.065506 | P07686 | P07686 |
| UPF0565 protein C2orf69 | C2orf69 | 0.0603805 | 0.0651964 | Q8N8R5 | Q8N8R5 |
| Insulin-like growth factor-binding protein 4 | IGFBP4 | 0.0511583 | 0.0635522 | P22692;P22692-2 | P22692;P22692-2 |
| Integrin beta-like protein 1 | ITGBL1 | 0.0549184 | 0.0630175 | O95965;O95965-3;O95965-2 | O95965;O95965-3;O95965-2 |
| Metalloproteinase inhibitor 2 | TIMP2 | 0.0604142 | 0.0622323 | P16035 | P16035 |
| Ectonucleoside triphosphate diphosphohydrolase 6 | ENTPD6 | 0.0758792 | 0.0621332 | O75354-2;O75354-3;O75354 | O75354-2;O75354-3;O75354 |
| Calnexin | CANX | 0.0815396 | 0.0605437 | P27824;P27824-2;P27824-3 | P27824;P27824-2;P27824-3 |
| Elongation factor Tu, mitochondrial | TUFM | 0.0265785 | 0.0604947 | P49411 | P49411 |
| Transmembrane emp24 domain-containing protein 4 | TMED4 | 0.0315532 | 0.0603701 | Q7Z7H5-3;Q7Z7H5-2;Q7Z7H5;Q9BVK6 | Q7Z7H5-3;Q7Z7H5-2;Q7Z7H5 |
| Peroxiredoxin-6 | PRDX6 | 0.154955 | 0.060318 | P30041 | P30041 |
| Cofilin-1 | CFL1 | 0.194594 | 0.0588328 | P23528 | P23528 |
| Ras-related protein Rab-13 | RAB13 | 0.234177 | 0.0575924 | P51153 | P51153 |
| Protein NOV homolog | NOV | 0.0436581 | 0.0568066 | P48745 | P48745 |
| Nuclear transport factor 2 | NUTF2 | 0.0543763 | 0.0549399 | P61970 | P61970 |
| Putative hydrolase RBBP9 | RBBP9 | 0.0364288 | 0.0546494 | O75884;O75884-2 | O75884;O75884-2 |
| Multivesicular body subunit 12A | MVB12A | 0.171839 | 0.0542119 | Q96EY5;Q96EY5-3;Q96EY5-2 | Q96EY5;Q96EY5-3;Q96EY5-2 |
| Prolyl endopeptidase FAP;Antiplasmin-cleaving enzyme FAP, soluble form | FAP | 0.0786427 | 0.0527668 | Q12884;Q12884-2 | Q12884 |
| DnaJ homolog subfamily C member 5 | DNAJC5 | 0.0354293 | 0.0513713 | Q9H3Z4-2;Q9H3Z4 | Q9H3Z4-2;Q9H3Z4 |
| Torsin-1B | TOR1B | 0.133525 | 0.0504557 | O14657 | O14657 |
| Bone morphogenetic protein 1 | BMP1 | 0.0657218 | 0.0502911 | P13497;P13497-5;P13497-2;P13497-6;P13497-4;P13497-3 | P13497;P13497-5;P13497-2;P13497-6;P13497-4 |
| Protein SEC13 homolog | SEC13 | 0.0504348 | 0.0496457 | P55735;P55735-3;P55735-2 | P55735;P55735-3;P55735-2 |
| Aflatoxin B1 aldehyde reductase member 2 | AKR7A2 | 0.0446723 | 0.0493768 | O43488;Q8NHP1;O95154 | O43488 |
| Neutral alpha-glucosidase AB | GANAB | 0.0622704 | 0.0492268 | Q14697;Q14697-2;Q14697-3 | Q14697;Q14697-2 |
| Ataxin-10 | ATXN10 | 0.0902702 | 0.0486927 | Q9UBB4;Q9UBB4-2 | Q9UBB4;Q9UBB4-2 |
| Proto-oncogene tyrosine-protein kinase Src | SRC | 0.0714668 | 0.048481 | P12931;P12931-2;P06241-3;P06241-2;P06241;P08631-3;P08631-2;P | P12931;P12931-2 |
| Annexin A11 | ANXA11 | 0.0711519 | 0.048076 | P50995-2;P50995 | P50995-2;P50995 |
| Creatine kinase U-type, mitochondrial | CKMT1A | 0.179551 | 0.0450643 | P12532;P17540 | P12532 |
| Protein phosphatase 1 regulatory subunit 7 | PPP1R7 | 0.0875322 | 0.0449454 | Q15435;Q15435-2;Q15435-3;Q15435-5;Q15435-4 | Q15435;Q15435-2;Q15435-3;Q15435-5;Q15435-4 |
| Programmed cell death protein 5 | PDCD5 | 0.0881366 | 0.0444921 | O14737;O14737-2 | O14737;O14737-2 |
| Exosome complex component RRP45 | EXOSC9 | 0.0300269 | 0.0423107 | Q06265;Q06265-2;Q06265-3;Q06265-4 | Q06265;Q06265-2;Q06265-3;Q06265-4 |
| Receptor-type tyrosine-protein phosphatase gamma | PTPRG | 0.0501864 | 0.0418797 | P23470-2;P23470 | P23470-2;P23470 |
| Cytosol aminopeptidase | LAP3 | 0.0205484 | 0.0407651 | P28838-2;P28838 | P28838-2;P28838 |
| CD9 antigen | CD9 | 0.0380499 | 0.0398299 | P21926 | P21926 |
| Hepatocyte growth factor receptor | MET | 0.0412056 | 0.0379639 | P08581;P08581-2;P08581-3;P42685-2;P42685 | P08581;P08581-2;P08581-3 |
| Glutathione S-transferase A3 | GSTA3 | 0.0407093 | 0.0377064 | Q16772 | Q16772 |
| Guanine nucleotide-binding protein-like 1 | GNL1 | 0.0487317 | 0.0363839 | P36915;P36915-2 | P36915;P36915-2 |
| Protein kinase C iota type | PRKCI | 0.0591595 | 0.0352306 | P41743 | P41743 |
| Alpha-taxilin | TXLNA | 0.0199857 | 0.0350577 | P40222 | P40222 |
| Integrin beta-1 | ITGB1 | 0.0489564 | 0.0346457 | P05556;P05556-2;P05556-5;P05556-4;P05556-3 | P05556;P05556-2;P05556-5;P05556-4;P05556-3 |
| Xylosyltransferase 1 | XYLT1 | 0.0161456 | 0.0309594 | Q86Y38 | Q86Y38 |
| Endothelial differentiation-related factor 1 | EDF1 | 0.0210782 | 0.0305532 | O60869-2;O60869-3;O60869 | O60869-2;O60869-3;O60869 |
| Alpha-enolase | ENO1 | 0.0298414 | 0.0278994 | P06733 | P06733 |
| Laminin subunit beta-2 | LAMB2 | 0.0307409 | 0.0253213 | P55268;A4D0S4-2;A4D0S4-3;A4D0S4 | P55268 |
| Cytosolic non-specific dipeptidase | CNDP2 | 0.0405729 | 0.0237103 | Q96KP4;Q96KP4-2 | Q96KP4;Q96KP4-2 |
| Protein NDRG1 | NDRG1 | 0.0246421 | 0.0235062 | Q92597;Q92597-3;Q92597-2 | Q92597;Q92597-3;Q92597-2 |
| Serine hydroxymethyltransferase, cytosolic | SHMT1 | 0.044067 | 0.0232633 | P34896-2;P34896;P34896-3;P34896-4 | P34896-2;P34896;P34896-3;P34896-4 |
| Prothymosin alpha;Prothymosin alpha, N-terminally processed;Thymosin alpha-1 | PTMA | 0.0071267 | 0.0222893 | P06454-2;P06454 | P06454-2;P06454 |
| Synaptobrevin homolog YKT6 | YKT6 | 0.018872 | 0.0218506 | O15498;O15498-2 | O15498;O15498-2 |
| Probable ATP-dependent RNA helicase DDX5 | DDX5 | 0.0203153 | 0.0202726 | P17844;P17844-2 | P17844;P17844-2 |
| Tyrosine-protein kinase Lyn | LYN | 0.022449 | 0.0172952 | P07948-2;P07948 | P07948-2;P07948 |
| Endoglin | ENG | 0.0187958 | 0.0157032 | P17813-2;P17813 | P17813-2;P17813 |
| Ras-related protein Rab-11B;Ras-related protein Rab-11A | RAB11B;RAB11A | 0.0193371 | 0.0156403 | Q15907;Q15907-2;P62491-2 | Q15907;Q15907-2;P62491-2 |
| Tissue-type plasminogen activator;Tissue-type plasminogen activator chain A;Tissue-type plasminogen activato | PLAT | 0.0146055 | 0.0154209 | P00750;P00750-3;P00750-4;P00750-2 | P00750;P00750-3;P00750-4 |
| Bifunctional ATP-dependent dihydroxyacetone kinase/FAD-AMP lyase (cyclizing);ATP-dependent dihydroxyacet | DAK | 0.0152452 | 0.0152613 | Q3LXA3;Q3LXA3-2 | Q3LXA3;Q3LXA3-2 |
| Peroxiredoxin-4 | PRDX4 | 0.0040312 | 0.00915591 | Q13162 | Q13162 |
| Epididymis-specific alpha-mannosidase | MAN2B2 | 0.00466554 | 0.00891113 | Q9Y2E5;Q9Y2E5-2 | Q9Y2E5;Q9Y2E5-2 |
| Mannose-1-phosphate guanyltransferase alpha | GMPPA | 0.00450668 | 0.00652377 | Q96IJ6;Q96IJ6-2 | Q96IJ6;Q96IJ6-2 |
| Lysosomal alpha-mannosidase;Lysosomal alpha-mannosidase A peptide;Lysosomal alpha-mannosidase B pepti | MAN2B1 | 0.00395849 | 0.00464058 | O00754-2;O00754 | O00754-2;O00754 |
| Acetyl-CoA acetyltransferase, mitochondrial | ACAT1 | 0.0116223 | 0.00460498 | P24752;P24752-2 | P24752 |
| C-1-tetrahydrofolate synthase, cytoplasmic;Methylenetetrahydrofolate dehydrogenase;Methenyltetrahydrofol | MTHFD1 | 0.007008 | 0.00402768 | P11586 | P11586 |
| Pyridoxal phosphate phosphatase | PDXP | 0.00422194 | 0.00342496 | Q96GD0 | Q96GD0 |
| Triosephosphate isomerase | TPI1 | 0.000202687 | -0.000257492 | P60174-1;P60174;P60174-4 | P60174-1;P60174;P60174-4 |
| Endothelin-converting enzyme 1 | ECE1 | 0.00144772 | -0.00151634 | P42892-3;P42892-2;P42892-4;P42892 | P42892-3;P42892-2;P42892-4;P42892 |
| Integrin alpha-6;Integrin alpha-6 heavy chain;Integrin alpha-6 light chain;Processed integrin alpha-6 | ITGA6 | 0.00352173 | -0.0023454 | P23229-4;P23229-2;P23229-5;P23229-3;P23229-9;P23229-6;P23229 | P23229-4;P23229-2;P23229-5;P23229-3;P23229-9;P2322 |
| Adenylate kinase 2, mitochondrial;Adenylate kinase 2, mitochondrial, N-terminally processed | AK2 | 0.00316165 | -0.00377909 | P54819-2;P54819;P54819-5;P54819-3;P54819-6;P54819-4 | P54819-2;P54819;P54819-5;P54819-3;P54819-6;P54819- |
| RelA-associated inhibitor | PPP1R13L | 0.00122283 | -0.00476646 | Q8WUF5 | Q8WUF5 |
| Complement component 1 Q subcomponent-binding protein, mitochondrial | C1QBP | 0.00336584 | -0.00537745 | Q07021 | Q07021 |
| CD44 antigen | CD44 | 0.00511642 | -0.00668844 | P16070-8;P16070-4;P16070-3;P16070-7;P16070-5;P16070;P16070-1 | P16070-8;P16070-4;P16070-3;P16070-7;P16070-5;P1607 |
| Contactin-1 | CNTN1 | 0.0211425 | -0.00830523 | Q12860-2;Q12860;Q12860-3 | Q12860-2;Q12860;Q12860-3 |
| Ubiquitin-fold modifier 1 | UFM1 | 0.0106291 | -0.00831604 | P61960;P61960-2 | P61960;P61960-2 |
| Soluble calcium-activated nucleotidase 1 | CANT1 | 0.0281377 | -0.00928688 | Q8WVQ1-3;Q8WVQ1;Q8WVQ1-2 | Q8WVQ1-3;Q8WVQ1;Q8WVQ1-2 |
| Transcription activator BRG1 | SMARCA4 | 0.0142546 | -0.0111319 | P51532-5;P51532-2;P51532-3;P51532-4;P51532 | P51532-5;P51532-2;P51532-3;P51532-4;P51532 |
| Crk-like protein | CRKL | 0.0210149 | -0.0123755 | P46109 | P46109 |
| Arginine--tRNA ligase, cytoplasmic | RARS | 0.0198988 | -0.012921 | P54136;P54136-2 | P54136;P54136-2 |
| Ubiquitin-conjugating enzyme E2 variant 1 | UBE2V1 | 0.0238831 | -0.01577 | Q13404;Q13404-8;Q13404-7;Q13404-2;Q13404-1;Q13404-6 | Q13404;Q13404-8;Q13404-7;Q13404-2;Q13404-1;Q1340 |
| Voltage-dependent anion-selective channel protein 1 | VDAC1 | 0.0105265 | -0.0165717 | P21796 | P21796 |

| | | | | | |
|---|---|---|---|---|---|
| Quinone oxidoreductase | CRYZ | 0.0132291 | -0.0166086 | Q08257;Q08257-3;Q08257-2 | Q08257;Q08257-3;Q08257-2 |
| Sorcin | SRI | 0.0110776 | -0.0176086 | P30626-2;P30626;P30626-3 | P30626-2;P30626;P30626-3 |
| ATPase ASNA1 | ASNA1 | 0.0100031 | -0.0185432 | O43681 | O43681 |
| Isocitrate dehydrogenase [NADP] cytoplasmic | IDH1 | 0.013305 | -0.0187829 | O75874 | O75874 |
| HLA class I histocompatibility antigen, Cw-12 alpha chain;HLA class I histocompatibility antigen, Cw-15 alpha cha | HLA-C | 0.0167655 | -0.0219765 | P30508;Q07000;P30510;P30505;P30504;Q9TNN7;P30501;P04222;C | P30508;Q07000;P30510;P30505;P30504;Q9TNN7;P3050 |
| Cathepsin Z | CTSZ | 0.0197726 | -0.0225487 | Q9UBR2 | Q9UBR2 |
| Ephrin type-A receptor 2 | EPHA2 | 0.056737 | -0.0235678 | P29317;P29317-2;Q15375-2;Q15375-4;Q15375;P54756-3;P29320;P | P29317 |
| Ras GTPase-activating-like protein IQGAP1 | IQGAP1 | 0.0296089 | -0.0252361 | P46940 | P46940 |
| Serine/threonine-protein kinase MARK2;MAP/microtubule affinity-regulating kinase 3 | MARK2;MARK3 | 0.0482722 | -0.0289694 | Q7KZI7-10;Q7KZI7-13;Q7KZI7-7;Q7KZI7-2;Q7KZI7-15;Q7KZI7-5;Q7K | Q7KZI7-10;Q7KZI7-13;Q7KZI7-7;Q7KZI7-2;Q7KZI7-15;Q7K |
| Tyrosine-protein kinase CSK | CSK | 0.0162108 | -0.0302785 | P41240;P42679-3;P42679;P42679-2 | P41240 |
| Cyclin-dependent kinase 1 | CDK1 | 0.0694706 | -0.0304661 | P06493;P06493-2 | P06493;P06493-2 |
| Phosphomannomutase 2 | PMM2 | 0.0667757 | -0.0314299 | O15305;O15305-2 | O15305;O15305-2 |
| Protein disulfide-isomerase A6 | PDIA6 | 0.0273733 | -0.0326271 | Q15084-3;Q15084;Q15084-4;Q15084-5;Q15084-2 | Q15084-3;Q15084;Q15084-4;Q15084-5;Q15084-2 |
| Macrophage migration inhibitory factor | MIF | 0.0163789 | -0.0327244 | P14174 | P14174 |
| Latent-transforming growth factor beta-binding protein 1 | LTBP1 | 0.032507 | -0.0341307 | Q14766;Q14766-4;Q14766-2;Q14766-5;Q14766-3 | Q14766;Q14766-4;Q14766-2;Q14766-5;Q14766-3 |
| Calcineurin B homologous protein 1 | CHP1 | 0.0149655 | -0.0352643 | Q99653 | Q99653 |
| Eukaryotic translation initiation factor 3 subunit H | EIF3H | 0.0497843 | -0.0367616 | O15372 | O15372 |
| Glucosamine-6-phosphate isomerase 1 | GNPDA1 | 0.0401381 | -0.0368137 | P46926;P46926-2 | P46926;P46926-2 |
| Translationally-controlled tumor protein | TPT1 | 0.0229595 | -0.0376835 | P13693;P13693-2;Q56UQ5 | P13693;P13693-2 |
| N-acetyllactosaminide beta-1,3-N-acetylglucosaminyltransferase 2 | B3GNT2 | 0.0523018 | -0.0381298 | Q9NY97-2;Q9NY97 | Q9NY97-2;Q9NY97 |
| Sorbitol dehydrogenase | SORD | 0.136878 | -0.0387319 | Q00796;Q00796-2 | Q00796 |
| Ephrin type-B receptor 4 | EPHB4 | 0.0473069 | -0.0395819 | P54760;P54760-2;P54760-4;P54760-3 | P54760;P54760-2;P54760-4 |
| N-acetylgalactosamine-6-sulfatase | GALNS | 0.0228643 | -0.0400855 | P34059 | P34059 |
| Transportin-1 | TNPO1 | 0.0343255 | -0.0407092 | Q92973-2;Q92973;Q92973-3 | Q92973-2;Q92973;Q92973-3 |
| TBC1 domain family member 15 | TBC1D15 | 0.0168556 | -0.040954 | Q8TC07-2;Q8TC07-3;Q8TC07 | Q8TC07-2;Q8TC07-3;Q8TC07 |
| Ubiquitin-conjugating enzyme E2 G1;Ubiquitin-conjugating enzyme E2 G1, N-terminally processed | UBE2G1 | 0.0792721 | -0.0419013 | P62253 | P62253 |
| Transmembrane protein 2 | TMEM2 | 0.019152 | -0.0423349 | Q9UHN6-2;Q9UHN6 | Q9UHN6-2;Q9UHN6 |
| Transgelin-2 | TAGLN2 | 0.149451 | -0.0426375 | P37802;P37802-2 | P37802;P37802-2 |
| ATP-dependent 6-phosphofructokinase, muscle type | PFKM | 0.0325723 | -0.0435047 | P08237-2;P08237;P08237-3 | P08237-2;P08237;P08237-3 |
| Probable serine carboxypeptidase CPVL | CPVL | 0.0222347 | -0.0441551 | Q9H3G5 | Q9H3G5 |
| High mobility group protein B1 | HMGB1 | 0.0892835 | -0.045627 | P09429;B2RPK0 | P09429 |
| Beta-1,4-galactosyltransferase 1;Lactose synthase A protein;N-acetyllactosamine synthase;Beta-N-acetylglucos | B4GALT1 | 0.0244766 | -0.0465546 | P15291-2;P15291 | P15291-2;P15291 |
| Pyruvate dehydrogenase E1 component subunit beta, mitochondrial | PDHB | 0.0216667 | -0.0475063 | P11177-2;P11177;P11177-3 | P11177-2;P11177;P11177-3 |
| Beta-galactosidase | GLB1 | 0.122901 | -0.0475763 | P16278-3;P16278;P16278-2 | P16278-3;P16278;P16278-2 |
| E3 ubiquitin-protein ligase TRIM11 | TRIM11 | 0.111155 | -0.0534223 | Q96F44-3;Q96F44;Q96F44-2;Q6AZZ1-2;O00481-4;Q6AZZ1;O00481;( | Q96F44-3;Q96F44;Q96F44-2 |
| Latent-transforming growth factor beta-binding protein 2 | LTBP2 | 0.0474333 | -0.0545692 | Q14767 | Q14767 |
| Fibulin-2 | FBLN2 | 0.0470005 | -0.0561848 | P98095;P98095-2 | P98095;P98095-2 |
| Myosin light polypeptide 6 | MYL6 | 0.0586707 | -0.0572643 | P60660-2 | P60660-2 |
| Collagen alpha-1(I) chain | COL1A1 | 0.0654295 | -0.0616811 | P02452;CON__Q862S4 | P02452 |
| Glucose-6-phosphate isomerase | GPI | 0.0640146 | -0.0618877 | P06744;P06744-2;Q8N196 | P06744;P06744-2 |
| Disks large homolog 3 | DLG3 | 0.0524377 | -0.0625807 | Q92796-3;Q92796-2;Q92796 | Q92796-3;Q92796-2;Q92796 |
| Ras suppressor protein 1 | RSU1 | 0.0420956 | -0.0630029 | Q15404;Q15404-2 | Q15404;Q15404-2 |
| V-type proton ATPase subunit d 1 | ATP6V0D1 | 0.0402936 | -0.0630849 | P61421 | P61421 |
| Proteasome subunit beta type-5 | PSMB5 | 0.0315219 | -0.0642802 | P28074;P28074-2 | P28074 |
| Hsp90 co-chaperone Cdc37;Hsp90 co-chaperone Cdc37, N-terminally processed | CDC37 | 0.04586 | -0.0643813 | Q16543 | Q16543 |
| Gamma-glutamyl hydrolase | GGH | 0.0517133 | -0.0644449 | Q92820 | Q92820 |
| Activator of 90 kDa heat shock protein ATPase homolog 1 | AHSA1 | 0.0609818 | -0.0646756 | O95433;O95433-2 | O95433;O95433-2 |
| Iron/zinc purple acid phosphatase-like protein | PAPL | 0.0466121 | -0.0649331 | Q6ZNF0 | Q6ZNF0 |
| Heterogeneous nuclear ribonucleoprotein A1;Heterogeneous nuclear ribonucleoprotein A1, N-terminally proce | HNRNPA1 | 0.0893863 | -0.0698853 | P09651;P09651-2;P09651-3 | P09651;P09651-2;P09651-3 |
| Succinyl-CoA ligase [ADP-forming] subunit beta, mitochondrial | SUCLA2 | 0.11265 | -0.0705751 | Q9P2R7-2;Q9P2R7 | Q9P2R7-2;Q9P2R7 |
| Netrin-4 | NTN4 | 0.0291611 | -0.0733058 | Q9HB63-3;Q9HB63-2;Q9HB63 | Q9HB63-3;Q9HB63-2;Q9HB63 |
| STE20-like serine/threonine-protein kinase | SLK | 0.0292607 | -0.0734227 | Q9H2G2;Q9H2G2-2 | Q9H2G2;Q9H2G2-2 |
| Phosphatidylethanolamine-binding protein 1;Hippocampal cholinergic neurostimulating peptide | PEBP1 | 0.0663259 | -0.0742505 | P30086 | P30086 |
| CD276 antigen | CD276 | 0.130443 | -0.0773627 | Q5ZPR3-4;Q5ZPR3;Q5ZPR3-3;Q5ZPR3-2 | Q5ZPR3-4;Q5ZPR3;Q5ZPR3-3;Q5ZPR3-2 |
| Myristoylated alanine-rich C-kinase substrate | MARCKS | 0.0503375 | -0.0779324 | P29966 | P29966 |
| Ly6/PLAUR domain-containing protein 3 | LYPD3 | 0.101106 | -0.0786463 | O95274 | O95274 |
| Thioredoxin domain-containing protein 16 | TXNDC16 | 0.0347154 | -0.0788472 | Q9P2K2 | Q9P2K2 |
| Tyrosine-protein phosphatase non-receptor type 23 | PTPN23 | 0.107464 | -0.0795752 | Q9H3S7 | Q9H3S7 |
| Proteasome subunit beta type-3 | PSMB3 | 0.0551889 | -0.0815722 | P49720 | P49720 |
| Cyclin-dependent kinase 6 | CDK6 | 0.109563 | -0.0818806 | Q00534;P11802-2;P11802;Q96Q40-2;Q96Q40-4;Q96Q40-3;O94921- | Q00534 |
| Nidogen-2 | NID2 | 0.0706181 | -0.0824617 | Q14112-2 | Q14112-2 |
| Fructose-bisphosphate aldolase C | ALDOC | 0.115299 | -0.0830059 | P09972 | P09972 |
| Enoyl-CoA hydratase, mitochondrial | ECHS1 | 0.0959476 | -0.0831051 | P30084 | P30084 |
| 72 kDa type IV collagenase;PEX | MMP2 | 0.114989 | -0.0845966 | P08253;P08253-3;P08253-2 | P08253;P08253-3;P08253-2 |
| Platelet-activating factor acetylhydrolase IB subunit gamma | PAFAH1B3 | 0.300248 | -0.084823 | Q15102 | Q15102 |
| Carbonic anhydrase 2 | CA2 | 0.103916 | -0.0852769 | P00918 | P00918 |
| Desmocollin-2 | DSC2 | 0.13329 | -0.0854715 | Q02487;Q02487-2 | Q02487;Q02487-2 |
| Threonine--tRNA ligase, cytoplasmic | TARS | 0.26952 | -0.0858854 | P26639;P26639-2;A2RTX5-2;A2RTX5 | P26639;P26639-2 |
| ATP-dependent RNA helicase DHX36 | DHX36 | 0.342636 | -0.086141 | Q9H2U1-2;Q9H2U1;Q9H2U1-3 | Q9H2U1-2;Q9H2U1;Q9H2U1-3 |
| Peptidyl-prolyl cis-trans isomerase B | PPIB | 0.0945592 | -0.0882142 | P23284 | P23284 |
| L-lactate dehydrogenase B chain | LDHB | 0.0920645 | -0.0897026 | P07195 | P07195 |
| Proteasome subunit beta type-6 | PSMB6 | 0.0725745 | -0.0902869 | P28072 | P28072 |
| Profilin-1 | PFN1 | 0.201418 | -0.0905094 | P07737 | P07737 |
| Proteasome activator complex subunit 1 | PSME1 | 0.231154 | -0.0912132 | Q06323;Q06323-3;Q06323-2 | Q06323;Q06323-3;Q06323-2 |
| Myosin-9 | MYH9 | 0.0712164 | -0.091732 | P35579;P35579-2;Q9UKX2 | P35579;P35579-2 |
| DNA damage-binding protein 1 | DDB1 | 0.0737106 | -0.0920868 | Q16531;Q16531-2 | Q16531 |
| Malate dehydrogenase, mitochondrial | MDH2 | 0.067153 | -0.0930875 | P40926;P40926-2 | P40926;P40926-2 |
| Syntaxin-3 | STX3 | 0.0902897 | -0.0938721 | Q13277-2;Q13277-3;Q13277 | Q13277-2;Q13277-3;Q13277 |
| Sepiapterin reductase | SPR | 0.13814 | -0.0946592 | P35270 | P35270 |
| Protein lin-7 homolog C | LIN7C | 0.0637691 | -0.0950534 | Q9NUP9 | Q9NUP9 |

| | | | | | |
|---|---|---|---|---|---|
| Alcohol dehydrogenase class-3 | ADH5 | 0.316059 | -0.0962823 | P11766 | P11766 |
| Vacuolar protein sorting-associated protein VTA1 homolog | VTA1 | 0.19503 | -0.0963828 | Q9NP79;Q9NP79-2 | Q9NP79 |
| Adenosine deaminase | ADA | 0.356756 | -0.0965252 | P00813 | P00813 |
| EF-hand domain-containing protein D2 | EFHD2 | 0.161938 | -0.0966511 | Q96C19;Q9BUP0-2;Q9BUP0 | Q96C19 |
| Arrestin domain-containing protein 1 | ARRDC1 | 0.192716 | -0.0971661 | Q8N5I2 | Q8N5I2 |
| Sorting nexin-3 | SNX3 | 0.151784 | -0.0978076 | O60493;O60493-2;O60493-4;O60493-3 | O60493;O60493-2;O60493-4;O60493-3 |
| Myeloid-derived growth factor | MYDGF | 0.13821 | -0.0982049 | Q969H8 | Q969H8 |
| Mannose-6-phosphate isomerase | MPI | 0.14354 | -0.0996564 | P34949;P34949-2 | P34949;P34949-2 |
| cAMP-dependent protein kinase type II-alpha regulatory subunit | PRKAR2A | 0.127934 | -0.100096 | P13861-2;P13861 | P13861-2;P13861 |
| Ubiquitin-conjugating enzyme E2 variant 2 | UBE2V2 | 0.11623 | -0.100404 | Q15819 | Q15819 |
| Formin-binding protein 1-like | FNBP1L | 0.126789 | -0.102098 | Q5T0N5-3;Q5T0N5-4;Q5T0N5-2;Q5T0N5-5;Q5T0N5 | Q5T0N5-3;Q5T0N5-4;Q5T0N5-2;Q5T0N5-5;Q5T0N5 |
| Golgi apparatus protein 1 | GLG1 | 0.117813 | -0.102592 | Q92896;Q92896-2;Q92896-3 | Q92896;Q92896-2;Q92896-3 |
| Alpha-amylase 1;Pancreatic alpha-amylase;Alpha-amylase 2B | AMY1A;AMY2A;AMY2B | 0.131991 | -0.102725 | P04745;P04746;P19961;P19961-2;P04746-2 | P04745;P04746;P19961;P19961-2 |
| Single-stranded DNA-binding protein, mitochondrial | SSBP1 | 0.0567831 | -0.103849 | Q04837 | Q04837 |
| Proteasome subunit beta type-4 | PSMB4 | 0.0747893 | -0.104593 | P28070 | P28070 |
| ADP-ribosylation factor-like protein 2 | ARL2 | 0.0788591 | -0.105225 | P36404;P36404-2 | P36404;P36404-2 |
| Adenosylhomocysteinase | AHCY | 0.142893 | -0.105321 | P23526;P23526-2 | P23526;P23526-2 |
| Peptidase M20 domain-containing protein 2 | PM20D2 | 0.398525 | -0.105565 | Q8IYS1 | Q8IYS1 |
| Serine/threonine-protein phosphatase PP1-alpha catalytic subunit | PPP1CA | 0.299206 | -0.106082 | P62136;P62136-2;P62136-3 | P62136;P62136-2;P62136-3 |
| Follistatin-related protein 1 | FSTL1 | 0.113727 | -0.108865 | Q12841;Q12841-2 | Q12841;Q12841-2 |
| Ubiquitin-conjugating enzyme E2 D2;Ubiquitin-conjugating enzyme E2 D3 | UBE2D2;UBE2D3 | 0.0392156 | -0.111153 | P62837-2;P62837;P61077;P61077-2;P61077-3 | P62837-2;P62837;P61077;P61077-2;P61077-3 |
| Retinoic acid-induced protein 3 | GPRC5A | 0.153607 | -0.111632 | Q8NFJ5 | Q8NFJ5 |
| GDP-L-fucose synthase | TSTA3 | 0.0846318 | -0.112082 | Q13630 | Q13630 |
| Alpha-mannosidase 2 | MAN2A1 | 0.197216 | -0.112426 | Q16706 | Q16706 |
| Mannose-1-phosphate guanyltransferase beta | GMPPB | 0.099951 | -0.113031 | Q9Y5P6;Q9Y5P6-2 | Q9Y5P6;Q9Y5P6-2 |
| Collagen alpha-1(VI) chain | COL6A1 | 0.237889 | -0.113148 | P12109 | P12109 |
| Ubiquitin-conjugating enzyme E2 N | UBE2N | 0.218839 | -0.113168 | P61088 | P61088 |
| Dipeptidyl peptidase 1;Dipeptidyl peptidase 1 exclusion domain chain;Dipeptidyl peptidase 1 heavy chain;Dipep | CTSC | 0.0990355 | -0.11341 | P53634;P53634-2;P53634-3 | P53634 |
| Heterogeneous nuclear ribonucleoprotein D0 | HNRNPD | 0.289153 | -0.114325 | Q14103-3;Q14103;Q14103-4;Q14103-2 | Q14103-3;Q14103;Q14103-4;Q14103-2 |
| Retinoic acid receptor responder protein 1 | RARRES1 | 0.0474552 | -0.114407 | P49788;P49788-2 | P49788;P49788-2 |
| Protein tyrosine phosphatase type IVA 2;Protein tyrosine phosphatase type IVA 1 | PTP4A2;PTP4A1 | 0.179974 | -0.117512 | Q12974;Q12974-3;Q12974-4;Q12974-2;Q93096 | Q12974;Q12974-3;Q12974-4;Q12974-2;Q93096 |
| Cathepsin B;Cathepsin B light chain;Cathepsin B heavy chain | CTSB | 0.0928905 | -0.117535 | P07858 | P07858 |
| U6 snRNA-associated Sm-like protein LSm6 | LSM6 | 0.0634008 | -0.11899 | P62312 | P62312 |
| Ribonuclease inhibitor | RNH1 | 0.201223 | -0.11972 | P13489 | P13489 |
| Galectin-1 | LGALS1 | 0.19034 | -0.119766 | P09382 | P09382 |
| Fumarate hydratase, mitochondrial | FH | 0.115501 | -0.120413 | P07954-2;P07954 | P07954-2;P07954 |
| Dihydropyrimidinase-related protein 2 | DPYSL2 | 0.145295 | -0.120754 | Q16555;Q16555-2;Q14195;Q14194;Q14194-2 | Q16555;Q16555-2 |
| Cullin-3 | CUL3 | 0.203715 | -0.124672 | Q13618-2;Q13618;Q13618-3 | Q13618-2;Q13618;Q13618-3 |
| Cilia- and flagella-associated protein 20 | CFAP20 | 0.0703219 | -0.125089 | Q9Y6A4 | Q9Y6A4 |
| Eukaryotic translation initiation factor 3 subunit K | EIF3K | 0.315882 | -0.125217 | Q9UBQ5;Q9UBQ5-2 | Q9UBQ5;Q9UBQ5-2 |
| UTP--glucose-1-phosphate uridylyltransferase | UGP2 | 0.264142 | -0.126542 | Q16851 | Q16851 |
| BRO1 domain-containing protein BROX | BROX | 0.561829 | -0.1266 | Q5VW32;Q5VW32-2 | Q5VW32;Q5VW32-2 |
| Complement factor H | CFH | 0.117667 | -0.128556 | P08603;P08603-2;Q02985-2;Q02985;Q03591 | P08603;P08603-2 |
| Bifunctional purine biosynthesis protein PURH;Phosphoribosylaminoimidazolecarboxamide formyltransferase;I | ATIC | 0.153592 | -0.130058 | P31939;P31939-2 | P31939;P31939-2 |
| Sulfhydryl oxidase 1 | QSOX1 | 0.28206 | -0.130552 | O00391;O00391-2 | O00391;O00391-2 |
| Calpain-2 catalytic subunit | CAPN2 | 0.452774 | -0.132218 | P17655;P17655-2 | P17655;P17655-2 |
| Vacuolar protein-sorting-associated protein 36 | VPS36 | 0.0431115 | -0.1328 | Q86VN1;Q86VN1-2 | Q86VN1;Q86VN1-2 |
| Annexin A2;Putative annexin A2-like protein | ANXA2;ANXA2P2 | 0.230334 | -0.13663 | P07355;P07355-2;A6NMY6 | P07355;P07355-2;A6NMY6 |
| Adenylosuccinate synthetase isozyme 2 | ADSS | 0.355692 | -0.138625 | P30520 | P30520 |
| Calponin-2 | CNN2 | 0.399285 | -0.138865 | Q99439;Q99439-2 | Q99439;Q99439-2 |
| F-actin-capping protein subunit alpha-2 | CAPZA2 | 0.179903 | -0.13984 | P47755;P47755-2 | P47755;P47755-2 |
| Ragulator complex protein LAMTOR2 | LAMTOR2 | 0.154803 | -0.139946 | Q9Y2Q5;Q9Y2Q5-2;Q9Y2Q5-3 | Q9Y2Q5;Q9Y2Q5-2 |
| Peptidyl-prolyl cis-trans isomerase FKBP1A | FKBP1A | 0.0727481 | -0.141279 | P62942 | P62942 |
| Transitional endoplasmic reticulum ATPase | VCP | 0.299782 | -0.14168 | P55072 | P55072 |
| Cofilin-2 | CFL2 | 0.198988 | -0.142396 | Q9Y281;Q9Y281-3 | Q9Y281;Q9Y281-3 |
| Chloride intracellular channel protein 1 | CLIC1 | 0.162168 | -0.14269 | O00299 | O00299 |
| IST1 homolog | IST1 | 0.270045 | -0.142935 | P53990-2;P53990;P53990-4;P53990-3;P53990-5;P53990-6 | P53990-2;P53990;P53990-4;P53990-3;P53990-5 |
| Tropomyosin alpha-4 chain | TPM4 | 0.146574 | -0.143541 | P67936 | P67936 |
| Fibulin-1 | FBLN1 | 0.183025 | -0.145398 | P23142 | P23142 |
| V-type proton ATPase subunit H | ATP6V1H | 0.151086 | -0.147735 | Q9UI12-2;Q9UI12 | Q9UI12-2;Q9UI12 |
| Cadherin-2 | CDH2 | 0.227051 | -0.149472 | P19022 | P19022 |
| Trifunctional purine biosynthetic protein adenosine-3;Phosphoribosylamine--glycine ligase;Phosphoribosylform | GART | 0.326573 | -0.151313 | P22102;P22102-2 | P22102;P22102-2 |
| COP9 signalosome complex subunit 8 | COPS8 | 0.440379 | -0.151669 | Q99627-2;Q99627 | Q99627-2;Q99627 |
| Peptidyl-prolyl cis-trans isomerase NIMA-interacting 4 | PIN4 | 0.145984 | -0.151813 | Q9Y237;Q9Y237-2;Q9Y237-3 | Q9Y237;Q9Y237-2 |
| Alpha/beta hydrolase domain-containing protein 14B | ABHD14B | 0.218453 | -0.154196 | Q96IU4;Q96IU4-2 | Q96IU4 |
| ATP-dependent 6-phosphofructokinase, platelet type | PFKP | 0.140906 | -0.154421 | Q01813;Q01813-2 | Q01813;Q01813-2 |
| Proteasome subunit alpha type-6 | PSMA6 | 0.123797 | -0.157444 | P60900;P60900-2;P60900-3 | P60900;P60900-2;P60900-3 |
| Inosine-5-monophosphate dehydrogenase 2 | IMPDH2 | 0.157309 | -0.16055 | P12268 | P12268 |
| Alpha-aminoadipic semialdehyde dehydrogenase | ALDH7A1 | 0.288826 | -0.163977 | P49419-2;P49419;P49419-4;P49419-3 | P49419-2;P49419;P49419-4 |
| 26S proteasome non-ATPase regulatory subunit 11 | PSMD11 | 0.594945 | -0.165002 | O00231;O00231-2 | O00231;O00231-2 |
| Rap1 GTPase-GDP dissociation stimulator 1 | RAP1GDS1 | 0.991651 | -0.16502 | P52306;P52306-5;P52306-4;P52306-2;P52306-6;P52306-3 | P52306;P52306-5;P52306-4;P52306-2;P52306-6;P52306- |
| Dual specificity mitogen-activated protein kinase kinase 1 | MAP2K1 | 0.260355 | -0.165955 | Q02750;Q02750-2 | Q02750;Q02750-2 |
| Lysophospholipase-like protein 1 | LYPLAL1 | 0.410991 | -0.166196 | Q5VWZ2-2;Q5VWZ2 | Q5VWZ2-2;Q5VWZ2 |
| CUB domain-containing protein 1 | CDCP1 | 0.25472 | -0.166538 | Q9H5V8;Q9H5V8-2 | Q9H5V8;Q9H5V8-2 |
| High mobility group protein B2 | HMGB2 | 0.178104 | -0.167384 | P26583 | P26583 |
| Elongation factor 1-gamma | EEF1G | 0.379704 | -0.168312 | P26641;P26641-2 | P26641;P26641-2 |
| Deoxycytidylate deaminase | DCTD | 0.241579 | -0.168884 | P32321;P32321-2 | P32321;P32321-2 |
| 26S proteasome non-ATPase regulatory subunit 7 | PSMD7 | 0.0994411 | -0.170282 | P51665 | P51665 |
| Heterogeneous nuclear ribonucleoprotein L | HNRNPL | 0.212117 | -0.17079 | P14866;P14866-2 | P14866;P14866-2 |
| 26S proteasome non-ATPase regulatory subunit 14 | PSMD14 | 0.127876 | -0.170911 | O00487 | O00487 |

| | | | | | | |
|---|---|---|---|---|---|---|
| Antithrombin-III | SERPINC1 | | 0.0903017 | -0.173222 | P01008 | P01008 |
| N-alpha-acetyltransferase 15, NatA auxiliary subunit | NAA15 | | 0.255974 | -0.174235 | Q9BXJ9;Q9BXJ9-4;Q6N069;Q6N069-5;Q6N069-4;Q6N069-3;Q6N069 | Q9BXJ9;Q9BXJ9-4 |
| Malate dehydrogenase, cytoplasmic | MDH1 | | 0.140901 | -0.178947 | P40925;P40925-3;P40925-2 | P40925;P40925-3;P40925-2 |
| ADP-ribose pyrophosphatase, mitochondrial | NUDT9 | | 0.185136 | -0.18002 | Q9BW91-2;Q9BW91 | Q9BW91-2;Q9BW91 |
| Poly(rC)-binding protein 2 | PCBP2 | | 0.345187 | -0.180333 | Q15366-3;Q15366-4;Q15366-6;Q15366-7;P57721-5;P57721-4;P5772 | Q15366-3;Q15366-4;Q15366-6;Q15366-7 |
| Rab GDP dissociation inhibitor beta | GDI2 | | 0.807804 | -0.181484 | P50395;P50395-2 | P50395;P50395-2 |
| cAMP-dependent protein kinase catalytic subunit alpha | PRKACA | | 0.173563 | -0.181955 | P17612-2;P17612;P22694-10;P22694-4;P22694-3;P22694;P22694-5; | P17612-2;P17612 |
| Glutaredoxin-3 | GLRX3 | | 0.364504 | -0.183675 | O76003 | O76003 |
| DCC-interacting protein 13-alpha | APPL1 | | 1.29719 | -0.184111 | Q9UKG1 | Q9UKG1 |
| Plexin-B2 | PLXNB2 | | 0.535485 | -0.1848 | O15031 | O15031 |
| Nascent polypeptide-associated complex subunit alpha;Nascent polypeptide-associated complex subunit alpha | NACA | | 0.299366 | -0.185064 | Q13765;E9PAV3-2;E9PAV3;Q9BZK3 | Q13765;E9PAV3-2;E9PAV3 |
| Thioredoxin domain-containing protein 5 | TXNDC5 | | 0.116624 | -0.187939 | Q8NBS9;Q8NBS9-2 | Q8NBS9;Q8NBS9-2 |
| Voltage-dependent anion-selective channel protein 2 | VDAC2 | | 0.132199 | -0.188971 | P45880-2;P45880;P45880-1 | P45880-2;P45880;P45880-1 |
| Phosphatidylinositol transfer protein beta isoform | PITPNB | | 0.182284 | -0.190083 | P48739-3;P48739;P48739-2 | P48739-3;P48739;P48739-2 |
| Heat shock protein HSP 90-beta | HSP90AB1 | | 0.482339 | -0.191728 | P08238;Q58FF7 | P08238 |
| Acyl-coenzyme A thioesterase 11 | ACOT11 | | 0.286879 | -0.195448 | Q8WXI4-2;Q8WXI4 | Q8WXI4-2;Q8WXI4 |
| UMP-CMP kinase | CMPK1 | | 0.362085 | -0.196836 | P30085;P30085-2 | P30085;P30085-2 |
| Phosphopantothenate--cysteine ligase | PPCS | | 0.52533 | -0.197824 | Q9HAB8;Q9HAB8-2 | Q9HAB8;Q9HAB8-2 |
| Eukaryotic translation initiation factor 2A;Eukaryotic translation initiation factor 2A, N-terminally processed | EIF2A | | 0.303001 | -0.198227 | Q9BY44;Q9BY44-3;Q9BY44-4;Q9BY44-2 | Q9BY44;Q9BY44-3;Q9BY44-4;Q9BY44-2 |
| Na(+)/H(+) exchange regulatory cofactor NHE-RF2 | SLC9A3R2 | | 0.22112 | -0.199115 | Q15599;Q15599-2;Q15599-3 | Q15599;Q15599-2;Q15599-3 |
| Alpha-1,3-mannosyl-glycoprotein 2-beta-N-acetylglucosaminyltransferase | MGAT1 | | 0.448574 | -0.201382 | P26572 | P26572 |
| Poliovirus receptor | PVR | | 0.175619 | -0.201981 | P15151-3;P15151-2;P15151-4;P15151 | P15151-3;P15151-2;P15151-4;P15151 |
| Phosphoglycerate kinase 1 | PGK1 | | 1.22813 | -0.202553 | P00558;P00558-2 | P00558;P00558-2 |
| Alpha-centractin | ACTR1A | | 0.539577 | -0.202775 | P61163 | P61163 |
| Omega-amidase NIT2 | NIT2 | | 0.531905 | -0.203986 | Q9NQR4 | Q9NQR4 |
| Roundabout homolog 1 | ROBO1 | | 0.578097 | -0.204422 | Q9Y6N7-6;Q9Y6N7-5;Q9Y6N7-4;Q9Y6N7-3;Q9Y6N7;Q9Y6N7-2;Q9H | Q9Y6N7-6;Q9Y6N7-5;Q9Y6N7-4;Q9Y6N7-3;Q9Y6N7;Q9Y |
| Heme-binding protein 1 | HEBP1 | | 0.772396 | -0.204582 | Q9NRV9 | Q9NRV9 |
| Rho GTPase-activating protein 1 | ARHGAP1 | | 0.43262 | -0.2056 | Q07960 | Q07960 |
| Trifunctional enzyme subunit beta, mitochondrial;3-ketoacyl-CoA thiolase | HADHB | | 0.100993 | -0.20609 | P55084-2;P55084 | P55084-2;P55084 |
| MAGUK p55 subfamily member 5 | MPP5 | | 0.183524 | -0.206291 | Q8N3R9-2;Q8N3R9 | Q8N3R9-2;Q8N3R9 |
| ATP-dependent 6-phosphofructokinase, liver type | PFKL | | 0.37872 | -0.206867 | P17858;P17858-2 | P17858;P17858-2 |
| Stress-70 protein, mitochondrial | HSPA9 | | 0.509333 | -0.20747 | P38646 | P38646 |
| Ubiquitin thioesterase OTUB1 | OTUB1 | | 0.609801 | -0.209665 | Q96FW1;Q96FW1-2 | Q96FW1 |
| Alpha-1,6-mannosylglycoprotein 6-beta-N-acetylglucosaminyltransferase A | MGAT5 | | 0.485258 | -0.211583 | Q09328 | Q09328 |
| Coiled-coil domain-containing protein 25 | CCDC25 | | 0.18529 | -0.2116 | Q86WR0;Q86WR0-2 | Q86WR0;Q86WR0-2 |
| L-xylulose reductase | DCXR | | 0.579723 | -0.212271 | Q7Z4W1 | Q7Z4W1 |
| Carbonyl reductase [NADPH] 1 | CBR1 | | 0.12213 | -0.212684 | P16152;P16152-2 | P16152;P16152-2 |
| Mannan-binding lectin serine protease 1;Mannan-binding lectin serine protease 1 heavy chain;Mannan-binding | MASP1 | | 0.176586 | -0.213324 | P48740-2;P48740-3;P48740;P48740-4 | P48740-2;P48740-3;P48740;P48740-4 |
| AP-1 complex subunit sigma-1A | AP1S1 | | 0.22206 | -0.213684 | P61966;P61966-2 | P61966;P61966-2 |
| Disks large homolog 1 | DLG1 | | 0.922326 | -0.216496 | Q12959-4;Q12959-2 | Q12959-4;Q12959-2 |
| Nuclear mitotic apparatus protein 1 | NUMA1 | | 0.686639 | -0.217766 | Q14980;Q14980-2;Q14980-4;Q14980-3;Q14980-5 | Q14980;Q14980-2;Q14980-4;Q14980-3 |
| Integrin beta-4 | ITGB4 | | 0.388804 | -0.218004 | P16144-4;P16144-2;P16144-3;P16144;P16144-5 | P16144-4;P16144-2;P16144-3;P16144;P16144-5 |
| Importin subunit alpha-3 | KPNA4 | | 0.223161 | -0.218334 | O00629 | O00629 |
| Myotubularin-related protein 2 | MTMR2 | | 0.0867606 | -0.219688 | Q13614-2;Q13614 | Q13614-2;Q13614 |
| Protein disulfide-isomerase A3 | PDIA3 | | 0.144152 | -0.220242 | P30101 | P30101 |
| Elongation factor 1-alpha 1;Putative elongation factor 1-alpha-like 3 | EEF1A1;EEF1A1P5 | | 0.809872 | -0.220266 | P68104;Q5VTE0;P68104-2 | P68104;Q5VTE0;P68104-2 |
| Ubiquitin carboxyl-terminal hydrolase isozyme L3 | UCHL3 | | 0.405823 | -0.220319 | P15374 | P15374 |
| Glycogenin-1 | GYG1 | | 0.342468 | -0.221128 | P46976-2;P46976;P46976-3 | P46976-2;P46976;P46976-3 |
| Phospholipid transfer protein | PLTP | | 0.0900741 | -0.221155 | P55058;P55058-3;P55058-2 | P55058;P55058-3;P55058-2 |
| Isoleucine--tRNA ligase, cytoplasmic | IARS | | 0.242968 | -0.221399 | P41252 | P41252 |
| Protein disulfide-isomerase | P4HB | | 0.243163 | -0.221406 | P07237 | P07237 |
| Gamma-glutamylcyclotransferase | GGCT | | 0.231464 | -0.225386 | O75223;O75223-3;O75223-2;O75223-4 | O75223;O75223-3 |
| UDP-glucose 4-epimerase | GALE | | 0.365823 | -0.225534 | Q14376;Q14376-2 | Q14376;Q14376-2 |
| Polypyrimidine tract-binding protein 1 | PTBP1 | | 0.246343 | -0.227914 | P26599;P26599-2;P26599-3 | P26599;P26599-2;P26599-3 |
| Ubiquitin-like modifier-activating enzyme 1 | UBA1 | | 0.431711 | -0.228242 | P22314-2 | P22314-2 |
| Cytoplasmic FMR1-interacting protein 1 | CYFIP1 | | 0.298102 | -0.228553 | Q7L576;Q7L576-2;Q7L576-3 | Q7L576;Q7L576-2 |
| Reticulon-4 | RTN4 | | 0.146137 | -0.229971 | Q9NQC3-5;Q9NQC3;Q9NQC3-2;Q9NQC3-4;Q9NQC3-6 | Q9NQC3-5;Q9NQC3;Q9NQC3-2;Q9NQC3-4;Q9NQC3-6 |
| Sorting nexin-2 | SNX2 | | 0.198331 | -0.233245 | O60749;O60749-2 | O60749;O60749-2 |
| Acyl-CoA-binding protein | DBI | | 0.439614 | -0.235281 | P07108;P07108-3;P07108-2;P07108-4;P07108-5;P07108-6 | P07108;P07108-3;P07108-2;P07108-4;P07108-5 |
| 26S proteasome non-ATPase regulatory subunit 1 | PSMD1 | | 0.452283 | -0.236177 | Q99460;Q99460-2 | Q99460;Q99460-2 |
| Coronin-1B | CORO1B | | 0.682157 | -0.236337 | Q9BR76 | Q9BR76 |
| Serine/threonine-protein phosphatase PP1-beta catalytic subunit | PPP1CB | | 0.236257 | -0.237382 | P62140 | P62140 |
| 3-ketoacyl-CoA thiolase, mitochondrial | ACAA2 | | 0.215311 | -0.23774 | P42765 | P42765 |
| Pyruvate kinase PKM | PKM | | 0.111396 | -0.239225 | P14618-2 | P14618-2 |
| Aspartate aminotransferase, cytoplasmic | GOT1 | | 0.20757 | -0.239251 | P17174;P17174-2 | P17174;P17174-2 |
| Latent-transforming growth factor beta-binding protein 3 | LTBP3 | | 0.352772 | -0.239738 | Q9NS15 | Q9NS15 |
| Glutathione S-transferase P | GSTP1 | | 0.749817 | -0.240639 | P09211 | P09211 |
| Syntaxin-binding protein 3 | STXBP3 | | 0.173996 | -0.243588 | O00186 | O00186 |
| Paired amphipathic helix protein Sin3a | SIN3A | | 0.200751 | -0.243707 | Q96ST3;O75182-2 | Q96ST3 |
| Rab3 GTPase-activating protein catalytic subunit | RAB3GAP1 | | 0.49118 | -0.244073 | Q15042;Q15042-3;Q15042-4 | Q15042;Q15042-3;Q15042-4 |
| COP9 signalosome complex subunit 6 | COPS6 | | 0.394766 | -0.244788 | Q7L5N1 | Q7L5N1 |
| E3 ubiquitin-protein ligase HUWE1 | HUWE1 | | 0.412856 | -0.247572 | Q7Z6Z7-3;Q7Z6Z7;Q7Z6Z7-2;A5YM72-2;A5YM72;A5YM72-4;A5YM7 | Q7Z6Z7-3;Q7Z6Z7;Q7Z6Z7-2 |
| Elongation factor 2 | EEF2 | + | 1.30924 | -0.249112 | P13639 | P13639 |
| ADP-ribosylation factor 4 | ARF4 | | 0.422382 | -0.250187 | P18085 | P18085 |
| Aldehyde dehydrogenase family 16 member A1 | ALDH16A1 | | 0.32072 | -0.254948 | Q8IZ83-3;Q8IZ83;Q8IZ83-2 | Q8IZ83-3;Q8IZ83 |
| Guanine nucleotide-binding protein subunit beta-2-like 1;Guanine nucleotide-binding protein subunit beta-2-lik | GNB2L1 | | 0.775644 | -0.256065 | P63244 | P63244 |
| UPF0587 protein C1orf123 | C1orf123 | | 0.319396 | -0.256297 | Q9NWV4 | Q9NWV4 |
| Heat shock protein 75 kDa, mitochondrial | TRAP1 | | 1.1931 | -0.257607 | Q12931-2;Q12931 | Q12931-2;Q12931 |
| Fatty acid synthase;[Acyl-carrier-protein] S-acetyltransferase;[Acyl-carrier-protein] S-malonyltransferase;3-oxoa | FASN | + | 1.46434 | -0.259537 | P49327 | P49327 |
| 26S protease regulatory subunit 10B | PSMC6 | | 0.373333 | -0.260369 | P62333 | P62333 |

| | | | | | | |
|---|---|---|---|---|---|---|
| Serine/threonine-protein kinase OSR1 | OXSR1 | | 0.21055 | -0.260539 | O95747 | O95747 |
| Dipeptidyl peptidase 3 | DPP3 | | 0.598377 | -0.261316 | Q9NY33;Q9NY33-4;Q9NY33-2 | Q9NY33;Q9NY33-4 |
| Dynamin-2 | DNM2 | | 0.174508 | -0.26154 | P50570-2;P50570-5;P50570;P50570-3;P50570-4;Q9UQ16-2;Q9UQ1 | P50570-2;P50570-5;P50570;P50570-3;P50570-4 |
| Thiopurine S-methyltransferase | TPMT | | 0.118171 | -0.262091 | P51580 | P51580 |
| Desmoplakin | DSP | | 0.226368 | -0.263187 | P15924;P15924-3;P15924-2 | P15924;P15924-3;P15924-2 |
| 26S proteasome non-ATPase regulatory subunit 13 | PSMD13 | | 0.389718 | -0.263575 | Q9UNM6;Q9UNM6-2 | Q9UNM6;Q9UNM6-2 |
| Citrate synthase, mitochondrial | CS | | 0.186304 | -0.263676 | O75390 | O75390 |
| Actin-related protein 2/3 complex subunit 5-like protein | ARPC5L | | 0.275136 | -0.264861 | Q9BPX5 | Q9BPX5 |
| CLIP-associating protein 1 | CLASP1 | | 0.251282 | -0.264967 | Q7Z460-2;Q7Z460-4;Q7Z460-5;Q7Z460-3;Q7Z460 | Q7Z460-2;Q7Z460-4;Q7Z460-5;Q7Z460-3;Q7Z460 |
| Glutamine--fructose-6-phosphate aminotransferase [isomerizing] 1 | GFPT1 | | 0.198862 | -0.265377 | Q06210-2;Q06210 | Q06210-2;Q06210 |
| Junction plakoglobin | JUP | | 0.435528 | -0.268123 | P14923 | P14923 |
| Anthrax toxin receptor 1 | ANTXR1 | | 0.276138 | -0.270689 | Q9H6X2-4;Q9H6X2-6;Q9H6X2-2;Q9H6X2-5;Q9H6X2;Q9H6X2-3 | Q9H6X2-4;Q9H6X2-6;Q9H6X2-2;Q9H6X2-5;Q9H6X2;Q9H |
| Melanotransferrin | MFI2 | | 0.337452 | -0.270927 | P08582;P08582-2 | P08582 |
| Nck-associated protein 1 | NCKAP1 | | 0.421468 | -0.271271 | Q9Y2A7;Q9Y2A7-2;P55160 | Q9Y2A7;Q9Y2A7-2 |
| Beta-1,4-galactosyltransferase 5 | B4GALT5 | | 0.135995 | -0.271808 | O43286;Q9UBX8-2;Q9UBX8 | O43286 |
| Structural maintenance of chromosomes protein 3 | SMC3 | | 0.358122 | -0.272345 | Q9UQE7 | Q9UQE7 |
| Isoamyl acetate-hydrolyzing esterase 1 homolog | IAH1 | | 0.387299 | -0.272559 | Q2TAA2;Q2TAA2-2 | Q2TAA2;Q2TAA2-2 |
| Structural maintenance of chromosomes protein 2 | SMC2 | | 0.700781 | -0.27637 | O95347;O95347-2 | O95347;O95347-2 |
| GDP-mannose 4,6 dehydratase | GMDS | | 0.560355 | -0.277512 | O60547-2;O60547 | O60547-2;O60547 |
| CD59 glycoprotein | CD59 | | 0.126014 | -0.277788 | P13987 | P13987 |
| ATP-citrate synthase | ACLY | | 1.13125 | -0.27958 | P53396;P53396-2;P53396-3 | P53396;P53396-2;P53396-3 |
| Bifunctional methylenetetrahydrofolate dehydrogenase/cyclohydrolase, mitochondrial;NAD-dependent methyl | MTHFD2 | | 0.149208 | -0.280896 | P13995-2;P13995 | P13995-2;P13995 |
| Destrin | DSTN | | 0.332301 | -0.282995 | P60981;P60981-2 | P60981;P60981-2 |
| Filamin-A | FLNA | + | 1.87645 | -0.284231 | P21333-2;P21333 | P21333-2;P21333 |
| Platelet-activating factor acetylhydrolase IB subunit alpha | PAFAH1B1 | | 0.766044 | -0.285187 | P43034;P43034-2 | P43034 |
| Epidermal growth factor receptor kinase substrate 8-like protein 2 | EPS8L2 | | 0.129766 | -0.285203 | Q9H6S3;Q9H6S3-3;Q9H6S3-2 | Q9H6S3;Q9H6S3-3;Q9H6S3-2 |
| NKG2D ligand 2 | ULBP2 | | 0.469843 | -0.287561 | Q9BZM5;Q6H3X3-2;Q5VY80;Q6H3X3 | Q9BZM5 |
| Adapter molecule crk | CRK | | 0.508728 | -0.289033 | P46108;P46108-2 | P46108;P46108-2 |
| Protein FAM49B | FAM49B | | 0.319975 | -0.291393 | Q9NUQ9;Q9NUQ9-2 | Q9NUQ9 |
| DnaJ homolog subfamily B member 6 | DNAJB6 | | 0.148881 | -0.291978 | O75190-3;O75190;O75190-2;Q8WWF6;O75190-4;Q7Z6W7 | O75190-3;O75190;O75190-2 |
| Endothelial protein C receptor | PROCR | | 0.408419 | -0.294839 | Q9UNN8 | Q9UNN8 |
| Vacuolar protein sorting-associated protein 35 | VPS35 | | 0.343044 | -0.296233 | Q96QK1 | Q96QK1 |
| Glutathione reductase, mitochondrial | GSR | | 0.562402 | -0.296391 | P00390-2;P00390;P00390-3;P00390-4;P00390-5 | P00390-2;P00390;P00390-3;P00390-4;P00390-5 |
| Peptidyl-prolyl cis-trans isomerase C | PPIC | | 0.195591 | -0.297729 | P45877 | P45877 |
| rRNA 2-O-methyltransferase fibrillarin | FBL | | 0.0648349 | -0.298199 | P22087 | P22087 |
| Granulins;Acrogranin;Paragranulin;Granulin-1;Granulin-2;Granulin-3;Granulin-4;Granulin-5;Granulin-6;Granulin | GRN | | 0.305221 | -0.29855 | P28799;P28799-2;P28799-3 | P28799;P28799-2;P28799-3 |
| Methionine--tRNA ligase, cytoplasmic | MARS | | 0.361887 | -0.298562 | P56192;P56192-2 | P56192;P56192-2 |
| Nucleoside diphosphate kinase B | NME2 | | 0.401099 | -0.299028 | P22392-2;P22392 | P22392-2;P22392 |
| Serpin B5 | SERPINB5 | | 0.346861 | -0.299819 | P36952;P36952-2 | P36952;P36952-2 |
| Delta-aminolevulinic acid dehydratase | ALAD | | 0.287462 | -0.300467 | P13716;P13716-2 | P13716;P13716-2 |
| Brain-specific serine protease 4 | PRSS22 | | 0.473607 | -0.300538 | Q9GZN4 | Q9GZN4 |
| Secretory carrier-associated membrane protein 3 | SCAMP3 | | 1.07691 | -0.300573 | O14828;O14828-2 | O14828;O14828-2 |
| Endonuclease domain-containing 1 protein | ENDOD1 | | 0.399033 | -0.301662 | O94919 | O94919 |
| ADP-sugar pyrophosphatase | NUDT5 | | 0.715879 | -0.302011 | Q9UKK9 | Q9UKK9 |
| 3(2),5-bisphosphate nucleotidase 1 | BPNT1 | | 0.518267 | -0.30342 | O95861;O95861-2;O95861-4 | O95861;O95861-2;O95861-4 |
| Vacuolar protein sorting-associated protein 26A | VPS26A | | 0.756137 | -0.303451 | O75436;O75436-2 | O75436;O75436-2 |
| C-Jun-amino-terminal kinase-interacting protein 4 | SPAG9 | | 0.367282 | -0.303878 | O60271;O60271-4;O60271-2;O60271-9;O60271-3;O60271-5;Q9UPT | O60271;O60271-4;O60271-2;O60271-9;O60271-3;O6027 |
| Twisted gastrulation protein homolog 1 | TWSG1 | | 0.210202 | -0.305577 | Q9GZX9;Q9GZX9-2 | Q9GZX9 |
| Membrane cofactor protein | CD46 | | 0.181589 | -0.305786 | P15529-7;P15529-4;P15529-6;P15529-3;P15529-5;P15529-2;P1552 | P15529-7;P15529-4;P15529-6;P15529-3;P15529-5;P1552 |
| Proteasome subunit beta type-1 | PSMB1 | | 0.181578 | -0.306164 | P20618 | P20618 |
| CCR4-NOT transcription complex subunit 1 | CNOT1 | | 0.39368 | -0.306754 | A5YKK6;A5YKK6-2;A5YKK6-3;A5YKK6-4 | A5YKK6;A5YKK6-2;A5YKK6-3;A5YKK6-4 |
| SH3 domain-binding glutamic acid-rich-like protein | SH3BGRL | | 0.365437 | -0.308285 | O75368 | O75368 |
| Adenylate kinase isoenzyme 1 | AK1 | | 0.364421 | -0.308584 | P00568;Q9Y6K8-2;Q9Y6K8-3;Q9Y6K8 | P00568 |
| Angiopoietin-related protein 4 | ANGPTL4 | | 0.31626 | -0.30934 | Q9BY76;Q9BY76-3;Q9BY76-2 | Q9BY76;Q9BY76-3;Q9BY76-2 |
| Talin-1 | TLN1 | | 0.522191 | -0.309602 | Q9Y490 | Q9Y490 |
| Mitogen-activated protein kinase 1 | MAPK1 | | 0.244845 | -0.310625 | P28482-2;P28482 | P28482-2;P28482 |
| Phosphoglucomutase-1 | PGM1 | | 0.621735 | -0.310791 | P36871;P36871-2;P36871-3 | P36871;P36871-2;P36871-3 |
| Xaa-Pro dipeptidase | PEPD | | 1.0453 | -0.311717 | P12955;P12955-2;P12955-3 | P12955;P12955-2;P12955-3 |
| Peptidyl-prolyl cis-trans isomerase FKBP4;Peptidyl-prolyl cis-trans isomerase FKBP4, N-terminally processed | FKBP4 | | 1.18145 | -0.312221 | Q02790 | Q02790 |
| S-formylglutathione hydrolase | ESD | | 0.540856 | -0.312338 | P10768 | P10768 |
| Protein-glutamine gamma-glutamyltransferase 2 | TGM2 | | 0.647481 | -0.318996 | P21980;P21980-2;P21980-3 | P21980;P21980-2 |
| Calreticulin | CALR | | 0.156264 | -0.319394 | P27797 | P27797 |
| Transforming growth factor beta-1;Latency-associated peptide | TGFB1 | | 0.439488 | -0.319672 | P01137 | P01137 |
| Serine/threonine-protein phosphatase 2A catalytic subunit alpha isoform | PPP2CA | | 0.77151 | -0.321784 | P67775;P67775-2 | P67775;P67775-2 |
| LanC-like protein 1 | LANCL1 | | 0.995162 | -0.321927 | O43813 | O43813 |
| Translocon-associated protein subunit delta | SSR4 | | 0.169332 | -0.32245 | P51571 | P51571 |
| CD166 antigen | ALCAM | | 0.679691 | -0.322507 | Q13740-2;Q13740;Q13740-4;Q13740-3 | Q13740-2;Q13740 |
| Myoferlin | MYOF | | 0.231813 | -0.3246 | Q9NZM1-6;Q9NZM1-3;Q9NZM1;Q9NZM1-5;Q9NZM1-2;Q9NZM1-7; | Q9NZM1-6;Q9NZM1-3;Q9NZM1;Q9NZM1-5;Q9NZM1-2 |
| Glucose-6-phosphate 1-dehydrogenase | G6PD | | 0.598411 | -0.326262 | P11413;P11413-2;P11413-3 | P11413;P11413-2;P11413-3 |
| Fructose-bisphosphate aldolase A | ALDOA | | 0.780735 | -0.330249 | P04075 | P04075 |
| Actin-related protein 2/3 complex subunit 4 | ARPC4 | + | 1.32129 | -0.331864 | P59998;P59998-3;P59998-4;P59998-2 | P59998;P59998-3;P59998-4;P59998-2 |
| Sorting nexin-12 | SNX12 | | 0.55796 | -0.333073 | Q9UMY4-2;Q9UMY4;Q9UMY4-3 | Q9UMY4-2;Q9UMY4;Q9UMY4-3 |
| Glyoxalase domain-containing protein 4 | GLOD4 | + | 1.34456 | -0.33526 | Q9HC38-2;Q9HC38;Q9HC38-3 | Q9HC38-2;Q9HC38;Q9HC38-3 |
| Aspartate--tRNA ligase, cytoplasmic | DARS | | 0.868958 | -0.335347 | P14868;P14868-2 | P14868;P14868-2 |
| Heat shock 70 kDa protein 4 | HSPA4 | | 1.24854 | -0.335568 | P34932;P34932-2 | P34932 |
| Tyrosine--tRNA ligase, cytoplasmic;Tyrosine--tRNA ligase, cytoplasmic, N-terminally processed | YARS | | 0.849735 | -0.335796 | P54577 | P54577 |
| Glycogen phosphorylase, liver form | PYGL | | 1.16306 | -0.33739 | P06737-2;P06737 | P06737-2;P06737 |
| Receptor-type tyrosine-protein phosphatase F | PTPRF | | 1.25523 | -0.33783 | P10586-2;P10586 | P10586-2;P10586 |
| Ras-related protein Rab-1A | RAB1A | | 0.166044 | -0.340892 | P62820;P62820-2;P62820-3 | P62820;P62820-2;P62820-3 |
| Actin-related protein 2/3 complex subunit 1A | ARPC1A | | 0.89235 | -0.340947 | Q92747;Q92747-2 | Q92747;Q92747-2 |

| | | | | | | |
|---|---|---|---|---|---|---|
| Cullin-associated NEDD8-dissociated protein 1 | CAND1 | + | 1.51945 | -0.342446 | Q86VP6;Q86VP6-2;Q86VP6-3;O75155-2;O75155 | Q86VP6;Q86VP6-2 |
| Phosphoglucomutase-2 | PGM2 | | 1.00191 | -0.342841 | Q96G03;Q96G03-2 | Q96G03 |
| Protein-L-isoaspartate(D-aspartate) O-methyltransferase | PCMT1 | | 0.373701 | -0.342934 | P22061;P22061-2 | P22061;P22061-2 |
| Neurogenic locus notch homolog protein 2;Notch 2 extracellular truncation;Notch 2 intracellular domain | NOTCH2 | | 0.288989 | -0.346342 | Q04721 | Q04721 |
| 6-phosphogluconolactonase | PGLS | | 0.85503 | -0.346358 | O95336 | O95336 |
| Thrombospondin type-1 domain-containing protein 4 | THSD4 | | 0.684236 | -0.347404 | Q6ZMP0;Q6ZMP0-4;Q6ZMP0-2 | Q6ZMP0;Q6ZMP0-4 |
| Serine/threonine-protein phosphatase 2A 65 kDa regulatory subunit A alpha isoform | PPP2R1A | + | 1.41296 | -0.347489 | P30153 | P30153 |
| Fibroblast growth factor-binding protein 1 | FGFBP1 | | 0.410499 | -0.347856 | Q14512 | Q14512 |
| Proteasome subunit alpha type-3 | PSMA3 | | 0.150152 | -0.348169 | P25788-2;P25788 | P25788-2;P25788 |
| Dipeptidyl peptidase 2 | DPP7 | | 0.239656 | -0.348207 | Q9UHL4 | Q9UHL4 |
| Proprotein convertase subtilisin/kexin type 5 | PCSK5 | | 0.186069 | -0.348291 | Q92824-2;Q92824 | Q92824-2;Q92824 |
| Extracellular serine/threonine protein kinase FAM20C | FAM20C | | 0.467781 | -0.348648 | Q8IXL6;Q8IXL6-2;Q96MK3 | Q8IXL6 |
| Podocalyxin | PODXL | | 0.12548 | -0.350081 | O00592-2;O00592 | O00592-2;O00592 |
| Lysine--tRNA ligase | KARS | | 0.676535 | -0.350387 | Q15046;Q15046-2 | Q15046;Q15046-2 |
| Radixin | RDX | | 1.27564 | -0.351782 | P35241;P35241-5;P35241-4;P35241-2 | P35241;P35241-5;P35241-4 |
| V-type proton ATPase subunit C 1 | ATP6V1C1 | | 0.740538 | -0.351858 | P21283 | P21283 |
| COP9 signalosome complex subunit 4 | COPS4 | + | 1.8884 | -0.355656 | Q9BT78;Q9BT78-2 | Q9BT78;Q9BT78-2 |
| Polypeptide N-acetylgalactosaminyltransferase 3 | GALNT3 | | 1.00666 | -0.357276 | Q14435;Q14435-2 | Q14435 |
| Putative adenosylhomocysteinase 2 | AHCYL1 | | 0.221848 | -0.357981 | O43865;O43865-2;Q96HN2-4;Q96HN2-3;Q96HN2-2;Q96HN2 | O43865;O43865-2 |
| Cystatin-A;Cystatin-A, N-terminally processed | CSTA | | 0.800858 | -0.359667 | P01040 | P01040 |
| NEDD8 | NEDD8 | | 0.104961 | -0.360248 | Q15843 | Q15843 |
| Xaa-Pro aminopeptidase 1 | XPNPEP1 | + | 2.83852 | -0.363378 | Q9NQW7-3;Q9NQW7;Q9NQW7-4;Q9NQW7-2 | Q9NQW7-3;Q9NQW7;Q9NQW7-4;Q9NQW7-2 |
| Alcohol dehydrogenase [NADP(+)] | AKR1A1 | | 0.357001 | -0.365993 | P14550 | P14550 |
| Prostaglandin reductase 1 | PTGR1 | | 0.936028 | -0.366514 | Q14914;Q14914-2 | Q14914;Q14914-2 |
| 7,8-dihydro-8-oxoguanine triphosphatase | NUDT1 | | 0.342167 | -0.366522 | P36639-2;P36639-4;P36639-3;P36639 | P36639-2;P36639-4;P36639-3;P36639 |
| Growth factor receptor-bound protein 2 | GRB2 | | 0.437803 | -0.366545 | P62993;P62993-2 | P62993;P62993-2 |
| Aminoacylase-1 | ACY1 | + | 1.36356 | -0.369913 | Q03154;Q03154-3;Q03154-4;Q03154-2 | Q03154;Q03154-3;Q03154-4;Q03154-2 |
| Cysteine and glycine-rich protein 1 | CSRP1 | + | 1.42011 | -0.371481 | P21291 | P21291 |
| Hepatocyte growth factor activator;Hepatocyte growth factor activator short chain;Hepatocyte growth factor a | HGFAC | | 0.165492 | -0.37181 | Q04756 | Q04756 |
| T-complex protein 1 subunit eta | CCT7 | + | 1.90238 | -0.372297 | Q99832;Q99832-3;Q99832-4;Q99832-2 | Q99832;Q99832-3;Q99832-4;Q99832-2 |
| Spectrin alpha chain, non-erythrocytic 1 | SPTAN1 | | 1.15338 | -0.374509 | Q13813 | Q13813 |
| Glutathione synthetase | GSS | | 0.431607 | -0.374633 | P48637;P48637-2 | P48637;P48637-2 |
| Cadherin-1;E-Cad/CTF1;E-Cad/CTF2;E-Cad/CTF3 | CDH1 | | 0.570226 | -0.375303 | P12830;P12830-2 | P12830;P12830-2 |
| Annexin A3 | ANXA3 | | 0.984762 | -0.377151 | P12429 | P12429 |
| Putative helicase MOV-10 | MOV10 | | 0.425719 | -0.37827 | Q9HCE1;Q9HCE1-2 | Q9HCE1;Q9HCE1-2 |
| Septin-11 | SEPT11 | | 0.578307 | -0.380753 | Q9NVA2;Q9NVA2-2 | Q9NVA2;Q9NVA2-2 |
| Myosin-14 | MYH14 | | 0.538221 | -0.381363 | Q7Z406;Q7Z406-6;Q7Z406-2;Q7Z406-5;Q7Z406-4 | Q7Z406;Q7Z406-6;Q7Z406-2;Q7Z406-5;Q7Z406-4 |
| Retinol-binding protein 4;Plasma retinol-binding protein(1-182);Plasma retinol-binding protein(1-181);Plasma re | RBP4 | | 0.0496661 | -0.381841 | P02753 | P02753 |
| Spectrin beta chain, non-erythrocytic 1 | SPTBN1 | | 0.453214 | -0.382428 | Q01082;Q01082-2 | Q01082;Q01082-2 |
| Aminopeptidase B | RNPEP | | 0.775208 | -0.382547 | Q9H4A4 | Q9H4A4 |
| Acylamino-acid-releasing enzyme | APEH | | 0.918641 | -0.384588 | P13798 | P13798 |
| NEDD8-conjugating enzyme Ubc12 | UBE2M | | 1.19511 | -0.392294 | P61081 | P61081 |
| Charged multivesicular body protein 1a | CHMP1A | | 0.128782 | -0.393323 | Q9HD42 | Q9HD42 |
| Alpha-soluble NSF attachment protein | NAPA | | 1.13799 | -0.394755 | P54920;Q9H115-3;Q9H115-2;Q9H115 | P54920 |
| Coatomer subunit alpha;Xenin;Proxenin | COPA | | 0.507372 | -0.394847 | P53621;P53621-2 | P53621;P53621-2 |
| Osteoclast-stimulating factor 1 | OSTF1 | | 0.221806 | -0.395777 | Q92882 | Q92882 |
| Leucine-rich alpha-2-glycoprotein | LRG1 | | 0.372715 | -0.395917 | P02750 | P02750 |
| Proteasome subunit beta type-2 | PSMB2 | | 0.268876 | -0.39657 | P49721 | P49721 |
| Glypican-6;Secreted glypican-6 | GPC6 | | 0.177455 | -0.396796 | Q9Y625;O75487-2;O75487 | Q9Y625 |
| Tumor necrosis factor receptor superfamily member 11B | TNFRSF11B | | 0.269733 | -0.398502 | O00300 | O00300 |
| Nectin-1 | PVRL1 | | 0.463379 | -0.400373 | Q15223;Q15223-3;Q15223-2 | Q15223;Q15223-3;Q15223-2 |
| Calpain small subunit 1 | CAPNS1 | | 0.571298 | -0.402213 | P04632;Q96L46 | P04632 |
| Synaptic vesicle membrane protein VAT-1 homolog | VAT1 | | 0.778228 | -0.403929 | Q99536;Q99536-3;Q99536-2 | Q99536;Q99536-3;Q99536-2 |
| Importin subunit beta-1 | KPNB1 | + | 2.23362 | -0.40469 | Q14974;Q14974-2 | Q14974;Q14974-2 |
| Delta(3,5)-Delta(2,4)-dienoyl-CoA isomerase, mitochondrial | ECH1 | | 0.241429 | -0.40527 | Q13011 | Q13011 |
| Tuftelin-interacting protein 11 | TFIP11 | | 0.216902 | -0.406042 | Q9UBB9;Q9UBB9-2 | Q9UBB9 |
| Protein S100-A6 | S100A6 | | 0.125247 | -0.406059 | P06703 | P06703 |
| Ezrin | EZR | | 0.804551 | -0.40606 | P15311 | P15311 |
| Proteasome subunit alpha type-4 | PSMA4 | | 0.49412 | -0.409369 | P25789;P25789-2 | P25789;P25789-2 |
| Mannosyl-oligosaccharide 1,2-alpha-mannosidase IA | MAN1A1 | + | 1.34877 | -0.409718 | P33908;P33908-2 | P33908;P33908-2 |
| Calcyclin-binding protein | CACYBP | + | 2.1888 | -0.410117 | Q9HB71;Q9HB71-3;Q9HB71-2 | Q9HB71;Q9HB71-3 |
| Kin of IRRE-like protein 1 | KIRREL | | 0.556591 | -0.410356 | Q96J84;Q96J84-2;Q96J84-3 | Q96J84;Q96J84-2;Q96J84-3 |
| Thioredoxin reductase 1, cytoplasmic | TXNRD1 | | 0.956987 | -0.410563 | Q16881-4;Q16881-3;Q16881 | Q16881-4;Q16881-3;Q16881 |
| Follistatin-related protein 3 | FSTL3 | | 0.307175 | -0.412599 | O95633-2;O95633 | O95633-2;O95633 |
| Ubiquitin-conjugating enzyme E2 K | UBE2K | + | 2.14688 | -0.413949 | P61086;P61086-2;P61086-3 | P61086;P61086-2;P61086-3 |
| Syntaxin-binding protein 2 | STXBP2 | | 0.546007 | -0.418629 | Q15833-2;Q15833;Q15833-3 | Q15833-2;Q15833;Q15833-3 |
| N-alpha-acetyltransferase 50 | NAA50 | | 0.393124 | -0.419142 | Q9GZZ1;Q9GZZ1-2 | Q9GZZ1;Q9GZZ1-2 |
| Complement C1r subcomponent-like protein | C1RL | | 0.220598 | -0.419575 | Q9NZP8 | Q9NZP8 |
| Electron transfer flavoprotein subunit alpha, mitochondrial | ETFA | | 0.466763 | -0.42139 | P13804;P13804-2 | P13804;P13804-2 |
| Abl interactor 1 | ABI1 | | 0.234174 | -0.421399 | Q8IZP0-10;Q8IZP0-8;Q8IZP0-7;Q8IZP0-2;Q8IZP0-4;Q8IZP0-3;Q8IZP0 | Q8IZP0-10;Q8IZP0-8;Q8IZP0-7;Q8IZP0-2;Q8IZP0-4;Q8IZP |
| Eukaryotic initiation factor 4A-I | EIF4A1 | | 0.808742 | -0.421652 | P60842;P60842-2 | P60842;P60842-2 |
| InaD-like protein | INADL | | 0.277316 | -0.42396 | Q8NI35-3;Q8NI35;Q8NI35-4;Q8NI35-2;Q8NI35-5 | Q8NI35-3;Q8NI35;Q8NI35-4;Q8NI35-2;Q8NI35-5 |
| Galectin-3-binding protein | LGALS3BP | | 0.643361 | -0.424375 | Q08380 | Q08380 |
| Eukaryotic translation initiation factor 4E | EIF4E | | 0.400506 | -0.426133 | P06730;P06730-2;P06730-3;A6NMX2 | P06730;P06730-2;P06730-3 |
| LIM and SH3 domain protein 1 | LASP1 | | 0.859699 | -0.426569 | Q14847;Q14847-2;Q14847-3 | Q14847;Q14847-2;Q14847-3 |
| AP-1 complex subunit mu-1 | AP1M1 | | 0.356435 | -0.428387 | Q9BXS5;Q9BXS5-2 | Q9BXS5;Q9BXS5-2 |
| S-phase kinase-associated protein 1 | SKP1 | | 1.16557 | -0.429548 | P63208;P63208-2 | P63208;P63208-2 |
| Synaptophysin-like protein 1 | SYPL1 | | 0.13084 | -0.429998 | Q16563-2;Q16563 | Q16563-2;Q16563 |
| Methionine adenosyltransferase 2 subunit beta | MAT2B | | 0.486409 | -0.432968 | Q9NZL9;Q9NZL9-4;Q9NZL9-2;Q9NZL9-3;Q9NZL9-5 | Q9NZL9;Q9NZL9-4;Q9NZL9-2;Q9NZL9-3 |
| Obg-like ATPase 1 | OLA1 | + | 2.06071 | -0.434794 | Q9NTK5;Q9NTK5-2 | Q9NTK5;Q9NTK5-2 |

| | | | | | | |
|---|---|---|---|---|---|---|
| Cytosolic acyl coenzyme A thioester hydrolase | ACOT7 | | 0.923711 | -0.436846 | O00154-4;O00154-6;O00154-7;O00154;O00154-5;O00154-2;O0015 | O00154-4;O00154-6;O00154-7;O00154;O00154-5;O0015 |
| Cytoskeleton-associated protein 4 | CKAP4 | | 0.839631 | -0.437103 | Q07065 | Q07065 |
| 3-hydroxyisobutyryl-CoA hydrolase, mitochondrial | HIBCH | | 0.464281 | -0.438179 | Q6NVY1;Q6NVY1-2 | Q6NVY1;Q6NVY1-2 |
| Chondroadherin | CHAD | | 0.538438 | -0.440312 | O15335 | O15335 |
| Twinfilin-2 | TWF2 | + | 3.27946 | -0.440466 | Q6IBS0;Q9NR96-4 | Q6IBS0 |
| Metalloproteinase inhibitor 3 | TIMP3 | | 0.649949 | -0.443235 | P35625 | P35625 |
| Transducin-like enhancer protein 3 | TLE3 | | 0.578718 | -0.44369 | Q04726-2;Q04726-7;Q04726-3;Q04726-6;Q04726-5;Q04726-4;Q047 | Q04726-2;Q04726-7;Q04726-3;Q04726-6;Q04726-5;Q04 |
| Eukaryotic translation initiation factor 2 subunit 1 | EIF2S1 | + | 1.46877 | -0.445976 | P05198 | P05198 |
| Tumor susceptibility gene 101 protein | TSG101 | + | 2.66215 | -0.446044 | Q99816;Q99816-2 | Q99816;Q99816-2 |
| Exportin-2 | CSE1L | | 0.818126 | -0.447002 | P55060-3;P55060;P55060-4;P55060-2 | P55060-3;P55060;P55060-4 |
| Serpin H1 | SERPINH1 | | 0.540995 | -0.448837 | P50454 | P50454 |
| Apolipoprotein B-100;Apolipoprotein B-48 | APOB | | 0.493092 | -0.449507 | P04114 | P04114 |
| Vascular endothelial growth factor A | VEGFA | | 0.235066 | -0.451277 | P15692-8;P15692-4;P15692-3;P15692-2;P15692;P15692-15;P15692- | P15692-8;P15692-4;P15692-3;P15692-2;P15692;P15692- |
| Tumor necrosis factor alpha-induced protein 2 | TNFAIP2 | | 0.292973 | -0.451511 | Q03169 | Q03169 |
| Catenin beta-1 | CTNNB1 | + | 1.47568 | -0.454746 | P35222;P35222-2 | P35222 |
| Purine nucleoside phosphorylase | PNP | | 0.453399 | -0.455106 | P00491 | P00491 |
| Cartilage intermediate layer protein 2;Cartilage intermediate layer protein 2 C1;Cartilage intermediate layer pro | CILP2 | | 1.22043 | -0.45588 | Q8IUL8 | Q8IUL8 |
| Ubiquitin carboxyl-terminal hydrolase 14 | USP14 | | 1.07775 | -0.457809 | P54578;P54578-2;P54578-3 | P54578;P54578-2;P54578-3 |
| T-complex protein 1 subunit epsilon | CCT5 | + | 1.3648 | -0.458191 | P48643;P48643-2 | P48643;P48643-2 |
| Peptidyl-prolyl cis-trans isomerase FKBP2 | FKBP2 | | 1.07087 | -0.460314 | P26885 | P26885 |
| Mitochondrial import receptor subunit TOM70 | TOMM70A | | 0.665121 | -0.460627 | O94826 | O94826 |
| Rabankyrin-5 | ANKFY1 | | 1.06064 | -0.461721 | Q9P2R3;Q9P2R3-2;Q9P2R3-4 | Q9P2R3;Q9P2R3-2;Q9P2R3-4 |
| Ribosyldihydronicotinamide dehydrogenase [quinone] | NQO2 | | 1.01757 | -0.462851 | P16083 | P16083 |
| Pseudouridine-5-phosphatase | HDHD1 | | 0.110703 | -0.462893 | Q08623-3;Q08623;Q08623-4;Q08623-2 | Q08623-3;Q08623;Q08623-4;Q08623-2 |
| Phosphoglycerate mutase 1 | PGAM1 | | 0.88716 | -0.463316 | P18669;P15259;Q8N0Y7 | P18669 |
| Acylphosphatase-1 | ACYP1 | | 0.285375 | -0.463805 | P07311;P07311-2 | P07311 |
| Major vault protein | MVP | | 0.391639 | -0.465055 | Q14764 | Q14764 |
| Hippocalcin-like protein 1 | HPCAL1 | | 0.980053 | -0.465907 | P37235;P84074 | P37235 |
| Diphosphoinositol polyphosphate phosphohydrolase 1 | NUDT3 | | 1.07733 | -0.466126 | O95989;Q96G61;Q8NFP7 | O95989 |
| Parkinson disease 7 domain-containing protein 1 | PDDC1 | | 0.639125 | -0.467033 | Q8NB37;Q8NB37-3;Q8NB37-2 | Q8NB37;Q8NB37-3;Q8NB37-2 |
| Staphylococcal nuclease domain-containing protein 1 | SND1 | | 1.10222 | -0.467915 | Q7KZF4 | Q7KZF4 |
| Matrix metalloproteinase-14 | MMP14 | + | 2.23184 | -0.469041 | P50281 | P50281 |
| 26S proteasome non-ATPase regulatory subunit 12 | PSMD12 | | 0.523781 | -0.471639 | O00232;O00232-2 | O00232;O00232-2 |
| Phospholipase A-2-activating protein | PLAA | + | 1.41346 | -0.475319 | Q9Y263 | Q9Y263 |
| NAD(P)H-hydrate epimerase | APOA1BP | | 0.551069 | -0.476625 | Q8NCW5;Q8NCW5-2 | Q8NCW5;Q8NCW5-2 |
| Calcium-binding protein 39 | CAB39 | | 0.772096 | -0.480082 | Q9Y376;Q9H9S4 | Q9Y376 |
| F-actin-capping protein subunit beta | CAPZB | | 1.07194 | -0.48026 | P47756-2;P47756 | P47756-2;P47756 |
| Proteasome subunit alpha type-2 | PSMA2 | | 0.400768 | -0.480422 | P25787 | P25787 |
| Vasodilator-stimulated phosphoprotein | VASP | + | 1.30423 | -0.484053 | P50552 | P50552 |
| T-complex protein 1 subunit delta | CCT4 | | 0.637474 | -0.486321 | P50991;P50991-2 | P50991;P50991-2 |
| Enoyl-CoA delta isomerase 1, mitochondrial | ECI1 | | 0.50511 | -0.487076 | P42126;P42126-2 | P42126;P42126-2 |
| Protein kinase C and casein kinase substrate in neurons protein 2 | PACSIN2 | | 0.669244 | -0.487244 | Q9UNF0-2;Q9UNF0 | Q9UNF0-2;Q9UNF0 |
| Caspase-14;Caspase-14 subunit p17, mature form;Caspase-14 subunit p10, mature form;Caspase-14 subunit p2 | CASP14 | | 0.436204 | -0.489419 | P31944 | P31944 |
| V-type proton ATPase subunit B, brain isoform | ATP6V1B2 | | 0.442936 | -0.490524 | P21281;P15313 | P21281 |
| Actin-related protein 2 | ACTR2 | | 0.95726 | -0.490589 | P61160;P61160-2 | P61160;P61160-2 |
| Proteasome subunit beta type-7 | PSMB7 | | 0.780474 | -0.49183 | Q99436;Q99436-2 | Q99436;Q99436-2 |
| Glutathione S-transferase omega-1 | GSTO1 | | 0.570569 | -0.4928 | P78417;P78417-3;P78417-2 | P78417;P78417-3;P78417-2 |
| HEAT repeat-containing protein 5B | HEATR5B | | 0.294009 | -0.493082 | Q9P2D3-2;Q9P2D3-3;Q9P2D3 | Q9P2D3-2;Q9P2D3-3;Q9P2D3 |
| Alpha-actinin-4 | ACTN4 | + | 2.10529 | -0.493472 | O43707;O43707-2;O43707-3 | O43707;O43707-2;O43707-3 |
| Proteasome subunit alpha type-5 | PSMA5 | | 0.458852 | -0.494062 | P28066;P28066-2 | P28066;P28066-2 |
| Leukotriene A-4 hydrolase | LTA4H | + | 1.50156 | -0.494639 | P09960;P09960-4;P09960-2;P09960-3 | P09960;P09960-4;P09960-2;P09960-3 |
| Carbonyl reductase [NADPH] 3 | CBR3 | | 0.936131 | -0.495181 | O75828 | O75828 |
| Prostasin;Prostasin light chain;Prostasin heavy chain | PRSS8 | | 0.190561 | -0.49615 | Q16651;Q16651-2 | Q16651;Q16651-2 |
| Protein ITFG3 | ITFG3 | | 0.596152 | -0.496701 | Q9H0X4;Q9H0X4-2 | Q9H0X4;Q9H0X4-2 |
| Pantetheinase | VNN1 | | 0.550381 | -0.496848 | O95497 | O95497 |
| Thioredoxin | TXN | | 0.371964 | -0.497038 | P10599;P10599-2 | P10599;P10599-2 |
| Deoxyribose-phosphate aldolase | DERA | | 1.18753 | -0.497823 | Q9Y315 | Q9Y315 |
| Heparan-sulfate 6-O-sulfotransferase 2 | HS6ST2 | | 0.788168 | -0.498966 | Q96MM7-2;Q96MM7-3;Q96MM7;Q96MM7-4 | Q96MM7-2;Q96MM7-3;Q96MM7;Q96MM7-4 |
| F-box only protein 2 | FBXO2 | | 1.28585 | -0.500305 | Q9UK22 | Q9UK22 |
| Olfactomedin-like protein 3 | OLFML3 | | 0.709773 | -0.50363 | Q9NRN5;Q9NRN5-2;Q9NRN5-3 | Q9NRN5;Q9NRN5-2 |
| MOB kinase activator 1A;MOB kinase activator 1B | MOB1A;MOB1B | | 0.329609 | -0.50418 | Q9H8S9;Q7L9L4;Q7L9L4-2;Q9H8S9-2 | Q9H8S9;Q7L9L4;Q7L9L4-2;Q9H8S9-2 |
| Ubiquitin-conjugating enzyme E2 L3 | UBE2L3 | + | 1.54048 | -0.506517 | P68036;P68036-3;P68036-2 | P68036;P68036-3;P68036-2 |
| Phosphatidylinositol transfer protein alpha isoform | PITPNA | | 0.281471 | -0.508685 | Q00169 | Q00169 |
| Protein kinase C-binding protein NELL2 | NELL2 | | 0.277365 | -0.509052 | Q99435-4;Q99435;Q99435-2;Q99435-3 | Q99435-4;Q99435;Q99435-2;Q99435-3 |
| Tenascin | TNC | | 0.402034 | -0.511128 | P24821;P24821-3 | P24821;P24821-3 |
| Sorting nexin-9 | SNX9 | | 0.506099 | -0.517483 | Q9Y5X1 | Q9Y5X1 |
| Nucleoside diphosphate kinase A | NME1 | | 0.350311 | -0.517897 | P15531;P15531-2 | P15531;P15531-2 |
| Protein BRICK1 | BRK1 | | 0.179841 | -0.518858 | Q8WUW1;Q8WUW1-2 | Q8WUW1;Q8WUW1-2 |
| Apoptosis regulator BAX | BAX | | 0.996956 | -0.521217 | Q07812-5;Q07812-8;Q07812;Q07812-2;Q07812-7;Q07812-4;Q0781 | Q07812-5;Q07812-8;Q07812;Q07812-2;Q07812-7 |
| Plastin-1 | PLS1 | | 0.780366 | -0.521489 | Q14651 | Q14651 |
| Ferritin light chain | FTL | | 0.255177 | -0.522064 | P02792 | P02792 |
| Cdc42-interacting protein 4 | TRIP10 | | 1.26114 | -0.52456 | Q15642-4;Q15642-2;Q15642-3;Q15642;Q15642-5 | Q15642-4;Q15642-2;Q15642-3;Q15642;Q15642-5 |
| 26S protease regulatory subunit 8 | PSMC5 | | 0.890077 | -0.524658 | P62195;P62195-2;Q8NB90-3;Q8NB90-2;Q8NB90 | P62195;P62195-2 |
| JmjC domain-containing protein 8 | JMJD8 | | 0.228368 | -0.524696 | Q96S16-2;Q96S16 | Q96S16-2;Q96S16 |
| Nitrilase homolog 1 | NIT1 | + | 2.09733 | -0.52782 | Q86X76-2;Q86X76-4;Q86X76;Q86X76-3;Q86X76-5 | Q86X76-2;Q86X76-4;Q86X76;Q86X76-3 |
| Hepatoma-derived growth factor | HDGF | | 1.24709 | -0.528108 | P51858;P51858-2;P51858-3 | P51858;P51858-2;P51858-3 |
| EH domain-containing protein 1 | EHD1 | | 0.485879 | -0.528617 | Q9H4M9 | Q9H4M9 |
| Flavin reductase (NADPH) | BLVRB | | 0.77783 | -0.531434 | P30043 | P30043 |
| F-actin-capping protein subunit alpha-1 | CAPZA1 | | 1.06119 | -0.531593 | P52907 | P52907 |
| Actin-related protein 2/3 complex subunit 5 | ARPC5 | | 0.748794 | -0.534572 | O15511;O15511-2 | O15511;O15511-2 |

| | | | | | | |
|---|---|---|---|---|---|---|
| Clathrin heavy chain 1 | CLTC | | 1.25902 | -0.537753 | Q00610-2;Q00610 | Q00610-2;Q00610 |
| Electron transfer flavoprotein subunit beta | ETFB | | 0.595945 | -0.538853 | P38117-2;P38117 | P38117-2;P38117 |
| Cytoplasmic dynein 1 heavy chain 1 | DYNC1H1 | | 0.949962 | -0.539417 | Q14204 | Q14204 |
| Kunitz-type protease inhibitor 2 | SPINT2 | | 0.489648 | -0.540133 | O43291;O43291-2 | O43291;O43291-2 |
| Protein CYR61 | CYR61 | | 0.728824 | -0.540255 | O00622 | O00622 |
| E3 ubiquitin-protein ligase NEDD4 | NEDD4 | | 0.251098 | -0.540318 | P46934-4;P46934-2;P46934;P46934-3 | P46934-4;P46934-2;P46934 |
| Proteasome subunit alpha type-7 | PSMA7 | + | 1.69182 | -0.541876 | O14818;O14818-4;Q8TAA3-2;Q8TAA3-5;Q8TAA3 | O14818 |
| Transketolase | TKT | | 0.812351 | -0.543564 | P29401;P29401-2 | P29401;P29401-2 |
| Eukaryotic translation initiation factor 2 subunit 3;Putative eukaryotic translation initiation factor 2 subunit 3-lik | EIF2S3;EIF2S3L | | 0.987776 | -0.545508 | P41091;Q2VIR3;Q2VIR3-2 | P41091;Q2VIR3;Q2VIR3-2 |
| Galectin-3 | LGALS3 | | 0.506732 | -0.548187 | P17931 | P17931 |
| Beta-2-microglobulin;Beta-2-microglobulin form pI 5.3 | B2M | | 0.134901 | -0.548678 | P61769 | P61769 |
| Collagen alpha-1(II) chain;Collagen alpha-1(II) chain;Chondrocalcin | COL2A1 | | 0.935643 | -0.548779 | P02458-3;P02458-1;P02458 | P02458-3;P02458-1;P02458 |
| Syndecan-4 | SDC4 | | 0.481934 | -0.554641 | P31431;P31431-2 | P31431;P31431-2 |
| Calmodulin | CALM1 | | 0.386845 | -0.554882 | P62158 | P62158 |
| PDZ domain-containing protein GIPC1 | GIPC1 | | 0.919917 | -0.555758 | O14908;O14908-2;Q8TF64 | O14908;O14908-2 |
| L-lactate dehydrogenase A chain | LDHA | | 0.704926 | -0.557631 | P00338;P00338-3;P00338-4;P00338-5;P00338-2;Q6ZMR3 | P00338;P00338-3;P00338-4;P00338-5;P00338-2 |
| Transforming growth factor-beta-induced protein ig-h3 | TGFBI | | 0.901304 | -0.559087 | Q15582 | Q15582 |
| SH3 domain-binding glutamic acid-rich-like protein 3 | SH3BGRL3 | | 0.656814 | -0.559813 | Q9H299 | Q9H299 |
| C1GALT1-specific chaperone 1 | C1GALT1C1 | | 0.294427 | -0.560267 | Q96EU7 | Q96EU7 |
| 6-phosphogluconate dehydrogenase, decarboxylating | PGD | + | 1.58192 | -0.562342 | P52209-2;P52209 | P52209-2;P52209 |
| Ganglioside GM2 activator;Ganglioside GM2 activator isoform short | GM2A | | 0.328111 | -0.562674 | P17900 | P17900 |
| Actin-related protein 3 | ACTR3 | | 0.742712 | -0.562828 | P61158;Q9C0K3;Q9P1U1-3 | P61158 |
| Plectin | PLEC | + | 1.48779 | -0.56348 | Q15149-2 | Q15149-2 |
| Argininosuccinate lyase | ASL | | 0.878681 | -0.564481 | P04424-2;P04424;P04424-3 | P04424-2;P04424;P04424-3 |
| Vacuolar protein sorting-associated protein 29 | VPS29 | | 0.759048 | -0.565422 | Q9UBQ0;Q9UBQ0-2 | Q9UBQ0;Q9UBQ0-2 |
| Thrombospondin-4 | THBS4 | | 0.617455 | -0.568146 | P35443 | P35443 |
| NAD(P)H dehydrogenase [quinone] 1 | NQO1 | | 0.809677 | -0.570733 | P15559-3;P15559-2;P15559 | P15559-3;P15559-2;P15559 |
| Mitogen-activated protein kinase 14 | MAPK14 | | 0.927164 | -0.572631 | Q16539;Q16539-2;Q16539-3;Q16539-5;Q16539-4 | Q16539;Q16539-2;Q16539-3;Q16539-5;Q16539-4 |
| Profilin-2 | PFN2 | | 1.02555 | -0.572711 | P35080-2 | P35080-2 |
| Serine/threonine-protein phosphatase 2B catalytic subunit gamma isoform | PPP3CC | | 0.160566 | -0.573718 | P48454-2;P48454;P48454-3 | P48454-2;P48454;P48454-3 |
| Actin-related protein 2/3 complex subunit 2 | ARPC2 | | 0.705514 | -0.574378 | O15144 | O15144 |
| Arylsulfatase A;Arylsulfatase A component B;Arylsulfatase A component C | ARSA | | 0.48985 | -0.574774 | P15289-2;P15289 | P15289-2;P15289 |
| Rho GDP-dissociation inhibitor 2 | ARHGDIB | | 1.10045 | -0.577962 | P52566 | P52566 |
| Vinculin | VCL | | 1.27025 | -0.578531 | P18206-2;P18206;P18206-3 | P18206-2;P18206 |
| Aspartate aminotransferase, mitochondrial | GOT2 | | 0.584312 | -0.579326 | P00505;P00505-2 | P00505;P00505-2 |
| Alpha-actinin-1 | ACTN1 | + | 1.39736 | -0.581653 | P12814;P12814-3;P12814-2;P12814-4 | P12814;P12814-3;P12814-2;P12814-4 |
| Mesencephalic astrocyte-derived neurotrophic factor | MANF | | 1.02235 | -0.581748 | P55145 | P55145 |
| Serine/threonine-protein kinase 10 | STK10 | | 0.237595 | -0.582423 | O94804 | O94804 |
| Lysozyme C | LYZ | | 0.170554 | -0.583816 | P61626 | P61626 |
| Niban-like protein 1 | FAM129B | | 0.976222 | -0.584056 | Q96TA1-2;Q96TA1 | Q96TA1-2;Q96TA1 |
| Interleukin-1 receptor antagonist protein | IL1RN | | 0.470408 | -0.591895 | P18510-2;P18510-4;P18510;P18510-3 | P18510-2;P18510-4;P18510;P18510-3 |
| Probable ATP-dependent RNA helicase DDX23 | DDX23 | | 0.68591 | -0.592381 | Q9BUQ8 | Q9BUQ8 |
| Beta-galactoside alpha-2,6-sialyltransferase 1 | ST6GAL1 | | 0.477425 | -0.593489 | P15907;P15907-2 | P15907;P15907-2 |
| Unconventional myosin-Ib | MYO1B | | 0.755947 | -0.594592 | O43795-2;O43795 | O43795-2;O43795 |
| Guanine nucleotide-binding protein G(i) subunit alpha-1 | GNAI1 | | 0.437411 | -0.595318 | P63096;P63096-2 | P63096;P63096-2 |
| Mannosyl-oligosaccharide glucosidase | MOGS | | 0.718004 | -0.598961 | Q13724-2;Q13724 | Q13724-2;Q13724 |
| Gamma-enolase | ENO2 | + | 1.69558 | -0.600765 | P09104;P09104-2 | P09104;P09104-2 |
| Very-long-chain 3-oxoacyl-CoA reductase | HSD17B12 | | 0.179372 | -0.602417 | Q53GQ0 | Q53GQ0 |
| Ubiquitin-conjugating enzyme E2 variant 3 | UEVLD | | 0.150716 | -0.602886 | Q8IX04-2;Q8IX04;Q8IX04-4;Q8IX04-3;Q8IX04-6;Q8IX04-7;Q8IX04-5 | Q8IX04-2;Q8IX04;Q8IX04-4;Q8IX04-3;Q8IX04-6;Q8IX04-7 |
| Moesin | MSN | + | 2.91496 | -0.605125 | P26038;REV__Q9H446-2;REV__Q9H446 | P26038 |
| Syntenin-1 | SDCBP | + | 1.72262 | -0.606747 | O00560;O00560-3 | O00560;O00560-3 |
| T-complex protein 1 subunit gamma | CCT3 | + | 1.60139 | -0.607475 | P49368;P49368-2 | P49368;P49368-2 |
| Programmed cell death 6-interacting protein | PDCD6IP | | 1.11376 | -0.607591 | Q8WUM4;Q8WUM4-2;Q8WUM4-3 | Q8WUM4;Q8WUM4-2 |
| Vesicle-fusing ATPase | NSF | | 0.626843 | -0.609786 | P46459;P46459-2 | P46459;P46459-2 |
| Syntaxin-7 | STX7 | | 0.715147 | -0.611944 | O15400-2;O15400 | O15400-2;O15400 |
| Golgin subfamily A member 4 | GOLGA4 | | 0.690454 | -0.611991 | Q13439-3;Q13439-4;Q13439;Q13439-5 | Q13439-3;Q13439-4;Q13439;Q13439-5 |
| ICOS ligand | ICOSLG | | 0.185566 | -0.614847 | O75144;O75144-2;O75144-3 | O75144;O75144-2;O75144-3 |
| Serpin B8 | SERPINB8 | + | 1.55234 | -0.615328 | P50452;P50452-2;P50452-3 | P50452;P50452-2 |
| Hydroxyacylglutathione hydrolase, mitochondrial | HAGH | | 0.324116 | -0.615626 | Q16775-2;Q16775;Q16775-3 | Q16775-2;Q16775 |
| Dystonin | DST | | 0.179568 | -0.616662 | Q03001;Q03001-8;Q03001-9;Q03001-13;Q03001-10 | Q03001;Q03001-8 |
| Adenylyl cyclase-associated protein 1 | CAP1 | + | 1.35821 | -0.621562 | Q01518-2 | Q01518-2 |
| Nuclear receptor 2C2-associated protein | NR2C2AP | | 0.798326 | -0.622456 | Q86WQ0;Q86WQ0-2 | Q86WQ0;Q86WQ0-2 |
| Hypoxanthine-guanine phosphoribosyltransferase | HPRT1 | | 0.775673 | -0.625255 | P00492 | P00492 |
| V-type proton ATPase catalytic subunit A | ATP6V1A | | 0.693518 | -0.625731 | P38606-2;P38606 | P38606-2;P38606 |
| Bleomycin hydrolase | BLMH | | 0.522182 | -0.628163 | Q13867 | Q13867 |
| Adipocyte plasma membrane-associated protein | APMAP | | 0.184496 | -0.629814 | Q9HDC9;Q9HDC9-2 | Q9HDC9;Q9HDC9-2 |
| Galectin-8 | LGALS8 | | 0.81267 | -0.631443 | O00214;O00214-2 | O00214;O00214-2 |
| T-complex protein 1 subunit beta | CCT2 | | 0.841095 | -0.631612 | P78371;P78371-2 | P78371;P78371-2 |
| Stanniocalcin-2 | STC2 | | 0.363056 | -0.632763 | O76061 | O76061 |
| T-complex protein 1 subunit theta | CCT8 | + | 1.78235 | -0.636568 | P50990;P50990-2;P50990-3 | P50990;P50990-2;P50990-3 |
| Aspartyl aminopeptidase | DNPEP | | 1.15431 | -0.64083 | Q9ULA0 | Q9ULA0 |
| 14 kDa phosphohistidine phosphatase | PHPT1 | | 0.717926 | -0.645281 | Q9NRX4;Q9NRX4-2 | Q9NRX4;Q9NRX4-2 |
| Trimethylguanosine synthase | TGS1 | | 0.147064 | -0.646936 | Q96RS0 | Q96RS0 |
| T-complex protein 1 subunit alpha | TCP1 | + | 1.45155 | -0.654202 | P17987 | P17987 |
| Protein XRP2 | RP2 | | 0.400215 | -0.655775 | O75695 | O75695 |
| Serpin B6 | SERPINB6 | + | 1.69391 | -0.657793 | P35237 | P35237 |
| 26S proteasome non-ATPase regulatory subunit 2 | PSMD2 | + | 1.56995 | -0.661743 | Q13200;Q13200-3;Q13200-2 | Q13200;Q13200-3;Q13200-2 |
| Succinate dehydrogenase [ubiquinone] iron-sulfur subunit, mitochondrial | SDHB | | 0.660884 | -0.663832 | P21912 | P21912 |
| Cell surface glycoprotein MUC18 | MCAM | | 1.21497 | -0.668236 | P43121;P43121-2 | P43121 |
| N(G),N(G)-dimethylarginine dimethylaminohydrolase 1 | DDAH1 | + | 1.51186 | -0.67108 | O94760;O94760-2 | O94760;O94760-2 |

| | | | | | | |
|---|---|---|---|---|---|---|
| Hepatocyte growth factor-regulated tyrosine kinase substrate | HGS | + | 1.47288 | -0.671343 | O14964;O14964-2 | O14964;O14964-2 |
| T-complex protein 1 subunit zeta | CCT6A | + | 1.8004 | -0.6726 | P40227;P40227-2;Q92526;Q92526-2;Q92526-3 | P40227;P40227-2 |
| Peptidyl-prolyl cis-trans isomerase-like 1 | PPIL1 | | 0.765583 | -0.674328 | Q9Y3C6 | Q9Y3C6 |
| ADP-dependent glucokinase | ADPGK | | 0.197994 | -0.675588 | Q9BRR6-2;Q9BRR6;Q9BRR6-4;Q9BRR6-3;Q9BRR6-5 | Q9BRR6-2;Q9BRR6;Q9BRR6-4;Q9BRR6-3 |
| Eukaryotic translation elongation factor 1 epsilon-1 | EEF1E1 | | 0.842313 | -0.676838 | O43324;O43324-2 | O43324;O43324-2 |
| Actin-related protein 2/3 complex subunit 3 | ARPC3 | + | 1.54665 | -0.67968 | O15145 | O15145 |
| Phosphatidylinositol 5-phosphate 4-kinase type-2 gamma | PIP4K2C | | 0.201247 | -0.679929 | Q8TBX8-3;Q8TBX8;Q8TBX8-2 | Q8TBX8-3;Q8TBX8;Q8TBX8-2 |
| Semaphorin-3A | SEMA3A | | 0.544012 | -0.683065 | Q14563 | Q14563 |
| Glutathione S-transferase theta-2B;Glutathione S-transferase theta-2 | GSTT2B;GSTT2 | + | 1.96884 | -0.683724 | P0CG30;P0CG29 | P0CG30;P0CG29 |
| DnaJ homolog subfamily C member 13 | DNAJC13 | | 1.03457 | -0.685371 | O75165 | O75165 |
| Serine/threonine-protein kinase MRCK beta | CDC42BPB | | 1.16701 | -0.6858 | Q9Y5S2 | Q9Y5S2 |
| 26S proteasome non-ATPase regulatory subunit 6 | PSMD6 | + | 1.48088 | -0.689058 | Q15008;Q15008-4;Q15008-3;Q15008-2 | Q15008;Q15008-4;Q15008-3;Q15008-2 |
| Plexin-A1 | PLXNA1 | | 0.381255 | -0.689319 | Q9UIW2 | Q9UIW2 |
| PDZ and LIM domain protein 2 | PDLIM2 | | 0.998617 | -0.690112 | Q96JY6;Q96JY6-5;Q96JY6-3;Q96JY6-4;Q96JY6-2 | Q96JY6;Q96JY6-5;Q96JY6-3;Q96JY6-4 |
| Anterior gradient protein 2 homolog | AGR2 | | 0.634042 | -0.69112 | O95994;Q8TD06 | O95994 |
| Serine/threonine-protein phosphatase 2A 65 kDa regulatory subunit A beta isoform | PPP2R1B | | 0.909386 | -0.69836 | P30154;P30154-3;P30154-2;P30154-4;P30154-5 | P30154;P30154-3;P30154-2;P30154-4;P30154-5 |
| STAM-binding protein | STAMBP | | 0.952286 | -0.699347 | O95630;O95630-2 | O95630;O95630-2 |
| Titin | TTN | | 0.591423 | -0.70342 | Q8WZ42-5;Q8WZ42-2;Q8WZ42;Q8WZ42-8;Q8WZ42-13;Q8WZ42-12 | Q8WZ42-5;Q8WZ42-2;Q8WZ42;Q8WZ42-8;Q8WZ42-13;( |
| Dual specificity protein phosphatase 3 | DUSP3 | | 0.241359 | -0.703684 | P51452-2 | P51452-2 |
| Dynein light chain 1, cytoplasmic | DYNLL1 | | 0.271881 | -0.707795 | P63167 | P63167 |
| Adenylyl cyclase-associated protein 1 | CAP1 | | 0.108464 | -0.719158 | Q01518 | Q01518 |
| 26S proteasome non-ATPase regulatory subunit 8 | PSMD8 | + | 1.51286 | -0.719961 | P48556 | P48556 |
| 5(3)-deoxyribonucleotidase, cytosolic type | NT5C | | 0.536456 | -0.721009 | Q8TCD5;Q8TCD5-2 | Q8TCD5;Q8TCD5-2 |
| Matrix metalloproteinase-15 | MMP15 | + | 1.30863 | -0.722004 | P51511;Q9Y5R2 | P51511 |
| WD repeat-containing protein 1 | WDR1 | + | 1.72119 | -0.725985 | O75083;O75083-3 | O75083;O75083-3 |
| Intercellular adhesion molecule 1 | ICAM1 | + | 2.52599 | -0.726014 | P05362 | P05362 |
| Interferon regulatory factor 6 | IRF6 | | 0.377743 | -0.727282 | O14896-2;O14896 | O14896-2;O14896 |
| Protein deglycase DJ-1 | PARK7 | + | 1.32605 | -0.728297 | Q99497 | Q99497 |
| Protein S100-A14 | S100A14 | | 0.399565 | -0.732166 | Q9HCY8 | Q9HCY8 |
| Spectrin beta chain, non-erythrocytic 2 | SPTBN2 | | 1.20743 | -0.738607 | O15020;O15020-2 | O15020;O15020-2 |
| Epithelial discoidin domain-containing receptor 1 | DDR1 | | 0.629026 | -0.739202 | Q08345-2;Q08345-6;Q08345;Q08345-5;Q08345-4 | Q08345-2;Q08345-6;Q08345;Q08345-5;Q08345-4 |
| Transcription elongation factor B polypeptide 2 | TCEB2 | | 0.532853 | -0.739371 | Q15370;Q15370-2 | Q15370;Q15370-2 |
| Neurogenic locus notch homolog protein 3;Notch 3 extracellular truncation;Notch 3 intracellular domain | NOTCH3 | + | 1.68544 | -0.73947 | Q9UM47 | Q9UM47 |
| Ubiquitin-like protein ISG15 | ISG15 | + | 2.31787 | -0.739841 | P05161 | P05161 |
| 26S proteasome non-ATPase regulatory subunit 3 | PSMD3 | + | 1.84603 | -0.740316 | O43242;O43242-2 | O43242;O43242-2 |
| Migration and invasion enhancer 1 | MIEN1 | | 0.355068 | -0.741835 | Q9BRT3 | Q9BRT3 |
| Mitochondrial-processing peptidase subunit alpha | PMPCA | | 0.207173 | -0.745553 | Q10713;Q10713-2 | Q10713;Q10713-2 |
| Glyoxylate reductase/hydroxypyruvate reductase | GRHPR | | 1.13389 | -0.745909 | Q9UBQ7 | Q9UBQ7 |
| Proteasome subunit alpha type-1 | PSMA1 | + | 2.19866 | -0.749501 | P25786;P25786-2 | P25786;P25786-2 |
| Sialic acid synthase | NANS | | 0.854919 | -0.751085 | Q9NR45 | Q9NR45 |
| Phospholipase D3 | PLD3 | | 0.416258 | -0.753145 | Q8IV08 | Q8IV08 |
| Aminoacyl tRNA synthase complex-interacting multifunctional protein 1;Endothelial monocyte-activating polyp | AIMP1 | | 1.03859 | -0.753566 | Q12904;Q12904-2 | Q12904;Q12904-2 |
| Puromycin-sensitive aminopeptidase | NPEPPS | | 1.19282 | -0.754814 | P55786;P55786-2;A6NEC2;A6NEC2-2;A6NEC2-3 | P55786;P55786-2 |
| N(G),N(G)-dimethylarginine dimethylaminohydrolase 2 | DDAH2 | | 0.957138 | -0.755954 | O95865 | O95865 |
| Tyrosine-protein phosphatase non-receptor type 6 | PTPN6 | | 0.775383 | -0.764127 | P29350;P29350-3;P29350-4;P29350-2 | P29350;P29350-3;P29350-4;P29350-2 |
| Nicotinate phosphoribosyltransferase | NAPRT | + | 2.18614 | -0.765801 | Q6XQN6;Q6XQN6-3;Q6XQN6-2 | Q6XQN6;Q6XQN6-3;Q6XQN6-2 |
| Kunitz-type protease inhibitor 1 | SPINT1 | | 0.716686 | -0.770717 | O43278 | O43278 |
| Cysteine-rich secretory protein LCCL domain-containing 2 | CRISPLD2 | | 0.536 | -0.771005 | Q9H0B8-3;Q9H0B8-2;Q9H0B8;O43692 | Q9H0B8-3;Q9H0B8-2;Q9H0B8 |
| Syntaxin-12 | STX12 | + | 1.82505 | -0.77666 | Q86Y82 | Q86Y82 |
| Glycolipid transfer protein | GLTP | | 0.550369 | -0.777514 | Q9NZD2 | Q9NZD2 |
| Macrophage-capping protein | CAPG | + | 2.85122 | -0.778807 | P40121;P40121-2 | P40121;P40121-2 |
| Cytosolic Fe-S cluster assembly factor NUBP1 | NUBP1 | | 0.619586 | -0.780479 | P53384-2;P53384 | P53384-2;P53384 |
| Syntaxin-binding protein 1 | STXBP1 | | 0.412327 | -0.783855 | P61764;P61764-2 | P61764;P61764-2 |
| Fumarylacetoacetase | FAH | | 0.661746 | -0.786798 | P16930;P16930-2 | P16930;P16930-2 |
| V-type proton ATPase subunit E 1 | ATP6V1E1 | | 0.841359 | -0.789042 | P36543;P36543-2 | P36543;P36543-2 |
| Galactokinase | GALK1 | | 0.921367 | -0.795251 | P51570;P51570-2 | P51570;P51570-2 |
| Annexin A8-like protein 2 | ANXA8L2 | | 0.559021 | -0.796061 | Q5VT79;Q5VT79-2 | Q5VT79;Q5VT79-2 |
| Transaldolase | TALDO1 | | 0.715712 | -0.79795 | P37837 | P37837 |
| AP-2 complex subunit mu | AP2M1 | | 0.814218 | -0.801189 | Q96CW1-2;Q96CW1 | Q96CW1-2;Q96CW1 |
| Low-density lipoprotein receptor-related protein 6 | LRP6 | | 1.09088 | -0.802299 | O75581 | O75581 |
| Insulin-like growth factor II;Insulin-like growth factor II;Insulin-like growth factor II Ala-25 Del;Preptin | IGF2 | | 0.785305 | -0.813304 | P01344;P01344-2;P01344-3 | P01344;P01344-2;P01344-3 |
| 1,4-alpha-glucan-branching enzyme | GBE1 | | 1.15984 | -0.814747 | Q04446 | Q04446 |
| Tubulin polymerization-promoting protein family member 3 | TPPP3 | | 0.544631 | -0.820274 | Q9BW30 | Q9BW30 |
| Coactosin-like protein | COTL1 | + | 2.6478 | -0.822316 | Q14019 | Q14019 |
| Vasorin | VASN | | 1.01789 | -0.822931 | Q6EMK4 | Q6EMK4 |
| Insulin-like growth factor-binding protein 6 | IGFBP6 | | 0.74837 | -0.825407 | P24592 | P24592 |
| Chondroitin sulfate proteoglycan 4 | CSPG4 | + | 1.51566 | -0.828871 | Q6UVK1 | Q6UVK1 |
| Heme oxygenase 1 | HMOX1 | | 0.286782 | -0.82989 | P09601 | P09601 |
| Collagenase 3 | MMP13 | | 0.965037 | -0.831727 | P45452 | P45452 |
| Ephrin-A1;Ephrin-A1, secreted form | EFNA1 | | 0.364967 | -0.833952 | P20827;P20827-2 | P20827;P20827-2 |
| Lysyl oxidase homolog 1 | LOXL1 | | 0.67638 | -0.834724 | Q08397 | Q08397 |
| Proteasome subunit beta type-9 | PSMB9 | + | 1.56738 | -0.836692 | P28065-2;P28065 | P28065-2;P28065 |
| Protein FAM162A | FAM162A | | 0.44237 | -0.836842 | Q96A26 | Q96A26 |
| Complement C3;Complement C3 beta chain;C3-beta-c;Complement C3 alpha chain;C3a anaphylatoxin;Acylatior | C3 | + | 1.85987 | -0.839137 | P01024 | P01024 |
| Breakpoint cluster region protein | BCR | | 0.504082 | -0.849439 | P11274-2;P11274 | P11274-2;P11274 |
| Chloride intracellular channel protein 4 | CLIC4 | | 1.02112 | -0.854155 | Q9Y696;O15247 | Q9Y696 |
| Lysosome membrane protein 2 | SCARB2 | | 0.384543 | -0.855055 | Q14108;Q14108-2 | Q14108;Q14108-2 |
| Spondin-1 | SPON1 | | 0.411566 | -0.855238 | Q9HCB6 | Q9HCB6 |
| Leucine-rich repeat-containing protein 16A | LRRC16A | | 0.484654 | -0.856474 | Q5VZK9-2;Q5VZK9;Q5VZK9-3;Q5VZK9-4 | Q5VZK9-2;Q5VZK9 |
| Cystatin-B | CSTB | + | 1.6832 | -0.861571 | P04080 | P04080 |

| | | | | | | |
|---|---|---|---|---|---|---|
| Intersectin-2 | ITSN2 | | 0.475674 | -0.867024 | Q9NZM3-4;Q9NZM3-3;Q9NZM3-2;Q9NZM3 | Q9NZM3-4;Q9NZM3-3;Q9NZM3-2;Q9NZM3 |
| Cartilage oligomeric matrix protein | COMP | | 0.670971 | -0.868481 | P49747;P49747-2 | P49747;P49747-2 |
| N-acetyl-D-glucosamine kinase | NAGK | + | 2.16113 | -0.872193 | Q9UJ70;Q9UJ70-2 | Q9UJ70;Q9UJ70-2 |
| Cytidine deaminase | CDA | + | 1.48891 | -0.87356 | P32320 | P32320 |
| Filamin-B | FLNB | + | 2.88673 | -0.876121 | O75369-8;O75369;O75369-2;O75369-9;O75369-3;O75369-6;O7536 | O75369-8;O75369;O75369-2;O75369-9;O75369-3;O7536 |
| Midkine | MDK | | 0.914115 | -0.876148 | P21741;P21741-2 | P21741 |
| Programmed cell death protein 10 | PDCD10 | | 0.741891 | -0.878911 | Q9BUL8 | Q9BUL8 |
| Biliverdin reductase A | BLVRA | | 1.16964 | -0.879613 | P53004 | P53004 |
| Leucine-rich repeat transmembrane protein FLRT3 | FLRT3 | + | 1.38905 | -0.881621 | Q9NZU0 | Q9NZU0 |
| Interleukin-1 receptor-like 1 | IL1RL1 | | 0.763404 | -0.889517 | Q01638-2;Q01638;Q01638-4;Q01638-3 | Q01638-2;Q01638;Q01638-4;Q01638-3 |
| Calpain-1 catalytic subunit | CAPN1 | + | 2.24806 | -0.891219 | P07384 | P07384 |
| Protein archease | ZBTB8OS | + | 1.94823 | -0.895805 | Q8IWT0;Q8IWT0-2 | Q8IWT0;Q8IWT0-2 |
| Glutaredoxin-1 | GLRX | | 1.19976 | -0.899164 | P35754 | P35754 |
| Protocadherin gamma-C3 | PCDHGC3 | | 0.176036 | -0.906384 | Q9UN70-2;Q9UN70 | Q9UN70-2;Q9UN70 |
| ADP-ribosylation factor-like protein 3 | ARL3 | | 0.326438 | -0.907213 | P36405 | P36405 |
| Cysteine-rich with EGF-like domain protein 2 | CRELD2 | | 0.430278 | -0.913537 | Q6UXH1-4;Q6UXH1;Q6UXH1-6;Q6UXH1-5;Q6UXH1-3;Q6UXH1-2 | Q6UXH1-4;Q6UXH1;Q6UXH1-6;Q6UXH1-5;Q6UXH1-3;Q6 |
| Charged multivesicular body protein 2a | CHMP2A | | 0.172013 | -0.917365 | O43633 | O43633 |
| Endoplasmic reticulum aminopeptidase 1 | ERAP1 | + | 2.14864 | -0.921822 | Q9NZ08;Q9NZ08-2 | Q9NZ08;Q9NZ08-2 |
| Serpin B7 | SERPINB7 | | 0.941154 | -0.924569 | O75635-2;O75635 | O75635-2;O75635 |
| Protein S100-A4 | S100A4 | + | 1.59877 | -0.942834 | P26447 | P26447 |
| Proteasome activator complex subunit 2 | PSME2 | + | 1.42375 | -0.952721 | Q9UL46 | Q9UL46 |
| WAP four-disulfide core domain protein 2 | WFDC2 | + | 1.76268 | -0.957677 | Q14508;Q14508-2;Q14508-3;Q14508-4;Q14508-5 | Q14508;Q14508-2;Q14508-3 |
| Protein CREG1 | CREG1 | | 0.280194 | -0.957758 | O75629 | O75629 |
| Protein S100-A16 | S100A16 | | 0.689003 | -0.961571 | Q96FQ6 | Q96FQ6 |
| Calpain-7 | CAPN7 | | 0.771568 | -0.96225 | Q9Y6W3 | Q9Y6W3 |
| Phosphatidylinositol 4-kinase type 2-alpha | PI4K2A | | 0.251438 | -0.969021 | Q9BTU6 | Q9BTU6 |
| Chloride intracellular channel protein 3 | CLIC3 | | 0.786341 | -0.974916 | O95833 | O95833 |
| Ubiquitin-fold modifier-conjugating enzyme 1 | UFC1 | | 1.28588 | -0.982649 | Q9Y3C8 | Q9Y3C8 |
| Succinyl-CoA ligase [ADP/GDP-forming] subunit alpha, mitochondrial | SUCLG1 | | 0.679581 | -0.986658 | P53597 | P53597 |
| Urokinase plasminogen activator surface receptor | PLAUR | + | 2.3837 | -1.00266 | Q03405;Q03405-2;Q03405-3 | Q03405;Q03405-2;Q03405-3 |
| Tumor necrosis factor receptor superfamily member 1A;Tumor necrosis factor receptor superfamily member 1 | TNFRSF1A | | 1.12503 | -1.00736 | P19438-5;P19438-4;P19438 | P19438-5;P19438-4;P19438 |
| Echinoderm microtubule-associated protein-like 2 | EML2 | | 1.15296 | -1.00821 | O95834;O95834-2;O95834-3;Q05BV3-4;Q05BV3-2;Q6ZMW3-2;Q6Z | O95834;O95834-2;O95834-3 |
| 15 kDa selenoprotein | SEP15 | | 0.413097 | -1.01414 | O60613;O60613-2 | O60613 |
| Proteasome subunit beta type-8 | PSMB8 | | 0.82371 | -1.01624 | P28062-2;P28062 | P28062-2;P28062 |
| Acetyl-CoA carboxylase 1;Biotin carboxylase | ACACA | | 0.29155 | -1.0168 | Q13085-4;Q13085-3;Q13085-2;Q13085;O00763-3;O00763-2;O0076 | Q13085-4;Q13085-3;Q13085-2;Q13085 |
| Arachidonate 15-lipoxygenase B | ALOX15B | | 0.652608 | -1.03409 | O15296-3;O15296;O15296-2;O15296-4 | O15296-3;O15296;O15296-2;O15296-4 |
| Programmed cell death protein 6 | PDCD6 | | 0.33797 | -1.03785 | O75340;O75340-2 | O75340;O75340-2 |
| 2-oxoglutarate and iron-dependent oxygenase domain-containing protein 3 | OGFOD3 | | 0.253593 | -1.04526 | Q6PK18;Q6PK18-2 | Q6PK18;Q6PK18-2 |
| Intraflagellar transport protein 25 homolog | HSPB11 | | 0.773602 | -1.04526 | Q9Y547 | Q9Y547 |
| DnaJ homolog subfamily B member 4 | DNAJB4 | | 0.805271 | -1.0476 | Q9UDY4 | Q9UDY4 |
| Coiled-coil domain-containing protein 80 | CCDC80 | | 0.871899 | -1.04856 | Q76M96;Q76M96-2;Q76M96-3 | Q76M96;Q76M96-2 |
| Laminin subunit alpha-3 | LAMA3 | | 1.24384 | -1.0504 | Q16787;Q16787-3 | Q16787;Q16787-3 |
| Protein MON2 homolog | MON2 | | 1.19878 | -1.05465 | Q7Z3U7-2;Q7Z3U7-6;Q7Z3U7-5;Q7Z3U7;Q7Z3U7-3 | Q7Z3U7-2;Q7Z3U7-6;Q7Z3U7-5;Q7Z3U7 |
| Argininosuccinate synthase | ASS1 | + | 1.46947 | -1.06138 | P00966 | P00966 |
| Complement C1q tumor necrosis factor-related protein 3 | C1QTNF3 | | 0.842838 | -1.06882 | Q9BXJ4;Q9BXJ4-3;Q9BXJ4-2 | Q9BXJ4;Q9BXJ4-3;Q9BXJ4-2 |
| Suppressor of tumorigenicity 14 protein | ST14 | + | 1.9773 | -1.0697 | Q9Y5Y6 | Q9Y5Y6 |
| V-type proton ATPase subunit F | ATP6V1F | | 0.366338 | -1.07022 | Q16864;Q16864-2 | Q16864;Q16864-2 |
| Follistatin | FST | + | 1.3407 | -1.07418 | P19883-2;P19883 | P19883-2;P19883 |
| Antileukoproteinase | SLPI | + | 1.81023 | -1.07494 | P03973 | P03973 |
| Tubulointerstitial nephritis antigen-like | TINAGL1 | + | 1.7474 | -1.07795 | Q9GZM7;Q9GZM7-3;Q9GZM7-2 | Q9GZM7;Q9GZM7-3 |
| Collagen alpha-1(XI) chain | COL11A1 | | 0.41213 | -1.09468 | P12107-4;P12107-3;P12107;P12107-2 | P12107-4;P12107-3;P12107;P12107-2 |
| Cellular retinoic acid-binding protein 2 | CRABP2 | | 0.334368 | -1.09531 | P29373;P29762 | P29373 |
| Pterin-4-alpha-carbinolamine dehydratase | PCBD1 | | 1.20049 | -1.11896 | P61457 | P61457 |
| MHC class I polypeptide-related sequence A | MICA | | 0.531536 | -1.143 | Q29983;Q29983-2 | Q29983 |
| Cytoplasmic aconitate hydratase | ACO1 | + | 2.71199 | -1.14423 | P21399 | P21399 |
| Macrophage colony-stimulating factor 1;Processed macrophage colony-stimulating factor 1 | CSF1 | | 1.11736 | -1.14829 | P09603;P09603-3;P09603-2 | P09603;P09603-3;P09603-2 |
| Serine/threonine-protein kinase Chk1 | CHEK1 | | 1.16552 | -1.14876 | O14757-2;O14757-3;O14757 | O14757-2;O14757-3;O14757 |
| Actin-related protein 2/3 complex subunit 1B | ARPC1B | + | 2.28622 | -1.15792 | O15143 | O15143 |
| BH3-interacting domain death agonist;BH3-interacting domain death agonist p15;BH3-interacting domain death | BID | + | 1.78068 | -1.16805 | P55957;P55957-2;P55957-4;P55957-3 | P55957;P55957-2 |
| UDP-glucose 6-dehydrogenase | UGDH | + | 1.47766 | -1.17251 | O60701;O60701-2 | O60701;O60701-2 |
| Protocadherin Fat 1;Protocadherin Fat 1, nuclear form | FAT1 | + | 2.68797 | -1.19326 | Q14517 | Q14517 |
| Latexin | LXN | | 0.656613 | -1.19835 | Q9BS40 | Q9BS40 |
| Serine/threonine-protein phosphatase 2A activator | PPP2R4 | | 0.896726 | -1.19839 | Q15257-3;Q15257-2;Q15257;Q15257-4 | Q15257-3;Q15257-2;Q15257;Q15257-4 |
| Coiled-coil and C2 domain-containing protein 1A | CC2D1A | | 0.391961 | -1.20665 | Q6P1N0;Q6P1N0-2 | Q6P1N0;Q6P1N0-2 |
| Acyl-protein thioesterase 2 | LYPLA2 | | 0.612055 | -1.21949 | O95372 | O95372 |
| SEC14-like protein 2 | SEC14L2 | + | 1.81844 | -1.22162 | O76054;O76054-5;O76054-4;Q9UDX4 | O76054;O76054-5;O76054-4 |
| Calmodulin-like protein 5 | CALML5 | | 0.524432 | -1.2226 | Q9NZT1 | Q9NZT1 |
| Protein kinase C delta type;Protein kinase C delta type regulatory subunit;Protein kinase C delta type catalytic s | PRKCD | | 0.351502 | -1.23573 | Q05655;Q05655-2 | Q05655;Q05655-2 |
| E3 ubiquitin-protein ligase LRSAM1 | LRSAM1 | | 0.720174 | -1.25374 | Q6UWE0-2;Q6UWE0 | Q6UWE0-2;Q6UWE0 |
| Tumor necrosis factor ligand superfamily member 13 | TNFSF13 | + | 1.63967 | -1.26718 | O43508-2;O75888-5;O75888-4;O75888;O75888-3 | O43508-2;O75888-5;O75888-4;O75888;O75888-3 |
| Ester hydrolase C11orf54 | C11orf54 | + | 1.94824 | -1.26884 | Q9H0W9-2;Q9H0W9;Q9H0W9-3;Q9H0W9-4 | Q9H0W9-2;Q9H0W9;Q9H0W9-3;Q9H0W9-4 |
| Aldo-keto reductase family 1 member B10 | AKR1B10 | | 1.249 | -1.2721 | O60218 | O60218 |
| Inhibin beta A chain | INHBA | + | 3.5301 | -1.28776 | P08476 | P08476 |
| NADH-cytochrome b5 reductase 3;NADH-cytochrome b5 reductase 3 membrane-bound form;NADH-cytochrom | CYB5R3 | | 0.414615 | -1.31499 | P00387-2;P00387;P00387-3 | P00387-2;P00387;P00387-3 |
| Insulin-like growth factor-binding protein 3 | IGFBP3 | + | 1.49458 | -1.32578 | P17936;P17936-2 | P17936;P17936-2 |
| Dual specificity protein phosphatase 23 | DUSP23 | | 0.373575 | -1.32601 | Q9BVJ7 | Q9BVJ7 |
| PDZ domain-containing protein 11 | PDZD11 | | 0.59981 | -1.33077 | Q5EBL8;Q5EBL8-2 | Q5EBL8;Q5EBL8-2 |
| Aldehyde dehydrogenase, mitochondrial | ALDH2 | + | 2.45221 | -1.33612 | P05091;P05091-2 | P05091;P05091-2 |
| Mitogen-activated protein kinase kinase kinase kinase 4 | MAP4K4 | | 0.37696 | -1.33743 | O95819-4;O95819-2;O95819;O95819-5;O95819-3;Q9UKE5-8;Q9UKE | O95819-4;O95819-2;O95819;O95819-5;O95819-3 |
| Coiled-coil domain-containing protein 132 | CCDC132 | + | 1.36909 | -1.33829 | Q96JG6;Q96JG6-3;Q96JG6-2 | Q96JG6;Q96JG6-3;Q96JG6-2 |

| | | | | | | |
|---|---|---|---|---|---|---|
| Pentraxin-related protein PTX3 | PTX3 | + | 1.84743 | -1.35994 | P26022 | P26022 |
| Interferon-induced guanylate-binding protein 1 | GBP1 | | 0.987581 | -1.36785 | P32455;Q9H0R5-4;Q9H0R5 | P32455 |
| Calcyphosin | CAPS | | 0.412067 | -1.37368 | Q13938;Q13938-3 | Q13938;Q13938-3 |
| Nectin-4;Processed poliovirus receptor-related protein 4 | PVRL4 | | 0.354113 | -1.38175 | Q96NY8;Q96NY8-2 | Q96NY8 |
| Desmoglein-3 | DSG3 | | 0.52073 | -1.38214 | P32926 | P32926 |
| Interleukin-36 gamma | IL36G | | 0.887552 | -1.38345 | Q9NZH8;Q9NZH8-2 | Q9NZH8;Q9NZH8-2 |
| Cystatin-M | CST6 | + | 1.59288 | -1.40995 | Q15828 | Q15828 |
| Complement C1r subcomponent;Complement C1r subcomponent heavy chain;Complement C1r subcomponent | C1R | + | 1.58271 | -1.41712 | P00736 | P00736 |
| Nephronectin | NPNT | | 0.782739 | -1.42062 | Q6UXI9;Q6UXI9-3;Q6UXI9-6;Q6UXI9-5;Q6UXI9-4;Q6UXI9-2 | Q6UXI9;Q6UXI9-3;Q6UXI9-6;Q6UXI9-5;Q6UXI9-4;Q6UXI9 |
| Ferritin heavy chain;Ferritin heavy chain, N-terminally processed | FTH1 | | 0.802557 | -1.42188 | P02794 | P02794 |
| Acyl-protein thioesterase 1 | LYPLA1 | + | 1.38556 | -1.42192 | O75608-2;O75608 | O75608-2;O75608 |
| Ras-related protein Rab-22A | RAB22A | | 0.42244 | -1.42198 | Q9UL26 | Q9UL26 |
| Plectin | PLEC | | 0.561709 | -1.46518 | Q15149-4;Q15149-7;Q15149-8;Q15149-9;Q15149-5;Q15149-6;Q151 | Q15149-4;Q15149-7;Q15149-8;Q15149-9;Q15149-5;Q15 |
| V-type proton ATPase subunit D | ATP6V1D | | 0.332174 | -1.48068 | Q9Y5K8 | Q9Y5K8 |
| Nesprin-2 | SYNE2 | | 0.360145 | -1.48334 | Q8WXH0;Q8WXH0-2;Q8WXH0-7;Q8WXH0-4;Q8WXH0-3;Q8WXH0-1 | Q8WXH0;Q8WXH0-2 |
| Interleukin-1 alpha | IL1A | | 0.551985 | -1.50277 | P01583 | P01583 |
| Alpha-1-antitrypsin;Short peptide from AAT | SERPINA1 | + | 1.69755 | -1.50367 | P01009;P01009-2;P01009-3 | P01009;P01009-2;P01009-3 |
| ADP-ribosylation factor-like protein 1 | ARL1 | | 0.588808 | -1.50562 | P40616-2;P40616 | P40616-2;P40616 |
| Secernin-2 | SCRN2 | | 1.00761 | -1.51541 | Q96FV2-2;Q96FV2 | Q96FV2-2;Q96FV2 |
| Nucleoside diphosphate kinase 3 | NME3 | | 0.612588 | -1.52048 | Q13232 | Q13232 |
| Putative peptidyl-tRNA hydrolase PTRHD1 | PTRHD1 | | 0.323685 | -1.57293 | Q6GMV3 | Q6GMV3 |
| Guanine deaminase | GDA | + | 1.47414 | -1.59998 | Q9Y2T3;Q9Y2T3-3;Q9Y2T3-2 | Q9Y2T3;Q9Y2T3-3;Q9Y2T3-2 |
| Laminin subunit beta-3 | LAMB3 | | 1.13397 | -1.6033 | Q13751 | Q13751 |
| Fibronectin;Anastellin;Ugl-Y1;Ugl-Y2;Ugl-Y3 | FN1 | + | 1.81343 | -1.61072 | P02751;P02751-3;P02751-8;P02751-17;P02751-14;P02751-9 | P02751;P02751-3;P02751-8;P02751-17;P02751-14;P0275 |
| Interleukin-6 receptor subunit beta | IL6ST | | 0.627837 | -1.6163 | P40189;P40189-3;P40189-2 | P40189;P40189-3 |
| Laminin subunit gamma-2 | LAMC2 | + | 1.5016 | -1.65974 | Q13753;Q13753-2 | Q13753;Q13753-2 |
| Quinone oxidoreductase PIG3 | TP53I3 | | 0.402212 | -1.69586 | Q53FA7;Q53FA7-2 | Q53FA7 |
| Tissue factor pathway inhibitor 2 | TFPI2 | | 0.582788 | -1.71499 | P48307-2;P48307 | P48307-2;P48307 |
| Bone morphogenetic protein 5 | BMP5 | | 0.545612 | -1.73124 | P22003-2;P22003 | P22003-2;P22003 |
| Protein-lysine 6-oxidase | LOX | + | 2.10998 | -1.75364 | P28300 | P28300 |
| Platelet-derived growth factor receptor-like protein | PDGFRL | | 0.692147 | -1.75718 | Q15198 | Q15198 |
| Peflin | PEF1 | | 1.03644 | -1.7895 | Q9UBV8 | Q9UBV8 |
| Semaphorin-4D | SEMA4D | | 0.884303 | -1.79126 | Q92854-2;Q92854 | Q92854-2;Q92854 |
| Fatty acid-binding protein, heart | FABP3 | + | 1.77865 | -1.80346 | P05413 | P05413 |
| Periplakin | PPL | | 0.797028 | -1.81015 | O60437 | O60437 |
| Matrilin-2 | MATN2 | | 0.493358 | -1.84445 | O00339-3;O00339-2;O00339;O00339-4 | O00339-3;O00339-2;O00339 |
| Protein Hikeshi | C11orf73 | | 0.599784 | -1.85657 | Q53FT3 | Q53FT3 |
| Monocyte differentiation antigen CD14;Monocyte differentiation antigen CD14, urinary form;Monocyte differe | CD14 | + | 1.73692 | -1.87093 | P08571 | P08571 |
| A disintegrin and metalloproteinase with thrombospondin motifs 1 | ADAMTS1 | | 0.730003 | -1.87378 | Q9UHI8;Q8TE58 | Q9UHI8 |
| Coronin-1A | CORO1A | + | 2.7237 | -1.87966 | P31146 | P31146 |
| Protein jagged-1 | JAG1 | + | 2.14597 | -1.89628 | P78504-2;P78504;Q9Y219-2;Q9Y219 | P78504-2;P78504 |
| Aldose reductase | AKR1B1 | + | 1.93015 | -1.91308 | P15121;C9JRZ8;C9JRZ8-2 | P15121 |
| Uridine diphosphate glucose pyrophosphatase | NUDT14 | + | 1.33708 | -1.92381 | O95848 | O95848 |
| Suprabasin | SBSN | | 1.17901 | -1.93296 | Q6UWP8;Q6UWP8-2 | Q6UWP8 |
| Serpin B9 | SERPINB9 | + | 1.66709 | -1.93672 | P50453 | P50453 |
| Hepatocyte growth factor-like protein;Hepatocyte growth factor-like protein alpha chain;Hepatocyte growth fa | MST1 | | 0.293722 | -1.96469 | P26927 | P26927 |
| Hyaluronan-binding protein 2;Hyaluronan-binding protein 2 50 kDa heavy chain;Hyaluronan-binding protein 2 5 | HABP2 | | 0.767976 | -1.98847 | Q14520-2;Q14520 | Q14520-2;Q14520 |
| Prenylcysteine oxidase 1 | PCYOX1 | | 0.764556 | -1.98905 | Q9UHG3-2;Q9UHG3 | Q9UHG3-2;Q9UHG3 |
| Desmoglein-1 | DSG1 | | 0.984192 | -2.00786 | Q02413;Q02413-2 | Q02413 |
| Fructose-1,6-bisphosphatase 1 | FBP1 | + | 2.04464 | -2.00796 | P09467;O00757 | P09467 |
| Leukocyte elastase inhibitor | SERPINB1 | + | 3.79014 | -2.01689 | P30740;P30740-2 | P30740;P30740-2 |
| Insulin-like growth factor-binding protein 2 | IGFBP2 | + | 2.81297 | -2.03366 | P18065 | P18065 |
| Involucrin | IVL | | 0.438197 | -2.05162 | P07476 | P07476 |
| Lysosome-associated membrane glycoprotein 1 | LAMP1 | | 0.546403 | -2.05447 | P11279;P11279-2 | P11279;P11279-2 |
| Nicotinamide phosphoribosyltransferase | NAMPT | + | 3.67303 | -2.0797 | P43490 | P43490 |
| C-X-C motif chemokine 16 | CXCL16 | | 1.17057 | -2.10522 | Q9H2A7 | Q9H2A7 |
| Ig gamma-1 chain C region | IGHG1 | | 0.40672 | -2.11611 | P01857 | P01857 |
| Dehydrogenase/reductase SDR family member 4 | DHRS4 | + | 2.04564 | -2.12048 | Q9BTZ2-3;Q9BTZ2-7;Q9BTZ2-2;Q9BTZ2-5;Q9BTZ2-8;Q9BTZ2-4;Q9B | Q9BTZ2-3;Q9BTZ2-7;Q9BTZ2-2;Q9BTZ2-5;Q9BTZ2-8;Q9B |
| Plakophilin-1 | PKP1 | | 0.618995 | -2.12808 | Q13835-2;Q13835 | Q13835-2;Q13835 |
| Neutrophil gelatinase-associated lipocalin | LCN2 | + | 1.57266 | -2.13162 | P80188 | P80188 |
| Protein Niban | FAM129A | + | 1.58419 | -2.13859 | Q9BZQ8 | Q9BZQ8 |
| Plastin-2 | LCP1 | | 1.26874 | -2.15594 | P13796;P13796-2 | P13796 |
| Cathepsin S | CTSS | + | 1.42745 | -2.25344 | P25774;P25774-2 | P25774;P25774-2 |
| CD82 antigen | CD82 | + | 2.95809 | -2.25786 | P27701-2;P27701 | P27701-2;P27701 |
| Pappalysin-1 | PAPPA | | 0.98081 | -2.30087 | Q13219 | Q13219 |
| Transmembrane protein 132A | TMEM132A | + | 2.49763 | -2.33846 | Q24JP5;Q24JP5-2;Q24JP5-4;Q24JP5-3 | Q24JP5;Q24JP5-2;Q24JP5-4 |
| Reticulon-3 | RTN3 | | 0.589153 | -2.35183 | O95197-3;O95197-4;O95197-7;O95197-2;O95197 | O95197-3;O95197-4;O95197-7;O95197-2;O95197 |
| Putative nucleoside diphosphate kinase | NME2P1 | | 0.525077 | -2.40268 | O60361 | O60361 |
| Transferrin receptor protein 1;Transferrin receptor protein 1, serum form | TFRC | + | 3.95717 | -2.41322 | P02786;Q9UP52-2;Q9UP52-3;Q9UP52 | P02786 |
| Serpin B3 | SERPINB3 | + | 2.41198 | -2.42901 | P29508;P29508-2 | P29508;P29508-2 |
| Ephrin type-B receptor 2 | EPHB2 | + | 1.6645 | -2.48489 | P29323-2;P29323-3;P29323;P54762-5;P54762 | P29323-2;P29323-3;P29323 |
| Signal recognition particle receptor subunit alpha | SRPR | | 0.666862 | -2.53684 | P08240;P08240-2 | P08240;P08240-2 |
| Oxidized low-density lipoprotein receptor 1;Oxidized low-density lipoprotein receptor 1, soluble form | OLR1 | + | 2.62553 | -2.61219 | P78380;P78380-2;P78380-3 | P78380;P78380-2;P78380-3 |
| Elafin | PI3 | + | 1.8916 | -2.70888 | P19957 | P19957 |
| Plasminogen activator inhibitor 1 | SERPINE1 | + | 1.79197 | -2.73163 | P05121;P05121-2 | P05121;P05121-2 |
| Aldo-keto reductase family 1 member C1 | AKR1C1 | | 0.932452 | -2.77494 | Q04828;P52895-2 | Q04828 |
| Tumor necrosis factor-inducible gene 6 protein | TNFAIP6 | + | 2.20357 | -2.80074 | P98066 | P98066 |
| Urokinase-type plasminogen activator;Urokinase-type plasminogen activator long chain A;Urokinase-type plasn | PLAU | + | 2.33625 | -3.00481 | P00749-2;P00749 | P00749-2;P00749 |
| Fibromodulin | FMOD | + | 1.45526 | -3.05074 | Q06828 | Q06828 |
| Alpha-2-macroglobulin-like protein 1 | A2ML1 | + | 1.94917 | -3.14194 | A8K2U0;A8K2U0-2 | A8K2U0;A8K2U0-2 |

| | | | | | | |
|---|---|---|---|---|---|---|
| Nectin-2 | PVRL2 | + | 2.58608 | -3.16731 | Q92692-2;Q92692 | Q92692-2;Q92692 |
| Superoxide dismutase [Mn], mitochondrial | SOD2 | + | 2.78011 | -3.18803 | P04179;P04179-4;P04179-2;P04179-3 | P04179;P04179-4;P04179-2;P04179-3 |
| Aldo-keto reductase family 1 member C3 | AKR1C3 | + | 3.56657 | -3.22993 | P42330;P42330-2;P17516 | P42330;P42330-2 |
| Alpha-1-antichymotrypsin;Alpha-1-antichymotrypsin His-Pro-less | SERPINA3 | + | 1.82144 | -3.33394 | P01011;P01011-2 | P01011 |
| Beta-actin-like protein 2 | ACTBL2 | | 0.241784 | -3.33421 | Q562R1 | Q562R1 |
| Plasminogen activator inhibitor 2 | SERPINB2 | + | 2.24401 | -3.37576 | P05120 | P05120 |
| Tenascin | TNC | | 0.965047 | -3.40287 | P24821-4;P24821-2;P24821-6 | P24821-4;P24821-2;P24821-6 |
| Growth-regulated alpha protein;GRO-alpha(4-73);GRO-alpha(5-73);GRO-alpha(6-73) | CXCL1 | + | 2.71754 | -3.66795 | P09341 | P09341 |
| Serpin B4 | SERPINB4 | + | 2.32533 | -3.67814 | P48594 | P48594 |
| Aldehyde dehydrogenase family 1 member A3 | ALDH1A3 | + | 4.50832 | -3.71605 | P47895;P30837 | P47895 |
| Platelet factor 4;Platelet factor 4, short form | PF4 | + | 1.31743 | -3.77294 | P02776 | P02776 |
| Complement C1s subcomponent;Complement C1s subcomponent heavy chain;Complement C1s subcomponen | C1S | + | 2.40954 | -3.84401 | P09871 | P09871 |
| Neuropilin-2 | NRP2 | + | 1.90656 | -4.05008 | O60462-4;O60462-5;O60462-2;O60462-3;O60462;O60462-6 | O60462-4;O60462-5;O60462-2;O60462-3;O60462;O6046 |
| Complement factor B;Complement factor B Ba fragment;Complement factor B Bb fragment | CFB | + | 3.08016 | -4.23605 | P00751;P00751-2 | P00751 |
| Interleukin-6 | IL6 | + | 2.87346 | -4.2843 | P05231 | P05231 |
| Leukemia inhibitory factor | LIF | + | 2.95337 | -4.62362 | P15018 | P15018 |
| Matrix metalloproteinase-9;67 kDa matrix metalloproteinase-9;82 kDa matrix metalloproteinase-9 | MMP9 | + | 2.94553 | -5.0757 | P14780 | P14780 |
| Ras GTPase-activating-like protein IQGAP2 | IQGAP2 | + | 4.03968 | -5.32723 | Q13576-3;Q13576-2;Q13576 | Q13576-3;Q13576-2;Q13576 |
| Cadherin-2 | CDH2 | | 1.05059 | -5.37665 | P19022-2 | P19022-2 |
| C-X-C motif chemokine 5;ENA-78(8-78);ENA-78(9-78) | CXCL5 | + | 3.35544 | -5.49615 | P42830 | P42830 |
| Platelet basic protein;Connective tissue-activating peptide III;TC-2;Connective tissue-activating peptide III(1-81) | PPBP | + | 2.36211 | -5.65301 | P02775 | P02775 |
| Interleukin-8;MDNCF-a;Interleukin-8;IL-8(5-77);IL-8(6-77);IL-8(7-77);IL-8(8-77);IL-8(9-77) | CXCL8 | | 1.02884 | -5.78774 | P10145;P10145-2 | P10145;P10145-2 |
| Protein S100-A9 | S100A9 | + | 3.36057 | -5.89133 | P06702 | P06702 |
| Protein S100-A7 | S100A7 | + | 1.9856 | -7.03533 | P31151;Q86SG5 | P31151 |
| Protein S100-A8;Protein S100-A8, N-terminally processed | S100A8 | + | 3.24581 | -7.12688 | P05109 | P05109 |
| C-C motif chemokine 20;CCL20(1-67);CCL20(1-64);CCL20(2-70) | CCL20 | + | 3.31661 | -7.98656 | P78556-2;P78556 | P78556-2;P78556 |
| C-X-C motif chemokine 3;GRO-gamma(5-73) | CXCL3 | + | 3.02168 | -9.17555 | P19876 | P19876 |

**Sup. Table 7: SUM149 1D annotation enrichment (NODAL versus EV)**

| C: Type | C: Name | N: Size | N: Score | N: P value | N: Benj. Hoch. FDR | N: Mean | N: Median |
|---|---|---|---|---|---|---|---|
| **GOBP name** | **chemokine-mediated signaling pathway** | **5** | **-0.957895** | **0.000238939** | **0.00897992** | **-5.55312** | **-5.49615** |
| GOBP name | cell chemotaxis | 12 | -0.810887 | 1.91E-06 | 0.000133647 | -4.61952 | -4.89023 |
| GOBP name | positive regulation of leukocyte chemotaxis | 9 | -0.82 | 2.77E-05 | 0.00148046 | -4.55077 | -4.2843 |
| GOBP name | positive regulation of leukocyte migration | 11 | -0.801708 | 6.32E-06 | 0.000415554 | -4.51541 | -4.2843 |
| GOBP name | regulation of leukocyte chemotaxis | 10 | -0.731104 | 8.40E-05 | 0.00398933 | -4.04054 | -4.02862 |
| GOBP name | regulation of leukocyte migration | 14 | -0.63293 | 6.30E-05 | 0.00312126 | -3.50137 | -3.72044 |
| GOBP name | response to lipopolysaccharide | 22 | -0.543554 | 2.15E-05 | 0.00118508 | -2.92077 | -2.95983 |
| GOBP name | positive regulation of response to external stimulus | 18 | -0.508209 | 0.00029654 | 0.0110235 | -2.91196 | -3.19979 |
| GOBP name | response to molecule of bacterial origin | 24 | -0.538889 | 1.16E-05 | 0.000710399 | -2.77474 | -2.31396 |
| GOBP name | regulation of response to external stimulus | 26 | -0.523512 | 9.98E-06 | 0.000632092 | -2.53814 | -2.76618 |
| **GOBP name** | **positive regulation of cell migration** | **24** | **-0.465497** | **0.000152319** | **0.00676536** | **-2.43325** | **-2.58032** |
| GOBP name | positive regulation of cell motility | 24 | -0.465497 | 0.000152319 | 0.00667862 | -2.43325 | -2.58032 |
| **GOBP name** | **inflammatory response** | **28** | **-0.547534** | **1.77E-06** | **0.000126304** | **-2.39666** | **-1.55719** |
| GOBP name | positive regulation of locomotion | 25 | -0.444507 | 0.000229083 | 0.00880297 | -2.31385 | -2.42901 |
| GOBP name | response to wounding | 31 | -0.545138 | 6.41E-07 | 5.35E-05 | -2.29644 | -1.61072 |
| GOBP name | response to inorganic substance | 21 | -0.487434 | 0.000192182 | 0.00801541 | -1.92384 | -1.50367 |
| GOBP name | regulation of cell migration | 32 | -0.401173 | 0.000202738 | 0.00815722 | -1.90932 | -1.60272 |
| GOBP name | regulation of endopeptidase activity | 24 | -0.540058 | 1.11E-05 | 0.000692351 | -1.87996 | -1.72019 |
| GOBP name | response to metal ion | 18 | -0.5063 | 0.000312499 | 0.0113697 | -1.84067 | -1.32395 |
| GOBP name | regulation of cell motility | 33 | -0.384936 | 0.000301586 | 0.0110906 | -1.83183 | -1.32578 |
| GOBP name | regulation of locomotion | 36 | -0.393773 | 0.000122915 | 0.00560492 | -1.78026 | -1.34286 |
| **GOBP name** | **positive regulation of immune system process** | **40** | **-0.415056** | **2.28E-05** | **0.00123861** | **-1.66507** | **-0.837914** |
| GOBP name | regulation of peptidase activity | 27 | -0.472025 | 5.08E-05 | 0.00255586 | -1.62439 | -1.40995 |
| GOBP name | negative regulation of endopeptidase activity | 16 | -0.553754 | 0.000191598 | 0.00808971 | -1.62128 | -1.72019 |
| GOBP name | immune response | 51 | -0.403405 | 5.31E-06 | 0.0003561 | -1.58381 | -0.836692 |
| GOBP name | negative regulation of hydrolase activity | 22 | -0.493506 | 0.000114474 | 0.00529054 | -1.55295 | -1.45681 |
| GOBP name | negative regulation of peptidase activity | 20 | -0.49308 | 0.000226251 | 0.00879295 | -1.46947 | -1.45681 |
| GOBP name | regulation of defense response | 30 | -0.419594 | 0.000159147 | 0.00688966 | -1.40411 | -0.793096 |
| GOBP name | positive regulation of response to stimulus | 58 | -0.364198 | 1.54E-05 | 0.000878011 | -1.35187 | -0.744908 |
| GOBP name | defense response | 61 | -0.39146 | 2.17E-06 | 0.00014873 | -1.34862 | -0.749501 |
| GOBP name | regulation of immune system process | 56 | -0.36773 | 1.66E-05 | 0.000932446 | -1.33465 | -0.744908 |
| GOBP name | regulation of cell proliferation | 42 | -0.354557 | 0.000221131 | 0.00869273 | -1.29699 | -1.21761 |
| GOBP name | regulation of response to stress | 40 | -0.40223 | 4.05E-05 | 0.00206823 | -1.275 | -0.793096 |
| GOCC name | extracellular region | 70 | -0.362821 | 3.90E-06 | 0.000776325 | -1.25868 | -1.07606 |
| GOBP name | immune system process | 65 | -0.318916 | 7.78E-05 | 0.00374877 | -1.22002 | -0.739841 |
| GOBP name | response to stress | 84 | -0.331746 | 7.24E-06 | 0.000467004 | -1.03079 | -0.727156 |
| GOBP name | negative regulation of catalytic activity | 37 | -0.364269 | 0.000324515 | 0.0116825 | -1.01317 | -0.728297 |
| GOBP name | regulation of response to stimulus | 91 | -0.313036 | 1.44E-05 | 0.000836011 | -0.96799 | -0.689058 |
| GOCC name | extracellular space | 93 | -0.27041 | 0.000163078 | 0.0049927 | -0.82741 | -0.726014 |

| | | | | | | | |
|---|---|---|---|---|---|---|---|
| GOBP name | positive regulation of cellular process | 113 | -0.270905 | 7.31E-05 | 0.0035694 | -0.777264 | -0.606747 |
| GOBP name | positive regulation of biological process | 135 | -0.274244 | 3.55E-05 | 0.00186889 | -0.661824 | -0.541876 |
| GOCC name | organelle | 282 | 0.398739 | 0.000619912 | 0.0117488 | 0.141802 | 0.450564 |
| GOCC name | membrane-bounded organelle | 281 | 0.400102 | 0.000480746 | 0.0106298 | 0.144938 | 0.451517 |
| GOCC name | membrane-bounded vesicle | 275 | 0.345027 | 0.00103074 | 0.0186471 | 0.149427 | 0.451517 |
| GOCC name | vesicle | 275 | 0.345027 | 0.00103074 | 0.0178363 | 0.149427 | 0.451517 |
| GOCC name | organelle part | 176 | 0.262816 | 7.63E-05 | 0.00379783 | 0.37663 | 0.638808 |
| GOCC name | intracellular organelle part | 174 | 0.274244 | 3.55E-05 | 0.00201955 | 0.395737 | 0.659781 |
| GOCC name | membrane | 173 | 0.293183 | 9.69E-06 | 0.000642845 | 0.398801 | 0.710527 |
| GOBP name | cellular macromolecule metabolic process | 119 | 0.241221 | 0.000359493 | 0.012807 | 0.401717 | 0.747575 |
| GOBP name | cellular component organization at cellular level | 128 | 0.292559 | 1.18E-05 | 0.000710004 | 0.493676 | 0.853663 |
| GOBP name | cellular component organization or biogenesis at cellular level | 128 | 0.292559 | 1.18E-05 | 0.000697762 | 0.493676 | 0.853663 |
| GOBP name | cellular component organization | 150 | 0.344822 | 1.62E-07 | 1.58E-05 | 0.529733 | 0.888187 |
| GOBP name | cellular component organization or biogenesis | 150 | 0.344822 | 1.62E-07 | 1.54E-05 | 0.529733 | 0.888187 |
| GOBP name | cellular biosynthetic process | 87 | 0.272859 | 0.000191183 | 0.00817309 | 0.549292 | 0.977729 |
| GOBP name | biosynthetic process | 89 | 0.278447 | 0.000126633 | 0.0056985 | 0.556772 | 0.966807 |
| GOBP name | nitrogen compound metabolic process | 92 | 0.264877 | 0.000231691 | 0.00880427 | 0.569716 | 0.822344 |
| GOCC name | macromolecular complex | 120 | 0.308113 | 5.00E-06 | 0.000663647 | 0.59424 | 0.729856 |
| GOBP name | cellular nitrogen compound metabolic process | 82 | 0.276351 | 0.000208051 | 0.00827366 | 0.600175 | 0.941646 |
| GOCC name | cell junction | 70 | 0.291811 | 0.000204785 | 0.00582174 | 0.65913 | 1.13268 |
| GOBP name | establishment of localization | 100 | 0.348325 | 7.28E-07 | 5.79E-05 | 0.661886 | 1.10313 |
| GOCC name | cell-substrate adherens junction | 57 | 0.290866 | 0.000605052 | 0.0126742 | 0.687598 | 1.16472 |
| GOCC name | focal adhesion | 57 | 0.290866 | 0.000605052 | 0.0120405 | 0.687598 | 1.16472 |
| GOBP name | transport | 98 | 0.362511 | 2.93E-07 | 2.57E-05 | 0.701357 | 1.11324 |
| GOCC name | adherens junction | 59 | 0.303322 | 0.000290161 | 0.00769893 | 0.715053 | 1.16472 |
| GOCC name | anchoring junction | 59 | 0.303322 | 0.000290161 | 0.00721775 | 0.715053 | 1.16472 |
| GOCC name | cell-substrate junction | 58 | 0.303613 | 0.000313693 | 0.00734412 | 0.715252 | 1.19276 |
| GOBP name | establishment of localization in cell | 79 | 0.373583 | 7.28E-07 | 5.92E-05 | 0.764035 | 1.25992 |
| GOBP name | macromolecule catabolic process | 57 | 0.33027 | 9.86E-05 | 0.00462113 | 0.805121 | 1.12287 |
| GOBP name | nucleobase-containing compound metabolic process | 67 | 0.38954 | 1.06E-06 | 7.74E-05 | 0.874148 | 1.32112 |
| GOBP name | cellular macromolecule catabolic process | 50 | 0.368649 | 3.68E-05 | 0.00190502 | 0.896987 | 1.24292 |
| GOBP name | macromolecule biosynthetic process | 58 | 0.416953 | 7.46E-07 | 5.80E-05 | 0.942658 | 1.32915 |
| GOBP name | gene expression | 56 | 0.421796 | 7.85E-07 | 5.97E-05 | 0.995939 | 1.35562 |
| GOBP name | cellular macromolecule biosynthetic process | 51 | 0.448092 | 4.27E-07 | 3.65E-05 | 1.01474 | 1.37536 |
| GOBP name | cellular process involved in reproduction | 46 | 0.453133 | 9.44E-07 | 7.02E-05 | 1.06143 | 1.35562 |
| GOBP name | protein complex subunit organization | 60 | 0.458501 | 3.53E-08 | 3.65E-06 | 1.08605 | 1.4447 |
| GOBP name | nucleic acid metabolic process | 59 | 0.487593 | 5.69E-09 | 6.09E-07 | 1.09435 | 1.38121 |
| GOBP name | macromolecular complex subunit organization | 65 | 0.477932 | 3.20E-09 | 3.77E-07 | 1.10874 | 1.43627 |
| GOBP name | cellular component disassembly | 50 | 0.460232 | 2.57E-07 | 2.38E-05 | 1.14336 | 1.37828 |
| GOBP name | cellular component disassembly at cellular level | 50 | 0.460232 | 2.57E-07 | 2.32E-05 | 1.14336 | 1.37828 |
| GOBP name | RNA metabolic process | 54 | 0.505592 | 5.33E-09 | 5.88E-07 | 1.15511 | 1.48411 |

| | | | | | | | |
|---|---|---|---|---|---|---|---|
| GOBP name | establishment of protein localization | 61 | 0.50119 | 1.33E-09 | 2.39E-07 | 1.1608 | 1.47085 |
| GOBP name | protein transport | 59 | 0.511322 | 1.00E-09 | 1.90E-07 | 1.19441 | 1.48332 |
| GOCC name | nucleolus | 25 | 0.456338 | 0.00015517 | 0.00561435 | 1.20053 | 1.59818 |
| GOBP name | RNA biosynthetic process | 50 | 0.528649 | 3.26E-09 | 3.71E-07 | 1.20653 | 1.52113 |
| GOBP name | viral reproduction | 55 | 0.473443 | 3.69E-08 | 3.72E-06 | 1.20966 | 1.37536 |
| GOBP name | intracellular transport | 57 | 0.529518 | 4.29E-10 | 1.33E-07 | 1.24235 | 1.48332 |
| GOBP name | intracellular protein transport | 49 | 0.544898 | 1.45E-09 | 2.47E-07 | 1.30156 | 1.48491 |
| GOBP name | cellular macromolecular complex subunit organization | 47 | 0.581939 | 2.12E-10 | 1.21E-07 | 1.38582 | 1.48491 |
| GOCC name | ribonucleoprotein complex | 40 | 0.580483 | 3.15E-09 | 1.25E-06 | 1.40243 | 1.52113 |
| GOBP name | viral reproductive process | 42 | 0.588193 | 8.93E-10 | 2.04E-07 | 1.40885 | 1.57777 |
| GOBP name | protein targeting | 44 | 0.590738 | 3.49E-10 | 1.19E-07 | 1.44417 | 1.57777 |
| **GOBP name** | **establishment of protein localization to organelle** | **38** | **0.603418** | **1.70E-09** | **2.65E-07** | **1.48738** | **1.57777** |
| GOCC name | ribosome | 14 | 0.59661 | 0.000162092 | 0.00537606 | 1.51725 | 1.65523 |
| GOBP name | translation | 34 | 0.626524 | 2.53E-09 | 3.09E-07 | 1.52386 | 1.57777 |
| GOBP name | viral infectious cycle | 38 | 0.649058 | 9.20E-11 | 3.15E-07 | 1.58895 | 1.62163 |
| GOBP name | translational initiation | 32 | 0.647112 | 2.05E-09 | 2.60E-07 | 1.59239 | 1.62163 |
| GOBP name | mRNA catabolic process | 32 | 0.649819 | 1.76E-09 | 2.62E-07 | 1.59547 | 1.62163 |
| GOBP name | nuclear-transcribed mRNA catabolic process | 32 | 0.649819 | 1.76E-09 | 2.51E-07 | 1.59547 | 1.62163 |
| GOBP name | nuclear-transcribed mRNA catabolic process, nonsense-mediated decay | 32 | 0.649819 | 1.76E-09 | 2.41E-07 | 1.59547 | 1.62163 |
| GOBP name | RNA catabolic process | 32 | 0.649819 | 1.76E-09 | 2.32E-07 | 1.59547 | 1.62163 |
| GOBP name | translational elongation | 32 | 0.651173 | 1.63E-09 | 2.65E-07 | 1.59855 | 1.62163 |
| GOBP name | protein targeting to membrane | 34 | 0.652834 | 5.30E-10 | 1.51E-07 | 1.6124 | 1.62163 |
| **GOBP name** | **mRNA metabolic process** | **35** | **0.6561** | **2.61E-10** | **9.90E-08** | **1.62027** | **1.64508** |
| GOCC name | cytosolic small ribosomal subunit | 18 | 0.623902 | 8.91E-06 | 0.000886787 | 1.62106 | 1.54154 |
| GOCC name | small ribosomal subunit | 18 | 0.623902 | 8.91E-06 | 0.00070943 | 1.62106 | 1.54154 |
| GOCC name | cytosolic large ribosomal subunit | 14 | 0.616949 | 9.61E-05 | 0.00424761 | 1.63036 | 1.85468 |
| GOCC name | large ribosomal subunit | 14 | 0.616949 | 9.61E-05 | 0.00382285 | 1.63036 | 1.85468 |
| GOBP name | cotranslational protein targeting to membrane | 32 | 0.665839 | 6.97E-10 | 1.83E-07 | 1.64082 | 1.62163 |
| GOBP name | SRP-dependent cotranslational protein targeting to membrane | 32 | 0.665839 | 6.97E-10 | 1.70E-07 | 1.64082 | 1.62163 |
| GOBP name | translational termination | 31 | 0.670225 | 9.31E-10 | 1.99E-07 | 1.65815 | 1.64508 |
| GOBP name | viral transcription | 31 | 0.670225 | 9.31E-10 | 1.87E-07 | 1.65815 | 1.64508 |
| GOBP name | establishment of protein localization in endoplasmic reticulum | 33 | 0.678305 | 1.91E-10 | 1.64E-07 | 1.67965 | 1.64508 |
| GOBP name | protein targeting to ER | 33 | 0.678305 | 1.91E-10 | 1.31E-07 | 1.67965 | 1.64508 |
| **GOBP name** | **cellular macromolecular complex disassembly** | **33** | **0.684673** | **1.29E-10** | **2.21E-07** | **1.73447** | **1.64508** |
| GOBP name | macromolecular complex disassembly | 33 | 0.684673 | 1.29E-10 | 1.47E-07 | 1.73447 | 1.64508 |
| GOBP name | cellular protein complex disassembly | 32 | 0.685469 | 2.17E-10 | 1.06E-07 | 1.74701 | 1.65523 |
| GOBP name | protein complex disassembly | 32 | 0.685469 | 2.17E-10 | 9.28E-08 | 1.74701 | 1.65523 |
| GOBP name | cellular component biogenesis | 10 | 0.654181 | 0.000433259 | 0.0152757 | 1.7874 | 1.78112 |
| GOBP name | cellular component biogenesis at cellular level | 10 | 0.654181 | 0.000433259 | 0.0151199 | 1.7874 | 1.78112 |
| GOBP name | ribonucleoprotein complex biogenesis | 10 | 0.654181 | 0.000433259 | 0.0149671 | 1.7874 | 1.78112 |
| GOBP name | rRNA metabolic process | 11 | 0.660769 | 0.000197937 | 0.00815596 | 1.79286 | 1.71861 |

| | | | | | | | |
|---|---|---|---|---|---|---|---|
| GOBP name | rRNA processing | 11 | 0.660769 | 0.000197937 | 0.00805887 | 1.79286 | 1.71861 |

**Sup. Table 8: Antibody information**

| Antibody | Vendor | Application | Dilution |
|---|---|---|---|
| Alpha-SMA (ab5694) Rabbit Polyclonal antibody | Abcam | Immunohistochemistry | 1:1000 |
| Nodal (clone 5C3) Mouse mAb | Abcam | Immunohistochemistry | 1:500 |
| Nodal (WS65) Mouse mAb | Santa Cruz Biotechnology | Western Blotting | 1:2500 |
| β-Actin (C4) Mouse mAb | Santa Cruz Biotechnology | Western Blotting | 1:5000 |
| Stat3 (F-2) Mouse mAb | Santa Cruz Biotechnology | Western Blotting | 1:2000 |
| p-Stat3 (B-7) Mouse mAb | Santa Cruz Biotechnology | Western Blotting | 1:2000 |
| Smad2/3 (D7G7) XP® Rabbit mAb | Cell Signaling Technology | Western Blotting | 1:5000 |
| Phospho-Smad2 (Ser465/467) (138D4) Rabbit mAb | Cell Signaling Technology | Western Blotting | 1:3000 |
| Phospho-Smad3 (Ser423/425) (C25A9) Rabbit mAb | Cell Signaling Technology | Western Blotting | 1:3000 |
| p44/42 MAPK (Erk1/2) (137F5) Rabbit mAb | Cell Signaling Technology | Western Blotting | 1:5000 |
| Phospho-p44/42 MAPK (Erk1/2) (Thr202/Tyr204) (D13.14.4E) XP® Rabbit mAb | Cell Signaling Technology | Western Blotting | 1:5000 |
| Phospho-p38 MAPK (Thr180/Tyr182) (D3F9) XP® Rabbit mAb | Cell Signaling Technology | Western Blotting | 1:3000 |
| Human Activin RIB/ALK-4 Mouse mAb | R&D systems | Western Blotting | 1:2000 |
| Goat Anti-Mouse IgG (H + L)-HRP Conjugate | Bio Rad | Western Blotting | 1:10000 |
| Goat Anti-Rabbit IgG (H + L)-HRP Conjugate | Bio Rad | Western Blotting | 1:10000 |
| Mouse IgG1 K Isotype Control Purified | eBioscience | Flow cytometry | 0.2µg/1e5 cells |
| Anti-Human CD126 (IL6R) Purified | eBioscience | Flow cytometry | 0.2-0.5µg/1e5 cells |
| Anti-Human CD182 (CXCR2) Purified IgG1, kappa | ebioscience | Flow cytometry | 0.5µg/1e5 cells |
| APC anti-Mouse IgG1 Secondary | eBioscience | Flow cytometry | 0.2µg/1e5 cells |
| FITC Mouse IgG2b, κ Isotype Ctrl | Biolegend | Flow cytometry | 0.2µg/1e5 cells |
| FITC anti-Human CD181 (CXCR1) | Biolegend | Flow cytometry | 0.2µg/1e5 cells |
| PE Mouse IgG1, κ Isotype Ctrl | Biolegend | Flow cytometry | 0.2µg/1e5 cells |
| PE anti-Human CD182 (CXCR2) | Biolegend | Flow cytometry | 0.2µg/1e5 cells |
| Human Cripto 1 PE-conjugated Antibody | R&D Systems | Flow cytometry | 2µL/1e5 cells |
| Human IL-6 mAb (Clone 1936) Monoclonal Mouse IgG2B Clone# 1936 | R&D Systems | Neutralization | 2.5µg/mL |
| Mouse IgG2B Isotype Control (Clone 20116), Mouse IgG2B | R&D Systems | Neutralization | 2.5µg/mL |

**Sup. Table 9: Mass-spectrometry instrument settings for data acquisition**

| **Parameter** | **Q Exactive** | **Orbitrap Elite** |
|---|---|---|
| Orbitrap resolution (MS1) | 70K | 120K |
| Mass range | 400-1450m/z | 400-1450m/z |
| MS1 Injection time | 250ms | 200ms |
| MS1 AGC taget | 1.00E+06 | 1.00E+06 |
| Lock mass | 445.120025 | 445.120025 |
| MS2 detection | FT | IT |
| MS2 resolution | 17.5K | rapid scan mode |
| MS2 minimum signal threshold | 5.00E+04 | 1.00E+03 |
| MS2 AGC target | 2.00E+05 | 1.00E+04 |
| MS2 injection time | 120ms | 50ms |
| MS2 scan events | 15 | 10 |
| Isolation width | 3 | 2 |
| MS2 Activation | HCD | rapid CID |
| Normalized Collision Energy | 23 | 35 |
| Dynamic exclusion | enabled | enabled |
| Exclusion duration | 30 | 30 |
| Exclusion mass width (Da) | +/- 1.5 | -0.5, +1.5 |
| Charge Exclusion | unassigned, 1, >8 | unassigned, 1, >8 |

**Sup. Table 10: TaqMan primer probe sets used for real-time PCR**

| Gene | Assay ID |
|---|---|
| Nodal | Hs00250630_s1 |
| ALK4 | Hs00244715_m1 |
| Cripto | Hs02339499_g1 |
| CXCR1 | Hs00174146_m1 |
| CXCR2 | Hs01011557_m1 |
| CTGF | Hs01026927_g1 |
| Desmin | Hs00157258_M1 |
| a-SMA | Hs00426835_g1 |
| HPRT1 | 4333768T |
| RPLPO | 4333761T |
